# Regional and cell type-specific afferent and efferent projections of the mouse claustrum

**DOI:** 10.1101/2022.02.23.481555

**Authors:** Quanxin Wang, Yun Wang, Peng Xie, Hsien-Chi Kuo, Karla E Hirokawa, Maitham Naeemi, Shenqin Yao, Ben Ouellette, Matt Mallory, Phil Lesnar, Xiuli Kuang, Yaoyao Li, Min Ye, Chao Chen, Wei Xiong, Leila Ahmadinia, Laila El-Hifnawi, Ali Cetin, Julie A Harris, Hongkui Zeng, Christof Koch

**Author notes:** Co-senior authors (Hongkui Zeng, Christof Koch). These authors contributed equally (Quanxin Wang, Yun Wang).

## Abstract

The claustrum (CLA) is a conspicuous subcortical structure interconnected with cortical and subcortical regions. However, its regional anatomy and cell-type-specific connections in the mouse remain largely undetermined. Here, we accurately delineated the boundary of the mouse CLA and quantitatively investigated its inputs and outputs brain-wide using anterograde and retrograde viral tracing and fully reconstructed single claustral principal neurons. At a population level, the CLA reciprocally connects with all isocortical modules. It also receives inputs from at least 35 subcortical structures but sends projections back to only a few of them. We found that cell types projecting to the CLA are differentiated by cortical areas and layers. We classified single CLA principal neurons into at least 9 cell types that innervate the diverse sets of functionally linked cortical targets. Axons of interneurons within the CLA arborize along almost its entire anteroposterior extent. Together, this detailed wiring diagram of the cell-type-specific connections of the mouse CLA lays a foundation for studying its functions.

## Introduction

The claustrum (CLA) is a thin, elongated and sheet-like subcortical structure located between the insular cortex and the striatum. It is well-conserved from reptiles to mammals (Bruguier et al., 2020; Norimoto et al., 2020), and is the most highly reciprocally connected structure with the cortex (Peng et al., 2021; Torgerson et al., 2015; Wang et al., 2017; Zingg et al., 2014). Although its function is not fully understood (Atilgan et al., 2021; Crick and Koch 2005), recent studies using a combination of optogenetics and electrophysiological recording or fiber photometry in transgenic mice suggest that the CLA is involved in sleep (Atlan et al., 2021; Narikiyo et al., 2020), inhibition of the medial prefrontal cortex (Jackson et al., 2018), selective attention (Atlan et al., 2018; Fodoulian et al., 2019; Terem et al., 2020), modulation of engagement with the external world (Atlan et al., 2021) and is required for shifting to a complex cognitive task (White et al., 2020). Furthermore, top-down signals from the anterior cingulate cortex may go through the CLA in a circuit that influences posterior sensory and association cortices (White and Mathur, 2018). Because it is amenable to advanced transgenic tools and newly developed technologies, the laboratory mouse has become a popular animal model for studying CLA functions (Jackson et al., 2020; Smith et al., 2020).

A crucial step to understanding the cell types, circuits, and function of a given structure, is to accurately delineate the structure using anatomical criteria. While the CLA has been a subject of study for many years, there are still discrepancies in how it is annotated in the literature, even between mouse brain atlases. Currently, there are four parcellation schemes for the mouse CLA. In one, the CLA is split into the dorsal and ventral subdivisions underneath the gustatory and agranular insular areas, respectively (Dillingham et al., 2019; Paxinos and Franklin’s, 2019). In a second scheme, the CLA is delineated largely underneath the gustatory, agranular insular, and visceral areas, but not subdivided. In a third scheme, the CLA is divided into core and shell subdivisions (Atlan et al., 2017, 2021; Binks et al., 2019; Erwin et al., 2021; Graf et al., 2020; Marriott et al., 2020; Real et al., 2003, 2006). A fourth scheme, advocated for here, avoids any core/shell or dorsal/ventral subdivision and defines the CLA as a single, densely packed group of neurons embedded in the agranular insular area (Dillingham et al., 2017; Wang et al., 2017, 2020). Reconciling the differences among these four anatomical definitions is necessary for correctly interpreting and quantifying tract-tracing data as well as electrophysiological recording and opto/chemogenetic manipulations.

Over the past several decades, the afferent and efferent connections of the CLA have been extensively studied in mammals (Crick and Koch, 2005; Druga, 2014; Edelstein and Denaro, 2004; Jackson et al., 2020; Smith et al., 2020). As in human, non-human primate and cat (Fernández-Miranda et al., 2008; Macchi et al., 1983; Milardi et al., 2015; Olsen and Graybiel 1980; Pearson et al., 1982; Reser et al., 2017; Torgerson et al., 2015), the CLA in mouse is reciprocally connected with frontal, parietal, temporal and occipital cortices (Atlan et al., 2018; Narikiyo et al., 2020; Wang et al., 2017; Zingg et al., 2018). These connections are organized in a crude topography and are stronger with the prefrontal cortex than the sensory cortex. Cortical inputs to the CLA are bilateral, with an ipsilateral predominance, except the cingulate and secondary motor cortices whose inputs are stronger to the contralateral CLA than to the ipsilateral one (Wang et al., 2017; Zingg et al., 2018). Additionally, the CLA has been reported to receive inputs from other brain regions, including olfactory areas, cortical subplate, striatum, pallidum, hippocampal formation, thalamus, hypothalamus, midbrain, and pons (Atlan et al., 2018; Narikiyo et al., 2020; Zingg et al., 2018).

The inputs to the CLA are predominant from cortical pyramidal neurons (Atlan et al., 2017; Wang et al., 2017; Zingg et al., 2018). These are classified into three major cell types: intra-telencephalic (IT) neurons in layer (L) 2-L6, extra-telencephalic (ET) neurons in L5, and corticothalamic (CT) neurons in L6, based on their downstream targets (Shepherd et al., 2013; Peng et al., 2021). However, a few studies have reported specific cell types involved in cortico-claustral circuits. In the primary visual cortex (VISp), one group of L6 pyramidal neurons projects to the dorsolateral geniculate nucleus (LGd) in the thalamus and another group to the CLA (Katz, 1987). These two groups of L6 neurons have different dendritic morphologies, projection targets, and intrinsic electrical properties, and belong to two different cell types: L6 CT neurons projecting to LGd and L6 IT neurons projecting to CLA (Baker et al., 2018; Cotel et al., 2018). Different from the VISp, in the secondary motor cortex (MOs), L2/3 IT and L5 IT neurons project to the CLA but L5 ET and L6 CT neurons do not (Smith et al., 2016). Given that different cortical cell types convey different kinds of information (Economo et al., 2018; Kim et al., 2015), comprehensively understanding cell-type-specific connections of the mouse CLA will facilitate its functional study in the future.

The CLA is composed of both excitatory and inhibitory neurons (Marriott et al., 2020; Graf et al., 2020). Its excitatory neurons (also called principal neurons) have recently been classified into four or five cell types based on their intrinsic electrical properties (Chia et al., 2017; Graf et al., 2020) and into four types based on axonal projections of fully reconstructed single CLA neurons (Peng et al., 2021). How its morphological cell types are related to the electrophysiological ones is unclear. CLA inhibitory neuron types expressing parvalbumin (Pvalb), somatostatin (SST), or vasoactive intestinal peptide (VIP) are distinguishable in their distribution patterns and intrinsic electrical properties (Graf et al., 2020). The Pvalb neurons are denser than the other two inhibitory cell types and can be activated by cortical input and send feedforward inhibition to principal neurons within the CLA (Day-Brown et al., 2017; Kim et al., 2016). How axons of CLA inhibitory neurons arborize is not known.

To address these issues, we first confirmed accurate delineation of the boundary of the CLA in the Allen 3D mouse brain atlas based on multimodal anatomical reference datasets (Wang et al., 2017, 2020). We then characterized the spatial position, brain-wide connectivity patterns, and full morphology of single principal neurons of the mouse CLA using systematic labeling, whole-brain imaging, and informatics approaches. Specifically, we performed new experiments by injecting monosynaptic retrograde rabies viral tracer into the CLA of the transgenic mice and combined these with complementary datasets on CLA outputs and inputs mapped with anterograde Cre-dependent AAV tracer injections from the Allen Mouse Brain Connectivity Atlas (Oh et al., 2014; Harris et al., 2019). We reconstructed full morphologies of individual single CLA principal neurons in sparsely labeled transgenic mice. We analyzed axonal arborization of CLA inhibitory neurons by injecting Cre-dependent anterograde AAV tracer into the pan-inhibitory neuron transgenic mouse, Gad2-IRES-Cre. By integrating all these datasets, our study provides a detailed wiring diagram of the afferent and efferent projections of the mouse CLA at regional, laminar, and cell type levels.

The companion paper (McBride *et al*. 2022) studies the function of these CLA projection neurons in the same transgenic mouse lines by optogenetically exciting them and recording postsynaptic neurons in multiple cortical areas using Neuropixels electrodes.

## Results

### Anatomical delineation of the mouse CLA and viral tracer injections

To correctly interpret and quantify tract-tracing data, we first confirmed accurate delineation of the boundary of the mouse CLA using multimodal anatomical reference datasets, including geno-, cyto-, myelo-, chemo-architecture and connectivity in our previous studies (Wang et al., 2017, 2020). All these reference datasets are openly accessible at our web portal (http://connectivity.brain-map.org). One of the key data modalities for CLA annotation is whole-brain serial two-photon tomography (STPT) imaging data of tdTomato reporter transgene expression in Cre driver transgenic mouse lines. Six of them have been used in our previous publication (Wang et al., 2017).

We here identified four new Cre driver lines with enriched gene expressions in the CLA but with less expression in adjacent structures; Esr2-IRES2-Cre, Tacr1-T2A-Cre, Gnb4-IRES2-CreERT2, and Grp-Cre_KH288 (**Figure 1A-1D**). The CLA is bordered dorsally by L6b of the gustatory cortical area (GU) (**Figure 1B and 1C**), ventrally by the dorsal endopiriform nucleus (EPd) (**Figures 1C, S1E, S1I and S1K**), laterally by L6 of the agranular insular cortex (AI) (**Figure S1K and S1L)** and medially by L6 of AI and the fiber bundle making up the external capsule (**Figure S1J and S1K**). Three other Cre driver lines Htr1a-IRES-Cre, Dlg-Cre_KG118 and Fezf2-CreER have enriched expressions in L6 of the isocortex, tapering off from GU to AI with little or no expression in the CLA (**Figures S1A-S1C and S1E-S1G**). The CLA is a more densely packed group of neurons compared to adjacent structures in the Nissl- (**Figure S1I**) and immunohistochemically (antibody against NeuN) stained specimens (**Figure S1K**) and in the transgenic mouse line Rorb-IRES2-Cre Ai110 with red nuclear-labeling (**Figure S1L**). The CLA is surrounded by densely labeled myelin fibers revealed by immunostaining with an antibody against SMI-99 (basic myelin protein, in red) (**Figure S1J**) and an antibody against NF-160 (Neurofilament-M, in green) (**Figure S1K),** a characteristic used to define the CLA in mammals (Pham et al., 2019; Wang et al., 2017). Similarly, it is also surrounded by more darkly stained AchE-positive fibers (data not shown but see the Allen Reference Data portal http://connectivity.brain-map.org/static/referencedata/experiment/thumbnails/100140478?image_type=ACHE&popup=true). Our connectivity data show dense projections to the CLA from cortical and subcortical structures such as the anterior cingulate area (ACA) (**Figure 3I and 3J**) and anteromedial thalamic nucleus (AM) (**Figure 4Q**). Because all anatomical reference datasets converge to the same conclusion, we believe that our current and previous studies accurately delineated the boundary of the mouse CLA in 3D (Wang et al., 2017, 2020).

**Figure 1.**
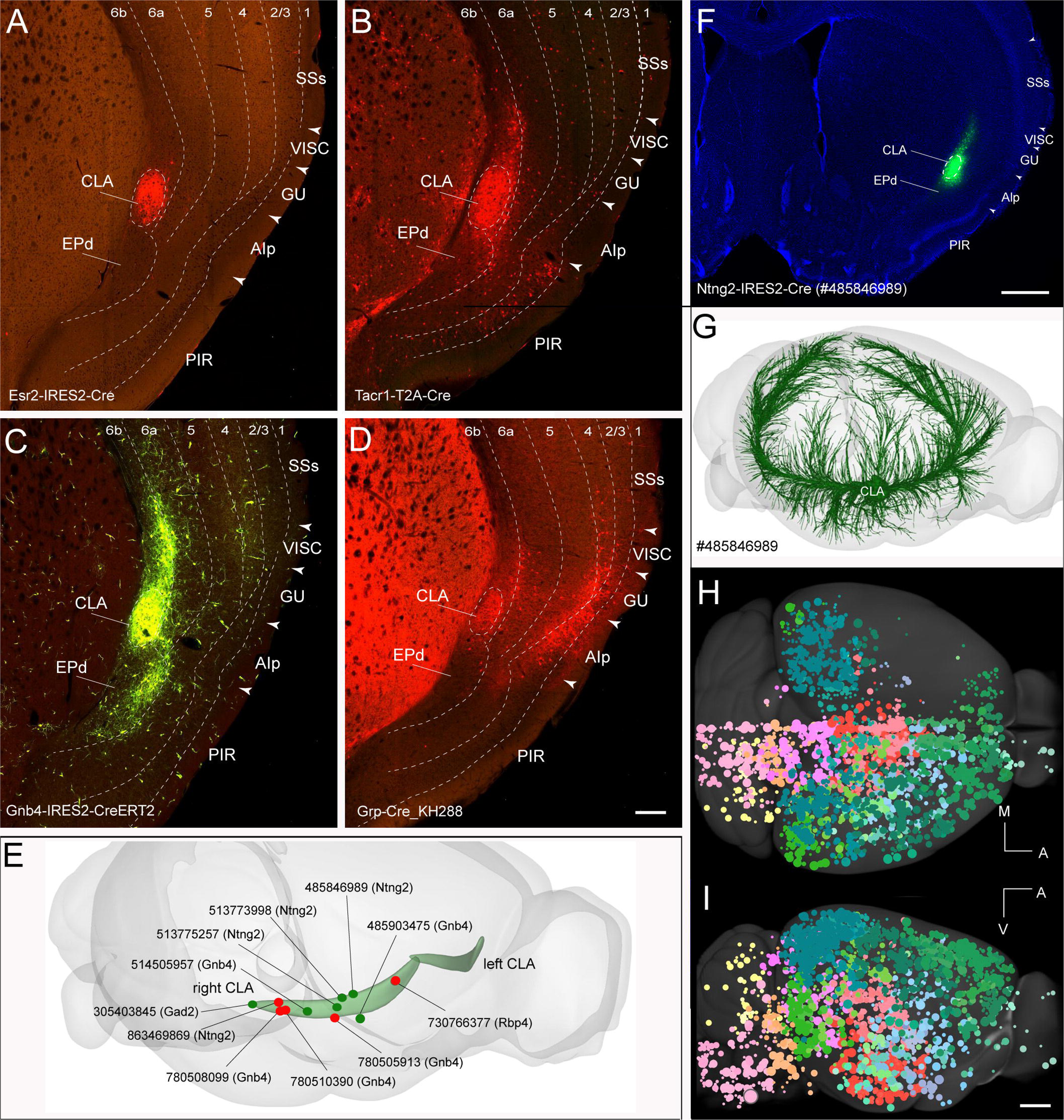
Delineation of the boundary of the mouse CLA and coverage of injection sites. (A-D) Four transgenic lines Esr2-IRES2-Cre (A), Tacr1-T2A-Cre (B), Gnb4-IRES2-CreERT2 (C) and Grp-Cre (D) have enriched gene expressions in the CLA but far less in its adjacent structures. Arrowheads indicate the borders between cortical areas. Dashed lines indicate the borders of the CLA and between cortical layers. These labels are the same in F. For abbreviations, see Supplementary Table 7. Scale bar, 150 μm. (E) Injection sites in the CLA (curved light green on both hemispheres) are shown in rotated lateral view with retrograde rabies tracer (in red dots) and anterograde Cre-dependent AAV tracer (in green dots). The dot sizes are not adjusted to the actual injection sizes. All these injection sites were made in the right side of the CLA. (F) A representative example of anterograde Cre-dependent AAV (in green) injections in the CLA counterstained with DAPI (in blue). Scale bar, 500 μm. (G) CLA output to the whole brain from injection in F is shown in the rotated dorsolateral view. (H-I) Cortical and subcortical injections into transgenic mouse lines and wild-type mice with anterograde viral tracer AAV are shown in the dorsal (H) and lateral (I) views. Different color dots represent injections in different major brain divisions, with their diameters proportional to the actual sizes of injection sites. A, anterior; M, medial; V, ventral. Scale bar, 1 mm.

Annotation of injection sites is a critical step for correctly interpreting tract-tracing data (see Methods for details). Anterograde (AAV) and retrograde (rabies virus) tracing experiments injected into the CLA were carefully inspected for targeting and infection results. We selected 5 out of 14 retrograde injections and 6 out of 9 anterograde injections for further analysis as they are spaced out along different anterior-posterior locations illustrated in the rotated lateral view (**Figure 1E**). A representative example of anterograde Cre-dependent AAV injections into the CLA of the transgenic mouse is shown (**Figure 1F)**: here, the informatically reconstructed axon skeletons from the injection site travel forward and backward to cortical and subcortical targets and are densely arborized in the ipsilateral midline and higher association cortical areas and retrohippocampal region, but sparsely in the sensory cortices (**Figure 1G**).

To validate retrograde tracing results and to fully characterize the cortical cell types projecting to the CLA, we also systematically reviewed for inclusion of more than 2,000 AAV injection experiments from the Allen Mouse Brain Connectivity Atlas dataset (http://connectivity.brain-map.org<u>, Oh et al., 2014, Harris et</u> <u>al., 2019</u>). This dataset consists of ∼1,000 isocortical tracing experiments injected in 40 of 43 isocortical areas, and ∼1,000 experiments injected with AAVs expressing GFP in 175 structures across 11 other major brain divisions of wild type and Cre driver transgenic mouse lines. The coverage of all these anterograde injections is illustrated in the dorsal and lateral views of the Allen 3D mouse brain template (**Figure 1H and 1I**). All anterograde and retrograde viral tracing experiments used in the current and previous CLA studies are listed in **Supplementary Table 1**.

### Whole-brain inputs to the CLA revealed with retrograde and anterograde viral tracing

To map whole-brain inputs to the CLA, we used the monosynaptic retrograde cell-type-specific rabies tracing method by injecting a Cre-dependent AAV helper virus (AAV1-DIO-TVA66T-dTom-N2cG) expressing EnvA TVA receptor, tdTomato and rabies glycoprotein, followed by injection of EnvA pseudotyped, glycoprotein-deleted rabies virus (RV-CVS-N2CdG-H2B-EGFP) expressing nuclear-localized EGFP (Yao et al., 2021) into the same location in the CLA of the transgenic mice Gnb4-IRES2-Cre, Ntng2-IRES2-Cre and Rbp4-Cre-KL100. These transgenic mouse lines have enriched transgene expressions in the CLA. The two sequential injections resulted in retrograde nuclear-EGFP labeling of the presynaptic neurons that form synaptic contacts with the starter cells infected with both Cre-dependent AAV helper virus and rabies virus. All starter cells were manually counted in each experiment (see details in the Methods). Most of the starter cells were found in the CLA and small numbers in its adjacent structures (**Supplementary Table 2)**. A representative example of the retrograde rabies injection sites is shown in the coronal section (**Figure 2A**). To enhance visualization of AAV helper virus-infected neurons, we performed immunostaining with an antibody against tdTomato and found red neurons labeled mostly in the CLA (**Figure 2B**). The presynaptic neurons were labeled in green (**Figure 2C**) and starter cells in yellow (**Figure 2D**). Starter cells were labeled mostly in the CLA but sparsely in the GU and EPd (**Figure 2D**). The presynaptic neurons to the CLA in the whole brain are shown in the rendered 3D images at three axes (**Figure 2E**) and are found more frequently in the ipsilateral than in the contralateral hemisphere (**Figure 2E and 2F**). Although the total numbers of starter cells vary more than 15 times from the most to the least labeled, overall presynaptic labeling patterns appeared similar across all five injection cases (**Figure 2F**). The fraction of total fluorescent signal detected as a proxy for presynaptic neurons in whole brains (n=5) is the highest in isocortex (60%), moderate in olfactory areas (14%), hippocampus (8%), cortical subplate (10%) and thalamus (6%), low in the striatum (1%) and pallidum (1%) and extremely low in hypothalamus, midbrain, pons, medulla, and cerebellum (all together less than 1%) (**Figure 2G**).

**Figure 2.**
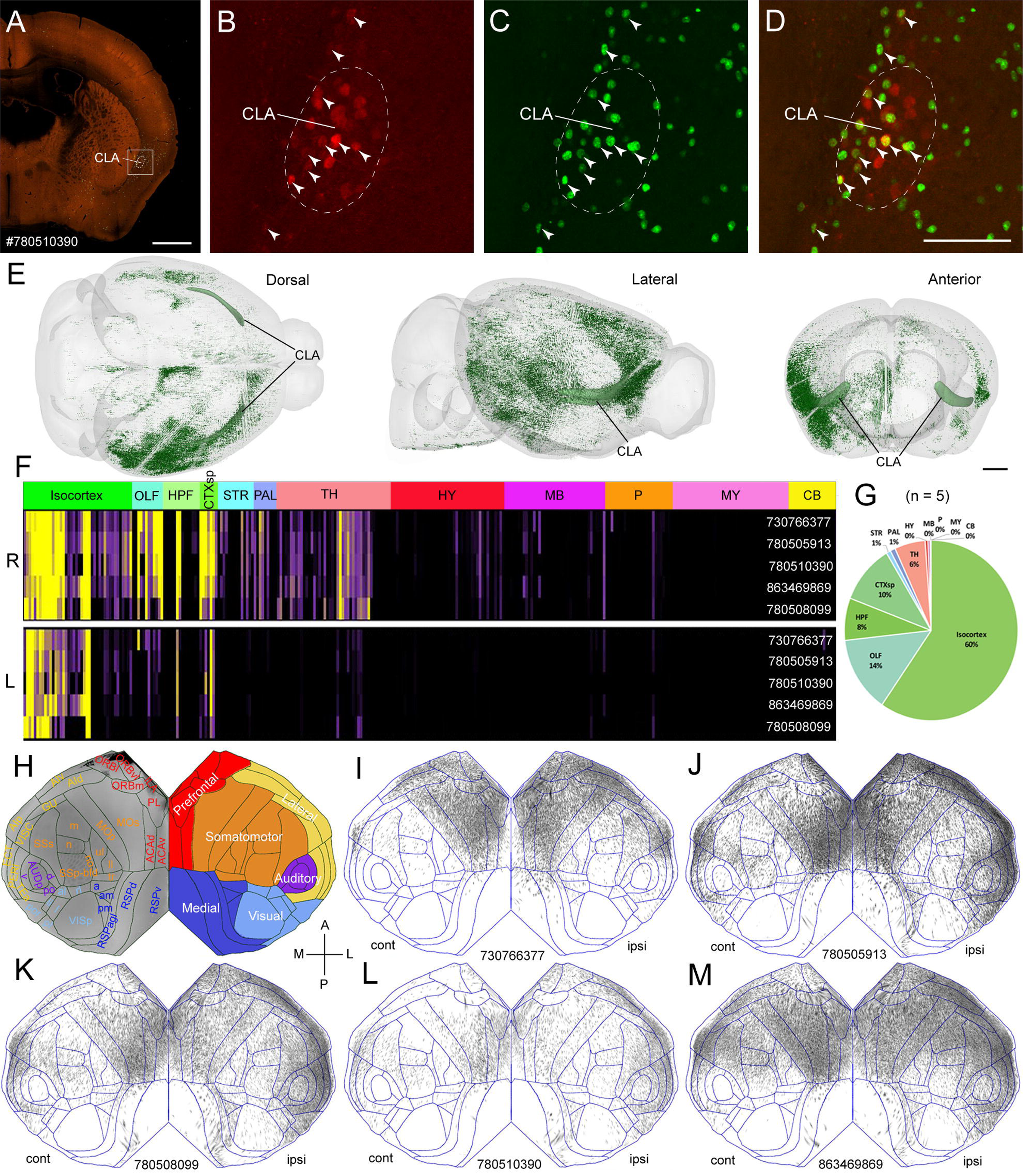
Whole-brain inputs to the CLA revealed with monosynaptic retrograde rabies tracing. (A) One representative example of retrograde rabies injection sites in coronal view. Scale bar, 1 mm. (B-D) An enlargement of rectangle in A is shown in high power image of injection site in red, green and merged channels. Neurons were infected with the AAV helper virus in red (B) and with rabies virus in green (C). The starter cells infected with both rabies and AAV helper virus are in yellow (D). The dashed lines indicate the border of the CLA. Arrowheads indicate starter cells. Scale bar, 200 μm. (E) Brain-wide presynaptic inputs to the CLA are shown in the dorsal, lateral and anterior views. The curved light green represents the CLA and dark-green dots represent segmented presynaptic neurons. Scale bar, 1 mm. (F) The presynaptic labeling matrix shows the fraction of total fluorescence signal in individual whole brains normalized by the starter cell and structural volume. Each experiment ID is shown on the right side along the Y-axis. Twelve major brain divisions at the top row are color-coded differently. R, right hemisphere; L, left hemisphere. For abbreviations, see Supplementary Table 7. (G) Pie chart shows the proportion of whole-brain presynaptic input to the CLA from 12 major brain divisions. (H) All iscocortical areas and the six cortical modules are labeled and are color-coded differently on the left side and right side of the isocortical flatmap, respectively. A, anterior; P, posterior; M, medial; and L, lateral. (I-M) The isocortical flatmaps reveal similar presynaptic labeling patterns across 5 CLA injection cases. The injections are arranged from the anterior (I) to the posterior (M). Each black dot in the flamap represents a presynaptic neuron. “ipsi” and “cont” stand for ipsilateral and contralateral respectively.

We further analyzed presynaptic neurons in isocortex and other major divisions at the fine structural level of the mouse ontology (316 structures in each hemisphere). In the isocortex, distribution of presynaptic neurons can be visualized in the flatmap by projecting them to the cortical surface along curved streamlines. In the flatmap, all isocortical areas and subdivisions were labeled on the left side (**Figure 2H)** and six cortical modules (prefrontal, lateral, medial, somatomotor, visual, and auditory) were color-coded on the right side (**Figure 2H**) (Harris et al., 2019). Presynaptic neurons were observed more in the ipsilateral prefrontal and lateral modules than in the medial, visual, somatomotor and auditory modules in the flatmaps (**Figures 2I-2M**). The contralateral labeling looks like a mirror image, but with fewer presynaptic neurons, except in the anterior cingulate (ACA), prelimbic (PL) and secondary motor areas (MOs) whose presynaptic neurons are more on the contralateral than on the ipsilateral side (**Figure 2I-2M**). This observation is confirmed by the quantitative analysis (**Figure S2**).

We found a rough topography of cortical presynaptic inputs to the CLA along the anteroposterior axis of the CLA. The anterior CLA injection (**Figure 2I**) shows strong bilateral labeling in the prefrontal module, with less or no labeling in the RSP, visual areas and the posterior part of the lateral module. By contrast, the posterior CLA injections show substantial presynaptic labeling in the RSP, visual areas and the posterior part of the lateral module in addition to strong presynaptic labeling in the prefrontal and anterior part of the lateral modules (**Figure 2J-2M**).

To reveal the laminar distribution of presynaptic neurons, we plotted the average pixels of segmented presynaptic neurons in each layer in individual cortical areas across the five injections (**Figure S3A**). Presynaptic neurons are denser in infragranular than in supragranular layers in both ipsilateral and contralateral cortical areas except one (ORBvl). Even in the infragranular layers, we found that most cortical areas contain more presynaptic neurons in L5 than in L6, while a few cortical areas show the opposite distribution pattern: more presynaptic neurons in L6 than in L5 (**Figure S3A**). **Figure 3A** shows an example of whole-brain presynaptic inputs to the CLA. In this injection, presynaptic neurons are denser in L5 than in L2/3 and L6 in the ipsilateral medial orbital area (ORBm), PL, infralimbic (ILA), ACA, MOs, primary motor area (MOp), posterior agranular insular area (AIp), visceral area (VISC), primary somatosensory area (SSp), supplemental somatosensory area (SSs), temporal association area (TEa), laterointermediate visual area (VISli), perirhinal area (PERl), ectorhinal area (ECT), ventral auditory area (AUDv) and primary auditory area (AUDp) (**Figure 3B-3F**), while they are denser in L6 than L5 in the primary visual area (VISp) and lateral visual area (VISl) (**Figure 3G**). There was no presynaptic labeling in L4 across any cortical areas (**Figure 3F-3G**).

**Figure 3.**
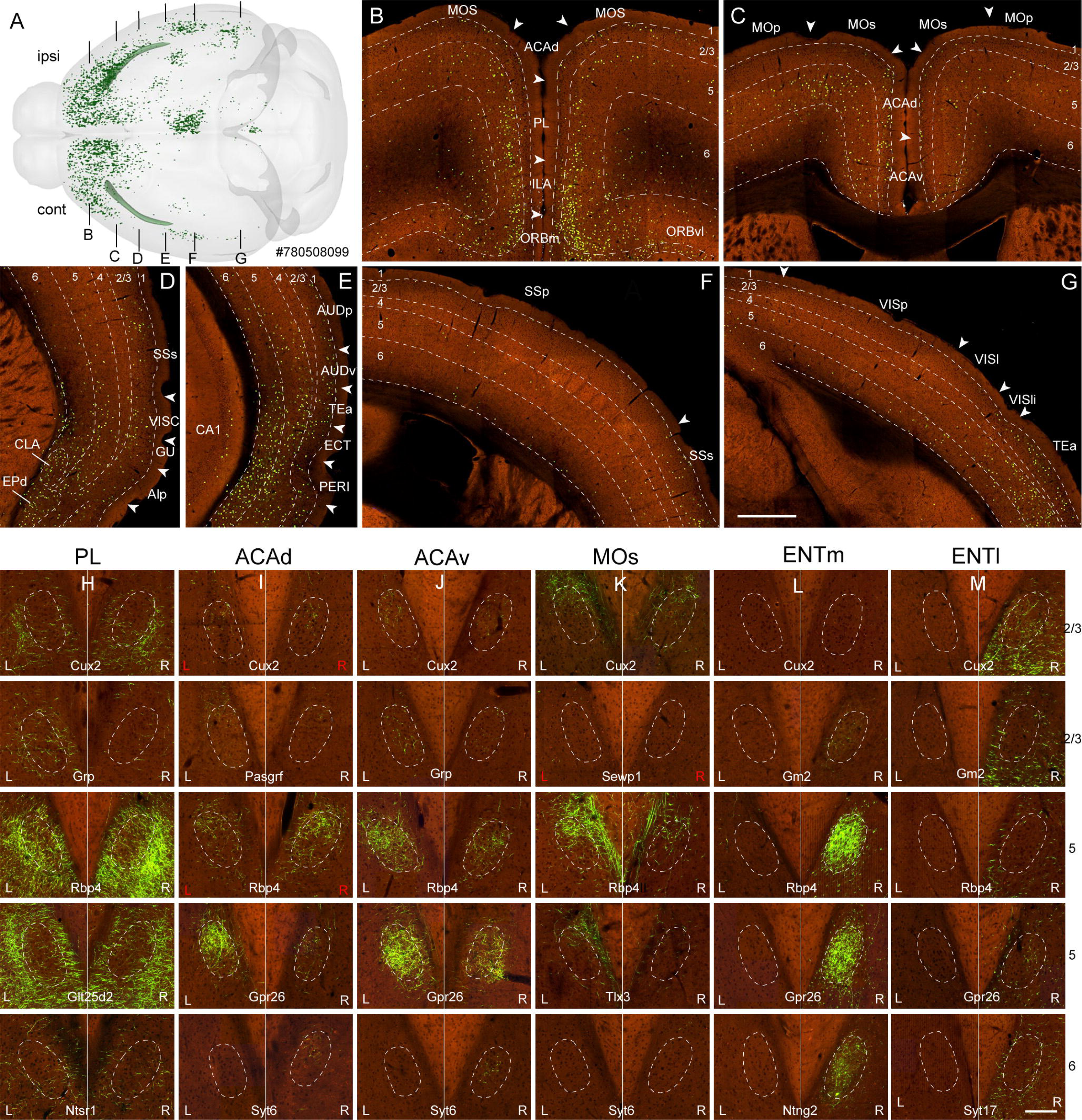
Laminar origin of pyramidal neurons and cell-type-specific cortical inputs to the CLA. (A) A dorsal view of a representative retrograde rabies injection shows whole-brain inputs to the CLA. Green dots represent presynaptic neurons. Black strip lines indicate approximately section levels in B-G. “ipsi” and “cont” stand for ipsilateral and contralateral respectively. (B-G) High power images show presynaptic neurons in frontal (B and C), temporal (D and E), parietal (F), and occipital (G) cortical regions. Arrowheads indicate the borders between cortical areas. Dashed lines indicate the borders of the cortical layer, CLA and EPd. For abbreviations, see Supplementary Table 7. Scale bar, 500 μm. (H-M) High power images showing axonal terminals in the CLA and its adjacent structures from injections into cell-type-specific layers 2/3, 5, and 6 of cortical areas PL (H), ACAd (I), ACAv (J) and MOs (K) and retrohippocampal regions ENTm (L), and ENTl (M) of the transgenic lines. The dashed line represents the border of the CLA. L, left; R, right. Color-coded L (in white) and R (in red) indicate injection sites in right and left cortical areas, respectively. Scale bar, 200 μm.

To characterize the cortical cell types projecting to the CLA, we analyzed anterograde Cre-dependent AAV injections labeling cell-type/class-specific neurons defined by the transgenic mouse lines (**Supplementary Table 3**) (Harris et al, 2014, 2019). We found that the prefrontal and lateral modules have more cell types (L2/3 IT, L5 IT, L5 IT ET, and a subset of L5 ET and L6 CT neurons) with denser projections to the CLA than the sensory cortical modules (L2/3 IT, L5 IT, L5 IT ET, L6 IT neurons) with few exceptions (**Figure S3B**). **Figure S4** shows examples of injection sites, in which the infected neurons are either in pan-layers or in L2/3 IT, L5 IT, L5 IT ET, L5ET and L6 CT of six cortical areas (PL, ACAd, ACAv, MOs, SSp and VISp). Injections in these cortical areas show two different projection patterns to the CLA and its adjacent structures. Some of them send denser projections to the surrounding structures than to the CLA (**Figure 3H**) and others do the opposite (**Figure 3I-3K**). The cortical areas PL, ACAd, ACAv and MOs sending strong projections to the CLA were selected as representative examples. In the PL, L5 IT (including L5 IT ET) neurons send denser projections to bilateral CLA than L2/3 IT, a subset of L5 ET, and L6 CT neurons with a bias toward deep layers of the AI over the CLA (**Figures 3H and S3B**). In the ACAv and ACAd, L5 IT neurons send denser projections to bilateral CLA than L2/3 IT, a subset of L5 ET and L6 CT, and L6 local neurons, with a contralateral predominance (**Figures 3I, 3J and S3B**). In the MOs, L5 IT neurons send denser projections to the dorsal part of bilateral CLA than L2/3, a subset of L5ET and L6 CT, and L6b local neurons with a contralateral predominance (**Figures 3K and S3B**). In contrast, injections into specific layers of MOp show no projections to the CLA (**Figure S3B**). In the RSPagl, L5 IT neurons send denser projections to bilateral CLA than L2/3 IT neurons with an ipsilateral dominance. In the RSPd, L5 IT neurons project to the ipsilateral CLA. In the RSPv, L5 IT neurons send denser projections to the CLA than L2/3 IT and L5 ET neurons, with an ipsilateral predominance (**Figure S3B**). In the SSp and SSs, L5 IT neurons send sparse projections to the ipsilateral CLA, but with exception of the subdivisions (SSp-n and SSp-ul) whose projections do not terminate in the CLA (**Figure S3B**). In the VISp and VISl, L6 IT neurons dominate projections to the CLA. In contrast, in higher visual areas, L5 IT neurons send denser projections to bilateral CLA than L2/3 IT neurons with an ipsilateral predominance (**Figure S3B**). In the auditory cortical areas, L5 IT neurons send denser projections to the CLA than L2/3 IT neurons with an ipsilateral predominance (**Figure S3B**). L5 ET and L6 CT neurons of the sensory cortical areas do not project to the CLA but with an exception, the VISpor (L6 CT, Syt6-Cre_KI148) whose sparse projections to the ipsilateral CLA. The injections made into interneurons and L4 neurons in cortical areas of the transgenic lines show no projection to the CLA (**Supplementary Table 3**). It should be noted that each transgenic line was selected for injecting certain cortical areas ranging from the most 33 cortical areas to the least 3 cortical areas. Due to the lack of subclass-specific transgenic lines (e.g., L6 IT) and also lack of injections into certain cell-type-specific layers of some cortical areas such as ECT and PERl, their inputs to the CLA need to be explored in the future.

Outside of isocortex, we found presynaptic neurons labeled in many structures across 11 major divisions and verified 35 across 10 major brain divisions using our anterograde tracing data. These validated structures are listed in **Figure 4A**, including the anterior olfactory nucleus (AON), dorsal peduncular area (DP), taenia tecta (TT) and piriform area (PIR) of the olfactory areas, the ventral CA1, medial entorhinal area (ENTm) and lateral entorhinal area (ENTl) of the hippocampal formation, the CLA, dorsal endopiriform nucleus (EPd), lateral amygdala (LA) and basolateral amygdala (BLA) of the cortical subplate, the medial amygdalar nucleus (MEA) of the striatum, the diagonal band nucleus (NDB) of the pallidum, the posterior limiting nucleus (POL), posterior intralaminar nucleus (PIL), parataenial nucleus (PT), central medial nucleus (CM), paraventricular nucleus (PVT), reunion nucleus (RE) and AM of the thalamus, the dorsomedial nucleus (DMH), ventromedial nucleus (VMH), supramammilary nucleus (SUM), lateral hypothalamic area (LHA), ventral tuberomammillary nucleus (TMv) and posterior hypothalamic nucleus (PH) of the hypothalamus, the ventral tegmental area (VTA), peduncularpontine nucleus (PPN) and dorsal raphe (DR) of the midbrain, the pontine central gray (PCG), superior central nucleus raphe (CS), locus ceruleus (LC), nucleus incertus (NI) and nucleus raphe pontis (RPO) of the pons and the interposed nucleus (IP) of the cerebellar nucleus. Presynaptic neurons in some of these structures, especially in the midline structures, were labeled bilaterally with an ipsilateral predominance. The most densely labeled structure is the ipsilateral CLA itself. **Figure 4B-4J** shows a few examples of presynaptic input structures in each of the 9 major brain divisions, including the AON of the olfactory areas, the ventral CA1 of the hippocampal formation, the EPd, LA and BLA of the cortical subplate, the NDB of the pallidum, the RE, PVT, AM and CM of the thalamus, the SUM of the hypothalamus, the VTA and DR of the midbrain, the LC of the pons and the cerebellar nucleus, contralateral IP. These 15 presynaptic structures were verified by injecting anterograde viral tracer into them (**Figure 4K-4Y**). The injections of the AON, BLA, PVT, RE, CM and DR show axon terminals labeled in bilateral CLA with an ipsilateral predominance (**Figure 4K, 4O, 4R-4T and 4W**), while the injections of the ventral part of CA1, EPd, LA, NDB, AM, SUM, VTA and LC show axon terminals only labeled in the ipsilateral CLA (**Figures 4L-4N, 4P, 4Q, 4U, 4V and 4X**). Consistent with retrograde rabies tracing, anterograde injections into the IP show sparse projections only to the contralateral CLA (**Figure 4Y**). As cortical input, some of the subcortical structures (AM, PVT, SUM, NDB, CM and IP) send preferential inputs to the CLA over the AI, while others (AON, ventral CA1, EPd, LA, BLA, RE, VTA, DR and LC) do the opposite. We also observed these differential projection patterns between the lateral entorhinal area (ENTl) and medial entorhinal area (ENTm) injections (**Figure 3L and 3M**). Injections in ENTm show denser projections to the CLA than to the AI, while injections in the ENTl show denser projections to the AI and EPd than to the CLA. In the ENTm injections, L5 IT and L5 IT ET neurons send denser projections to ipsilateral CLA than L2/3 IT and L6 IT neurons (**Figure 3L**), while in the ENTl injections, L2/3 IT, L5 IT, L5 IT ET and L6 IT neurons send denser ipsilateral projections to the AI and EPd than to the CLA (**Figure 3M**). One anterograde injection into ENTm of the transgenic mouse Nos1-CreERT2 (ID# 299403823), an interneuron transgenic line, shows long-range projections to the CLA. All anterograde tracing experiments in the major brain divisions, outside of the isocortex, are listed in **Supplementary Table 4**.

**Figure 4.**
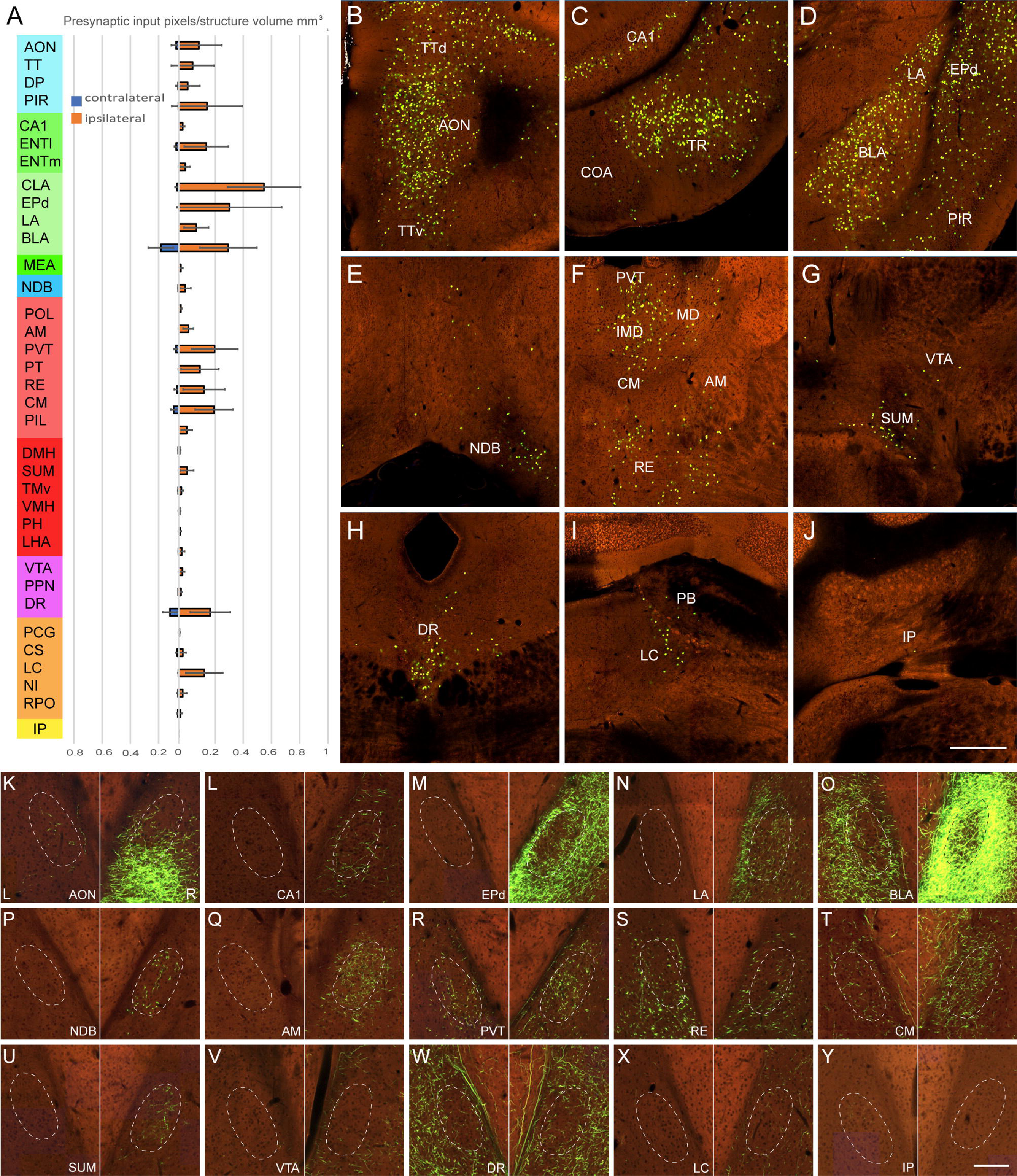
Inputs to the CLA from structures, outside of the isocortex, revealed with both retrograde and anterograde viral tracing. (A) A histogram showing the fraction of total presynaptic input pixels averaged from five injections. Along the Y-axis, ten major brain divisions are color-coded differently, including olfactory areas, hippocampal formation, cortical subplate, striatum, pallidum, thalamus, hypothalamus, midbrain, pons, and cerebellum. These presynaptic input structures were verified using anterograde AAV tracing results. Red and blue bars represent presynaptic labeling on the ipsilateral and contralateral sides, respectively. For abbreviations, see Supplementary Table 7. Bar histogram shows Mean ± SD (standard deviation). (B-J) High power coronal images showing examples of presynaptic neurons labeled in fifteen structures, including AON (B) of the olfactory areas, ventral CA1 (C) of the hippocampal formation, EPd, LA and BLA (D) of cortical subplate, NDB (E) of pallidum, AM, PVT, RE and CM (F) of thalamic nuclei, SUM (G) of the hypothalamus, VTA and DR (H) of the midbrain, LC (I) of pons and IP (J) of the cerebellum. Scale bar, 500 μm. (K-V) High power images of axon terminals in the CLA and its adjacent structures from injections into the above presynaptic fifteen structures, including AON (K) of the olfactory areas, ventral CA1 (L) of the hippocampal formation, EPd (M), LA (N) and BLA (O) of cortical subplate, NDB (P) of pallidum, AM (Q), PVT (R), RE (S) and CM (T) of the thalamic nuclei, SUM (U) of the hypothalamus, VTA (V) and DR (W) of the midbrain, LC (X) of pons and IP (Y) of the cerebellum. These injections were made in the right hemisphere. The dashed line represents the boundary of the CLA. Scale bar, 200 μm.

In addition to the validated structures, there are some unvalidated presynaptically labeled structures, most of which contain very few presynaptic neurons. These could be false positives due to contamination of injection sites and, to a lesser extent, lack of anterograde injections for verification. It should be noted that we used both anterograde and retrograde tracing data to validate each other. For example, presynaptic neurons were observed in the parabrachial nucleus (PB). However, anterograde injections into PB show no axonal terminals in the CLA but in its adjacent structures GU and AI (Grady et al., 2020; current study). Conversely, injections with anterograde tracer into the medial mammillary body (MM) show axon terminals in the CLA **(Supplementary Table 4**); yet we did not find presynaptic neurons in the MM from retrograde rabies injections into the CLA. The anterograde labeling in the CLA in the MM injections can be explained by contaminated SUM because injection tracks entering the MM must pass through the SUM, which does send projections to the CLA (Barbier et al.,2019; Zingg et al., 2018). There is another scenario. For example, retrograde injections into the CLA show presynaptic labeling in the POL. The POL is a very narrow thalamic nucleus, medially to the lateral posterior thalamic nucleus (LP) and suprageniculate nucleus (SGN) and laterally to the anterior pretectal nucleus (APN). Anterograde injections into the LP and SGN show projections to the CLA, which were only seen with POL contamination but not without it. By integrating both anterograde and retrograde datasets, we can conclude that the POL is the structure sending projections to the CLA, rather than the LP and SGN (**Supplementary Table 4**).

### CLA outputs revealed with anterograde AAV tracing and single neuron tracing

To reveal brain-wide outputs, we analyzed both bulk anterograde tracing and single neuron tracing datasets that are complementary to each other. The former reveals the union of projection targets from many infected neurons but cannot reveal the projection diversity of individual neurons. Furthermore, they can be contaminated by nearby neurons. The latter, on the other hand, can show projection diversity of individual neurons without the contamination issue but is limited in the number of fully reconstructed cells. A representative example of the bulk anterograde AAV injections into the CLA of the transgenic lines is shown in the dorsal and lateral views (**Figure 5A and 5B)**, with claustrocortical projections terminating in lower L1 to L6 with different densities across isocortical areas **(Figure 5C-5F**). Axons frequently ramify in lower L1, L2/3 and L5a and L6 but not in L4 and L5b. This qualitative observation is confirmed by our quantitative analysis results, which show that axons are denser in L2/3 and L5 than in other layers in most cortical areas (**Figure 5G**). In a few cortical areas the GU, VISC, SSs and SSp, axons are denser in L6 than in other layers. In the VISp, axons are denser in L2/3 than in other layers (**Figure 5F and 5G**). It should be noted that L5 is treated as one layer for quantification due to no sublayers being divided in the Allen Common Coordinate Framework (CCFv3). In the retrohippocampal region, claustral outputs are much denser in the deep layers than in the superficial layers of the ENTm and ENTI, while they are denser in the superficial layers than in the deep layers of the postsubiculum (POST), presubiculum (PRE) and parasubiculum (PAR), together called subiculum complex (**Figure 5F**). To reveal axonal distribution in the entire isocortex, we projected claustrocortical projections to the flatmap of the isocortex as in **Figure 2H-M**. In these flatmaps, we found similar projection patterns across these injections.

**Figure 5.**
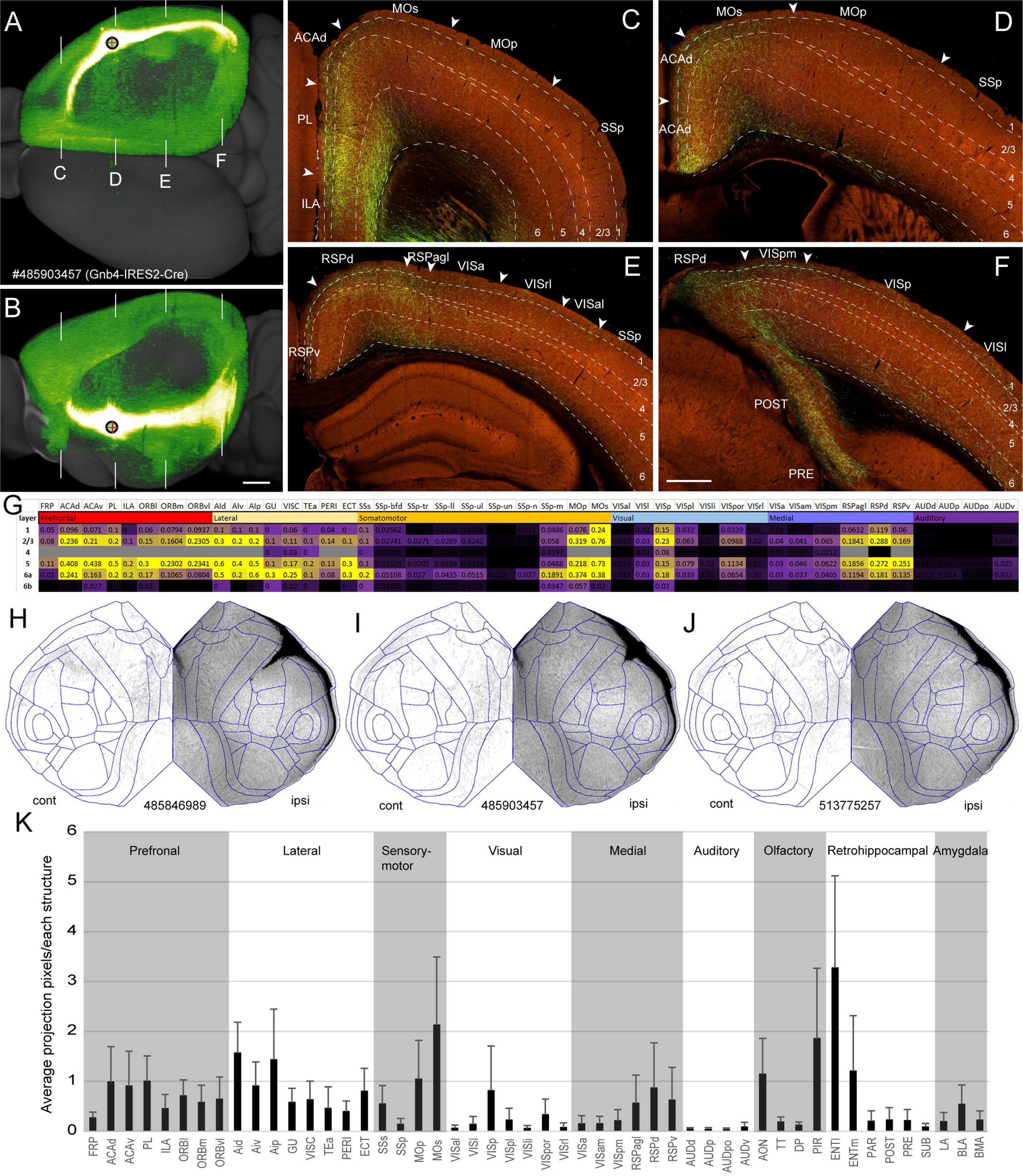
Efferent projections of the CLA revealed with bulk anterograde Cre-dependent AAV tracing. (A-B) A representative example of bulk anterograde Cre-dependent AAV injections into the CLA shows strong projections to the midline and association cortical areas in the dorsal (A) and lateral (B) views. White strips in A and B indicate coronal section levels shown in C-F. Scale bar = 1 mm. (C-F) High power images shows axonal terminals in different cortical areas at the frontal (C), parietal (D), parietal-occipital transition (E) and occipital (F) levels. Arrowhead indicates the border between cortical areas. Dash-line indicates the border between cortical layers. For abbreviations see supplementary Table 7. Scale bar, 400 μm. (G) Laminar distribution of claustrocortical projections in the ipsilateral cortical areas is shown by averaged axon pixels in each layer of individual cortical areas from five anterograde Cre-dependent AAV injections into the CLA. (H-J) The flamaps of the isocortex show similar claustrocortical projection patterns from three Cre-dependent anterograde AAV injection cases. “ipsi” and “cont” stand for ipsilateral and contralateral respectively. (K) A histogram shows averaged pixels of axonal projections in the ipsilateral isocortical and subcortical structures from five CLA injection cases. Mean = SD (Standard Deviation).

Claustrocortical projections were labeled in all ipsilateral isocortial areas, preferentially in the prefrontal and lateral modules and motor areas as well as in the VISp, SSs, RSPd and RSPagl, and sparsely in the higher visual, primary somatosensory and auditory cortical areas (**Figure 5H-5J**). Topography of claustral outputs to cortex can be seen between the anterior and posterior CLA injections. The anterior CLA injections (**Figure 5H**) show fewer projections to the RSPv compared with the posterior CLA injections (**Figure 5I and 5J**). The qualitative observation of claustrocortical projections is confirmed by our quantitative analysis results (**Figure 5K).** In addition to the isocortex and retrohippocampal region, claustral projections were found in the olfactory areas (AON, DP, TT and PIR) and cortical subplate (LA, BLA and BMA) (**Figure 5K**). Sparse contralateral projections were consistently found in the ACA, MOs and RSP (**Figure 5H-5J**). Due to the sparseness, we did not analyze the laminar distribution of claustrocortical projections in the contralateral cortical areas.

To reveal projection diversity, we reconstructed the full morphology of 52 single CLA principal neurons using sparsely labeled three transgenic Gnb4-IRES2-CreERT2 mice. 35 neurons of them were reconstructed in the left CLA and 17 neurons in the right CLA. For comparison and visualization purposes, we flipped the 35 neurons from the left to the right side. Their somas are distributed throughout the anteroposterior extent of the CLA (**Figure S5A**) and their dendritic arbors are very narrow mediolaterally without typical apical dendrites (Watakabe et al., 2014; Peng et al., 2021). These CLA neurons can be differentiated from neurons in the deep layers of its adjacent structure AI by both local and long-range projections (Peng et al., 2021). One such exemplar cell (**Figure S5B**) sends dense axonal projections to the midline cortical targets and ENT. Synaptic boutons in axon branches are present in the targets but barely in the axon shafts within the CLA. These axon shafts travel rostrally or caudally or in both directions, but with no, or only a few, local collaterals within the CLA (**Figures 6C-6K, S5C**). Individual neurons send projections to the diverse sets of cortical and subcortical targets with different projection densities and frequencies. The most densely and frequently targeted structures are the PL, ACA, MOs, RSP and VISp, and ENT (**Figure S5D-S5F**). Anterior CLA neurons (from 3.3 to 4.7 mm) send dense projections to the bilateral or ipsilateral midline cortical targets with fewer projections to the visual areas, subiculum complex and ENT, while posterior CLA neurons (5.6-6.6 mm) send dense projections only to the ipsilateral targets with substantial projections to the visual cortical areas, subiculum complex and ENT. Like the posterior neurons, the middle CLA neurons (4.7-5.6 mm) send projections to the ipsilateral cortical targets with projections to the ORB, PL and ILA (**Figure S5D**). The combined projections of these principal neurons recapitulate our bulk anterograde tracing results, showing CLA projections to all 43 ipsilateral cortical areas and to several contralateral cortical areas that mirror the strongly projected ipsilateral cortical ones, including ACA, MOs, and/or RSP (**Figure S6A**).

**Figure 6.**
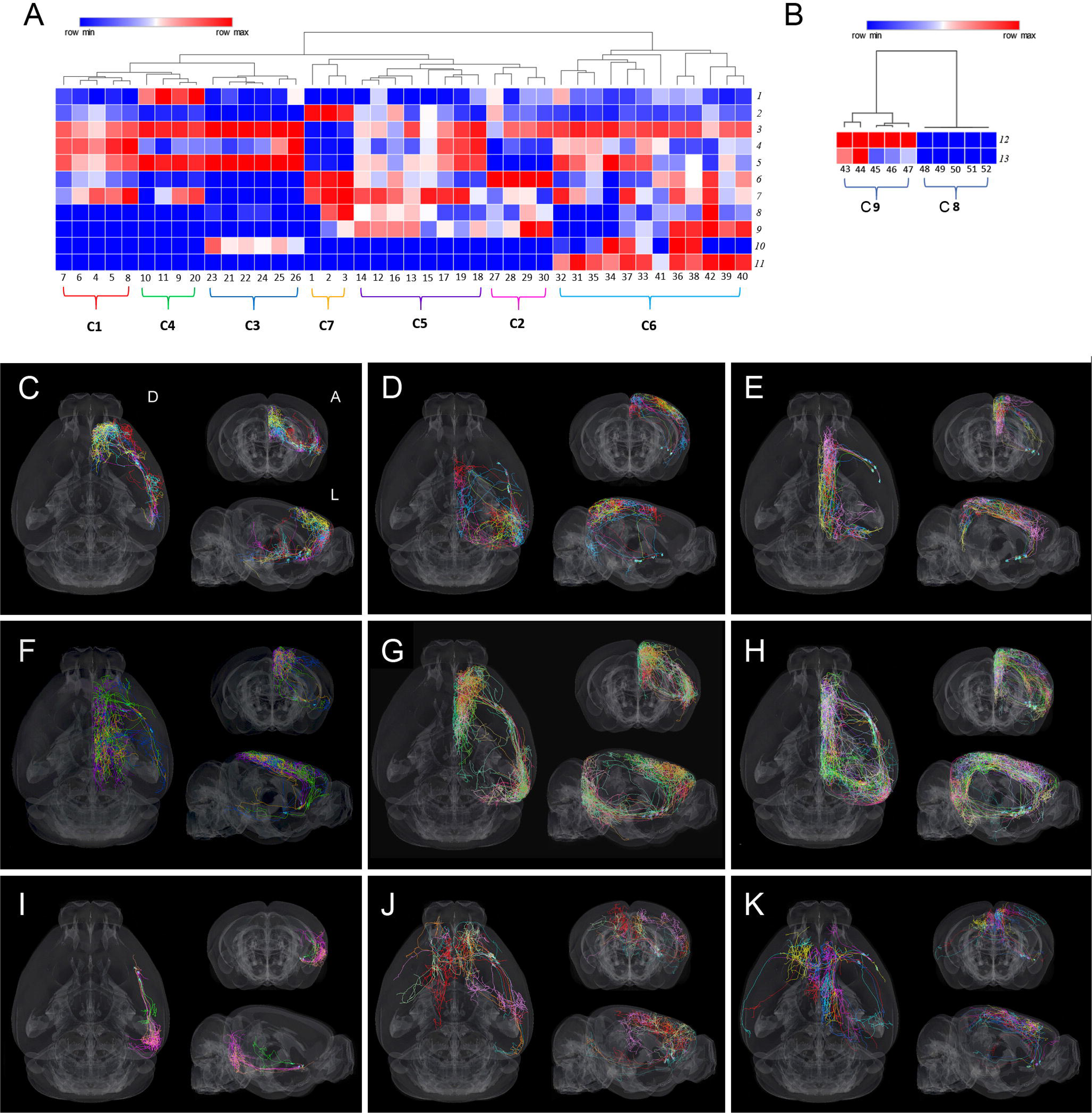
Diverse cell types of CLA principal neurons revealed by single neuron tracing. (A-B) Unsupervised hierarchical clustering analysis shows 7 clusters of ipsilateral projecting CLA neurons in (A) and 2 clusters of the bilateral projecting CLA principal neurons in (B). (C-K) Individual CLA principal neurons color-coded differently in each of the 9 clusters are shown in the dorsal, lateral and anterior views. D, dorsal; L, lateral; A, anterior.

To classify the morphological cell types, we first manually separated the 52 CLA principal neurons into two major classes: the ipsilateral (n = 42 neurons) and bilateral (n = 10 neurons) projecting classes. Using unsupervised hierarchical clustering analysis, we further divided the ipsilateral and bilateral projecting classes into 7 (**Figure 6A**) and 2 clusters (**Figure 6B**) based on 11 and 2 morphological features, respectively (see Methods for the details). All neurons in 9 clusters are matched to our manual annotation except a neuron (# 20), which is in cluster 4 but is manually annotated in cluster 5 (**Supplementary Table 5**). Neurons in each cluster are shown in the dorsal view of the brain (**Figure 6C-6K**). Since the dorsal view only shows 64% of the isocortex and leaves out cortical areas that receive strong and frequent inputs from the CLA not well visualized at the periphery, we projected axonal projections of individual neurons to the flatmap of the isocortex (**Figure S6B-S6J**). It should be noted that this flatmap shows CLA outputs to the entire isocortex but cannot visualize structures that are in other major brain divisions, such as the ENT, subiculum complex and amygdala.

In the ipsilateral projecting class, C1 neurons (n = 5) project to the prefrontal module and the lateral module densely, with weak projections to the somatosensory and auditory areas but do not project to the RSP, ENT and visual areas (**Figures 6C and S6B**); C2 neurons (n = 4) project densely to the visual areas (VISp, VISl, VISal and VISli) as well as to the RSP or ACAd, with sparse projections to the rest of the visual areas, somatosensory and auditory areas but do not project to the frontal areas (ORB, PL and ILA) (**Figures 6D and S6C**); C3 neurons (n = 6) send anteriorly a single axon shaft arborizing densely in the ACA, MOs and RSP, with sparse projections to the VISp, PL, ILA and subiculum complex but not to somatosensory, auditory and ORB areas (**Figures 6E and S6D**); C4 neurons (n = 4) send the anterior axon shaft arborizing densely in the ACAd, MOs and RSP, with sparse axonal terminals in the SSp, dorsal stream visual areas (VISa, VISam, VISpm) and VISp, and the posterior axon shaft arborizing sparsely in the PERl and ECT (**Figures 6F and S6E**); C5 neurons (n = 8) send the anterior axon shaft arborizing densely in the ACA, MOs and ORB, and extend to the anterior part of the RSP, with sparse projections to the SSp, and the posterior axon shaft arborizing densely in the ENT, but with weak projections in to the visual areas (**Figures 6G and S6F**); C6 neurons (n = 12), also called “crown neurons” (Peng et al., 2021), send the anterior axon shaft arborizing densely in the ACA, MOs and RSP, and the posterior axon shaft arborizing densely in the ENT and subiculum complex, with sparse terminals in the visual areas, ORB, PL, ILA, and somatosensory and auditory areas (**Figures 6H and S6G**); and C7 neurons (n = 3) only project posteriorly to the ENT and/or VISpor, TEa, PERl and ECT, but not to ACA, RSP and MOs (**Figures 6I and S6H**). In the bilateral projecting class, C8 neurons (n = 5) send the anterior axon shaft arborizing densely in the ipsilateral ACA and MOs, with sparse projections in ORB as well as in the contralateral ACA and RSP, or MOs, and MOp and SSp, but not in ipsilateral the RSP, and the posterior axon shaft arborizing sparsely in the SSp, SSs, auditory areas, or TEa and ENT (**Figures 6J and S6I**), and C9 neurons (n = 5) send the anterior axon shaft arborizing densely in the ipsilateral ACA, MOs and extends to the RSP as well as in the contralateral ACA, or MOs, or RSP, and the posterior shaft arborizing sparsely in the VISp and dorsal stream visual areas (VISal, VISrl, VISam, VISpm and VISa) (**Figures 6K and S6J**).

We used the violin plot to reveal the laminar distribution by averaging axon total lengths in each layer of cortical areas from all these single CLA neurons (**Figure S6K**). CLA axons terminate in the lower L1 - L6 across cortical areas but with different densities, similar to our bulk anterograde tracing results. In the frontal pole (FRP), PL, ORB, and TEa, axons are denser in L5 than in other layers. In the AI, GU and ECT, axons are denser in L6 than in other layers. In the ACA, MOs, and RSP, axons are dense across all layers. In the VISp, axons are denser in the lower L1 and upper L2/3 than in other layers. In the ENTl and ENTm, axons are denser in the infragranular layers than in the supragranular layers. We show examples of the laminar distribution of axons of individual CLA neurons in the strongest recipient structures PL, MOs, ACA, RPS, VISp and ENTl (**Figure S6L-S6P)**.

### Projections of interneurons within the CLA

To reveal axonal projections of interneurons, we analyzed one anterograde Cre-dependent AAV injection into the CLA of the pan-inhibitory neuron transgenic mouse, Gad2-IRES-Cre (**Figure S7A and S7B**). This injection site is in the posterior part of the CLA with contamination of nearby structures EPd and AI (**Figure S7C and S7D**). Axons derived from infected interneurons arborize along almost the entire anteroposterior extent of the CLA (**Figure S7E-S7J**). The inset in **Figure S7F** shows axons bearing *boutons en passage* within the CLA. Axons are densely arborized around the injection site and decrease density gradually with increasing distance from the injection site (**Figure S7E-S7J**). We also observed axons labeled densely in the EPd, ventral to the CLA, and in the deep layers of AI, but sparsely in its superficial layers (**Figure S7E-S7J**). It seems likely that axons in EPd and AI are derived from contaminated interneurons within EPd and AI, respectively. We did not find long-range axonal projections of CLA interneurons.

## DISCUSSION

In the current and in our previous studies, we accurately delineated the boundary of the mouse CLA (Wang et al., 2017, 2020). By systematically analyzing retrograde and anterograde viral tract-tracing data and fully morphology of single CLA principal neurons, we demonstrate that the CLA reciprocally connects with all cortical modules, preferentially with the prefrontal module (**Figure 7A and 7B**). It also receives inputs from at least 35 subcortical structures across 10 other major divisions and sends projections back to only a few of these (**Figure 7C and 7D**). We found that cell types projecting to the CLA are determined by cortical areas and layers. We classified CLA principal neurons into at least 9 cell types that innervate diverse sets of anatomically and functionally linked targets. Axons of interneurons arborize along almost the entire anteroposterior extent within the CLA. Altogether, this study provides a detailed wiring diagram of cell-type-specific afferent and efferent projections of the mouse CLA.

**Figure 7.**
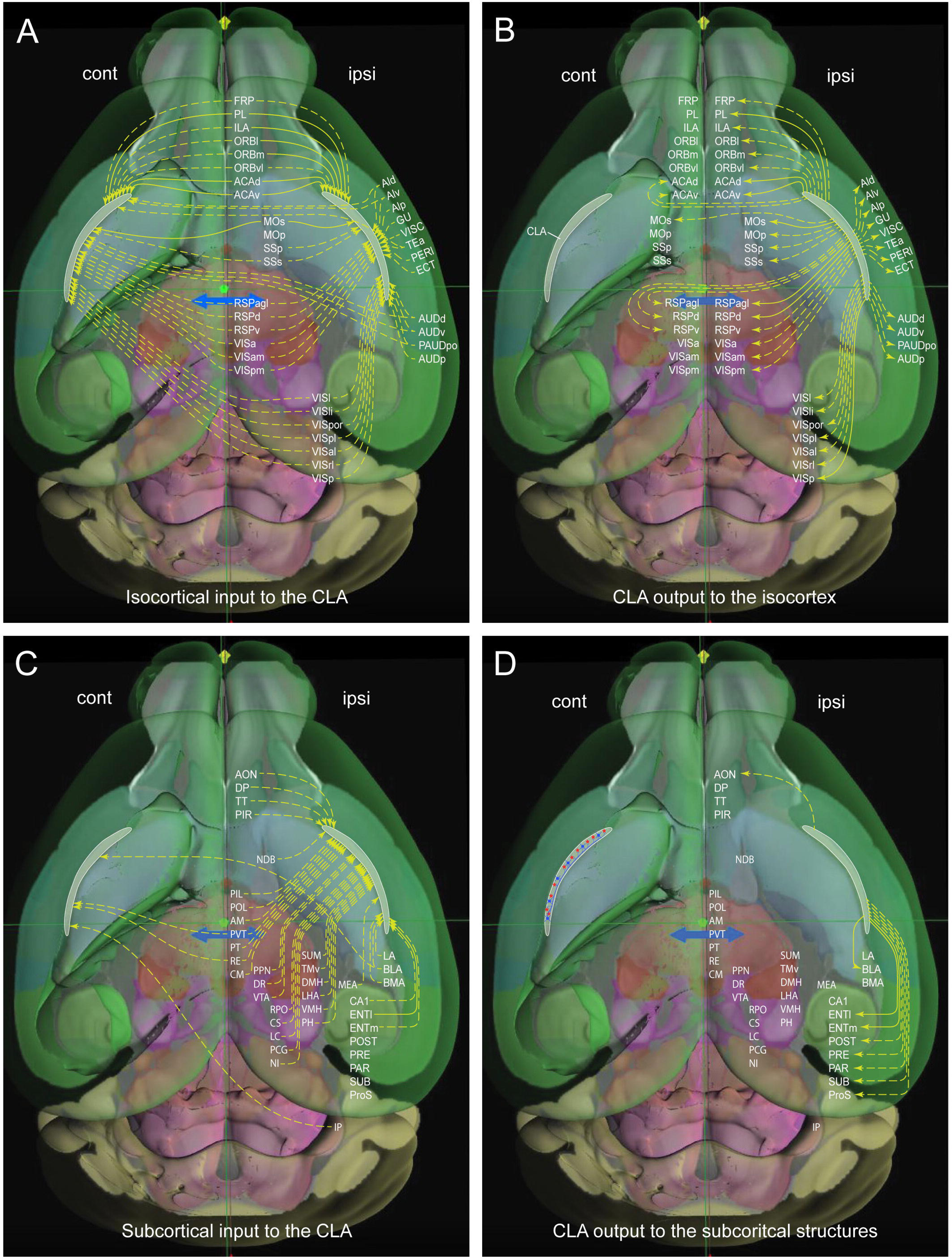
A wiring diagram of the input-output connections of the mouse CLA. (A-B) Cortical modular inputs to the mouse CLA (A) and CLA outputs to the modular cortical areas (B). Solid- and dash-arrows represent strong and weak projections, respectively. Major brain divisions were color-coded differently, with the CLA in grey. “ipsi” and “cont” stand for ipsilateral and contralateral respectively. These are the same in C and D. (C-D) Subcortical structures send inputs to bilateral, ipsilateral, or contralateral CLA (C), and only some of them receive inputs from the CLA (D). In D of right-side CLA, blue dots and red stars within the CLA represent inhibitory and excitatory neurons, respectively.

### Delineation of the boundary of the mouse CLA

How to accurately delineate the mouse CLA is open to debate. As described in the Introduction, there are four distinct parcellation schemes for mouse CLA, unlike in large mammals, such as human, non-human primate and cat whose CLA is easily identifiable as a thin sheet of densely packed neurons embedded between two fiber tracts, the *external* and *extreme* capsules (Baizer et al., 2014; LeVay and Shek, 1981; Reser et al., 2014). By registering multimodal anatomical reference datasets to the Allen 3D framework, our studies demonstrated that the mouse CLA is restricted to the “core” or “ventral” CLA and that the “shell” or “dorsal” CLA belongs to L6 of the AI and GU and the EPd (Wang et al., 2017, 2020). The CLA and its adjacent structures are different in terms of the cyto-, myelio-, geno-, chemo-architecture and connectivity. Such differences in connectivity, geno-architecture, and a combination of both have also been reported in recent studies but are interpreted as the “core” sending strong projections to the ACA and RSP and the dorsal and ventral “shell” sending strong projections to MOp and ENTl, respectively (Erwin et al., 2021; Marriott et al., 2020). This interpretation is different from ours. Our tract-tracing data show that the MOp and ENTl have weaker reciprocal connections with the CLA compared with L6 of the AI and GU, or L6 of the Al and the EPd, respectively. The marker genes, such as Pcp4, nnat, and slc30a3 used to define the “shell” of the CLA (Erwin et al., 2021), are preferentially expressed in L6 of the AI, or the EPd. That is, the “dorsal” or “shell” of the CLA should be interpreted as L6 of the insular cortex and the EPd as the mouse CLA lacks a clear extreme capsule. Our results show that the CLA and AI have differential connections with the ACA and ORB (Gehrlach et al., 2020; Park et al., 2012), which underlies their physiological and functional differences (Altan et al., 2021; Gogolla, 2017). Our definition of the mouse CLA meets anatomical criteria and is consistent with a developmental study, which uses gene expression patterns to mark only one group of neurons underneath of the AI (Watson and Puelles, 2017), and with a combination of tract-tracing and immunohistochemical studies (Barbier et al.,2019; Qadir et al., 2018; Suárez et al., 2018), similar to what has been used for rat CLA (Mathur et al., 2009, 2014). The accurate delineation of the boundary of the mouse CLA studies lays a solid foundation for a better understanding of its cell types, circuits, and functions as in the companion paper (McBride et al., 2022).

### The efferent and afferent connections between the CLA and isocortex

Precisely deciphering CLA circuitry is essential to understanding how information is transmitted to and from it. A few recent studies have systematically investigated the efferent and afferent projections of the mouse CLA by injecting anterograde and retrograde viral tracers (Atlan et al., 2018; Narikiyo et al., 2020; Zingg et al., 2018). There are discrepancies in terms of some input and output projections of the CLA between these previous studies. For comparison purposes, their quantitative tract-tracing results and ours are listed in **Supplementary Table 6**. For descending cortico-claustral projections, these reports are not only different in the absence of inputs from six cortical areas (FRP, MOp, SSp, GU, VISC and AIp) to four (FRP, SSp, GU and AI) to three (ILA, RSP and RSPagl) but also in preferential inputs from six cortical areas (ORB, ACA, PL, ILA, Ald and TEa/ECT/PERl) to three (ORB, MOp and MOs) to two (ORB and AI) (Atlan et al., 2018; Narikiyo et al., 2020; Zingg et al., 2018). In contrast, our retrograde data showed that the CLA receives inputs from all cortical modules, preferentially from the prefrontal and lateral modules. This finding was largely confirmed by our anterograde tracing results, but with sparse inputs from the SSp-bfd, SSp-ll, SSp-tr, SSp-m, AI, GU, VISC and RSPv to the CLA and no input from the MOp, SSp-n and SSp-ul. Our finding is in line with electrophysiological *in vivo* and *in vitro* studies, in which CLA neurons were activated mostly by the frontal input but barely by single sensory input (Chia et al., 2020; Reus-García et al., 2021), providing an anatomical basis that cortical areas with different projection strengths influence CLA neuron activity differently.

For ascending claustro-cortical projections, discrepancies are not only seen in the absence of outputs to six cortical areas (FRP, SSp, SSs, GU, VISC and ORBl) to one (FRP) or presence of projections to all ipsilateral cortical areas but also in preferential outputs to the cortical areas among the FRP, ORB, RSP and posterior parietal association areas (PTLp), AI, GU, VISC, MOp, MOs, SSp, SSs and PERl (Atlan et al., 2018; Narikiyo et al., 2020; Zingg et al., 2018). In our hands, anterograde tracing shows that the CLA sends outputs to all cortical modules, but preferentially to prefrontal and lateral modules as well as to the motor cortices and RSP. Our single neuron tracing data recapitulates these bulk anterograde tracing results, but with sparse projections to the lateral modules and MOp. The difference in projection strengths between our single neuron tracing results and bulk anterograde tracing ones and the difference in labeling between our anterograde and retrograde viral tracing results and others can be largely explained by the injection site contamination. This also explains the different results between the tract-tracing studies described below. Therefore, using multiple tracing datasets for cross-validation is crucial. Our finding that the CLA strongly and reciprocally connects with the prefrontal module is consistent with other studies (Chia et al., 2020; Qadir et al., 2018; White et al., 2017), supporting the notion that the CLA is involved in cognitive control but not in sensorimotor processing (Atlan et al., 2021; Krimmel et al., 2019a; White et al., 2018).

### Cell types and laminar origin of cortical pyramidal neurons projecting to the CLA

Cortical pyramidal neurons residing in L2-L6 determine the direction of information flow through their local and long-range axonal projections. They are classified into three major types: L2-L6 IT, L5 ET and L6 CT neurons based on downstream projection targets (Harris and Shepherd, 2015; Shepherd, 2013; Peng et al., 2021). Few studies have investigated cortical cell-type-specific projections to the CLA. One study reported that L2/3 IT and L5 IT neurons in the MOs project to the CLA but L5 ET and L6 CT neurons do not (Smith et al., 2016), while another reported that L6 IT neurons in the VISp send projections to the CLA (Baker et al., 2018). We here comprehensively demonstrate that cell types projecting to the CLA are determined by cortical areas and layers. Specifically, the prefrontal module has more cell types with stronger projections to the CLA than do the lateral, medial and sensory modules. In most cortical areas, L5 IT and L5 IT ET neurons send stronger projections to the bilateral CLA than other cell types/layers, while L6 IT neurons in VISp and VISl predominate projections to the ipsilateral CLA. These anterograde tracing results are consistent with our retrograde tracing ones and others, in which cortical neurons projecting to the CLA are denser in the infragranular layers than in the supragranular layer across cortical areas (Gutierrez-Ibarluzea et al., 1999; Zhang et al., 2001; Zingg et al., 2018). It has been reported that L5 cell types in the visual and motor cortices not only differ in brain-wide connectivity but also in function (Economo et al., 2018; Kim et al., 2015). Cortical cell types with different projection densities in various cortical areas and layers may influence the activity of CLA neurons differently.

### Cell types of CLA principal neurons and their axonal distribution in cortical areas

The heterogeneity of the principal neurons within the CLA is revealed by injecting multiple retrograde tracers into different combinations of the cortical areas (Li et al., 1986; Macchi et al., 1983; Marriott et al., 2020; Minciacchi et al., 1985; Zingg et al., 2018). Some cortical injections show high numbers of double-, triple- and quadruple-labeled neurons in the CLA but others do not. These retrograde tracing studies are unable to reveal the projection diversity of single CLA neurons due to numerous projection targets. Recently, principal neurons in the mouse CLA have been classified into different cell types based on their intrinsic electrical properties, or long-range axonal projections. Using whole-cell patch-clamp recording combined with retrograde tracing (White and Mathur, 2018) classified principal neurons into 2 cell types projecting to cortical areas ACA, PTLP, VISp, and higher visual areas in different proportions. A similar study classified ACA-projecting principal neurons into 4 cell types at different locations within the CLA (Chia et al., 2017). This classification may relate to our 4 morphological cell types: two ipsilateral ACA-projecting cell types (with and without projections to the ipsilateral RSP) and two bilateral ACA-projecting cell types (with and without projections to the ipsilateral RSP). Based on their intrinsic electrical properties, principal neurons are classified into 5 cell types: 4 to cortical targets and 1 to subcortical structures (thalamus, habenula and hippocampus) (Graf et al., 2020). However, CLA output to the thalamus remains controversial in the previous tract-tracing studies. While one study reported the presence of such output (Atlan et al., 2018), other studies failed to do so (Narikiyo et al., 2020; Peng et al., 2021 Zingg et al., 2018). Our single neuron tracing results are consistent with the latter showing absence of such output. This discrepancy could be caused by injection site contamination because the AI, adjacent to the CLA, does project to the thalamus.

In our previous study, we classified 29 CLA principal neurons into four broad cell types: one bilateral and three ipsilateral projecting types (Peng et al., 2021). By nearly doubling the number of reconstructed single neurons to 52, we not only confirm the rough topographical organization of individual CLA neurons but also find more cell types (nine versus four). Individual principal neurons project strongly and frequently to the five cortical areas (PL, ACA, MOs, RSP and VISp) and one of the retrohippocampal formation (ENT) in combinations with weak projections to different sensory cortical areas and subcortical structures. Based on projection trajectories and targets, we classified these cells into at least 9 morphological cell types that innervate anatomically and functionally linked targets (Harris et al., 2019; Ho et al., 2014; Holschneider et al., 2014; Vanni et al., 2017). Interestingly, all these morphologically diverse CLA cell types belong to the same car3 transcriptomic cell type (Peng et al., 2021). It will be interesting to know when and how adult-like principal cell types emerge in the CLA and to establish the relationship between the morphological and electrophysiological cell types.

In addition to projection strength, the laminar distribution of claustrocortical projections is another key factor influencing postsynaptic neurons. Although it has been investigated in a few cortical areas of different species, these reports are inconsistent. Early tract-tracing studies in cats reported that CLA axons predominantly terminate in L4 and L6 of V1 (LeVay and Sherk, 1981; LeVay, 1986; Sherk, 1986), whereas others reported heavy axon terminals in L1 (Carey et al., 1979). A recent study combining anterograde tracing with electron microscopy in cat V1 demonstrated that claustrocortical projections form synapses in all layers, preferentially in L2/3 and L5 (da Costa et al., 2010). On the other hand, claustrocortical projections in the cat’s frontal cortex terminate in L3b/4, L6, and L1, but are absent in L5 (Clascá et al., 1992). Similarly, differences are also seen in the anterograde tract-tracing studies in the mouse CLA. A study reported that CLA axons are denser in the superficial layers than the infragranular layers of the PL, ACA, ORBm, RSP, MO and SSp, while they are denser in L1 than in other layers of VISp and VISam (Fodoulian et al., 2020). In contrast, another study reported that CLA axons are denser in L6 than in L5 and L2/3 of the PL, ACA, ILA and MOs (Jackson et al., 2018). The third study reported that CLA axons are denser in L1 and L5 than in L2/3 and L6 of the ACA, PL, ILA, RSPd and PTLp, and they are denser in L2/3 and L5 than in L6 of the ORBvl, and they are denser in L1 than in other layers of the VISp (Zingg et al., 2018). Our single neuron tracing results recapitulate our anterograde tracing ones, showing different laminar distribution patterns across cortical areas. CLA axons are denser in L5 and L2/3 than in L1 and L6 of the prefrontal module and are denser in L6 than in other layers of the lateral module (AI and GU), and are denser in the lower L1 and L2/3 than in other layers of the sensory modules but avoid branches in L4. These discrepancies between previous studies and ours could be due to contamination of bulk anterograde tracer injections into the CLA, rather than species differences. Our single neuron tracing data are much better than the bulk tracing technique to identify the laminar distribution of their cortical targets without contamination. Different laminar distributions between claustrocortical projections and thalamocortical projections influence different cell types in the cortex (Brennan et al., 2021). It may be underly the mechanisms that neuron responses differ in cortical areas, layers and cell types after optical activating the CLA neurons (McBride et al., 2022)

### Intrinsic connections of inhibitory and excitatory neurons within CLA

A conventional retrograde tract-tracing study found extensive retrograde connections within the rat CLA itself, suggesting intrinsic CLA connections (Smith et al., 2010). This technique has two limitations. First, it cannot exclude the possibility of tracer uptake by damaged passing fibers. Second, it cannot reveal what cell type gives rise to the intrinsic connections. The newly developed retrograde viral tract-tracing method overcomes these limitations (Miyamichi et al., 2011; Yao et al., 2021). Using this new method, we also found substantial presynaptic neurons within the CLA. Some of these may be derived from CLA inhibitory neurons since they send extensive axonal arbors within the CLA (**Figure S7**) and form synapses with CLA principal neurons (Kim et al., 2016). Unexpectedly, we found that axon shafts of individual CLA principal neurons travel with few or no collaterals within the CLA. This finding is in line with a study in acute slice recordings, in which very few synaptic contacts were recorded between principal neurons but many between interneurons (Kim et al., 2016). However, another study in oblique slices that contain a large portion of rat CLA tissue, reported synchronous activity between CLA principal neurons incubated with the GABA_A_ receptor antagonist bicuculline (Orman, 2015). It is possible that both interneurons and principal neurons contribute to intrinsic CLA connections. The inhibitory intra-claustral connections could selectively gate information flow to functionally linked cortical areas through feedforward inhibition (Day-Brown et al., 2017).

### The connections between CLA and other major brain regions

The mouse CLA receives inputs from the olfactory areas, subcortical plate, hippocampal formation, striatum, pallidum, thalamus, hypothalamus, midbrain, and pons but only sends output back to a few of these structures (Altlan et al., 2018; Narikiyo et al., 2020; Zingg et al., 2018). These reported results are inconsistent regarding the presence or absence of some input-output connections of the CLA (**Supplementary Table 6**). Here, we discuss CLA connections with three regions: thalamus, neuromodulatory systems, and cerebellum.

In the previous studies, inputs to the CLA are inconsistently reported from the thalamic nuclei, including the parafascicular nucleus (PF), PVT, PT, RE, AM, mediodorsal nucleus (MD), CM, central lateral nucleus (CL), intermediodorsal nucleus (IMD), posterior complex (PO), ventral medial nucleus (VM), lateral dorsal nucleus (LD), subparafascicular nucleus (SPF) and PIL (Altlan et al., 2018; Narikiyo et al., 2020; Zingg et al., 2018). By integrating both anterograde and retrograde tracing results, we confirmed six of them, including PVT, PT, RE, CM, AM and PIL, and extended one (POL) to the list (**Figure 7C**). Collectively, these thalamic nuclei have reciprocal connections with the midline and higher association cortices (Oh et al., 2014; Harris et al., 2019) and have been suggested to involve arousal and higher cognitive functions (Saalmann, 2014; Van der Werf et al., 2002; Wolff and Vann, 2019). In addition to the direct thalamocortical pathways, our results clearly showed the existence of indirect thalamo-CLA-cortical pathways, which could also be part of the higher cognitive processing networks.

CLA neurons receive inputs from cholinergic (Selden et al., 1998), noradrenergic, dopaminergic, and serotonergic nuclei (Baizer, 2001; Pirone et al., 2018) and have corresponding receptors to these neuromodulators (Sitte et al., 2017; Terem et al., 2020). We confirmed inputs to the CLA from the NDB (cholinergic neurons), CS and DR (serotoninergic neurons), VTA (dopaminergic neurons) and LC (noradrenergic neurons) and extended new inputs from PPN (cholinergic neurons) and RPO (serotoninergic neurons) to the list (**Figure 7C**). Our finding of dopaminergic innervation of the mouse CLA is consistent with the tract-tracing and immunohistochemical studies (Atlan et al., 2018; Pirone et al., 2018; Zingg et al., 2018), but disagrees with others (Narikiyo et al., 2020; Barbier et al., 2016). What causes this difference remains to be determined. It may be due to projections being too sparse to be detected from the LC to the CLA (Plummer et al., 2020). Cholinergic and serotonergic inputs to the CLA in acute slice recording studies have been suggested to influence CLA neuron activities differently (Nair et al., 2021; Wong et al., 2021). Neuromodulation via CLA influencing animal behaviors has recently been explored in humans. Psilocybin, a partial serotonin 2A (5-HT_2A_) receptor agonist, has been suggested to alter the functional connectivity of the CLA to cortical networks that support perception, memory, and attention (Barrett et al., 2020; Doss et al., 2021).

A recent study reported projections from the superior cerebellar peduncle to the CLA by injecting conventional tracers into the superior cerebellar peduncle but did not specify projections from which part of the cerebellum (Cavdar et al., 2018). In this study, we showed sparse projections from the IP to the contralateral CLA (**Figure 7C**) but not from the dentate and fastigial nuclei. This finding suggests that the IP signal could reach the prefrontal module and other cortical modules through the CLA. Therefore, in addition to controlling motor executive function, this IP-CLA-cortical pathway could involve higher cognitive processing (Balsters et al., 2014; Brooks et al., 2015; Gao et al., 2018; Stoodley, 2012).

### Functional implications

In this study, we demonstrate that the prefrontal module, especially ACA, has more cell types projecting strongly to and receives stronger input from the CLA compared with other cortical modules. These reciprocal connections are symmetrical in input-output projection densities with most cortical areas such as the ACA but not with a few others such as the RSP and VISp, which receive denser input from the CLA than sending output to it. Many subcortical structures, such as higher-order thalamic nuclei, send projections to the CLA which are not reciprocated (**Figure 7C and 7D**). As a unique hub within the cortical network, the CLA has been suggested to be involved in higher cognitive processing (Crick and Koch, 2005; Krimmel et al., 2019; Reus-García et al., 2021). This notion is supported by investigating different CLA-cortical pathways in mice. Optogenetic silencing ENTm-projecting CLA neurons impaired memory retrieval during contextual fear conditioning, suggesting that the CLA-ENTm pathway modulates the function of contextual memory (Kitanishi and Matsuo, 2017). Furthermore, perturbing the CLA-medial prefrontal pathway suggest that this pathway is involved in regulating impulsivity (Lui et al., 2019), attentional set-shifting (Fodoulian et al., 2020), and salience detection (Atlan et al., 2018; Terem et al., 2020), while optogenetic interrogating of the CLA-ACA pathway suggests that this pathway modulates engagement with the external world (Atlan et al., 2021). Optically activating CLA principal neurons of transgenic lines in the awake head-fixed mice on freely moving-wheel shows that neuronal responses differ in different layers and cell types of various cortical areas (prefrontal cortex, ACA, MOs and RSP) (McBride et al., 2022).

Our findings showed that the diverse CLA-cortical pathways that receive cortical and subcortical inputs transmit information back to functionally linked targets, after competitive “gating” via CLA inhibitory neurons. We look toward future studies of cell-type-specific inputs and outputs of the CLA in mice engaged in tasks that require competitive attentional and/or cognitive control.

## Supporting information

Supplementary Stable 1

## Acknowledgments

We are grateful to the Transgenic Colony Management, Neurosurgery and Behavior, Lab Animal Services, Molecular Genetics, Imaging, Histology, Technology, and Project Management teams at the Allen Institute for technical and management support. We thank Thomas R Reardon, Andrew J Murray and Ian Wickersham for providing cell lines and plasmids for the establishment of rabies virus production at the Allen Institute. This work was supported by the Allen Institute for Brain Science and by the National Institute of Mental Health (NIMH) of the National Institutes of Health (NIH) under award number U19MH114830 to H.Z. We also acknowledge funding from the Tiny Blue Dot Foundation. The content is solely the responsibility of the authors and does not necessarily represent the official views of NIH and its subsidiary institutes. We thank the Allen Institute founder, Paul G. Allen, for his vision, encouragement, and support.

## Author Contributions

Conceptualization: H.Z, J.A.H, A.C. Q.W; Transgenic mice: K.E.H, J.A.H, and H.Z; Project Administration: J.A.H; S.S; Single-cell reconstruction: Y.W, X.K, Y.L, M.Y, C.C, W.X, P.L, L.A and L.E; Single-cell data analyses: P.X, H.K, M.M, Y.W, Q.W; Retrograde data generation: S.Y, A.C; Anterograde and retrograde data analyses: K.E.H, Q.W; Data visualization: M.N; The original draft was written by Q.W, with input from C.K, J.A.H, and S.S. Supervision: C.K., H.Z. All authors discussed and commented on the manuscript.

## Supplemental Figure Legends

**Figure S1.**
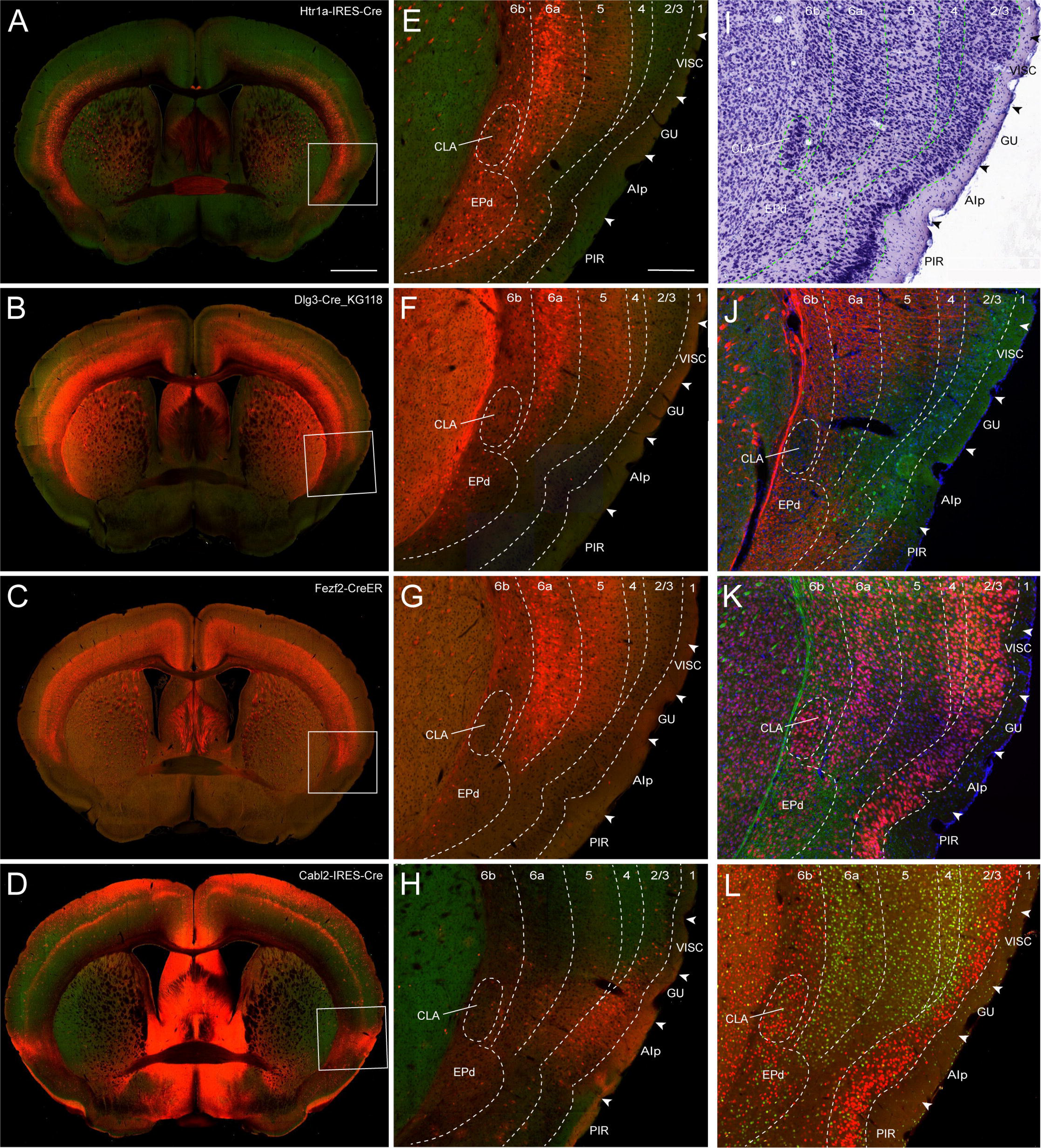
Delineation of the boundary of the mouse CLA, related to Figure 1. (A-C) Three transgenic lines Htr1a-IRES-Cre (A), Dig3-Cre (B) and Fezf2-CreER (C) show enriched gene and poor expression in layer 6a and layer 6b of isocortical areas respectively, with less or absence of gene expression in the CLA. The boxes in A-C are enlarged in E-G. Scale bar, 1 mm in A-D. (D) Transgenic line Calb2-IRES-Cre shows enriched gene expression in layers 2/3 and layer 5 of the isocortex but less expression in the CLA. The box in D is enlarged in H. (E-G) Enlargements of boxes are from A-C. Dash lines indicate the borders between cortical layers as well as for CLA and EPd. Arrowheads indicate borders between the cortical areas. Dash-lines and arrowheads indicate the same in H-L below. For abbreviations, see Supplementary Table 7. Scale bar = 200 μm in E-L. (H) Enlargement of the box in D. (I) Nissl specimen shows the CLA as a densely packed group of neurons darkly labeled underneath layer 6 of the agranular insular area. (J) Dual immunostaining with antibodies against SMI-99 (in red) and Calb1 (in green) shows less myelination of CLA compared to its adjacent structures. (K) Dual immunostaining with antibodies against NeuN (in red) and NF-160 (in green) reveals the CLA as a densely packed group of neurons underneath layer 6 of the agranular insular area with less myelination compared to its adjacent structures. (L) Transgenic line Rorb-IRES2-Cre shows that red neurons are more in cortical layers 2/3 and 6b than layers 5 and 6a, while the green neurons do the opposite: more in layers 4. 5 and 6a than in layers 2/3 and 6b. Red neurons in layer 6b extend to and become relatively densely packed in the CLA.

**Figure S2.**
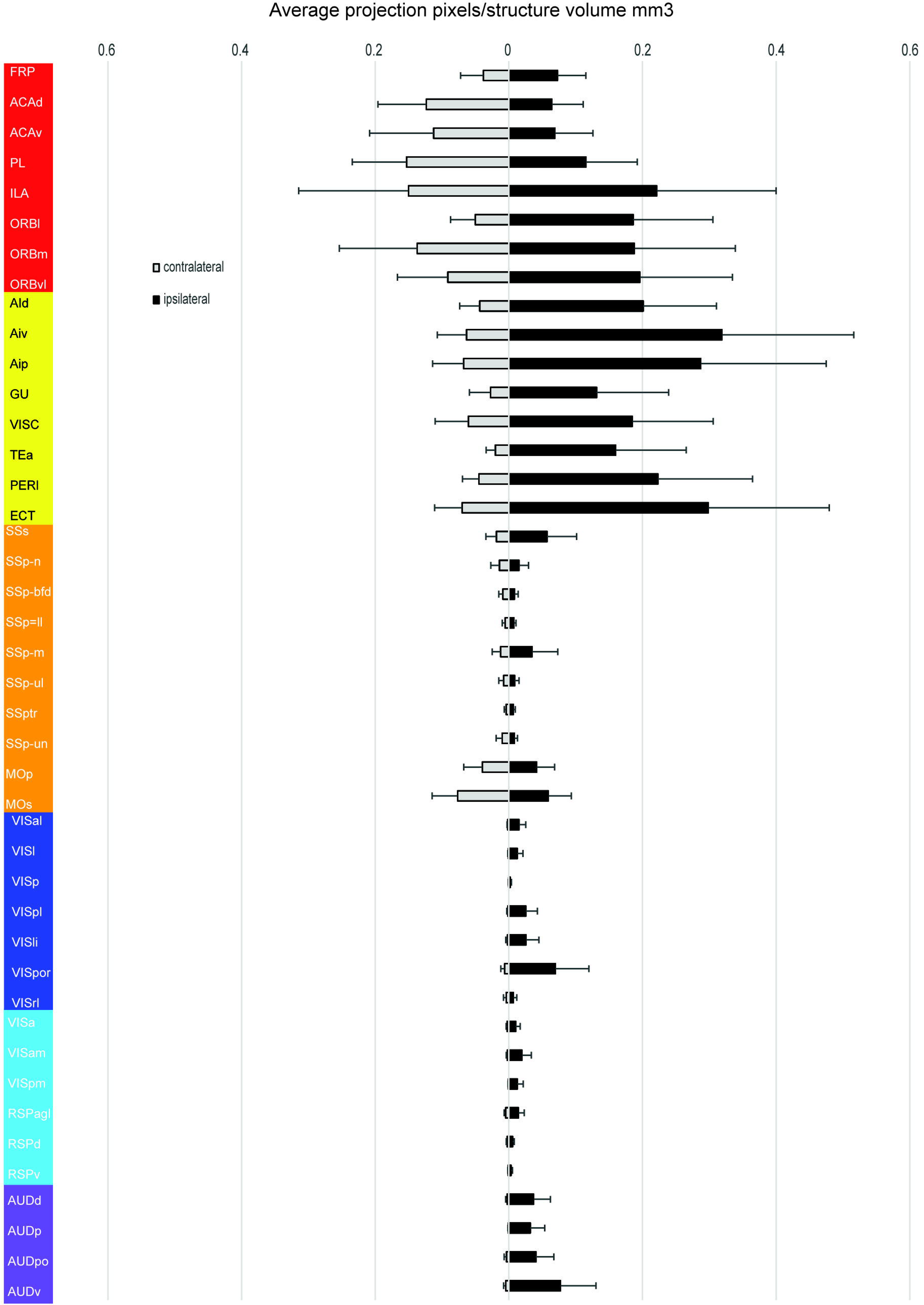
Presynaptic neuron distribution in the isocortex, related to Figure 2. The histogram shows the fraction of total presynaptic labeling for each cortical area normalized by starter cells and structural volume. Y-axis shows presynaptic input from six cortical modules color-coded differently. Black and shaded bars represent presynaptic labeling in the ipsilateral and contralateral cortex, respectively. For abbreviations see Supplementary Table 7. Mean ± SD (standard deviation).

**Figure S3.**
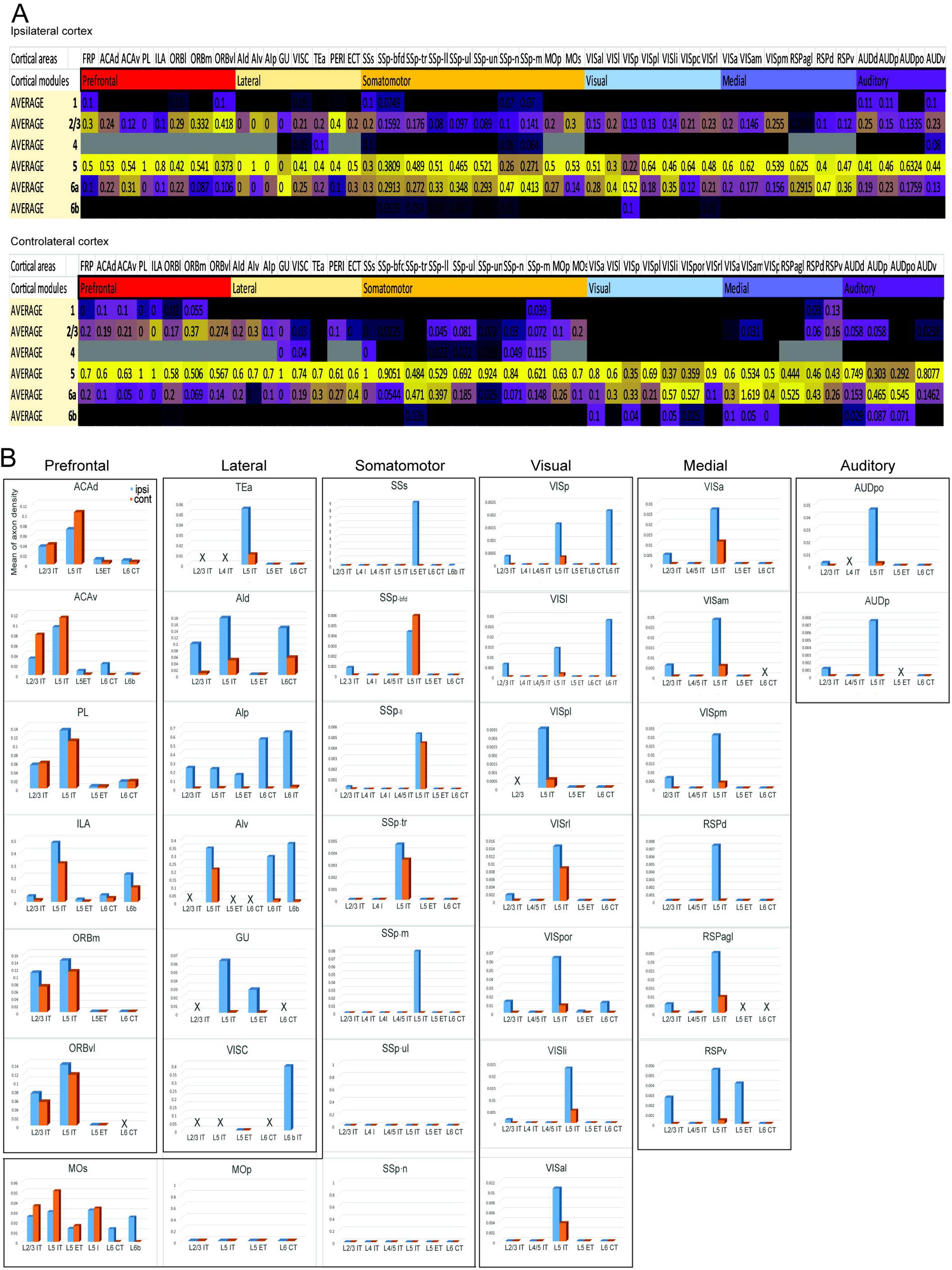
Laminar origin and cell type-specific cortical inputs to the CLA, related to Figure 3. (A) Laminar origin of cortical neurons sending inputs to the CLA are predominately from L5 in the ipsilateral and contralateral cortical areas averaged from five retrograde rabies injection cases. Cortical modules are color-coded differently along the X-axis. Yeller and blue colors represent dense and sparse presynaptic input to the CLA from individual layers of cortical areas. For abbreviations see Supplementary Table 7. (B) Cortical areas in the prefrontal and lateral modules have more cell types and send stronger projections to the CAL than sensory and medial modules. However, the MOp and SSp-n and SSp-ul send no projections to the CLA. Bule and red bars in the histogram represent ipsilateral and contralateral projections respectively. “X” indicates an absence of injection. Note that L5 IT ET neurons are included in L5 IT neurons for each cortical area and that axonal density in Y-axis differs in order of magnitude across cortical areas.

**Figure S4.**
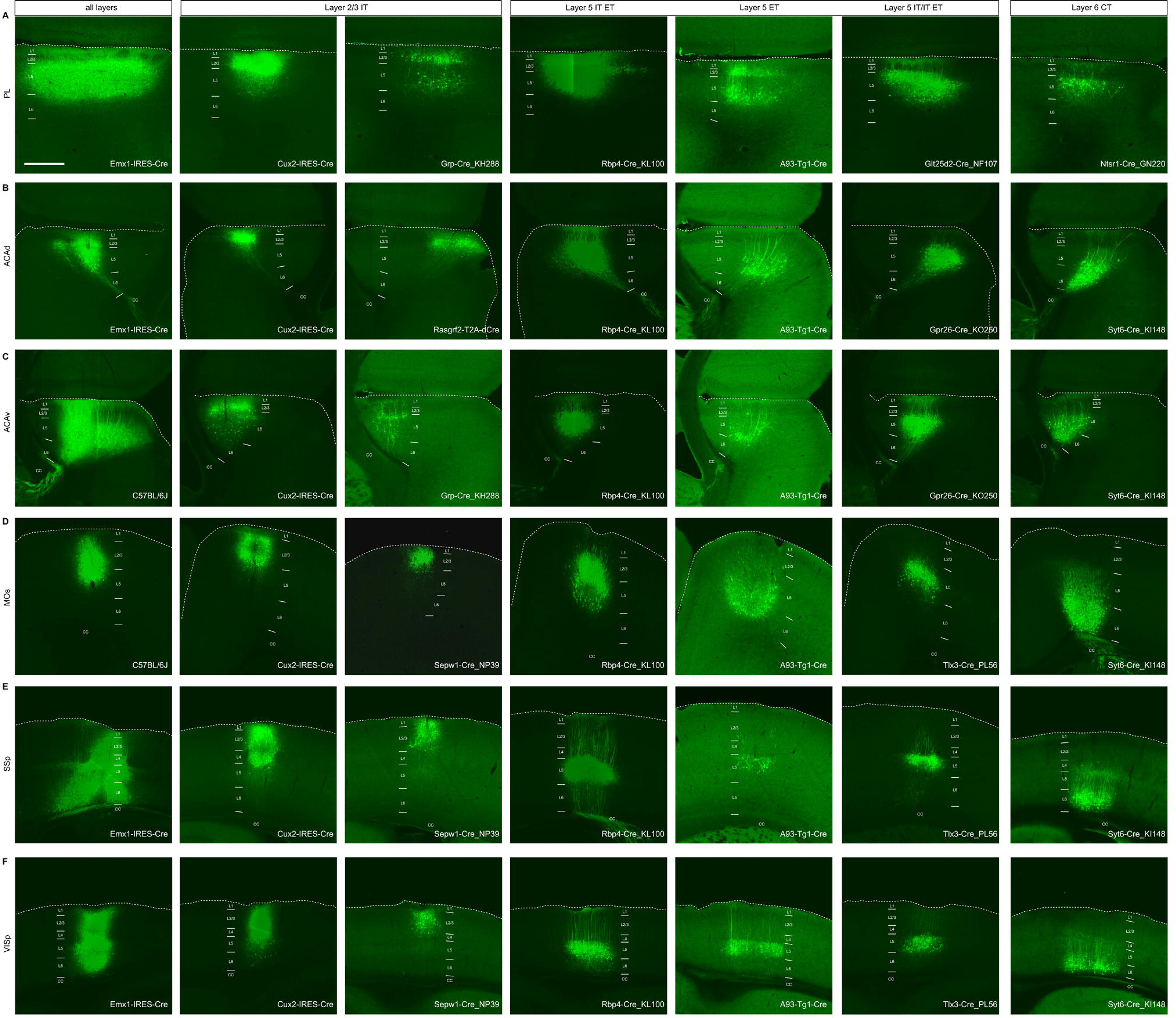
Examples of injection sites in different layers of the cortical areas, related to Figure 3. (A) A row shows injection sites in the PL of the pan-neural transgenic line and in layers 2/3 IT, 5 IT, 5 IT ET, 5 ET and 6 CT defined by transgenic lines. White strips indicate the border between layers. Scale bar, 500 μm in A-F. (B) B row shows injection sites in the ACAd of the pan-neural transgenic line and in layers 2/3 IT, 5 IT ET, 5 ET and 6 CT defined by transgenic lines. (C) C row shows injection sites in the ACAv of wide-type mice and layers 2/3 IT, L5 IT ET, 5 ET and 6 CT defined by transgenic lines. (D) D row shows injection sites in the MOs of wide-type mice and layers 2/3 IT, 5 IT, 5 IT ET, 5 ET and 6 CT defined by transgenic lines. (E) E row shows injection sites in the SSp of the pan-neural transgenic line and layers 2/3 IT, 5 IT, 5 IT ET, 5 ET and 6 CT defined by transgenic lines. (F) F row shows injection sites in the VISp of the pan-neural transgenic line and layers 2/3 IT, 5 IT, 5 IT ET, 5 ET and 6 CT defined by transgenic lines.

**Figure S5.**
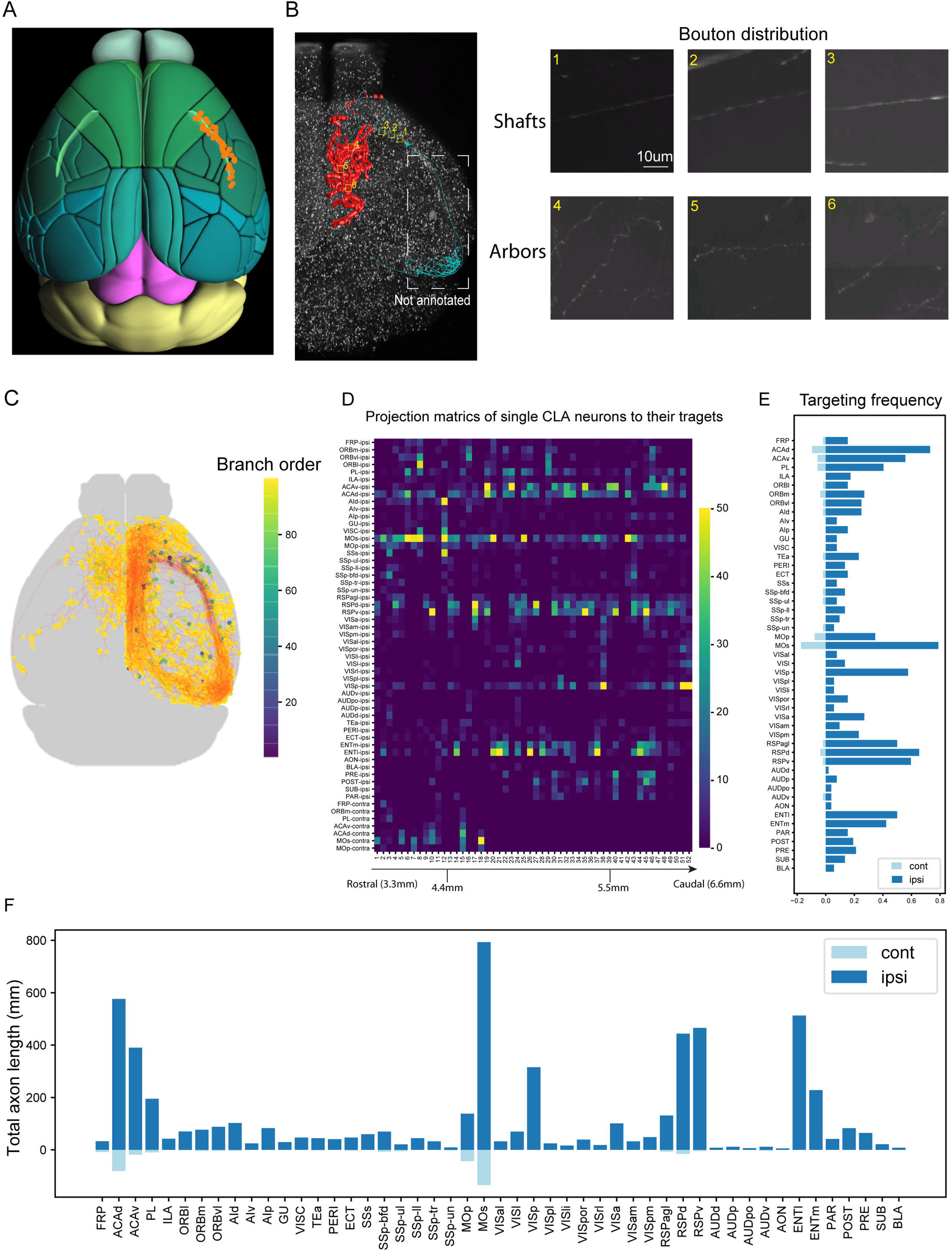
Individual CLA neurons project to the diverse subsets of the cortical targets, related to Figure 6. (A) Soma locations of individual principal neurons (in brown) within the CLA (in light green) are shown in the dorsal view. 35 principal neurons reconstructed from the left CLA were flipped to the right CLA. (B) Synaptic bouton distribution from a representative example of individual principal neurons is shown in the dorsal view. Manually plotted synaptic boutons were marked by red in the midline cortical areas, but they are not plotted in the entorhinal cortex (in cyan). The insets on the left panel of B are enlarged on the right panel of B. The enlarged insets 1, 2, and 3 show axon shafts without boutons within the CLA, and insets 4, 5, and 6 show axonal arbors bearing synaptic boutons in the targets. Scale bar, 10 μm in the insets. (C) Axon branch points of all 52 reconstructed principal neurons are scatter-plotted in the dorsal view. Branching levels are calculated by the sizes of descendent trees. The color key indicates axon branch levels. Yellow indicates high branches and blue indicates fewer branches. (D) The matrix shows the total lengths of projections of principal neurons in its projection targets as a proxy for projection strengths. Projection strength is indicated by the color key: strong in yellow and weak in blue. Along X-axis, these neurons were arranged from the rostral (left) to the caudal (right). Y-axis shows projection targets of individual principal neurons. (E) Histogram shows projection frequency to targets by individual principal neurons. (F) Histogram shows the total axonal projection lengths of all 52 principal neurons in individual targets.

**Figure S6.**
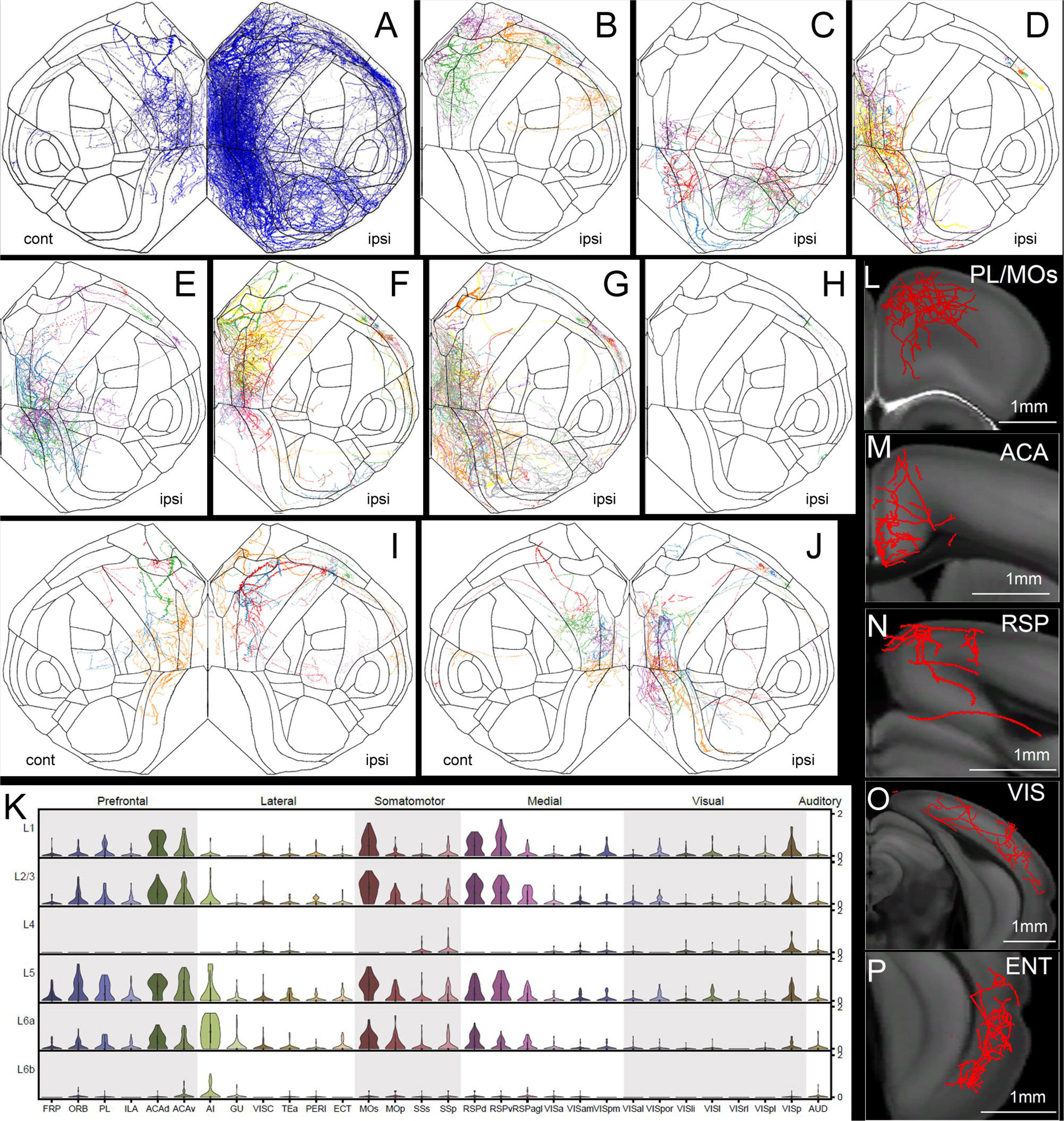
Distinct cell types of CLA principal neurons and their axonal distribution in cortical areas, related to Figure 6. (A) The flatmap reveals the combined axonal distribution of all reconstructed CLA principal neurons in cortical areas. “ipsi” and “cont” are abbreviations for ipsilateral and coroplateral respectively. These are the same in B-J. (B-H) The flatmap shows 7 clusters of the ipsilateral projecting principal neurons. Individual neurons were color-coded differently. (I-J) The flatmap shows 2 clusters of the bilateral projecting principal neurons. Individual neurons were color-coded differently. (K) The violin plot shows the laminar distribution of claustrocortical projections across cortical areas from all principal CLA neurons. For abbreviations see supplementary Table 7. (L-P) Examples of the laminar distribution of ipsilateral claustrocortical projections in the strongest recipient cortical areas PL/MOs (L), ACA (M), RSP (N), VIS (O), and ENTl (P). Scale bar, 1 mm in L-P.

**Figure S7.**
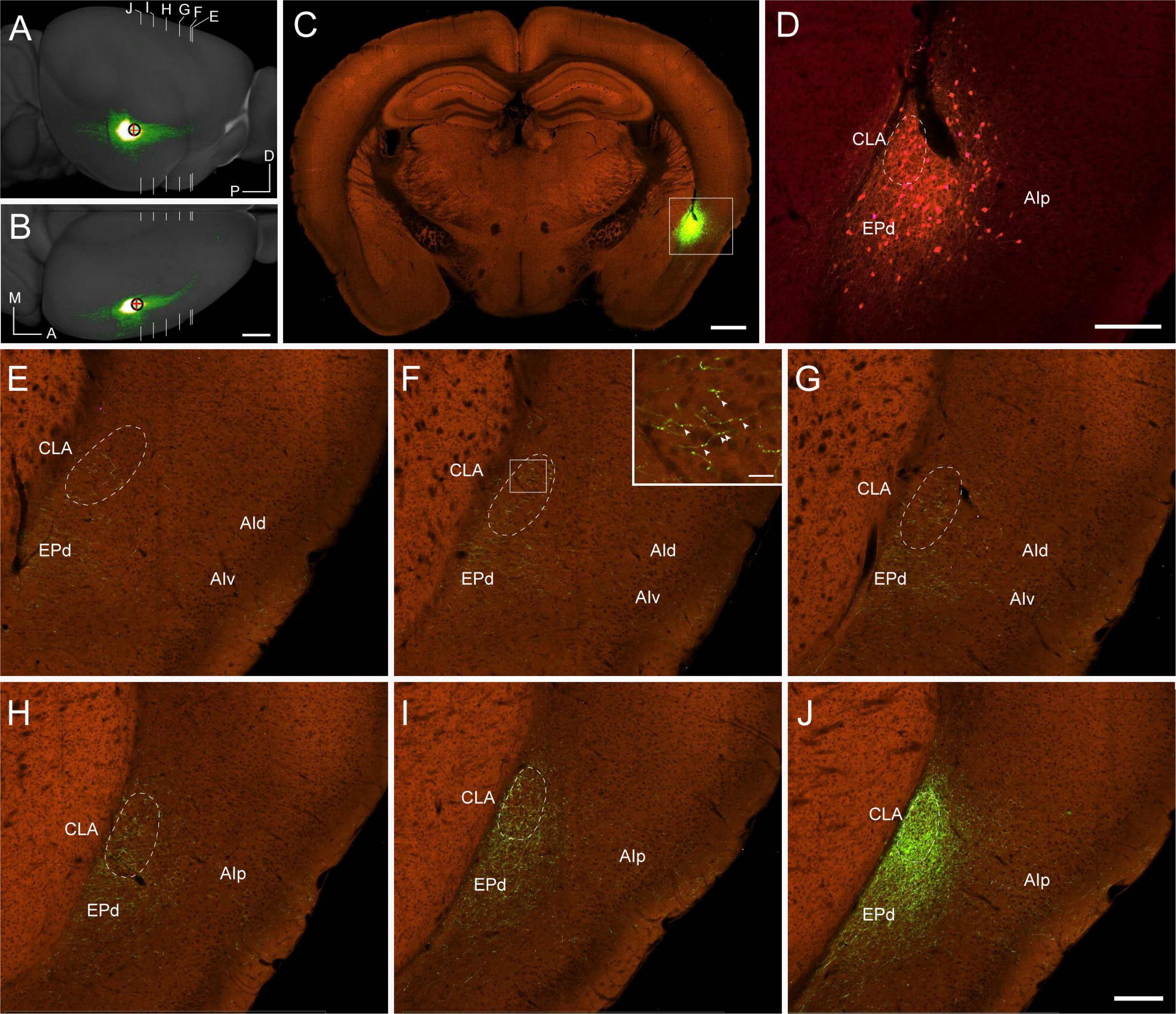
Axonal arborization of inhibitory neurons within the CLA, related to Figure 7. (A-B) The injection into the CLA of the pan-inhibitory neuron transgenic mouse, Gad2-IRES-Cre, is shown in the dorsal (A) and lateral (B) views. White strip lines indicate coronal section levels from E to J shown below. The red cross inside the black circle indicates the center of the injection site in A and B. A, anterior; P, posterior; L, lateral; D, dorsal. Scale bar, 1 mm. (C) Injection site is shown in the coronal section marked in A and B with the red cross. Scale bar, 700 μm. (D) The injection site in the inset (C) was enlarged in (D). The intensity of the green and red channels was reduced in order to see infected neurons in the injection site. The dash-line indicates the border of the CLA. For abbreviations see Supplementary Table 7. Scale bar, 200 μm. (E-J) Axons of inhibitory neurons travel distantly from the caudal to the rostral within the CLA at different levels indicated in A. The inset in F shows axons with beads-like synaptic boutons within the CLA. Dash-line indicates the border of the CLA. Scale bar, 200 μm in E-J, 20 μm in the inset of F.

## STAR*METHODS

Detailed methods are provided in the online version of this paper and include the following:

### CONTACT FOR REAGENT AND RESOURCE SHARING

Further information and requests for resources and reagents should be directed to and will be fulfilled by the Lead Contact H.Z.

### EXPERIMENTAL MODEL AND SUBJECT DETAILS

#### Animal care and use

All experimental procedures were approved by the Allen Institute Institutional Animal Care and Use Committee (IACUC) and conform to NIH guidelines. Both male and female wild-type (C57BL/6J) and transgenic mice ≥P56 were used in this study. All animals were housed 3-5 per cage, under constant temperature, humidity, and light conditions (12 hours light/dark cycles) and given food and water ad libitum.

### METHOD DETAILS

#### Histology and immunohistochemistry

The histology and double immunohistochemistry (IHC) procedures are detailed at the Allen Mouse Brain Connectivity Atlas documentation page (http://help.brain-map.org//display/mouseconnectivity/Documentation) and in our previous publications (Wang et al., 2017, 2020). Briefly, adult mice were anesthetized with 5% isoflurane and intracardially perfused with 10 ml saline (0.9% NaCl) followed by 50 ml freshly prepared 4% paraformaldehyde (PFA). Brains were rapidly dissected, postfixed, and transferred to a 30% sucrose solution. After sunk, brains were embedded in OCT and sectioned at 25 μm on a Leica 3050 S cryostat equipped with an Instrumedics Tape Transfer System (plus UV light polymerization chamber and warming pad). A series of alternative sections from each brain were collected for Nissl-staining, or double IHC. Nissl-stained sections were delipidated with the xylene substitute Formula 83 (CBG Biotech, Columbus, OH; catalog No. CH0104) and ethanol rehydrated. After several washes in water, the sections were stained in 0.21% thionine for 3 minutes and dehydrated by sequential immersion in increasing concentrations of ethanol. Differentiation and monitoring were performed at 95% ethanol before completion with pure ethanol. Dehydrated sections were subsequently incubated in Formula 83 and coverslipped with the Curemount mounting medium (Instrumedics, Hackensack, NJ; catalog No. 475232). For imaging, sections were scanned with the x10 objective on ScanScope, an automated image capture platform (Aperio Technologies, Vista, CA). For double IHC, sections were washed after antigen retrieval with 10 mM sodium citrate and then incubated in blocking solution (4% normal goat serum plus 0.3% Triton X-100 in PBS) for 1 hour. After brief rinsing, each series of sections was incubated with one of the following primary antibody pairs overnight: Calbindin1 (Calb1; Swant, catalog No. CB38.Rabbit final dilution 1:2,000 in blocking solution) and SMI-99 (Covance, Berkeley, CA; catalog No. SMI-99P, RRID?; 1:1,000), or NeuN (Millipore, Bedford, MA; catalog No. MAB377, RRID:AB_10048713; 1:1,000) and NF-160 (Abcam, Cambridge, MA; catalog No. ab9034, RRID:AB_306956; 1:1,000). After rinsing primary antibodies thoroughly, each series of sections were incubated in a pair of the secondary antibodies: goat anti-rabbit-488 (final dilution 1:1,000 in blocking solution for Calb1, and 1:500 for NF160) and goat anti-mouse-594 (1:500 for SMI-99 and NeuN) overnight. After rinsing, sections were counterstained with DAPI (Invitrogen, Carlsbad, CA; catalog No. D1306) and coverslipped with Fluoromount G medium (Southern Biotechnology, Birmingham, AL; catalog No. 0100-01). Sections were scanned on a fully automated, high-speed multichannel epifluorescence scanning system, VS110/ 120 (Olympus, Center Valley, PA) with x10 objectives.

#### Tracer injections

Detailed procedure for anterograde viral tracing has been described elsewhere (Harris et al., 2019; Oh et al., 2014). Surgery procedures for all experiments were the same with different tracers and mice used. A pan-neuronal AAV vector expressing EGFP under the human synapsin I promoter (AAV2/1.pSynI.EGFP.WPRE.bGH, Penn Vector Core, AV-1-PV1696, Addgene ID 105539) was as an anterograde tracer for injections into wild-type mice and a Cre-dependent AAV vector expressing EGFP (AAV2/1.pCAG.FLEX.EGFP.WPRE.bGH, Penn Vector Core, AV-1-ALL854, Addgene ID 51502), or a Cre-dependent AAV virus expressing a synaptophysin-EGFP fusion protein (AAV2/1.pCAG.FLEX.sypEGFP.WPRE.bGH, Penn Vector Core) was used as an anterograde tracer for injections into Cre driver mice. All adult mice were anesthetized with 5% isoflurane briefly and secured to a stereotaxic frame (Model# 1900, Kopf, Tujunga, CA) before surgery. During surgery, anesthesia was maintained at 2% isoflurane. After the skin incision, a small divot was made on the skull surface with a fine drill burr. To reveal the brain surface, a tiny thin layer of bone was removed with miniature forceps. For injection, a glass pipette (inner tip diameter of 10-20 µm) loaded with AAV was inserted into the desired brain depth from the pial surface. The coordinates for injections into cortical and subcortical structures were listed on our portal (connectivity.brain-map.org) based on the mouse brain atlas (Paxinos and Franklin, 2001). These viral tracers were delivered by iontophoresis (current 3 µA, and 7s on / 7s off duty cycle) for 5 min. Most of the injections were made in the right hemisphere and a small number of injections in the left hemisphere.

The procedure for monosynaptic retrograde rabies tracing has been described in detail elsewhere (Yao et al., 2021). First, Cre-dependent anterograde viral tracer AAV2/1.pSyn.CVS-N2C-g.VTA.tdTomato as helper virus was delivered to the CLA of transgenic mice (Gnb4-IRES2-Cre, Ntgn2-IRES2-Cre, Rbp4-Cre or Glt25d2-Cre) by iontophoresis with the parameters as described above. Two weeks later, EnvA-pseudotyped CVS-N2C-histoneGFP rabies virus (pRbV) (500nl) was pressure-injected into the same location in the CLA. The injection coordinates for CLA were based on the mouse brain atlas (Paxinos and Franklin, 2001) at different anterior-posterior locations: AP -0.34 – 1.78mm from Bregma, ML 1.85–3.5mm from the midline, DV 2.83–3.10mm from pia surface. The rabies viral vector can only infect cells that express TVA. Since the Cre-dependent AAV provides Cre+ cells with a source of rabies glycoprotein, newly formed rabies virus particles can spread retrogradely from these Cre+ neurons to their directly connected presynaptic neurons. The presynaptic neurons do not express TVA or rabies glycoprotein which prevents the rabies virus from spreading further. This method effectively restricts rabies virus infection to Cre+ cells and their direct, monosynaptic presynaptic neurons, and several control experiments were shown elsewhere (Yao et al., 2021). Thus, the “starter cell” contains both red and green fluorophores and appears yellow after merging two channels, while the nucleus of each presynaptic neuron is labeled only in green, and neuron infected with the helper virus is labeled in red.

#### Whole-brain imaging

After the survival of two weeks for anterograde AAV injections, or one week for retrograde rabies injections, mice were anesthetized with 5% isoflurane and intracardially perfused with 10 ml of saline (0.9% NaCl) followed by 50 ml of freshly prepared 4% PFA. Brains were rapidly dissected, post-fixed in 4% PFA at room temperature for 3-6 hours and overnight at 4°C, then rinsed briefly with PBS and stored in PBS with 0.1% sodium azide. Before scanning, residual moisture on the brain surfaces was removed with Kimwipes. The brains were subsequently placed in 4.5% oxidized agarose (made by stirring 10 mM NaIO_4_ in agarose), transferred to a phosphate buffer solution, and put in a grid-lined embedding mold for standardized orientation in an aligned coordinate space. High resolution 140 block-face coronal images (20x) for the Allen Mouse Brain Connectivity Atlas were scanned with serial two-photon tomography (STPT) (TissueCyte 1000 system, TissueVision, Cambridge, MA).

The transgenic dataset used in this study was prepared and imaged in the same way as described above without any injections (Wang et al., 2017, 2020).

#### Annotation of injection sites

In the Allen Mouse Brain Connectivity Atlas, all anterograde injection sites are listed on our web portal (http://connectivity.brain-map.org/). The largest infected volume of a given brain structure measured was automatically assigned as the primary injection site, and if any, the smaller infected volume of the structure(s) was named as the secondary injection site. Due to some experiments were not perfectly registered to the CCFv3, we manually inspected all injection sites to ensure that they are assigned correctly. Most of the automatically assigned primary injection sites are correct but a small portion of injections was re-assigned based on their projection targets and relative geographic locations to their neighbors (**Supplemental Tables 1**). We also carefully inspected injection sites of retrograde rabies injections. Retrograde rabies injection sites were done as above for anterograde injection sites. A structure containing the most volume of the starter cells is named as a primary injection site and the smaller volume of structure(s), if any, were named as the secondary injection site. In this study, we not only carefully check the primary injection sites but also the secondary injection sites because the afferent and efferent projections of the CLA can derive from the secondary injection sites. All anterograde tracing experiments in cortical and subcortical structures are listed in **Supplementary Tables 3 and 4**, respectively.

#### Data processing and quantification

STPT images were processed using the informatics data pipeline (IDP), which manages the processing and organization of the image and quantified data for analysis and display in the web application as previously described (Kaun et al., 2015; Oh et al., 2014; Harris et al., 2019). The IDP contains two key algorithms: image alignment and signal detection. The global alignment process between the Cre mouse brains or AAV-injected brains and the average template consists of three steps: 1) a coarse registration initialized by matching the image moments of the image stack and template, 2) a rigid registration (rotation plus translation), and 3) a 12-parameter affine registration. Each step was based on maximizing the image similarity metric between the transgenic image stack or injection image stack and the template using a multiresolution algorithm. To increase alignment accuracy further, local registration was then performed. As with the global alignment, local registration was conducted sequentially from coarse to fine at four resolution levels with decreasing smoothness constraints. The signal detection algorithm was applied to each image to segment positive fluorescent signals from the background. Image intensity was first rescaled by square root transform to remove second-order effects followed by histogram matching at the midpoint to a template profile. Median filtering and large kernel low pass filter were then applied to remove noise. Signal detection on the processed image was based on a combination of adaptive edge/line detection and morphological processing. Two variations of the algorithm were employed, depending on the virus used for that experiment; one was tuned for EGFP, and one for SypEGFP detection. High-threshold edge information was combined with spatial distance conditioned low-threshold edge results to form candidate signal object sets. The candidate objects were then filtered based on their morphological attributes such as length and area using connected component labeling. For the SypEGFP data, filters were tuned to detect smaller objects (punctate terminal boutons vs long fibers). In addition, high-intensity pixels near the detected objects were included in the signal pixel set. Detected objects near hyper-intense artifacts occurring in multiple channels were removed. We developed an additional filtering step using a supervised decision tree classifier to filter out surface segmentation artifacts, based on morphological measurements, location context, and the normalized intensities of all three channels. The output is a full resolution mask that classifies each 0.35 μm × 0.35 μm pixel as either signal or background. An isotropic 3-D summary of each brain is constructed by dividing each image into 10 μm × 10 μm grid voxels. The total signal is computed for each voxel by summing the number of signal positive pixels in that voxel. Each image stack is registered in a multi-step process using both global affine and local deformable registration to the CCFv3 as described above. Segmentation and registration results are combined to quantify the signal for each voxel in the reference space and for each structure in the reference atlas ontology by combining voxels from the same structure.

To quantify retrogradely labeled neurons in the whole brain, we modified the above algorithm to automatically detect fluorescence signal of nucleus-localized H2B-EGFP over the background and measured the pixel intensity in each anatomical structure (Yao et al., 2021). Since the signal detection algorithm was optimized to detect sparse retrograde labeling with high sensitivity, the automatically detected retrograde labeling voxel can have false positives where high background signal is falsely identified as retrograde neurons. To remove this type of artifact, we identified a set of 92 negative brains that were processed through the pipeline without rabies-mediated GFP expression and used this negative dataset to calculate the threshold of false-positive signal, i.e., the value of mean retrograde labeling pixel plus 6 standard deviations for each of the 316 ipsilateral and 316 contralateral structures of the brain. Retrograde labeling pixel not passing this threshold was set to “0”. A manually validated binary mask was then applied to further remove artifacts from informatically-derived measures.

We compared the quantification of retrograde labeling in 6 structures between automatic detection and manual count (Yao et al., 2021) and found strong positive linear correlations between them. Therefore, we used the automatically quantified retrograde labeling pixels as a proxy for the number of retrogradely labeled neurons in each structure.

#### Counting starter cells

All sections putatively containing the injection site were collected for immunostaining with an anti-RFP antibody (rabbit, Rockland Antibodies and Assays, 600-401-379) to amplify the tdTomato signal for the AAV1-pSyn-CVS-N2C-g-TVA-tdTomato virus. After immunostaining, all these sections containing tdTomato-positive cells were imaged by confocal microscopy (SP8, Leica) using a 10x objective, 4 μm z-step size stacks. Maximum intensity images of each section were generated and used to manually count starter cells using ImageJ. Cells co-labeled for both tdTomato (in red) and native histone-GFP (in green) signals were identified as starter neurons (in yellow). The CLA was identified as a densely labeled cluster of neurons counterstained with DAPI. The starter cells were identified as being most in the CLA but a small amount in adjacent structures L6 of the AI and VISC/GU latera-dorsally, and/or EPd ventrally to the CLA.

#### Sparse neuron labeling for single CLA principal neuron tracing

The procedure of sparse neuron labeling has been described in detail in our previous study (Peng et al., 2021). Briefly, a viral load of approximately 1×10^8^ particles was diluted into 50 μl sterile PBS and injected into the retroorbital sinus of anesthetized Gnb4-CreERT2 driver mice using a 31G insulin syringe. Following a single dose of tamoxifen induction administered 5-7 days, mice were survived three weeks.

#### Tissue preparation for single CLA principal neuron tracing

All tissue preparation procedures have been described in the previous publication (Peng et al., 2021). In brief, anesthetized mice were fixed on the operating floor and then intracardially perfused with 50 ml of 0.01 M PBS (Sigma-Aldrich Inc., St. Louis, US), followed by the same volume of 4% paraformaldehyde (PFA) and 2.5% sucrose in 0.01 M PBS. The infusion speeds were strictly controlled to avoid bubbles in the brain, which seriously affect imaging quality. The brains were removed from the skull and immersed in 4% PFA at 4°C for 24 hours. For embedding resin tissue, each intact brain was dehydrated by immersion in a graded ethanol series and then impregnated with HM20 working solution series (Electron Microscopy Sciences, cat. no. 14340)., each intact brain was rinsed three times in 12 hours at 4°C in a 0.01 M PBS solution (Sigma-Aldrich Inc., St. Louis, US). Then the brain was subsequently dehydrated via immersion in a graded series of ethanol mixtures (50%, 70%, and 95% ethanol solutions in distilled water) and the absolute ethanol solution three times for 2 hours each at 4°C. After dehydration, the whole brain was impregnated with Lowicryl HM20 Resin Kits (Electron Microscopy Sciences, cat.no. 14340) by sequential immersions in 50, 75, 100 and 100% embedding medium in ethanol, 2 hours each for the first three solutions and 72 hours for the final solution. Finally, each brain was embedded in a gelatin capsule that had been filled with HM20 and polymerized at 50°C for 24 hours.

#### Whole-brain imaging for single CLA principal neuron tracing

The whole-brain images were scanned with a fluorescence microscopic optical sectioning tomography (fMOST) system (Peng et al., 2021). This system runs in a wide-field block-face mode but is updated with a new principle to get better image contrast and speed and thus enables high throughput imaging of the fluorescence protein labeled sample. A block-face fluorescence imaging across the whole coronal plane (X-Y axes) was scanned and a section (Z-axis) was removed by a diamond knife, and the same procedures were repeated. The thickness of each section is 1.0 μm. To cover a whole coronal plane, multiple images were stitched together for making a montage. These procedures were repeated for a whole-brain dataset. The objective used is 40X WI with a numerical aperture (NA) 0.8 to provide a designed optical resolution (at 520 nm) of 0.35 μm in XY axes. The final resolution of images is 0.35 x 0.35 x 1.0 μ m voxel. The imaging parameters for GFP are 488 nm for an excitation wavelength and 510-550 nm for an emission filter with a passing band.

#### Full morphological reconstruction of single CLA principal neurons

The full morphological reconstruction procedure has been described previously (Peng et al., 2021). To reconstruct the complete morphology of a single CLA neuron in whole-brain, we used Vaa3D, open-source software for visualization, reconstruction, and analysis. The new modules TeraFly and TeraVR which improved both efficiency and precision of the reconstruction were incorporated into Vaa3D. TeraFly supports visualization and annotation of multidimensional imaging data with virtually unlimited scales. The out-of-core data management of TeraFly allows the software to smoothly deal with terabyte-scale of data even on a portable workstation with normal RAM size. Driven by virtual reality (VR) technologies, TeraVR is an annotation tool for immersive neuron reconstruction, which has been proved to be critical for achieving precision and efficiency in 3D data reconstruction. It creates stereo visualization for image volumes and reconstructions and offers an intuitive interface for the user to interact with large-scale data. Initially reconstructed several CLA neurons with Neurolucida 360 (NL360, MBF Bioscience, USA) were loaded to Vaa3D and some misconnected segments were corrected for those cells. In some cases, however, two or multiple similar-looking segments were intertwined or bundled together so closely that could not be separated even using Tera-VR. For these special cases, an extending segment and its branches were kept if it was continued reaching all ends or given up if it was eventually connected to a main axonal stem that connect to another soma. A neuron was considered fully reconstructed in the case that it was completed with all ends that typically had very well labeled, enlarged boutons. Finally, a quality control (QC) procedure was performed by an experienced annotator using TeraVR by double-checking the entire reconstruction of each reconstructed neuron, at high magnification by paying special attention to the proximal axonal part or a main axonal trunk of an axon cluster where axonal collaterals often emerged and branches but more frequently missed due to the local image environment being composed of crowded high contrasting structures. Auto-refinement fits the tracing to the center of fluorescent signals as the last step. The final reconstruction is a single tree without breaks, loops, multiple branches from a single point. In total, 52 single principal neurons within the CLA were reconstructed and 29 of them have been used in our previous study (Peng et al., 2021). All these single neurons were analyzed and visualized in the current study.

#### Image registration of reconstructed single CLA principal neurons to the CCFv3

3D registration from fMOST images (subject) to average mouse brain template of CCFv3 (target) were performed step by step: (1) Original fMOST images were down-sampled to 64x64x16 (X, Y, Z) to roughly match the size of the target brain; (2) 2D stripe-removal of fMOST images was performed using frequency notch filters; (3) A dozen matching landmark pairs between subject and target were manually added to ensure a correct affine transformation that approximately aligned the orientation and scales; (4) Affine transformation was applied to minimize the sum of squared difference of intensity between target and subject images; (5) Intensity was normalized by matching the local average intensity of subject image to that of target image; (6) A candidate list of landmarks across CCF space was generated by grid search (grid size = 16 pixels); (7) The BrainAligner software was used for searching corresponding landmarks in the subject image and performing the local alignment.

#### Quantification of axon terminals of single CLA principal neurons

SWC files were processed and examined with Vaa3D plugins to ensure topological correctness: sorted single tree with root node as soma. SWC files were resampled with a step size of 5 microns. In this study, we used CCFv3 ontology, a manually curated set of 316 non-overlapping structures named “summary structures”. Ipsi- and contra- lateral sides of brain regions to the somas were calculated separately. To quantitatively analyze the distribution of voxels of axons in brain-wide targets, fMOST images, and 3D reconstructed individual single CLA neurons were registered into the CCFv3. The axonal lengths in targets derived from single CLA principal neurons were measured as a proxy for projection strengths.

#### Branch analysis of single CLA principal neurons

A branch point is defined as a node with more than 1 child, except for the soma. The level of a branch point is the size of its descendent tree, measured by the percentage of the complete neuron.

#### Hierarchical clustering analysis of single CLA principal neurons

First, we manually separated 52 CLA single neurons into 2 major classes: ipsilateral (n = 42) and bilateral projecting neurons (n = 10). We then used the unsupervised hierarchical clustering analysis (Morpheus (broadinstitute.org)) to further divide them into 7 and 2 clusters based on 13 morphological features, respectively. These morphological features are intuitive to show the similarity and dissimilarity among individual single neurons. Feature 1 measures the lateral extension ratio of axons between the ACA/RSP and somatosensory/dorsal stream visual areas. Features 2, 3 and 4 measure the length ratio of axons from the left side, right side and anterior side of the somas, respectively. Feature 5 measures the anterior projection length ratio. Feature 6 measures posterior projection ratio, which is calculated by a formula = 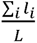, where *l_i_* denotes the axon length from projection cluster, *i* and *L* denote the total axon length of the cell. Feature 7 measures posterior projection efficacy, which is calculated by a formula = 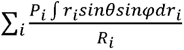, where *P_i_* denotes the *i^th^* posterior projection ratio (feature 7); *R_i_* denotes the *i^th^* projecting path length: 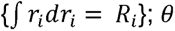 denotes the angle formed by projecting vectors with regard to soma on transverse plane; φ denotes the angle formed by projecting vectors with regard to soma on sagittal plane. Note that a straight projecting path toward posterior side yields the highest value for this feature due to *sinθ* and *sinφ* both equal to 1. Features 8 and 9 measure axon length ratios and axonal arbor width of the VISl-VISpl-ENT, respectively. Feature 10 measures cluster width that is extended from RSP. Feature 11 measures the “crown” forming length. This feature looks for the largest circle formed by one anterior projection and one posterior projection. The circle is formed when the two projecting clusters meet. Feature 12 reveals whether projections to the RSP. Feature 13 measures the length ratio of axons in the RSP. Each feature was normalized to the same scale so that Euclidean distance can be used to make these clusters better discerned throughout various heights in the dendrogram.

Features 1-11 were used for clustering the ipsilateral projecting neurons in the dendrogram (**Figure 6A**) and Features 12 and 13 were used for clustering the bilateral projecting neurons (**Figure 6B**).

### DATA AND SOFTWARE AVAILABILITY

Anterograde viral tracing data from The Allen Mouse Brain Connectivity Atlas are publicly available at our portal (http://portal.brain-map.org/).

Multimodal Reference Datasets, including images of the transgenic lines, are publicly available at our portal (https://connectivity.brain-map.org/static/referencedata).

CCFv3 is available at the Allen Brain Reference Atlases portal (http://atlas.brain-map.org/). The average template and associated annotation images can be manually downloaded using the Atlas Viewer, while images can be downloaded en masse using the AllenSDK (https://allensdk.readthedocs.io/en/latest/_static/examples/nb/image_download.html).

The AllenSDK is hosted on Github (https://github.com/alleninstitute/allensdk) and its documentation on Read The Docs (https://allensdk.rtfd.io). Registration code is available at (https://github.com/AllenInstitute/stpt_registration); see critical documentation at (https://github.com/AllenInstitute/stpt_registration/blob/master/README.md).

Our web API is documented at http://help.brain-map.org/. Other data files include the transgenic lines registered and downsampled to 25 μm resolution.

Rabies injections and reconstructed single CLA neurons are publicly available at the BIL.

## ADDITIONAL RESOURCES

Detailed protocol descriptions for all procedures can be found at the Allen Mouse Brain Connectivity Atlas documentation page (http://help.brain-map.org//display/mouseconnectivity/Documentation).

## KEY RESOURCES TABLE

For Bacterial and Virus Strains, Chemicals, Peptides, and Recombinant Proteins, Experimental Models: Organisms/Strains, Software and Algorithms, and Other, please see Key Resources Table.

