## Supplementary Stable 1 for "Regional and cell type-specific afferent and efferent projections of the mouse claustrum"

| review | image-id | mouse-line | dataset-name | tracer-type | graph-order |
|---|---|---|---|---|---|
| | 100141214 | C57Bl/6J | mca_classic | AAV | 473 |
| | 100141222 | C57Bl/6J | mca_classic | AAV | 380 |
| | 100141223 | C57Bl/6J | mca_classic | AAV | 649 |
| | 100141434 | C57Bl/6J | mca_classic | AAV | 792 |
| | 100141435 | C57Bl/6J | mca_classic | AAV | 588 |
| | 100141596 | C57Bl/6J | mca_classic | AAV | 812 |
| | 100141597 | C57Bl/6J | mca_classic | AAV | 617 |
| | 100141598 | C57Bl/6J | mca_classic | AAV | 662 |
| | 100141993 | C57Bl/6J | mca_classic | AAV | 822 |
| | 100142569 | C57Bl/6J | mca_classic | AAV | 675 |
| | 100142580 | C57Bl/6J | mca_classic | AAV | 573 |
| | 100147785 | C57Bl/6J | mca_classic | AAV | 669 |
| | 100147861 | C57Bl/6J | mca_classic | AAV | 457 |
| | 100148143 | C57Bl/6J | mca_classic | AAV | 473 |
| | 100148443 | C57Bl/6J | mca_classic | AAV | 457 |
| | 100148554 | C57Bl/6J | mca_classic | AAV | 523 |
| | 112167395 | C57Bl/6J | mca_classic | AAV | 473 |
| | 112228391 | C57Bl/6J | mca_classic | AAV | 801 |
| | 112307046 | C57Bl/6J | mca_classic | AAV | 416 |
| | 112307754 | C57Bl/6J | mca_classic | AAV | 573 |
| | 112308468 | C57Bl/6J | mca_classic | AAV | 467 |
| | 112372418 | C57Bl/6J | mca_classic | AAV | 794 |
| | 112424102 | C57Bl/6J | mca_classic | AAV | 1033 |
| | 112425523 | C57Bl/6J | mca_classic | AAV | 792 |
| | 112458831 | C57Bl/6J | mca_classic | AAV | 573 |
| | 112459547 | C57Bl/6J | mca_classic | AAV | 595 |
| | 112460257 | C57Bl/6J | mca_classic | AAV | 1067 |
| Y | 112514202 | C57BL/6J | mca_classic | AAV | 232 |
| Y | 112596790 | C57BL/6J | mca_classic | AAV | 279 |
| | 112597496 | C57Bl/6J | mca_classic | AAV | 410 |
| | 112672268 | C57Bl/6J | mca_classic | AAV | 473 |
| | 112672974 | C57Bl/6J | mca_classic | AAV | 507 |
| | 112745073 | C57Bl/6J | mca_classic | AAV | 467 |
| | 112745787 | C57Bl/6J | mca_classic | AAV | 473 |
| | 112789899 | C57Bl/6J | mca_classic | AAV | 942 |
| | 112826458 | C57Bl/6J | mca_classic | AAV | 956 |
| | 112827164 | C57Bl/6J | mca_classic | AAV | 808 |
| | 112827872 | C57Bl/6J | mca_classic | AAV | 1071 |
| | 112951097 | Oxt-IRES-Cre | mca_classic | AAV | 720 |
| | 113037759 | Oxt-IRES-Cre | mca_classic | AAV | 801 |
| | 113095845 | C57Bl/6J | mca_classic | AAV | 802 |
| | 113096571 | C57Bl/6J | mca_classic | AAV | 838 |
| | 113144533 | C57Bl/6J | mca_classic | AAV | 562 |
| | 113165340 | Etv1-CreERT2 | mca_classic | AAV | 671 |
| | 113166056 | C57Bl/6J | mca_classic | AAV | 467 |
| | 113225519 | Lepr-IRES-Cre | mca_classic | AAV | 794 |

| | | | | |
|---|---|---|---|---|
| 113226232 | C57Bl/6J | mca_classic | AAV | 507 |
| 113313632 | C57Bl/6J | mca_classic | AAV | 794 |
| 113314337 | Lepr-IRES-Cre | mca_classic | AAV | 739 |
| 113368085 | C57Bl/6J | mca_classic | AAV | 812 |
| 113368898 | Drd1a-Cre_EY2( | mca_classic | AAV | 535 |
| 113369603 | C57Bl/6J | mca_classic | AAV | 794 |
| 113399428 | C57Bl/6J | mca_classic | AAV | 826 |
| 113400134 | C57Bl/6J | mca_classic | AAV | 826 |
| 113442864 | C57Bl/6J | mca_classic | AAV | 848 |
| 113443572 | C57Bl/6J | mca_classic | AAV | 390 |
| 113444277 | Pmch-Cre | mca_classic | AAV | 794 |
| 113504763 | C57Bl/6J | mca_classic | AAV | 694 |
| 113505468 | C57Bl/6J | mca_classic | AAV | 573 |
| 113506174 | Pmch-Cre | mca_classic | AAV | 794 |
| 113553300 | C57Bl/6J | mca_classic | AAV | 795 |
| 113554008 | Gal-Cre_KI87 | mca_classic | AAV | 682 |
| 113554719 | C57Bl/6J | mca_classic | AAV | 744 |
| 113696423 | Gal-Cre_KI87 | mca_classic | AAV | 617 |
| 113766038 | C57Bl/6J | mca_classic | AAV | 573 |
| 113766744 | Grik4-Cre | mca_classic | AAV | 967 |
| 113780276 | Pomc-Cre_ST | mca_classic | AAV | 473 |
| 113783321 | C57Bl/6J | mca_classic | AAV | 589 |
| 113784293 | C57Bl/6J | mca_classic | AAV | 682 |
| 113846682 | C57Bl/6J | mca_classic | AAV | 844 |
| 113884251 | C57Bl/6J | mca_classic | AAV | 644 |
| 113888575 | Scnn1a-Tg3-Cre | mca_classic | AAV | 1067 |
| 113933871 | Grik4-Cre | mca_classic | AAV | 473 |
| 113934579 | Scnn1a-Tg3-Cre | mca_classic | AAV | 527 |
| 113935285 | C57Bl/6J | mca_classic | AAV | 467 |
| 113935990 | C57Bl/6J | mca_classic | AAV | 566 |
| 113936696 | C57Bl/6J | mca_classic | AAV | 1067 |
| 114008220 | C57Bl/6J | mca_classic | AAV | 473 |
| 114009636 | Grik4-Cre | mca_classic | AAV | 473 |
| 114045733 | C57Bl/6J | mca_classic | AAV | 770 |
| 114046440 | C57Bl/6J | mca_classic | AAV | 794 |
| 114047146 | Grik4-Cre | mca_classic | AAV | 473 |
| 114155190 | Slc6a4-CreERT2 | mca_classic | AAV | 881 |
| 114155923 | C57Bl/6J | mca_classic | AAV | 838 |
| 114248377 | Slc6a4-CreERT2 | mca_classic | AAV | 881 |
| 114249084 | C57Bl/6J | mca_classic | AAV | 431 |
| 114290225 | Nr5a1-Cre | mca_classic | AAV | 787 |
| 114291646 | C57Bl/6J | mca_classic | AAV | 685 |
| 114399224 | C57Bl/6J | mca_classic | AAV | 473 |
| 114400640 | C57Bl/6J | mca_classic | AAV | 945 |
| 114402050 | C57Bl/6J | mca_classic | AAV | 954 |
| 114427219 | C57Bl/6J | mca_classic | AAV | 675 |
| 114428632 | Etv1-CreERT2 | mca_classic | AAV | 226 |

| | | | | |
|---|---|---|---|---|
| 114429338 | C57Bl/6J | mca_classic | AAV | 467 |
| 114430043 | C57Bl/6J | mca_classic | AAV | 473 |
| 114472145 | C57Bl/6J | mca_classic | AAV | 494 |
| 114472860 | C57Bl/6J | mca_classic | AAV | 776 |
| 114473794 | C57Bl/6J | mca_classic | AAV | 1033 |
| 114474520 | C57Bl/6J | mca_classic | AAV | 826 |
| 114475228 | Etv1-CreERT2 | mca_classic | AAV | 232 |
| 114752488 | Nr5a1-Cre | mca_classic | AAV | 122 |
| 114754390 | C57Bl/6J | mca_classic | AAV | 808 |
| 114755099 | C57Bl/6J | mca_classic | AAV | 390 |
| 114755811 | C57Bl/6J | mca_classic | AAV | 889 |
| 115956702 | Nr5a1-Cre | mca_classic | AAV | 51 |
| 115958115 | C57Bl/6J | mca_classic | AAV | 566 |
| 116900714 | C57Bl/6J | mca_classic | AAV | 457 |
| 116904684 | Gal-Cre_KI87 | mca_classic | AAV | 685 |
| 116905391 | C57Bl/6J | mca_classic | AAV | 812 |
| 117302771 | C57Bl/6J | mca_classic | AAV | 794 |
| 117312486 | C57Bl/6J | mca_classic | AAV | 621 |
| 117316260 | C57Bl/6J | mca_classic | AAV | 703 |
| 119846129 | C57Bl/6J | mca_classic | AAV | 885 |
| 119846838 | C57Bl/6J | mca_classic | AAV | 744 |
| 119847544 | C57Bl/6J | mca_classic | AAV | 1088 |
| 119848249 | C57Bl/6J | mca_classic | AAV | 1024 |
| 120280191 | C57Bl/6J | mca_classic | AAV | 776 |
| 120280939 | C57Bl/6J | mca_classic | AAV | 523 |
| 120281646 | Erbb4-T2A-CreE | mca_classic | AAV | 595 |
| 120282354 | C57Bl/6J | mca_classic | AAV | 562 |
| 120436274 | C57Bl/6J | mca_classic | AAV | 457 |
| 120436988 | C57Bl/6J | mca_classic | AAV | 587 |
| 120438416 | C57Bl/6J | mca_classic | AAV | 1088 |
| 120493315 | C57Bl/6J | mca_classic | AAV | 1098 |
| 120494024 | Ntsr1-Cre_GN2 | mca_classic | AAV | 100 |
| 120494729 | C57Bl/6J | mca_classic | AAV | 457 |
| 120571672 | C57Bl/6J | mca_classic | AAV | 838 |
| 120572378 | C57Bl/6J | mca_classic | AAV | 823 |
| 120584036 | Ntsr1-Cre_GN2 | mca_classic | AAV | 24 |
| 120760759 | C57Bl/6J | mca_classic | AAV | 826 |
| 120761491 | C57Bl/6J | mca_classic | AAV | 838 |
| 120811946 | C57Bl/6J | mca_classic | AAV | 853 |
| 120812686 | C57Bl/6J | mca_classic | AAV | 386 |
| 120814821 | Rbp4-Cre_KL10 | mca_classic | AAV | 24 |
| 120874405 | C57Bl/6J | mca_classic | AAV | 1062 |
| 120875111 | C57Bl/6J | mca_classic | AAV | 692 |
| 120875816 | Rbp4-Cre_KL10 | mca_classic | AAV | 44 |
| 120916102 | Rbp4-Cre_KL10 | mca_classic | AAV | 100 |
| 121145045 | C57Bl/6J | mca_classic | AAV | 918 |
| 121145750 | C57Bl/6J | mca_classic | AAV | 975 |

| | | | | |
|---|---|---|---|---|
| 121146455 | C57Bl/6J | mca_classic | AAV | 1003 |
| 121509005 | C57Bl/6J | mca_classic | AAV | 826 |
| 121509711 | C57Bl/6J | mca_classic | AAV | 390 |
| 122640358 | C57Bl/6J | mca_classic | AAV | 985 |
| 122641078 | C57Bl/6J | mca_classic | AAV | 444 |
| 122641784 | C57Bl/6J | mca_classic | AAV | 457 |
| 122642490 | Syt6-Cre_KI148 | mca_classic | AAV | 24 |
| 123662982 | C57Bl/6J | mca_classic | AAV | 693 |
| 123663689 | C57Bl/6J | mca_classic | AAV | 967 |
| 123664417 | Syt6-Cre_KI148 | mca_classic | AAV | 44 |
| 124059700 | C57Bl/6J | mca_classic | AAV | 573 |
| 124060405 | Syt6-Cre_KI148 | mca_classic | AAV | 136 |
| 125361005 | C57Bl/6J | mca_classic | AAV | 600 |
| 125363160 | C57Bl/6J | mca_classic | AAV | 844 |
| 125435783 | Syt6-Cre_KI148 | mca_classic | AAV | 185 |
| 125436508 | C57Bl/6J | mca_classic | AAV | 613 |
| 125437216 | C57Bl/6J | mca_classic | AAV | 913 |
| 125437921 | C57Bl/6J | mca_classic | AAV | 975 |
| 125801033 | Syt6-Cre_KI148 | mca_classic | AAV | 232 |
| 125801739 | C57Bl/6J | mca_classic | AAV | 1080 |
| 125831616 | C57Bl/6J | mca_classic | AAV | 1054 |
| 125832322 | C57Bl/6J | mca_classic | AAV | 566 |
| 125833030 | Rbp4-Cre_KL10 | mca_classic | AAV | 226 |
| 126115436 | C57Bl/6J | mca_classic | AAV | 554 |
| 126116142 | C57Bl/6J | mca_classic | AAV | 757 |
| 126116848 | C57Bl/6J | mca_classic | AAV | 494 |
| 126117554 | Rbp4-Cre_KL10 | mca_classic | AAV | 232 |
| 126188607 | C57Bl/6J | mca_classic | AAV | 978 |
| 126189322 | C57Bl/6J | mca_classic | AAV | 988 |
| 126190033 | Syt6-Cre_KI148 | mca_classic | AAV | 232 |
| 126190743 | C57Bl/6J | mca_classic | AAV | 802 |
| 126351299 | C57Bl/6J | mca_classic | AAV | 1004 |
| 126352037 | C57Bl/6J | mca_classic | AAV | 826 |
| 126352744 | C57Bl/6J | mca_classic | AAV | 955 |
| 126353451 | Syt6-Cre_KI148 | mca_classic | AAV | 258 |
| 126522350 | C57Bl/6J | mca_classic | AAV | 852 |
| 126523066 | C57Bl/6J | mca_classic | AAV | 830 |
| 126523791 | C57Bl/6J | mca_classic | AAV | 535 |
| 126646502 | C57Bl/6J | mca_classic | AAV | 830 |
| 126653015 | C57Bl/6J | mca_classic | AAV | 449 |
| 126709328 | C57Bl/6J | mca_classic | AAV | 978 |
| 126710034 | C57Bl/6J | mca_classic | AAV | 975 |
| 126710740 | C57Bl/6J | mca_classic | AAV | 764 |
| 126711445 | Chat-IRES-Cre-n | mca_classic | AAV | 613 |
| 126841788 | Chat-IRES-Cre-n | mca_classic | AAV | 613 |
| 126843200 | C57Bl/6J | mca_classic | AAV | 693 |
| 126843905 | C57Bl/6J | mca_classic | AAV | 975 |

| | | | | |
|---|---|---|---|---|
| 126853068 | Kcnc2-Cre | mca_classic | AAV | 573 |
| 126863090 | Kcnc2-Cre | mca_classic | AAV | 561 |
| 126908712 | Slc6a5-Cre_KF1 | mca_classic | AAV | 954 |
| 127041126 | C57Bl/6J | mca_classic | AAV | 978 |
| 127041832 | C57Bl/6J | mca_classic | AAV | 975 |
| 127042540 | C57Bl/6J | mca_classic | AAV | 992 |
| 127083591 | C57Bl/6J | mca_classic | AAV | 844 |
| 127085005 | Pvalb-IRES-Cre | mca_classic | AAV | 703 |
| 127085717 | C57Bl/6J | mca_classic | AAV | 890 |
| 127090378 | Pvalb-IRES-Cre | mca_classic | AAV | 703 |
| 127091083 | C57Bl/6J | mca_classic | AAV | 817 |
| 127139568 | C57Bl/6J | mca_classic | AAV | 507 |
| 127140981 | Drd2-Cre_ER44 | mca_classic | AAV | 573 |
| 127223428 | C57Bl/6J | mca_classic | AAV | 931 |
| 127224133 | Drd2-Cre_ER44 | mca_classic | AAV | 573 |
| 127255254 | C57Bl/6J | mca_classic | AAV | 575 |
| 127255962 | C57Bl/6J | mca_classic | AAV | 692 |
| 127349111 | C57Bl/6J | mca_classic | AAV | 992 |
| 127350480 | C57Bl/6J | mca_classic | AAV | 1007 |
| 127353220 | C57Bl/6J | mca_classic | AAV | 978 |
| 127396051 | C57Bl/6J | mca_classic | AAV | 702 |
| 127396760 | Sim1-Cre_KJ18 | mca_classic | AAV | 764 |
| 127397469 | C57Bl/6J | mca_classic | AAV | 494 |
| 127398177 | C57Bl/6J | mca_classic | AAV | 1024 |
| 127468854 | C57Bl/6J | mca_classic | AAV | 703 |
| 127469566 | C57Bl/6J | mca_classic | AAV | 890 |
| 127470271 | C57Bl/6J | mca_classic | AAV | 794 |
| 127470976 | Ucn3-Cre_KF43 | mca_classic | AAV | 720 |
| 127557915 | C57Bl/6J | mca_classic | AAV | 826 |
| 127560043 | Syt17-Cre_NO1 | mca_classic | AAV | 272 |
| 127649713 | C57Bl/6J | mca_classic | AAV | 757 |
| 127650431 | C57Bl/6J | mca_classic | AAV | 1097 |
| 127651139 | Syt17-Cre_NO1 | mca_classic | AAV | 823 |
| 127710392 | C57Bl/6J | mca_classic | AAV | 792 |
| 127711098 | C57Bl/6J | mca_classic | AAV | 523 |
| 127711803 | C57Bl/6J | mca_classic | AAV | 573 |
| 127761449 | C57Bl/6J | mca_classic | AAV | 595 |
| 127762867 | C57Bl/6J | mca_classic | AAV | 573 |
| 127796728 | C57Bl/6J | mca_classic | AAV | 823 |
| 127797441 | C57Bl/6J | mca_classic | AAV | 645 |
| 127798146 | Syt17-Cre_NO1 | mca_classic | AAV | 823 |
| 127865687 | C57Bl/6J | mca_classic | AAV | 826 |
| 127867804 | Erbb4-T2A-CreE | mca_classic | AAV | 823 |
| 127907465 | C57Bl/6J | mca_classic | AAV | 416 |
| 127908173 | C57Bl/6J | mca_classic | AAV | 996 |
| 127908879 | Erbb4-T2A-CreE | mca_classic | AAV | 24 |
| 127909584 | C57Bl/6J | mca_classic | AAV | 776 |

| | | | | | |
|---|---|---|---|---|---|
| | 127991964 | C57Bl/6J | mca_classic | AAV | 826 |
| | 127993375 | Erbb4-T2A-CreE | mca_classic | AAV | 44 |
| | 128001349 | C57Bl/6J | mca_classic | AAV | 830 |
| | 128002057 | C57Bl/6J | mca_classic | AAV | 838 |
| | 128003477 | Slc6a5-Cre_KF1 | mca_classic | AAV | 948 |
| | 128055110 | Slc6a4-Cre_ET3 | mca_classic | AAV | 881 |
| | 128056535 | C57Bl/6J | mca_classic | AAV | 1004 |
| | 129567943 | C57Bl/6J | mca_classic | AAV | 1004 |
| Y | 129573239 | C57Bl/6J | mca_classic | AAV | 431 |
| | 131068390 | C57Bl/6J | mca_classic | AAV | 416 |
| | 133286030 | C57Bl/6J | mca_classic | AAV | 398 |
| | 133286781 | Slc6a4-Cre_ET3 | mca_classic | AAV | 1013 |
| | 138058320 | C57Bl/6J | mca_classic | AAV | 613 |
| | 138059031 | C57Bl/6J | mca_classic | AAV | 738 |
| | 139311530 | C57Bl/6J | mca_classic | AAV | 826 |
| Y | 139519496 | C57BL/6J | mca_classic | AAV | 122 |
| | 141603895 | Slc6a4-Cre_ET3 | mca_classic | AAV | 1013 |
| | 142653395 | Pmch-Cre | mca_classic | AAV | 794 |
| | 142654100 | C57Bl/6J | mca_classic | AAV | 494 |
| | 142654808 | C57Bl/6J | mca_classic | AAV | 1071 |
| | 142655513 | C57Bl/6J | mca_classic | AAV | 911 |
| | 142656218 | C57Bl/6J | mca_classic | AAV | 494 |
| | 143512399 | Pnmt-Cre | mca_classic | AAV | 1003 |
| | 143515102 | C57Bl/6J | mca_classic | AAV | 942 |
| | 146012184 | C57Bl/6J | mca_classic | AAV | 1024 |
| | 146012934 | C57Bl/6J | mca_classic | AAV | 566 |
| | 146045723 | C57Bl/6J | mca_classic | AAV | 613 |
| | 146046430 | C57Bl/6J | mca_classic | AAV | 675 |
| | 146078721 | C57Bl/6J | mca_classic | AAV | 830 |
| | 146470726 | C57Bl/6J | mca_classic | AAV | 575 |
| | 146553266 | C57Bl/6J | mca_classic | AAV | 573 |
| | 146553971 | C57Bl/6J | mca_classic | AAV | 494 |
| | 146554676 | Agrp-IRES-Cre | mca_classic | AAV | 733 |
| | 146658170 | C57Bl/6J | mca_classic | AAV | 676 |
| | 146658879 | C57Bl/6J | mca_classic | AAV | 668 |
| | 146659588 | C57Bl/6J | mca_classic | AAV | 781 |
| | 146660293 | C57Bl/6J | mca_classic | AAV | 757 |
| | 146660999 | Agrp-IRES-Cre | mca_classic | AAV | 733 |
| | 146747721 | C57Bl/6J | mca_classic | AAV | 621 |
| | 146794439 | C57Bl/6J | mca_classic | AAV | 890 |
| | 146795148 | C57Bl/6J | mca_classic | AAV | 595 |
| | 146856593 | C57Bl/6J | mca_classic | AAV | 954 |
| | 146857301 | C57Bl/6J | mca_classic | AAV | 416 |
| | 146859480 | Cdhr1-Cre_KG6 | mca_classic | AAV | 380 |
| | 146921849 | Cdhr1-Cre_KG6 | mca_classic | AAV | 380 |
| | 146983504 | Oxt-IRES-Cre | mca_classic | AAV | 720 |
| | 146984209 | C57Bl/6J | mca_classic | AAV | 416 |

| | | | | |
|---|---|---|---|---|
| 146984915 | C57Bl/6J | mca_classic | AAV | 531 |
| 146985623 | C57Bl/6J | mca_classic | AAV | 600 |
| 146986331 | C57Bl/6J | mca_classic | AAV | 800 |
| 147049515 | C57Bl/6J | mca_classic | AAV | 621 |
| 147051682 | Oxt-IRES-Cre | mca_classic | AAV | 782 |
| 147134401 | C57Bl/6J | mca_classic | AAV | 1024 |
| 147135107 | C57Bl/6J | mca_classic | AAV | 698 |
| 147135812 | C57Bl/6J | mca_classic | AAV | 918 |
| 147136518 | Oxt-IRES-Cre | mca_classic | AAV | 720 |
| 147159899 | C57Bl/6J | mca_classic | AAV | 975 |
| 147162027 | C57Bl/6J | mca_classic | AAV | 812 |
| 147162736 | C57Bl/6J | mca_classic | AAV | 590 |
| 147212977 | C57Bl/6J | mca_classic | AAV | 707 |
| 147214398 | C57Bl/6J | mca_classic | AAV | 967 |
| 147215105 | C57Bl/6J | mca_classic | AAV | 954 |
| 147353537 | C57Bl/6J | mca_classic | AAV | 713 |
| 147354242 | C57Bl/6J | mca_classic | AAV | 826 |
| 147354951 | C57Bl/6J | mca_classic | AAV | 988 |
| 147632458 | C57Bl/6J | mca_classic | AAV | 613 |
| 147633169 | Pcp2-Cre_GN13 | mca_classic | AAV | 1071 |
| 147633885 | C57Bl/6J | mca_classic | AAV | 913 |
| 147635309 | C57Bl/6J | mca_classic | AAV | 838 |
| 147706228 | C57Bl/6J | mca_classic | AAV | 913 |
| 147708352 | Pcp2-Cre_GN13 | mca_classic | AAV | 1024 |
| 147787606 | C57Bl/6J | mca_classic | AAV | 689 |
| 147789031 | C57Bl/6J | mca_classic | AAV | 844 |
| 147790181 | C57Bl/6J | mca_classic | AAV | 978 |
| 147790922 | Oxt-IRES-Cre | mca_classic | AAV | 717 |
| 147968866 | Oxt-IRES-Cre | mca_classic | AAV | 717 |
| 148197327 | C57Bl/6J | mca_classic | AAV | 764 |
| 148198052 | C57Bl/6J | mca_classic | AAV | 573 |
| 152995635 | C57Bl/6J | mca_classic | AAV | 931 |
| 155735108 | Pnmt-Cre | mca_classic | AAV | 948 |
| 155735826 | Slc6a5-Cre_KF1 | mca_classic | AAV | 679 |
| 155736539 | Drd2-Cre_ER44 | mca_classic | AAV | 573 |
| 156195758 | Sim1-Cre_KJ18 | mca_classic | AAV | 794 |
| 156198187 | Scnn1a-Tg2-Cre | mca_classic | AAV | 662 |
| 156202979 | Scnn1a-Tg2-Cre | mca_classic | AAV | 649 |
| 156252954 | Gad2-IRES-Cre | mca_classic | AAV | 703 |
| 156253662 | Pnmt-Cre | mca_classic | AAV | 988 |
| 156254369 | Gad2-IRES-Cre | mca_classic | AAV | 703 |
| 156255078 | Scnn1a-Tg3-Cre | mca_classic | AAV | 1033 |
| 156313344 | Scnn1a-Tg3-Cre | mca_classic | AAV | 885 |
| 156314054 | Scnn1a-Tg3-Cre | mca_classic | AAV | 527 |
| 156314762 | Th-IRES-CreER | mca_classic | AAV | 823 |
| 156315468 | Gad2-IRES-Cre | mca_classic | AAV | 802 |
| 156393801 | Gal-Cre_KI87 | mca_classic | AAV | 676 |

| | | | | |
|---|---|---|---|---|
| 156394513 | A930038C07Rik | mca_classic | AAV | 24 |
| 156395997 | A930038C07Rik | mca_classic | AAV | 100 |
| 156492394 | Etv1-CreERT2 | mca_classic | AAV | 24 |
| 156493815 | Etv1-CreERT2 | mca_classic | AAV | 122 |
| 156494530 | Etv1-CreERT2 | mca_classic | AAV | 185 |
| 156545918 | Ntsr1-Cre_GN2: | mca_classic | AAV | 185 |
| 156546627 | Ntsr1-Cre_GN2: | mca_classic | AAV | 232 |
| 156670520 | Drd2-Cre_ER44 | mca_classic | AAV | 573 |
| 156671933 | Ntsr1-Cre_GN2: | mca_classic | AAV | 171 |
| 156741826 | Rbp4-Cre_KL10 | mca_classic | AAV | 258 |
| 156742543 | Erbb4-T2A-CreE | mca_classic | AAV | 226 |
| 156743264 | Erbb4-T2A-CreE | mca_classic | AAV | 100 |
| 156784823 | Syt6-Cre_KI148 | mca_classic | AAV | 185 |
| 156785529 | Syt6-Cre_KI148 | mca_classic | AAV | 226 |
| 156786234 | Syt6-Cre_KI148 | mca_classic | AAV | 18 |
| 156786939 | Syt6-Cre_KI148 | mca_classic | AAV | 213 |
| 156819600 | Chat-IRES-Cre-n | mca_classic | AAV | 617 |
| 156872043 | C57Bl/6J | mca_classic | AAV | 890 |
| 156929391 | C57Bl/6J | mca_classic | AAV | 925 |
| 156930126 | C57Bl/6J | mca_classic | AAV | 885 |
| 156931568 | C57Bl/6J | mca_classic | AAV | 685 |
| 156978574 | C57Bl/6J | mca_classic | AAV | 826 |
| 156979283 | C57Bl/6J | mca_classic | AAV | 838 |
| 156979988 | C57Bl/6J | mca_classic | AAV | 889 |
| 157063074 | C57Bl/6J | mca_classic | AAV | 645 |
| 157063781 | C57Bl/6J | mca_classic | AAV | 544 |
| 157549402 | C57Bl/6J | mca_classic | AAV | 787 |
| 157550122 | C57Bl/6J | mca_classic | AAV | 600 |
| 157654069 | C57Bl/6J | mca_classic | AAV | 416 |
| 157659671 | C57Bl/6J | mca_classic | AAV | 613 |
| 157711043 | Syt6-Cre_KI148 | mca_classic | AAV | 332 |
| 157712456 | Chat-IRES-Cre-n | mca_classic | AAV | 967 |
| 157765542 | Pdzk1ip1-Cre_K | mca_classic | AAV | 707 |
| 157766259 | Pdzk1ip1-Cre_K | mca_classic | AAV | 885 |
| 157767683 | Nr5a1-Cre | mca_classic | AAV | 185 |
| 157768393 | Ntsr1-Cre_GN2: | mca_classic | AAV | 65 |
| 157769139 | Syt6-Cre_KI148 | mca_classic | AAV | 346 |
| 157826227 | Cort-T2A-Cre | mca_classic | AAV | 24 |
| 157909001 | Ntsr1-Cre_GN2: | mca_classic | AAV | 18 |
| 157911126 | Slc6a5-Cre_KF1 | mca_classic | AAV | 1003 |
| 157911832 | Gad2-IRES-Cre | mca_classic | AAV | 573 |
| 157952068 | Hdc-Cre_IM1 | mca_classic | AAV | 781 |
| 157952778 | Hdc-Cre_IM1 | mca_classic | AAV | 763 |
| 157954927 | Gng7-Cre_KH71 | mca_classic | AAV | 136 |
| 157955639 | Gng7-Cre_KH71 | mca_classic | AAV | 185 |
| 158017916 | Gng7-Cre_KH71 | mca_classic | AAV | 44 |
| 158018630 | Gng7-Cre_KH71 | mca_classic | AAV | 226 |

| | | | | |
|---|---|---|---|---|
| 158019342 | Drd2-Cre_ER44 | mca_classic | AAV | 573 |
| 158020947 | Drd2-Cre_ER44 | mca_classic | AAV | 573 |
| 158138440 | Pvalb-IRES-Cre | mca_classic | AAV | 24 |
| 158139151 | Pvalb-IRES-Cre | mca_classic | AAV | 93 |
| 158139883 | Ntsr1-Cre_GN2: | mca_classic | AAV | 24 |
| 158141324 | Syt6-Cre_KI148 | mca_classic | AAV | 213 |
| 158142090 | Pomc-Cre_ST | mca_classic | AAV | 733 |
| 158255941 | A930038C07Rik | mca_classic | AAV | 185 |
| 158257355 | C57Bl/6J | mca_classic | AAV | 618 |
| 158258062 | C57Bl/6J | mca_classic | AAV | 787 |
| 158314987 | C57Bl/6J | mca_classic | AAV | 764 |
| 158315810 | C57Bl/6J | mca_classic | AAV | 744 |
| 158321996 | C57Bl/6J | mca_classic | AAV | 559 |
| 158373181 | C57Bl/6J | mca_classic | AAV | 794 |
| 158373958 | C57Bl/6J | mca_classic | AAV | 610 |
| 158374671 | C57Bl/6J | mca_classic | AAV | 416 |
| 158375425 | C57Bl/6J | mca_classic | AAV | 649 |
| 158376179 | C57Bl/6J | mca_classic | AAV | 838 |
| 158434409 | C57Bl/6J | mca_classic | AAV | 911 |
| 158435826 | C57Bl/6J | mca_classic | AAV | 1003 |
| 158737454 | C57Bl/6J | mca_classic | AAV | 826 |
| 158738180 | C57Bl/6J | mca_classic | AAV | 744 |
| 158738894 | C57Bl/6J | mca_classic | AAV | 1088 |
| 158838128 | C57Bl/6J | mca_classic | AAV | 913 |
| 158840459 | C57Bl/6J | mca_classic | AAV | 676 |
| 158841171 | C57Bl/6J | mca_classic | AAV | 698 |
| 158914182 | C57Bl/6J | mca_classic | AAV | 822 |
| 158915602 | C57Bl/6J | mca_classic | AAV | 846 |
| 158916311 | C57Bl/6J | mca_classic | AAV | 573 |
| 159021559 | C57Bl/6J | mca_classic | AAV | 985 |
| 159024474 | C57Bl/6J | mca_classic | AAV | 985 |
| 159097209 | Etv1-CreERT2 | mca_classic | AAV | 332 |
| 159151138 | Sim1-Cre_KJ18 | mca_classic | AAV | 421 |
| 159209586 | Syt6-Cre_KI148 | mca_classic | AAV | 264 |
| 159222295 | Sim1-Cre_KJ18 | mca_classic | AAV | 792 |
| 159223001 | Drd1a-Cre_EY2( | mca_classic | AAV | 573 |
| 159223769 | Syt6-Cre_KI148 | mca_classic | AAV | 51 |
| 159224478 | Syt6-Cre_KI148 | mca_classic | AAV | 178 |
| 159258618 | Pdzk1ip1-Cre_K | mca_classic | AAV | 653 |
| 159319654 | A930038C07Rik | mca_classic | AAV | 232 |
| 159320367 | Ntsr1-Cre_GN2: | mca_classic | AAV | 206 |
| 159321097 | Ntsr1-Cre_GN2: | mca_classic | AAV | 51 |
| 159321806 | Ntsr1-Cre_GN2: | mca_classic | AAV | 213 |
| 159322514 | Ntsr1-Cre_GN2: | mca_classic | AAV | 279 |
| 159329308 | Drd2-Cre_ER44 | mca_classic | AAV | 573 |
| 159330030 | Syt6-Cre_KI148 | mca_classic | AAV | 185 |
| 159330754 | Syt6-Cre_KI148 | mca_classic | AAV | 226 |

| | | | | |
|---|---|---|---|---|
| 159331462 | Cck-IRES-Cre | mca_classic | AAV | 693 |
| 159373612 | Ntsr1-Cre_GN2: | mca_classic | AAV | 24 |
| 159375743 | Sim1-Cre_KJ18 | mca_classic | AAV | 757 |
| 159432479 | Cck-IRES-Cre | mca_classic | AAV | 693 |
| 159433187 | Cck-IRES-Cre | mca_classic | AAV | 621 |
| 159433905 | Syt6-Cre_KI148 | mca_classic | AAV | 285 |
| 159508775 | Syt6-Cre_KI148 | mca_classic | AAV | 100 |
| 159510205 | Dbh-Cre_KH212 | mca_classic | AAV | 929 |
| 159511623 | Ntsr1-Cre_GN2: | mca_classic | AAV | 245 |
| 159550125 | Ntsr1-Cre_GN2: | mca_classic | AAV | 185 |
| 159551564 | Ntsr1-Cre_GN2: | mca_classic | AAV | 226 |
| 159552290 | Ntsr1-Cre_GN2: | mca_classic | AAV | 573 |
| 159554010 | Ntsr1-Cre_GN2: | mca_classic | AAV | 325 |
| 159602992 | Cart-Tg1-Cre | mca_classic | AAV | 51 |
| 159648854 | C57Bl/6J | mca_classic | AAV | 948 |
| 159649643 | C57Bl/6J | mca_classic | AAV | 792 |
| 159650350 | C57Bl/6J | mca_classic | AAV | 613 |
| 159651060 | Ntsr1-Cre_GN2: | mca_classic | AAV | 18 |
| 159651770 | Ntsr1-Cre_GN2: | mca_classic | AAV | 72 |
| 159750477 | Ntsr1-Cre_GN2: | mca_classic | AAV | 72 |
| 159751184 | Pomc-Cre_ST | mca_classic | AAV | 733 |
| 159753308 | Rbp4-Cre_KL10 | mca_classic | AAV | 171 |
| 159832064 | Rbp4-Cre_KL10 | mca_classic | AAV | 332 |
| 159887627 | Nr5a1-Cre | mca_classic | AAV | 258 |
| 159888336 | Scnn1a-Tg3-Cre | mca_classic | AAV | 51 |
| 159941339 | Drd1a-Cre_EY2( | mca_classic | AAV | 573 |
| 159942097 | Drd1a-Cre_EY2( | mca_classic | AAV | 610 |
| 159942862 | Dbh-Cre_KH212 | mca_classic | AAV | 890 |
| 159945113 | Gabra6-IRES-Cr | mca_classic | AAV | 1067 |
| 159994767 | Syt17-Cre_NO1 | mca_classic | AAV | 390 |
| 159995481 | Pdzk1ip1-Cre_K | mca_classic | AAV | 826 |
| 159996191 | Pomc-Cre_BL | mca_classic | AAV | 473 |
| 160079360 | Pomc-Cre_BL | mca_classic | AAV | 473 |
| 160080068 | Pvalb-IRES-Cre | mca_classic | AAV | 122 |
| 160080778 | Slc6a5-Cre_KF1 | mca_classic | AAV | 975 |
| 160081484 | Slc6a5-Cre_KF1 | mca_classic | AAV | 975 |
| 160150861 | Slc6a5-Cre_KF1 | mca_classic | AAV | 955 |
| 160151570 | Slc6a5-Cre_KF1 | mca_classic | AAV | 955 |
| 160152987 | Slc6a5-Cre_KF1 | mca_classic | AAV | 941 |
| 160153696 | Slc6a5-Cre_KF1 | mca_classic | AAV | 978 |
| 160294327 | Sst-Cre | mca_classic | AAV | 587 |
| 160296448 | Lepr-IRES-Cre | mca_classic | AAV | 739 |
| 160317628 | Lepr-IRES-Cre | mca_classic | AAV | 739 |
| 160398593 | Gnrh1-Cre | mca_classic | AAV | 588 |
| 160399309 | Gnrh1-Cre | mca_classic | AAV | 776 |
| 160537018 | Drd1a-Cre_EY2( | mca_classic | AAV | 573 |
| 160537796 | Drd1a-Cre_EY2( | mca_classic | AAV | 573 |

| | | | | |
|---|---|---|---|---|
| 160538548 | Slc6a3-Cre | mca_classic | AAV | 838 |
| 160539283 | Slc6a3-Cre | mca_classic | AAV | 823 |
| 160540013 | Drd2-Cre_ER44 | mca_classic | AAV | 573 |
| 160540751 | Slc6a3-Cre | mca_classic | AAV | 823 |
| 161176690 | Gabrr3-Cre_KC: | mca_classic | AAV | 573 |
| 161178152 | Syt17-Cre_NO1 | mca_classic | AAV | 386 |
| 161458737 | Rbp4-Cre_KL10 | mca_classic | AAV | 232 |
| 161460153 | Chat-IRES-Cre-n | mca_classic | AAV | 967 |
| 161460864 | Chat-IRES-Cre-n | mca_classic | AAV | 613 |
| 161463521 | Ntsr1-Cre_GN2: | mca_classic | AAV | 238 |
| 162018169 | Scnn1a-Tg2-Cre | mca_classic | AAV | 650 |
| 162018879 | Scnn1a-Tg2-Cre | mca_classic | AAV | 802 |
| 162019589 | Scnn1a-Tg2-Cre | mca_classic | AAV | 919 |
| 162020630 | Gad2-IRES-Cre | mca_classic | AAV | 838 |
| 164985329 | Gad2-IRES-Cre | mca_classic | AAV | 880 |
| 164986046 | Sim1-Cre_KJ18 | mca_classic | AAV | 792 |
| 165034344 | Sim1-Cre_KJ18 | mca_classic | AAV | 600 |
| 165035106 | Sim1-Cre_KJ18 | mca_classic | AAV | 794 |
| 165644051 | Cort-T2A-Cre | mca_classic | AAV | 226 |
| 165972951 | Erbb4-T2A-CreE | mca_classic | AAV | 566 |
| 165973664 | Chat-IRES-Cre-n | mca_classic | AAV | 1007 |
| 165974379 | Ucn3-Cre_KF43 | mca_classic | AAV | 797 |
| 165975096 | Erbb4-T2A-CreE | mca_classic | AAV | 823 |
| 165975810 | Cort-T2A-Cre | mca_classic | AAV | 136 |
| 166052101 | Chat-IRES-Cre-n | mca_classic | AAV | 919 |
| 166054222 | Pdzk1ip1-Cre_K | mca_classic | AAV | 823 |
| 166054929 | Rbp4-Cre_KL10 | mca_classic | AAV | 332 |
| 166082128 | Rbp4-Cre_KL10 | mca_classic | AAV | 24 |
| 166082842 | Rbp4-Cre_KL10 | mca_classic | AAV | 18 |
| 166083557 | Rbp4-Cre_KL10 | mca_classic | AAV | 213 |
| 166153483 | Rbp4-Cre_KL10 | mca_classic | AAV | 291 |
| 166154193 | Slc6a3-Cre | mca_classic | AAV | 380 |
| 166156241 | Slc6a3-Cre | mca_classic | AAV | 380 |
| 166264185 | Grm2-Cre_MR9 | mca_classic | AAV | 685 |
| 166267651 | Grm2-Cre_MR9 | mca_classic | AAV | 668 |
| 166268376 | Scnn1a-Tg3-Cre | mca_classic | AAV | 136 |
| 166269090 | Scnn1a-Tg3-Cre | mca_classic | AAV | 332 |
| 166269819 | Scnn1a-Tg3-Cre | mca_classic | AAV | 185 |
| 166271142 | A930038C07Rik | mca_classic | AAV | 332 |
| 166323186 | A930038C07Rik | mca_classic | AAV | 65 |
| 166323896 | A930038C07Rik | mca_classic | AAV | 226 |
| 166324604 | A930038C07Rik | mca_classic | AAV | 185 |
| 166326029 | Scnn1a-Tg3-Cre | mca_classic | AAV | 79 |
| 166326736 | Scnn1a-Tg3-Cre | mca_classic | AAV | 185 |
| 166458363 | Scnn1a-Tg3-Cre | mca_classic | AAV | 325 |
| 166459070 | Scnn1a-Tg3-Cre | mca_classic | AAV | 51 |
| 166459778 | Scnn1a-Tg3-Cre | mca_classic | AAV | 164 |

| | | | | |
|---|---|---|---|---|
| 166460484 | A930038C07Rik | mca_classic | AAV | 24 |
| 166461193 | A930038C07Rik | mca_classic | AAV | 18 |
| 166461899 | A930038C07Rik | mca_classic | AAV | 178 |
| 166532512 | Oxt-IRES-Cre | mca_classic | AAV | 782 |
| 166533924 | Ntsr1-Cre_GN2: | mca_classic | AAV | 51 |
| 166562678 | Pvalb-IRES-Cre | mca_classic | AAV | 185 |
| 166566678 | Nr5a1-Cre | mca_classic | AAV | 65 |
| 167025578 | Ucn3-Cre_KF43 | mca_classic | AAV | 694 |
| 167026321 | Erbb4-T2A-CreE | mca_classic | AAV | 621 |
| 167029528 | Cnnm2-Cre_KD | mca_classic | AAV | 398 |
| 167117360 | Agrp-IRES-Cre | mca_classic | AAV | 749 |
| 167118084 | Pdzk1ip1-Cre_K | mca_classic | AAV | 830 |
| 167200755 | Chat-IRES-Cre-n | mca_classic | AAV | 618 |
| 167201465 | Pvalb-IRES-Cre | mca_classic | AAV | 1071 |
| 167202174 | Pvalb-IRES-Cre | mca_classic | AAV | 1033 |
| 167211503 | Pvalb-IRES-Cre | mca_classic | AAV | 942 |
| 167212217 | Slc6a4-CreERT2 | mca_classic | AAV | 1013 |
| 167212932 | Otof-Cre | mca_classic | AAV | 494 |
| 167213641 | Crh-IRES-Cre_BI | mca_classic | AAV | 941 |
| 167373923 | Crh-IRES-Cre_BI | mca_classic | AAV | 693 |
| 167374643 | Cort-T2A-Cre | mca_classic | AAV | 51 |
| 167375352 | Cort-T2A-Cre | mca_classic | AAV | 185 |
| 167376068 | Cnnm2-Cre_KD | mca_classic | AAV | 913 |
| 167439900 | Grik4-Cre | mca_classic | AAV | 668 |
| 167440619 | Gad2-IRES-Cre | mca_classic | AAV | 24 |
| 167441329 | Gad2-IRES-Cre | mca_classic | AAV | 51 |
| 167442042 | Syt17-Cre_NO1 | mca_classic | AAV | 890 |
| 167511639 | Gad2-IRES-Cre | mca_classic | AAV | 185 |
| 167512351 | Gad2-IRES-Cre | mca_classic | AAV | 226 |
| 167513067 | Gad2-IRES-Cre | mca_classic | AAV | 457 |
| 167569313 | Etv1-CreERT2 | mca_classic | AAV | 24 |
| 167570021 | Etv1-CreERT2 | mca_classic | AAV | 51 |
| 167571459 | Grik4-Cre | mca_classic | AAV | 676 |
| 167654019 | Slc17a6-IRES-Cr | mca_classic | AAV | 527 |
| 167656152 | Lepr-IRES-Cre | mca_classic | AAV | 781 |
| 167665302 | Crh-IRES-Cre_BI | mca_classic | AAV | 977 |
| 167791990 | Nr5a1-Cre | mca_classic | AAV | 24 |
| 167793416 | Wfs1-Tg3-CreEI | mca_classic | AAV | 507 |
| 167794131 | Rbp4-Cre_KL10 | mca_classic | AAV | 164 |
| 167902586 | Rbp4-Cre_KL10 | mca_classic | AAV | 272 |
| 167904255 | Drd2-Cre_ER44 | mca_classic | AAV | 577 |
| 167904966 | Drd2-Cre_ER44 | mca_classic | AAV | 575 |
| 168002073 | Cux2-IRES-Cre | mca_classic | AAV | 24 |
| 168002780 | Gal-Cre_KI87 | mca_classic | AAV | 685 |
| 168003640 | Cux2-IRES-Cre | mca_classic | AAV | 51 |
| 168004394 | Gal-Cre_KI87 | mca_classic | AAV | 685 |
| 168005102 | Gal-Cre_KI87 | mca_classic | AAV | 749 |

| | | | | |
|---|---|---|---|---|
| 168094300 | Gnrh1-Cre | mca_classic | AAV | 588 |
| 168095041 | Gal-Cre_KI87 | mca_classic | AAV | 681 |
| 168095756 | Syt6-Cre_KI148 | mca_classic | AAV | 285 |
| 168096467 | Syt6-Cre_KI148 | mca_classic | AAV | 44 |
| 168097187 | Syt6-Cre_KI148 | mca_classic | AAV | 185 |
| 168162771 | Rbp4-Cre_KL10 | mca_classic | AAV | 24 |
| 168163498 | Rbp4-Cre_KL10 | mca_classic | AAV | 100 |
| 168164230 | Rbp4-Cre_KL10 | mca_classic | AAV | 298 |
| 168164972 | A930038C07Rik | mca_classic | AAV | 258 |
| 168165712 | A930038C07Rik | mca_classic | AAV | 346 |
| 168227692 | Slc6a4-CreERT2 | mca_classic | AAV | 1013 |
| 168229113 | Syt6-Cre_KI148 | mca_classic | AAV | 18 |
| 168229820 | Syt6-Cre_KI148 | mca_classic | AAV | 72 |
| 168230532 | Pcp2-Cre_GN13 | mca_classic | AAV | 1071 |
| 168299316 | Pcp2-Cre_GN13 | mca_classic | AAV | 1067 |
| 168300027 | Slc6a4-Cre_ET3 | mca_classic | AAV | 881 |
| 168300739 | Grik4-Cre | mca_classic | AAV | 685 |
| 168301446 | Grik4-Cre | mca_classic | AAV | 685 |
| 168302154 | Grik4-Cre | mca_classic | AAV | 431 |
| 168361750 | Gad2-IRES-Cre | mca_classic | AAV | 847 |
| 168362462 | Gad2-IRES-Cre | mca_classic | AAV | 749 |
| 168363168 | Syt6-Cre_KI148 | mca_classic | AAV | 332 |
| 168363874 | Cart-Tg1-Cre | mca_classic | AAV | 600 |
| 168364580 | Cart-Tg1-Cre | mca_classic | AAV | 764 |
| 168401109 | Pvalb-IRES-Cre | mca_classic | AAV | 232 |
| 168454779 | Calb2-IRES-Cre | mca_classic | AAV | 24 |
| 168455487 | Calb2-IRES-Cre | mca_classic | AAV | 136 |
| 168513750 | Calb2-IRES-Cre | mca_classic | AAV | 51 |
| 168515225 | Slc6a5-Cre_KF1 | mca_classic | AAV | 931 |
| 168515938 | Calb2-IRES-Cre | mca_classic | AAV | 185 |
| 168614604 | Drd1a-Cre_EY2 | mca_classic | AAV | 575 |
| 168616111 | Chat-IRES-Cre-n | mca_classic | AAV | 965 |
| 168616827 | Hdc-Cre_IM1 | mca_classic | AAV | 792 |
| 168663472 | Slc17a6-IRES-Cr | mca_classic | AAV | 802 |
| 168664192 | Gabrr3-Cre_KC | mca_classic | AAV | 1098 |
| 170261951 | Ntsr1-Cre_GN2 | mca_classic | AAV | 213 |
| 170263370 | Gabrr3-Cre_KC | mca_classic | AAV | 802 |
| 170784358 | Crh-IRES-Cre_B | mca_classic | AAV | 694 |
| 170785775 | Drd1a-Cre_EY2 | mca_classic | AAV | 575 |
| 170858675 | Gabrr3-Cre_KC | mca_classic | AAV | 802 |
| 170859382 | Gabrr3-Cre_KC | mca_classic | AAV | 645 |
| 170860092 | Sim1-Cre_KJ18 | mca_classic | AAV | 794 |
| 170860801 | Sim1-Cre_KJ18 | mca_classic | AAV | 600 |
| 170946889 | Sim1-Cre_KJ18 | mca_classic | AAV | 600 |
| 170947610 | Pdzk1ip1-Cre_K | mca_classic | AAV | 844 |
| 170949055 | Syt17-Cre_NO1 | mca_classic | AAV | 390 |
| 171019004 | Etv1-CreERT2 | mca_classic | AAV | 457 |

| | | | | | |
|---|---|---|---|---|---|
| | 171019710 | Slc32a1-IRES-Cr | mca_classic | AAV | 870 |
| | 171020416 | Pmch-Cre | mca_classic | AAV | 794 |
| | 171021122 | Slc6a3-Cre | mca_classic | AAV | 973 |
| | 171021829 | Cck-IRES-Cre | mca_classic | AAV | 823 |
| | 171064488 | Slc32a1-IRES-Cr | mca_classic | AAV | 703 |
| | 171065200 | Slc32a1-IRES-Cr | mca_classic | AAV | 703 |
| | 171065906 | Slc32a1-IRES-Cr | mca_classic | AAV | 802 |
| | 171066613 | Slc32a1-IRES-Cr | mca_classic | AAV | 802 |
| | 171067319 | Crh-IRES-Cre_ZJ | mca_classic | AAV | 693 |
| | 171274191 | Prkcd-GluCla-CF | mca_classic | AAV | 467 |
| | 171275617 | A930038C07Rik | mca_classic | AAV | 226 |
| | 171276330 | A930038C07Rik | mca_classic | AAV | 44 |
| | 171409520 | Scnn1a-Tg2-Cre | mca_classic | AAV | 992 |
| | 171482142 | Lepr-IRES-Cre | mca_classic | AAV | 733 |
| | 171485060 | Cart-Tg1-Cre | mca_classic | AAV | 416 |
| | 173206592 | C57Bl/6J | mca_classic | AAV | 685 |
| Y | 174361746 | C57BL/6J | mca_classic | AAV | 114 |
| | 174583187 | C57Bl/6J | mca_classic | AAV | 826 |
| | 174583904 | C57Bl/6J | mca_classic | AAV | 826 |
| | 174736554 | C57Bl/6J | mca_classic | AAV | 645 |
| | 174781014 | C57Bl/6J | mca_classic | AAV | 669 |
| | 174788109 | C57Bl/6J | mca_classic | AAV | 802 |
| | 174957972 | C57Bl/6J | mca_classic | AAV | 694 |
| | 175018829 | C57Bl/6J | mca_classic | AAV | 802 |
| | 175019536 | C57Bl/6J | mca_classic | AAV | 694 |
| | 175072215 | C57Bl/6J | mca_classic | AAV | 573 |
| | 175106053 | C57Bl/6J | mca_classic | AAV | 757 |
| | 175106769 | C57Bl/6J | mca_classic | AAV | 787 |
| | 175142304 | C57Bl/6J | mca_classic | AAV | 992 |
| | 175158132 | C57Bl/6J | mca_classic | AAV | 830 |
| | 175158844 | C57Bl/6J | mca_classic | AAV | 870 |
| | 175263063 | C57Bl/6J | mca_classic | AAV | 822 |
| | 175263771 | C57Bl/6J | mca_classic | AAV | 744 |
| | 175264587 | C57Bl/6J | mca_classic | AAV | 913 |
| | 175265301 | C57Bl/6J | mca_classic | AAV | 978 |
| | 175372863 | C57Bl/6J | mca_classic | AAV | 593 |
| | 175373569 | C57Bl/6J | mca_classic | AAV | 575 |
| | 175374275 | C57Bl/6J | mca_classic | AAV | 792 |
| | 175374982 | C57Bl/6J | mca_classic | AAV | 694 |
| | 175718634 | Etv1-CreERT2 | mca_classic | AAV | 457 |
| | 175719341 | Calb2-IRES-Cre | mca_classic | AAV | 226 |
| | 175728165 | Cdhr1-Cre_KG6 | mca_classic | AAV | 380 |
| | 175732001 | Drd2-Cre_ER44 | mca_classic | AAV | 575 |
| | 175732996 | Drd2-Cre_ER44 | mca_classic | AAV | 573 |
| | 175736945 | Hdc-Cre_IM1 | mca_classic | AAV | 823 |
| | 175738378 | Pomc-Cre_BL | mca_classic | AAV | 733 |
| | 175739085 | Slc6a3-Cre | mca_classic | AAV | 621 |

| | | | | |
|---|---|---|---|---|
| 175739791 | Grm2-Cre_MR9 | mca_classic | AAV | 685 |
| 175740500 | Grm2-Cre_MR9 | mca_classic | AAV | 685 |
| 175816975 | Gal-Cre_KI87 | mca_classic | AAV | 948 |
| 175817683 | Gal-Cre_KI87 | mca_classic | AAV | 682 |
| 175818392 | Gal-Cre_KI87 | mca_classic | AAV | 675 |
| 175819113 | Gpr26-Cre_KO2 | mca_classic | AAV | 51 |
| 176430283 | Cux2-IRES-Cre | mca_classic | AAV | 122 |
| 176431817 | Gal-Cre_KI87 | mca_classic | AAV | 733 |
| 176432524 | Gal-Cre_KI87 | mca_classic | AAV | 720 |
| 176433237 | A930038C07Rik | mca_classic | AAV | 24 |
| 176496282 | Pvalb-IRES-Cre | mca_classic | AAV | 93 |
| 176497015 | Syt6-Cre_KI148 | mca_classic | AAV | 332 |
| 176881134 | Nr5a1-Cre | mca_classic | AAV | 51 |
| 176882966 | Nr5a1-Cre | mca_classic | AAV | 185 |
| 176886958 | Esr1-2A-Cre | mca_classic | AAV | 787 |
| 176887774 | Esr1-2A-Cre | mca_classic | AAV | 794 |
| 176888661 | Esr1-2A-Cre | mca_classic | AAV | 801 |
| 176897793 | Cart-Tg1-Cre | mca_classic | AAV | 693 |
| 176898557 | Cart-Tg1-Cre | mca_classic | AAV | 792 |
| 176899332 | Kcnc2-Cre | mca_classic | AAV | 410 |
| 176900059 | Pdzk1ip1-Cre_K | mca_classic | AAV | 808 |
| 176901480 | Nr5a1-Cre | mca_classic | AAV | 185 |
| 177319236 | Syt6-Cre_KI148 | mca_classic | AAV | 65 |
| 177322838 | Nr5a1-Cre | mca_classic | AAV | 44 |
| 177460028 | Chat-IRES-Cre-n | mca_classic | AAV | 854 |
| 177605425 | Chat-IRES-Cre-n | mca_classic | AAV | 618 |
| 177606140 | Chat-IRES-Cre-n | mca_classic | AAV | 1007 |
| 177606858 | Grik4-Cre | mca_classic | AAV | 467 |
| 177780284 | Grik4-Cre | mca_classic | AAV | 467 |
| 177781006 | Nr5a1-Cre | mca_classic | AAV | 185 |
| 177781745 | Nr5a1-Cre | mca_classic | AAV | 185 |
| 177782493 | Nr5a1-Cre | mca_classic | AAV | 72 |
| 177783204 | Syt6-Cre_KI148 | mca_classic | AAV | 226 |
| 177783918 | Syt6-Cre_KI148 | mca_classic | AAV | 100 |
| 177889243 | Syt6-Cre_KI148 | mca_classic | AAV | 245 |
| 177890246 | Syt6-Cre_KI148 | mca_classic | AAV | 6 |
| 177890956 | Sim1-Cre_KJ18 | mca_classic | AAV | 801 |
| 177892379 | Nr5a1-Cre | mca_classic | AAV | 72 |
| 177893658 | Nr5a1-Cre | mca_classic | AAV | 18 |
| 177903648 | Pnmt-Cre | mca_classic | AAV | 941 |
| 177904363 | Scnn1a-Tg2-Cre | mca_classic | AAV | 1007 |
| 177905562 | Scnn1a-Tg2-Cre | mca_classic | AAV | 967 |
| 177907797 | A930038C07Rik | mca_classic | AAV | 51 |
| 178282527 | Gal-Cre_KI87 | mca_classic | AAV | 733 |
| 178283239 | Gal-Cre_KI87 | mca_classic | AAV | 739 |
| 178283952 | Gal-Cre_KI87 | mca_classic | AAV | 992 |
| 178284661 | Cux2-IRES-Cre | mca_classic | AAV | 65 |

| | | | | | |
|---|---|---|---|---|---|
| | 178382220 | Grik4-Cre | mca_classic | AAV | 467 |
| | 178486024 | Cart-Tg1-Cre | mca_classic | AAV | 588 |
| | 178487444 | Syt6-Cre_KI148 | mca_classic | AAV | 51 |
| | 178488152 | Oxt-IRES-Cre | mca_classic | AAV | 717 |
| | 178488859 | Satb2-Cre_MO2 | mca_classic | AAV | 507 |
| | 178489574 | Prkcd-GluCla-CF | mca_classic | AAV | 658 |
| | 179640955 | Scnn1a-Tg3-Cre | mca_classic | AAV | 18 |
| | 179641666 | Syt6-Cre_KI148 | mca_classic | AAV | 18 |
| | 179642401 | Syt6-Cre_KI148 | mca_classic | AAV | 380 |
| | 179643824 | Grik4-Cre | mca_classic | AAV | 467 |
| | 179644545 | Grp-Cre_KH288 | mca_classic | AAV | 467 |
| | 179902073 | Grp-Cre_KH288 | mca_classic | AAV | 467 |
| | 179902786 | Ucn3-Cre_KF43 | mca_classic | AAV | 684 |
| | 179904203 | Lepr-IRES-Cre | mca_classic | AAV | 781 |
| | 179904912 | Vip-IRES-Cre | mca_classic | AAV | 24 |
| | 180075597 | Vip-IRES-Cre | mca_classic | AAV | 44 |
| | 180297139 | C57Bl/6J | mca_classic | AAV | 494 |
| | 180405830 | Vipr2-Cre_KE2 | mca_classic | AAV | 457 |
| | 180520257 | C57Bl/6J | mca_classic | AAV | 658 |
| | 180520968 | Vipr2-Cre_KE2 | mca_classic | AAV | 457 |
| | 180522266 | Erbb4-T2A-CreE | mca_classic | AAV | 826 |
| | 180523704 | C57Bl/6J | mca_classic | AAV | 467 |
| | 180524412 | Tac1-IRES2-Cre | mca_classic | AAV | 838 |
| | 180525136 | Tac1-IRES2-Cre | mca_classic | AAV | 948 |
| | 180568155 | C57Bl/6J | mca_classic | AAV | 693 |
| | 180628971 | C57Bl/6J | mca_classic | AAV | 649 |
| | 180674463 | C57Bl/6J | mca_classic | AAV | 757 |
| | 180707817 | C57Bl/6J | mca_classic | AAV | 645 |
| | 180708524 | C57Bl/6J | mca_classic | AAV | 669 |
| Y | 180981417 | C57Bl/6J | mca_classic | AAV | 600 |
| | 181057754 | C57Bl/6J | mca_classic | AAV | 757 |
| | 181058463 | C57Bl/6J | mca_classic | AAV | 992 |
| | 181116850 | Gabra6-IRES-Cr | mca_classic | AAV | 1071 |
| | 181128406 | Gabra6-IRES-Cr | mca_classic | AAV | 1033 |
| | 181180790 | Ntsr1-Cre_GN2 | mca_classic | AAV | 93 |
| | 181258571 | Ntsr1-Cre_GN2 | mca_classic | AAV | 185 |
| | 181259279 | Tac1-IRES2-Cre | mca_classic | AAV | 24 |
| | 181371331 | Tac1-IRES2-Cre | mca_classic | AAV | 44 |
| | 181372049 | Tac1-IRES2-Cre | mca_classic | AAV | 100 |
| | 181598954 | Gpr26-Cre_KO2 | mca_classic | AAV | 24 |
| | 181599674 | Gpr26-Cre_KO2 | mca_classic | AAV | 44 |
| | 181600380 | Gpr26-Cre_KO2 | mca_classic | AAV | 136 |
| | 181601101 | Gpr26-Cre_KO2 | mca_classic | AAV | 185 |
| | 181777177 | Sim1-Cre_KJ18 | mca_classic | AAV | 457 |
| | 181786681 | Sim1-Cre_KJ18 | mca_classic | AAV | 621 |
| | 181818339 | Scnn1a-Tg3-Cre | mca_classic | AAV | 44 |
| | 181819064 | Scnn1a-Tg3-Cre | mca_classic | AAV | 72 |

| | | | | |
|---|---|---|---|---|
| 181858761 | Etv1-CreERT2 | mca_classic | AAV | 226 |
| 181859467 | Cux2-CreERT2 | mca_classic | AAV | 507 |
| 181860173 | Scnn1a-Tg3-Cre | mca_classic | AAV | 79 |
| 181860879 | Scnn1a-Tg3-Cre | mca_classic | AAV | 332 |
| 181889764 | Etv1-CreERT2 | mca_classic | AAV | 595 |
| 181890477 | Etv1-CreERT2 | mca_classic | AAV | 595 |
| 181891892 | Lepr-IRES-Cre | mca_classic | AAV | 733 |
| 181895006 | Slc17a6-IRES-Cr | mca_classic | AAV | 685 |
| 182029174 | Grik4-Cre | mca_classic | AAV | 473 |
| 182029881 | Pdzk1ip1-Cre_K | mca_classic | AAV | 849 |
| 182030590 | Pdzk1ip1-Cre_K | mca_classic | AAV | 975 |
| 182031296 | Pdzk1ip1-Cre_K | mca_classic | AAV | 847 |
| 182040934 | Vip-IRES-Cre | mca_classic | AAV | 136 |
| 182041643 | Cart-Tg1-Cre | mca_classic | AAV | 600 |
| 182042349 | Slc6a3-Cre | mca_classic | AAV | 380 |
| 182043055 | Nos1-CreERT2 | mca_classic | AAV | 51 |
| 182089608 | Scnn1a-Tg3-Cre | mca_classic | AAV | 185 |
| 182090318 | Rbp4-Cre_KL10 | mca_classic | AAV | 136 |
| 182144176 | Syt6-Cre_KI148 | mca_classic | AAV | 838 |
| 182182936 | Grm2-Cre_MR9 | mca_classic | AAV | 764 |
| 182183683 | Erbb4-T2A-CreE | mca_classic | AAV | 226 |
| 182184486 | Erbb4-T2A-CreE | mca_classic | AAV | 185 |
| 182185289 | Grm2-Cre_MR9 | mca_classic | AAV | 684 |
| 182224715 | Crh-IRES-Cre_ZJ | mca_classic | AAV | 911 |
| 182226133 | Otof-Cre | mca_classic | AAV | 494 |
| 182226839 | Vipr2-Cre_KE2 | mca_classic | AAV | 457 |
| 182280207 | Vipr2-Cre_KE2 | mca_classic | AAV | 838 |
| 182280916 | Slc6a5-Cre_KF1 | mca_classic | AAV | 679 |
| 182293273 | Vip-IRES-Cre | mca_classic | AAV | 185 |
| 182294687 | Rbp4-Cre_KL10 | mca_classic | AAV | 431 |
| 182295393 | Rbp4-Cre_KL10 | mca_classic | AAV | 473 |
| 182336846 | Nr5a1-Cre | mca_classic | AAV | 739 |
| 182337561 | Nr5a1-Cre | mca_classic | AAV | 787 |
| 182338356 | Cux2-IRES-Cre | mca_classic | AAV | 332 |
| 182341627 | Syt6-Cre_KI148 | mca_classic | AAV | 619 |
| 182458849 | Gal-Cre_KI87 | mca_classic | AAV | 992 |
| 182459635 | Gal-Cre_KI87 | mca_classic | AAV | 744 |
| 182460343 | Gal-Cre_KI87 | mca_classic | AAV | 776 |
| 182465608 | Syt6-Cre_KI148 | mca_classic | AAV | 889 |
| 182467026 | A930038C07Rik | mca_classic | AAV | 51 |
| 182467736 | Cux2-IRES-Cre | mca_classic | AAV | 325 |
| 182515576 | Gal-Cre_KI87 | mca_classic | AAV | 669 |
| 182613651 | Pvalb-IRES-Cre | mca_classic | AAV | 830 |
| 182615063 | Grp-Cre_KH288 | mca_classic | AAV | 467 |
| 182615771 | Grp-Cre_KH288 | mca_classic | AAV | 467 |
| 182616478 | Cux2-IRES-Cre | mca_classic | AAV | 18 |
| 182617186 | Wfs1-Tg3-CreEI | mca_classic | AAV | 523 |

| | | | | |
|---|---|---|---|---|
| 182685522 | Grik4-Cre | mca_classic | AAV | 1088 |
| 182686228 | Cdhr1-Cre_KG6 | mca_classic | AAV | 386 |
| 182686935 | Trib2-F2A-CreEI | mca_classic | AAV | 136 |
| 182793477 | Rbp4-Cre_KL10 | mca_classic | AAV | 24 |
| 182794184 | Rbp4-Cre_KL10 | mca_classic | AAV | 494 |
| 182795499 | Grm2-Cre_MR9 | mca_classic | AAV | 24 |
| 182803137 | Rbp4-Cre_KL10 | mca_classic | AAV | 65 |
| 182804552 | Pvalb-IRES-Cre | mca_classic | AAV | 919 |
| 182805258 | Vipr2-Cre_KE2 | mca_classic | AAV | 671 |
| 182805965 | Crh-IRES-Cre_BI | mca_classic | AAV | 675 |
| 182842391 | Sim1-Cre_KJ18 | mca_classic | AAV | 757 |
| 182843806 | Kcnc2-Cre | mca_classic | AAV | 457 |
| 182886547 | Sst-IRES-Cre | mca_classic | AAV | 136 |
| 182887258 | Calb2-IRES-Cre | mca_classic | AAV | 589 |
| 182888003 | Calb2-IRES-Cre | mca_classic | AAV | 698 |
| 182890735 | Calb2-IRES-Cre | mca_classic | AAV | 992 |
| 182892855 | Slc18a2-Cre_OZ | mca_classic | AAV | 669 |
| 182896517 | Etv1-CreERT2 | mca_classic | AAV | 199 |
| 182933935 | Kcnc2-Cre | mca_classic | AAV | 467 |
| 182934777 | Gad2-IRES-Cre | mca_classic | AAV | 136 |
| 182935487 | Vip-IRES-Cre | mca_classic | AAV | 24 |
| 183009881 | Chat-IRES-Cre-n | mca_classic | AAV | 573 |
| 183011353 | Grik4-Cre | mca_classic | AAV | 669 |
| 183057424 | Grp-Cre_KH288 | mca_classic | AAV | 698 |
| 183058837 | Grp-Cre_KH288 | mca_classic | AAV | 685 |
| 183071513 | Grp-Cre_KH288 | mca_classic | AAV | 699 |
| 183103805 | Crh-IRES-Cre_BI | mca_classic | AAV | 18 |
| 183104511 | Crh-IRES-Cre_BI | mca_classic | AAV | 226 |
| 183105218 | Crh-IRES-Cre_BI | mca_classic | AAV | 136 |
| 183171679 | Rbp4-Cre_KL10 | mca_classic | AAV | 51 |
| 183172820 | Crh-IRES-Cre_BI | mca_classic | AAV | 24 |
| 183173527 | Rbp4-Cre_KL10 | mca_classic | AAV | 185 |
| 183174303 | Rbp4-Cre_KL10 | mca_classic | AAV | 668 |
| 183175010 | Grik4-Cre | mca_classic | AAV | 668 |
| 183225124 | Grm2-Cre_MR9 | mca_classic | AAV | 494 |
| 183225830 | Grm2-Cre_MR9 | mca_classic | AAV | 692 |
| 183282261 | Calb2-IRES-Cre | mca_classic | AAV | 668 |
| 183282970 | Calb2-IRES-Cre | mca_classic | AAV | 668 |
| 183284388 | Calb2-IRES-Cre | mca_classic | AAV | 890 |
| 183329222 | Calb2-IRES-Cre | mca_classic | AAV | 685 |
| 183329991 | Gal-Cre_KI87 | mca_classic | AAV | 658 |
| 183330908 | Rbp4-Cre_KL10 | mca_classic | AAV | 285 |
| 183374804 | Scnn1a-Tg3-Cre | mca_classic | AAV | 51 |
| 183375545 | Pdzk1ip1-Cre_K | mca_classic | AAV | 812 |
| 183376269 | Pdzk1ip1-Cre_K | mca_classic | AAV | 830 |
| 183376982 | Rbp4-Cre_KL10 | mca_classic | AAV | 473 |
| 183459175 | Avp-IRES2-Cre | mca_classic | AAV | 720 |

| | | | | |
|---|---|---|---|---|
| 183461297 | Cux2-IRES-Cre | mca_classic | AAV | 232 |
| 183470468 | Cux2-IRES-Cre | mca_classic | AAV | 232 |
| 183471174 | Cux2-IRES-Cre | mca_classic | AAV | 258 |
| 183471884 | Cux2-IRES-Cre | mca_classic | AAV | 185 |
| 183472596 | Grp-Cre_KH288 | mca_classic | AAV | 226 |
| 183562831 | Cck-IRES-Cre | mca_classic | AAV | 881 |
| 183617432 | Htr2a-Cre_KM2 | mca_classic | AAV | 24 |
| 183618845 | Htr2a-Cre_KM2 | mca_classic | AAV | 272 |
| 183901489 | Chat-IRES-Cre-n | mca_classic | AAV | 890 |
| 184073678 | Scnn1a-Tg3-Cre | mca_classic | AAV | 353 |
| 184074388 | Scnn1a-Tg3-Cre | mca_classic | AAV | 185 |
| 184075100 | Scnn1a-Tg3-Cre | mca_classic | AAV | 213 |
| 184075807 | Scnn1a-Tg3-Cre | mca_classic | AAV | 523 |
| 184157585 | Calb2-IRES-Cre | mca_classic | AAV | 692 |
| 184158290 | Calb2-IRES-Cre | mca_classic | AAV | 694 |
| 184158996 | Calb2-IRES-Cre | mca_classic | AAV | 692 |
| 184159706 | Cux2-IRES-Cre | mca_classic | AAV | 136 |
| 184167484 | Cux2-IRES-Cre | mca_classic | AAV | 171 |
| 184168193 | Cux2-IRES-Cre | mca_classic | AAV | 332 |
| 184168899 | Grp-Cre_KH288 | mca_classic | AAV | 279 |
| 184212995 | Ntrk1-IRES-Cre | mca_classic | AAV | 870 |
| 184213701 | Grp-Cre_KH288 | mca_classic | AAV | 232 |
| 184259031 | Syt17-Cre_NO1 | mca_classic | AAV | 416 |
| 187269162 | Adcyap1-2A-Cre | mca_classic | AAV | 421 |
| 194947823 | Cux2-CreERT2 | mca_classic | AAV | 507 |
| 194948535 | Erbb4-T2A-CreE | mca_classic | AAV | 621 |
| 204832205 | Adcyap1-2A-Cre | mca_classic | AAV | 694 |
| 204832917 | Crh-IRES-Cre_ZJ | mca_classic | AAV | 621 |
| 204907355 | Crh-IRES-Cre_ZJ | mca_classic | AAV | 595 |
| 204908781 | Gal-Cre_KI87 | mca_classic | AAV | 692 |
| 232310521 | Kiss1-Cre | mca_classic | AAV | 733 |
| 232311236 | Kiss1-Cre | mca_classic | AAV | 733 |
| 232311959 | Rbp4-Cre_KL10 | mca_classic | AAV | 494 |
| 241278553 | Tac2-IRES2-Cre | mca_classic | AAV | 733 |
| 241279261 | Tac2-IRES2-Cre | mca_classic | AAV | 595 |
| 241279971 | Sim1-Cre_KJ18 | mca_classic | AAV | 621 |
| 241280698 | Sim1-Cre_KJ18 | mca_classic | AAV | 621 |
| 249327301 | Rbp4-Cre_KL10 | mca_classic | AAV | 44 |
| 249396394 | Rbp4-Cre_KL10 | mca_classic | AAV | 332 |
| 249402048 | Rbp4-Cre_KL10 | mca_classic | AAV | 107 |
| 249402769 | Syt17-Cre_NO1 | mca_classic | AAV | 467 |
| 257636467 | Scnn1a-Tg3-Cre | mca_classic | AAV | 100 |
| 257666283 | Scnn1a-Tg3-Cre | mca_classic | AAV | 72 |
| 257667830 | Scnn1a-Tg3-Cre | mca_classic | AAV | 199 |
| 258315443 | Slc6a5-Cre_KF1 | mca_classic | AAV | 885 |
| 258316155 | Gpr26-Cre_KO2 | mca_classic | AAV | 226 |
| 258316862 | Pvalb-IRES-Cre | mca_classic | AAV | 913 |

| | | | | |
|---|---|---|---|---|
| 258914806 | Pvalb-IRES-Cre | mca_classic | AAV | 826 |
| 258915564 | Htr2a-Cre_KM2 | mca_classic | AAV | 24 |
| 258916270 | Htr2a-Cre_KM2 | mca_classic | AAV | 264 |
| 262188772 | Vip-IRES-Cre | mca_classic | AAV | 838 |
| 262215150 | Ntrk1-IRES-Cre | mca_classic | AAV | 911 |
| 262536037 | Htr2a-Cre_KM2 | mca_classic | AAV | 279 |
| 263106036 | Rbp4-Cre_KL10 | mca_classic | AAV | 238 |
| 263106751 | Ntrk1-IRES-Cre | mca_classic | AAV | 692 |
| 263241470 | Scnn1a-Tg2-Cre | mca_classic | AAV | 662 |
| 263242463 | Rbp4-Cre_KL10 | mca_classic | AAV | 24 |
| 263369222 | Pomc-Cre_BL | mca_classic | AAV | 733 |
| 263370720 | Slc18a2-Cre_OZ | mca_classic | AAV | 51 |
| 263553163 | Slc18a2-Cre_OZ | mca_classic | AAV | 86 |
| 263553934 | Scnn1a-Tg2-Cre | mca_classic | AAV | 981 |
| 263780018 | Slc6a3-Cre | mca_classic | AAV | 781 |
| 263780729 | Cux2-IRES-Cre | mca_classic | AAV | 185 |
| 263781454 | Cux2-IRES-Cre | mca_classic | AAV | 24 |
| 263784128 | Syt17-Cre_NO1 | mca_classic | AAV | 973 |
| 263785543 | Slc17a6-IRES-Cr | mca_classic | AAV | 701 |
| 263974698 | Cux2-IRES-Cre | mca_classic | AAV | 494 |
| 263976175 | Gabrr3-Cre_KC | mca_classic | AAV | 925 |
| 264076081 | Slc32a1-IRES-Cr | mca_classic | AAV | 844 |
| 264076854 | Cux2-IRES-Cre | mca_classic | AAV | 279 |
| 264077561 | Slc32a1-IRES-Cr | mca_classic | AAV | 535 |
| 264078267 | Slc32a1-IRES-Cr | mca_classic | AAV | 749 |
| 264095536 | Slc32a1-IRES-Cr | mca_classic | AAV | 573 |
| 264096244 | Syt6-Cre_KI148 | mca_classic | AAV | 954 |
| 264096952 | Ntsr1-Cre_GN2 | mca_classic | AAV | 1098 |
| 264097661 | Pdzk1ip1-Cre_K | mca_classic | AAV | 844 |
| 264248605 | Esr1-2A-Cre | mca_classic | AAV | 801 |
| 264249312 | Esr1-2A-Cre | mca_classic | AAV | 298 |
| 264319363 | Esr1-2A-Cre | mca_classic | AAV | 787 |
| 264320076 | Esr1-2A-Cre | mca_classic | AAV | 801 |
| 264320859 | Syt6-Cre_KI148 | mca_classic | AAV | 279 |
| 264321572 | Chat-IRES-Cre-n | mca_classic | AAV | 978 |
| 264564375 | Chat-IRES-Cre-n | mca_classic | AAV | 1007 |
| 264565965 | Chat-IRES-Cre-n | mca_classic | AAV | 890 |
| 264566672 | Chat-IRES-Cre-n | mca_classic | AAV | 867 |
| 264629246 | A930038C07Rik | mca_classic | AAV | 65 |
| 264630019 | A930038C07Rik | mca_classic | AAV | 206 |
| 264631432 | A930038C07Rik | mca_classic | AAV | 380 |
| 264696942 | A930038C07Rik | mca_classic | AAV | 797 |
| 264697714 | A930038C07Rik | mca_classic | AAV | 802 |
| 264707643 | Crh-IRES-Cre_Bl | mca_classic | AAV | 685 |
| 264708349 | Grm2-Cre_MR9 | mca_classic | AAV | 258 |
| 264709761 | Grm2-Cre_MR9 | mca_classic | AAV | 473 |
| 264872385 | Grm2-Cre_MR9 | mca_classic | AAV | 473 |

| | | | | |
|---|---|---|---|---|
| 264873092 | Cux2-CreERT2 | mca_classic | AAV | 507 |
| 264873809 | Cux2-CreERT2 | mca_classic | AAV | 507 |
| 264874516 | Calb2-IRES-Cre | mca_classic | AAV | 390 |
| 264945172 | Grik4-Cre | mca_classic | AAV | 535 |
| 265125894 | Oxt-IRES-Cre | mca_classic | AAV | 794 |
| 265135682 | Ntrk1-IRES-Cre | mca_classic | AAV | 942 |
| 265136608 | Ntrk1-IRES-Cre | mca_classic | AAV | 575 |
| 265138021 | Tac2-IRES2-Cre | mca_classic | AAV | 621 |
| 265286700 | Cart-Tg1-Cre | mca_classic | AAV | 694 |
| 265287564 | Tac2-IRES2-Cre | mca_classic | AAV | 712 |
| 265288825 | Grp-Cre_KH288 | mca_classic | AAV | 232 |
| 265289624 | Grp-Cre_KH288 | mca_classic | AAV | 24 |
| 265291552 | Grp-Cre_KH288 | mca_classic | AAV | 232 |
| 265292679 | Grp-Cre_KH288 | mca_classic | AAV | 24 |
| 265293391 | Grp-Cre_KH288 | mca_classic | AAV | 844 |
| 265487421 | Grp-Cre_KH288 | mca_classic | AAV | 890 |
| 265633461 | Slc18a2-Cre_OZ | mca_classic | AAV | 985 |
| 265648940 | Slc18a2-Cre_OZ | mca_classic | AAV | 595 |
| 265712971 | Gpr26-Cre_KO2 | mca_classic | AAV | 232 |
| 265713683 | Gpr26-Cre_KO2 | mca_classic | AAV | 258 |
| 265813096 | Ins2-Cre_25 | mca_classic | AAV | 739 |
| 265820216 | Syt6-Cre_KI148 | mca_classic | AAV | 264 |
| 265820951 | Slc18a2-Cre_OZ | mca_classic | AAV | 51 |
| 265928489 | Gpr26-Cre_KO2 | mca_classic | AAV | 1098 |
| 265929196 | Nr5a1-Cre | mca_classic | AAV | 24 |
| 265929968 | Ntrk1-IRES-Cre | mca_classic | AAV | 573 |
| 265930674 | Ntrk1-IRES-Cre | mca_classic | AAV | 913 |
| 265943460 | Ntrk1-IRES-Cre | mca_classic | AAV | 610 |
| 265944167 | Ntrk1-IRES-Cre | mca_classic | AAV | 610 |
| 265944939 | Ntrk1-IRES-Cre | mca_classic | AAV | 617 |
| 265945645 | Prkcd-GluCla-CF | mca_classic | AAV | 595 |
| 265946352 | Prkcd-GluCla-CF | mca_classic | AAV | 588 |
| 265947058 | Prkcd-GluCla-CF | mca_classic | AAV | 694 |
| 266099165 | Pvalb-IRES-Cre | mca_classic | AAV | 838 |
| 266100645 | Nr5a1-Cre | mca_classic | AAV | 185 |
| 266172624 | Trib2-F2A-CreEI | mca_classic | AAV | 51 |
| 266173339 | Gal-Cre_KI87 | mca_classic | AAV | 473 |
| 266174045 | Gal-Cre_KI87 | mca_classic | AAV | 676 |
| 266174751 | Cart-Tg1-Cre | mca_classic | AAV | 739 |
| 266175461 | Nr5a1-Cre | mca_classic | AAV | 100 |
| 266176167 | Trib2-F2A-CreEI | mca_classic | AAV | 24 |
| 266177248 | Nr5a1-Cre | mca_classic | AAV | 185 |
| 266248065 | Gal-Cre_KI87 | mca_classic | AAV | 792 |
| 266248776 | Gal-Cre_KI87 | mca_classic | AAV | 662 |
| 266249483 | Rbp4-Cre_KL10 | mca_classic | AAV | 79 |
| 266250195 | Rbp4-Cre_KL10 | mca_classic | AAV | 185 |
| 266409949 | Rbp4-Cre_KL10 | mca_classic | AAV | 473 |

| | | | | |
|---|---|---|---|---|
| 266486371 | Cux2-IRES-Cre | mca_classic | AAV | 51 |
| 266487079 | Cux2-IRES-Cre | mca_classic | AAV | 178 |
| 266488504 | Gad2-IRES-Cre | mca_classic | AAV | 467 |
| 266489212 | Slc32a1-IRES-Cr | mca_classic | AAV | 787 |
| 266490034 | Gad2-IRES-Cre | mca_classic | AAV | 757 |
| 266500714 | Th-Cre_FI172 | mca_classic | AAV | 838 |
| 266501422 | Th-Cre_FI172 | mca_classic | AAV | 826 |
| 266563321 | Gad2-IRES-Cre | mca_classic | AAV | 386 |
| 266564027 | Grp-Cre_KH288 | mca_classic | AAV | 272 |
| 266582782 | Grp-Cre_KH288 | mca_classic | AAV | 238 |
| 266585624 | C57Bl/6J | mca_classic | AAV | 668 |
| 266643194 | Slc18a2-Cre_OZ | mca_classic | AAV | 65 |
| 266644610 | Cux2-IRES-Cre | mca_classic | AAV | 65 |
| 266646036 | Gad2-IRES-Cre | mca_classic | AAV | 844 |
| 266693274 | C57Bl/6J | mca_classic | AAV | 701 |
| 266816894 | Htr2a-Cre_KM2 | mca_classic | AAV | 238 |
| 266836749 | Crh-IRES-Cre_Bl | mca_classic | AAV | 185 |
| 266837456 | Sim1-Cre_KJ18 | mca_classic | AAV | 792 |
| 266839077 | Crh-IRES-Cre_Bl | mca_classic | AAV | 693 |
| 266839784 | Crh-IRES-Cre_Bl | mca_classic | AAV | 941 |
| 266840498 | Crh-IRES-Cre_Bl | mca_classic | AAV | 720 |
| 266913240 | Crh-IRES-Cre_Bl | mca_classic | AAV | 44 |
| 266962653 | Slc17a6-IRES-Cr | mca_classic | AAV | 431 |
| 266963362 | Ntsr1-Cre_GN2 | mca_classic | AAV | 136 |
| 266964075 | Ntsr1-Cre_GN2 | mca_classic | AAV | 226 |
| 266964788 | Ntsr1-Cre_GN2 | mca_classic | AAV | 65 |
| 266966337 | Pdzk1ip1-Cre_K | mca_classic | AAV | 812 |
| 267029447 | Pdzk1ip1-Cre_K | mca_classic | AAV | 838 |
| 267030155 | Pvalb-IRES-Cre | mca_classic | AAV | 838 |
| 267032286 | Syt6-Cre_KI148 | mca_classic | AAV | 298 |
| 267101850 | Scnn1a-Tg3-Cre | mca_classic | AAV | 226 |
| 267103498 | Slc17a7-IRES2-C | mca_classic | AAV | 390 |
| 267104205 | Slc17a7-IRES2-C | mca_classic | AAV | 457 |
| 267106046 | Slc17a7-IRES2-C | mca_classic | AAV | 1067 |
| 267150949 | Slc17a7-IRES2-C | mca_classic | AAV | 431 |
| 267151656 | Chat-IRES-Cre-n | mca_classic | AAV | 929 |
| 267152406 | Crh-IRES-Cre_ZJ | mca_classic | AAV | 595 |
| 267153115 | Crh-IRES-Cre_ZJ | mca_classic | AAV | 985 |
| 267211671 | Pcdh9-Cre_NP2 | mca_classic | AAV | 507 |
| 267213087 | Calb2-IRES-Cre | mca_classic | AAV | 794 |
| 267213793 | Calb2-IRES-Cre | mca_classic | AAV | 792 |
| 267396430 | Avp-IRES2-Cre | mca_classic | AAV | 794 |
| 267397226 | Avp-IRES2-Cre | mca_classic | AAV | 801 |
| 267397941 | A930038C07Rik | mca_classic | AAV | 279 |
| 267398651 | Avp-IRES2-Cre | mca_classic | AAV | 781 |
| 267493760 | Calb2-IRES-Cre | mca_classic | AAV | 668 |
| 267494468 | C57Bl/6J | mca_classic | AAV | 685 |

| | | | | |
|---|---|---|---|---|
| 267538006 | Slc32a1-IRES-Cr | mca_classic | AAV | 707 |
| 267538735 | Erbb4-T2A-CreE | mca_classic | AAV | 870 |
| 267540168 | Fezf1-T2A-dCre | mca_classic | AAV | 787 |
| 267547788 | Fezf1-T2A-dCre | mca_classic | AAV | 600 |
| 267607635 | Prkcd-GluCla-CF | mca_classic | AAV | 685 |
| 267608343 | Prkcd-GluCla-CF | mca_classic | AAV | 682 |
| 267609756 | Prkcd-GluCla-CF | mca_classic | AAV | 675 |
| 267610466 | Prkcd-GluCla-CF | mca_classic | AAV | 685 |
| 267611175 | Prkcd-GluCla-CF | mca_classic | AAV | 1084 |
| 267657327 | Gpr26-Cre_KO2 | mca_classic | AAV | 171 |
| 267658040 | Gpr26-Cre_KO2 | mca_classic | AAV | 332 |
| 267658747 | Gpr26-Cre_KO2 | mca_classic | AAV | 65 |
| 267659565 | Efr3a-Cre_NO1( | mca_classic | AAV | 24 |
| 267660272 | Efr3a-Cre_NO1( | mca_classic | AAV | 51 |
| 267661018 | Efr3a-Cre_NO1( | mca_classic | AAV | 332 |
| 267703239 | Slc6a5-Cre_KF1 | mca_classic | AAV | 975 |
| 267704653 | Efr3a-Cre_NO1( | mca_classic | AAV | 185 |
| 267705378 | Efr3a-Cre_NO1( | mca_classic | AAV | 226 |
| 267749107 | Efr3a-Cre_NO1( | mca_classic | AAV | 232 |
| 267749821 | Chrna2-Cre_OE | mca_classic | AAV | 44 |
| 267750528 | Chrna2-Cre_OE | mca_classic | AAV | 24 |
| 267760731 | Efr3a-Cre_NO1( | mca_classic | AAV | 386 |
| 267761438 | Efr3a-Cre_NO1( | mca_classic | AAV | 346 |
| 267762146 | Oxtr-Cre_ON66 | mca_classic | AAV | 573 |
| 267762859 | Oxtr-Cre_ON66 | mca_classic | AAV | 291 |
| 267763584 | Ppp1r17-Cre_N | mca_classic | AAV | 621 |
| 267764292 | Ppp1r17-Cre_N | mca_classic | AAV | 621 |
| 267810394 | Ppp1r17-Cre_N | mca_classic | AAV | 588 |
| 267811103 | Ppp1r17-Cre_N | mca_classic | AAV | 1067 |
| 267813224 | Nos1-CreERT2 | mca_classic | AAV | 24 |
| 267813932 | Etv1-CreERT2 | mca_classic | AAV | 566 |
| 267928135 | Ppp1r17-Cre_N | mca_classic | AAV | 757 |
| 267928844 | Prkcd-GluCla-CF | mca_classic | AAV | 685 |
| 267929554 | Prkcd-GluCla-CF | mca_classic | AAV | 645 |
| 267930268 | Nos1-CreERT2 | mca_classic | AAV | 136 |
| 267930978 | Nos1-CreERT2 | mca_classic | AAV | 24 |
| 267931685 | Nos1-CreERT2 | mca_classic | AAV | 185 |
| 267958444 | Slc6a4-Cre_ET3 | mca_classic | AAV | 925 |
| 267959197 | Vipr2-Cre_KE2 | mca_classic | AAV | 701 |
| 267997620 | Avp-IRES2-Cre | mca_classic | AAV | 720 |
| 267998328 | Grik4-Cre | mca_classic | AAV | 507 |
| 267999034 | Grik4-Cre | mca_classic | AAV | 507 |
| 267999740 | Grik4-Cre | mca_classic | AAV | 669 |
| 268038262 | Scnn1a-Tg3-Cre | mca_classic | AAV | 232 |
| 268038969 | Scnn1a-Tg3-Cre | mca_classic | AAV | 171 |
| 268039675 | Scnn1a-Tg3-Cre | mca_classic | AAV | 226 |
| 268041088 | Scnn1a-Tg3-Cre | mca_classic | AAV | 535 |

| | | | | |
|---|---|---|---|---|
| 268042502 | Slc18a2-Cre_OZ | mca_classic | AAV | 100 |
| 268076421 | Vipr2-Cre_KE2 | mca_classic | AAV | 685 |
| 268163228 | Vipr2-Cre_KE2 | mca_classic | AAV | 689 |
| 268165349 | Ppp1r17-Cre_N | mca_classic | AAV | 1067 |
| 268204599 | Ppp1r17-Cre_N | mca_classic | AAV | 792 |
| 268205344 | Ppp1r17-Cre_N | mca_classic | AAV | 645 |
| 268206050 | Ppp1r17-Cre_N | mca_classic | AAV | 649 |
| 268208632 | Ppp1r17-Cre_N | mca_classic | AAV | 792 |
| 268321221 | Erbb4-T2A-CreE | mca_classic | AAV | 945 |
| 268321927 | Chat-IRES-Cre-n | mca_classic | AAV | 712 |
| 268323342 | Ppp1r17-Cre_N | mca_classic | AAV | 948 |
| 268389532 | C57Bl/6J | mca_classic | AAV | 1098 |
| 268399145 | Ppp1r17-Cre_N | mca_classic | AAV | 685 |
| 268399868 | Ppp1r17-Cre_N | mca_classic | AAV | 649 |
| 268415561 | C57Bl/6J | mca_classic | AAV | 890 |
| 272404772 | C57Bl/6J | mca_classic | AAV | 467 |
| 272414403 | C57Bl/6J | mca_classic | AAV | 494 |
| 272697238 | Th-Cre_FI172 | mca_classic | AAV | 573 |
| 272698650 | Rbp4-Cre_KL10 | mca_classic | AAV | 65 |
| 272699357 | Th-Cre_FI172 | mca_classic | AAV | 838 |
| 272700063 | C57Bl/6J | mca_classic | AAV | 931 |
| 272735030 | Rbp4-Cre_KL10 | mca_classic | AAV | 51 |
| 272735744 | Cux2-IRES-Cre | mca_classic | AAV | 332 |
| 272736450 | Cux2-IRES-Cre | mca_classic | AAV | 213 |
| 272738620 | C57Bl/6J | mca_classic | AAV | 955 |
| 272781246 | Cux2-IRES-Cre | mca_classic | AAV | 264 |
| 272819994 | Adcyap1-2A-Cre | mca_classic | AAV | 600 |
| 272821309 | Cux2-IRES-Cre | mca_classic | AAV | 185 |
| 272822110 | Cux2-IRES-Cre | mca_classic | AAV | 24 |
| 272825299 | C57Bl/6J | mca_classic | AAV | 942 |
| 272826010 | C57Bl/6J | mca_classic | AAV | 913 |
| 272827141 | Cux2-IRES-Cre | mca_classic | AAV | 285 |
| 272829024 | Pcdh9-Cre_NP2 | mca_classic | AAV | 919 |
| 272829745 | C57Bl/6J | mca_classic | AAV | 838 |
| 272830456 | C57Bl/6J | mca_classic | AAV | 685 |
| 272873704 | C57Bl/6J | mca_classic | AAV | 671 |
| 272874417 | C57Bl/6J | mca_classic | AAV | 692 |
| 272875132 | C57Bl/6J | mca_classic | AAV | 685 |
| 272875838 | C57Bl/6J | mca_classic | AAV | 685 |
| 272917631 | C57Bl/6J | mca_classic | AAV | 588 |
| 272918345 | C57Bl/6J | mca_classic | AAV | 386 |
| 272929308 | Rbp4-Cre_KL10 | mca_classic | AAV | 353 |
| 272967913 | C57Bl/6J | mca_classic | AAV | 682 |
| 272968624 | C57Bl/6J | mca_classic | AAV | 701 |
| 272969333 | C57Bl/6J | mca_classic | AAV | 682 |
| 272970747 | C57Bl/6J | mca_classic | AAV | 685 |
| 273025166 | C57Bl/6J | mca_classic | AAV | 689 |

| | | | | |
|---|---|---|---|---|
| 273025872 | C57Bl/6J | mca_classic | AAV | 645 |
| 273055501 | C57Bl/6J | mca_classic | AAV | 764 |
| 273065264 | C57Bl/6J | mca_classic | AAV | 826 |
| 273068974 | C57Bl/6J | mca_classic | AAV | 985 |
| 277615922 | C57Bl/6J | mca_classic | AAV | 787 |
| 277618054 | C57Bl/6J | mca_classic | AAV | 1003 |
| 277618762 | C57Bl/6J | mca_classic | AAV | 1003 |
| 277710753 | C57Bl/6J | mca_classic | AAV | 562 |
| 277799582 | Ntrk1-IRES-Cre | mca_classic | AAV | 911 |
| 277800288 | Ntrk1-IRES-Cre | mca_classic | AAV | 744 |
| 277800995 | Sst-IRES-Cre | mca_classic | AAV | 24 |
| 277801701 | Sst-IRES-Cre | mca_classic | AAV | 51 |
| 277849256 | Cck-IRES-Cre | mca_classic | AAV | 685 |
| 277849965 | Nr5a1-Cre | mca_classic | AAV | 178 |
| 277851379 | Nr5a1-Cre | mca_classic | AAV | 353 |
| 277852088 | Scnn1a-Tg3-Cre | mca_classic | AAV | 72 |
| 277853501 | Crh-IRES-Cre_Bl | mca_classic | AAV | 911 |
| 277854208 | Erbb4-T2A-CreE | mca_classic | AAV | 787 |
| 277854916 | Erbb4-T2A-CreE | mca_classic | AAV | 787 |
| 277855624 | Crh-IRES-Cre_Bl | mca_classic | AAV | 264 |
| 277856332 | Crh-IRES-Cre_Bl | mca_classic | AAV | 595 |
| 277906744 | Crh-IRES-Cre_Bl | mca_classic | AAV | 981 |
| 277907462 | Efr3a-Cre_NO1( | mca_classic | AAV | 86 |
| 277908168 | Efr3a-Cre_NO1( | mca_classic | AAV | 226 |
| 277956496 | Htr2a-Cre_KM2 | mca_classic | AAV | 264 |
| 277957908 | Htr2a-Cre_KM2 | mca_classic | AAV | 18 |
| 277958616 | Pvalb-IRES-Cre | mca_classic | AAV | 649 |
| 277959325 | Pvalb-IRES-Cre | mca_classic | AAV | 812 |
| 278066728 | Pvalb-IRES-Cre | mca_classic | AAV | 984 |
| 278067445 | Prkcd-GluCla-CF | mca_classic | AAV | 685 |
| 278068196 | Chrna2-Cre_OE | mca_classic | AAV | 185 |
| 278069982 | Chrna2-Cre_OE | mca_classic | AAV | 226 |
| 278070717 | Gal-Cre_KI87 | mca_classic | AAV | 662 |
| 278171908 | Oxtr-Cre_ON66 | mca_classic | AAV | 258 |
| 278173743 | Chrna2-Cre_OE | mca_classic | AAV | 232 |
| 278174451 | Chrna2-Cre_OE | mca_classic | AAV | 258 |
| 278175580 | Chrnb4-Cre_OL | mca_classic | AAV | 346 |
| 278178382 | Chrna2-Cre_OE | mca_classic | AAV | 823 |
| 278179088 | Crh-IRES-Cre_Bl | mca_classic | AAV | 595 |
| 278179794 | Efr3a-Cre_NO1( | mca_classic | AAV | 325 |
| 278180500 | Crh-IRES-Cre_Bl | mca_classic | AAV | 934 |
| 278181912 | Efr3a-Cre_NO1( | mca_classic | AAV | 24 |
| 278257366 | Efr3a-Cre_NO1( | mca_classic | AAV | 65 |
| 278258073 | Efr3a-Cre_NO1( | mca_classic | AAV | 24 |
| 278258780 | Efr3a-Cre_NO1( | mca_classic | AAV | 24 |
| 278259822 | Oxtr-Cre_ON66 | mca_classic | AAV | 617 |
| 278260569 | Oxtr-Cre_ON66 | mca_classic | AAV | 792 |

| | | | | |
|---|---|---|---|---|
| 278261300 | Oxtr-Cre_ON66 | mca_classic | AAV | 653 |
| 278317239 | Cux2-IRES-Cre | mca_classic | AAV | 494 |
| 278317945 | Cux2-IRES-Cre | mca_classic | AAV | 72 |
| 278318653 | Ppp1r17-Cre_N | mca_classic | AAV | 467 |
| 278319402 | Ppp1r17-Cre_N | mca_classic | AAV | 467 |
| 278398243 | Ppp1r17-Cre_N | mca_classic | AAV | 467 |
| 278398949 | Ppp1r17-Cre_N | mca_classic | AAV | 707 |
| 278399657 | Ppp1r17-Cre_N | mca_classic | AAV | 467 |
| 278400363 | Ppp1r17-Cre_N | mca_classic | AAV | 685 |
| 278401072 | Slc17a6-IRES-Cr | mca_classic | AAV | 890 |
| 278401778 | Htr2a-Cre_KM2 | mca_classic | AAV | 844 |
| 278433737 | Cux2-IRES-Cre | mca_classic | AAV | 238 |
| 278434443 | Penk-2A-CreER | mca_classic | AAV | 573 |
| 278435152 | Wfs1-Tg2-CreE | mca_classic | AAV | 507 |
| 278435864 | Wfs1-Tg2-CreE | mca_classic | AAV | 507 |
| 278500868 | Htr2a-Cre_KM2 | mca_classic | AAV | 694 |
| 278501857 | Htr2a-Cre_KM2 | mca_classic | AAV | 611 |
| 278502708 | Scnn1a-Tg3-Cre | mca_classic | AAV | 232 |
| 278503555 | Ppp1r17-Cre_N | mca_classic | AAV | 457 |
| 278504263 | Ppp1r17-Cre_N | mca_classic | AAV | 473 |
| 278508779 | Ppp1r17-Cre_N | mca_classic | AAV | 794 |
| 278510197 | Ppp1r17-Cre_N | mca_classic | AAV | 794 |
| 278510903 | Ppp1r17-Cre_N | mca_classic | AAV | 692 |
| 278511717 | Ppp1r17-Cre_N | mca_classic | AAV | 830 |
| 278513131 | Ppp1r17-Cre_N | mca_classic | AAV | 830 |
| 281459203 | A930038C07Rik | mca_classic | AAV | 86 |
| 283017324 | A930038C07Rik | mca_classic | AAV | 178 |
| 283019341 | A930038C07Rik | mca_classic | AAV | 245 |
| 283020912 | A930038C07Rik | mca_classic | AAV | 185 |
| 284665639 | Grm2-Cre_MR9 | mca_classic | AAV | 685 |
| 286299886 | Cux2-IRES-Cre | mca_classic | AAV | 6 |
| 286300594 | Cux2-IRES-Cre | mca_classic | AAV | 79 |
| 286301303 | Cux2-IRES-Cre | mca_classic | AAV | 353 |
| 286303000 | Slc6a3-Cre | mca_classic | AAV | 600 |
| 286313491 | Rbp4-Cre_KL10 | mca_classic | AAV | 245 |
| 286314623 | Rbp4-Cre_KL10 | mca_classic | AAV | 100 |
| 286317619 | Slc17a7-IRES2-C | mca_classic | AAV | 457 |
| 286318327 | Ins2-Cre_25 | mca_classic | AAV | 733 |
| 286319033 | Ins2-Cre_25 | mca_classic | AAV | 787 |
| 286319739 | Ntrk1-IRES-Cre | mca_classic | AAV | 618 |
| 286417464 | Rbp4-Cre_KL10 | mca_classic | AAV | 192 |
| 286482701 | Rbp4-Cre_KL10 | mca_classic | AAV | 232 |
| 286483411 | Rbp4-Cre_KL10 | mca_classic | AAV | 494 |
| 286484879 | Syt17-Cre_NO1 | mca_classic | AAV | 507 |
| 286485585 | Syt17-Cre_NO1 | mca_classic | AAV | 685 |
| 286486329 | Gpr26-Cre_KO2 | mca_classic | AAV | 575 |
| 286553311 | Gpr26-Cre_KO2 | mca_classic | AAV | 675 |

| | | | | |
|---|---|---|---|---|
| 286554017 | Gpr26-Cre_KO2 | mca_classic | AAV | 457 |
| 286554724 | Nr5a1-Cre | mca_classic | AAV | 206 |
| 286556208 | Slc17a6-IRES-Cr | mca_classic | AAV | 787 |
| 286556914 | Slc17a6-IRES-Cr | mca_classic | AAV | 941 |
| 286608092 | Slc17a6-IRES-Cr | mca_classic | AAV | 1099 |
| 286609510 | Slc17a6-IRES-Cr | mca_classic | AAV | 975 |
| 286610216 | Syt17-Cre_NO1 | mca_classic | AAV | 457 |
| 286610923 | Syt17-Cre_NO1 | mca_classic | AAV | 457 |
| 286646170 | Syt17-Cre_NO1 | mca_classic | AAV | 685 |
| 286646877 | Tac1-IRES2-Cre | mca_classic | AAV | 226 |
| 286647583 | Slc32a1-IRES-Cr | mca_classic | AAV | 984 |
| 286648290 | Efr3a-Cre_NO1( | mca_classic | AAV | 264 |
| 286648997 | Efr3a-Cre_NO1( | mca_classic | AAV | 51 |
| 286649703 | Ppp1r17-Cre_N | mca_classic | AAV | 573 |
| 286662551 | Gad2-IRES-Cre | mca_classic | AAV | 802 |
| 286725359 | Pvalb-IRES-Cre | mca_classic | AAV | 992 |
| 286726065 | Avp-IRES2-Cre | mca_classic | AAV | 717 |
| 286726777 | Th-Cre_FI172 | mca_classic | AAV | 733 |
| 286727483 | Ppp1r17-Cre_N | mca_classic | AAV | 757 |
| 286728190 | Cux2-CreERT2 | mca_classic | AAV | 416 |
| 286728896 | Th-Cre_FI172 | mca_classic | AAV | 781 |
| 286772650 | Etv1-CreERT2 | mca_classic | AAV | 51 |
| 286773358 | Efr3a-Cre_NO1( | mca_classic | AAV | 185 |
| 286774064 | Etv1-CreERT2 | mca_classic | AAV | 595 |
| 286774770 | A930038C07Rik | mca_classic | AAV | 226 |
| 286775476 | A930038C07Rik | mca_classic | AAV | 285 |
| 286834976 | A930038C07Rik | mca_classic | AAV | 100 |
| 286835688 | Efr3a-Cre_NO1( | mca_classic | AAV | 178 |
| 286838022 | Oxtr-Cre_ON66 | mca_classic | AAV | 100 |
| 286882342 | Ppp1r17-Cre_N | mca_classic | AAV | 794 |
| 287027773 | Ppp1r17-Cre_N | mca_classic | AAV | 561 |
| 287028480 | Rbp4-Cre_KL10 | mca_classic | AAV | 507 |
| 287029186 | Dbh-Cre_KH212 | mca_classic | AAV | 988 |
| 287029894 | Dbh-Cre_KH212 | mca_classic | AAV | 992 |
| 287036898 | Ntsr1-Cre_GN2 | mca_classic | AAV | 178 |
| 287037604 | Ntsr1-Cre_GN2 | mca_classic | AAV | 100 |
| 287039061 | Slc18a2-Cre_OZ | mca_classic | AAV | 100 |
| 287044088 | Avp-IRES2-Cre | mca_classic | AAV | 720 |
| 287095785 | Slc18a2-Cre_OZ | mca_classic | AAV | 932 |
| 287096578 | Slc18a2-Cre_OZ | mca_classic | AAV | 973 |
| 287097454 | Grm2-Cre_MR9 | mca_classic | AAV | 473 |
| 287171983 | Grm2-Cre_MR9 | mca_classic | AAV | 473 |
| 287172689 | Grm2-Cre_MR9 | mca_classic | AAV | 531 |
| 287173396 | Grm2-Cre_MR9 | mca_classic | AAV | 764 |
| 287174103 | Rorb-IRES2-Cre | mca_classic | AAV | 51 |
| 287223629 | Rorb-IRES2-Cre | mca_classic | AAV | 136 |
| 287224335 | Rorb-IRES2-Cre | mca_classic | AAV | 185 |

| | | | | |
|---|---|---|---|---|
| 287225041 | Rorb-IRES2-Cre | mca_classic | AAV | 199 |
| 287225747 | Adcyap1-2A-Cre | mca_classic | AAV | 416 |
| 287246555 | Adcyap1-2A-Cre | mca_classic | AAV | 776 |
| 287247978 | Adcyap1-2A-Cre | mca_classic | AAV | 838 |
| 287248684 | Gpr26-Cre_KO2 | mca_classic | AAV | 226 |
| 287446625 | Grik4-Cre | mca_classic | AAV | 703 |
| 287447332 | Grik4-Cre | mca_classic | AAV | 647 |
| 287448038 | Slc17a6-IRES-Cr | mca_classic | AAV | 812 |
| 287452840 | Slc17a6-IRES-Cr | mca_classic | AAV | 812 |
| 287458189 | Slc18a2-Cre_OZ | mca_classic | AAV | 662 |
| 287458895 | Slc18a2-Cre_OZ | mca_classic | AAV | 662 |
| 287459601 | Ppp1r17-Cre_N | mca_classic | AAV | 808 |
| 287460307 | Cck-IRES-Cre | mca_classic | AAV | 954 |
| 287461719 | A930038C07Rik | mca_classic | AAV | 18 |
| 287492899 | Slc18a2-Cre_OZ | mca_classic | AAV | 802 |
| 287494320 | Cux2-IRES-Cre | mca_classic | AAV | 238 |
| 287495026 | Cux2-IRES-Cre | mca_classic | AAV | 178 |
| 287533790 | Th-Cre_FI172 | mca_classic | AAV | 575 |
| 287538943 | Gad2-IRES-Cre | mca_classic | AAV | 739 |
| 287539649 | Gad2-IRES-Cre | mca_classic | AAV | 1067 |
| 287598261 | Gad2-IRES-Cre | mca_classic | AAV | 826 |
| 287599685 | Trib2-F2A-CreEI | mca_classic | AAV | 226 |
| 287600392 | Trib2-F2A-CreEI | mca_classic | AAV | 232 |
| 287601100 | Trib2-F2A-CreEI | mca_classic | AAV | 332 |
| 287601808 | Trib2-F2A-CreEI | mca_classic | AAV | 171 |
| 287633753 | Syt6-Cre_KI148 | mca_classic | AAV | 24 |
| 287664997 | Gad2-IRES-Cre | mca_classic | AAV | 941 |
| 287665706 | Avp-IRES2-Cre | mca_classic | AAV | 752 |
| 287667137 | Fezf1-T2A-dCre | mca_classic | AAV | 794 |
| 287668550 | Nxph4-T2A-CreI | mca_classic | AAV | 902 |
| 287712779 | Nxph4-T2A-CreI | mca_classic | AAV | 838 |
| 287713485 | Nxph4-T2A-CreI | mca_classic | AAV | 1024 |
| 287714197 | Nxph4-T2A-CreI | mca_classic | AAV | 812 |
| 287714903 | Htr2a-Cre_KM2 | mca_classic | AAV | 232 |
| 287715650 | Htr2a-Cre_KM2 | mca_classic | AAV | 390 |
| 287769286 | Rbp4-Cre_KL10 | mca_classic | AAV | 272 |
| 287769992 | Scnn1a-Tg3-Cre | mca_classic | AAV | 494 |
| 287772112 | Scnn1a-Tg3-Cre | mca_classic | AAV | 531 |
| 287807030 | Sim1-Cre_KJ18 | mca_classic | AAV | 24 |
| 287807743 | Sim1-Cre_KJ18 | mca_classic | AAV | 24 |
| 287808449 | Slc17a6-IRES-Cr | mca_classic | AAV | 880 |
| 287809155 | Gad2-IRES-Cre | mca_classic | AAV | 1033 |
| 287876212 | Gad2-IRES-Cre | mca_classic | AAV | 975 |
| 287879384 | Grm2-Cre_MR9 | mca_classic | AAV | 1071 |
| 287880102 | Rorb-IRES2-Cre | mca_classic | AAV | 65 |
| 287880821 | Calb2-IRES-Cre | mca_classic | AAV | 702 |
| 287950390 | Calb2-IRES-Cre | mca_classic | AAV | 619 |

| | | | | |
|---|---|---|---|---|
| 287951098 | Cux2-IRES-Cre | mca_classic | AAV | 100 |
| 287952517 | Cux2-IRES-Cre | mca_classic | AAV | 494 |
| 287953929 | Cux2-IRES-Cre | mca_classic | AAV | 213 |
| 287993060 | Ntrk1-IRES-Cre | mca_classic | AAV | 575 |
| 287994474 | Ntrk1-IRES-Cre | mca_classic | AAV | 573 |
| 287995180 | Ntrk1-IRES-Cre | mca_classic | AAV | 573 |
| 287996596 | Efr3a-Cre_NO1( | mca_classic | AAV | 100 |
| 288168426 | Efr3a-Cre_NO1( | mca_classic | AAV | 185 |
| 288169135 | Efr3a-Cre_NO1( | mca_classic | AAV | 18 |
| 288169842 | Tlx3-Cre_PL56 | mca_classic | AAV | 24 |
| 288170549 | Tlx3-Cre_PL56 | mca_classic | AAV | 51 |
| 288171964 | Tlx3-Cre_PL56 | mca_classic | AAV | 185 |
| 288261928 | Sst-IRES-Cre | mca_classic | AAV | 185 |
| 288262635 | Sst-IRES-Cre | mca_classic | AAV | 226 |
| 288263341 | Sst-IRES-Cre | mca_classic | AAV | 272 |
| 288264047 | Sst-IRES-Cre | mca_classic | AAV | 171 |
| 288264753 | Sst-IRES-Cre | mca_classic | AAV | 332 |
| 288321385 | Sst-IRES-Cre | mca_classic | AAV | 507 |
| 288322797 | Syt17-Cre_NO1 | mca_classic | AAV | 473 |
| 288324211 | Grp-Cre_KH288 | mca_classic | AAV | 258 |
| 292034715 | Grp-Cre_KH288 | mca_classic | AAV | 24 |
| 292035484 | Slc17a6-IRES-Cr | mca_classic | AAV | 757 |
| 292123352 | Slc17a6-IRES-Cr | mca_classic | AAV | 744 |
| 292124058 | Chrna2-Cre_OE | mca_classic | AAV | 332 |
| 292124765 | Chrna2-Cre_OE | mca_classic | AAV | 86 |
| 292125472 | Chrna2-Cre_OE | mca_classic | AAV | 226 |
| 292126180 | Chrna2-Cre_OE | mca_classic | AAV | 185 |
| 292172100 | Chrna2-Cre_OE | mca_classic | AAV | 332 |
| 292172846 | Chrna2-Cre_OE | mca_classic | AAV | 24 |
| 292173552 | Chrna2-Cre_OE | mca_classic | AAV | 79 |
| 292174264 | Chrna2-Cre_OE | mca_classic | AAV | 206 |
| 292174974 | Chrna2-Cre_OE | mca_classic | AAV | 24 |
| 292208876 | Chrna2-Cre_OE | mca_classic | AAV | 360 |
| 292209592 | Chrna2-Cre_OE | mca_classic | AAV | 279 |
| 292210312 | Chrna2-Cre_OE | mca_classic | AAV | 24 |
| 292211026 | Fezf1-T2A-dCre | mca_classic | AAV | 787 |
| 292211743 | Gad2-IRES-Cre | mca_classic | AAV | 808 |
| 292212456 | Grik4-Cre | mca_classic | AAV | 669 |
| 292318449 | Efr3a-Cre_NO1( | mca_classic | AAV | 72 |
| 292319155 | Prkcd-GluCla-CF | mca_classic | AAV | 941 |
| 292319865 | Prkcd-GluCla-CF | mca_classic | AAV | 668 |
| 292320572 | Prkcd-GluCla-CF | mca_classic | AAV | 662 |
| 292321278 | Prkcd-GluCla-CF | mca_classic | AAV | 675 |
| 292372636 | Oxtr-Cre_ON66 | mca_classic | AAV | 114 |
| 292374068 | A930038C07Rik | mca_classic | AAV | 24 |
| 292374777 | A930038C07Rik | mca_classic | AAV | 238 |
| 292375491 | A930038C07Rik | mca_classic | AAV | 107 |

| | | | | |
|---|---|---|---|---|
| 292476595 | A930038C07Rik | mca_classic | AAV | 24 |
| 292477301 | Grp-Cre_KH288 | mca_classic | AAV | 264 |
| 292478008 | Grp-Cre_KH288 | mca_classic | AAV | 675 |
| 292479421 | Ppp1r17-Cre_N | mca_classic | AAV | 457 |
| 292480129 | Ppp1r17-Cre_N | mca_classic | AAV | 792 |
| 292530653 | Wfs1-Tg2-CreEI | mca_classic | AAV | 531 |
| 292531359 | Grm2-Cre_MR9 | mca_classic | AAV | 1033 |
| 292532065 | Ppp1r17-Cre_N | mca_classic | AAV | 457 |
| 292532771 | Ppp1r17-Cre_N | mca_classic | AAV | 457 |
| 292533477 | Ppp1r17-Cre_N | mca_classic | AAV | 830 |
| 292620251 | Trib2-F2A-CreEI | mca_classic | AAV | 65 |
| 292620968 | Cux2-CreERT2 | mca_classic | AAV | 573 |
| 292622743 | Crh-IRES-Cre_ZJ | mca_classic | AAV | 981 |
| 292623457 | Grm2-Cre_MR9 | mca_classic | AAV | 713 |
| 292624169 | Grm2-Cre_MR9 | mca_classic | AAV | 838 |
| 292790567 | Grm2-Cre_MR9 | mca_classic | AAV | 890 |
| 292791310 | Syt6-Cre_KI148 | mca_classic | AAV | 206 |
| 292792016 | A930038C07Rik | mca_classic | AAV | 79 |
| 292792724 | A930038C07Rik | mca_classic | AAV | 353 |
| 292793967 | A930038C07Rik | mca_classic | AAV | 185 |
| 292794673 | Slc18a2-Cre_OZ | mca_classic | AAV | 1013 |
| 292958638 | Slc18a2-Cre_OZ | mca_classic | AAV | 823 |
| 292959343 | Chat-IRES-Cre-n | mca_classic | AAV | 573 |
| 292960052 | Ntrk1-IRES-Cre | mca_classic | AAV | 703 |
| 292960764 | Ntrk1-IRES-Cre | mca_classic | AAV | 1003 |
| 292961470 | Ntrk1-IRES-Cre | mca_classic | AAV | 618 |
| 292962283 | Ntrk1-IRES-Cre | mca_classic | AAV | 577 |
| 293008559 | Ntrk1-IRES-Cre | mca_classic | AAV | 575 |
| 293009265 | Syt17-Cre_NO1 | mca_classic | AAV | 494 |
| 293009970 | Syt17-Cre_NO1 | mca_classic | AAV | 535 |
| 293011049 | Syt17-Cre_NO1 | mca_classic | AAV | 473 |
| 293011759 | Syt17-Cre_NO1 | mca_classic | AAV | 473 |
| 293114113 | Crh-IRES-Cre_BI | mca_classic | AAV | 735 |
| 293115527 | Syt17-Cre_NO1 | mca_classic | AAV | 473 |
| 293116943 | Nos1-CreERT2 | mca_classic | AAV | 185 |
| 293252166 | Pvalb-IRES-Cre | mca_classic | AAV | 867 |
| 293254286 | Tac2-IRES2-Cre | mca_classic | AAV | 794 |
| 293255030 | Htr2a-Cre_KM2 | mca_classic | AAV | 792 |
| 293365328 | Plxnd1-Cre_OG | mca_classic | AAV | 685 |
| 293366035 | Plxnd1-Cre_OG | mca_classic | AAV | 573 |
| 293366741 | Plxnd1-Cre_OG | mca_classic | AAV | 573 |
| 293367448 | Plxnd1-Cre_OG | mca_classic | AAV | 685 |
| 293368154 | Plxnd1-Cre_OG | mca_classic | AAV | 794 |
| 293431163 | Vip-IRES-Cre | mca_classic | AAV | 752 |
| 293431869 | Tlx3-Cre_PL56 | mca_classic | AAV | 226 |
| 293432575 | Tlx3-Cre_PL56 | mca_classic | AAV | 226 |
| 293433283 | Tlx3-Cre_PL56 | mca_classic | AAV | 272 |

| | | | | | |
|---|---|---|---|---|---|
| | 293433996 | Htr2a-Cre_KM2 | mca_classic | AAV | 6 |
| | 293469501 | Htr2a-Cre_KM2 | mca_classic | AAV | 600 |
| | 293471629 | Cux2-IRES-Cre | mca_classic | AAV | 264 |
| | 293472335 | Cux2-IRES-Cre | mca_classic | AAV | 185 |
| | 293546902 | Slc17a6-IRES-Cr | mca_classic | AAV | 386 |
| | 293548317 | Nr5a1-Cre | mca_classic | AAV | 353 |
| | 293549729 | Gad2-IRES-Cre | mca_classic | AAV | 776 |
| | 293550435 | Gad2-IRES-Cre | mca_classic | AAV | 1084 |
| | 293700454 | Rorb-IRES2-Cre | mca_classic | AAV | 65 |
| | 293701770 | Drd3-Cre_KI196 | mca_classic | AAV | 544 |
| | 293728197 | Drd3-Cre_KI196 | mca_classic | AAV | 51 |
| | 293750063 | Calb1-T2A-dgCr | mca_classic | AAV | 507 |
| | 293752263 | Crh-IRES-Cre_BI | mca_classic | AAV | 913 |
| | 293787288 | Slc17a7-IRES2-C | mca_classic | AAV | 662 |
| | 293787994 | Slc17a7-IRES2-C | mca_classic | AAV | 473 |
| | 293788700 | Slc17a7-IRES2-C | mca_classic | AAV | 473 |
| | 293820681 | Sim1-Cre_KJ18 | mca_classic | AAV | 136 |
| | 293821389 | Sim1-Cre_KJ18 | mca_classic | AAV | 171 |
| | 293888501 | Nos1-CreERT2 | mca_classic | AAV | 473 |
| | 293889216 | Nos1-CreERT2 | mca_classic | AAV | 258 |
| | 293889922 | Chrna2-Cre_OE | mca_classic | AAV | 51 |
| | 293914056 | Slc17a6-IRES-Cr | mca_classic | AAV | 668 |
| | 293914766 | Slc17a6-IRES-Cr | mca_classic | AAV | 662 |
| | 293942188 | Slc17a6-IRES-Cr | mca_classic | AAV | 795 |
| | 293942897 | Gad2-IRES-Cre | mca_classic | AAV | 662 |
| | 294005186 | Gad2-IRES-Cre | mca_classic | AAV | 744 |
| | 294040662 | Gad2-IRES-Cre | mca_classic | AAV | 694 |
| | 294173584 | Slc17a6-IRES-Cr | mca_classic | AAV | 988 |
| | 294174290 | Slc17a6-IRES-Cr | mca_classic | AAV | 830 |
| | 294174996 | Slc17a6-IRES-Cr | mca_classic | AAV | 421 |
| | 294175704 | Slc17a6-IRES-Cr | mca_classic | AAV | 847 |
| | 294199114 | Slc17a6-IRES-Cr | mca_classic | AAV | 817 |
| | 294266031 | Efr3a-Cre_NO1( | mca_classic | AAV | 178 |
| | 294311830 | Oxtr-Cre_ON66 | mca_classic | AAV | 258 |
| | 294312537 | Oxtr-Cre_ON66 | mca_classic | AAV | 185 |
| | 294313246 | Tac1-IRES2-Cre | mca_classic | AAV | 390 |
| | 294314037 | Tac1-IRES2-Cre | mca_classic | AAV | 535 |
| | 294316542 | Tac1-IRES2-Cre | mca_classic | AAV | 781 |
| | 294355509 | Tac1-IRES2-Cre | mca_classic | AAV | 776 |
| | 294356216 | Tac1-IRES2-Cre | mca_classic | AAV | 890 |
| | 294356922 | Slc17a6-IRES-Cr | mca_classic | AAV | 685 |
| | 294396492 | Cux2-IRES-Cre | mca_classic | AAV | 238 |
| | 294397199 | Cux2-IRES-Cre | mca_classic | AAV | 72 |
| | 294397907 | Cux2-IRES-Cre | mca_classic | AAV | 544 |
| | 294399325 | Oxtr-Cre_ON66 | mca_classic | AAV | 245 |
| Y | 294434161 | Cux2-IRES-Cre | mca_classic | AAV | 232 |
| | 294434867 | Cux2-IRES-Cre | mca_classic | AAV | 171 |

| | | | | | |
|---|---|---|---|---|---|
| | 294435580 | Cux2-IRES-Cre | mca_classic | AAV | 51 |
| | 294481346 | Rbp4-Cre_KL10 | mca_classic | AAV | 199 |
| | 294516943 | Tlx3-Cre_PL56 | mca_classic | AAV | 213 |
| | 294525229 | Tlx3-Cre_PL56 | mca_classic | AAV | 279 |
| | 294525944 | Tlx3-Cre_PL56 | mca_classic | AAV | 24 |
| | 294532700 | Tlx3-Cre_PL56 | mca_classic | AAV | 24 |
| | 294533406 | Tlx3-Cre_PL56 | mca_classic | AAV | 346 |
| | 296047084 | Chrna2-Cre_OE | mca_classic | AAV | 100 |
| Y | 296047806 | Cux2-IRES-Cre | mca_classic | AAV | 279 |
| | 296048512 | Rbp4-Cre_KL10 | mca_classic | AAV | 279 |
| | 296052133 | Tlx3-Cre_PL56 | mca_classic | AAV | 185 |
| | 296052839 | Tlx3-Cre_PL56 | mca_classic | AAV | 65 |
| | 297225422 | Tlx3-Cre_PL56 | mca_classic | AAV | 178 |
| | 297231636 | Tlx3-Cre_PL56 | mca_classic | AAV | 164 |
| | 297233422 | Tlx3-Cre_PL56 | mca_classic | AAV | 171 |
| | 297592527 | Tlx3-Cre_PL56 | mca_classic | AAV | 185 |
| | 297593235 | Tlx3-Cre_PL56 | mca_classic | AAV | 185 |
| | 297593942 | Tlx3-Cre_PL56 | mca_classic | AAV | 178 |
| | 297628576 | Ntsr1-Cre_GN2 | mca_classic | AAV | 51 |
| | 297629282 | Ntsr1-Cre_GN2 | mca_classic | AAV | 185 |
| | 297629988 | Ntsr1-Cre_GN2 | mca_classic | AAV | 178 |
| | 297652799 | Rbp4-Cre_KL10 | mca_classic | AAV | 65 |
| | 297654263 | Tlx3-Cre_PL56 | mca_classic | AAV | 51 |
| | 297668898 | Plxnd1-Cre_OG | mca_classic | AAV | 51 |
| | 297669605 | Plxnd1-Cre_OG | mca_classic | AAV | 24 |
| | 297670312 | Drd3-Cre_KI19 | mca_classic | AAV | 171 |
| | 297710633 | Sim1-Cre_KJ18 | mca_classic | AAV | 51 |
| | 297711339 | Sim1-Cre_KJ18 | mca_classic | AAV | 18 |
| | 297712045 | Sim1-Cre_KJ18 | mca_classic | AAV | 353 |
| | 297713365 | Sim1-Cre_KJ18 | mca_classic | AAV | 44 |
| | 297714071 | Sim1-Cre_KJ18 | mca_classic | AAV | 65 |
| | 297854981 | Sim1-Cre_KJ18 | mca_classic | AAV | 24 |
| | 297855879 | Cux2-IRES-Cre | mca_classic | AAV | 100 |
| | 297858011 | Chrna2-Cre_OE | mca_classic | AAV | 285 |
| | 297892130 | Tlx3-Cre_PL56 | mca_classic | AAV | 24 |
| | 297892843 | Chrna2-Cre_OE | mca_classic | AAV | 272 |
| | 297894255 | Chrna2-Cre_OE | mca_classic | AAV | 457 |
| | 297945448 | Chrna2-Cre_OE | mca_classic | AAV | 100 |
| | 297946154 | Chrna2-Cre_OE | mca_classic | AAV | 199 |
| | 297946935 | Sim1-Cre_KJ18 | mca_classic | AAV | 24 |
| | 297947641 | Sim1-Cre_KJ18 | mca_classic | AAV | 24 |
| | 297948420 | Sim1-Cre_KJ18 | mca_classic | AAV | 72 |
| | 297951732 | Sim1-Cre_KJ18 | mca_classic | AAV | 79 |
| | 297985755 | Sim1-Cre_KJ18 | mca_classic | AAV | 72 |
| | 297987980 | Chrnb4-Cre_OL | mca_classic | AAV | 24 |
| | 298000880 | Chrnb4-Cre_OL | mca_classic | AAV | 332 |
| | 298001595 | Lypd6-Cre_KL1 | mca_classic | AAV | 685 |

| | | | | |
|---|---|---|---|---|
| 298003295 | Htr2a-Cre_KM2 | mca_classic | AAV | 662 |
| 298004028 | Htr2a-Cre_KM2 | mca_classic | AAV | 662 |
| 298048079 | Hcrt-Cre | mca_classic | AAV | 794 |
| 298048787 | Hcrt-Cre | mca_classic | AAV | 794 |
| 298049545 | Lypd6-Cre_KL15 | mca_classic | AAV | 744 |
| 298050269 | Lypd6-Cre_KL15 | mca_classic | AAV | 692 |
| 298078515 | Lypd6-Cre_KL15 | mca_classic | AAV | 794 |
| 298079222 | Lypd6-Cre_KL15 | mca_classic | AAV | 867 |
| 298079928 | Lypd6-Cre_KL15 | mca_classic | AAV | 838 |
| 298104533 | Lypd6-Cre_KL15 | mca_classic | AAV | 763 |
| 298105299 | Ins2-Cre_25 | mca_classic | AAV | 733 |
| 298106713 | Cux2-IRES-Cre | mca_classic | AAV | 79 |
| 298178204 | Rbp4-Cre_KL10 | mca_classic | AAV | 213 |
| 298178912 | Ins2-Cre_25 | mca_classic | AAV | 781 |
| 298179622 | Rbp4-Cre_KL10 | mca_classic | AAV | 100 |
| 298182842 | Tlx3-Cre_PL56 | mca_classic | AAV | 86 |
| 298230624 | Efr3a-Cre_NO1( | mca_classic | AAV | 258 |
| 298232793 | A930038C07Rik | mca_classic | AAV | 473 |
| 298272589 | Chrna2-Cre_OE | mca_classic | AAV | 245 |
| 298273313 | Chrna2-Cre_OE | mca_classic | AAV | 18 |
| 298274021 | Chrna2-Cre_OE | mca_classic | AAV | 245 |
| 298275548 | Chrna2-Cre_OE | mca_classic | AAV | 332 |
| 298324391 | A930038C07Rik | mca_classic | AAV | 199 |
| 298325807 | Chrna2-Cre_OE | mca_classic | AAV | 18 |
| 298326521 | A930038C07Rik | mca_classic | AAV | 44 |
| 298350212 | Efr3a-Cre_NO1( | mca_classic | AAV | 185 |
| 298350922 | A930038C07Rik | mca_classic | AAV | 213 |
| 298351630 | A930038C07Rik | mca_classic | AAV | 185 |
| 298352336 | Efr3a-Cre_NO1( | mca_classic | AAV | 79 |
| 298404154 | Gpr26-Cre_KO2 | mca_classic | AAV | 86 |
| 298404860 | Gpr26-Cre_KO2 | mca_classic | AAV | 185 |
| 298718072 | Rasgrf2-T2A-dC | mca_classic | AAV | 24 |
| 298718778 | Rasgrf2-T2A-dC | mca_classic | AAV | 51 |
| 298719484 | Rasgrf2-T2A-dC | mca_classic | AAV | 136 |
| 298720191 | Gpr26-Cre_KO2 | mca_classic | AAV | 325 |
| 298720898 | Rbp4-Cre_KL10 | mca_classic | AAV | 416 |
| 298757426 | Rorb-IRES2-Cre | mca_classic | AAV | 178 |
| 298758841 | Rasgrf2-T2A-dC | mca_classic | AAV | 226 |
| 298759552 | Drd3-Cre_KI196 | mca_classic | AAV | 185 |
| 298760261 | Sim1-Cre_KJ18 | mca_classic | AAV | 51 |
| 298796577 | A930038C07Rik | mca_classic | AAV | 72 |
| 298797288 | A930038C07Rik | mca_classic | AAV | 232 |
| 298797996 | A930038C07Rik | mca_classic | AAV | 494 |
| 298798846 | A930038C07Rik | mca_classic | AAV | 494 |
| 298830161 | Cux2-IRES-Cre | mca_classic | AAV | 65 |
| 298830868 | Ntsr1-Cre_GN2: | mca_classic | AAV | 213 |
| 298831574 | A930038C07Rik | mca_classic | AAV | 535 |

| | | | | | |
|---|---|---|---|---|---|
| | 298833033 | Calb2-IRES-Cre | mca_classic | AAV | 682 |
| | 298833739 | Cart-Tg1-Cre | mca_classic | AAV | 739 |
| | 298835152 | Efr3a-Cre_NO1( | mca_classic | AAV | 613 |
| | 298835859 | Drd3-Cre_KI196 | mca_classic | AAV | 621 |
| | 299245589 | Gad2-IRES-Cre | mca_classic | AAV | 613 |
| | 299247009 | Kiss1-Cre | mca_classic | AAV | 744 |
| | 299403823 | Nos1-CreERT2 | mca_classic | AAV | 507 |
| | 299404532 | Ntrk1-IRES-Cre | mca_classic | AAV | 613 |
| | 299408890 | Oxtr-Cre_ON66 | mca_classic | AAV | 757 |
| | 299447153 | Scnn1a-Tg2-Cre | mca_classic | AAV | 507 |
| | 299447873 | Scnn1a-Tg3-Cre | mca_classic | AAV | 72 |
| | 299448592 | Oxtr-Cre_ON66 | mca_classic | AAV | 692 |
| | 299623085 | Slc32a1-IRES-Cr | mca_classic | AAV | 431 |
| | 299623794 | Sst-IRES-Cre | mca_classic | AAV | 707 |
| | 299624500 | Syt17-Cre_NO1 | mca_classic | AAV | 684 |
| | 299653551 | Syt17-Cre_NO1 | mca_classic | AAV | 410 |
| | 299654968 | Tac1-IRES2-Cre | mca_classic | AAV | 781 |
| | 299656382 | Vip-IRES-Cre | mca_classic | AAV | 457 |
| | 299732033 | Efr3a-Cre_NO1( | mca_classic | AAV | 1033 |
| | 299732738 | Efr3a-Cre_NO1( | mca_classic | AAV | 701 |
| | 299733445 | Syt6-Cre_KI148 | mca_classic | AAV | 199 |
| | 299759175 | Efr3a-Cre_NO1( | mca_classic | AAV | 844 |
| | 299759881 | Efr3a-Cre_NO1( | mca_classic | AAV | 782 |
| | 299760587 | Lypd6-Cre_KL1! | mca_classic | AAV | 802 |
| | 299761293 | Nos1-CreERT2 | mca_classic | AAV | 812 |
| | 299761999 | Nos1-CreERT2 | mca_classic | AAV | 870 |
| | 299782273 | Chrna2-Cre_OE | mca_classic | AAV | 206 |
| | 299782983 | Chrna2-Cre_OE | mca_classic | AAV | 232 |
| Y | 299783689 | Cux2-IRES-Cre | mca_classic | AAV | 279 |
| | 299784429 | Sim1-Cre_KJ18 | mca_classic | AAV | 86 |
| | 299820770 | Syt6-Cre_KI148 | mca_classic | AAV | 353 |
| | 299828473 | Syt6-Cre_KI148 | mca_classic | AAV | 192 |
| | 299829181 | Syt6-Cre_KI148 | mca_classic | AAV | 72 |
| | 299829892 | Syt6-Cre_KI148 | mca_classic | AAV | 232 |
| | 299856390 | Cck-IRES-Cre | mca_classic | AAV | 838 |
| | 299857096 | Chrna2-Cre_OE | mca_classic | AAV | 494 |
| | 299857813 | Lypd6-Cre_KL1! | mca_classic | AAV | 473 |
| | 299859225 | Plxnd1-Cre_OG | mca_classic | AAV | 185 |
| | 299894738 | Pvalb-IRES-Cre | mca_classic | AAV | 703 |
| | 299895444 | Pvalb-IRES-Cre | mca_classic | AAV | 822 |
| | 299896150 | Sim1-Cre_KJ18 | mca_classic | AAV | 600 |
| | 299897573 | Vip-IRES-Cre | mca_classic | AAV | 416 |
| | 299995638 | A930038C07Rik | mca_classic | AAV | 975 |
| | 299996344 | Calb2-IRES-Cre | mca_classic | AAV | 658 |
| | 300076066 | Dbh-Cre_KH212 | mca_classic | AAV | 838 |
| | 300078194 | Drd3-Cre_KI196 | mca_classic | AAV | 507 |
| | 300078901 | Drd3-Cre_KI196 | mca_classic | AAV | 575 |

| | | | | |
|---|---|---|---|---|
| 300109663 | Drd3-Cre_KI196 | mca_classic | AAV | 416 |
| 300111087 | Drd3-Cre_KI196 | mca_classic | AAV | 613 |
| 300111793 | Drd3-Cre_KI196 | mca_classic | AAV | 838 |
| 300164356 | Efr3a-Cre_NO1( | mca_classic | AAV | 588 |
| 300166697 | Gabrr3-Cre_KC: | mca_classic | AAV | 838 |
| 300167479 | Gad2-IRES-Cre | mca_classic | AAV | 757 |
| 300208016 | Gad2-IRES-Cre | mca_classic | AAV | 988 |
| 300209456 | Gal-Cre_KI87 | mca_classic | AAV | 977 |
| 300235349 | Grm2-Cre_MR9 | mca_classic | AAV | 24 |
| 300236056 | Grm2-Cre_MR9 | mca_classic | AAV | 712 |
| 300236763 | Htr2a-Cre_KM2 | mca_classic | AAV | 621 |
| 300237470 | Grik4-Cre | mca_classic | AAV | 644 |
| 300318924 | Htr2a-Cre_KM2 | mca_classic | AAV | 610 |
| 300319630 | Htr2a-Cre_KM2 | mca_classic | AAV | 494 |
| 300622004 | Nos1-CreERT2 | mca_classic | AAV | 1067 |
| 300623424 | Oxtr-Cre_ON66 | mca_classic | AAV | 527 |
| 300624130 | Rbp4-Cre_KL10 | mca_classic | AAV | 853 |
| 300641829 | Rbp4-Cre_KL10 | mca_classic | AAV | 421 |
| 300642574 | Rbp4-Cre_KL10 | mca_classic | AAV | 701 |
| 300644693 | Slc17a6-IRES-Cr | mca_classic | AAV | 380 |
| 300687607 | Slc17a6-IRES-Cr | mca_classic | AAV | 1003 |
| 300688721 | Slc18a2-Cre_OZ | mca_classic | AAV | 671 |
| 300689439 | Slc18a2-Cre_OZ | mca_classic | AAV | 925 |
| 300691325 | Slc6a5-Cre_KF1 | mca_classic | AAV | 913 |
| 300841699 | Sst-IRES-Cre | mca_classic | AAV | 588 |
| 300842406 | Sst-IRES-Cre | mca_classic | AAV | 1003 |
| 300843120 | Sst-IRES-Cre | mca_classic | AAV | 945 |
| 300843826 | Tac1-IRES2-Cre | mca_classic | AAV | 712 |
| 300888673 | Tac1-IRES2-Cre | mca_classic | AAV | 720 |
| 300889379 | Tac1-IRES2-Cre | mca_classic | AAV | 685 |
| 300923916 | Tac1-IRES2-Cre | mca_classic | AAV | 782 |
| 300927483 | Tac2-IRES2-Cre | mca_classic | AAV | 792 |
| 300929973 | Rorb-IRES2-Cre | mca_classic | AAV | 185 |
| 301016175 | Th-Cre_FI172 | mca_classic | AAV | 890 |
| 301016900 | Adcyap1-2A-Cre | mca_classic | AAV | 890 |
| 301057735 | Calb2-IRES-Cre | mca_classic | AAV | 713 |
| 301060890 | Calb2-IRES-Cre | mca_classic | AAV | 685 |
| 301061596 | Calb2-IRES-Cre | mca_classic | AAV | 776 |
| 301062306 | Calb2-IRES-Cre | mca_classic | AAV | 823 |
| 301063301 | Calb2-IRES-Cre | mca_classic | AAV | 702 |
| 301120618 | Drd3-Cre_KI198 | mca_classic | AAV | 467 |
| 301122593 | Drd3-Cre_KI198 | mca_classic | AAV | 473 |
| 301130077 | Drd3-Cre_KI198 | mca_classic | AAV | 457 |
| 301178953 | Efr3a-Cre_NO1( | mca_classic | AAV | 44 |
| 301179679 | Efr3a-Cre_NO1( | mca_classic | AAV | 232 |
| 301180385 | Efr3a-Cre_NO1( | mca_classic | AAV | 573 |
| 301209502 | Efr3a-Cre_NO1( | mca_classic | AAV | 692 |

| | | | | |
|---|---|---|---|---|
| 301210208 | Gal-Cre_KI87 | mca_classic | AAV | 890 |
| 301210923 | Gpr26-Cre_KO2 | mca_classic | AAV | 178 |
| 301264977 | Gpr26-Cre_KO2 | mca_classic | AAV | 494 |
| 301265683 | Gpr26-Cre_KO2 | mca_classic | AAV | 494 |
| 301323451 | Gpr26-Cre_KO2 | mca_classic | AAV | 272 |
| 301324157 | Grm2-Cre_MR9 | mca_classic | AAV | 1071 |
| 301324895 | Grp-Cre_KH288 | mca_classic | AAV | 685 |
| 301325601 | Nos1-CreERT2 | mca_classic | AAV | 457 |
| 301326316 | Nos1-CreERT2 | mca_classic | AAV | 467 |
| 301327022 | Nos1-CreERT2 | mca_classic | AAV | 473 |
| 301421253 | Oxtr-Cre_ON66 | mca_classic | AAV | 703 |
| 301421960 | Oxtr-Cre_ON66 | mca_classic | AAV | 467 |
| 301422682 | Oxtr-Cre_ON66 | mca_classic | AAV | 473 |
| 301464820 | Plxnd1-Cre_OG | mca_classic | AAV | 967 |
| 301466249 | Plxnd1-Cre_OG | mca_classic | AAV | 668 |
| 301538025 | Plxnd1-Cre_OG | mca_classic | AAV | 698 |
| 301539438 | Pvalb-IRES-Cre | mca_classic | AAV | 802 |
| 301540850 | Rbp4-Cre_KL10 | mca_classic | AAV | 838 |
| 301541824 | Rbp4-Cre_KL10 | mca_classic | AAV | 838 |
| 301580339 | Rorb-IRES2-Cre | mca_classic | AAV | 100 |
| 301581763 | Rorb-IRES2-Cre | mca_classic | AAV | 72 |
| 301583889 | Rorb-IRES2-Cre | mca_classic | AAV | 44 |
| 301616660 | Rorb-IRES2-Cre | mca_classic | AAV | 185 |
| 301617370 | Rorb-IRES2-Cre | mca_classic | AAV | 178 |
| 301618122 | Rorb-IRES2-Cre | mca_classic | AAV | 199 |
| 301618828 | Slc17a7-IRES2-C | mca_classic | AAV | 467 |
| 301620241 | Slc18a2-Cre_OZ | mca_classic | AAV | 573 |
| 301671287 | Slc18a2-Cre_OZ | mca_classic | AAV | 838 |
| 301672044 | Slc18a2-Cre_OZ | mca_classic | AAV | 587 |
| 301672755 | Slc18a2-Cre_OZ | mca_classic | AAV | 588 |
| 301673462 | Slc18a2-Cre_OZ | mca_classic | AAV | 757 |
| 301674988 | Slc18a2-Cre_OZ | mca_classic | AAV | 669 |
| 301732962 | Slc18a2-Cre_OZ | mca_classic | AAV | 869 |
| 301735080 | Slc18a2-Cre_OZ | mca_classic | AAV | 650 |
| 301735795 | Slc18a2-Cre_OZ | mca_classic | AAV | 662 |
| 301765327 | Calb2-IRES-Cre | mca_classic | AAV | 880 |
| 301800314 | Chat-IRES-Cre-n | mca_classic | AAV | 967 |
| 301875208 | Crh-IRES-Cre_ZJ | mca_classic | AAV | 975 |
| 301875966 | Gabrr3-Cre_KC | mca_classic | AAV | 912 |
| 301877386 | Grik4-Cre | mca_classic | AAV | 647 |
| 301947600 | Htr2a-Cre_KM2 | mca_classic | AAV | 621 |
| 301988879 | Htr2a-Cre_KM2 | mca_classic | AAV | 494 |
| 301989585 | Kiss1-Cre | mca_classic | AAV | 776 |
| 301991007 | Nxph4-T2A-Crel | mca_classic | AAV | 890 |
| 301991713 | Plxnd1-Cre_OG | mca_classic | AAV | 707 |
| 302014692 | Scnn1a-Tg2-Cre | mca_classic | AAV | 507 |
| 302016107 | Slc17a6-IRES-Cr | mca_classic | AAV | 978 |

| | | | | |
|---|---|---|---|---|
| 302016815 | Gad2-IRES-Cre | mca_classic | AAV | 795 |
| 302050617 | Gad2-IRES-Cre | mca_classic | AAV | 948 |
| 302053755 | Gad2-IRES-Cre | mca_classic | AAV | 838 |
| 302079862 | Gad2-IRES-Cre | mca_classic | AAV | 1033 |
| 302084009 | Slc17a6-IRES-Cr | mca_classic | AAV | 531 |
| 302085421 | Slc17a6-IRES-Cr | mca_classic | AAV | 985 |
| 302086846 | Gad2-IRES-Cre | mca_classic | AAV | 792 |
| 302087552 | Slc17a6-IRES-Cr | mca_classic | AAV | 792 |
| 302217570 | Gad2-IRES-Cre | mca_classic | AAV | 531 |
| 302221478 | Gad2-IRES-Cre | mca_classic | AAV | 720 |
| 302222934 | Gad2-IRES-Cre | mca_classic | AAV | 984 |
| 302723204 | Gad2-IRES-Cre | mca_classic | AAV | 826 |
| 302736304 | Gad2-IRES-Cre | mca_classic | AAV | 613 |
| 302739608 | Slc17a6-IRES-Cr | mca_classic | AAV | 613 |
| 302740314 | Gad2-IRES-Cre | mca_classic | AAV | 431 |
| 303468314 | Slc32a1-IRES-Cr | mca_classic | AAV | 984 |
| 303474463 | Gad2-IRES-Cre | mca_classic | AAV | 870 |
| 303478748 | Gad2-IRES-Cre | mca_classic | AAV | 573 |
| 303534443 | Gad2-IRES-Cre | mca_classic | AAV | 822 |
| 303535149 | Gad2-IRES-Cre | mca_classic | AAV | 890 |
| 303535867 | Gad2-IRES-Cre | mca_classic | AAV | 917 |
| 303536573 | Gad2-IRES-Cre | mca_classic | AAV | 1007 |
| 303537993 | Gad2-IRES-Cre | mca_classic | AAV | 589 |
| 303578324 | Slc17a6-IRES-Cr | mca_classic | AAV | 600 |
| 303580293 | Gad2-IRES-Cre | mca_classic | AAV | 617 |
| 303582161 | Tlx3-Cre_PL56 | mca_classic | AAV | 136 |
| 303614706 | Tlx3-Cre_PL56 | mca_classic | AAV | 72 |
| 303615412 | Tlx3-Cre_PL56 | mca_classic | AAV | 79 |
| 303616127 | Tlx3-Cre_PL56 | mca_classic | AAV | 353 |
| 303616833 | Tlx3-Cre_PL56 | mca_classic | AAV | 100 |
| 303617548 | Ntsr1-Cre_GN2: | mca_classic | AAV | 72 |
| 303708513 | Adcyap1-2A-Cre | mca_classic | AAV | 787 |
| 303709219 | Chrna2-Cre_OE | mca_classic | AAV | 1003 |
| 303710632 | Lypd6-Cre_KL15 | mca_classic | AAV | 698 |
| 303713313 | Rasgrf2-T2A-dC | mca_classic | AAV | 86 |
| 303783015 | Rasgrf2-T2A-dC | mca_classic | AAV | 232 |
| 303783725 | Rasgrf2-T2A-dC | mca_classic | AAV | 258 |
| 303785454 | Rbp4-Cre_KL10 | mca_classic | AAV | 72 |
| 304335875 | Sst-IRES-Cre | mca_classic | AAV | 51 |
| 304337288 | Th-Cre_FI172 | mca_classic | AAV | 823 |
| 304338014 | Rorb-IRES2-Cre | mca_classic | AAV | 808 |
| 304473503 | Rorb-IRES2-Cre | mca_classic | AAV | 751 |
| 304474221 | Slc17a6-IRES-Cr | mca_classic | AAV | 1098 |
| 304537794 | Slc32a1-IRES-Cr | mca_classic | AAV | 1098 |
| 304543821 | Slc17a6-IRES-Cr | mca_classic | AAV | 1033 |
| 304612686 | Nxph4-T2A-Cre | mca_classic | AAV | 973 |
| 304617742 | Slc17a6-IRES-Cr | mca_classic | AAV | 739 |

| | | | | |
|---|---|---|---|---|
| 304641170 | Ntsr1-Cre_GN2: | mca_classic | AAV | 86 |
| 304674547 | Slc32a1-IRES-Cr | mca_classic | AAV | 794 |
| 304675254 | Gad2-IRES-Cre | mca_classic | AAV | 1099 |
| 304675961 | Slc32a1-IRES-Cr | mca_classic | AAV | 1088 |
| 304694156 | Htr2a-Cre_KM2 | mca_classic | AAV | 527 |
| 304694870 | Nos1-CreERT2 | mca_classic | AAV | 600 |
| 304719312 | Chrna2-Cre_OE | mca_classic | AAV | 390 |
| 304720034 | Nos1-CreERT2 | mca_classic | AAV | 764 |
| 304720741 | Nxph4-T2A-Crel | mca_classic | AAV | 764 |
| 304721447 | Rbp4-Cre_KL10 | mca_classic | AAV | 569 |
| 304760810 | Sst-IRES-Cre | mca_classic | AAV | 621 |
| 304761539 | Th-Cre_FI172 | mca_classic | AAV | 826 |
| 304762245 | Chrna2-Cre_OE | mca_classic | AAV | 613 |
| 304857992 | Chrna2-Cre_OE | mca_classic | AAV | 812 |
| 304858700 | Drd3-Cre_KI196 | mca_classic | AAV | 802 |
| 304859407 | Drd3-Cre_KI196 | mca_classic | AAV | 870 |
| 304947087 | Gpr26-Cre_KO2 | mca_classic | AAV | 380 |
| 304947804 | Nos1-CreERT2 | mca_classic | AAV | 770 |
| 304948510 | Sst-IRES-Cre | mca_classic | AAV | 801 |
| 304949216 | Syt6-Cre_KI148 | mca_classic | AAV | 838 |
| 304969906 | Th-Cre_FI172 | mca_classic | AAV | 981 |
| 304970618 | Chrna2-Cre_OE | mca_classic | AAV | 595 |
| 304996627 | Gpr26-Cre_KO2 | mca_classic | AAV | 1099 |
| 304997333 | Nos1-CreERT2 | mca_classic | AAV | 1092 |
| 304998039 | Nos1-CreERT2 | mca_classic | AAV | 782 |
| 305024724 | Rbp4-Cre_KL10 | mca_classic | AAV | 611 |
| 305025440 | Nos1-CreERT2 | mca_classic | AAV | 830 |
| 305026146 | Gpr26-Cre_KO2 | mca_classic | AAV | 621 |
| 305026861 | Drd3-Cre_KI196 | mca_classic | AAV | 613 |
| 305071285 | Nos1-CreERT2 | mca_classic | AAV | 523 |
| 305092904 | Lypd6-Cre_KL15 | mca_classic | AAV | 757 |
| 305124396 | Gad2-IRES-Cre | mca_classic | AAV | 600 |
| 305125123 | Slc17a6-IRES-Cr | mca_classic | AAV | 693 |
| 305235787 | Gad2-IRES-Cre | mca_classic | AAV | 844 |
| 305270515 | Gad2-IRES-Cre | mca_classic | AAV | 776 |
| 305293222 | Gad2-IRES-Cre | mca_classic | AAV | 1062 |
| 305293928 | Slc32a1-IRES-Cr | mca_classic | AAV | 978 |
| 305294666 | Slc32a1-IRES-Cr | mca_classic | AAV | 416 |
| 305319461 | Slc32a1-IRES-Cr | mca_classic | AAV | 954 |
| 305320171 | Slc32a1-IRES-Cr | mca_classic | AAV | 473 |
| 305321177 | Slc32a1-IRES-Cr | mca_classic | AAV | 562 |
| 305321883 | Gad2-IRES-Cre | mca_classic | AAV | 802 |
| 305379705 | Slc32a1-IRES-Cr | mca_classic | AAV | 794 |
| 305380411 | Gad2-IRES-Cre | mca_classic | AAV | 457 |
| 305381127 | Slc32a1-IRES-Cr | mca_classic | AAV | 457 |
| 305400137 | Gad2-IRES-Cre | mca_classic | AAV | 473 |
| 305403845 | Gad2-IRES-Cre | mca_classic | AAV | 559 |

| | | | | |
|---|---|---|---|---|
| 305404551 | Slc17a6-IRES-Cr | mca_classic | AAV | 890 |
| 305425490 | Slc17a6-IRES-Cr | mca_classic | AAV | 682 |
| 305426208 | Slc17a6-IRES-Cr | mca_classic | AAV | 844 |
| 305446876 | Sst-IRES-Cre | mca_classic | AAV | 830 |
| 305447664 | Sst-IRES-Cre | mca_classic | AAV | 812 |
| 305449231 | Slc17a6-IRES-Cr | mca_classic | AAV | 693 |
| 305487234 | Htr3a-Cre_NO1 | mca_classic | AAV | 44 |
| 305487964 | Htr3a-Cre_NO1 | mca_classic | AAV | 24 |
| 305488671 | Htr3a-Cre_NO1 | mca_classic | AAV | 136 |
| 305489377 | Htr3a-Cre_NO1 | mca_classic | AAV | 185 |
| 305525773 | Htr3a-Cre_NO1 | mca_classic | AAV | 226 |
| 305618479 | Glt25d2-Cre_NI | mca_classic | AAV | 238 |
| 305645132 | Htr3a-Cre_NO1 | mca_classic | AAV | 600 |
| 305645843 | Htr3a-Cre_NO1 | mca_classic | AAV | 621 |
| 305677409 | Htr3a-Cre_NO1 | mca_classic | AAV | 757 |
| 305973112 | Penk-IRES2-Cre | mca_classic | AAV | 494 |
| 306268688 | Penk-IRES2-Cre | mca_classic | AAV | 44 |
| 306270474 | Th-Cre_FI172 | mca_classic | AAV | 802 |
| 306271212 | Erbb4-T2A-CreE | mca_classic | AAV | 739 |
| 306444486 | Cck-IRES-Cre | mca_classic | AAV | 645 |
| 306491185 | Adcyap1-2A-Cre | mca_classic | AAV | 332 |
| 306959448 | Slc18a2-Cre_OZ | mca_classic | AAV | 72 |
| 306990880 | Slc18a2-Cre_OZ | mca_classic | AAV | 79 |
| 307655867 | Adcyap1-2A-Cre | mca_classic | AAV | 764 |
| 307656580 | Ppp1r17-Cre_N | mca_classic | AAV | 830 |
| 307691605 | Ppp1r17-Cre_N | mca_classic | AAV | 830 |
| 307692311 | Chat-IRES-Cre-n | mca_classic | AAV | 573 |
| 307742524 | Nxph4-T2A-Crel | mca_classic | AAV | 1098 |
| 307743960 | Nxph4-T2A-Crel | mca_classic | AAV | 890 |
| 307766627 | Etv1-CreERT2 | mca_classic | AAV | 258 |
| 307909888 | Chat-IRES-Cre-n | mca_classic | AAV | 573 |
| 307910595 | Chat-IRES-Cre-n | mca_classic | AAV | 573 |
| 308027576 | Nos1-CreERT2 | mca_classic | AAV | 332 |
| 308385516 | Nos1-CreERT2 | mca_classic | AAV | 185 |
| 308386222 | Rbp4-Cre_KL10 | mca_classic | AAV | 621 |
| 308395312 | Ntrk1-IRES-Cre | mca_classic | AAV | 573 |
| 308549214 | Htr2a-Cre_KM2 | mca_classic | AAV | 559 |
| 308550233 | Ppp1r17-Cre_N | mca_classic | AAV | 473 |
| 308641549 | Ppp1r17-Cre_N | mca_classic | AAV | 787 |
| 308642254 | Pcdh9-Cre_NP2 | mca_classic | AAV | 507 |
| 308721177 | Pcdh9-Cre_NP2 | mca_classic | AAV | 523 |
| 308721884 | Trib2-F2A-CreEI | mca_classic | AAV | 332 |
| 308879859 | Trib2-F2A-CreEI | mca_classic | AAV | 185 |
| 308981706 | Gad2-IRES-Cre | mca_classic | AAV | 988 |
| 309385637 | Sst-IRES-Cre | mca_classic | AAV | 885 |
| 309386361 | Fezf1-T2A-dCre | mca_classic | AAV | 600 |
| 309427991 | Drd3-Cre_KI196 | mca_classic | AAV | 72 |

| | | | | |
|---|---|---|---|---|
| 309428697 | Drd3-Cre_KI196 | mca_classic | AAV | 79 |
| 309514434 | Drd3-Cre_KI196 | mca_classic | AAV | 44 |
| 309515141 | Efr3a-Cre_NO1( | mca_classic | AAV | 171 |
| 309564818 | Efr3a-Cre_NO1( | mca_classic | AAV | 178 |
| 309580102 | Slc32a1-IRES-Cr | mca_classic | AAV | 272 |
| 309580808 | Slc17a6-IRES-Cr | mca_classic | AAV | 701 |
| 309701202 | Slc17a6-IRES-Cr | mca_classic | AAV | 931 |
| 309702727 | Slc17a6-IRES-Cr | mca_classic | AAV | 682 |
| 309739641 | Ntrk1-IRES-Cre | mca_classic | AAV | 573 |
| 309740347 | Ntrk1-IRES-Cre | mca_classic | AAV | 575 |
| 309794438 | Ntrk1-IRES-Cre | mca_classic | AAV | 735 |
| 310175667 | Ntrk1-IRES-Cre | mca_classic | AAV | 573 |
| 310176384 | Calb2-IRES-Cre | mca_classic | AAV | 880 |
| 310193233 | Slc17a6-IRES-Cr | mca_classic | AAV | 644 |
| 310194040 | Cux2-CreERT2 | mca_classic | AAV | 18 |
| 310377782 | Sst-IRES-Cre | mca_classic | AAV | 934 |
| 310436191 | Cux2-CreERT2 | mca_classic | AAV | 416 |
| 310437091 | Cux2-CreERT2 | mca_classic | AAV | 561 |
| 310439724 | Cux2-CreERT2 | mca_classic | AAV | 416 |
| 310695955 | Crh-IRES-Cre_ZJ | mca_classic | AAV | 925 |
| 310853453 | Rasgrf2-T2A-dC | mca_classic | AAV | 948 |
| 310976160 | Rasgrf2-T2A-dC | mca_classic | AAV | 694 |
| 311845265 | Htr3a-Cre_NO1 | mca_classic | AAV | 854 |
| 311845972 | Htr3a-Cre_NO1 | mca_classic | AAV | 890 |
| 311846681 | Th-Cre_FI172 | mca_classic | AAV | 954 |
| 312240825 | Slc17a6-IRES-Cr | mca_classic | AAV | 649 |
| 313141786 | Rasgrf2-T2A-dC | mca_classic | AAV | 890 |
| 313324664 | Rasgrf2-T2A-dC | mca_classic | AAV | 692 |
| 313325371 | Rasgrf2-T2A-dC | mca_classic | AAV | 787 |
| 313327028 | Glt25d2-Cre_NI | mca_classic | AAV | 279 |
| 313327735 | Glt25d2-Cre_NI | mca_classic | AAV | 258 |
| 477037203 | Sepw1-Cre_NP3 | mca_classic | AAV | 24 |
| 477269754 | Sepw1-Cre_NP3 | mca_classic | AAV | 185 |
| 477271169 | Sepw1-Cre_NP3 | mca_classic | AAV | 44 |
| 477475894 | Ctgf-T2A-dgCre | mca_classic | AAV | 185 |
| 477834984 | Sepw1-Cre_NP3 | mca_classic | AAV | 122 |
| 477835774 | Sepw1-Cre_NP3 | mca_classic | AAV | 24 |
| 477836675 | Calb1-T2A-dgCr | mca_classic | AAV | 18 |
| 477924853 | Penk-IRES2-Cre | mca_classic | AAV | 573 |
| 477926293 | Dlg3-Cre_KG118 | mca_classic | AAV | 467 |
| 477927130 | Rasgrf2-T2A-dC | mca_classic | AAV | 823 |
| 478095541 | Rasgrf2-T2A-dC | mca_classic | AAV | 787 |
| 478096249 | Rasgrf2-T2A-dC | mca_classic | AAV | 822 |
| 478097069 | Rasgrf2-T2A-dC | mca_classic | AAV | 822 |
| 478097779 | Ctgf-T2A-dgCre | mca_classic | AAV | 122 |
| 478780588 | Sepw1-Cre_NP3 | mca_classic | AAV | 232 |
| 478782005 | Sepw1-Cre_NP3 | mca_classic | AAV | 346 |

| | | | | | |
|---|---|---|---|---|---|
| | 478818791 | Dlg3-Cre_KG11ā | mca_classic | AAV | 918 |
| | 478819500 | Dlg3-Cre_KG11ā | mca_classic | AAV | 918 |
| | 485022109 | Calb1-T2A-dgCr | mca_classic | AAV | 51 |
| | 485022815 | Calb1-T2A-dgCr | mca_classic | AAV | 136 |
| | 485105033 | Dlg3-Cre_KG11ā | mca_classic | AAV | 985 |
| | 485105742 | Dlg3-Cre_KG11ā | mca_classic | AAV | 562 |
| | 485239207 | Rasgrf2-T2A-dC | mca_classic | AAV | 794 |
| | 485846989 | Ntng2-IRES2-Cr | mca_classic | AAV | 573 |
| | 485847695 | Ntng2-IRES2-Cr | mca_classic | AAV | 285 |
| | 485875903 | Gnb4-IRES2-Cre | mca_classic | AAV | 416 |
| | 485903475 | Gnb4-IRES2-Cre | mca_classic | AAV | 557 |
| | 506117918 | Gpr26-Cre_KO2 | mca_classic | AAV | 416 |
| | 506426038 | Glt25d2-Cre_NI | mca_classic | AAV | 494 |
| | 508357710 | Rorb-IRES2-Cre | mca_classic | AAV | 808 |
| | 508373923 | Sim1-Cre_KJ18 | mca_classic | AAV | 114 |
| | 508539001 | Sim1-Cre_KJ18 | mca_classic | AAV | 802 |
| | 508775240 | Sim1-Cre_KJ18 | mca_classic | AAV | 566 |
| | 509463587 | Htr2a-Cre_KM2 | mca_classic | AAV | 473 |
| | 509602066 | Htr2a-Cre_KM2 | mca_classic | AAV | 531 |
| | 509880387 | Rorb-IRES2-Cre | mca_classic | AAV | 808 |
| | 510124187 | Tac2-IRES2-Cre | mca_classic | AAV | 713 |
| | 510125001 | Tac1-IRES2-Cre | mca_classic | AAV | 713 |
| | 511234217 | Sepw1-Cre_NP3 | mca_classic | AAV | 185 |
| | 511234957 | Sepw1-Cre_NP3 | mca_classic | AAV | 51 |
| | 511549277 | Scnn1a-Tg2-Cre | mca_classic | AAV | 802 |
| | 511942270 | Pvalb-IRES-Cre | mca_classic | AAV | 610 |
| | 511971714 | Th-Cre_FI172 | mca_classic | AAV | 890 |
| | 512130198 | Npr3-IRES2-Cre | mca_classic | AAV | 18 |
| | 512130913 | Npr3-IRES2-Cre | mca_classic | AAV | 44 |
| Y | 513773998 | Ntng2-IRES2-Cr | mca_classic | AAV | 291 |
| | 513775257 | Ntng2-IRES2-Cr | mca_classic | AAV | 573 |
| | 514505957 | Gnb4-IRES2-Cre | mca_classic | AAV | 573 |
| | 514506712 | Gnb4-IRES2-Cre | mca_classic | AAV | 291 |
| | 514513838 | Gal-Cre_KI87 | mca_classic | AAV | 662 |
| | 515198413 | Sim1-Cre_KJ18 | mca_classic | AAV | 757 |
| | 515410820 | Sim1-Cre_KJ18 | mca_classic | AAV | 794 |
| | 515418047 | Slc6a5-Cre_KF1 | mca_classic | AAV | 913 |
| | 516278420 | Chrnb4-Cre_OL | mca_classic | AAV | 494 |
| | 516491813 | Tlx3-Cre_PL56 | mca_classic | AAV | 171 |
| | 516529585 | Chrnb4-Cre_OL | mca_classic | AAV | 535 |
| | 516545903 | Drd3-Cre_KI196 | mca_classic | AAV | 44 |
| | 516838033 | Drd3-Cre_KI196 | mca_classic | AAV | 332 |
| | 516846757 | Cux2-IRES-Cre | mca_classic | AAV | 561 |
| | 516848198 | Drd3-Cre_KI196 | mca_classic | AAV | 226 |
| | 518207061 | Drd3-Cre_KI196 | mca_classic | AAV | 494 |
| | 518745077 | Gpr26-Cre_KO2 | mca_classic | AAV | 416 |
| | 518745840 | Grp-Cre_KH288 | mca_classic | AAV | 507 |

| | | | | |
|---|---|---|---|---|
| 519164644 | Hdc-Cre_IM1 | mca_classic | AAV | 770 |
| 520336173 | Hdc-Cre_IM1 | mca_classic | AAV | 775 |
| 520342605 | Hdc-Cre_IM1 | mca_classic | AAV | 781 |
| 520619072 | Hdc-Cre_IM1 | mca_classic | AAV | 763 |
| 520728084 | Sim1-Cre_KJ18 | mca_classic | AAV | 100 |
| 521264566 | Drd3-Cre_KI196 | mca_classic | AAV | 332 |
| 521402511 | Drd3-Cre_KI196 | mca_classic | AAV | 353 |
| 521600217 | Sepw1-Cre_NP3 | mca_classic | AAV | 258 |
| 522078446 | Prkcd-GluCla-CF | mca_classic | AAV | 621 |
| 522774776 | Ctgf-T2A-dgCre | mca_classic | AAV | 51 |
| 523178542 | Rasgrf2-T2A-dC | mca_classic | AAV | 830 |
| 523705737 | Rasgrf2-T2A-dC | mca_classic | AAV | 757 |
| 524070052 | Ctgf-T2A-dgCre | mca_classic | AAV | 24 |
| 524268045 | Calb1-T2A-dgCr | mca_classic | AAV | 24 |
| 525796603 | Calb1-T2A-dgCr | mca_classic | AAV | 707 |
| 526502961 | Sepw1-Cre_NP3 | mca_classic | AAV | 332 |
| 527051458 | Slc17a6-IRES-Cr | mca_classic | AAV | 535 |
| 527393818 | Slc17a6-IRES-Cr | mca_classic | AAV | 507 |
| 527577056 | Drd3-Cre_KI196 | mca_classic | AAV | 213 |
| 527808181 | Ndnf-IRES2-dgC | mca_classic | AAV | 587 |
| 530008580 | Drd3-Cre_KI196 | mca_classic | AAV | 178 |
| 530018580 | Drd3-Cre_KI196 | mca_classic | AAV | 178 |
| 538078619 | Drd3-Cre_KI196 | mca_classic | AAV | 332 |
| 538833505 | Calb1-T2A-dgCr | mca_classic | AAV | 694 |
| 538834292 | Calb1-T2A-dgCr | mca_classic | AAV | 645 |
| 539010002 | Lypd6-Cre_KL15 | mca_classic | AAV | 929 |
| 539028976 | Tlx3-Cre_PL56 | mca_classic | AAV | 890 |
| 540685246 | Pdyn-T2A-CreEF | mca_classic | AAV | 720 |
| 543875354 | Pdyn-T2A-CreEF | mca_classic | AAV | 595 |
| 543876073 | Pdyn-T2A-CreEF | mca_classic | AAV | 890 |
| 543880631 | Kcng4-Cre | mca_classic | AAV | 838 |
| 543881679 | Kcng4-Cre | mca_classic | AAV | 853 |
| 544455391 | Kcng4-Cre | mca_classic | AAV | 800 |
| 545415593 | Npr3-IRES2-Cre | mca_classic | AAV | 569 |
| 545427588 | Npr3-IRES2-Cre | mca_classic | AAV | 890 |
| 545428296 | Slc17a8-IRES2-C | mca_classic | AAV | 693 |
| 545434921 | Tlx3-Cre_PL56 | mca_classic | AAV | 238 |
| 547509190 | Slc17a8-IRES2-C | mca_classic | AAV | 930 |
| 547510030 | Slc17a8-IRES2-C | mca_classic | AAV | 881 |
| 549203025 | Oxtr-T2A-Cre | mca_classic | AAV | 457 |
| 549361039 | Oxtr-T2A-Cre | mca_classic | AAV | 600 |
| 549362997 | Oxtr-T2A-Cre | mca_classic | AAV | 613 |
| 549805072 | Cart-IRES2-Cre | mca_classic | AAV | 600 |
| 550892495 | Trib2-F2A-CreEI | mca_classic | AAV | 535 |
| 551350026 | Adcyap1-2A-Cre | mca_classic | AAV | 416 |
| 551351756 | Htr2a-Cre_KM2 | mca_classic | AAV | 713 |
| 551738231 | Adcyap1-2A-Cre | mca_classic | AAV | 792 |

| | | | | |
|---|---|---|---|---|
| 552280478 | Htr2a-Cre_KM2 | mca_classic | AAV | 649 |
| 552283801 | Htr2a-Cre_KM2 | mca_classic | AAV | 1099 |
| 552284594 | Adcyap1-2A-Cre | mca_classic | AAV | 890 |
| 552759734 | Adcyap1-2A-Cre | mca_classic | AAV | 787 |
| 553079356 | Ctgf-T2A-dgCre | mca_classic | AAV | 232 |
| 553446684 | Gnb4-IRES2-Cre | mca_classic | AAV | 100 |
| 554021622 | Tacr1-T2A-Cre | mca_classic | AAV | 617 |
| 554022330 | Tacr1-T2A-Cre | mca_classic | AAV | 595 |
| 554023317 | Tacr1-T2A-Cre | mca_classic | AAV | 852 |
| 554650901 | Cart-IRES2-Cre | mca_classic | AAV | 416 |
| 554651619 | Cart-IRES2-Cre | mca_classic | AAV | 703 |
| 555011865 | Cart-IRES2-Cre | mca_classic | AAV | 792 |
| 555739999 | Slc17a8-IRES2-C | mca_classic | AAV | 948 |
| 556341848 | Tacr1-T2A-Cre | mca_classic | AAV | 1007 |
| 557199437 | Ntng2-IRES2-Cr | mca_classic | AAV | 507 |
| 557200148 | Ntng2-IRES2-Cr | mca_classic | AAV | 178 |
| 557972433 | Nos1-CreERT2 | mca_classic | AAV | 398 |
| 557973149 | Pdyn-T2A-CreEI | mca_classic | AAV | 792 |
| 558579356 | Adcyap1-2A-Cre | mca_classic | AAV | 945 |
| 558580065 | C57Bl/6J | mca_classic | AAV | 507 |
| 558673113 | Adcyap1-2A-Cre | mca_classic | AAV | 764 |
| 558675974 | Adcyap1-2A-Cre | mca_classic | AAV | 1003 |
| 558697990 | Foxp2-IRES-Cre | mca_classic | AAV | 685 |
| 561916513 | Htr1a-IRES2-Cre | mca_classic | AAV | 494 |
| 562963492 | Slc17a8-IRES2-C | mca_classic | AAV | 925 |
| 564356750 | Slc17a8-IRES2-C | mca_classic | AAV | 185 |
| 564357489 | Esr2-IRES2-Cre | mca_classic | AAV | 600 |
| 564685751 | Htr1a-IRES2-Cre | mca_classic | AAV | 890 |
| 564688610 | Foxp2-IRES-Cre | mca_classic | AAV | 800 |
| 565721603 | Tacr1-T2A-Cre | mca_classic | AAV | 890 |
| 569264117 | Slc17a8-IRES2-C | mca_classic | AAV | 24 |
| 573639461 | Foxp2-IRES-Cre | mca_classic | AAV | 595 |
| 574363940 | Nos1-CreERT2 | mca_classic | AAV | 889 |
| 574572418 | Nos1-CreERT2 | mca_classic | AAV | 947 |
| 574950390 | Npr3-IRES2-Cre | mca_classic | AAV | 185 |
| 575216368 | Npr3-IRES2-Cre | mca_classic | AAV | 24 |
| 575772121 | Penk-IRES2-Cre | mca_classic | AAV | 185 |
| 581327676 | Gnb4-IRES2-Cre | mca_classic | AAV | 100 |
| 581328778 | Nos1-CreERT2 | mca_classic | AAV | 826 |
| 581350498 | Nos1-CreERT2 | mca_classic | AAV | 770 |
| 581641279 | Nos1-CreERT2 | mca_classic | AAV | 720 |
| 581642026 | Nos1-CreERT2 | mca_classic | AAV | 948 |
| 582609848 | Esr2-IRES2-Cre | mca_classic | AAV | 881 |
| 582814018 | Glt25d2-Cre_NI | mca_classic | AAV | 232 |
| 583290390 | Grik4-Cre | mca_classic | AAV | 507 |
| 583749274 | Syt6-Cre_KI148 | mca_classic | AAV | 24 |
| 584513749 | Syt6-Cre_KI148 | mca_classic | AAV | 24 |

| | | | | |
|---|---|---|---|---|
| 584902900 | Tlx3-Cre_PL56 | mca_classic | AAV | 65 |
| 585026021 | C57Bl/6J | mca_classic | AAV | 507 |
| 585051446 | C57Bl/6J | mca_classic | AAV | 507 |
| 585756865 | Ntng2-IRES2-Cr | mca_classic | AAV | 494 |
| 585775993 | Grik4-Cre | mca_classic | AAV | 507 |
| 585910380 | Penk-IRES2-Cre | mca_classic | AAV | 185 |
| 585911240 | Npr3-IRES2-Cre | mca_classic | AAV | 232 |
| 585931259 | Penk-IRES2-Cre | mca_classic | AAV | 185 |
| 585935640 | Npr3-IRES2-Cre | mca_classic | AAV | 258 |
| 586041882 | Npr3-IRES2-Cre | mca_classic | AAV | 346 |
| 586042591 | Npr3-IRES2-Cre | mca_classic | AAV | 332 |
| 586052938 | Npr3-IRES2-Cre | mca_classic | AAV | 86 |
| 586054033 | Penk-IRES2-Cre | mca_classic | AAV | 185 |
| 586054741 | Adcyap1-2A-Cre | mca_classic | AAV | 702 |
| 586377476 | Adcyap1-2A-Cre | mca_classic | AAV | 954 |
| 586447435 | Pomc-Cre_BL | mca_classic | AAV | 733 |
| 586448498 | Gabrr3-Cre_KC | mca_classic | AAV | 912 |
| 587060515 | Ppp1r17-Cre_N | mca_classic | AAV | 776 |
| 587294457 | Avp-IRES2-Cre | mca_classic | AAV | 717 |
| 591612976 | Tlx3-Cre_PL56 | mca_classic | AAV | 18 |
| 591622344 | Tlx3-Cre_PL56 | mca_classic | AAV | 65 |
| 592391705 | Npr3-IRES2-Cre | mca_classic | AAV | 226 |
| 592513871 | Npr3-IRES2-Cre | mca_classic | AAV | 122 |
| 596571282 | Oxtr-T2A-Cre | mca_classic | AAV | 185 |
| 596798450 | Sim1-Cre_KJ18 | mca_classic | AAV | 24 |
| 596828910 | Oxtr-T2A-Cre | mca_classic | AAV | 185 |
| 597007858 | Sim1-Cre_KJ18 | mca_classic | AAV | 18 |
| 598604288 | Rbp4-Cre_KL10 | mca_classic | AAV | 24 |
| 605657236 | Ctgf-T2A-dgCre | mca_classic | AAV | 185 |
| 605660419 | Ctgf-T2A-dgCre | mca_classic | AAV | 192 |
| 606250170 | Sim1-Cre_KJ18 | mca_classic | AAV | 18 |
| 606570335 | Sim1-Cre_KJ18 | mca_classic | AAV | 24 |
| 606785720 | Rbp4-Cre_KL10 | mca_classic | AAV | 24 |
| 614436425 | Slc17a8-iCre | mca_classic | AAV | 24 |
| 614735393 | Tlx3-Cre_PL56 | mca_classic | AAV | 24 |
| 623855077 | Rorb-IRES2-Cre | mca_classic | AAV | 185 |
| 626107114 | C57Bl/6J | mca_classic | AAV | 507 |
| 626118851 | Ctgf-T2A-dgCre | mca_classic | AAV | 185 |
| 627869431 | Ctgf-T2A-dgCre | mca_classic | AAV | 185 |
| 637365535 | Pvalb-T2A-CreE | mca_classic | AAV | 185 |
| 638313933 | Pvalb-T2A-CreE | mca_classic | AAV | 185 |
| 638978767 | Penk-IRES2-Cre | mca_classic | AAV | 185 |
| 640081415 | C57Bl/6J | mca_classic | AAV | 494 |
| 640282128 | C57Bl/6J | mca_classic | AAV | 507 |
| 640285199 | C57Bl/6J | mca_classic | AAV | 535 |
| 642177206 | Scnn1a-Tg2-Cre | mca_classic | AAV | 662 |
| 642180077 | Vipr2-IRES2-Cre | mca_classic | AAV | 662 |

| | | | | |
|---|---|---|---|---|
| 642480973 | Vipr2-IRES2-Cre | mca_classic | AAV | 662 |
| 642811309 | Scnn1a-Tg2-Cre | mca_classic | AAV | 662 |
| 642967852 | C57Bl/6J | mca_classic | AAV | 494 |
| 648048922 | Rorb-IRES2-Cre- | mca_classic | AAV | 185 |
| 648051857 | Rorb-IRES2-Cre- | mca_classic | AAV | 185 |
| 648318728 | Slc17a8-iCre | mca_classic | AAV | 185 |
| 656657344 | Slc17a8-IRES2-( | mca_classic | AAV | 185 |
| 656842599 | Rorb-IRES2-Cre | mca_classic | AAV | 185 |
| 670657874 | Penk-IRES2-Cre- | mca_classic | AAV | 185 |
| 671030719 | Penk-IRES2-Cre- | mca_classic | AAV | 185 |
| 671033786 | Penk-IRES2-Cre- | mca_classic | AAV | 185 |
| 672056591 | Slc17a8-IRES2-( | mca_classic | AAV | 185 |
| 672516002 | Slc17a8-IRES2-( | mca_classic | AAV | 185 |
| 674524470 | Rorb-IRES2-Cre | mca_classic | AAV | 185 |
| 503035875 | Sst-IRES-Cre | T504 Connectivity M | AAV | 566 |
| 503036583 | Oxtr-T2A-Cre | T504 Connectivity M | AAV | 787 |
| 503324388 | Nkx2-1-CreERT2 | T504 Connectivity M | AAV | 787 |
| 505790715 | Tlx3-Cre_PL56 | T504 Connectivity M | AAV | 24 |
| 507798246 | Sim1-Cre_KJ18 | T504 Connectivity M | AAV | 24 |
| 508273699 | Sim1-Cre_KJ18 | T504 Connectivity M | AAV | 24 |
| 513498584 | Sst-IRES-Cre | T504 Connectivity M | AAV | 595 |
| 515520455 | Oxtr-T2A-Cre | T504 Connectivity M | AAV | 781 |
| 517082898 | Tlx3-Cre_PL56 | T504 Connectivity M | AAV | 24 |
| 517314004 | Tlx3-Cre_PL56 | T504 Connectivity M | AAV | 24 |
| 518015408 | Nkx2-1-CreERT2 | T504 Connectivity M | AAV | 739 |
| 518023283 | Cux2-IRES-Cre | T504 Connectivity M | AAV | 24 |
| 518218546 | Ntsr1-Cre_GN2 | T504 Connectivity M | AAV | 24 |
| 518222748 | Sim1-Cre_KJ18 | T504 Connectivity M | AAV | 24 |
| 522255595 | Sst-IRES-Cre | T504 Connectivity M | AAV | 802 |
| 539498984 | Sst-IRES-Cre | T504 Connectivity M | AAV | 611 |
| 539641136 | Sst-IRES-Cre | T504 Connectivity M | AAV | 595 |
| 540139629 | Nkx2-1-CreERT2 | T504 Connectivity M | AAV | 787 |
| 540140513 | Tlx3-Cre_PL56 | T504 Connectivity M | AAV | 24 |
| 547202505 | Foxp2-IRES-Cre | T504 Connectivity M | AAV | 801 |
| 551756337 | Foxp2-IRES-Cre | T504 Connectivity M | AAV | 801 |
| 552757477 | Syt6-Cre_KI148 | T504 Connectivity M | AAV | 24 |
| 552758186 | Syt6-Cre_KI148 | T504 Connectivity M | AAV | 24 |
| 552973699 | Sst-IRES-Cre | T504 Connectivity M | AAV | 703 |
| 602828622 | Penk-IRES2-Cre- | T504 Connectivity M | AAV | 24 |
| 603211567 | Penk-IRES2-Cre- | T504 Connectivity M | AAV | 24 |
| 603331422 | Penk-IRES2-Cre- | T504 Connectivity M | AAV | 24 |
| 603574321 | Rorb-IRES2-Cre | T504 Connectivity M | AAV | 24 |
| 603891334 | Rorb-IRES2-Cre | T504 Connectivity M | AAV | 24 |
| 614737677 | Glt25d2-Cre_NF | T504 Connectivity M | AAV | 24 |
| 615543017 | Rorb-IRES2-Cre- | T504 Connectivity M | AAV | 24 |
| 616676552 | Sepw1-Cre_NP3 | T504 Connectivity M | AAV | 24 |
| 616856399 | Rorb-IRES2-Cre- | T504 Connectivity M | AAV | 24 |

| | | | | | |
|---|---|---|---|---|---|
| | 616857128 | Glt25d2-Cre_NF | T504 Connectivity N | AAV | 24 |
| | 636803027 | Pvalb-T2A-CreE | T504 Connectivity N | AAV | 24 |
| | 636967182 | Pvalb-T2A-CreE | T504 Connectivity N | AAV | 24 |
| | 639813139 | Penk-IRES2-Cre- | T504 Connectivity N | AAV | 24 |
| | 647576768 | Rorb-IRES2-Cre- | T504 Connectivity N | AAV | 24 |
| | 647807416 | Slc17a8-iCre | T504 Connectivity N | AAV | 24 |
| | 671463542 | Penk-IRES2-Cre- | T504 Connectivity N | AAV | 24 |
| | 671464291 | Penk-IRES2-Cre- | T504 Connectivity N | AAV | 24 |
| | 673383135 | Sepw1-Cre_NP3 | T504 Connectivity N | AAV | 24 |
| | 673386769 | Rorb-IRES2-Cre | T504 Connectivity N | AAV | 24 |
| | 673668737 | Rorb-IRES2-Cre- | T504 Connectivity N | AAV | 24 |
| | 679157574 | Rorb-IRES2-Cre | T504 Connectivity N | AAV | 24 |
| | 478256439 | A930038C07Rik | T601.3a anterograd | AAV | 185 |
| | 478257959 | Ntsr1-Cre_GN2 | T601.3a anterograd | AAV | 185 |
| | 478491090 | Emx1-IRES-Cre | T601.3a anterograd | AAV | 226 |
| | 478491810 | Chat-IRES-Cre-n | T601.3a anterograd | AAV | 613 |
| | 478581080 | Prkcd-GluCla-CF | T601.3a anterograd | AAV | 649 |
| | 478582494 | Emx1-IRES-Cre | T601.3a anterograd | AAV | 264 |
| | 479085232 | Scnn1a-Tg3-Cre | T601.3a anterograd | AAV | 185 |
| | 479267539 | Prkcd-GluCla-CF | T601.3a anterograd | AAV | 675 |
| | 479673174 | Emx1-IRES-Cre | T601.3a anterograd | AAV | 178 |
| | 479673887 | Emx1-IRES-Cre | T601.3a anterograd | AAV | 178 |
| | 479700629 | Emx1-IRES-Cre | T601.3a anterograd | AAV | 178 |
| | 479701339 | Emx1-IRES-Cre | T601.3a anterograd | AAV | 185 |
| | 479755622 | Emx1-IRES-Cre | T601.3a anterograd | AAV | 185 |
| | 479756361 | Emx1-IRES-Cre | T601.3a anterograd | AAV | 185 |
| | 479891303 | Scnn1a-Tg2-Cre | T601.3a anterograd | AAV | 662 |
| | 480074702 | Slc6a4-Cre_ET3 | T601.3a anterograd | AAV | 881 |
| | 480689656 | Dbh-Cre_KH212 | T601.3a anterograd | AAV | 1003 |
| | 480692170 | Slc17a6-IRES-Cr | T601.3a anterograd | AAV | 662 |
| | 480703321 | Slc17a6-IRES-Cr | T601.3a anterograd | AAV | 685 |
| | 482578964 | Rbp4-Cre_KL10 | T601.3a anterograd | AAV | 185 |
| | 483013787 | Cux2-IRES-Cre | T601.3a anterograd | AAV | 185 |
| | 485237081 | Rbp4-Cre_KL10 | T601.3a anterograd | AAV | 199 |
| | 495344543 | Cux2-IRES-Cre | T601.3a anterograd | AAV | 178 |
| Y | 495345251 | Cux2-IRES-Cre | T601.3a anterograd | AAV | 199 |
| | 495562600 | Rbp4-Cre_KL10 | T601.3a anterograd | AAV | 178 |
| | 496554237 | Rbp4-Cre_KL10 | T601.3a anterograd | AAV | 264 |
| | 496576666 | Rbp4-Cre_KL10 | T601.3a anterograd | AAV | 226 |
| | 502966396 | Emx1-IRES-Cre | T601.3a anterograd | AAV | 185 |
| | 503813164 | Scnn1a-Tg3-Cre | T601.3a anterograd | AAV | 199 |
| | 504519805 | Chat-IRES-Cre-n | T601.3a anterograd | AAV | 613 |
| | 504521113 | Chat-IRES-Cre-n | T601.3a anterograd | AAV | 613 |
| | 507708083 | Prkcd-GluCla-CF | T601.3a anterograd | AAV | 668 |
| | 512314723 | Emx1-IRES-Cre | T601.3a anterograd | AAV | 199 |
| | 514333422 | Slc17a6-IRES-Cr | T601.3a anterograd | AAV | 676 |
| | 517975511 | Chat-IRES-Cre-n | T601.3a anterograd | AAV | 795 |

| | | | |
|---|---|---|---|
| 518605900 | Emx1-IRES-Cre | T601.3a anterograd AAV | 164 |
| 518606617 | Emx1-IRES-Cre | T601.3a anterograd AAV | 353 |
| 518742338 | Emx1-IRES-Cre | T601.3a anterograd AAV | 171 |
| 519186737 | Tlx3-Cre_PL56 | T601.3a anterograd AAV | 199 |
| 520012330 | A930038C07Rik | T601.3a anterograd AAV | 346 |
| 520019334 | A930038C07Rik | T601.3a anterograd AAV | 185 |
| 520341802 | A930038C07Rik | T601.3a anterograd AAV | 178 |
| 520750912 | Tlx3-Cre_PL56 | T601.3a anterograd AAV | 178 |
| 520985971 | A930038C07Rik | T601.3a anterograd AAV | 185 |
| 520996382 | Cux2-IRES-Cre | T601.3a anterograd AAV | 164 |
| 522409371 | Emx1-IRES-Cre | T601.3a anterograd AAV | 185 |
| 522635991 | Tlx3-Cre_PL56 | T601.3a anterograd AAV | 353 |
| 522636747 | Tlx3-Cre_PL56 | T601.3a anterograd AAV | 185 |
| 522773270 | Cux2-IRES-Cre | T601.3a anterograd AAV | 272 |
| 522774029 | Cux2-IRES-Cre | T601.3a anterograd AAV | 226 |
| 523714193 | Ntsr1-Cre_GN2 | T601.3a anterograd AAV | 185 |
| 523718075 | Rbp4-Cre_KL10 | T601.3a anterograd AAV | 325 |
| 524656772 | Cux2-IRES-Cre | T601.3a anterograd AAV | 185 |
| 524666904 | Rbp4-Cre_KL10 | T601.3a anterograd AAV | 164 |
| 524667618 | Rbp4-Cre_KL10 | T601.3a anterograd AAV | 171 |
| 526924964 | A930038C07Rik | T601.3a anterograd AAV | 185 |
| 528509838 | Tlx3-Cre_PL56 | T601.3a anterograd AAV | 164 |
| 528510546 | Tlx3-Cre_PL56 | T601.3a anterograd AAV | 171 |
| 528963283 | Ntsr1-Cre_GN2 | T601.3a anterograd AAV | 171 |
| 528963991 | Tlx3-Cre_PL56 | T601.3a anterograd AAV | 185 |
| 529692491 | Tlx3-Cre_PL56 | T601.3a anterograd AAV | 185 |
| 530574594 | Rbp4-Cre_KL10 | T601.3a anterograd AAV | 185 |
| 531441947 | Tlx3-Cre_PL56 | T601.3a anterograd AAV | 185 |
| 535692871 | Emx1-IRES-Cre | T601.3a anterograd AAV | 185 |
| 536298009 | A930038C07Rik | T601.3a anterograd AAV | 185 |
| 536298726 | Emx1-IRES-Cre | T601.3a anterograd AAV | 185 |
| 536299435 | Emx1-IRES-Cre | T601.3a anterograd AAV | 185 |
| 538079359 | Ntsr1-Cre_GN2 | T601.3a anterograd AAV | 51 |
| 539521918 | Cux2-IRES-Cre | T601.3a anterograd AAV | 226 |
| 539738599 | Ntsr1-Cre_GN2 | T601.3a anterograd AAV | 185 |
| 539739321 | Ntsr1-Cre_GN2 | T601.3a anterograd AAV | 185 |
| 540145406 | Rbp4-Cre_KL10 | T601.3a anterograd AAV | 353 |
| 540146149 | Tlx3-Cre_PL56 | T601.3a anterograd AAV | 185 |
| 543679575 | Ntsr1-Cre_GN2 | T601.3a anterograd AAV | 185 |
| 543680289 | Ntsr1-Cre_GN2 | T601.3a anterograd AAV | 325 |
| 544488252 | Ntsr1-Cre_GN2 | T601.3a anterograd AAV | 185 |
| 544488964 | Ntsr1-Cre_GN2 | T601.3a anterograd AAV | 164 |
| 552279683 | Ntsr1-Cre_GN2 | T601.3a anterograd AAV | 346 |
| 552430870 | A930038C07Rik | T601.3a anterograd AAV | 178 |
| 553092069 | Ntsr1-Cre_GN2 | T601.3a anterograd AAV | 136 |
| 553747363 | Slc17a6-IRES-Cr | T601.3a anterograd AAV | 685 |
| 554333581 | A930038C07Rik | T601.3a anterograd AAV | 298 |

| | | | |
|---|---|---|---|
| 554334685 | Rbp4-Cre_KL10 | T601.3a anterograd AAV | 213 |
| 554420321 | Tlx3-Cre_PL56 | T601.3a anterograd AAV | 206 |
| 554421791 | Emx1-IRES-Cre | T601.3a anterograd AAV | 143 |
| 555745687 | Tlx3-Cre_PL56 | T601.3a anterograd AAV | 192 |
| 556922099 | Cux2-IRES-Cre | T601.3a anterograd AAV | 143 |
| 557826519 | Ntsr1-Cre_GN2 | T601.3a anterograd AAV | 51 |
| 562061175 | Emx1-IRES-Cre | T601.3a anterograd AAV | 185 |
| 562520963 | Emx1-IRES-Cre | T601.3a anterograd AAV | 178 |
| 562521707 | Emx1-IRES-Cre | T601.3a anterograd AAV | 325 |
| 562671482 | Emx1-IRES-Cre | T601.3a anterograd AAV | 136 |
| 562674923 | Emx1-IRES-Cre | T601.3a anterograd AAV | 298 |
| 566244185 | Cux2-IRES-Cre | T601.3a anterograd AAV | 171 |
| 566454054 | Cux2-IRES-Cre | T601.3a anterograd AAV | 353 |
| 566730846 | Cux2-IRES-Cre | T601.3a anterograd AAV | 143 |
| 566992832 | Rbp4-Cre_KL10 | T601.3a anterograd AAV | 185 |
| 568768472 | Sim1-Cre_KJ18 | T601.3a anterograd AAV | 794 |
| 569994739 | Cux2-IRES-Cre | T601.3a anterograd AAV | 136 |
| 570460301 | Cux2-IRES-Cre | T601.3a anterograd AAV | 199 |
| 571823196 | Cux2-IRES-Cre | T601.3a anterograd AAV | 232 |
| 572388249 | Prkcd-GluCla-CF | T601.3a anterograd AAV | 668 |
| 572588941 | Emx1-IRES-Cre | T601.3a anterograd AAV | 199 |
| 573035760 | Prkcd-GluCla-CF | T601.3a anterograd AAV | 676 |
| 573624241 | Slc17a6-IRES-Cr | T601.3a anterograd AAV | 792 |
| 576332845 | Emx1-IRES-Cre | T601.3a anterograd AAV | 136 |
| 576660608 | A930038C07Rik | T601.3a anterograd AAV | 353 |
| 576684233 | A930038C07Rik | T601.3a anterograd AAV | 164 |
| 577773267 | Cux2-IRES-Cre | T601.3a anterograd AAV | 100 |
| 583320505 | A930038C07Rik | T601.3a anterograd AAV | 812 |
| 583748537 | Emx1-IRES-Cre | T601.3a anterograd AAV | 51 |
| 584194481 | Emx1-IRES-Cre | T601.3a anterograd AAV | 226 |
| 584511119 | A930038C07Rik | T601.3a anterograd AAV | 185 |
| 584895127 | Emx1-IRES-Cre | T601.3a anterograd AAV | 325 |
| 584896065 | A930038C07Rik | T601.3a anterograd AAV | 143 |
| 585021827 | Ntsr1-Cre_GN2 | T601.3a anterograd AAV | 298 |
| 585760423 | Ntsr1-Cre_GN2 | T601.3a anterograd AAV | 298 |
| 590988082 | Emx1-IRES-Cre | T601.3a anterograd AAV | 213 |
| 591223633 | Tlx3-Cre_PL56 | T601.3a anterograd AAV | 325 |
| 591535205 | Emx1-IRES-Cre | T601.3a anterograd AAV | 332 |
| 593016678 | Emx1-IRES-Cre | T601.3a anterograd AAV | 185 |
| 593018150 | Rbp4-Cre_KL10 | T601.3a anterograd AAV | 298 |
| 593277684 | Rbp4-Cre_KL10 | T601.3a anterograd AAV | 192 |
| 593407064 | Ntsr1-Cre_GN2 | T601.3a anterograd AAV | 213 |
| 595884140 | Tlx3-Cre_PL56 | T601.3a anterograd AAV | 298 |
| 596252304 | Tlx3-Cre_PL56 | T601.3a anterograd AAV | 185 |
| 602094339 | Sim1-Cre_KJ18 | T601.3a anterograd AAV | 185 |
| 602166541 | Sim1-Cre_KJ18 | T601.3a anterograd AAV | 185 |
| 603208501 | Scnn1a-Tg3-Cre | T601.3a anterograd AAV | 164 |

| | | | |
|---|---|---|---|
| 603890605 | Ntsr1-Cre_GN2: | T601.3a anterograd AAV | 360 |
| 605661910 | Tlx3-Cre_PL56 | T601.3a anterograd AAV | 171 |
| 606100558 | Rbp4-Cre_KL10 | T601.3a anterograd AAV | 143 |
| 636803957 | Scnn1a-Tg3-Cre | T601.3a anterograd AAV | 199 |
| 642970691 | A930038C07Rik | T601.3a anterograd AAV | 192 |
| 643166866 | A930038C07Rik | T601.3a anterograd AAV | 192 |
| 643749624 | A930038C07Rik | T601.3a anterograd AAV | 51 |
| 646214807 | Tlx3-Cre_PL56 | T601.3a anterograd AAV | 213 |
| 646216224 | A930038C07Rik | T601.3a anterograd AAV | 213 |
| 646525156 | Rbp4-Cre_KL10 | T601.3a anterograd AAV | 298 |
| 647574144 | Ntrk1-IRES-Cre | T601.3a anterograd AAV | 610 |
| 647806688 | Rbp4-Cre_KL10 | T601.3a anterograd AAV | 51 |
| 648048130 | Tlx3-Cre_PL56 | T601.3a anterograd AAV | 213 |
| 648050335 | Rbp4-Cre_KL10 | T601.3a anterograd AAV | 213 |
| 648253235 | Rbp4-Cre_KL10 | T601.3a anterograd AAV | 51 |
| 649183036 | Ntsr1-Cre_GN2: | T601.3a anterograd AAV | 213 |
| 649361916 | Ntsr1-Cre_GN2: | T601.3a anterograd AAV | 199 |
| 651041703 | Tlx3-Cre_PL56 | T601.3a anterograd AAV | 346 |
| 656688345 | Gnb4-IRES2-Cre | T601.3a anterograd AAV | 44 |
| 657042668 | Emx1-IRES-Cre | T601.3a anterograd AAV | 185 |
| 657162589 | Scnn1a-Tg3-Cre | T601.3a anterograd AAV | 178 |
| 666909423 | Scnn1a-Tg3-Cre | T601.3a anterograd AAV | 185 |
| 666936463 | Scnn1a-Tg3-Cre | T601.3a anterograd AAV | 171 |
| 667294910 | Scnn1a-Tg3-Cre | T601.3a anterograd AAV | 353 |
| 680910929 | Scnn1a-Tg3-Cre | T601.3a anterograd AAV | 136 |
| 681573722 | Scnn1a-Tg3-Cre | T601.3a anterograd AAV | 206 |
| 681881558 | Scnn1a-Tg3-Cre | T601.3a anterograd AAV | 298 |
| 690735403 | Tlx3-Cre_PL56 | T601.3a anterograd AAV | 360 |
| 694306983 | Scnn1a-Tg3-Cre | T601.3a anterograd AAV | 346 |
| 694812317 | Scnn1a-Tg3-Cre | T601.3a anterograd AAV | 213 |
| 695974330 | Scnn1a-Tg3-Cre | T601.3a anterograd AAV | 199 |
| 696461404 | Scnn1a-Tg3-Cre | T601.3a anterograd AAV | 51 |
| 478257195 | A930038C07Rik | T601.3b anterograd AAV | 185 |
| 478258719 | Prkcd-GluCla-CF | T601.3b anterograd AAV | 662 |
| 478376197 | Emx1-IRES-Cre | T601.3b anterograd AAV | 238 |
| 478376911 | Emx1-IRES-Cre | T601.3b anterograd AAV | 226 |
| 478490368 | Chat-IRES-Cre-n | T601.3b anterograd AAV | 613 |
| 478581786 | Chat-IRES-Cre-n | T601.3b anterograd AAV | 613 |
| 479268685 | Prkcd-GluCla-CF | T601.3b anterograd AAV | 662 |
| 479670988 | Slc17a6-IRES-Cr | T601.3b anterograd AAV | 662 |
| 479671695 | Scnn1a-Tg2-Cre | T601.3b anterograd AAV | 662 |
| 479980810 | Emx1-IRES-Cre | T601.3b anterograd AAV | 178 |
| 479981981 | Emx1-IRES-Cre | T601.3b anterograd AAV | 185 |
| 479982715 | Emx1-IRES-Cre | T601.3b anterograd AAV | 185 |
| 479983421 | Emx1-IRES-Cre | T601.3b anterograd AAV | 185 |
| 479984127 | Emx1-IRES-Cre | T601.3b anterograd AAV | 185 |
| 480069939 | Emx1-IRES-Cre | T601.3b anterograd AAV | 185 |

| | | | | |
|---|---|---|---|---|
| | 480994108 | Rbp4-Cre_KL10 | T601.3b anterograd AAV | 272 |
| | 482580380 | Rbp4-Cre_KL10 | T601.3b anterograd AAV | 178 |
| Y | 482581199 | Rbp4-Cre_KL10 | T601.3b anterograd AAV | 325 |
| | 483014695 | Cux2-IRES-Cre | T601.3b anterograd AAV | 185 |
| | 484503464 | Cux2-IRES-Cre | T601.3b anterograd AAV | 178 |
| | 484504171 | Slc17a6-IRES-Cr | T601.3b anterograd AAV | 685 |
| | 495345959 | Cux2-IRES-Cre | T601.3b anterograd AAV | 298 |
| | 495753765 | Ntsr1-Cre_GN2 | T601.3b anterograd AAV | 185 |
| | 495754475 | Ntsr1-Cre_GN2 | T601.3b anterograd AAV | 178 |
| | 496097187 | Ntsr1-Cre_GN2 | T601.3b anterograd AAV | 199 |
| | 496097913 | Cux2-IRES-Cre | T601.3b anterograd AAV | 245 |
| | 496113850 | Rbp4-Cre_KL10 | T601.3b anterograd AAV | 332 |
| | 496114558 | Dbh-Cre_KH212 | T601.3b anterograd AAV | 838 |
| | 496150781 | Cux2-IRES-Cre | T601.3b anterograd AAV | 24 |
| | 496152279 | A930038C07Rik | T601.3b anterograd AAV | 185 |
| | 496252297 | A930038C07Rik | T601.3b anterograd AAV | 185 |
| | 500836105 | Emx1-IRES-Cre | T601.3b anterograd AAV | 178 |
| | 500836840 | Emx1-IRES-Cre | T601.3b anterograd AAV | 185 |
| | 500837552 | Emx1-IRES-Cre | T601.3b anterograd AAV | 185 |
| | 501004474 | Emx1-IRES-Cre | T601.3b anterograd AAV | 185 |
| | 501006221 | Emx1-IRES-Cre | T601.3b anterograd AAV | 199 |
| | 501115762 | Emx1-IRES-Cre | T601.3b anterograd AAV | 185 |
| | 501116471 | Cux2-IRES-Cre | T601.3b anterograd AAV | 185 |
| | 501117182 | Cux2-IRES-Cre | T601.3b anterograd AAV | 185 |
| | 501483880 | Emx1-IRES-Cre | T601.3b anterograd AAV | 226 |
| | 501484658 | Emx1-IRES-Cre | T601.3b anterograd AAV | 332 |
| | 502074651 | Emx1-IRES-Cre | T601.3b anterograd AAV | 199 |
| | 502140653 | Tlx3-Cre_PL56 | T601.3b anterograd AAV | 185 |
| | 502180994 | Tlx3-Cre_PL56 | T601.3b anterograd AAV | 199 |
| | 503017948 | Tlx3-Cre_PL56 | T601.3b anterograd AAV | 185 |
| | 503018656 | Tlx3-Cre_PL56 | T601.3b anterograd AAV | 178 |
| | 503069254 | Rbp4-Cre_KL10 | T601.3b anterograd AAV | 185 |
| | 503809372 | Rbp4-Cre_KL10 | T601.3b anterograd AAV | 185 |
| | 504099316 | Scnn1a-Tg3-Cre | T601.3b anterograd AAV | 185 |
| | 504100025 | Prkcd-GluCla-CF | T601.3b anterograd AAV | 668 |
| | 504173156 | Chat-IRES-Cre-n | T601.3b anterograd AAV | 618 |
| | 506890448 | Cux2-IRES-Cre | T601.3b anterograd AAV | 185 |
| | 506891241 | Cux2-IRES-Cre | T601.3b anterograd AAV | 185 |
| | 506947040 | Prkcd-GluCla-CF | T601.3b anterograd AAV | 676 |
| | 508889322 | A930038C07Rik | T601.3b anterograd AAV | 185 |
| | 510581751 | Tlx3-Cre_PL56 | T601.3b anterograd AAV | 185 |
| | 510582887 | Tlx3-Cre_PL56 | T601.3b anterograd AAV | 185 |
| | 510834706 | Rbp4-Cre_KL10 | T601.3b anterograd AAV | 164 |
| | 510848595 | Tlx3-Cre_PL56 | T601.3b anterograd AAV | 164 |
| | 511817919 | Ntsr1-Cre_GN2 | T601.3b anterograd AAV | 185 |
| | 512315551 | Emx1-IRES-Cre | T601.3b anterograd AAV | 185 |
| | 514313871 | Ntsr1-Cre_GN2 | T601.3b anterograd AAV | 185 |

| | | | | |
|---|---|---|---|---|
| | 514314707 | Ntsr1-Cre_GN2: | T601.3b anterograd AAV | 185 |
| | 514332492 | Ntsr1-Cre_GN2: | T601.3b anterograd AAV | 185 |
| | 516273371 | Ntsr1-Cre_GN2: | T601.3b anterograd AAV | 325 |
| | 516274127 | Ntsr1-Cre_GN2: | T601.3b anterograd AAV | 164 |
| | 517072832 | Rbp4-Cre_KL10 | T601.3b anterograd AAV | 185 |
| | 517078399 | Cux2-IRES-Cre | T601.3b anterograd AAV | 206 |
| | 517325325 | Cux2-IRES-Cre | T601.3b anterograd AAV | 353 |
| | 517326050 | Cux2-IRES-Cre | T601.3b anterograd AAV | 171 |
| | 517974408 | Ntsr1-Cre_GN2: | T601.3b anterograd AAV | 51 |
| | 518619451 | Tlx3-Cre_PL56 | T601.3b anterograd AAV | 171 |
| | 518741580 | Tlx3-Cre_PL56 | T601.3b anterograd AAV | 353 |
| | 519487627 | A930038C07Rik | T601.3b anterograd AAV | 164 |
| | 519735413 | Dbh-Cre_KH212 | T601.3b anterograd AAV | 929 |
| | 520018181 | A930038C07Rik | T601.3b anterograd AAV | 325 |
| | 521600943 | Emx1-IRES-Cre | T601.3b anterograd AAV | 353 |
| | 522408611 | Tlx3-Cre_PL56 | T601.3b anterograd AAV | 185 |
| | 523714940 | Ntsr1-Cre_GN2: | T601.3b anterograd AAV | 171 |
| Y | 523718823 | Rbp4-Cre_KL10 | T601.3b anterograd AAV | 199 |
| | 526783054 | Rbp4-Cre_KL10 | T601.3b anterograd AAV | 226 |
| | 534175823 | A930038C07Rik | T601.3b anterograd AAV | 164 |
| | 546103149 | Emx1-IRES-Cre | T601.3b anterograd AAV | 185 |
| | 552544756 | Cux2-IRES-Cre | T601.3b anterograd AAV | 213 |
| | 553743594 | Ntsr1-Cre_GN2: | T601.3b anterograd AAV | 192 |
| | 553746532 | Ntsr1-Cre_GN2: | T601.3b anterograd AAV | 298 |
| | 557187751 | Cux2-IRES-Cre | T601.3b anterograd AAV | 298 |
| | 557341233 | Ntsr1-Cre_GN2: | T601.3b anterograd AAV | 143 |
| | 557342452 | Ntsr1-Cre_GN2: | T601.3b anterograd AAV | 199 |
| | 557347149 | Ntsr1-Cre_GN2: | T601.3b anterograd AAV | 346 |
| | 557827228 | Ntsr1-Cre_GN2: | T601.3b anterograd AAV | 298 |
| | 559878074 | Rbp4-Cre_KL10 | T601.3b anterograd AAV | 171 |
| | 560799305 | Ntsr1-Cre_GN2: | T601.3b anterograd AAV | 185 |
| | 560800029 | Ntsr1-Cre_GN2: | T601.3b anterograd AAV | 325 |
| | 563179067 | Emx1-IRES-Cre | T601.3b anterograd AAV | 143 |
| | 571100135 | Emx1-IRES-Cre | T601.3b anterograd AAV | 171 |
| | 572770444 | Tlx3-Cre_PL56 | T601.3b anterograd AAV | 178 |
| | 572969377 | Tlx3-Cre_PL56 | T601.3b anterograd AAV | 122 |
| | 573330828 | Slc17a6-IRES-Cr | T601.3b anterograd AAV | 787 |
| | 574941472 | Emx1-IRES-Cre | T601.3b anterograd AAV | 185 |
| | 575782182 | Emx1-IRES-Cre | T601.3b anterograd AAV | 298 |
| | 576036240 | Cux2-IRES-Cre | T601.3b anterograd AAV | 298 |
| | 576340889 | Rbp4-Cre_KL10 | T601.3b anterograd AAV | 213 |
| | 576341623 | Rbp4-Cre_KL10 | T601.3b anterograd AAV | 298 |
| | 579203888 | Emx1-IRES-Cre | T601.3b anterograd AAV | 143 |
| | 579628905 | A930038C07Rik | T601.3b anterograd AAV | 185 |
| | 580150240 | Emx1-IRES-Cre | T601.3b anterograd AAV | 185 |
| | 583747816 | Emx1-IRES-Cre | T601.3b anterograd AAV | 226 |
| | 584507699 | A930038C07Rik | T601.3b anterograd AAV | 185 |

| | | | | | |
|---|---|---|---|---|---|
| | 584511827 | Emx1-IRES-Cre | T601.3b anterograd | AAV | 185 |
| | 587345518 | Cux2-IRES-Cre | T601.3b anterograd | AAV | 143 |
| | 590548119 | Tlx3-Cre_PL56 | T601.3b anterograd | AAV | 171 |
| | 590983389 | Tlx3-Cre_PL56 | T601.3b anterograd | AAV | 213 |
| | 590987294 | Tlx3-Cre_PL56 | T601.3b anterograd | AAV | 199 |
| | 591224520 | Tlx3-Cre_PL56 | T601.3b anterograd | AAV | 192 |
| | 591233395 | Tlx3-Cre_PL56 | T601.3b anterograd | AAV | 325 |
| | 592392441 | Cux2-IRES-Cre | T601.3b anterograd | AAV | 213 |
| | 592540591 | Emx1-IRES-Cre | T601.3b anterograd | AAV | 332 |
| | 593296997 | Ntrk1-IRES-Cre | T601.3b anterograd | AAV | 613 |
| | 595858299 | Tlx3-Cre_PL56 | T601.3b anterograd | AAV | 206 |
| | 597007143 | Rbp4-Cre_KL10 | T601.3b anterograd | AAV | 346 |
| | 598605738 | Emx1-IRES-Cre | T601.3b anterograd | AAV | 185 |
| | 599136295 | Ntsr1-Cre_GN2 | T601.3b anterograd | AAV | 298 |
| | 601268292 | Ntsr1-Cre_GN2 | T601.3b anterograd | AAV | 51 |
| | 601476074 | Emx1-IRES-Cre | T601.3b anterograd | AAV | 185 |
| | 603468246 | Emx1-IRES-Cre | T601.3b anterograd | AAV | 507 |
| | 603717226 | Ntsr1-Cre_GN2 | T601.3b anterograd | AAV | 51 |
| | 606930364 | A930038C07Rik | T601.3b anterograd | AAV | 171 |
| | 642809043 | Rbp4-Cre_KL10 | T601.3b anterograd | AAV | 143 |
| | 644250774 | A930038C07Rik | T601.3b anterograd | AAV | 51 |
| | 648252402 | Rbp4-Cre_KL10 | T601.3b anterograd | AAV | 213 |
| | 649181539 | Slc6a4-Cre_ET3 | T601.3b anterograd | AAV | 881 |
| | 649484051 | Ntsr1-Cre_GN2 | T601.3b anterograd | AAV | 213 |
| | 650408717 | A930038C07Rik | T601.3b anterograd | AAV | 206 |
| | 651702588 | Tlx3-Cre_PL56 | T601.3b anterograd | AAV | 346 |
| | 652753758 | Tlx3-Cre_PL56 | T601.3b anterograd | AAV | 213 |
| | 653191449 | Cux2-IRES-Cre | T601.3b anterograd | AAV | 346 |
| | 656839070 | Gnb4-IRES2-Cre | T601.3b anterograd | AAV | 285 |
| | 656959718 | Rbp4-Cre_KL10 | T601.3b anterograd | AAV | 346 |
| | 657041814 | Emx1-IRES-Cre | T601.3b anterograd | AAV | 51 |
| | 657172447 | Emx1-IRES-Cre | T601.3b anterograd | AAV | 213 |
| | 657334568 | Emx1-IRES-Cre | T601.3b anterograd | AAV | 353 |
| | 670228985 | Tlx3-Cre_PL56 | T601.3b anterograd | AAV | 51 |
| | 594021642 | Gnb4-IRES2-Cre | trans-synaptic | rabies | 573 |
| | 653020645 | Gnb4-IRES2-Cre | trans-synaptic | rabies | 573 |
| Y, currently failed in LIMS | 717571579 | Glt25d2-Cre_NF | trans-synaptic | rabies | 557 |
| Y | 730766377 | Rbp4-Cre_KL10 | trans-synaptic | rabies | 557 |
| | 766654105 | Gnb4-IRES2-Cre | trans-synaptic | rabies | 416 |
| | 767100712 | Gnb4-IRES2-Cre | trans-synaptic | rabies | 100 |
| | 767101425 | Gnb4-IRES2-Cre | trans-synaptic | rabies | 559 |
| | 780505913 | Gnb4-IRES2-Cre | trans-synaptic | rabies | 559 |
| | 780508099 | Gnb4-IRES2-Cre | trans-synaptic | rabies | 557 |
| | 780510390 | Gnb4-IRES2-Cre | trans-synaptic | rabies | 557 |
| | 863469869 | Ntng2-IRES2-Cr | trans-synaptic | rabies | 285 |
| | 865328287 | Ntng2-IRES2-Cr | trans-synaptic | rabies | 100 |
| | 907255727 | Gnb4-IRES2-Cre | trans-synaptic | rabies | 291 |

| | | | | |
|---|---|---|---|---|
| 908029900 | Gnb4-IRES2-Cre | trans-synaptic | rabies | 291 |

| primary-injection-struc | secondary-injection-structures |
|---|---|
| DG | CA3 |
| MOB | AON |
| VPM | VAL\|VPLpc\|VPMpc\|PO |
| PH | VM\|VPMpc\|SPFm\|MD\|SMT\|PR\|PCN\| |
| LSr | CP\|LSc\|LSv\|SF |
| IC | |
| MS | LSr\|LSv\|SF\|MEPO |
| LGd | LP |
| SNr | SNc |
| AV | VAL\|AM\|LD\|RT |
| CP | |
| PO | DG\|LP\|POL\|PF\|MRN\|APN |
| CA1 | SSp-ll\|SSp-ul\|SSp-tr\|CA2\|CA3 |
| DG | CA1\|CA3 |
| CA1 | SUB\|ProS |
| PAR | ENTm |
| DG | |
| TU | ARH\|PVp\|TMv\|PMv\|VMH |
| PIR | EPd |
| CP | GPe |
| CA3 | CA1\|CA2 |
| LHA | RT\|PVHd\|PeF\|ZI |
| CUL | DEC |
| PH | VM\|SPFm\|ZI |
| CP | |
| CEA | LA\|BLA |
| SIM | CUL |
| ACAv | MOs\|ACAd |
| AId | MOp\|GU\|CLA |
| DP | ILA\|ORBm\|TT |
| DG | CA3 |
| ENTm | ENTl\|PAR\|PRE\|SUB |
| CA3 | |
| DG | |
| VCO | FL |
| SPVO | NTS\|SPVI\|ISN\|PARN |
| SCs | SCm |
| AN | |
| PVH | AHN\|PVHd |
| TU | VMH |
| ZI | VM\|VPLpc\|VPM\|PoT\|SPFp |
| PAG | MRN\|SCm |
| BLA | CP\|CEA |
| SGN | PoT\|MG\|LP\|POL\|PIL\|NB\|MRN |
| CA3 | DG |
| LHA | DMH\|PeF |

| | |
|---|---|
| ENTm | ENTl |
| LHA | DMH\|AHN\|PVHd\|VMH\|PeF\|ZI |
| DMH | PVi\|VMH\|PH |
| IC | |
| SUB | RSPv\|CA1\|FC\|ProS |
| LHA | AHN\|VMH\|PeF\|TU |
| MRN | SCm |
| MRN | VTA\|RR\|PAG\|RN\|MA3\|EW\|IF\|RL\|CLI |
| OP | MPT\|NOT\|NPC\|PPT |
| AON | PIR\|EPd |
| LHA | DMH\|AHN\|PVHd\|PH\|PST\|PSTN\|PeF |
| RE | PR\|Xi\|PVHd |
| CP | PIR\|ACB\|FS |
| LHA | AHN\|PVHd\|PST\|PeF\|ZI |
| LPO | SI\|NDB\|BST\|AVP\|MPO\|VLPO\|LHA |
| LD | AD |
| MPO | NDB\|BST\|AVP\|PS\|VLPO\|AHN\|MPN\|LI |
| MS | LSr\|LSv\|SF |
| CP | SSp-n\|SSp-ul\|SSp-un |
| VII | IC\|IRN\|PARN |
| DG | |
| LSv | ACB\|MS\|BST\|ADP\|PS |
| LD | CA3\|LP |
| APN | VISp\|VISpm\|DG\|PoT\|LP\|POL\|SGN\|NB |
| VAL | VPL\|VPM\|PO\|RT |
| SIM | CUL |
| DG | CA3 |
| POST | VISp |
| CA3 | DG |
| BMA | COAa\|COAp\|BLA\|PA\|CEA\|IA\|MEA |
| SIM | CUL |
| DG | CA1\|SUB\|ProS |
| DG | CA3 |
| SUM | MM\|PMd\|PH |
| LHA | MOp\|SSp-ll\|VM\|RT\|PVHd\|PST\|PSTN\| |
| DG | CA3 |
| DR | PAG\|IV\|CLI |
| PAG | SCm\|IV\|DR |
| DR | PAG |
| COAp | |
| VMH | ARH\|DMH\|PMv |
| MD | AM\|IAM\|IAD\|IMD\|PVT\|PT\|CM\|PCN |
| DG | CA3\|LP |
| GR | |
| SPVC | Pa5 |
| AV | AD |
| ACAd | MOs\|ACAv\|PL |

| | |
|---|---|
| CA3 | CA1\|CA2 |
| DG | CA3 |
| ENTl | PERI\|CA1 |
| MPN | AVPV\|MPO\|PVpo\|SBPV\|VMPO\|AHN |
| CUL | |
| MRN | VISpm\|IC\|NB\|SCm |
| ACAv | MOs\|ACAd |
| AUDd | SSs\|AUDp |
| SCs | RSPv\|SCm |
| AON | TT\|ACB\|OT\|SI |
| PSV | PB\|SUT\|I5\|VCO\|CUL |
| SSp-bfd | |
| BMA | COAa\|COAp\|BLA\|PA\|IA\|MEA |
| CA1 | VISam\|RSPagl\|SUB\|ProS |
| MD | PCN\|CL\|LH |
| IC | SCm\|PAG |
| LHA | CEA\|MEA\|GPi\|SI\|RT\|ZI |
| BST | LSv\|BAC |
| RT | GPe\|GPi\|VPL |
| NLL | SAG\|PBG\|CUN\|PB |
| MPO | LSv\|MS\|BST\|ADP\|AVP\|AVPV\|MEPO\|[illegible] |
| PFL | CUL\|SIM\|AN\|COPY |
| CENT | |
| MPN | ACAd\|ACAv\|LSc\|BST\|MPO\|PD\|PVpo\|[illegible] |
| PAR | VISp |
| CEA | |
| BLA | LA\|CP\|CEA\|GPe |
| CA1 | |
| LSc | ACAv\|LSr |
| PFL | AN\|PRM\|COPY |
| IP | |
| SSs | AUDd |
| CA1 | |
| PAG | MRN\|CUN |
| VTA | SUM\|PH\|PN\|MRN\|PAG\|MA3\|EW\|IF\| |
| MOs | |
| MRN | PAG\|RN |
| PAG | SCm |
| RN | VTA\|MRN |
| AOB | ORBl\|ORBvl\|MOB\|AON |
| MOs | MOp |
| NOD | LING\|CUL\|UVU\|FN |
| PVT | MD\|MH\|LH\|PAG |
| SSp-n | SSp-bfd\|SSp-un |
| SSs | SSp-bfd\|AUDd |
| TRN | PG\|PRNc\|PRNr |
| GRN | MARN\|RM |

| | |
|---|---|
| MV | SPIV |
| MRN | SCs\|SCm\|PAG\|RN\|III\|IV\|Pa4\|RL\|CLI\|[illegible] |
| AON | MOB |
| MDRN | SPVC\|IRN\|MDRNd |
| PAA | PIR\|COAa\|EPv\|BMA |
| CA1 | CA2\|CA3\|DG |
| MOs | MOp |
| PT | LSv\|SF\|MS\|TRS\|BST\|AM\|IAD\|PVT\|RE |
| VII | PB\|V\|P5\|SPVO\|IRN\|PARN\|LAV\|MV\|S |
| SSp-n | SSp-bfd\|SSp-un |
| CP | MOs |
| AUDp | AUDd |
| MEA | BA\|SI\|LHA |
| APN | PF\|MRN\|NPC |
| VISp | |
| SI | CP\|ACB\|OT\|MA |
| PRNc | IC\|MRN\|PAG\|PCG\|LDT\|PRNr\|SLC\|SL[illegible] |
| GRN | IRN\|MARN\|PGRNl |
| ACAv | |
| PRM | |
| PYR | FOTU |
| BMA | COAp\|BLA\|PA\|IA |
| ACAd | MOs\|ACAv |
| APr | VISp\|RSPd\|POST |
| AHN | BST\|MPO\|SBPV\|MPN\|LHA\|LPO\|PeF |
| ENTl | |
| ACAv | ACAd |
| IRN | GRN\|LIN\|PARN\|XII |
| PARN | SPVI\|SPVO\|AMB\|IRN\|LIN\|PGRNl |
| ACAv | |
| ZI | VM\|VPLpc\|LHA\|PST\|PSTN\|STN |
| SPIV | LAV\|MV |
| MRN | PAG\|CUN\|PPN |
| SPVI | SPVC\|Pa5 |
| ORBl | |
| CUN | MRN\|SCm\|PAG\|PPN\|PB |
| SCm | MRN\|PAG\|CUN |
| SUB | RSPv\|ProS\|SCm\|APN\|NOT\|OP\|PPT |
| SCm | SCs |
| TR | |
| IRN | NTS\|SPVI\|GRN\|LIN\|PARN\|SPIV |
| GRN | IRN\|LIN\|LRN\|MARN\|PGRNl |
| MM | LM\|SUM\|TMv\|PMd\|PMv\|PH\|LHA\|VT |
| SI | GPi |
| SI | MA\|LPO |
| PT | IAD\|MD\|PVT |
| GRN | IRN\|LIN\|PGRNl |

| | |
|---|---|
| CP | AIv\|CLA\|EPd |
| LA | VISC\|AIp\|ECT\|EPd\|CP |
| SPVC | CU\|SPVI\|MDRN\|MDRNd |
| IRN | CU\|NTS\|GRN\|LIN\|LRN\|MDRN\|MDRN[illegible] |
| GRN | IO\|IRN\|LRN\|MARN\|MDRN\|MDRNv\|P[illegible] |
| PGRNl | SPVC\|VII\|GRN\|IRN\|LRN\|MARN\|MDR[illegible] |
| APN | SCm |
| RT | |
| PB | MEV\|PAG\|B\|PCG\|LC\|LDT\|CENT |
| RT | VAL\|VPL\|VPM\|PO |
| SAG | VISp\|IC\|NB\|PBG\|MRN\|CUN |
| ENTm | ENTl\|PAR\|PRE\|SUB |
| CP | |
| PRNr | MRN\|SCm\|PAG\|PPN\|PRNc |
| CP | |
| ACB | LSr\|LSv |
| PVT | CA2\|CA3\|DG\|IMD\|MD\|MH\|LH |
| PGRNl | GRN\|LRN\|MARN |
| XII | NTS\|DMX\|IRN\|MDRN\|MDRNd\|MDRN |
| IRN | AMB\|LRN\|MDRN\|MDRNd\|MDRNv\|P[illegible] |
| PIL | DG\|PoT\|SPFp\|PP\|MG\|LP\|POL\|SGN\|[illegible] |
| MM | LM\|SUM\|PH\|VTA\|MRN |
| ENTl | PERI\|ECT\|PIR |
| CENT | IC |
| RT | GPe\|GPi |
| PB | CUN\|PPN |
| LHA | VM\|LM\|MM\|SUM\|PMd\|PH\|PSTN\|ZI |
| PVH | BST\|RE |
| MRN | SAG\|SNr\|RR\|SNc |
| ORBvl | FRP\|ORBl |
| AHN | RT\|ASO\|SBPV\|PVHd\|VMH\|LHA\|PeF\|[illegible] |
| FN | CUL |
| VTA | RR\|IF\|RL\|CLI |
| PH | RSPd\|RSPv\|DMH\|MM\|SUM\|TMd\|PM |
| PAR | VISp\|APr |
| CP | FS\|GPe |
| CEA | SSp-bfd\|SSp-un\|BMA\|CP\|AAA\|IA\|ME[illegible] |
| CP | SSp-ul |
| VTA | PH\|PN\|MRN\|PAG\|IF\|IPN\|RL |
| VM | MOp\|SSp-ll\|SSp-tr\|RSPagl\|VAL\|RT\|ZI |
| VTA | PN\|RR\|MT\|IF\|RL\|CLI |
| MRN | RSPv\|PAG\|NPC\|RN |
| VTA | RSPv\|PH\|MRN\|PAG\|RN\|IF\|IPN |
| PIR | CLA\|EPd |
| PRP | NTS\|DMX\|GRN\|PGRNd\|NR\|MV\|XII |
| MOs | MOp |
| MPN | ACAv\|LSc\|LSr\|LSv\|SF\|MS\|TRS\|BST\|A[illegible] |

| | |
|---|---|
| MRN | VISp\|VISpm\|RSPv\|SCs\|IC\|SAG\|SCm |
| SSp-n | SSp-bfd |
| SCm | RSPagl\|RSPd\|SCs |
| PAG | MRN\|Pa4\|VTN\|DR\|LDT |
| NTS | AP\|CU\|GR\|DMX\|IRN\|MDRN\|MDRNd\| |
| DR | PAG\|IV\|CLI |
| SPIV | MV\|x |
| SPIV | LAV\|MV |
| COAp | BLA\|BMA\|PA\|CEA\|IA |
| PIR | EPv\|CP |
| TT | MOs\|ACAd\|PL\|ILA\|ORBm\|AON\|DP |
| RO | IO |
| SI | SSp-ul\|NLOT\|FS\|AAA\|CEA |
| AVPV | AVP\|MEPO\|MPO\|OV\|PVpo\|MPN |
| MRN | SCm\|PAG\|APN\|NPC\|RN |
| AUDd | SSs\|AUDp |
| RO | IO\|RPA |
| LHA | DMH\|AHN\|PVHd\|PeF\|ZI |
| ENTl | VISpor\|PERI\|ECT\|SUB |
| AN | |
| PCG | MRN\|VTN\|AT\|DR\|PDTg\|CS\|NI\|RPO |
| ENTl | VISpor\|PERI\|ECT\|SUB\|ProS |
| MV | NTS\|IRN\|PARN\|PGRNd |
| VCO | DCO\|PFL\|FL\|DN |
| CENT | |
| BMA | COAa\|BLA\|CEA\|IA\|MEA |
| SI | MOs\|CP\|ACB\|OT\|MA\|NDB\|LPO |
| AV | VAL\|AM\|AD\|IAD\|LD\|MD\|PCN\|CL |
| SCm | VISpm\|POST |
| ACB | LSr\|LSv\|SI\|MS\|NDB\|BST\|ADP\|MPO\|P |
| CP | MOs |
| ENTl | PERI\|ECT\|EPd |
| ARH | VMH\|ME |
| AM | IAM\|IAD\|MD\|PVT\|PT\|CM\|PCN |
| LP | DG\|PoT\|PO\|POL\|SGN\|PF\|MRN\|APN |
| PMv | VPMpc\|SPFm\|PF\|ARH\|DMH\|PVp\|MM |
| AHN | BST\|MPO\|MPN\|LHA\|LPO\|PeF |
| ARH | VMH |
| BST | SI\|LPO |
| PB | PSV\|DCO\|VCO\|LAV\|MV\|SPIV\|SUV\|y |
| CEA | MEA\|GPe\|GPi\|SI |
| SPVC | MDRN\|MDRNd |
| PIR | EPd\|EPv |
| MOB | |
| MOB | |
| PVH | SBPV\|PVHd |
| PIR | ENTl\|EPd |

<table>
<tr><td>PRE</td><td>DG|ENTm|PAR|SUB</td></tr>
<tr><td>MEA</td><td>COAa|COAp|BMA</td></tr>
<tr><td>STN</td><td>GPi</td></tr>
<tr><td>BST</td><td>AM|RT|ZI</td></tr>
<tr><td>PVHd</td><td>PVH</td></tr>
<tr><td>CENT</td><td>LING</td></tr>
<tr><td>CM</td><td>IAM|IMD|MD|RH</td></tr>
<tr><td>TRN</td><td>PG|PRNc|CS|RM</td></tr>
<tr><td>PVH</td><td></td></tr>
<tr><td>GRN</td><td>MARN|RM|RO</td></tr>
<tr><td>IC</td><td></td></tr>
<tr><td>SF</td><td>LSr|LSv|MS|TRS|PVT|PT|RE|MEPO</td></tr>
<tr><td>LGv</td><td>LGd|IGL|IntG|ZI</td></tr>
<tr><td>VII</td><td>SPVO|IRN|PARN</td></tr>
<tr><td>SPVC</td><td>SPVI|MDRN|MDRNd</td></tr>
<tr><td>LH</td><td>CA1|DG|MH</td></tr>
<tr><td>MRN</td><td>RSPd|RSPv|SUB|PF|PAG|APN|NPC|O</td></tr>
<tr><td>PARN</td><td>SPVO|IRN|ISN|LAV|MV|SUV</td></tr>
<tr><td>SI</td><td>CP|ACB|OT|MA|NDB</td></tr>
<tr><td>AN</td><td>DEC|SIM</td></tr>
<tr><td>PRNc</td><td>IC|PAG|P5|PRNr|IRN|PARN</td></tr>
<tr><td>PAG</td><td>MRN|CUN|PPN|PB</td></tr>
<tr><td>PRNc</td><td>PCG|NI|PRNr|SLD|GRN|IRN|MARN|R</td></tr>
<tr><td>CENT</td><td></td></tr>
<tr><td>SMT</td><td>VM|VPMpc|MD|RE|RH|CM|PCN</td></tr>
<tr><td>APN</td><td>RSPagl|DG|SCm|NOT|PPT</td></tr>
<tr><td>IRN</td><td>SPVC|MDRN|MDRNd|MDRNv|PARN|[illegible]</td></tr>
<tr><td>SO</td><td>LHA</td></tr>
<tr><td>SO</td><td></td></tr>
<tr><td>MM</td><td>PVp|SUM|TMd|PMd|PMv</td></tr>
<tr><td>CP</td><td>MOp</td></tr>
<tr><td>PRNr</td><td></td></tr>
<tr><td>NTS</td><td>CU|GR|DMX|MDRN|MDRNd|XII</td></tr>
<tr><td>AD</td><td>IAD|MD|PVT|PT</td></tr>
<tr><td>CP</td><td></td></tr>
<tr><td>LHA</td><td>LM|TMv|PMd|PH|PSTN</td></tr>
<tr><td>LGd</td><td>LP</td></tr>
<tr><td>VPM</td><td>VPLpc|PO</td></tr>
<tr><td>RT</td><td>GPe|VAL|VPL</td></tr>
<tr><td>PARN</td><td>NTS|SPVI|IRN|LIN|MV</td></tr>
<tr><td>RT</td><td>VPL</td></tr>
<tr><td>CUL</td><td>NLL</td></tr>
<tr><td>NLL</td><td>VISp|PSV|PB|SUT|FL</td></tr>
<tr><td>POST</td><td>VISp|PRE|APr</td></tr>
<tr><td>VTA</td><td>PN|IF|CLI</td></tr>
<tr><td>ZI</td><td>VM|RT|PVHd|LHA</td></tr>
<tr><td>AM</td><td>VAL|VM|SMT|PR|RE|Xi|PVH|PVHd|Z</td></tr>
</table>

| | |
|---|---|
| MOs | |
| SSs | SSp-bfd\|AUDd\|AUDp |
| MOs | |
| AUDd | SSs\|AUDp |
| VISp | |
| VISp | |
| ACAv | ACAd |
| CP | |
| VISam | VISa |
| ORBl | FRP\|ORBvl |
| ACAd | ACAv |
| SSs | AUDd |
| VISp | |
| ACAd | ACAv\|PL |
| MOp | SSp-ll\|SSp-ul |
| VISpor | TEa |
| MS | LSr\|LSv\|SF |
| PB | MEV\|PAG\|PCG\|LC\|LING\|CENT\|NOD\| |
| CS | MRN\|VTN\|AT\|IPN\|CLI\|TRN\|PRNr\|RPC |
| NLL | IC\|SAG\|MRN\|SCm\|PPN\|PSV\|PB\|SUT\| |
| MD | IMD\|PVT\|MH\|LH |
| MRN | NB\|SAG\|SCm |
| PAG | MH\|LH\|SCO\|NPC |
| PSV | SAG\|PPN\|NLL\|PB\|SUT\|V\|P5 |
| VM | LHA\|ZI |
| ProS | CA1\|SUB |
| VMH | AHN |
| MEA | NLOT\|BMA |
| PIR | AIp\|EPd |
| SI | CP\|ACB\|OT\|MA\|NDB\|LPO |
| RSPv | RSPd |
| VII | AMB\|IRN\|PARN\|PGRNl |
| LGv | LGd\|IGL |
| NLL | |
| VISp | |
| SSp-ll | SSp-ul |
| VISa | VISam |
| MOs | MOp |
| MOp | SSp-ll |
| MV | SPIV |
| CP | |
| PMv | PVp\|LM\|MM\|TMv\|PMd |
| LM | SUM\|LHA |
| AUDp | AUDd |
| VISp | |
| SSp-n | SSp-bfd\|SSp-un |
| ACAd | ACAv |

<table>
<tr><td>CP</td><td></td></tr>
<tr><td>CP</td><td>ACB</td></tr>
<tr><td>MOs</td><td>MOp</td></tr>
<tr><td>SSp-un</td><td>SSp-n|SSp-bfd|SSp-ul</td></tr>
<tr><td>MOs</td><td>ACAd</td></tr>
<tr><td>VISpor</td><td>ECT|ENTl</td></tr>
<tr><td>ARH</td><td>ME</td></tr>
<tr><td>VISp</td><td></td></tr>
<tr><td>NDB</td><td>ACAv|ACB|LSr|LSv|SI|MS|BST|ADP|A</td></tr>
<tr><td>VMH</td><td>PVi|ARH|DMH|AHN|TMv|PMv|PH|LH</td></tr>
<tr><td>MM</td><td>LM|SUM|PMd|PH|LHA|VTA|MT</td></tr>
<tr><td>MPO</td><td>BST|AVP|PD|PS|MPN|LPO</td></tr>
<tr><td>EPd</td><td>AIp|PIR</td></tr>
<tr><td>LHA</td><td>SSp-ll|CEA|MEA|GPi|SI|ZI</td></tr>
<tr><td>GPe</td><td>RT</td></tr>
<tr><td>PIR</td><td>SSp-bfd|VISC|EPd|EPv|LA|BLA</td></tr>
<tr><td>VPM</td><td>SSp-tr|CA3|VPL|LGd|LP|PO|LD</td></tr>
<tr><td>PAG</td><td>MRN|III|EW|IF|IPN|RL|CLI|CS</td></tr>
<tr><td>PCG</td><td>MRN|VTN|B|DTN|PDTg|LDT|NI|SLD</td></tr>
<tr><td>MV</td><td>MRN|PB|B|PCG|PRNc|LC|LDT|NI|PR</td></tr>
<tr><td>MRN</td><td>RSPd|PF|PAG|APN|NPC|RN</td></tr>
<tr><td>MPO</td><td>SI|NDB|BST|AVP|AVPV|PS|VMPO|VL</td></tr>
<tr><td>PFL</td><td></td></tr>
<tr><td>PRNc</td><td>VI|GRN|IRN|PARN|PGRNd|MV</td></tr>
<tr><td>AM</td><td>AV|AD|IAM|IAD|MD|PVT|PT|RE|RH</td></tr>
<tr><td>CM</td><td>VM|VPMpc|SPFm|SPA|IMD|MD|SMT</td></tr>
<tr><td>SNr</td><td>RSPv|VTA|RR|MRN|RN|MT|SNc|PRN</td></tr>
<tr><td>NOT</td><td>RSPv|SUB|SCm|MPT|OP|PPT</td></tr>
<tr><td>CP</td><td>MOp</td></tr>
<tr><td>MDRN</td><td>AMB|GRN|IO|IRN|LRN|MARN|MDRN</td></tr>
<tr><td>MDRN</td><td>SPVC|SPVI|AMB|GRN|IRN|LIN|LRN|M</td></tr>
<tr><td>RSPv</td><td>RSPd</td></tr>
<tr><td>NLOT</td><td>COAa|BMA|AAA|MEA</td></tr>
<tr><td>ORBm</td><td>PL|ILA|ORBvl</td></tr>
<tr><td>PH</td><td>DMH|AHN|PVHd</td></tr>
<tr><td>CP</td><td></td></tr>
<tr><td>SSp-bfd</td><td>SSp-un</td></tr>
<tr><td>VISl</td><td>VISal|VISli</td></tr>
<tr><td>SPFm</td><td>SPA|PH|PAG</td></tr>
<tr><td>ACAv</td><td>ACAd</td></tr>
<tr><td>VISli</td><td>VISl</td></tr>
<tr><td>SSp-bfd</td><td></td></tr>
<tr><td>VISpor</td><td>TEa</td></tr>
<tr><td>AId</td><td>AIv</td></tr>
<tr><td>CP</td><td></td></tr>
<tr><td>VISp</td><td></td></tr>
<tr><td>ACAd</td><td>MOs|ACAv</td></tr>
</table>

| | |
|---|---|
| PT | AV\|AM\|IAD\|PVT\|RE |
| MOs | ACAd |
| AHN | BST\|ASO\|PVH\|MPO\|SBPV\|MPN\|PVH( |
| PT | AM\|IAM\|IAD\|PVT\|RE\|RH\|CM |
| BST | BAC |
| AIp | VISC\|PIR\|CLA\|EPd |
| SSs | SSp-n |
| LDT | MEV\|MRN\|PAG\|PB\|PCG\|LC |
| ILA | PL\|ORBm |
| VISp | |
| ACAd | MOs |
| CP | SSs\|VISC |
| RSPd | VISp\|RSPagl |
| SSp-bfd | |
| NTS | CU\|GR\|DMX |
| PH | PVi\|DMH\|MM\|SUM\|TMd\|TMv\|PMd\| |
| SI | MOp\|CP\|ACB\|FS\|OT\|AAA\|CEA\|MA |
| MOp | |
| SSp-m | |
| SSp-m | SSs\|GU |
| ARH | PVi\|DMH\|PVp\|PMv\|VMH |
| VISam | VISpm |
| RSPv | |
| ORBl | ORBvl |
| SSp-bfd | SSp-n\|SSp-un |
| CP | SSp-ul |
| GPe | SSp-ul\|SSp-un\|CP |
| PB | MEV\|PCG\|LC\|LDT\|SLC\|MV |
| SIM | AN |
| AON | PL\|ORBm\|ORBvl\|MOB\|TT |
| MRN | IC\|NB\|SCm |
| DG | CA1 |
| DG | CA3 |
| AUDd | SSs\|AUDp |
| GRN | PRNc\|VII\|IRN\|MARN\|PGRNl\|PPY\|RM |
| GRN | MARN\|PGRNd\|RM\|RO |
| SPVI | DCO\|SPVO\|PARN |
| SPVI | SPVC\|Pa5 |
| DCO | LAV\|SUV\|y |
| IRN | SPVO\|VII\|ACVII\|GRN\|PARN |
| LSc | LSr |
| DMH | PMv\|VMH\|PH\|LHA\|TU |
| DMH | PH\|LHA |
| LSr | LSc\|LSv\|NDB\|ADP\|AVP\|AVPV\|MPO\|P |
| MPN | PVpo\|SBPV\|AHN |
| CP | MOp\|MOs |
| CP | ACB\|LSr\|LSv |

| | |
|---|---|
| PAG | DR |
| VTA | ZI\|SNr\|MRN\|MT\|SNc |
| CP | |
| VTA | RR\|MT |
| CP | |
| AOB | MOB\|AON |
| ACAv | ACAd |
| VII | IRN\|PGRNl |
| SI | CP\|ACB\|FS\|MA\|NDB\|BST\|LPO |
| PL | ILA |
| VPMpc | VM\|VPLpc\|SPFm\|PO\|MD\|SMT\|PCN\|[illegible] |
| ZI | VAL\|VM\|VPL\|VPLpc\|VPM\|VPMpc\|PO |
| V | MRN\|PPN\|PB\|SUT\|P5\|PRNr |
| PAG | MRN\|SCm |
| CLI | VTA\|MRN\|PAG\|RN\|EW\|IV\|IF\|RL\|DR |
| PH | SPFm\|SPA\|MM\|SUM\|LHA\|PAG |
| MEA | BMA\|CEA\|GPe\|GPi\|SI\|LHA |
| LHA | DMH\|PVHd\|PH\|PST\|PSTN\|PeF\|ZI |
| ACAd | MOs\|ACAv |
| BMA | NLOT\|COAa\|EPv\|BLA\|AAA\|CEA\|IA\|M |
| XII | NTS\|DMX\|MDRN\|MDRNv\|NR |
| PSTN | SUM\|LHA\|STN |
| VTA | PH\|PN\|MRN\|PAG\|MA3\|EW\|IF\|IPN\|R |
| AUDp | AUDd\|AUDv |
| V | PB\|SUT\|P5\|CENT |
| VTA | RR\|MRN\|PAG\|RN\|MA3\|RL\|CLI |
| RSPv | RSPd |
| MOs | MOp |
| MOp | SSp-ll |
| VISpor | VISpl\|VISli\|TEa |
| AIv | MOp\|AId\|CLA |
| MOB | |
| MOB | AOB |
| MD | VAL\|AM\|IAM\|IAD\|IMD\|SMT\|RH\|CM\| |
| LP | PoT\|POL\|SGN\|APN |
| AUDp | AUDd |
| RSPv | RSPd |
| VISp | |
| RSPv | |
| SSp-ll | SSp-ul |
| ACAd | MOs\|ACAv\|PL |
| VISp | |
| SSp-ul | SSp-ll |
| VISp | |
| RSPd | RSPv |
| SSp-bfd | SSp-un |
| VISal | VISl |

<table>
<tr><td>MOs</td><td>ACAd|PL</td></tr>
<tr><td>MOp</td><td>SSp-ll|SSp-ul</td></tr>
<tr><td>VISl</td><td>VISli</td></tr>
<tr><td>PVHd</td><td>PVH</td></tr>
<tr><td>SSp-bfd</td><td>VISrl</td></tr>
<tr><td>VISp</td><td></td></tr>
<tr><td>SSp-ll</td><td>SSp-ul|SSp-tr</td></tr>
<tr><td>RE</td><td>PR|Xi|PVH|AHN|PVHd|ZI</td></tr>
<tr><td>BST</td><td>LSv|SF|MS|TRS|PVT|PT</td></tr>
<tr><td>TT</td><td>ACAv|DP|IG|LSr|SH|MS</td></tr>
<tr><td>PVp</td><td>ARH|MM|TMd|PMd|PMv</td></tr>
<tr><td>SCm</td><td>MRN|APN|NOT|OP|PPT</td></tr>
<tr><td>NDB</td><td>SI|MPO|LPO</td></tr>
<tr><td>AN</td><td>DEC|SIM</td></tr>
<tr><td>CUL</td><td>CENT</td></tr>
<tr><td>VCO</td><td>DCO</td></tr>
<tr><td>RO</td><td>GRN|IO|RPA</td></tr>
<tr><td>ENTl</td><td></td></tr>
<tr><td>DCO</td><td>LAV|y|IP</td></tr>
<tr><td>PT</td><td>AM|IAD|MD|PVT</td></tr>
<tr><td>SSp-bfd</td><td>SSp-n|SSp-un</td></tr>
<tr><td>VISp</td><td></td></tr>
<tr><td>PRNc</td><td>VI|GRN|RM</td></tr>
<tr><td>LP</td><td>CA3|DG|PO|POL</td></tr>
<tr><td>MOs</td><td>MOp</td></tr>
<tr><td>SSp-bfd</td><td>SSp-un</td></tr>
<tr><td>PB</td><td>PAG|CUN</td></tr>
<tr><td>VISp</td><td></td></tr>
<tr><td>ACAd</td><td>MOs|ACAv</td></tr>
<tr><td>CA1</td><td>RSPv|FC|SUB|ProS</td></tr>
<tr><td>MOs</td><td>MOp</td></tr>
<tr><td>SSp-bfd</td><td>SSp-n|SSp-un</td></tr>
<tr><td>AM</td><td>AV|AD|IAD|MD|PT</td></tr>
<tr><td>POST</td><td>VISp|PAR|PRE|APr</td></tr>
<tr><td>PMv</td><td>PVp|LM|SUM|TMv</td></tr>
<tr><td>IO</td><td>RPA</td></tr>
<tr><td>MOs</td><td></td></tr>
<tr><td>ENTm</td><td>ENTl</td></tr>
<tr><td>VISal</td><td>VISl|VISp|VISrl</td></tr>
<tr><td>ORBvl</td><td>PL|ORBl|ORBm</td></tr>
<tr><td>OT</td><td>CP|ACB|SI</td></tr>
<tr><td>ACB</td><td>CP|OT|SI</td></tr>
<tr><td>MOs</td><td></td></tr>
<tr><td>MD</td><td>DG|AD|LD|CL|LH</td></tr>
<tr><td>SSp-bfd</td><td>SSp-n|SSp-un</td></tr>
<tr><td>MD</td><td>RSPv|CA1|CA2|CA3|DG|PVT|CM|PCN</td></tr>
<tr><td>PVp</td><td>DMH|TMd|TMv|PMd|PMv</td></tr>
</table>

| | |
|---|---|
| LSr | MS |
| IAD | AM\|MD\|CM\|PCN |
| AIp | CLA |
| SSp-n | SSs |
| VISp | VISpm\|RSPagl |
| MOs | ACAd |
| SSs | SSp-n |
| RSPagl | VISp\|RSPd |
| ORBl | ORBvl |
| VISa | VISam |
| RO | IO |
| MOp | |
| SSp-m | |
| AN | |
| SIM | AN |
| DR | MRN\|IF\|IPN\|CLI\|CS |
| MD | LH |
| MD | PVT\|PT\|LH |
| COAp | BMA\|PA |
| NPC | SCm\|PAG\|MPT\|NOT\|OP\|PPT |
| PVp | ARH\|DMH\|TMd\|TMv\|PMv |
| RSPv | |
| MEA | CEA\|SI |
| MM | SUM\|PMd\|PH |
| ACAv | ACAd |
| MOs | |
| AUDp | AUDd |
| SSp-bfd | SSp-n\|SSp-un |
| PRNr | MRN\|PAG\|PRNc\|SUT\|P5 |
| VISp | |
| ACB | SI\|BST |
| VI | |
| PH | DMH\|LHA |
| ZI | VM\|VPL\|VPM\|LP\|PO\|LHA\|PST\|PSTN\| |
| IP | PB\|ICB\|SUV\|CENT\|CUL\|VeCB |
| VISpor | VISli\|ENTl |
| ZI | GPi\|VM\|RT\|LHA\|STN |
| RE | VAL\|VM\|AM\|PR\|Xi\|PVH\|PVHd |
| ACB | OT\|SI |
| ZI | STN |
| VM | SMT\|PR\|PH\|ZI |
| LHA | DMH\|AHN\|PVHd\|PH\|PeF\|ZI |
| MEA | SI\|LHA |
| MEA | COAp\|PA |
| APN | POL\|MRN\|SCm |
| AON | ORBvl\|MOB\|TT |
| CA1 | SSp-ll\|SSp-tr\|CA2\|CA3 |

| | |
|---|---|
| IPN | VTA\|PN\|MRN\|EW\|IF\|RL\|CLI |
| LHA | AHN\|PVHd\|PeF\|ZI |
| DMX | NTS\|XII |
| VTA | RR\|MRN\|IF\|CLI |
| RT | |
| RT | VPL |
| ZI | |
| ZI | RSPagl\|CA1\|VM\|PH\|LHA |
| PT | AV\|AM\|IAD\|PR\|PVT\|RE\|RT |
| CA3 | CA2\|DG |
| ACAd | MOs |
| SSp-n | SSp-bfd\|SSp-un |
| PGRNl | AMB\|GRN\|IRN\|LRN\|MDRN\|MDRNd\| |
| ARH | PVp\|ME |
| PIR | |
| MD | MOs\|ACAd\|RSPagl\|RSPd\|RSPv\|CA3\|D |
| VISC | SSs\|AIp |
| MRN | VISp\|PoT\|SPFp\|PP\|MG\|POL\|SGN\|PIL |
| MRN | RSPd\|RSPv\|CA1\|SUB\|ProS\|SPFp\|PF\|P |
| VM | RSPd\|RSPv\|VPMpc\|SMT\|ZI |
| PO | DG\|LP |
| ZI | SSp-tr\|VISa\|STN\|SNr\|SNc |
| RE | AM\|IAM\|IAD\|SMT\|PR\|PVT\|PT\|Xi\|RH |
| ZI | VISam\|VISpm\|PoT\|SPFp\|PP\|MG\|LP\|P |
| RE | RSPv\|VAL\|VM\|AM\|IAM\|IAD\|MD\|SM |
| CP | SSp-n\|SSp-m |
| AHN | PR\|RE\|Xi\|PVH\|PVa\|PVi\|DMH\|SBPV\| |
| VMH | MOs\|PVi\|ARH\|AHN\|RCH\|TU |
| PGRNl | VII\|AMB\|GRN\|IO\|IRN\|LRN\|MARN\|PP |
| SCm | NB\|MRN |
| IPN | PN\|IF\|CS\|PRNr |
| SNr | SNc |
| MPO | NDB\|AVP\|VLPO\|AHN\|MPN\|LPO |
| PRNc | NTB\|GRN\|RM |
| IRN | IC\|PB\|PRNc\|SPVO\|GRN\|PARN\|PGRNc |
| AAA | NLOT\|COAa\|BMA\|FS\|MEA\|SI\|MA |
| ACB | CP\|BST |
| PH | RSPv\|VM\|VPMpc\|MD\|SMT\|PR\|RE\|Xi |
| RE | PR\|Xi\|CM\|PVH |
| CA1 | CA2\|CA3 |
| ACAd | MOs\|ACAv |
| MOB | |
| ACB | |
| CP | |
| VTA | LM\|SUM\|LHA\|SNr\|MT\|SNc |
| ARH | DMH\|PVp\|TMd\|PMv |
| BST | |

| | |
|---|---|
| MD | IMD\|PVT\|PT\|CM\|MH |
| MD | DG\|LP\|AV\|AD\|LD\|CL\|LH |
| NTS | CU\|GR\|DMX |
| LD | LP\|AD\|MD\|CL\|LH |
| AV | AM\|AD\|IAD |
| SSp-bfd | SSp-tr |
| AUDd | SSs\|AUDp |
| ARH | PVi\|DMH\|PVp\|TMd\|PMv\|VMH\|TU |
| PVH | IAM\|PR\|RE\|DMH\|AHN\|PVHd\|PeF\|ZI |
| MOs | MOp\|ACAd |
| SSp-un | SSp-n\|SSp-bfd |
| RSPv | MOs\|ACAd\|ACAv\|RSPagl\|RSPd |
| SSp-bfd | SSp-tr\|VISa\|VISrl |
| VISp | VISal\|VISl\|VISrl |
| VMH | ARH\|DMH\|PVp\|TMv\|PMv\|TU |
| LHA | RT\|SO\|AHN\|PVHd\|VMH\|PeF\|RCH\|TU |
| TU | ARH\|VMH |
| PT | PVT\|RE\|RT |
| PH | MM\|SUM\|IF |
| DP | PL\|ORBm\|ORBvl\|AON\|TT |
| SCs | |
| VISp | |
| SSp-ll | SSp-ul |
| SSp-n | SSs |
| III | MRN\|PAG\|EW\|IV\|Pa4\|RL |
| NDB | LSr\|MS\|AVPV |
| XII | DMX\|NR |
| CA3 | CA1\|CA2\|DG |
| CA3 | CA1\|CA2\|DG |
| VISp | |
| VISp | |
| SSp-m | |
| ACAd | ACAv |
| SSs | SSp-m\|GU |
| ILA | PL\|DP |
| FRP | MOs\|ORBl |
| TU | SO\|AHN\|VMH\|LHA\|PeF\|RCH |
| SSp-m | SSs\|GU |
| MOp | SSp-ul |
| DCO | VCO |
| XII | IRN\|MDRN\|MDRNd\|MDRNv\|NR |
| VII | IRN\|PARN |
| SSp-bfd | SSp-un |
| ARH | PVi\|DMH\|PVp\|PMv\|VMH\|TU |
| DMH | PVi\|PH |
| PGRNl | GRN\|IO\|LRN\|MARN |
| SSp-ll | SSp-tr |

| | |
|---|---|
| CA3 | CA2 |
| LSr | LSc\|SF\|TRS |
| SSp-bfd | |
| SO | |
| ENTm | PAR\|PRE\|SUB |
| MG | DG\|PP\|PIL |
| MOp | SSp-ll |
| MOp | SSp-ul |
| MOB | |
| CA3 | CA1\|CA2 |
| CA3 | |
| CA3 | CA1\|CA2 |
| IMD | MD\|PVT\|CM |
| PMv | ARH\|DMH\|PVp\|TMd\|TMv\|VMH\|TU |
| MOs | MOp |
| SSp-n | SSp-un |
| ENTl | VISpor\|PERI |
| CA1 | VISam\|VISp\|VISpm\|VISa |
| MG | PP\|PIL |
| CA1 | |
| MRN | PH\|VTA\|PAG\|RN\|MA3\|EW\|RL |
| CA3 | CA2 |
| PAG | VTA\|MRN\|SCm\|NPC\|RN\|MA3\|RL |
| NTS | CU\|GR\|DMX |
| PT | PVT |
| VPM | VPLpc\|PoT\|MG\|PO |
| AHN | BST\|PVH\|PVa\|PVi\|MPO\|PVpo\|SBPV\|I |
| VM | ZI |
| PO | |
| MEA | NLOT\|BMA\|AAA\|CEA\|IA\|SI |
| AHN | PR\|PT\|RE\|RT\|PVH\|PVi\|ARH\|SBPV\|SC |
| PGRNl | VII\|AMB\|GRN\|IO\|IRN\|LRN\|MARN\|PA |
| AN | |
| CUL | |
| SSp-un | SSp-n\|SSp-bfd\|SSp-ul |
| VISp | |
| MOs | MOp |
| SSp-n | SSp-bfd\|SSp-un |
| SSs | AUDd |
| MOs | MOp |
| SSp-n | SSp-bfd\|SSp-un |
| AUDp | AUDd |
| VISp | |
| CA1 | |
| BST | PVH\|MPO\|PD\|MPN |
| SSp-n | SSp-bfd\|SSs |
| SSp-m | |

| | |
|---|---|
| ACAd | MOs\|ACAv |
| ENTm | VISpl |
| SSp-ul | |
| RSPv | |
| CEA | CP\|FS\|GPe\|SI |
| CEA | BLA\|BMA\|AAA\|IA\|MEA\|GPi\|SI\|LHA |
| ARH | DMH\|PVp\|TMv\|PMv\|VMH\|ME |
| MD | AV\|AD\|IAD\|LD\|CL\|MH\|LH |
| DG | CA3\|ENTm\|SUB\|ProS\|HATA |
| PPT | SCs\|SCm\|APN\|MPT\|NOT\|NPC\|OP |
| GRN | PRNc\|VII\|IRN\|MARN\|PGRNl |
| NPC | PAG\|APN\|MPT\|NOT\|OP\|PPT |
| AUDp | |
| MEA | NLOT\|BMA\|AAA\|CEA\|GPe\|GPi\|SI\|MA |
| MOB | AON |
| SSp-bfd | SSp-un |
| VISp | |
| AUDp | AUDd |
| PAG | SPA\|PVT\|PF\|LH |
| MM | SUM\|PMd\|PH |
| ACAd | MOs\|ACAv |
| VISp | |
| IMD | SPA\|MD\|PVT\|PF |
| PCG | MRN\|PPN\|PB\|B\|PRNc\|LC\|LDT\|SLC\|SL |
| ENTl | |
| CA1 | SUB\|ProS |
| PAG | MRN\|III\|IV\|Pa4\|CLI\|DR |
| AD | CA3\|DG\|AV\|IAD\|LD\|MD\|CL |
| VISp | |
| COAp | BLA\|BMA\|PA |
| DG | CA1 |
| DMH | PVi\|TMd\|PMd\|PMv\|VMH\|PH |
| VMH | DMH\|PMv\|TU |
| RSPv | RSPd |
| TRS | MOs\|ACAd\|ACAv\|LSr\|SF |
| PGRNl | GRN\|IO\|LRN\|MARN |
| MPO | BST\|PVH\|ADP\|AVPV\|MEPO\|PD\|PVpo |
| MPN | PVH\|MPO\|SBPV\|AHN |
| PSV | |
| SSp-bfd | |
| RSPd | RSPv |
| PO | VPM |
| SCm | SCs\|PAG |
| CA3 | CA1\|CA2 |
| CA3 | |
| MOp | SSp-ll |
| PAR | ENTm\|PRE |

| | |
|---|---|
| PFL | FL |
| AOB | AON |
| AUDp | |
| MOs | MOp\|ACAd |
| ENTl | VISpor\|ENTm |
| MOs | FRP |
| SSp-ll | MOp\|SSp-ul |
| V | PSV\|SUT\|P5\|Acs5\|PC5\|I5\|SPVO\|IRN\| |
| SGN | PoT\|MG\|LP\|POL |
| AV | LP\|AD\|IAD\|LD\|MD\|PCN\|CL |
| AHN | PVH\|DMH\|SBPV\|PVHd\|PeF\|ZI |
| CA1 | DG |
| AUDp | AUDd\|AUDv |
| LSv | LSr\|SF\|TRS\|BST\|PT |
| CM | IMD\|MD\|PCN |
| PGRNl | IO |
| PO | LP |
| VISpm | VISam\|VISp\|VISa |
| CA3 | VISp\|VISrl\|CA1\|DG |
| AUDp | AUDd |
| MOs | ACAd |
| CP | |
| PO | LP |
| CM | IMD\|MD\|PCN |
| MD | CL\|PF\|LH |
| PCN | VAL\|PO\|MD\|CL |
| MOp | MOs |
| ACAd | MOs |
| AUDp | |
| SSp-bfd | SSp-un |
| MOs | ACAd |
| VISp | |
| LP | RSPagl\|CL\|PF |
| LP | CA3\|DG\|LD |
| ENTl | PERI\|ECT |
| PVT | IMD\|MD\|PT\|CM |
| LP | CA3\|DG\|LD |
| LP | POL |
| PB | CENT |
| MD | PVT\|PCN\|CL\|MH\|LH |
| MG | CA3\|DG\|LGd\|LP\|SGN |
| AIp | SSs\|VISC\|CLA\|EPd |
| SSp-bfd | |
| IC | PAG\|CUN |
| SCm | SCs\|MPT\|NOT\|PPT |
| DG | |
| PVH | PVHd |

| | |
|---|---|
| ACAv | ACAd |
| ACAv | MOs\|ACAd |
| ORBl | ORBvl |
| VISp | |
| ACAd | MOs |
| DR | PAG\|CLI |
| MOs | MOp |
| ORBvl | FRP\|MOs\|ORBl |
| PB | MEV\|LC |
| VISrl | SSp-bfd\|VISp |
| VISp | |
| VISpor | VISli\|TEa |
| PAR | ENTm\|POST\|PRE |
| PVT | IMD\|MD\|CM |
| RE | IAM\|PVT\|PT\|Xi\|RH\|CM |
| PVT | SPA\|IMD\|MD\|PF\|LH\|PAG |
| AUDp | AUDd |
| VISam | VISpm\|VISa |
| RSPv | |
| AId | AIv |
| IPN | IF |
| ACAv | |
| PIR | EPd\|EPv |
| NLOT | COAa\|BMA\|AAA\|MEA |
| ENTm | |
| BST | BAC\|PT\|RE\|PVH |
| RE | VM\|AM\|IAD\|PR\|PT\|RT\|PVH\|PVHd\|ZI |
| BST | LSv\|BAC\|PT\|RE\|RT\|PVH\|ZI |
| CEA | MEA |
| PVT | MD\|PT |
| ARH | PVp |
| ARH | DMH\|PVp\|PMv\|VMH\|TU |
| ENTl | VISpor\|ENTm |
| ARH | DMH\|PVp\|PMv\|VMH |
| CEA | MEA\|GPi\|SI |
| BST | CP\|AV\|RT |
| BST | PVH\|PD |
| SSp-n | SSp-m\|SSp-un |
| RSPv | RSPd\|SCs\|SCm |
| GU | SSp-m\|SSs\|AId |
| CA3 | DG |
| SSs | |
| SSp-m | SSp-n\|SSs |
| VISpm | VISp |
| NLL | PSV\|PB\|SUT\|V\|P5\|PC5\|PRNr |
| ACAd | MOs |
| PRNc | TRN\|PRNr |

| | |
|---|---|
| MRN | SCs\|SCm\|PAG\|RN |
| MOs | MOp |
| ORBm | MOs\|PL\|ILA\|ORBvl |
| PAG | SCm\|DR |
| PCG | MRN\|PAG\|VTN\|DTN\|LDT |
| AId | MOp\|AIv\|CLA |
| PL | ACAd\|ILA\|ORBm |
| PVT | IMD\|MD |
| LGd | VPM\|LP |
| MOs | FRP |
| ARH | VMH\|TU\|ME |
| SSp-bfd | |
| SSp-tr | SSp-ll\|SSp-ul\|VISa |
| LRN | IRN\|MDRN\|MDRNv\|PGRNl |
| PMv | PVp\|TMd\|TMv\|PMd\|LHA |
| VISp | |
| MOs | MOp |
| DMX | CU\|GR\|NTS\|IRN\|MDRN\|MDRNd\|XII |
| PF | VPMpc\|SPFm\|SPFp\|PH\|ZI\|VTA\|MRN\| |
| ENTl | ENTm |
| CS | IPN\|PG\|TRN |
| APN | PoT\|SPFp\|POL\|PIL\|MRN |
| AId | MOp\|AIv |
| SUB | RSPv\|CA1\|FC\|ProS |
| PVp | ARH\|DMH\|TMd\|PMv\|VMH |
| CP | GPe |
| SPVC | |
| IP | ICB |
| APN | PAG\|NPC\|OP\|PPT |
| TU | ARH\|TMv\|PMv\|VMH |
| RSPagl | SSp-tr |
| VMH | RSPagl\|ARH\|DMH\|TMv\|PMv\|RCH\|TU |
| TU | ARH\|DMH\|PVp\|TMv\|PMv\|VMH\|PH\|L |
| AId | MOp\|CLA |
| IRN | AMB\|MDRN\|MDRNd\|PGRNl |
| XII | NTS\|DMX\|IRN\|MDRN\|MDRNv\|NR |
| PB | |
| PPN | MRN |
| SSp-ll | SSp-tr |
| VISli | VISl\|VISpor\|TEa |
| MOB | |
| PSTN | LHA\|PST\|STN\|ZI\|SNr\|SNc |
| ZI (PSTN) | VAL\|VM\|VPM\|PO\|LHA\|PST\|PSTN\|STN |
| MD | IAM\|IAD\|IMD\|PVT\|PT\|CM |
| ORBl | FRP\|MOs\|ORBvl |
| DG | |
| DG | VISp\|CA3 |

<table>
<tr><td>ENTm</td><td></td></tr>
<tr><td>ENTm</td><td>ENTl</td></tr>
<tr><td>AON</td><td>ACB</td></tr>
<tr><td>SUB</td><td>ProS</td></tr>
<tr><td>LHA</td><td>SO|PeF</td></tr>
<tr><td>VCO</td><td></td></tr>
<tr><td>ACB</td><td>LSr|LSv</td></tr>
<tr><td>BST</td><td>LHA|LPO</td></tr>
<tr><td>RE</td><td>PR|Xi|PVH|ZI</td></tr>
<tr><td>MH</td><td>PVT|LH</td></tr>
<tr><td>ACAv</td><td>MOs|ACAd</td></tr>
<tr><td>MOs</td><td>MOp</td></tr>
<tr><td>ACAv</td><td></td></tr>
<tr><td>MOs</td><td>MOp|ACAd</td></tr>
<tr><td>APN</td><td></td></tr>
<tr><td>PB</td><td>CUN|PPN|SUT</td></tr>
<tr><td>MDRN</td><td>SPVC|AMB|IRN|LRN|MDRNd|MDRNv</td></tr>
<tr><td>CEA</td><td>MEA|GPe|GPi|SI</td></tr>
<tr><td>ACAv</td><td>ACAd</td></tr>
<tr><td>ORBl</td><td>FRP|ORBvl|AOB</td></tr>
<tr><td>DMH</td><td>PVi|AHN|VMH</td></tr>
<tr><td>ORBm</td><td>ORBvl</td></tr>
<tr><td>SSp-bfd</td><td></td></tr>
<tr><td>IP</td><td>SUV</td></tr>
<tr><td>MOs</td><td>MOp</td></tr>
<tr><td>CP</td><td></td></tr>
<tr><td>PRNc</td><td>MRN|PCG|CS|NI|PRNr|RPO|SLD</td></tr>
<tr><td>GPe</td><td>GPi</td></tr>
<tr><td>GPe</td><td>GPi|SI</td></tr>
<tr><td>MS</td><td>LSr|LSv|SF</td></tr>
<tr><td>CEA</td><td></td></tr>
<tr><td>LSr</td><td>LSc|SF|MS|TRS</td></tr>
<tr><td>RE</td><td>IAM|MD|SMT|PR|Xi|RH|CM</td></tr>
<tr><td>PAG</td><td>MRN|III|IV|Pa4|DR</td></tr>
<tr><td>VISp</td><td></td></tr>
<tr><td>SSp-bfd</td><td>SSp-n|SSp-un</td></tr>
<tr><td>DG</td><td>CA3</td></tr>
<tr><td>AM</td><td>AV|AD|IAD|PT</td></tr>
<tr><td>DMH</td><td>VM|MD|SMT|PR|RE|CM|PCN|AHN|[illegible]</td></tr>
<tr><td>SSs</td><td></td></tr>
<tr><td>MOs</td><td>MOp</td></tr>
<tr><td>VISp</td><td>VISl</td></tr>
<tr><td>PH</td><td>DMH|SUM|PMd|LHA</td></tr>
<tr><td>LGd</td><td>VPM|MG|LP|SGN</td></tr>
<tr><td>SSp-ul</td><td>SSp-bfd|SSp-un</td></tr>
<tr><td>VISp</td><td></td></tr>
<tr><td>DG</td><td>CA3</td></tr>
</table>

| | |
|---|---|
| SSp-bfd | |
| VISl | VISal\|VISp |
| CA3 | |
| VMH | PVi\|ARH\|DMH\|SBPV\|AHN\|RCH |
| AHN | BST\|PR\|RE\|RT\|ASO\|PVH\|MPO\|SBPV\| |
| PAG | MRN\|IV\|CLI\|DR |
| MRN | SPFp\|PP\|PIL\|ZI\|SNr\|SNc |
| AOB | ORBl\|ORBvl\|MOB\|AON |
| ORBvl | PL\|ORBm |
| PL | MOs\|ACAd\|ILA |
| LP | DG\|PO\|LD\|CL |
| SSp-ll | MOp\|SSp-ul |
| SSp-ll | SSp-ul |
| APN | MRN\|SCm\|NOT\|PPT |
| PF | RSPd\|RSPv\|VPMpc\|SPFp\|MRN |
| PL | ACAv\|ILA\|DP |
| VISp | |
| PH | ARH\|DMH\|PMv\|VMH\|LHA\|TU |
| PT | LSv\|BST\|AM\|IAD\|RE |
| DCO | LAV\|y\|IP\|DN |
| PVH | PR\|RE\|AHN\|PVHd\|ZI |
| SSp-n | SSp-bfd |
| COAp | BMA\|PA |
| AUDp | AUDd |
| ACAd | MOs\|PL |
| SSp-ll | |
| IC | |
| PAG | MRN\|SCm\|RN |
| PAG | SCs\|MRN\|SCm |
| RSPagl | VISp\|VISpl\|RSPd\|POST |
| ACAd | MOs |
| AON | MOB\|TT |
| CA1 | CA2\|CA3\|DG |
| SIM | CUL |
| COAp | COAa\|BLA\|BMA\|PA\|MEA |
| LDT | B\|PCG |
| CEA | MEA\|SI |
| MDRN | GRN\|IO\|MDRNv\|NR\|XII\|RO |
| ENTm | ENTl |
| LHA | DMH\|PVHd\|PH\|PST\|PeF |
| PH | SPFm\|VTA |
| LHA | TU |
| TU | LHA\|PeF |
| AId | |
| PMv | DMH\|TMv\|PMd\|PH\|TU |
| LP | VISam\|VISpm\|MG\|SGN |
| MD | RSPd\|RSPv\|DG\|PF\|LH |

| | |
|---|---|
| LGv | CA1\|CA2\|CA3\|LGd\|RT\|IGL\|IntG\|ZI |
| IPN | VTA\|PN\|MRN\|PAG\|MA3\|EW\|IF\|RL\|C |
| VMH | PVi\|ARH\|DMH\|TU |
| MEA | NLOT\|AAA\|SI\|MA\|SO\|LHA |
| MD | IMD\|PVT\|PF |
| LD | DG\|LP |
| AV | VAL\|AD\|LD\|CL |
| MD | VAL\|AV\|AM\|IAD\|PCN\|CL |
| COPY | DCO\|PFL |
| VISam | VISa |
| RSPv | RSPd |
| SSp-ll | |
| MOs | MOp |
| SSp-bfd | SSp-n\|SSp-un |
| RSPv | RSPd |
| GRN | NTS\|IRN\|MARN\|PARN\|PGRNd\|PGRNl |
| VISp | |
| ACAd | MOs |
| ACAv | ACAd |
| SSp-n | SSp-un |
| MOs | |
| AOB | ORBl\|ORBvl\|MOB\|AON |
| VISa | VISam |
| CP | MOp\|AIv\|CLA\|EPd |
| AIv | AId\|CLA\|EPd\|CP |
| BST | LSv\|AM\|RT\|ZI |
| BST | ACB\|LSv\|PVT\|PT\|ADP\|PS |
| LSr | LSv |
| SIM | |
| MOs | |
| BMA | PA\|CEA\|IA\|MEA |
| AHN | ASO\|PVH\|DMH\|PVHd\|VMH\|LHA\|PeF |
| MD | DG\|LP\|AV\|AD\|LD\|CL\|LH |
| VM | VAL\|VPLpc\|VPM\|VPMpc\|PO\|MD\|SM |
| AUDp | AUDd |
| MOs | ACAd |
| VISp | |
| CS | MRN\|DR\|PCG\|NI\|RPO |
| PF | LP\|MD\|CL\|LH\|MRN\|PAG\|APN\|NPC\|C |
| PVH | SMT\|PR\|RE\|PVHd |
| ENTm | VISl\|VISpl\|VISpor\|ENTl |
| ENTm | ENTl\|SUB |
| PO | DG\|PoT\|LP\|POL |
| ACAv | ACAd |
| VISam | VISpm\|VISa |
| ACAd | |
| SUB | VISpor\|ENTl\|ENTm |

| | |
|---|---|
| SSs | SSp-n |
| MD | LP\|PVT\|CL\|PF\|LH |
| SMT | VM\|VPMpc\|MD\|RH\|CM\|PCN |
| SIM | |
| PH | SPFm\|SUM\|PAG |
| VM | VAL\|PO\|MD\|SMT\|PCN\|CL |
| VPM | VPL |
| PH | VM\|VPMpc\|SPFm\|SPA\|IMD\|MD\|PF\|[illegible] |
| GR | CU\|NTS\|DMX |
| MH | CA1\|DG\|LH |
| NTS | CU\|GR\|DMX\|XII |
| IP | SIM\|COPY\|DN |
| MD | PVT\|MH\|LH |
| VPM | VPLpc\|PO |
| PB | CUN\|PPN\|SUT |
| CA3 | SSp-bfd\|VISrl\|CA1\|CA2 |
| ENTl | VISpor\|PERI\|ECT |
| CP | SSp-n\|SSp-bfd\|SSs |
| SSp-ll | SSp-ul |
| PAG | CLI\|DR |
| PRNr | PPN\|PRNc\|SUT\|P5 |
| SSp-bfd | |
| RSPv | RSPd |
| VISpor | VISl\|VISli\|TEa |
| SPVI | SPVC |
| ORBm | PL\|ORBvl\|MOB |
| MEA | NLOT\|BA |
| VISp | |
| MOs | |
| VCO | DCO |
| PRNc | TRN\|NTB\|GRN\|RM |
| AIp | SSs\|VISC\|CLA\|CP |
| V | SUT |
| PAG | VTA\|MRN\|NPC\|RN\|MA3\|EW\|IF\|RL\|C |
| MD | MOs\|ACAd\|PVT\|PT\|MH\|LH |
| SGN | VISp\|VISpm\|PoT\|MG\|LP\|POL |
| PVT | MD\|PT |
| MD | MOp\|SSp-ll\|DG\|IMD\|PVT\|PT\|CM\|LH |
| MD | RSPv\|DG\|PVT\|MH\|LH |
| LSr | LSc |
| AOB | MOB |
| VISrl | SSp-bfd\|VISa |
| LD | LP\|PO |
| PF | DG\|VPMpc\|PO\|MD\|PCN\|CL\|LH |
| LD | |
| MD | |
| SMT | VM\|VPMpc\|SPFm\|MD\|PR\|RE\|RH\|CM |

| | |
|---|---|
| VM | RSPd\|VPMpc\|SPFm\|SMT\|PR\|RE\|PH\|Z |
| MM | SUM\|PMd\|PH |
| MRN | RSPv\|SCm\|RN |
| MDRN | SPVC\|GRN\|IO\|IRN\|LRN\|MARN\|MDRN |
| VMH | AHN\|PeF\|TU |
| MV | SPVO\|PARN\|LAV\|SPIV |
| MV | IC\|MRN\|PPN\|PB\|B\|PCG\|PRNc\|SUT\|L |
| BLA | SSp-n\|SSp-ul\|SSp-un\|CP\|CEA |
| PCG | MRN\|VTN\|PDTg\|PRNc\|CS\|NI\|RPO\|SL |
| MPO | MS\|NDB\|ADP\|AVP\|AVPV\|MEPO |
| MOs | MOp |
| SSp-bfd | SSp-n\|SSp-un |
| MD | VM\|VPMpc\|SMT\|PR\|RE\|CM\|PCN\|DM |
| VISl | VISal\|VISp |
| VISrl | SSp-bfd |
| SSp-m | GU\|AId |
| PCG | MEV\|MRN\|PB\|B\|LC\|LDT\|SLC\|SLD\|MV |
| VMH | PVi\|ARH\|DMH\|AHN\|PVHd\|RCH\|TU |
| VMH | ARH\|DMH\|TMv\|PMv\|PH\|LHA\|TU |
| ORBm | PL\|ILA\|ORBvl |
| CEA | BLA\|BMA\|IA\|MEA |
| LRN | IO\|PGRNl |
| SSp-tr | SSp-ll\|VISa |
| ACAd | MOs |
| ORBm | PL\|ILA\|ORBvl |
| MOp | SSp-ul |
| VPM | GPi\|VAL\|VM\|VPL\|VPLpc\|VPMpc\|PO\| |
| IC | |
| MARN | VII\|GRN\|PGRNl\|PPY |
| MD | VM\|VPMpc\|IMD\|SMT\|PR\|PVT\|RE\|RH |
| VISp | |
| ACAd | ACAv |
| LGd | VPL\|VPM\|IGL\|LGv\|ZI |
| ORBl | MOs\|AId\|AIv\|CLA\|CP |
| ACAv | ACAd |
| ORBl | FRP\|MOs\|ORBvl |
| VISa | VISam |
| VTA | PN\|MRN\|PAG\|RN\|IF\|IPN\|RL |
| CEA | MEA\|GPe\|GPi\|SI\|LHA |
| RSPd | RSPv |
| SLD | MRN\|B\|PCG\|PRNc\|SLC |
| MOs | ACAd |
| SSp-ll | SSp-ul |
| MOs | MOp\|ACAd |
| MOs | |
| MS | LSr\|LSv\|NDB |
| PH | SPFm\|SPA\|IMD\|SUM\|PAG |

| | |
|---|---|
| SPFm | VPMpc\|SPFp\|SPA\|PF\|PH\|PAG |
| ENTl | ENTm |
| SSp-m | SSp-n |
| CA3 | |
| CA3 | SSp-ll\|SSp-ul\|CA1\|CA2 |
| CA3 | SSp-bfd\|CA1\|CA2 |
| LGv | CA3 |
| CA3 | SSp-bfd\|CA1\|CA2 |
| MD | LH |
| PB | CUN\|PPN |
| APN | MRN\|SCm\|RN |
| PL | ACAd\|ILA |
| CP | |
| ENTm | ENTl |
| ENTm | VISl\|VISpor\|ENTl\|SUB |
| RE | AM\|IAM\|IAD\|PVT\|PT\|Xi\|RH\|CM\|RT |
| GPi | GPe\|RT\|STN\|ZI |
| ACAv | ACAd |
| CA1 | CA3\|DG\|FC\|SUB |
| DG | CA3 |
| LHA | BST\|MPO\|AHN\|LPO\|PeF |
| LHA | AHN\|PVHd\|PeF\|TU\|ZI |
| PVT | IAM\|MD\|PT\|CM |
| SCm | SCs |
| SCm | SCs |
| SSp-tr | MOp\|SSp-ll |
| VISl | VISal |
| ILA | PL\|ORBm |
| VISp | |
| MD | IMD\|CM\|PCN\|LH |
| FRP | MOs\|ORBl |
| SSp-ul | MOp |
| VISrl | VISp\|VISa |
| MEA | CEA |
| ILA | ACAv\|PL\|DP |
| SSs | SSp-n |
| CA1 | DG\|SUB\|ProS |
| ARH | PVi\|DMH\|VMH\|ME |
| VMH | ARH\|DMH\|PMv\|TU |
| NDB | SI\|MA\|VLPO\|LPO |
| VISpl | VISp |
| ACAv | MOs\|ACAd |
| ENTl | PERI |
| ENTm | |
| MD | AD\|IAD\|IMD\|CL\|LH |
| ACB | AON\|SI |
| AV | |

| | |
|---|---|
| CA1 | |
| VISli | VISpor |
| VMH | DMH\|PH |
| DCO | VCO |
| DN | IP |
| GRN | IRN |
| CA1 | ProS |
| CA1 | ProS |
| MD | DG\|IMD\|PVT\|CM\|PCN\|MH\|LH |
| ACAd | MOs |
| MARN | PRNc\|GRN\|PGRNl\|PPY\|RM\|RPA |
| ORBm | PL\|ORBvl |
| SSp-bfd | |
| CP | SSp-bfd\|LA |
| ZI | PoT\|SPFp\|MG\|PIL\|MRN\|SNc |
| PGRNl | LRN |
| SO | |
| ARH | DMH\|VMH\|TU\|ME |
| AHN | MPO\|PVpo\|SBPV\|SCH\|VMPO\|MPN\|V |
| PIR | SSs\|GU\|VISC\|AIp\|EPd |
| PMv | ARH\|DMH\|PVp\|TMv\|PMd\|VMH\|PH\| |
| SSp-bfd | |
| VISp | VISa |
| CEA | CP\|MEA\|GPe\|GPi\|SI |
| ACAd | MOs\|ACAv |
| AIp | SSs\|GU\|VISC\|PIR\|EPd |
| SSs | VISC |
| VISl | VISal\|VISp\|VISrl |
| SSs | SSp-n\|SSp-m\|CLA\|CP |
| LHA | PeF\|TU |
| LA | AIp\|EPd |
| ENTm | VISp\|PAR\|PRE\|SUB |
| PARN | AMB\|IRN\|LIN\|LRN\|PGRNl |
| PGRNl | MEV\|PB\|LC\|SLC\|VII\|GRN\|IRN\|MV |
| VISl | VISal |
| SSs | SSp-n |
| SSs | SSp-n\|SSp-bfd |
| PVH | PR\|RE\|DMH\|AHN\|PVHd\|ZI |
| RPO | MRN\|DR\|PCG\|PRNc\|CS\|NI |
| DMX | NTS\|XII |
| DG | |
| DG | |
| PRE | DG\|ENTm\|PAR\|SUB |
| MM | SUM |
| SSp-bfd | SSp-n\|SSp-un |
| AUDp | AUDd |
| VISp | |

| | |
|---|---|
| VISpm | VISam |
| PIR | EPd\|EPv |
| MPN | BST\|PT\|RE\|PVH\|ADP\|AVPV\|MPO\|PD |
| PAG | DR |
| ACAd | MOs |
| RT | VAL\|VPL |
| VPL | VPM\|RT |
| IC | |
| IC | |
| LGd | CA3\|IGL\|LGv |
| LGd | VISa\|VPM\|LP\|PO\|LD\|LGv |
| SCs | SCm |
| SPVC | MDRN\|MDRNd |
| MOp | |
| ZI | SPFp\|LM\|SUM\|LHA\|PSTN\|VTA\|SNc |
| PL | MOs\|ACAv\|ILA\|TT\|DP |
| VISl | VISli |
| ACB | LSr\|LSv |
| DMH | PVi\|TMd\|PMd\|PMv\|VMH\|PH\|ZI |
| SIM | CUL |
| MRN | PAG\|RN\|III\|RL |
| ACAd | MOs |
| ACAv | ACAd |
| RSPv | |
| VISam | VISa |
| MOs | |
| DCO | SUV\|y\|SIM\|COPY\|PFL\|FL\|IP\|DN |
| SCH | PVa\|PVi\|ARH\|SBPV\|AHN\|VMH\|RCH\|[illegible] |
| LHA | MEA\|SI\|PeF\|TU\|ZI |
| SOC | VII |
| PAG | MRN\|IV\|DR |
| CENT | CUL |
| IC | NB\|MRN\|SCm |
| ACAv | ACAd |
| AON | |
| ORBvl | ORBl |
| ENTl | ENTm |
| PRE | PAR\|POST\|APr |
| MOs | |
| MOs | |
| CLI | IF\|IPN\|RL |
| CUL | CENT |
| GRN | IRN\|MARN\|PGRNd |
| AN | SIM |
| SSp-ll | |
| PIL | PoT\|SPFp\|PP\|LP\|POL\|MRN\|SNc |
| TRS | LSc\|LSr\|SF |

| | |
|---|---|
| SSs | SSp-n\|SSp-m |
| ENTl | TEa\|PERI\|ECT |
| VISpor | VISl\|VISpl\|VISli |
| ACB | |
| CP | |
| CP | |
| SSs | SSp-n\|SSp-m\|CP |
| VISp | VISpm\|RSPagl\|RSPd |
| MOp | MOs |
| MOs | |
| SSp-bfd | SSp-n\|SSp-un |
| VISp | |
| VISp | |
| ACAd | MOs |
| ORBvl | FRP\|MOs\|ORBl |
| VISam | VISpm\|VISa |
| RSPv | RSPd |
| ENTm | VISl\|VISpl\|ENTl\|SUB |
| DG | |
| ORBl | ORBvl\|AOB |
| MOs | FRP |
| AHN | AM\|PR\|RE\|RT\|LHA\|PeF\|ZI |
| MPO | BST\|AVP\|PD\|MPN |
| RSPv | |
| SSp-tr | SSp-ll\|RSPagl\|VISa |
| ACAd | ACAv\|PL |
| VISp | |
| RSPv | RSPd |
| MOs | ACAd\|PL |
| SSp-ul | MOp\|SSp-ll |
| VISli | VISl\|TEa |
| MOs | ACAd |
| TEa | VISpor |
| AId | AIv\|CLA |
| MOs | |
| VMH | DMH\|AHN\|PeF\|TU |
| SCs | RSPd\|IC\|SCm |
| PO | LP |
| SSp-m | SSp-n\|SSp-ul\|SSp-un |
| DCO | VCO\|SIM |
| LP | DG\|VPM\|LGd\|PO |
| LGd | CA3\|VPL\|VPM\|LP\|PO\|LD\|RT\|LGv |
| AV | VAL\|AM\|AD\|RT |
| VISC | SSp-bfd\|AIp |
| MOs | MOp |
| PL | ACAv\|ILA |
| GU | MOp\|SSp-m\|AId\|CP |

| | |
|---|---|
| MOs | |
| ORBm | ACAd\|PL\|ILA |
| AV | AD |
| CA1 | CA3 |
| PH | VM\|SPFm\|SMT\|PR\|RE |
| PRE | DG\|ENTm\|PAR\|SUB\|HATA |
| CUL | CENT |
| CA1 | SUB\|ProS |
| CA1 | DG\|SUB\|ProS |
| SCm | MRN\|APN |
| SSp-ll | SSp-tr |
| CP | EPd |
| LRN | IO\|IRN\|MDRN\|MDRNd\|MDRNv\|PGRN |
| LH | DG\|LP\|CL\|MH |
| PAG | MRN\|SCm |
| PB | IC\|CUN\|PPN |
| VISli | VISl\|VISpor\|TEa |
| SSp-ul | MOp |
| VISrl | SSp-bfd\|VISp\|VISa |
| VISp | VISl |
| RO | |
| VTA | SNr\|RR\|MRN\|MT\|SNc |
| CP | |
| RT | GPi\|STN\|ZI |
| MV | NTS\|PGRNd\|PRP |
| NDB | SI\|MA\|AVP\|MPO\|VLPO\|LPO |
| OT | SI |
| ACB | CP\|OT\|SI |
| ENTl | PERI\|ECT |
| SUB | DG\|POST\|ProS |
| DG | |
| DG | |
| ADP | ACAv\|LSc\|LSv\|MS\|BST\|MEPO\|PS |
| DG | CA3 |
| VISp | |
| PPN | CUN\|NLL\|PB |
| LHA | BST\|LPO |
| PH | MM\|SUM\|VTA\|IF\|IPN |
| MD | IMD\|CM\|LH |
| CP | |
| CP | |
| MD | AM\|IAD\|CM\|PCN\|CL\|LH |
| LHA | LM\|MM\|TMv\|PMd\|PMv\|PH |
| SCH | PVa |
| ACAd | MOs |
| ACAd | MOs\|ACAv |
| ORBvl | FRP\|MOs\|ORBl |

| | |
|---|---|
| FRP | MOs\|ORBl |
| MEA | SI\|VPL\|VPM\|LHA |
| ORBm | PL\|ILA |
| VISp | |
| AOB | ORBvl\|MOB\|AON |
| VISrl | SSp-bfd\|VISp |
| MPN | PVH\|PVa\|MPO\|PD\|PVpo\|SBPV\|AHN |
| COPY | IP\|DN |
| SSp-ll | MOp\|SSp-ul |
| ProS | CA1\|SUB |
| SSp-bfd | |
| ENTm | VISl\|ENTl\|SUB |
| PRNc | |
| LGd | VISa\|CA3\|DG\|VPM\|LP\|PO |
| DG | CA3 |
| DG | SUB\|ProS\|MG\|LP\|SGN\|PIL |
| AUDp | AUDd |
| VISam | VISpm\|VISa |
| DG | PoT\|MG\|LP\|SGN |
| ORBl | FRP\|MOs\|ORBvl |
| SSp-bfd | CA1 |
| LP | LGd |
| LGd | |
| LPO | SI\|NDB\|BST\|AVP\|MPO\|PS\|VLPO |
| LGd | RT\|IGL\|IntG\|LGv\|ZI |
| MPO | SI\|BST\|AVP\|PS\|VLPO\|MPN\|LPO |
| RE | PVT\|PT\|PVH\|MEPO |
| PARN | IRN |
| SCm | IC |
| NLOT | |
| NPC | SCm\|PAG\|MPT\|PPT |
| SAG | IC\|NB\|PBG\|MRN |
| VISl | VISp |
| ORBl | FRP\|MOs\|ORBvl |
| VISp | |
| AON | ORBl\|ORBvl\|PIR |
| SUB | ENTm\|ProS |
| PMv | DMH\|PVp\|MM\|SUM\|TMd\|PMd\|PH\|[illegible] |
| MPN | BST\|PVH\|AVP\|AVPV\|MPO\|PD\|PS\|PV[illegible] |
| PB | MV\|CENT |
| MD | LH |
| PL | ILA |
| SSp-m | SSs\|GU |
| ProS | VISli\|CA1\|SUB |
| ILA | PL\|DP |
| ACAv | ACAd\|PL |
| VISam | VISpm\|VISa |

| | |
|---|---|
| SSp-bfd | VISrl |
| VISpm | VISp |
| VISpor | VISl\|VISpl |
| AId | MOp\|AIv |
| MOs | |
| MOs | ACAd |
| VISa | VISam |
| SSs | AUDd |
| AId | MOp\|GU\|AIv\|CLA\|CP |
| AId | MOp\|AIv\|CLA |
| VISp | |
| SSp-ll | SSp-ul |
| VISl | VISp |
| VISal | VISl\|VISp\|VISrl |
| VISam | VISpm |
| VISp | |
| VISp | |
| VISl | VISpl |
| SSp-bfd | |
| VISp | |
| VISl | VISp\|VISpl |
| SSp-ll | SSp-ul |
| SSp-bfd | |
| SSp-bfd | SSp-n\|SSp-un |
| MOs | MOp |
| VISam | VISpm |
| SSp-bfd | |
| MOp | MOs |
| VISrl | SSp-bfd\|VISp\|VISa |
| SSp-n | SSp-bfd\|SSp-un\|SSs |
| SSp-ll | MOp\|SSp-ul |
| MOs | MOp |
| SSs | SSp-n |
| AIp | SSp-n\|SSs\|GU\|VISC\|CLA\|CP |
| MOs | |
| ORBvl | FRP\|PL\|ORBm |
| CA1 | ProS |
| SSs | SSp-n |
| VISpm | VISam\|RSPagl |
| MOs | |
| MOs | |
| SSp-m | SSp-n\|SSp-ul\|SSp-un |
| SSp-ul | MOp |
| SSp-m | SSp-n\|SSs |
| MOs | |
| RSPv | RSPd\|CA3\|DG\|MH\|LH |
| MD | IMD\|PVT\|CM |

| | |
|---|---|
| LGd | VISam\|VISa\|VPM\|PoT\|MG\|LP\|PO\|SG |
| LGd | LP\|LD\|RT\|LGv |
| LHA | VAL\|VM\|RT\|PVHd\|PeF\|ZI |
| LHA | VM\|PH\|PST\|PSTN\|ZI |
| MPO | BST\|PD\|AHN\|MPN\|LPO |
| PVT | IMD\|MD\|PT\|CM |
| LHA | LM\|SUM\|PMd\|PH\|PSTN |
| PPN | MRN\|PB |
| PAG | LDT |
| LM | SUM\|TMv\|PMv\|PH\|LHA\|PSTN |
| ARH | DMH\|PVp\|PMv\|VMH\|TU |
| SSp-ul | |
| VISpor | VISli\|TEa |
| PMv | ARH\|PVp\|TMv\|PMd\|VMH\|PH\|TU |
| SSs | SSp-bfd |
| SSp-tr | SSp-ll\|VISa |
| ORBl | ORBvl |
| DG | RSPv\|CA1\|SUB\|ProS |
| ILA | ACAv\|PL\|DP |
| MOp | MOs |
| ILA | PL\|ORBm |
| RSPv | RSPd |
| VISpm | VISp\|RSPv |
| MOp | |
| SSp-n | SSp-m\|SSs |
| VISp | |
| VISpor | TEa\|ECT\|ENTl |
| VISp | |
| SSp-ul | MOp |
| SSp-tr | SSp-ll\|VISa |
| VISp | RSPagl\|POST |
| MOs | MOp |
| SSp-bfd | SSp-n\|SSp-un |
| AUDp | AUDd\|AUDv |
| RSPd | RSPv |
| PIR | VISC\|AIp\|EPd |
| VISl | VISal |
| ACAd | MOs |
| VISp | VISpl\|RSPagl\|RSPd\|APr |
| SSp-bfd | SSp-n\|SSp-un |
| SSp-m | SSp-ul\|SSp-un |
| ACAv | MOs\|ACAd |
| ENTl | ENTm\|SUB |
| ENTl | CA1\|EPd |
| SSp-ll | SSp-ul\|SSp-tr |
| VISpor | |
| SUB | ENTl\|ENTm\|ProS |

<table>
<tr><td>LD</td><td></td></tr>
<tr><td>DMH</td><td>VM|VPMpc|MD|PCN|ARH|PMv|PH</td></tr>
<tr><td>SI</td><td>MOp|MA|LPO</td></tr>
<tr><td>BST</td><td>LSr|LSv|SF|MS|TRS|AM|PVT|PT|RE|[illegible]</td></tr>
<tr><td>SI</td><td>BMA|AAA|CEA|MEA|MA|LHA</td></tr>
<tr><td>MPO</td><td>ADP|AVP|AVPV|PS|PVpo|MPN</td></tr>
<tr><td>ENTm</td><td></td></tr>
<tr><td>SI</td><td>GPe|BST|LHA|LPO</td></tr>
<tr><td>AHN</td><td>ASO|PVH|PVa|PVpo|SBPV|MPN</td></tr>
<tr><td>ENTm</td><td></td></tr>
<tr><td>SSp-m</td><td>GU</td></tr>
<tr><td>PVT</td><td>IMD|MD|LH</td></tr>
<tr><td>COAp</td><td>BMA|PA</td></tr>
<tr><td>LGv</td><td>RT|IGL|IntG|ZI</td></tr>
<tr><td>IMD</td><td>MD|PVT|CM</td></tr>
<tr><td>DP</td><td>ACAv|PL|ILA|TT</td></tr>
<tr><td>PMv</td><td>DMH|PVp|TMd|TMv|PMd|VMH|PH|[illegible]</td></tr>
<tr><td>CA1</td><td>COAp|TR|BLA|BMA|PA</td></tr>
<tr><td>CUL</td><td>SIM</td></tr>
<tr><td>PF</td><td>LP|MD|CL|MRN</td></tr>
<tr><td>VISpm</td><td>VISam</td></tr>
<tr><td>APN</td><td></td></tr>
<tr><td>PVHd</td><td>RE|PVH|PVi|ZI</td></tr>
<tr><td>ZI</td><td></td></tr>
<tr><td>IC</td><td>PAG|CENT</td></tr>
<tr><td>IPN</td><td>VTA|PN|MRN|MA3|IF|RL|CS|PRNr</td></tr>
<tr><td>VISli</td><td>VISl|VISpl|VISpor</td></tr>
<tr><td>ACAv</td><td>ACAd</td></tr>
<tr><td>AId</td><td>SSp-m|GU|AIp|AIv|CLA</td></tr>
<tr><td>SSp-tr</td><td>SSp-ll|RSPagl|VISa</td></tr>
<tr><td>VISrl</td><td>SSp-bfd</td></tr>
<tr><td>VISpl</td><td>VISl|VISpor</td></tr>
<tr><td>SSp-m</td><td>SSp-n|SSs</td></tr>
<tr><td>ACAv</td><td>ACAd</td></tr>
<tr><td>PAG</td><td>MRN|DR</td></tr>
<tr><td>ENTl</td><td>SUB</td></tr>
<tr><td>DG</td><td>RSPd|RSPv</td></tr>
<tr><td>VISp</td><td></td></tr>
<tr><td>RT</td><td>GPi|VAL|VPL</td></tr>
<tr><td>SNr</td><td>SNc</td></tr>
<tr><td>MEA</td><td>COAp</td></tr>
<tr><td>PIR</td><td>EPd|EPv</td></tr>
<tr><td>GRN</td><td>MDRN|MDRNv|NR|XII|RO</td></tr>
<tr><td>MG</td><td>SGN</td></tr>
<tr><td>PAG</td><td>DR|LDT</td></tr>
<tr><td>ENTm</td><td>ENTl|PAR|PRE|SUB</td></tr>
<tr><td>ACB</td><td></td></tr>
</table>

| | |
|---|---|
| PIR | EPv |
| SI | CEA\|MEA\|GPe\|GPi |
| PAG | MRN\|EW\|IV\|RL\|DR |
| LSr | ACAv |
| PAG | SUM\|PH\|VTA\|MRN\|IF\|IPN |
| AHN | ASO\|MPO\|SBPV\|SCH\|MPN\|RCH |
| PARN | P5\|SPVO\|IRN |
| IO | GRN\|MARN\|MDRN\|MDRNv\|NR\|XII\|R |
| MOs | FRP |
| MH | DG\|PVT\|LH |
| BST | LSv\|AM\|PT\|RT |
| VAL | VM\|VPL\|LD\|RT |
| GPe | GPi\|VAL\|VPL\|VPM\|RT |
| ENTl | ENTm |
| SIM | |
| POST | VISp\|SUB\|APr |
| RN | MRN |
| NLOT | BMA\|AAA\|CEA\|MEA\|SI |
| PF | MRN\|APN\|NPC |
| MOB | |
| MV | DCO\|ICB\|LAV\|SPIV\|SUV\|x\|y\|IP |
| SGN | DG\|PoT\|MG\|LP\|POL |
| CS | MRN\|PAG\|AT\|DR\|PCG\|NI\|RPO |
| PRNc | SOC\|P5\|PRNr\|VII\|IRN |
| LSr | LSc\|SF |
| MV | NTS\|GRN\|LAV\|SPIV |
| GR | |
| MH | RSPv\|CA1\|DG\|LH |
| PVH | PVT\|PT\|RE\|Xi\|PVpo\|MPN |
| MD | PVT\|MH\|LH |
| PVHd | PR\|PVH\|ZI |
| PH | VM\|DMH\|PMd\|PMv\|LHA |
| VISp | |
| PB | MEV\|MRN\|PAG\|CUN\|PPN\|PCG\|LC\|L[illegible] |
| PB | PSV\|SUT\|V\|P5\|I5\|PARN\|MV |
| LH | MD\|PVT\|CL\|PF\|MH |
| MD | DG\|LP\|CL\|LH |
| MPN | BST\|PT\|RE\|PVH\|PVpo |
| VTA | LHA\|PSTN\|STN\|SNr\|SNc |
| PIL | PoT\|SPFp\|PP\|POL\|MRN |
| CA3 | DG |
| DG | CA1\|CA2\|CA3 |
| CA1 | |
| SSp-n | SSp-m\|SSs |
| ACAv | MOs\|ACAd |
| CP | SSp-bfd |
| PVT | IMD\|MD |

| | |
|---|---|
| PB | MEV\|B\|PCG\|LC\|LDT |
| VISl | VISal |
| ENTl | VISpor\|ENTm |
| ENTl | VISpor\|PERI\|ECT |
| ORBvl | ORBm |
| AN | |
| MD | IMD\|CM\|PCN |
| CA1 | |
| CA3 | |
| DG | CA1\|CA3 |
| RT | SSp-bfd\|SSp-tr\|CA1\|CA2\|IGL\|LGv |
| CA3 | DG |
| DG | CA3 |
| VII | IRN\|PARN |
| LP | LGd\|SGN |
| CM | AM\|IAM\|IAD\|SMT\|RE\|RH\|PCN |
| ZI | |
| PAG | SCm\|III\|EW\|DR |
| PAG | SCm\|DR |
| SSs | SSp-n\|SSp-bfd |
| SSp-m | |
| SSp-n | |
| VISp | |
| VISl | VISp |
| VISpm | VISam\|RSPagl |
| CA3 | CA1\|CA2\|VPL\|LGd\|LP\|PO\|LD |
| CP | |
| PAG | MRN\|LDT |
| LSc | LSr\|SF\|TRS |
| LSr | LSc\|LSv\|SF\|MS\|TRS |
| AHN | ASO\|PVH\|PVi\|DMH\|SBPV\|PVHd\|VMH |
| PO | VPM\|PoT\|MG\|LP |
| IF | VTA\|PN\|IPN\|RL\|CLI\|CS\|PRNr |
| VPMpc | VM\|SPFm\|MD\|PCN\|PF\|PH |
| LGd | CA3\|VPM\|MG\|LP\|PO\|IGL\|LGv |
| CLI | AT\|IF\|IPN\|TRN\|CS\|RM |
| VII | SOC\|PRNc\|IRN |
| GRN | IO\|MARN\|PGRNd\|NR\|PRP\|MV\|XII\|RO |
| PG | TRN |
| VPL | VPM\|PO\|LD\|RT |
| BST | AM\|RT\|LHA\|ZI |
| ENTl | PERI\|ECT |
| MPN | PVH\|ADP\|AVPV\|MPO\|PD\|PVpo |
| PB | MEV\|PAG\|PCG\|LC\|LDT\|MV\|CENT |
| LGv | CA3\|VPM\|LGd\|LP\|IGL |
| ENTm | |
| IRN | GRN\|MARN\|PARN\|PGRNd\|PGRNl |

| | |
|---|---|
| LPO | NDB\|BST\|MPO\|AHN\|LHA |
| NTS | DMX |
| PAG | SCs\|SCm\|DR\|LDT\|CENT |
| CUL | SIM\|PFL\|FL |
| PRE | PAR |
| MDRN | GRN\|IRN\|LRN\|MDRNv |
| PH | SPFm |
| PH | VTA\|MRN\|IF |
| PRE | VISp\|POST |
| PVH | BST\|RE\|MPO\|PD\|SBPV\|AHN\|MPN\|ZI |
| MARN | GRN\|PPY\|RM\|RPA |
| MRN | PAG\|RN |
| SI | MA\|NDB\|BST\|LPO |
| SI | MOp\|MOs\|LPO |
| COAp | COAa\|PAA\|BLA\|BMA\|PA |
| MARN | GRN\|RM |
| IPN | VTA\|PN\|EW\|IF\|RL |
| CP | CEA\|GPe |
| SNr | RR\|MRN\|SNc\|NLL\|PRNr |
| PB | MV\|SUV\|CENT |
| SUT | RSPagl\|RSPd\|PSV\|PB\|V\|P5\|PC5\|PRNr |
| XII | NTS\|DMX\|MDRN\|MDRNd |
| LSv | LSc\|LSr\|BST |
| MEA | NLOT\|SO\|LHA |
| MS | LSr\|LSv\|SF\|TRS\|MEPO |
| AUDp | |
| SSp-m | SSp-n\|SSp-ul\|SSp-un |
| SSp-ul | MOp |
| VISrl | SSp-bfd\|VISa |
| SSs | SSp-n\|SSp-m |
| SSp-m | SSp-n\|SSs |
| VMH | SMT\|PR\|PVi\|ARH\|DMH\|AHN\|PVHd\|P |
| MV | B\|PCG\|SLD |
| CM | VAL\|IAM\|IMD\|MD\|SMT\|RH\|PCN |
| SSp-tr | SSp-ll |
| ACAv | MOs\|ACAd |
| ORBl | FRP\|MOs\|ORBvl |
| SSp-m | |
| SSp-bfd | |
| VTA | MRN\|PAG\|RN |
| SCs | SCm |
| SBPV | SCH\|VMPO\|AHN\|MPN\|RCH |
| IP | ICB\|SUV\|NOD\|FN\|VeCB |
| IP | ICB\|SUV\|VeCB |
| CUL | CENT |
| DMX | NTS\|XII |
| DMH | VM\|MD\|SMT\|PR\|RE\|Xi\|CM\|PVH\|PVi |

| | |
|---|---|
| SSp-tr | SSp-ll\|RSPagl\|VISa |
| LHA | PVHd\|PH\|PST\|PSTN\|PeF\|ZI |
| DN | DCO\|IP |
| PFL | DCO\|AN\|FL\|DN |
| POST | VISp\|RSPagl\|RSPd\|APr\|IC |
| MEA | |
| AON | AOB |
| MM | SUM\|PH\|VTA |
| MM | SUM\|PMd\|PH\|LHA |
| PA | COAp\|BMA |
| BST | SSp-ul\|SSp-un\|CP\|RT\|LHA |
| MRN | SNr\|VTA\|MT\|SNc |
| SI | CP\|ACB\|OT\|MA |
| IC | |
| ZI | LGv\|SubG |
| IPN | PH\|VTA\|IF\|RL |
| MOB | |
| SUM | MM\|PH\|VTA\|IPN |
| TU | DMH\|TMv\|PMv\|VMH\|PH\|LHA |
| PAG | SCm\|EW\|DR |
| LRN | PGRNl |
| CEA | GPe\|GPi\|SI |
| DN | COPY\|IP |
| FL | PFL |
| PVHd | RE\|PVH\|DMH\|AHN\|PeF\|ZI |
| GPi | ZI |
| SCm | SCs\|PAG |
| BST | MPO\|PD\|MPN |
| SI | CP\|MA\|NDB\|LPO |
| PAR | ENTm\|APr |
| AHN | SBPV\|SCH\|VMPO\|RCH\|TU |
| MEA | COAa\|COAp\|BMA\|PA |
| PT | AM\|IAD\|MD\|PVT |
| APN | RSPagl\|DG\|POL\|PF\|MRN |
| MPN | BST\|ADP\|AVP\|AVPV\|MPO\|PD\|PS\|PV[illegible] |
| NOD | MV |
| IRN | GRN\|LIN\|LRN\|MARN\|PARN\|PGRNl |
| PIR | VISC\|AIp\|EPd |
| SPVC | MDRN\|MDRNd |
| DG | CA1\|CA2\|CA3 |
| BLA | COAa\|COAp\|PAA\|EPv\|BMA\|PA\|CP\|CI |
| ZI | LP\|PSTN\|STN\|SNr |
| LHA | VM\|PST\|PSTN\|STN\|ZI |
| CA1 | VISa\|DG |
| CA1 | SSp-bfd\|VISrl\|CA2\|DG |
| DG | CA1\|CA3 |
| EPd | AIp\|PIR\|LA\|CP |

| | |
|---|---|
| PB | MEV\|PAG\|B\|PCG\|LC |
| LD | VPL\|LP\|PO |
| APN | MRN\|SCm |
| SCm | SCs\|IC |
| IC | PB |
| PT | PVT\|RE |
| SSp-n | SSp-bfd\|SSp-un |
| MOs | MOp |
| AUDp | AUDd |
| VISp | |
| ACAd | MOs |
| PL | ILA\|ORBm |
| MEA | NLOT\|COAa\|BMA\|BA |
| BST | BAC |
| AHN | BST\|MPO\|PVHd\|LHA\|PeF\|ZI |
| ENTl | ENTm\|SUB\|ProS |
| SSp-n | SSp-bfd\|SSp-un |
| ZI | |
| DMH | PR\|PVi\|ARH\|PVp\|TMd\|PMv\|VMH\|PH |
| VM | VPMpc\|SMT\|PCN\|PH |
| RSPv | RSPd\|DG\|MH\|LH |
| SSp-m | |
| SSp-ul | |
| MM | SUM\|PH\|PAG\|IF |
| SCm | POST\|SCs |
| SCm | SCs\|PAG |
| CP | |
| IP | DCO\|SUV\|NOD\|VeCB |
| PB | MEV\|PCG\|LC\|LDT\|CENT\|FN |
| ORBl | FRP\|MOs\|ORBvl |
| CP | GPe |
| CP | GPe |
| RSPv | RSPd |
| VISp | |
| BST | PT |
| CP | |
| EPd | VISC\|AIp\|PIR\|LA |
| DG | CA1\|CA2\|CA3 |
| VMH | PVi\|ARH\|DMH\|AHN |
| ENTm | |
| PAR | ENTm |
| RSPv | |
| VISp | |
| PARN | NTS\|SPVO\|IRN\|ISN |
| NLL | PPN\|PB\|SUT\|PRNr |
| MEA | SO |
| SSp-m | |

| | |
|---|---|
| SSp-ul | MOp |
| SSp-n | SSp-bfd |
| VISam | RSPagl |
| VISl | VISli |
| ORBvl | ORBl |
| PF | SPA\|IMD\|MD\|PVT\|CL\|LH\|PAG |
| PRNr | PRNc\|TRN\|CS |
| LD | VAL\|AV |
| CP | ACB |
| ACB | |
| ADP | LSv\|MS\|MPO\|PS |
| CP | |
| CLI | VTA\|PN\|IF\|IPN |
| VAL | VM |
| MOp | MOs\|CP |
| SLD | PCG\|PRNc\|NI\|MV |
| PIR | SSp-m\|EPd\|CP |
| LA | AIp\|EPd\|CP |
| PIR | EPd\|EPv\|BLA\|IA |
| CS | PAG\|PRNc\|PRNr |
| NTS | GR\|DMX\|XII |
| RE | PR\|Xi\|PVH\|PVHd\|ZI |
| III | MRN\|PAG\|EW |
| PB | MEV\|MRN\|PAG\|PPN\|LDT |
| SPVC | |
| VPM | VPL |
| PB | MEV\|PAG\|PCG\|LC\|CENT |
| PVT | IMD\|MD |
| VMH | PVi\|ARH\|DMH\|PVp\|TMd\|TMv\|PMd\|I |
| AId | AIv |
| ORBl | FRP\|MOs\|ORBvl |
| MOs | MOp |
| VISp | |
| SSp-n | SSp-bfd\|SSp-un |
| VISp | POST |
| AUDd | SSs\|AUDp |
| MOs | ACAd |
| MOp | MOs |
| CP | SSp-bfd |
| CA3 | |
| VTA | RR\|MRN\|PAG\|RN\|MA3\|EW\|IF\|RL\|CL |
| VMH | PVi\|ARH\|SCH\|RCH\|TU\|ME |
| SNr | PP\|PIL\|MRN\|SNc |
| SNr | VTA\|RR\|MRN\|MT\|SNc |
| AUDd | SSs\|AUDp\|AUDv |
| ACAv | ACAd |
| VISa | |

| | |
|---|---|
| TRN | |
| TRN | PG\|PRNc\|PRNr\|RM |
| SSp-bfd | SSp-n\|SSp-un |
| AUD and MOs | AUDd |
| MDRN | SPVC\|AMB\|GRN\|IRN\|LRN\|MARN\|MD |
| BLA | LA\|BMA |
| LHA | SO\|AHN\|PeF\|RCH\|TU |
| CP | SSp-m\|GU\|AId\|AIv\|CLA\|EPd |
| AIp | AId\|AIv |
| PIR | AIv\|EPd |
| CLA | GU\|AIp\|EPd\|CP |
| PIR | AIv |
| ENTl | ENTm |
| SCs | SCm |
| VISC | SSs |
| ZI | PH\|VTA |
| BMA | COAa\|PA\|CEA\|IA\|MEA |
| DG | VISp\|CA3 |
| PRE | PAR |
| SCs | SCm |
| LH | DG\|AD\|LD\|MD\|CL\|MH |
| LH | DG\|MD\|CL\|MH |
| VISp | |
| SSp-bfd | |
| ZI | |
| GPe | |
| PB | MEV\|B\|PCG\|LC\|LDT\|SLC\|MV\|VeCB |
| MOp | |
| SSp-n | SSp-bfd |
| AIv | GU\|AId\|AIp\|PIR\|CLA\|EPd |
| CP | GU\|AId\|AIp\|AIv\|CLA\|EPd |
| CP | SSp-m\|AId\|AIv\|CLA |
| AIv | SSp-m\|GU\|AId\|PIR |
| LGd | VPM\|IGL\|LGv |
| AHN | BST\|MPO\|MPN\|LHA\|PeF |
| LHA | PVHd\|PH\|PSTN\|PeF\|ZI |
| PRNc | SOC\|V\|P5\|PC5\|PRNr\|IRN\|PARN |
| ENTl | |
| VISam | VISpm\|VISa |
| SUB | VISl\|VISp\|VISpl\|ENTm\|PAR\|PRE |
| SSp-n | SSp-m\|SSp-un\|SSs |
| RSPv | RSPd |
| LA | AIp\|EPd\|BLA\|CP |
| ACAd | MOs\|ACAv |
| ENTl | VISpor\|PERI |
| PIR | |
| ENTm | ENTl\|PAR\|PRE\|SUB |

| | |
|---|---|
| SUM | LM\|MM\|TMv\|PMv |
| TMv | LM\|MM\|SUM\|PMv |
| PMv | LM\|MM\|SUM\|TMv |
| LM | MM\|SUM\|TMv\|PMv |
| SSs | VISC |
| RSPv | |
| VISrl | SSp-bfd\|VISa |
| ORBl | AON |
| BST | |
| SSp-bfd | SSp-n\|SSp-un |
| SCm | MRN\|PAG\|CUN |
| AHN | BST\|PT\|RE\|PVH\|MPO\|SBPV\|MPN\|PV |
| MOs | MOp |
| MOs | ACAd |
| LGv | VISa\|CA1\|CA2\|MG\|LGd\|LP\|SGN\|IGL\| |
| RSPv | RSPd |
| SUB | DG\|ENTl\|ProS |
| ENTm | |
| VISpor | |
| LSc | LSr |
| VISl | VISli\|VISpor |
| VISl | VISpl |
| RSPv | RSPd |
| RE | IAM\|SMT\|PR\|Xi\|RH\|CM |
| VM | VAL\|LD\|MD\|SMT\|PCN\|CL\|ZI |
| LDT | MRN\|PCG |
| PB | MEV\|PPN |
| PVH | SBPV\|AHN\|PVHd |
| CEA | BLA\|IA\|MEA |
| PB | |
| PAG | SCs\|MRN\|SCm\|RN\|III\|EW\|IV\|Pa4\|RL |
| RN | VTA\|MRN |
| STN | GPi\|VPL\|VPM\|PO\|LD\|ZI |
| PA | COAp\|BMA |
| PB | MEV\|PCG\|LC\|SLC\|MV |
| PT | PVT |
| PL | MOs\|ORBm\|ORBvl |
| NI | MRN\|DR\|PDTg\|PCG\|CS\|RPO |
| DR | PAG |
| CA1 | VISp\|VISrl\|CA2\|CA3\|HATA\|PA |
| MEA | SSp-tr\|COAp\|CA1\|CA3\|DG\|PA |
| SI | SSp-ul\|AAA\|CEA\|MEA\|MA\|LHA |
| MEA | BMA\|CEA\|IA |
| SUB | ProS |
| PIR | AIv\|CLA\|EPd |
| LH | LP\|CL\|PF |
| PH | SUM |

| | |
|---|---|
| VPM | LP\|PO\|LD |
| DN | DCO\|SIM\|AN\|COPY\|PFL |
| PB | SUT\|LAV\|MV\|SUV\|y |
| VMH | AHN\|PVHd\|PeF\|RCH\|TU |
| ACAv | |
| SSs | |
| MS | LSr\|LSv\|SF\|TRS\|PT\|MEPO |
| CEA | BMA\|AAA\|IA\|MEA\|GPe |
| CUN | IC\|PPN\|PB |
| PIR | EPv |
| RT | BST\|VM\|AV\|AM\|PR\|PT\|ZI |
| PH | PVi\|DMH\|MM\|SUM\|TMd\|PMd\|PMv |
| NTS | GR\|DMX |
| XII | CU\|NTS\|DMX\|IRN\|MDRN\|MDRNd\|M |
| ENTm | ENTl |
| VISl | VISp\|VISpl\|VISli\|VISpor\|ENTm |
| TT | |
| PH | MD\|PCN\|DMH |
| GR | CU\|NTS\|MDRN\|MDRNd |
| ENTm | ENTl |
| MM | LM\|SUM\|LHA |
| MV | NTS\|GRN\|IRN\|PGRNd\|PRP |
| MD | PVT\|MH\|LH |
| ENTl | ENTm\|SUB |
| CS | IPN\|NI\|RPO |
| VISp | RSPagl |
| MEA | CEA\|SI |
| PB | MEV\|PAG\|PCG\|MV\|SUV |
| STN | SI\|VPL\|VPM\|LHA\|ZI |
| PB | IC\|PAG\|CUN |
| MOs | |
| CEA | BLA\|BMA\|IA\|MEA\|GPe\|SI |
| PSV | SPVO\|VII\|PARN |
| NTB | SOC\|PRNc\|GRN\|MARN\|PGRNl\|PPY |
| VISp | VISpl\|RSPagl\|RSPd\|POST\|APr |
| MOs | FRP |
| VISp | VISl |
| SSs | |
| MRN | SCm\|PAG\|APN |
| SUM | MM\|PH\|LHA\|VTA\|MRN |
| PVH | BST\|PT\|RE |
| NTS | CU\|GR\|DMX |
| DR | PAG |
| ACAv | MOs\|ACAd\|PL |
| ENTm | COAp\|ENTl\|PA |
| MOs | |
| MOs | |

| | |
|---|---|
| SSp-ll | MOp |
| ENTm | ENTl\|SUB |
| ENTm | ENTl\|SUB |
| ENTl | CA1\|ENTm\|SUB\|ProS |
| ENTm | ENTl\|SUB |
| VISp | VISl\|VISpl\|RSPagl |
| ACAv | ACAd |
| VISp | |
| ORBl | FRP\|ORBvl |
| VISa | VISam\|VISp\|VISpm\|VISrl |
| RSPv | RSPd\|SUB |
| SSp-tr | SSp-ll |
| VISp | VISl\|VISpl |
| PIL | PoT\|SPFp\|PP\|MG\|POL\|MRN |
| SPVC | |
| ARH | DMH\|PVp\|TMd\|PMv\|VMH |
| PG | TRN |
| MPN | MPO\|PVpo\|SBPV\|SCH\|VMPO\|AHN\|R |
| SO | MEA |
| MOp | MOs |
| SSp-ll | MOp |
| ACAd | MOs |
| AUDd | AUDp |
| VISp | |
| MOs | |
| VISp | |
| MOp | |
| MOs | MOp |
| VISp | |
| VISpl | VISp\|POST\|APr |
| MOp | SSp-ll |
| MOs | MOp |
| MOs | MOp |
| MOs | FRP |
| MOs | MOp |
| VISp | |
| ENTm | VISp\|DG\|PAR\|PRE\|SUB |
| VISp | |
| VISp | |
| VISp | |
| VISp | |
| VISp | |
| ENTl | VISpor\|ENTm |
| ENTm | ENTl\|SUB\|ProS |
| SUB | ProS |
| LGd | |
| LGd | CA1\|CA3 |

| | |
|---|---|
| LGd | |
| LGd | LP |
| ENTl | VISpor |
| VISp | |
| VISp | |
| VISp | |
| VISp | |
| VISp | |
| VISp | |
| VISp | |
| VISp | |
| VISp | |
| VISp | |
| VISp | |
| BMA | CA1\|CA2\|CA3\|PA\|IA |
| VMH | ARH\|DMH\|PMv\|TU |
| VMH | DMH\|PMv\|PH\|TU |
| MOs | |
| MOs | |
| MOs | |
| CEA | MEA |
| PMv | ARH\|DMH\|PVp\|TMv\|VMH\|PH\|TU |
| MOs | |
| MOs | |
| DMH | PVi\|ARH\|PVp\|TMd\|PMv\|VMH\|PH |
| MOs | |
| MOs | FRP\|PL |
| MOs | |
| ZI | SSp-bfd\|CA1\|CA2\|CA3\|SI\|RT\|LGv\|Sub |
| GPi | AAA\|CEA\|MEA\|SI\|RT\|LHA |
| CEA | GPi\|SI |
| VMH | TU |
| MOs | FRP |
| TU | ARH\|VMH\|LHA\|PeF\|RCH |
| TU | VMH\|RCH |
| MOs | |
| MOs | FRP |
| RT | GPi\|SI\|VPL\|ZI |
| MOs | |
| MOs | |
| MOs | |
| MOs | |
| MOs | FRP |
| MOs | |
| MOs | |
| MOs | |
| MOs | |

| | |
|---|---|
| MOs | |
| MOs | |
| MOs | |
| MOs | |
| MOs | |
| MOs | FRP |
| MOs | |
| MOs | FRP\|ORBl |
| MOs | |
| MOs | |
| MOs | |
| MOs | |
| VISp | |
| VISp | |
| ACAd | MOs\|ACAv |
| SI | AAA\|MA\|NDB\|LHA\|LPO |
| VPM | CA3\|LGd\|LP |
| ORBm | PL\|ORBvl |
| VISp | |
| AV | VAL\|AM\|AD\|MD |
| VISl | VISal\|VISp\|VISpl\|VISli\|VISpor |
| VISl | VISp\|VISli |
| VISl | |
| VISp | |
| VISp | |
| VISp | |
| LGd | LGv |
| DR | PAG\|IV |
| MV | PB\|B\|PCG\|PRNc\|SUT\|P5\|LC\|SLC\|SLD |
| LGd | MG\|LP\|IGL |
| MD | VAL\|AV\|AM\|IAD\|IMD\|CM\|PCN\|CL |
| VISp | |
| VISp | |
| VISpm | VISp |
| VISl | VISli |
| VISpm | VISp\|RSPagl\|RSPd\|RSPv |
| VISl | |
| ORBm | MOs\|ACAd\|PL\|ILA\|ORBvl |
| ACAd | MOs\|ACAv |
| VISp | |
| VISpm | VISp |
| SI | GPi\|RT\|LHA |
| SI | GPe\|MA\|BST\|LHA |
| LP | MG\|LGd\|SGN |
| VISpm | VISp\|RSPagl |
| AM | AV\|AD\|IAD\|MD |
| LPO | SI\|NDB\|BST\|PS |

| | |
|---|---|
| VISal | VISl\|VISli |
| VISrl | SSp-bfd |
| VISam | VISpm\|VISa |
| VISpm | VISp\|RSPagl |
| VISa | VISam\|VISpm |
| VISp | VISl |
| VISl | VISal\|VISp\|VISli |
| VISl | VISp |
| VISp | |
| VISal | AUDpo\|VISl\|VISli |
| VISp | |
| VISrl | SSp-bfd |
| VISp | |
| ORBvl | MOs\|ACAd\|PL\|ORBm |
| ACAd | |
| VISp | RSPagl\|RSPd |
| RSPd | VISp\|RSPagl |
| VISp | |
| VISal | VISl |
| VISam | VISpm\|RSPagl\|VISa |
| VISp | |
| VISal | AUDpo\|VISl\|VISli |
| VISam | VISa |
| VISam | VISa\|CA1\|DG\|ProS |
| VISp | |
| VISp | |
| VISp | |
| VISp | |
| VISp | |
| VISp | RSPagl\|RSPd |
| VISp | RSPagl\|RSPd |
| VISp | |
| SSp-bfd | |
| ACAd | ACAv |
| VISp | |
| VISp | |
| VISrl | SSp-bfd\|VISp\|VISa |
| VISp | |
| VISp | |
| RSPd | VISp\|RSPagl |
| VISp | RSPagl |
| VISal | VISl |
| VISa | VISam\|VISpm |
| VISl | VISp |
| AUDp | AUDd\|AUDpo |
| MD | LH |
| RSPagl | VISam\|VISp\|VISpm\|RSPd\|RSPv |

| | |
|---|---|
| VISpor | VISli |
| VISli | VISal\|VISl\|TEa |
| AUDpo | AUDp\|VISal\|VISli\|TEa |
| VISpl | VISp |
| AUDpo | VISal\|VISli\|TEa |
| SSp-bfd | |
| VISp | |
| VISl | VISp |
| RSPd | RSPagl\|RSPv |
| AUDp | AUDd\|AUDpo |
| RSPagl | VISam |
| VISam | VISa |
| VISrl | VISp\|VISa |
| AUDpo | AUDp\|VISal\|TEa |
| VISp | |
| LHA | PSTN\|TU |
| AUDp | AUDpo\|AUDv\|TEa |
| VISpm | VISam\|RSPagl |
| ACAv | ACAd |
| LP | PO\|LD |
| VISpm | VISam\|RSPagl\|RSPd\|RSPv |
| AM | IAM\|IAD\|MD\|PT |
| PH | DMH\|MM\|SUM\|PMd\|PMv\|LHA |
| AUDp | AUDpo\|AUDv\|TEa |
| VISrl | VISp |
| VISal | AUDd\|AUDpo\|VISl\|VISli\|TEa |
| SSs | SSp-bfd |
| IC | |
| SSp-bfd | SSs\|AUDd |
| ACAd | MOs\|ACAv |
| VISp | VISam\|VISpm\|VISa |
| RSPd | RSPv |
| AUDpo | AUDd\|AUDp\|VISal\|TEa |
| RSPagl | VISam |
| RSPagl | RSPd\|RSPv |
| VISpor | |
| RSPd | RSPagl |
| RSPv | RSPd |
| VISp | VISpl\|RSPagl |
| RSPagl | VISam\|VISpm |
| VISpl | VISl\|VISp\|RSPagl\|RSPd\|PAR\|APr |
| VISpor | VISli |
| RSPagl | VISpm\|RSPd |
| VISp | |
| VISp | |
| VISp | VISrl |
| VISal | VISl\|VISli |

| | |
|---|---|
| TEa | AUDpo\|VISli\|VISpor |
| VISam | VISa |
| AUDpo | AUDd\|AUDp\|TEa |
| VISpm | VISam\|VISp\|VISa |
| VISpl | VISl\|VISli\|VISpor |
| VISpl | VISp |
| SSp-bfd | SSs |
| VISpor | VISl\|VISpl\|TEa |
| VISpor | ENTl |
| RSPagl | VISam\|RSPd\|VISa |
| GPe | GPi\|SI |
| SSp-bfd | |
| VISpor | ENTl |
| VISpor | VISli\|TEa |
| SSp-bfd | |
| VISpor | VISli |
| VISpm | VISam |
| VISa | SSp-tr |
| SSp-n | SSp-m |
| VISp | VISam\|VISpm\|VISa\|VISrl |
| VISl | VISpl\|VISli\|VISpor |
| VISp | |
| VISam | VISa |
| VISrl | SSp-bfd\|VISp\|VISa\|CA1 |
| AUDp | AUDd\|AUDpo |
| VISli | VISl\|VISpor\|TEa |
| RSPagl | VISam\|RSPd |
| TEa | AUDpo\|VISal\|VISli |
| VISa | VISp\|VISrl |
| VISpor | VISli |
| VISpm | VISam\|RSPagl |
| SSp-bfd | |
| VISp | |
| LGd | VPM\|PoT\|MG\|LP\|PO\|SGN |
| PL | ACAd\|ILA\|ORBm\|ORBvl |
| ACAd | ACAv |
| SI | AAA\|CEA\|MEA\|GPe\|GPi\|MA\|LHA\|LP( |
| SI | AAA\|CEA\|GPe\|MA\|LHA |
| LGd | VPM\|PoT\|MG\|LP\|PO\|SGN |
| LGd | |
| LGd | VPM\|LP |
| VISl | VISp |
| VISp | VISl |
| VISp | VISl\|CA1\|SUB\|ProS |
| VISp | |
| VISp | RSPagl\|RSPd |
| VISp | |

| | |
|---|---|
| ORBvl | PL\|ORBl\|ORBm\|MOB\|AON |
| VISl | VISal\|VISp |
| RSPd | VISp\|VISpm\|RSPagl\|RSPv\|POST\|APr |
| VISp | |
| VISl | VISal\|VISp |
| MD | AM\|IAM\|IAD\|PVT\|PT\|RE\|RH\|CM |
| RSPagl | VISpm |
| VISp | |
| VISl | |
| VISpm | RSPagl\|RSPd\|RSPv |
| ILA | MOs\|ACAd\|PL\|ORBm |
| RSPv | VISpm\|RSPagl\|RSPd |
| PAG | MRN\|PPN\|PB\|LDT |
| MOs | MOp\|ACAd\|ACAv |
| VISp | VISl |
| VISp | |
| VISl | VISal\|VISli |
| VISp | |
| VISp | |
| VISp | |
| VISpm | VISp\|RSPagl\|RSPd\|RSPv |
| VISp | RSPagl\|RSPd |
| VISp | |
| VISp | |
| ACAd | ACAv |
| RSPv | RSPd\|SUB |
| VISpm | VISam\|VISp |
| VISp | |
| VISpm | RSPagl |
| VISp | |
| VISl | VISp |
| VISp | |
| VISp | |
| VISp | |
| LP | VPM\|LGd\|PO |
| NDB | SI\|MA\|MPO\|VLPO\|LPO |
| VISp | |
| VISp | |
| AM | AV\|IAD\|MD\|PT |
| VISp | RSPagl |
| VISp | |
| VISp | |
| VISal | AUDpo\|VISl\|VISli |
| VISal | VISl\|VISrl |
| VISp | VISam\|VISpm\|VISa\|VISrl |
| VISp | VISl\|VISpl\|RSPagl |
| VISp | |

| | |
|---|---|
| VISp | |
| VISp | RSPagl\|RSPd |
| RSPd | VISp\|RSPagl |
| VISal | AUDpo\|VISl\|VISli |
| VISp | |
| VISli | AUDpo\|VISal\|VISl\|TEa |
| VISrl | SSp-bfd\|VISp\|VISa |
| VISam | VISpm\|VISa |
| SSp-bfd | VISrl |
| VISam | VISpm\|VISa |
| VISrl | SSp-bfd\|VISp |
| VISal | AUDd\|AUDpo\|VISl\|VISli\|TEa |
| LDT | MEV\|MRN\|PAG\|PB\|PCG\|LC |
| RSPd | VISpm\|RSPagl\|RSPv |
| VISrl | SSp-bfd |
| VISp | |
| VISam | VISpm\|VISa |
| VISpm | VISp\|RSPagl\|RSPd\|RSPv |
| ACAd | ACAv |
| VISal | SSp-bfd\|VISl\|VISp\|VISli |
| VISp | RSPagl\|RSPd |
| VISpor | VISl\|VISpl\|VISli |
| VISpl | VISl\|VISp |
| RSPagl | VISam\|RSPd |
| RSPagl | VISam |
| AUDpo | AUDp\|TEa |
| VISpm | VISam\|RSPagl\|RSPd |
| VISa | VISam\|VISp\|VISpm\|VISrl |
| RSPagl | VISam\|RSPd |
| VISam | VISpm\|VISa |
| VISp | RSPagl\|RSPd |
| RSPd | RSPagl\|RSPv |
| AUDpo | AUDd |
| VISam | VISpm\|RSPagl\|RSPd\|RSPv |
| VISl | VISpl\|VISli\|VISpor |
| AUDd | AUDp\|AUDpo |
| VMH | PR\|PVi\|ARH\|DMH\|TMd\|PMv\|PH |
| VISp | VISpl |
| RSPagl | VISam\|RSPd\|VISa |
| RSPagl | VISam\|VISpm\|RSPd\|RSPv |
| VISpor | VISli\|TEa |
| RSPagl | VISam\|VISpm\|RSPd |
| AUDpo | AUDp\|VISal\|VISli\|TEa |
| VISp | VISpm |
| VISp | RSPagl |
| ACAd | MOs\|ACAv\|PL |
| VISp | |

<table>
<tr><td>VISp</td><td>VISpl|VISpm|RSPagl|RSPd</td></tr>
<tr><td>AUDpo</td><td>AUDd|AUDp|VISal|TEa</td></tr>
<tr><td>VISam</td><td>RSPagl|VISa</td></tr>
<tr><td>VISpor</td><td>TEa</td></tr>
<tr><td>VISpm</td><td>VISam|VISp|RSPagl</td></tr>
<tr><td>VISpl</td><td>VISp|RSPagl|RSPd|APr</td></tr>
<tr><td>RSPd</td><td>RSPagl</td></tr>
<tr><td>VISpor</td><td>VISli|TEa</td></tr>
<tr><td>RSPv</td><td>RSPd</td></tr>
<tr><td>SI</td><td>CEA|GPe|GPi</td></tr>
<tr><td>VISli</td><td>VISpor|TEa</td></tr>
<tr><td>VISa</td><td>VISam|VISp|VISpm|RSPagl</td></tr>
<tr><td>VISp</td><td>VISpm|VISa|VISrl|CA1</td></tr>
<tr><td>RSPagl</td><td>RSPd|RSPv</td></tr>
<tr><td>SSp-bfd</td><td>VISam|VISp|VISpm|VISa|VISrl</td></tr>
<tr><td>VISp</td><td>VISpl|RSPagl|RSPd|POST|APr</td></tr>
<tr><td>ENTm</td><td>VISl|VISp|VISpl|VISpor|ENTl|PAR</td></tr>
<tr><td>SSp-bfd</td><td></td></tr>
<tr><td>VISam</td><td>VISpm|VISa</td></tr>
<tr><td>AUDpo</td><td>AUDp|VISpor|TEa|CA1</td></tr>
<tr><td>SSp-bfd</td><td></td></tr>
<tr><td>VISpor</td><td>TEa</td></tr>
<tr><td>DR</td><td>PAG|IV|CLI</td></tr>
<tr><td>VISpor</td><td>TEa</td></tr>
<tr><td>VISli</td><td>VISl|VISpor|TEa</td></tr>
<tr><td>VISa</td><td>SSp-bfd|SSp-tr|VISrl</td></tr>
<tr><td>VISpor</td><td>VISl|VISli|TEa</td></tr>
<tr><td>VISa</td><td>VISam|VISpm</td></tr>
<tr><td>AIp</td><td>SSs|PIR|EPd</td></tr>
<tr><td>VISa</td><td>SSp-bfd|SSp-tr|VISrl</td></tr>
<tr><td>SSp-bfd</td><td></td></tr>
<tr><td>VISpor</td><td>TEa|SUB|ProS</td></tr>
<tr><td>VISrl</td><td>SSp-bfd|VISp|VISa</td></tr>
<tr><td>SSp-bfd</td><td>VISrl</td></tr>
<tr><td>CP</td><td>AId|AIv|CLA|EPd</td></tr>
<tr><td>CP</td><td>AIv|EPd|CLA|ORBl</td></tr>
<tr><td>CLA</td><td>MOp|AId|AIv</td></tr>
<tr><td>CLA</td><td>AId|AIv</td></tr>
<tr><td>PIR</td><td>EPd|AIp|EPv|CLA|SSs|SSp-n</td></tr>
<tr><td>SSs</td><td>AIp|CLA|EPd|VISC|CP|GU|SSp-n|SSp-</td></tr>
<tr><td>EPd</td><td>AIv|CLA|PIR|SSp-m|AId|AIp|SSs|CP|[illegible]</td></tr>
<tr><td>CLA</td><td>EPd|CLA|AIv|AId|AIp|PIR|CP</td></tr>
<tr><td>CLA</td><td>EPd|AIp|GU</td></tr>
<tr><td>CLA</td><td>EPd|AIp</td></tr>
<tr><td>CLA</td><td>AIp|GU|EPd|VISC|CP</td></tr>
<tr><td>SSs</td><td>SSp-n|AIp|EPd|CLA|VISC</td></tr>
<tr><td>AIv</td><td>AId|CLA|MOp|CP</td></tr>
</table>

| | |
|---|---|
| AIv | AId\|CLA\|MOp\|EPd\|CP\|PIR\|GU |

**QW-analysis-spreadsheet**
Copy_of_normalized_projection_volume-brain_subcortical_for_CLA
Copy_of_normalized_projection_volume-brain_subcortical_for_CLA
Copy_of_normalized_projection_volume-brain_subcortical_for_CLA
Copy_of_normalized_projection_volume-brain_subcortical_for_CLA
Copy_of_normalized_projection_volume-brain_subcortical_for_CLA
Copy_of_normalized_projection_volume-brain_subcortical_for_CLA
Copy_of_normalized_projection_volume-brain_subcortical_for_CLA, best_injection_list
Copy_of_normalized_projection_volume-brain_subcortical_for_CLA
Copy_of_normalized_projection_volume-brain_subcortical_for_CLA
Copy_of_normalized_projection_volume-brain_subcortical_for_CLA
Copy_of_normalized_projection_volume-brain_subcortical_for_CLA
Copy_of_normalized_projection_volume-brain_subcortical_for_CLA
Copy_of_normalized_projection_volume-brain_subcortical_for_CLA
Copy_of_normalized_projection_volume-brain_subcortical_for_CLA
Copy_of_normalized_projection_volume-brain_subcortical_for_CLA, best_injection_list
Copy_of_normalized_projection_volume-brain_subcortical_for_CLA
Copy_of_normalized_projection_volume-brain_subcortical_for_CLA
Copy_of_normalized_projection_volume-brain_subcortical_for_CLA
Copy_of_normalized_projection_volume-brain_subcortical_for_CLA, best_injection_list
Copy_of_normalized_projection_volume-brain_subcortical_for_CLA
Copy_of_normalized_projection_volume-brain_subcortical_for_CLA
Copy_of_normalized_projection_volume-brain_subcortical_for_CLA
Copy_of_normalized_projection_volume-brain_subcortical_for_CLA
Copy_of_normalized_projection_volume-brain_subcortical_for_CLA
Copy_of_normalized_projection_volume-brain_subcortical_for_CLA
Copy_of_normalized_projection_volume-brain_subcortical_for_CLA
Copy_of_normalized_projection_volume-brain_subcortical_for_CLA
Copy_of_normalized_projection_volume_brain_for_CLA, best_injection_list
Copy_of_normalized_projection_volume_brain_for_CLA, best_injection_list
Copy_of_normalized_projection_volume-brain_subcortical_for_CLA, best_injection_list
Copy_of_normalized_projection_volume-brain_subcortical_for_CLA
Copy_of_normalized_projection_volume-brain_subcortical_for_CLA
Copy_of_normalized_projection_volume-brain_subcortical_for_CLA
Copy_of_normalized_projection_volume-brain_subcortical_for_CLA
Copy_of_normalized_projection_volume-brain_subcortical_for_CLA
Copy_of_normalized_projection_volume-brain_subcortical_for_CLA
Copy_of_normalized_projection_volume-brain_subcortical_for_CLA
Copy_of_normalized_projection_volume-brain_subcortical_for_CLA
Copy_of_normalized_projection_volume-brain_subcortical_for_CLA
Copy_of_normalized_projection_volume-brain_subcortical_for_CLA
Copy_of_normalized_projection_volume-brain_subcortical_for_CLA
Copy_of_normalized_projection_volume-brain_subcortical_for_CLA
Copy_of_normalized_projection_volume-brain_subcortical_for_CLA, best_injection_list
Copy_of_normalized_projection_volume-brain_subcortical_for_CLA, best_injection_list
Copy_of_normalized_projection_volume-brain_subcortical_for_CLA
Copy_of_normalized_projection_volume-brain_subcortical_for_CLA

Copy_of_normalized_projection_volume-brain_subcortical_for_CLA
Copy_of_normalized_projection_volume-brain_subcortical_for_CLA
Copy_of_normalized_projection_volume-brain_subcortical_for_CLA
Copy_of_normalized_projection_volume-brain_subcortical_for_CLA
Copy_of_normalized_projection_volume-brain_subcortical_for_CLA
Copy_of_normalized_projection_volume-brain_subcortical_for_CLA
Copy_of_normalized_projection_volume-brain_subcortical_for_CLA
Copy_of_normalized_projection_volume-brain_subcortical_for_CLA
Copy_of_normalized_projection_volume-brain_subcortical_for_CLA
Copy_of_normalized_projection_volume-brain_subcortical_for_CLA, best_injection_list
Copy_of_normalized_projection_volume-brain_subcortical_for_CLA, best_injection_list
Copy_of_normalized_projection_volume-brain_subcortical_for_CLA
Copy_of_normalized_projection_volume-brain_subcortical_for_CLA
Copy_of_normalized_projection_volume-brain_subcortical_for_CLA, best_injection_list
Copy_of_normalized_projection_volume-brain_subcortical_for_CLA
Copy_of_normalized_projection_volume-brain_subcortical_for_CLA
Copy_of_normalized_projection_volume-brain_subcortical_for_CLA
Copy_of_normalized_projection_volume-brain_subcortical_for_CLA
Copy_of_normalized_projection_volume-brain_subcortical_for_CLA
Copy_of_normalized_projection_volume-brain_subcortical_for_CLA
Copy_of_normalized_projection_volume-brain_subcortical_for_CLA
Copy_of_normalized_projection_volume-brain_subcortical_for_CLA
Copy_of_normalized_projection_volume-brain_subcortical_for_CLA
Copy_of_normalized_projection_volume-brain_subcortical_for_CLA
Copy_of_normalized_projection_volume-brain_subcortical_for_CLA
Copy_of_normalized_projection_volume-brain_subcortical_for_CLA
Copy_of_normalized_projection_volume-brain_subcortical_for_CLA
Copy_of_normalized_projection_volume-brain_subcortical_for_CLA
Copy_of_normalized_projection_volume-brain_subcortical_for_CLA
Copy_of_normalized_projection_volume-brain_subcortical_for_CLA
Copy_of_normalized_projection_volume-brain_subcortical_for_CLA
Copy_of_normalized_projection_volume-brain_subcortical_for_CLA
Copy_of_normalized_projection_volume-brain_subcortical_for_CLA
Copy_of_normalized_projection_volume-brain_subcortical_for_CLA
Copy_of_normalized_projection_volume-brain_subcortical_for_CLA
Copy_of_normalized_projection_volume-brain_subcortical_for_CLA
Copy_of_normalized_projection_volume-brain_subcortical_for_CLA, best_injection_list
Copy_of_normalized_projection_volume-brain_subcortical_for_CLA
Copy_of_normalized_projection_volume-brain_subcortical_for_CLA
Copy_of_normalized_projection_volume-brain_subcortical_for_CLA
Copy_of_normalized_projection_volume-brain_subcortical_for_CLA
Copy_of_normalized_projection_volume-brain_subcortical_for_CLA
Copy_of_normalized_projection_volume-brain_subcortical_for_CLA
Copy_of_normalized_projection_volume-brain_subcortical_for_CLA
Copy_of_normalized_projection_volume-brain_subcortical_for_CLA
Copy_of_normalized_projection_volume-brain_subcortical_for_CLA
Copy_of_normalized_projection_volume_brain_for_CLA, best_injection_list

Copy_of_normalized_projection_volume-brain_subcortical_for_CLA
Copy_of_normalized_projection_volume-brain_subcortical_for_CLA
Copy_of_normalized_projection_volume-brain_subcortical_for_CLA
Copy_of_normalized_projection_volume-brain_subcortical_for_CLA
Copy_of_normalized_projection_volume-brain_subcortical_for_CLA
Copy_of_normalized_projection_volume-brain_subcortical_for_CLA
Copy_of_normalized_projection_volume_brain_for_CLA, best_injection_list
Copy_of_normalized_projection_volume_brain_for_CLA
Copy_of_normalized_projection_volume-brain_subcortical_for_CLA
Copy_of_normalized_projection_volume-brain_subcortical_for_CLA
Copy_of_normalized_projection_volume-brain_subcortical_for_CLA
Copy_of_normalized_projection_volume_brain_for_CLA
Copy_of_normalized_projection_volume-brain_subcortical_for_CLA
Copy_of_normalized_projection_volume-brain_subcortical_for_CLA
Copy_of_normalized_projection_volume-brain_subcortical_for_CLA
Copy_of_normalized_projection_volume-brain_subcortical_for_CLA
Copy_of_normalized_projection_volume-brain_subcortical_for_CLA
Copy_of_normalized_projection_volume-brain_subcortical_for_CLA
Copy_of_normalized_projection_volume-brain_subcortical_for_CLA
Copy_of_normalized_projection_volume-brain_subcortical_for_CLA
Copy_of_normalized_projection_volume-brain_subcortical_for_CLA
Copy_of_normalized_projection_volume-brain_subcortical_for_CLA
Copy_of_normalized_projection_volume-brain_subcortical_for_CLA
Copy_of_normalized_projection_volume-brain_subcortical_for_CLA
Copy_of_normalized_projection_volume-brain_subcortical_for_CLA
Copy_of_normalized_projection_volume-brain_subcortical_for_CLA
Copy_of_normalized_projection_volume-brain_subcortical_for_CLA, best_injection_list
Copy_of_normalized_projection_volume-brain_subcortical_for_CLA
Copy_of_normalized_projection_volume-brain_subcortical_for_CLA
Copy_of_normalized_projection_volume-brain_subcortical_for_CLA
Copy_of_normalized_projection_volume-brain_subcortical_for_CLA
Copy_of_normalized_projection_volume_brain_for_CLA
Copy_of_normalized_projection_volume-brain_subcortical_for_CLA
Copy_of_normalized_projection_volume-brain_subcortical_for_CLA
Copy_of_normalized_projection_volume-brain_subcortical_for_CLA
Copy_of_normalized_projection_volume_brain_for_CLA
Copy_of_normalized_projection_volume-brain_subcortical_for_CLA
Copy_of_normalized_projection_volume-brain_subcortical_for_CLA
Copy_of_normalized_projection_volume-brain_subcortical_for_CLA
Copy_of_normalized_projection_volume-brain_subcortical_for_CLA
Copy_of_normalized_projection_volume_brain_for_CLA, best_injection_list
Copy_of_normalized_projection_volume-brain_subcortical_for_CLA
Copy_of_normalized_projection_volume-brain_subcortical_for_CLA
Copy_of_normalized_projection_volume_brain_for_CLA
Copy_of_normalized_projection_volume_brain_for_CLA, best_injection_list
Copy_of_normalized_projection_volume-brain_subcortical_for_CLA
Copy_of_normalized_projection_volume-brain_subcortical_for_CLA

Copy_of_normalized_projection_volume-brain_subcortical_for_CLA
Copy_of_normalized_projection_volume-brain_subcortical_for_CLA
Copy_of_normalized_projection_volume-brain_subcortical_for_CLA, best_injection_list
Copy_of_normalized_projection_volume-brain_subcortical_for_CLA
Copy_of_normalized_projection_volume-brain_subcortical_for_CLA, best_injection_list
Copy_of_normalized_projection_volume-brain_subcortical_for_CLA, best_injection_list
Copy_of_normalized_projection_volume_brain_for_CLA
Copy_of_normalized_projection_volume-brain_subcortical_for_CLA, best_injection_list
Copy_of_normalized_projection_volume-brain_subcortical_for_CLA
Copy_of_normalized_projection_volume_brain_for_CLA
Copy_of_normalized_projection_volume-brain_subcortical_for_CLA
Copy_of_normalized_projection_volume_brain_for_CLA
Copy_of_normalized_projection_volume-brain_subcortical_for_CLA
Copy_of_normalized_projection_volume-brain_subcortical_for_CLA
Copy_of_normalized_projection_volume_brain_for_CLA
Copy_of_normalized_projection_volume-brain_subcortical_for_CLA
Copy_of_normalized_projection_volume-brain_subcortical_for_CLA
Copy_of_normalized_projection_volume-brain_subcortical_for_CLA
Copy_of_normalized_projection_volume_brain_for_CLA, best_injection_list
Copy_of_normalized_projection_volume-brain_subcortical_for_CLA
Copy_of_normalized_projection_volume-brain_subcortical_for_CLA
Copy_of_normalized_projection_volume-brain_subcortical_for_CLA
Copy_of_normalized_projection_volume_brain_for_CLA, best_injection_list
Copy_of_normalized_projection_volume-brain_subcortical_for_CLA
Copy_of_normalized_projection_volume-brain_subcortical_for_CLA
Copy_of_normalized_projection_volume-brain_subcortical_for_CLA
Copy_of_normalized_projection_volume_brain_for_CLA, best_injection_list
Copy_of_normalized_projection_volume-brain_subcortical_for_CLA
Copy_of_normalized_projection_volume-brain_subcortical_for_CLA
Copy_of_normalized_projection_volume_brain_for_CLA, best_injection_list
Copy_of_normalized_projection_volume-brain_subcortical_for_CLA
Copy_of_normalized_projection_volume-brain_subcortical_for_CLA
Copy_of_normalized_projection_volume-brain_subcortical_for_CLA
Copy_of_normalized_projection_volume-brain_subcortical_for_CLA
Copy_of_normalized_projection_volume_brain_for_CLA
Copy_of_normalized_projection_volume-brain_subcortical_for_CLA
Copy_of_normalized_projection_volume-brain_subcortical_for_CLA
Copy_of_normalized_projection_volume-brain_subcortical_for_CLA
Copy_of_normalized_projection_volume-brain_subcortical_for_CLA
Copy_of_normalized_projection_volume-brain_subcortical_for_CLA
Copy_of_normalized_projection_volume-brain_subcortical_for_CLA
Copy_of_normalized_projection_volume-brain_subcortical_for_CLA
Copy_of_normalized_projection_volume-brain_subcortical_for_CLA
Copy_of_normalized_projection_volume-brain_subcortical_for_CLA
Copy_of_normalized_projection_volume-brain_subcortical_for_CLA, best_injection_list
Copy_of_normalized_projection_volume-brain_subcortical_for_CLA
Copy_of_normalized_projection_volume-brain_subcortical_for_CLA

Copy_of_normalized_projection_volume-brain_subcortical_for_CLA
Copy_of_normalized_projection_volume-brain_subcortical_for_CLA
Copy_of_normalized_projection_volume-brain_subcortical_for_CLA
Copy_of_normalized_projection_volume-brain_subcortical_for_CLA
Copy_of_normalized_projection_volume-brain_subcortical_for_CLA
Copy_of_normalized_projection_volume-brain_subcortical_for_CLA
Copy_of_normalized_projection_volume-brain_subcortical_for_CLA
Copy_of_normalized_projection_volume-brain_subcortical_for_CLA
Copy_of_normalized_projection_volume-brain_subcortical_for_CLA
Copy_of_normalized_projection_volume-brain_subcortical_for_CLA
Copy_of_normalized_projection_volume-brain_subcortical_for_CLA
Copy_of_normalized_projection_volume-brain_subcortical_for_CLA
Copy_of_normalized_projection_volume-brain_subcortical_for_CLA
Copy_of_normalized_projection_volume-brain_subcortical_for_CLA
Copy_of_normalized_projection_volume-brain_subcortical_for_CLA
Copy_of_normalized_projection_volume-brain_subcortical_for_CLA
Copy_of_normalized_projection_volume-brain_subcortical_for_CLA, best_injection_list
Copy_of_normalized_projection_volume-brain_subcortical_for_CLA
Copy_of_normalized_projection_volume-brain_subcortical_for_CLA
Copy_of_normalized_projection_volume-brain_subcortical_for_CLA
Copy_of_normalized_projection_volume-brain_subcortical_for_CLA
Copy_of_normalized_projection_volume-brain_subcortical_for_CLA
Copy_of_normalized_projection_volume-brain_subcortical_for_CLA
Copy_of_normalized_projection_volume-brain_subcortical_for_CLA
Copy_of_normalized_projection_volume-brain_subcortical_for_CLA
Copy_of_normalized_projection_volume-brain_subcortical_for_CLA
Copy_of_normalized_projection_volume-brain_subcortical_for_CLA
Copy_of_normalized_projection_volume-brain_subcortical_for_CLA
Copy_of_normalized_projection_volume-brain_subcortical_for_CLA
Copy_of_normalized_projection_volume_brain_for_CLA
Copy_of_normalized_projection_volume-brain_subcortical_for_CLA
Copy_of_normalized_projection_volume-brain_subcortical_for_CLA
Copy_of_normalized_projection_volume-brain_subcortical_for_CLA, best_injection_list
Copy_of_normalized_projection_volume-brain_subcortical_for_CLA
Copy_of_normalized_projection_volume-brain_subcortical_for_CLA
Copy_of_normalized_projection_volume-brain_subcortical_for_CLA
Copy_of_normalized_projection_volume-brain_subcortical_for_CLA
Copy_of_normalized_projection_volume-brain_subcortical_for_CLA
Copy_of_normalized_projection_volume-brain_subcortical_for_CLA
Copy_of_normalized_projection_volume-brain_subcortical_for_CLA
Copy_of_normalized_projection_volume-brain_subcortical_for_CLA, best_injection_list
Copy_of_normalized_projection_volume-brain_subcortical_for_CLA
Copy_of_normalized_projection_volume-brain_subcortical_for_CLA
Copy_of_normalized_projection_volume-brain_subcortical_for_CLA
Copy_of_normalized_projection_volume-brain_subcortical_for_CLA
Copy_of_normalized_projection_volume_brain_for_CLA
Copy_of_normalized_projection_volume-brain_subcortical_for_CLA

Copy_of_normalized_projection_volume-brain_subcortical_for_CLA
Copy_of_normalized_projection_volume_brain_for_CLA
Copy_of_normalized_projection_volume-brain_subcortical_for_CLA
Copy_of_normalized_projection_volume-brain_subcortical_for_CLA
Copy_of_normalized_projection_volume-brain_subcortical_for_CLA
Copy_of_normalized_projection_volume-brain_subcortical_for_CLA, best_injection_list
Copy_of_normalized_projection_volume-brain_subcortical_for_CLA
Copy_of_normalized_projection_volume-brain_subcortical_for_CLA
Copy_of_normalized_projection_volume-brain_subcortical_for_CLA, best_injection_list
Copy_of_normalized_projection_volume-brain_subcortical_for_CLA
Copy_of_normalized_projection_volume-brain_subcortical_for_CLA, best_injection_list
Copy_of_normalized_projection_volume-brain_subcortical_for_CLA
Copy_of_normalized_projection_volume-brain_subcortical_for_CLA
Copy_of_normalized_projection_volume-brain_subcortical_for_CLA
Copy_of_normalized_projection_volume-brain_subcortical_for_CLA
Copy_of_normalized_projection_volume_brain_for_CLA, best_injection_list
Copy_of_normalized_projection_volume-brain_subcortical_for_CLA
Copy_of_normalized_projection_volume-brain_subcortical_for_CLA
Copy_of_normalized_projection_volume-brain_subcortical_for_CLA
Copy_of_normalized_projection_volume-brain_subcortical_for_CLA
Copy_of_normalized_projection_volume-brain_subcortical_for_CLA, best_injection_list
Copy_of_normalized_projection_volume-brain_subcortical_for_CLA
Copy_of_normalized_projection_volume-brain_subcortical_for_CLA
Copy_of_normalized_projection_volume-brain_subcortical_for_CLA
Copy_of_normalized_projection_volume-brain_subcortical_for_CLA
Copy_of_normalized_projection_volume-brain_subcortical_for_CLA, best_injection_list
Copy_of_normalized_projection_volume-brain_subcortical_for_CLA
Copy_of_normalized_projection_volume-brain_subcortical_for_CLA
Copy_of_normalized_projection_volume-brain_subcortical_for_CLA
Copy_of_normalized_projection_volume-brain_subcortical_for_CLA
Copy_of_normalized_projection_volume-brain_subcortical_for_CLA
Copy_of_normalized_projection_volume-brain_subcortical_for_CLA
Copy_of_normalized_projection_volume-brain_subcortical_for_CLA
Copy_of_normalized_projection_volume-brain_subcortical_for_CLA, best_injection_list
Copy_of_normalized_projection_volume-brain_subcortical_for_CLA
Copy_of_normalized_projection_volume-brain_subcortical_for_CLA
Copy_of_normalized_projection_volume-brain_subcortical_for_CLA
Copy_of_normalized_projection_volume-brain_subcortical_for_CLA
Copy_of_normalized_projection_volume-brain_subcortical_for_CLA
Copy_of_normalized_projection_volume-brain_subcortical_for_CLA
Copy_of_normalized_projection_volume-brain_subcortical_for_CLA
Copy_of_normalized_projection_volume-brain_subcortical_for_CLA
Copy_of_normalized_projection_volume-brain_subcortical_for_CLA
Copy_of_normalized_projection_volume-brain_subcortical_for_CLA
Copy_of_normalized_projection_volume-brain_subcortical_for_CLA
Copy_of_normalized_projection_volume-brain_subcortical_for_CLA
Copy_of_normalized_projection_volume-brain_subcortical_for_CLA

Copy_of_normalized_projection_volume-brain_subcortical_for_CLA
Copy_of_normalized_projection_volume-brain_subcortical_for_CLA
Copy_of_normalized_projection_volume-brain_subcortical_for_CLA
Copy_of_normalized_projection_volume-brain_subcortical_for_CLA
Copy_of_normalized_projection_volume-brain_subcortical_for_CLA
Copy_of_normalized_projection_volume-brain_subcortical_for_CLA
Copy_of_normalized_projection_volume-brain_subcortical_for_CLA, best_injection_list
Copy_of_normalized_projection_volume-brain_subcortical_for_CLA
Copy_of_normalized_projection_volume-brain_subcortical_for_CLA
Copy_of_normalized_projection_volume-brain_subcortical_for_CLA
Copy_of_normalized_projection_volume-brain_subcortical_for_CLA
Copy_of_normalized_projection_volume-brain_subcortical_for_CLA
Copy_of_normalized_projection_volume-brain_subcortical_for_CLA
Copy_of_normalized_projection_volume-brain_subcortical_for_CLA
Copy_of_normalized_projection_volume-brain_subcortical_for_CLA
Copy_of_normalized_projection_volume-brain_subcortical_for_CLA
Copy_of_normalized_projection_volume-brain_subcortical_for_CLA
Copy_of_normalized_projection_volume-brain_subcortical_for_CLA
Copy_of_normalized_projection_volume-brain_subcortical_for_CLA
Copy_of_normalized_projection_volume-brain_subcortical_for_CLA
Copy_of_normalized_projection_volume-brain_subcortical_for_CLA
Copy_of_normalized_projection_volume-brain_subcortical_for_CLA
Copy_of_normalized_projection_volume-brain_subcortical_for_CLA
Copy_of_normalized_projection_volume-brain_subcortical_for_CLA
Copy_of_normalized_projection_volume-brain_subcortical_for_CLA, best_injection_list
Copy_of_normalized_projection_volume-brain_subcortical_for_CLA
Copy_of_normalized_projection_volume-brain_subcortical_for_CLA
Copy_of_normalized_projection_volume-brain_subcortical_for_CLA
Copy_of_normalized_projection_volume-brain_subcortical_for_CLA
Copy_of_normalized_projection_volume-brain_subcortical_for_CLA
Copy_of_normalized_projection_volume-brain_subcortical_for_CLA
Copy_of_normalized_projection_volume-brain_subcortical_for_CLA
Copy_of_normalized_projection_volume-brain_subcortical_for_CLA
Copy_of_normalized_projection_volume-brain_subcortical_for_CLA
Copy_of_normalized_projection_volume-brain_subcortical_for_CLA
Copy_of_normalized_projection_volume-brain_subcortical_for_CLA
Copy_of_normalized_projection_volume-brain_subcortical_for_CLA
Copy_of_normalized_projection_volume-brain_subcortical_for_CLA
Copy_of_normalized_projection_volume-brain_subcortical_for_CLA
Copy_of_normalized_projection_volume-brain_subcortical_for_CLA
Copy_of_normalized_projection_volume-brain_subcortical_for_CLA
Copy_of_normalized_projection_volume-brain_subcortical_for_CLA
Copy_of_normalized_projection_volume-brain_subcortical_for_CLA
Copy_of_normalized_projection_volume-brain_subcortical_for_CLA
Copy_of_normalized_projection_volume-brain_subcortical_for_CLA
Copy_of_normalized_projection_volume-brain_subcortical_for_CLA
Copy_of_normalized_projection_volume-brain_subcortical_for_CLA

Copy_of_normalized_projection_volume_brain_for_CLA
Copy_of_normalized_projection_volume_brain_for_CLA
Copy_of_normalized_projection_volume_brain_for_CLA, best_injection_list
Copy_of_normalized_projection_volume_brain_for_CLA, best_injection_list
Copy_of_normalized_projection_volume_brain_for_CLA
Copy_of_normalized_projection_volume_brain_for_CLA
Copy_of_normalized_projection_volume_brain_for_CLA
Copy_of_normalized_projection_volume-brain_subcortical_for_CLA
Copy_of_normalized_projection_volume_brain_for_CLA, best_injection_list
Copy_of_normalized_projection_volume_brain_for_CLA, best_injection_list
Copy_of_normalized_projection_volume_brain_for_CLA
Copy_of_normalized_projection_volume_brain_for_CLA
Copy_of_normalized_projection_volume_brain_for_CLA
Copy_of_normalized_projection_volume_brain_for_CLA, best_injection_list
Copy_of_normalized_projection_volume_brain_for_CLA
Copy_of_normalized_projection_volume_brain_for_CLA
Copy_of_normalized_projection_volume-brain_subcortical_for_CLA
Copy_of_normalized_projection_volume-brain_subcortical_for_CLA
Copy_of_normalized_projection_volume-brain_subcortical_for_CLA, best_injection_list
Copy_of_normalized_projection_volume-brain_subcortical_for_CLA
Copy_of_normalized_projection_volume-brain_subcortical_for_CLA, best_injection_list
Copy_of_normalized_projection_volume-brain_subcortical_for_CLA
Copy_of_normalized_projection_volume-brain_subcortical_for_CLA
Copy_of_normalized_projection_volume-brain_subcortical_for_CLA
Copy_of_normalized_projection_volume-brain_subcortical_for_CLA
Copy_of_normalized_projection_volume-brain_subcortical_for_CLA
Copy_of_normalized_projection_volume-brain_subcortical_for_CLA
Copy_of_normalized_projection_volume-brain_subcortical_for_CLA
Copy_of_normalized_projection_volume-brain_subcortical_for_CLA
Copy_of_normalized_projection_volume-brain_subcortical_for_CLA
Copy_of_normalized_projection_volume_brain_for_CLA
Copy_of_normalized_projection_volume-brain_subcortical_for_CLA
Copy_of_normalized_projection_volume-brain_subcortical_for_CLA
Copy_of_normalized_projection_volume-brain_subcortical_for_CLA
Copy_of_normalized_projection_volume_brain_for_CLA
Copy_of_normalized_projection_volume_brain_for_CLA
Copy_of_normalized_projection_volume_brain_for_CLA
Copy_of_normalized_projection_volume_brain_for_CLA
Copy_of_normalized_projection_volume_brain_for_CLA
Copy_of_normalized_projection_volume-brain_subcortical_for_CLA
Copy_of_normalized_projection_volume-brain_subcortical_for_CLA
Copy_of_normalized_projection_volume-brain_subcortical_for_CLA
Copy_of_normalized_projection_volume-brain_subcortical_for_CLA
Copy_of_normalized_projection_volume_brain_for_CLA, best_injection_list
Copy_of_normalized_projection_volume_brain_for_CLA
Copy_of_normalized_projection_volume_brain_for_CLA
Copy_of_normalized_projection_volume_brain_for_CLA, best_injection_list

Copy_of_normalized_projection_volume-brain_subcortical_for_CLA
Copy_of_normalized_projection_volume-brain_subcortical_for_CLA
Copy_of_normalized_projection_volume_brain_for_CLA
Copy_of_normalized_projection_volume_brain_for_CLA
Copy_of_normalized_projection_volume_brain_for_CLA
Copy_of_normalized_projection_volume_brain_for_CLA
Copy_of_normalized_projection_volume-brain_subcortical_for_CLA
Copy_of_normalized_projection_volume_brain_for_CLA
Copy_of_normalized_projection_volume-brain_subcortical_for_CLA
Copy_of_normalized_projection_volume-brain_subcortical_for_CLA
Copy_of_normalized_projection_volume-brain_subcortical_for_CLA
Copy_of_normalized_projection_volume-brain_subcortical_for_CLA
Copy_of_normalized_projection_volume-brain_subcortical_for_CLA, best_injection_list
Copy_of_normalized_projection_volume-brain_subcortical_for_CLA
Copy_of_normalized_projection_volume-brain_subcortical_for_CLA, best_injection_list
Copy_of_normalized_projection_volume-brain_subcortical_for_CLA
Copy_of_normalized_projection_volume-brain_subcortical_for_CLA
Copy_of_normalized_projection_volume-brain_subcortical_for_CLA
Copy_of_normalized_projection_volume-brain_subcortical_for_CLA, best_injection_list
Copy_of_normalized_projection_volume-brain_subcortical_for_CLA
Copy_of_normalized_projection_volume-brain_subcortical_for_CLA
Copy_of_normalized_projection_volume-brain_subcortical_for_CLA
Copy_of_normalized_projection_volume-brain_subcortical_for_CLA
Copy_of_normalized_projection_volume-brain_subcortical_for_CLA
Copy_of_normalized_projection_volume-brain_subcortical_for_CLA, best_injection_list
Copy_of_normalized_projection_volume-brain_subcortical_for_CLA
Copy_of_normalized_projection_volume-brain_subcortical_for_CLA
Copy_of_normalized_projection_volume-brain_subcortical_for_CLA
Copy_of_normalized_projection_volume-brain_subcortical_for_CLA
Copy_of_normalized_projection_volume-brain_subcortical_for_CLA
Copy_of_normalized_projection_volume-brain_subcortical_for_CLA
Copy_of_normalized_projection_volume_brain_for_CLA, best_injection_list
Copy_of_normalized_projection_volume-brain_subcortical_for_CLA, best_injection_list
Copy_of_normalized_projection_volume_brain_for_CLA
Copy_of_normalized_projection_volume-brain_subcortical_for_CLA
Copy_of_normalized_projection_volume-brain_subcortical_for_CLA
Copy_of_normalized_projection_volume_brain_for_CLA
Copy_of_normalized_projection_volume_brain_for_CLA
Copy_of_normalized_projection_volume-brain_subcortical_for_CLA
Copy_of_normalized_projection_volume_brain_for_CLA
Copy_of_normalized_projection_volume_brain_for_CLA
Copy_of_normalized_projection_volume_brain_for_CLA
Copy_of_normalized_projection_volume_brain_for_CLA
Copy_of_normalized_projection_volume_brain_for_CLA
Copy_of_normalized_projection_volume-brain_subcortical_for_CLA
Copy_of_normalized_projection_volume_brain_for_CLA
Copy_of_normalized_projection_volume_brain_for_CLA, best_injection_list

Copy_of_normalized_projection_volume-brain_subcortical_for_CLA
Copy_of_normalized_projection_volume_brain_for_CLA
Copy_of_normalized_projection_volume-brain_subcortical_for_CLA
Copy_of_normalized_projection_volume-brain_subcortical_for_CLA
Copy_of_normalized_projection_volume-brain_subcortical_for_CLA
Copy_of_normalized_projection_volume_brain_for_CLA
Copy_of_normalized_projection_volume_brain_for_CLA
Copy_of_normalized_projection_volume-brain_subcortical_for_CLA
Copy_of_normalized_projection_volume_brain_for_CLA, best_injection_list
Copy_of_normalized_projection_volume_brain_for_CLA
Copy_of_normalized_projection_volume_brain_for_CLA
Copy_of_normalized_projection_volume-brain_subcortical_for_CLA
Copy_of_normalized_projection_volume_brain_for_CLA
Copy_of_normalized_projection_volume_brain_for_CLA
Copy_of_normalized_projection_volume-brain_subcortical_for_CLA
Copy_of_normalized_projection_volume-brain_subcortical_for_CLA
Copy_of_normalized_projection_volume-brain_subcortical_for_CLA
Copy_of_normalized_projection_volume_brain_for_CLA
Copy_of_normalized_projection_volume_brain_for_CLA
Copy_of_normalized_projection_volume_brain_for_CLA
Copy_of_normalized_projection_volume-brain_subcortical_for_CLA
Copy_of_normalized_projection_volume_brain_for_CLA
Copy_of_normalized_projection_volume_brain_for_CLA, best_injection_list
Copy_of_normalized_projection_volume_brain_for_CLA
Copy_of_normalized_projection_volume_brain_for_CLA
Copy_of_normalized_projection_volume-brain_subcortical_for_CLA
Copy_of_normalized_projection_volume-brain_subcortical_for_CLA
Copy_of_normalized_projection_volume-brain_subcortical_for_CLA, best_injection_list
Copy_of_normalized_projection_volume-brain_subcortical_for_CLA
Copy_of_normalized_projection_volume-brain_subcortical_for_CLA, best_injection_list
Copy_of_normalized_projection_volume-brain_subcortical_for_CLA
Copy_of_normalized_projection_volume-brain_subcortical_for_CLA
Copy_of_normalized_projection_volume-brain_subcortical_for_CLA
Copy_of_normalized_projection_volume_brain_for_CLA
Copy_of_normalized_projection_volume-brain_subcortical_for_CLA
Copy_of_normalized_projection_volume-brain_subcortical_for_CLA
Copy_of_normalized_projection_volume-brain_subcortical_for_CLA
Copy_of_normalized_projection_volume-brain_subcortical_for_CLA
Copy_of_normalized_projection_volume-brain_subcortical_for_CLA
Copy_of_normalized_projection_volume-brain_subcortical_for_CLA
Copy_of_normalized_projection_volume-brain_subcortical_for_CLA
Copy_of_normalized_projection_volume-brain_subcortical_for_CLA
Copy_of_normalized_projection_volume-brain_subcortical_for_CLA
Copy_of_normalized_projection_volume-brain_subcortical_for_CLA
Copy_of_normalized_projection_volume-brain_subcortical_for_CLA
Copy_of_normalized_projection_volume-brain_subcortical_for_CLA
Copy_of_normalized_projection_volume-brain_subcortical_for_CLA

Copy_of_normalized_projection_volume-brain_subcortical_for_CLA
Copy_of_normalized_projection_volume-brain_subcortical_for_CLA
Copy_of_normalized_projection_volume-brain_subcortical_for_CLA
Copy_of_normalized_projection_volume-brain_subcortical_for_CLA
Copy_of_normalized_projection_volume-brain_subcortical_for_CLA
Copy_of_normalized_projection_volume-brain_subcortical_for_CLA
Copy_of_normalized_projection_volume_brain_for_CLA, best_injection_list
Copy_of_normalized_projection_volume-brain_subcortical_for_CLA
Copy_of_normalized_projection_volume-brain_subcortical_for_CLA
Copy_of_normalized_projection_volume_brain_for_CLA, best_injection_list
Copy_of_normalized_projection_volume-brain_subcortical_for_CLA
Copy_of_normalized_projection_volume-brain_subcortical_for_CLA
Copy_of_normalized_projection_volume-brain_subcortical_for_CLA
Copy_of_normalized_projection_volume-brain_subcortical_for_CLA
Copy_of_normalized_projection_volume-brain_subcortical_for_CLA
Copy_of_normalized_projection_volume-brain_subcortical_for_CLA
Copy_of_normalized_projection_volume-brain_subcortical_for_CLA, best_injection_list
Copy_of_normalized_projection_volume-brain_subcortical_for_CLA
Copy_of_normalized_projection_volume_brain_for_CLA
Copy_of_normalized_projection_volume-brain_subcortical_for_CLA
Copy_of_normalized_projection_volume-brain_subcortical_for_CLA
Copy_of_normalized_projection_volume-brain_subcortical_for_CLA
Copy_of_normalized_projection_volume-brain_subcortical_for_CLA
Copy_of_normalized_projection_volume_brain_for_CLA
Copy_of_normalized_projection_volume-brain_subcortical_for_CLA
Copy_of_normalized_projection_volume-brain_subcortical_for_CLA
Copy_of_normalized_projection_volume_brain_for_CLA
Copy_of_normalized_projection_volume_brain_for_CLA
Copy_of_normalized_projection_volume_brain_for_CLA, best_injection_list
Copy_of_normalized_projection_volume_brain_for_CLA
Copy_of_normalized_projection_volume_brain_for_CLA, best_injection_list
Copy_of_normalized_projection_volume-brain_subcortical_for_CLA
Copy_of_normalized_projection_volume-brain_subcortical_for_CLA
Copy_of_normalized_projection_volume-brain_subcortical_for_CLA
Copy_of_normalized_projection_volume-brain_subcortical_for_CLA, best_injection_list
Copy_of_normalized_projection_volume_brain_for_CLA
Copy_of_normalized_projection_volume_brain_for_CLA, best_injection_list
Copy_of_normalized_projection_volume_brain_for_CLA
Copy_of_normalized_projection_volume_brain_for_CLA, best_injection_list
Copy_of_normalized_projection_volume_brain_for_CLA
Copy_of_normalized_projection_volume_brain_for_CLA
Copy_of_normalized_projection_volume_brain_for_CLA
Copy_of_normalized_projection_volume_brain_for_CLA
Copy_of_normalized_projection_volume_brain_for_CLA
Copy_of_normalized_projection_volume_brain_for_CLA
Copy_of_normalized_projection_volume_brain_for_CLA
Copy_of_normalized_projection_volume_brain_for_CLA

Copy_of_normalized_projection_volume_brain_for_CLA
Copy_of_normalized_projection_volume_brain_for_CLA
Copy_of_normalized_projection_volume_brain_for_CLA
Copy_of_normalized_projection_volume-brain_subcortical_for_CLA
Copy_of_normalized_projection_volume_brain_for_CLA
Copy_of_normalized_projection_volume_brain_for_CLA
Copy_of_normalized_projection_volume_brain_for_CLA
Copy_of_normalized_projection_volume-brain_subcortical_for_CLA
Copy_of_normalized_projection_volume-brain_subcortical_for_CLA
Copy_of_normalized_projection_volume-brain_subcortical_for_CLA
Copy_of_normalized_projection_volume-brain_subcortical_for_CLA
Copy_of_normalized_projection_volume-brain_subcortical_for_CLA
Copy_of_normalized_projection_volume-brain_subcortical_for_CLA, best_injection_list
Copy_of_normalized_projection_volume-brain_subcortical_for_CLA
Copy_of_normalized_projection_volume-brain_subcortical_for_CLA
Copy_of_normalized_projection_volume-brain_subcortical_for_CLA
Copy_of_normalized_projection_volume-brain_subcortical_for_CLA
Copy_of_normalized_projection_volume-brain_subcortical_for_CLA, best_injection_list
Copy_of_normalized_projection_volume-brain_subcortical_for_CLA
Copy_of_normalized_projection_volume-brain_subcortical_for_CLA, best_injection_list
Copy_of_normalized_projection_volume_brain_for_CLA
Copy_of_normalized_projection_volume_brain_for_CLA
Copy_of_normalized_projection_volume-brain_subcortical_for_CLA
Copy_of_normalized_projection_volume-brain_subcortical_for_CLA
Copy_of_normalized_projection_volume_brain_for_CLA
Copy_of_normalized_projection_volume_brain_for_CLA
Copy_of_normalized_projection_volume-brain_subcortical_for_CLA
Copy_of_normalized_projection_volume_brain_for_CLA
Copy_of_normalized_projection_volume_brain_for_CLA
Copy_of_normalized_projection_volume-brain_subcortical_for_CLA
Copy_of_normalized_projection_volume_brain_for_CLA, best_injection_list
Copy_of_normalized_projection_volume_brain_for_CLA
Copy_of_normalized_projection_volume-brain_subcortical_for_CLA, best_injection_list
Copy_of_normalized_projection_volume-brain_subcortical_for_CLA
Copy_of_normalized_projection_volume-brain_subcortical_for_CLA
Copy_of_normalized_projection_volume-brain_subcortical_for_CLA
Copy_of_normalized_projection_volume_brain_for_CLA
Copy_of_normalized_projection_volume-brain_subcortical_for_CLA
Copy_of_normalized_projection_volume_brain_for_CLA
Copy_of_normalized_projection_volume_brain_for_CLA
Copy_of_normalized_projection_volume-brain_subcortical_for_CLA
Copy_of_normalized_projection_volume-brain_subcortical_for_CLA
Copy_of_normalized_projection_volume_brain_for_CLA, best_injection_list
Copy_of_normalized_projection_volume-brain_subcortical_for_CLA
Copy_of_normalized_projection_volume_brain_for_CLA
Copy_of_normalized_projection_volume-brain_subcortical_for_CLA
Copy_of_normalized_projection_volume-brain_subcortical_for_CLA

Copy_of_normalized_projection_volume-brain_subcortical_for_CLA
Copy_of_normalized_projection_volume-brain_subcortical_for_CLA
Copy_of_normalized_projection_volume_brain_for_CLA
Copy_of_normalized_projection_volume_brain_for_CLA
Copy_of_normalized_projection_volume_brain_for_CLA
Copy_of_normalized_projection_volume_brain_for_CLA
Copy_of_normalized_projection_volume_brain_for_CLA
Copy_of_normalized_projection_volume_brain_for_CLA
Copy_of_normalized_projection_volume_brain_for_CLA, best_injection_list
Copy_of_normalized_projection_volume_brain_for_CLA
Copy_of_normalized_projection_volume-brain_subcortical_for_CLA
Copy_of_normalized_projection_volume_brain_for_CLA
Copy_of_normalized_projection_volume_brain_for_CLA
Copy_of_normalized_projection_volume-brain_subcortical_for_CLA
Copy_of_normalized_projection_volume-brain_subcortical_for_CLA
Copy_of_normalized_projection_volume-brain_subcortical_for_CLA
Copy_of_normalized_projection_volume-brain_subcortical_for_CLA, best_injection_list
Copy_of_normalized_projection_volume-brain_subcortical_for_CLA, best_injection_list
Copy_of_normalized_projection_volume-brain_subcortical_for_CLA, best_injection_list
Copy_of_normalized_projection_volume-brain_subcortical_for_CLA
Copy_of_normalized_projection_volume-brain_subcortical_for_CLA
Copy_of_normalized_projection_volume_brain_for_CLA
Copy_of_normalized_projection_volume-brain_subcortical_for_CLA
Copy_of_normalized_projection_volume-brain_subcortical_for_CLA
Copy_of_normalized_projection_volume_brain_for_CLA
Copy_of_normalized_projection_volume_brain_for_CLA
Copy_of_normalized_projection_volume_brain_for_CLA
Copy_of_normalized_projection_volume_brain_for_CLA
Copy_of_normalized_projection_volume-brain_subcortical_for_CLA
Copy_of_normalized_projection_volume_brain_for_CLA
Copy_of_normalized_projection_volume-brain_subcortical_for_CLA
Copy_of_normalized_projection_volume-brain_subcortical_for_CLA
Copy_of_normalized_projection_volume-brain_subcortical_for_CLA
Copy_of_normalized_projection_volume-brain_subcortical_for_CLA
Copy_of_normalized_projection_volume-brain_subcortical_for_CLA
Copy_of_normalized_projection_volume_brain_for_CLA
Copy_of_normalized_projection_volume-brain_subcortical_for_CLA
Copy_of_normalized_projection_volume-brain_subcortical_for_CLA
Copy_of_normalized_projection_volume-brain_subcortical_for_CLA
Copy_of_normalized_projection_volume-brain_subcortical_for_CLA
Copy_of_normalized_projection_volume-brain_subcortical_for_CLA
Copy_of_normalized_projection_volume-brain_subcortical_for_CLA
Copy_of_normalized_projection_volume-brain_subcortical_for_CLA, best_injection_list
Copy_of_normalized_projection_volume-brain_subcortical_for_CLA
Copy_of_normalized_projection_volume-brain_subcortical_for_CLA
Copy_of_normalized_projection_volume-brain_subcortical_for_CLA
Copy_of_normalized_projection_volume-brain_subcortical_for_CLA

Copy_of_normalized_projection_volume-brain_subcortical_for_CLA
Copy_of_normalized_projection_volume-brain_subcortical_for_CLA
Copy_of_normalized_projection_volume-brain_subcortical_for_CLA
Copy_of_normalized_projection_volume-brain_subcortical_for_CLA
Copy_of_normalized_projection_volume-brain_subcortical_for_CLA
Copy_of_normalized_projection_volume-brain_subcortical_for_CLA
Copy_of_normalized_projection_volume-brain_subcortical_for_CLA
Copy_of_normalized_projection_volume-brain_subcortical_for_CLA
Copy_of_normalized_projection_volume-brain_subcortical_for_CLA
Copy_of_normalized_projection_volume-brain_subcortical_for_CLA
Copy_of_normalized_projection_volume_brain_for_CLA
Copy_of_normalized_projection_volume_brain_for_CLA
Copy_of_normalized_projection_volume-brain_subcortical_for_CLA
Copy_of_normalized_projection_volume-brain_subcortical_for_CLA
Copy_of_normalized_projection_volume-brain_subcortical_for_CLA
Copy_of_normalized_projection_volume-brain_subcortical_for_CLA
Copy_of_normalized_projection_volume_brain_for_CLA, best_injection_list
Copy_of_normalized_projection_volume-brain_subcortical_for_CLA
Copy_of_normalized_projection_volume-brain_subcortical_for_CLA
Copy_of_normalized_projection_volume-brain_subcortical_for_CLA
Copy_of_normalized_projection_volume-brain_subcortical_for_CLA
Copy_of_normalized_projection_volume-brain_subcortical_for_CLA
Copy_of_normalized_projection_volume-brain_subcortical_for_CLA
Copy_of_normalized_projection_volume-brain_subcortical_for_CLA
Copy_of_normalized_projection_volume-brain_subcortical_for_CLA
Copy_of_normalized_projection_volume-brain_subcortical_for_CLA
Copy_of_normalized_projection_volume-brain_subcortical_for_CLA
Copy_of_normalized_projection_volume-brain_subcortical_for_CLA
Copy_of_normalized_projection_volume-brain_subcortical_for_CLA
Copy_of_normalized_projection_volume-brain_subcortical_for_CLA
Copy_of_normalized_projection_volume-brain_subcortical_for_CLA
Copy_of_normalized_projection_volume-brain_subcortical_for_CLA
Copy_of_normalized_projection_volume-brain_subcortical_for_CLA
Copy_of_normalized_projection_volume-brain_subcortical_for_CLA
Copy_of_normalized_projection_volume-brain_subcortical_for_CLA
Copy_of_normalized_projection_volume-brain_subcortical_for_CLA, best_injection_list
Copy_of_normalized_projection_volume-brain_subcortical_for_CLA
Copy_of_normalized_projection_volume-brain_subcortical_for_CLA
Copy_of_normalized_projection_volume-brain_subcortical_for_CLA, best_injection_list
Copy_of_normalized_projection_volume-brain_subcortical_for_CLA, best_injection_list
Copy_of_normalized_projection_volume_brain_for_CLA
Copy_of_normalized_projection_volume-brain_subcortical_for_CLA
Copy_of_normalized_projection_volume-brain_subcortical_for_CLA
Copy_of_normalized_projection_volume-brain_subcortical_for_CLA
Copy_of_normalized_projection_volume-brain_subcortical_for_CLA
Copy_of_normalized_projection_volume-brain_subcortical_for_CLA
Copy_of_normalized_projection_volume-brain_subcortical_for_CLA

Copy_of_normalized_projection_volume-brain_subcortical_for_CLA
Copy_of_normalized_projection_volume-brain_subcortical_for_CLA
Copy_of_normalized_projection_volume-brain_subcortical_for_CLA
Copy_of_normalized_projection_volume-brain_subcortical_for_CLA
Copy_of_normalized_projection_volume-brain_subcortical_for_CLA
Copy_of_normalized_projection_volume_brain_for_CLA
Copy_of_normalized_projection_volume_brain_for_CLA, best_injection_list
Copy_of_normalized_projection_volume-brain_subcortical_for_CLA
Copy_of_normalized_projection_volume-brain_subcortical_for_CLA
Copy_of_normalized_projection_volume_brain_for_CLA
Copy_of_normalized_projection_volume_brain_for_CLA
Copy_of_normalized_projection_volume_brain_for_CLA
Copy_of_normalized_projection_volume_brain_for_CLA
Copy_of_normalized_projection_volume_brain_for_CLA
Copy_of_normalized_projection_volume-brain_subcortical_for_CLA
Copy_of_normalized_projection_volume-brain_subcortical_for_CLA
Copy_of_normalized_projection_volume-brain_subcortical_for_CLA
Copy_of_normalized_projection_volume-brain_subcortical_for_CLA
Copy_of_normalized_projection_volume-brain_subcortical_for_CLA
Copy_of_normalized_projection_volume-brain_subcortical_for_CLA, best_injection_list
Copy_of_normalized_projection_volume-brain_subcortical_for_CLA
Copy_of_normalized_projection_volume_brain_for_CLA
Copy_of_normalized_projection_volume_brain_for_CLA
Copy_of_normalized_projection_volume_brain_for_CLA
Copy_of_normalized_projection_volume-brain_subcortical_for_CLA
Copy_of_normalized_projection_volume-brain_subcortical_for_CLA
Copy_of_normalized_projection_volume-brain_subcortical_for_CLA
Copy_of_normalized_projection_volume-brain_subcortical_for_CLA
Copy_of_normalized_projection_volume-brain_subcortical_for_CLA
Copy_of_normalized_projection_volume_brain_for_CLA
Copy_of_normalized_projection_volume_brain_for_CLA
Copy_of_normalized_projection_volume_brain_for_CLA
Copy_of_normalized_projection_volume_brain_for_CLA, best_injection_list
Copy_of_normalized_projection_volume_brain_for_CLA
Copy_of_normalized_projection_volume_brain_for_CLA
Copy_of_normalized_projection_volume_brain_for_CLA
Copy_of_normalized_projection_volume-brain_subcortical_for_CLA
Copy_of_normalized_projection_volume_brain_for_CLA
Copy_of_normalized_projection_volume_brain_for_CLA
Copy_of_normalized_projection_volume-brain_subcortical_for_CLA
Copy_of_normalized_projection_volume-brain_subcortical_for_CLA
Copy_of_normalized_projection_volume-brain_subcortical_for_CLA
Copy_of_normalized_projection_volume_brain_for_CLA
Copy_of_normalized_projection_volume-brain_subcortical_for_CLA
Copy_of_normalized_projection_volume-brain_subcortical_for_CLA
Copy_of_normalized_projection_volume-brain_subcortical_for_CLA
Copy_of_normalized_projection_volume_brain_for_CLA

Copy_of_normalized_projection_volume-brain_subcortical_for_CLA
Copy_of_normalized_projection_volume-brain_subcortical_for_CLA
Copy_of_normalized_projection_volume_brain_for_CLA
Copy_of_normalized_projection_volume-brain_subcortical_for_CLA
Copy_of_normalized_projection_volume-brain_subcortical_for_CLA, best_injection_list
Copy_of_normalized_projection_volume-brain_subcortical_for_CLA
Copy_of_normalized_projection_volume_brain_for_CLA
Copy_of_normalized_projection_volume_brain_for_CLA
Copy_of_normalized_projection_volume-brain_subcortical_for_CLA
Copy_of_normalized_projection_volume-brain_subcortical_for_CLA
Copy_of_normalized_projection_volume-brain_subcortical_for_CLA
Copy_of_normalized_projection_volume-brain_subcortical_for_CLA
Copy_of_normalized_projection_volume-brain_subcortical_for_CLA
Copy_of_normalized_projection_volume-brain_subcortical_for_CLA
Copy_of_normalized_projection_volume_brain_for_CLA
Copy_of_normalized_projection_volume_brain_for_CLA
Copy_of_normalized_projection_volume-brain_subcortical_for_CLA
Copy_of_normalized_projection_volume-brain_subcortical_for_CLA
Copy_of_normalized_projection_volume-brain_subcortical_for_CLA
Copy_of_normalized_projection_volume-brain_subcortical_for_CLA
Copy_of_normalized_projection_volume-brain_subcortical_for_CLA
Copy_of_normalized_projection_volume-brain_subcortical_for_CLA
Copy_of_normalized_projection_volume-brain_subcortical_for_CLA
Copy_of_normalized_projection_volume-brain_subcortical_for_CLA
Copy_of_normalized_projection_volume-brain_subcortical_for_CLA
Copy_of_normalized_projection_volume-brain_subcortical_for_CLA
Copy_of_normalized_projection_volume-brain_subcortical_for_CLA
Copy_of_normalized_projection_volume-brain_subcortical_for_CLA
Copy_of_normalized_projection_volume-brain_subcortical_for_CLA
Copy_of_normalized_projection_volume-brain_subcortical_for_CLA, best_injection_list
Copy_of_normalized_projection_volume-brain_subcortical_for_CLA
Copy_of_normalized_projection_volume-brain_subcortical_for_CLA
Copy_of_normalized_projection_volume-brain_subcortical_for_CLA
Copy_of_normalized_projection_volume-brain_subcortical_for_CLA
Copy_of_normalized_projection_volume_brain_for_CLA
Copy_of_normalized_projection_volume_brain_for_CLA
Copy_of_normalized_projection_volume_brain_for_CLA
Copy_of_normalized_projection_volume_brain_for_CLA
Copy_of_normalized_projection_volume_brain_for_CLA
Copy_of_normalized_projection_volume_brain_for_CLA, best_injection_list
Copy_of_normalized_projection_volume_brain_for_CLA
Copy_of_normalized_projection_volume_brain_for_CLA
Copy_of_normalized_projection_volume_brain_for_CLA
Copy_of_normalized_projection_volume-brain_subcortical_for_CLA
Copy_of_normalized_projection_volume-brain_subcortical_for_CLA
Copy_of_normalized_projection_volume_brain_for_CLA
Copy_of_normalized_projection_volume_brain_for_CLA

Copy_of_normalized_projection_volume_brain_for_CLA, best_injection_list
Copy_of_normalized_projection_volume-brain_subcortical_for_CLA
Copy_of_normalized_projection_volume_brain_for_CLA
Copy_of_normalized_projection_volume_brain_for_CLA, best_injection_list
Copy_of_normalized_projection_volume-brain_subcortical_for_CLA
Copy_of_normalized_projection_volume-brain_subcortical_for_CLA
Copy_of_normalized_projection_volume-brain_subcortical_for_CLA
Copy_of_normalized_projection_volume-brain_subcortical_for_CLA
Copy_of_normalized_projection_volume-brain_subcortical_for_CLA
Copy_of_normalized_projection_volume-brain_subcortical_for_CLA
Copy_of_normalized_projection_volume-brain_subcortical_for_CLA
Copy_of_normalized_projection_volume-brain_subcortical_for_CLA
Copy_of_normalized_projection_volume_brain_for_CLA
Copy_of_normalized_projection_volume-brain_subcortical_for_CLA
Copy_of_normalized_projection_volume-brain_subcortical_for_CLA
Copy_of_normalized_projection_volume_brain_for_CLA
Copy_of_normalized_projection_volume_brain_for_CLA
Copy_of_normalized_projection_volume_brain_for_CLA, best_injection_list
Copy_of_normalized_projection_volume-brain_subcortical_for_CLA
Copy_of_normalized_projection_volume-brain_subcortical_for_CLA
Copy_of_normalized_projection_volume_brain_for_CLA
Copy_of_normalized_projection_volume_brain_for_CLA
Copy_of_normalized_projection_volume-brain_subcortical_for_CLA
Copy_of_normalized_projection_volume-brain_subcortical_for_CLA
Copy_of_normalized_projection_volume-brain_subcortical_for_CLA, best_injection_list
Copy_of_normalized_projection_volume-brain_subcortical_for_CLA, best_injection_list
Copy_of_normalized_projection_volume-brain_subcortical_for_CLA
Copy_of_normalized_projection_volume-brain_subcortical_for_CLA
Copy_of_normalized_projection_volume_brain_for_CLA
Copy_of_normalized_projection_volume-brain_subcortical_for_CLA, best_injection_list
Copy_of_normalized_projection_volume-brain_subcortical_for_CLA
Copy_of_normalized_projection_volume-brain_subcortical_for_CLA
Copy_of_normalized_projection_volume-brain_subcortical_for_CLA
Copy_of_normalized_projection_volume_brain_for_CLA
Copy_of_normalized_projection_volume-brain_subcortical_for_CLA
Copy_of_normalized_projection_volume-brain_subcortical_for_CLA
Copy_of_normalized_projection_volume-brain_subcortical_for_CLA
Copy_of_normalized_projection_volume-brain_subcortical_for_CLA
Copy_of_normalized_projection_volume-brain_subcortical_for_CLA
Copy_of_normalized_projection_volume_brain_for_CLA
Copy_of_normalized_projection_volume_brain_for_CLA
Copy_of_normalized_projection_volume-brain_subcortical_for_CLA
Copy_of_normalized_projection_volume-brain_subcortical_for_CLA
Copy_of_normalized_projection_volume-brain_subcortical_for_CLA
Copy_of_normalized_projection_volume-brain_subcortical_for_CLA
Copy_of_normalized_projection_volume_brain_for_CLA
Copy_of_normalized_projection_volume-brain_subcortical_for_CLA

Copy_of_normalized_projection_volume-brain_subcortical_for_CLA
Copy_of_normalized_projection_volume-brain_subcortical_for_CLA
Copy_of_normalized_projection_volume_brain_for_CLA
Copy_of_normalized_projection_volume_brain_for_CLA
Copy_of_normalized_projection_volume-brain_subcortical_for_CLA, best_injection_list
Copy_of_normalized_projection_volume_brain_for_CLA
Copy_of_normalized_projection_volume_brain_for_CLA, best_injection_list
Copy_of_normalized_projection_volume-brain_subcortical_for_CLA
Copy_of_normalized_projection_volume-brain_subcortical_for_CLA, best_injection_list
Copy_of_normalized_projection_volume-brain_subcortical_for_CLA
Copy_of_normalized_projection_volume-brain_subcortical_for_CLA
Copy_of_normalized_projection_volume-brain_subcortical_for_CLA
Copy_of_normalized_projection_volume_brain_for_CLA
Copy_of_normalized_projection_volume-brain_subcortical_for_CLA
Copy_of_normalized_projection_volume-brain_subcortical_for_CLA
Copy_of_normalized_projection_volume-brain_subcortical_for_CLA
Copy_of_normalized_projection_volume-brain_subcortical_for_CLA
Copy_of_normalized_projection_volume_brain_for_CLA
Copy_of_normalized_projection_volume-brain_subcortical_for_CLA
Copy_of_normalized_projection_volume_brain_for_CLA
Copy_of_normalized_projection_volume_brain_for_CLA
Copy_of_normalized_projection_volume-brain_subcortical_for_CLA
Copy_of_normalized_projection_volume-brain_subcortical_for_CLA
Copy_of_normalized_projection_volume-brain_subcortical_for_CLA
Copy_of_normalized_projection_volume-brain_subcortical_for_CLA
Copy_of_normalized_projection_volume-brain_subcortical_for_CLA
Copy_of_normalized_projection_volume_brain_for_CLA
Copy_of_normalized_projection_volume_brain_for_CLA
Copy_of_normalized_projection_volume_brain_for_CLA
Copy_of_normalized_projection_volume_brain_for_CLA
Copy_of_normalized_projection_volume_brain_for_CLA
Copy_of_normalized_projection_volume_brain_for_CLA
Copy_of_normalized_projection_volume-brain_subcortical_for_CLA
Copy_of_normalized_projection_volume-brain_subcortical_for_CLA
Copy_of_normalized_projection_volume-brain_subcortical_for_CLA, best_injection_list
Copy_of_normalized_projection_volume-brain_subcortical_for_CLA, best_injection_list
Copy_of_normalized_projection_volume-brain_subcortical_for_CLA
Copy_of_normalized_projection_volume-brain_subcortical_for_CLA, best_injection_list
Copy_of_normalized_projection_volume-brain_subcortical_for_CLA
Copy_of_normalized_projection_volume-brain_subcortical_for_CLA
Copy_of_normalized_projection_volume-brain_subcortical_for_CLA
Copy_of_normalized_projection_volume_brain_for_CLA, best_injection_list
Copy_of_normalized_projection_volume_brain_for_CLA
Copy_of_normalized_projection_volume-brain_subcortical_for_CLA
Copy_of_normalized_projection_volume-brain_subcortical_for_CLA
Copy_of_normalized_projection_volume-brain_subcortical_for_CLA
Copy_of_normalized_projection_volume-brain_subcortical_for_CLA

Copy_of_normalized_projection_volume_brain_for_CLA
Copy_of_normalized_projection_volume_brain_for_CLA, best_injection_list
Copy_of_normalized_projection_volume_brain_for_CLA, best_injection_list
Copy_of_normalized_projection_volume_brain_for_CLA
Copy_of_normalized_projection_volume_brain_for_CLA, best_injection_list
Copy_of_normalized_projection_volume-brain_subcortical_for_CLA
Copy_of_normalized_projection_volume_brain_for_CLA
Copy_of_normalized_projection_volume_brain_for_CLA
Copy_of_normalized_projection_volume-brain_subcortical_for_CLA
Copy_of_normalized_projection_volume_brain_for_CLA
Copy_of_normalized_projection_volume_brain_for_CLA
Copy_of_normalized_projection_volume_brain_for_CLA
Copy_of_normalized_projection_volume-brain_subcortical_for_CLA
Copy_of_normalized_projection_volume-brain_subcortical_for_CLA, best_injection_list
Copy_of_normalized_projection_volume-brain_subcortical_for_CLA
Copy_of_normalized_projection_volume-brain_subcortical_for_CLA
Copy_of_normalized_projection_volume_brain_for_CLA, best_injection_list
Copy_of_normalized_projection_volume_brain_for_CLA, best_injection_list
Copy_of_normalized_projection_volume_brain_for_CLA
Copy_of_normalized_projection_volume_brain_for_CLA, best_injection_list
Copy_of_normalized_projection_volume-brain_subcortical_for_CLA
Copy_of_normalized_projection_volume_brain_for_CLA, best_injection_list
Copy_of_normalized_projection_volume-brain_subcortical_for_CLA, best_injection_list
Copy_of_normalized_projection_volume-brain_subcortical_for_CLA, best_injection_list
Copy_of_normalized_projection_volume-brain_subcortical_for_CLA
Copy_of_normalized_projection_volume-brain_subcortical_for_CLA
Copy_of_normalized_projection_volume-brain_subcortical_for_CLA
Copy_of_normalized_projection_volume-brain_subcortical_for_CLA
Copy_of_normalized_projection_volume-brain_subcortical_for_CLA
Copy_of_normalized_projection_volume-brain_subcortical_for_CLA, best_injection_list
Copy_of_normalized_projection_volume-brain_subcortical_for_CLA
Copy_of_normalized_projection_volume-brain_subcortical_for_CLA
Copy_of_normalized_projection_volume-brain_subcortical_for_CLA
Copy_of_normalized_projection_volume-brain_subcortical_for_CLA
Copy_of_normalized_projection_volume-brain_subcortical_for_CLA
Copy_of_normalized_projection_volume-brain_subcortical_for_CLA
Copy_of_normalized_projection_volume-brain_subcortical_for_CLA
Copy_of_normalized_projection_volume_brain_for_CLA
Copy_of_normalized_projection_volume_brain_for_CLA
Copy_of_normalized_projection_volume_brain_for_CLA
Copy_of_normalized_projection_volume-brain_subcortical_for_CLA
Copy_of_normalized_projection_volume_brain_for_CLA
Copy_of_normalized_projection_volume_brain_for_CLA
Copy_of_normalized_projection_volume_brain_for_CLA
Copy_of_normalized_projection_volume-brain_subcortical_for_CLA
Copy_of_normalized_projection_volume_brain_for_CLA, best_injection_list
Copy_of_normalized_projection_volume-brain_subcortical_for_CLA

Copy_of_normalized_projection_volume-brain_subcortical_for_CLA
Copy_of_normalized_projection_volume_brain_for_CLA
Copy_of_normalized_projection_volume_brain_for_CLA
Copy_of_normalized_projection_volume-brain_subcortical_for_CLA
Copy_of_normalized_projection_volume-brain_subcortical_for_CLA
Copy_of_normalized_projection_volume_brain_for_CLA, best_injection_list
Copy_of_normalized_projection_volume_brain_for_CLA, best_injection_list
Copy_of_normalized_projection_volume-brain_subcortical_for_CLA, best_injection_list
Copy_of_normalized_projection_volume-brain_subcortical_for_CLA
Copy_of_normalized_projection_volume_brain_for_CLA
Copy_of_normalized_projection_volume-brain_subcortical_for_CLA
Copy_of_normalized_projection_volume_brain_for_CLA
Copy_of_normalized_projection_volume_brain_for_CLA
Copy_of_normalized_projection_volume-brain_subcortical_for_CLA
Copy_of_normalized_projection_volume-brain_subcortical_for_CLA
Copy_of_normalized_projection_volume_brain_for_CLA
Copy_of_normalized_projection_volume_brain_for_CLA, best_injection_list
Copy_of_normalized_projection_volume-brain_subcortical_for_CLA
Copy_of_normalized_projection_volume-brain_subcortical_for_CLA
Copy_of_normalized_projection_volume-brain_subcortical_for_CLA, best_injection_list
Copy_of_normalized_projection_volume-brain_subcortical_for_CLA
Copy_of_normalized_projection_volume-brain_subcortical_for_CLA
Copy_of_normalized_projection_volume_brain_for_CLA, best_injection_list
Copy_of_normalized_projection_volume-brain_subcortical_for_CLA
Copy_of_normalized_projection_volume-brain_subcortical_for_CLA
Copy_of_normalized_projection_volume-brain_subcortical_for_CLA
Copy_of_normalized_projection_volume-brain_subcortical_for_CLA
Copy_of_normalized_projection_volume-brain_subcortical_for_CLA
Copy_of_normalized_projection_volume-brain_subcortical_for_CLA
Copy_of_normalized_projection_volume-brain_subcortical_for_CLA
Copy_of_normalized_projection_volume_brain_for_CLA
Copy_of_normalized_projection_volume-brain_subcortical_for_CLA
Copy_of_normalized_projection_volume-brain_subcortical_for_CLA
Copy_of_normalized_projection_volume_brain_for_CLA
Copy_of_normalized_projection_volume-brain_subcortical_for_CLA
Copy_of_normalized_projection_volume-brain_subcortical_for_CLA
Copy_of_normalized_projection_volume-brain_subcortical_for_CLA
Copy_of_normalized_projection_volume-brain_subcortical_for_CLA, best_injection_list
Copy_of_normalized_projection_volume_brain_for_CLA
Copy_of_normalized_projection_volume_brain_for_CLA
Copy_of_normalized_projection_volume-brain_subcortical_for_CLA
Copy_of_normalized_projection_volume-brain_subcortical_for_CLA, best_injection_list
Copy_of_normalized_projection_volume-brain_subcortical_for_CLA
Copy_of_normalized_projection_volume-brain_subcortical_for_CLA
Copy_of_normalized_projection_volume_brain_for_CLA, best_injection_list
Copy_of_normalized_projection_volume-brain_subcortical_for_CLA
Copy_of_normalized_projection_volume-brain_subcortical_for_CLA

Copy_of_normalized_projection_volume-brain_subcortical_for_CLA
Copy_of_normalized_projection_volume-brain_subcortical_for_CLA
Copy_of_normalized_projection_volume-brain_subcortical_for_CLA
Copy_of_normalized_projection_volume-brain_subcortical_for_CLA
Copy_of_normalized_projection_volume-brain_subcortical_for_CLA
Copy_of_normalized_projection_volume-brain_subcortical_for_CLA
Copy_of_normalized_projection_volume-brain_subcortical_for_CLA
Copy_of_normalized_projection_volume-brain_subcortical_for_CLA
Copy_of_normalized_projection_volume-brain_subcortical_for_CLA, best_injection_list
Copy_of_normalized_projection_volume-brain_subcortical_for_CLA
Copy_of_normalized_projection_volume_brain_for_CLA
Copy_of_normalized_projection_volume_brain_for_CLA
Copy_of_normalized_projection_volume_brain_for_CLA, best_injection_list
Copy_of_normalized_projection_volume_brain_for_CLA
Copy_of_normalized_projection_volume-brain_subcortical_for_CLA
Copy_of_normalized_projection_volume-brain_subcortical_for_CLA
Copy_of_normalized_projection_volume-brain_subcortical_for_CLA
Copy_of_normalized_projection_volume-brain_subcortical_for_CLA
Copy_of_normalized_projection_volume_brain_for_CLA, best_injection_list
Copy_of_normalized_projection_volume_brain_for_CLA
Copy_of_normalized_projection_volume-brain_subcortical_for_CLA
Copy_of_normalized_projection_volume_brain_for_CLA
Copy_of_normalized_projection_volume_brain_for_CLA
Copy_of_normalized_projection_volume-brain_subcortical_for_CLA
Copy_of_normalized_projection_volume_brain_for_CLA
Copy_of_normalized_projection_volume-brain_subcortical_for_CLA
Copy_of_normalized_projection_volume-brain_subcortical_for_CLA
Copy_of_normalized_projection_volume-brain_subcortical_for_CLA, best_injection_list
Copy_of_normalized_projection_volume-brain_subcortical_for_CLA, best_injection_list
Copy_of_normalized_projection_volume-brain_subcortical_for_CLA
Copy_of_normalized_projection_volume-brain_subcortical_for_CLA
Copy_of_normalized_projection_volume-brain_subcortical_for_CLA
Copy_of_normalized_projection_volume-brain_subcortical_for_CLA, best_injection_list
Copy_of_normalized_projection_volume-brain_subcortical_for_CLA
Copy_of_normalized_projection_volume_brain_for_CLA
Copy_of_normalized_projection_volume_brain_for_CLA
Copy_of_normalized_projection_volume-brain_subcortical_for_CLA
Copy_of_normalized_projection_volume-brain_subcortical_for_CLA, best_injection_list
Copy_of_normalized_projection_volume-brain_subcortical_for_CLA
Copy_of_normalized_projection_volume_brain_for_CLA
Copy_of_normalized_projection_volume_brain_for_CLA, best_injection_list
Copy_of_normalized_projection_volume_brain_for_CLA
Copy_of_normalized_projection_volume-brain_subcortical_for_CLA
Copy_of_normalized_projection_volume-brain_subcortical_for_CLA
Copy_of_normalized_projection_volume_brain_for_CLA
Copy_of_normalized_projection_volume_brain_for_CLA
Copy_of_normalized_projection_volume-brain_subcortical_for_CLA

Copy_of_normalized_projection_volume_brain_for_CLA
Copy_of_normalized_projection_volume_brain_for_CLA
Copy_of_normalized_projection_volume-brain_subcortical_for_CLA
Copy_of_normalized_projection_volume-brain_subcortical_for_CLA
Copy_of_normalized_projection_volume-brain_subcortical_for_CLA
Copy_of_normalized_projection_volume-brain_subcortical_for_CLA
Copy_of_normalized_projection_volume-brain_subcortical_for_CLA
Copy_of_normalized_projection_volume-brain_subcortical_for_CLA
Copy_of_normalized_projection_volume_brain_for_CLA, best_injection_list
Copy_of_normalized_projection_volume_brain_for_CLA, best_injection_list
Copy_of_normalized_projection_volume-brain_subcortical_for_CLA
Copy_of_normalized_projection_volume_brain_for_CLA
Copy_of_normalized_projection_volume_brain_for_CLA
Copy_of_normalized_projection_volume-brain_subcortical_for_CLA
Copy_of_normalized_projection_volume-brain_subcortical_for_CLA
Copy_of_normalized_projection_volume_brain_for_CLA
Copy_of_normalized_projection_volume_brain_for_CLA
Copy_of_normalized_projection_volume-brain_subcortical_for_CLA
Copy_of_normalized_projection_volume-brain_subcortical_for_CLA, best_injection_list
Copy_of_normalized_projection_volume-brain_subcortical_for_CLA
Copy_of_normalized_projection_volume-brain_subcortical_for_CLA
Copy_of_normalized_projection_volume_brain_for_CLA
Copy_of_normalized_projection_volume-brain_subcortical_for_CLA
Copy_of_normalized_projection_volume_brain_for_CLA
Copy_of_normalized_projection_volume_brain_for_CLA, best_injection_list
Copy_of_normalized_projection_volume_brain_for_CLA
Copy_of_normalized_projection_volume-brain_subcortical_for_CLA
Copy_of_normalized_projection_volume-brain_subcortical_for_CLA
Copy_of_normalized_projection_volume-brain_subcortical_for_CLA
Copy_of_normalized_projection_volume_brain_for_CLA
Copy_of_normalized_projection_volume_brain_for_CLA
Copy_of_normalized_projection_volume-brain_subcortical_for_CLA, best_injection_list
Copy_of_normalized_projection_volume-brain_subcortical_for_CLA
Copy_of_normalized_projection_volume-brain_subcortical_for_CLA
Copy_of_normalized_projection_volume-brain_subcortical_for_CLA, best_injection_list
Copy_of_normalized_projection_volume-brain_subcortical_for_CLA
Copy_of_normalized_projection_volume-brain_subcortical_for_CLA
Copy_of_normalized_projection_volume-brain_subcortical_for_CLA
Copy_of_normalized_projection_volume-brain_subcortical_for_CLA
Copy_of_normalized_projection_volume-brain_subcortical_for_CLA
Copy_of_normalized_projection_volume-brain_subcortical_for_CLA
Copy_of_normalized_projection_volume-brain_subcortical_for_CLA
Copy_of_normalized_projection_volume-brain_subcortical_for_CLA
Copy_of_normalized_projection_volume_brain_for_CLA
Copy_of_normalized_projection_volume-brain_subcortical_for_CLA
Copy_of_normalized_projection_volume-brain_subcortical_for_CLA
Copy_of_normalized_projection_volume-brain_subcortical_for_CLA

Copy_of_normalized_projection_volume-brain_subcortical_for_CLA
Copy_of_normalized_projection_volume-brain_subcortical_for_CLA
Copy_of_normalized_projection_volume-brain_subcortical_for_CLA
Copy_of_normalized_projection_volume-brain_subcortical_for_CLA
Copy_of_normalized_projection_volume-brain_subcortical_for_CLA, best_injection_list
Copy_of_normalized_projection_volume-brain_subcortical_for_CLA
Copy_of_normalized_projection_volume-brain_subcortical_for_CLA, best_injection_list
Copy_of_normalized_projection_volume-brain_subcortical_for_CLA
Copy_of_normalized_projection_volume-brain_subcortical_for_CLA
Copy_of_normalized_projection_volume_brain_for_CLA, best_injection_list
Copy_of_normalized_projection_volume_brain_for_CLA, best_injection_list
Copy_of_normalized_projection_volume_brain_for_CLA
Copy_of_normalized_projection_volume_brain_for_CLA
Copy_of_normalized_projection_volume_brain_for_CLA
Copy_of_normalized_projection_volume_brain_for_CLA
Copy_of_normalized_projection_volume-brain_subcortical_for_CLA
Copy_of_normalized_projection_volume_brain_for_CLA
Copy_of_normalized_projection_volume_brain_for_CLA
Copy_of_normalized_projection_volume_brain_for_CLA
Copy_of_normalized_projection_volume_brain_for_CLA
Copy_of_normalized_projection_volume_brain_for_CLA
Copy_of_normalized_projection_volume-brain_subcortical_for_CLA
Copy_of_normalized_projection_volume_brain_for_CLA
Copy_of_normalized_projection_volume-brain_subcortical_for_CLA
Copy_of_normalized_projection_volume_brain_for_CLA
Copy_of_normalized_projection_volume-brain_subcortical_for_CLA
Copy_of_normalized_projection_volume-brain_subcortical_for_CLA
Copy_of_normalized_projection_volume-brain_subcortical_for_CLA
Copy_of_normalized_projection_volume-brain_subcortical_for_CLA
Copy_of_normalized_projection_volume_brain_for_CLA
Copy_of_normalized_projection_volume-brain_subcortical_for_CLA
Copy_of_normalized_projection_volume-brain_subcortical_for_CLA
Copy_of_normalized_projection_volume-brain_subcortical_for_CLA
Copy_of_normalized_projection_volume-brain_subcortical_for_CLA
Copy_of_normalized_projection_volume_brain_for_CLA
Copy_of_normalized_projection_volume_brain_for_CLA
Copy_of_normalized_projection_volume_brain_for_CLA
Copy_of_normalized_projection_volume-brain_subcortical_for_CLA
Copy_of_normalized_projection_volume-brain_subcortical_for_CLA
Copy_of_normalized_projection_volume-brain_subcortical_for_CLA
Copy_of_normalized_projection_volume-brain_subcortical_for_CLA
Copy_of_normalized_projection_volume-brain_subcortical_for_CLA, best_injection_list
Copy_of_normalized_projection_volume-brain_subcortical_for_CLA
Copy_of_normalized_projection_volume_brain_for_CLA
Copy_of_normalized_projection_volume_brain_for_CLA
Copy_of_normalized_projection_volume_brain_for_CLA
Copy_of_normalized_projection_volume-brain_subcortical_for_CLA

Copy_of_normalized_projection_volume_brain_for_CLA
Copy_of_normalized_projection_volume-brain_subcortical_for_CLA
Copy_of_normalized_projection_volume-brain_subcortical_for_CLA, best_injection_list
Copy_of_normalized_projection_volume-brain_subcortical_for_CLA
Copy_of_normalized_projection_volume-brain_subcortical_for_CLA
Copy_of_normalized_projection_volume-brain_subcortical_for_CLA
Copy_of_normalized_projection_volume-brain_subcortical_for_CLA
Copy_of_normalized_projection_volume-brain_subcortical_for_CLA
Copy_of_normalized_projection_volume-brain_subcortical_for_CLA
Copy_of_normalized_projection_volume-brain_subcortical_for_CLA
Copy_of_normalized_projection_volume-brain_subcortical_for_CLA
Copy_of_normalized_projection_volume-brain_subcortical_for_CLA, best_injection_list
Copy_of_normalized_projection_volume-brain_subcortical_for_CLA
Copy_of_normalized_projection_volume-brain_subcortical_for_CLA
Copy_of_normalized_projection_volume-brain_subcortical_for_CLA
Copy_of_normalized_projection_volume-brain_subcortical_for_CLA
Copy_of_normalized_projection_volume-brain_subcortical_for_CLA
Copy_of_normalized_projection_volume-brain_subcortical_for_CLA
Copy_of_normalized_projection_volume_brain_for_CLA
Copy_of_normalized_projection_volume-brain_subcortical_for_CLA
Copy_of_normalized_projection_volume-brain_subcortical_for_CLA
Copy_of_normalized_projection_volume_brain_for_CLA
Copy_of_normalized_projection_volume_brain_for_CLA, best_injection_list
Copy_of_normalized_projection_volume_brain_for_CLA, best_injection_list
Copy_of_normalized_projection_volume-brain_subcortical_for_CLA
Copy_of_normalized_projection_volume_brain_for_CLA, best_injection_list
Copy_of_normalized_projection_volume-brain_subcortical_for_CLA
Copy_of_normalized_projection_volume_brain_for_CLA
Copy_of_normalized_projection_volume_brain_for_CLA, best_injection_list
Copy_of_normalized_projection_volume-brain_subcortical_for_CLA
Copy_of_normalized_projection_volume-brain_subcortical_for_CLA
Copy_of_normalized_projection_volume_brain_for_CLA
Copy_of_normalized_projection_volume-brain_subcortical_for_CLA
Copy_of_normalized_projection_volume-brain_subcortical_for_CLA
Copy_of_normalized_projection_volume-brain_subcortical_for_CLA
Copy_of_normalized_projection_volume-brain_subcortical_for_CLA
Copy_of_normalized_projection_volume-brain_subcortical_for_CLA, best_injection_list
Copy_of_normalized_projection_volume-brain_subcortical_for_CLA
Copy_of_normalized_projection_volume-brain_subcortical_for_CLA, best_injection_list
Copy_of_normalized_projection_volume-brain_subcortical_for_CLA
Copy_of_normalized_projection_volume-brain_subcortical_for_CLA
Copy_of_normalized_projection_volume_brain_for_CLA, best_injection_list
Copy_of_normalized_projection_volume-brain_subcortical_for_CLA
Copy_of_normalized_projection_volume-brain_subcortical_for_CLA
Copy_of_normalized_projection_volume-brain_subcortical_for_CLA
Copy_of_normalized_projection_volume-brain_subcortical_for_CLA
Copy_of_normalized_projection_volume-brain_subcortical_for_CLA

Copy_of_normalized_projection_volume-brain_subcortical_for_CLA
Copy_of_normalized_projection_volume-brain_subcortical_for_CLA
Copy_of_normalized_projection_volume-brain_subcortical_for_CLA
Copy_of_normalized_projection_volume-brain_subcortical_for_CLA
Copy_of_normalized_projection_volume-brain_subcortical_for_CLA
Copy_of_normalized_projection_volume-brain_subcortical_for_CLA
Copy_of_normalized_projection_volume-brain_subcortical_for_CLA
Copy_of_normalized_projection_volume-brain_subcortical_for_CLA
Copy_of_normalized_projection_volume-brain_subcortical_for_CLA
Copy_of_normalized_projection_volume-brain_subcortical_for_CLA
Copy_of_normalized_projection_volume_brain_for_CLA
Copy_of_normalized_projection_volume_brain_for_CLA
Copy_of_normalized_projection_volume-brain_subcortical_for_CLA
Copy_of_normalized_projection_volume_brain_for_CLA
Copy_of_normalized_projection_volume_brain_for_CLA
Copy_of_normalized_projection_volume_brain_for_CLA
Copy_of_normalized_projection_volume-brain_subcortical_for_CLA
Copy_of_normalized_projection_volume-brain_subcortical_for_CLA
Copy_of_normalized_projection_volume-brain_subcortical_for_CLA
Copy_of_normalized_projection_volume_brain_for_CLA
Copy_of_normalized_projection_volume-brain_subcortical_for_CLA
Copy_of_normalized_projection_volume-brain_subcortical_for_CLA
Copy_of_normalized_projection_volume_brain_for_CLA
Copy_of_normalized_projection_volume_brain_for_CLA
Copy_of_normalized_projection_volume_brain_for_CLA
Copy_of_normalized_projection_volume_brain_for_CLA
Copy_of_normalized_projection_volume-brain_subcortical_for_CLA
Copy_of_normalized_projection_volume-brain_subcortical_for_CLA
Copy_of_normalized_projection_volume-brain_subcortical_for_CLA
Copy_of_normalized_projection_volume-brain_subcortical_for_CLA
Copy_of_normalized_projection_volume_brain_for_CLA
Copy_of_normalized_projection_volume_brain_for_CLA
Copy_of_normalized_projection_volume-brain_subcortical_for_CLA
Copy_of_normalized_projection_volume_brain_for_CLA
Copy_of_normalized_projection_volume_brain_for_CLA
Copy_of_normalized_projection_volume_brain_for_CLA
Copy_of_normalized_projection_volume_brain_for_CLA
Copy_of_normalized_projection_volume-brain_subcortical_for_CLA
Copy_of_normalized_projection_volume-brain_subcortical_for_CLA
Copy_of_normalized_projection_volume_brain_for_CLA
Copy_of_normalized_projection_volume-brain_subcortical_for_CLA
Copy_of_normalized_projection_volume_brain_for_CLA
Copy_of_normalized_projection_volume_brain_for_CLA
Copy_of_normalized_projection_volume_brain_for_CLA
Copy_of_normalized_projection_volume_brain_for_CLA
Copy_of_normalized_projection_volume-brain_subcortical_for_CLA, best_injection_list
Copy_of_normalized_projection_volume-brain_subcortical_for_CLA

Copy_of_normalized_projection_volume-brain_subcortical_for_CLA
Copy_of_normalized_projection_volume-brain_subcortical_for_CLA, best_injection_list
Copy_of_normalized_projection_volume_brain_for_CLA
Copy_of_normalized_projection_volume-brain_subcortical_for_CLA
Copy_of_normalized_projection_volume-brain_subcortical_for_CLA
Copy_of_normalized_projection_volume-brain_subcortical_for_CLA
Copy_of_normalized_projection_volume-brain_subcortical_for_CLA
Copy_of_normalized_projection_volume-brain_subcortical_for_CLA
Copy_of_normalized_projection_volume-brain_subcortical_for_CLA
Copy_of_normalized_projection_volume-brain_subcortical_for_CLA
Copy_of_normalized_projection_volume-brain_subcortical_for_CLA
Copy_of_normalized_projection_volume_brain_for_CLA, best_injection_list
Copy_of_normalized_projection_volume-brain_subcortical_for_CLA
Copy_of_normalized_projection_volume-brain_subcortical_for_CLA
Copy_of_normalized_projection_volume-brain_subcortical_for_CLA
Copy_of_normalized_projection_volume-brain_subcortical_for_CLA
Copy_of_normalized_projection_volume-brain_subcortical_for_CLA
Copy_of_normalized_projection_volume_brain_for_CLA
Copy_of_normalized_projection_volume-brain_subcortical_for_CLA
Copy_of_normalized_projection_volume-brain_subcortical_for_CLA
Copy_of_normalized_projection_volume-brain_subcortical_for_CLA
Copy_of_normalized_projection_volume-brain_subcortical_for_CLA
Copy_of_normalized_projection_volume-brain_subcortical_for_CLA, best_injection_list
Copy_of_normalized_projection_volume-brain_subcortical_for_CLA
Copy_of_normalized_projection_volume-brain_subcortical_for_CLA
Copy_of_normalized_projection_volume_brain_for_CLA
Copy_of_normalized_projection_volume_brain_for_CLA
Copy_of_normalized_projection_volume_brain_for_CLA, best_injection_list
Copy_of_normalized_projection_volume_brain_for_CLA
Copy_of_normalized_projection_volume-brain_subcortical_for_CLA
Copy_of_normalized_projection_volume_brain_for_CLA
Copy_of_normalized_projection_volume_brain_for_CLA
Copy_of_normalized_projection_volume_brain_for_CLA
Copy_of_normalized_projection_volume-brain_subcortical_for_CLA
Copy_of_normalized_projection_volume_brain_for_CLA, best_injection_list
Copy_of_normalized_projection_volume_brain_for_CLA
Copy_of_normalized_projection_volume-brain_subcortical_for_CLA
Copy_of_normalized_projection_volume-brain_subcortical_for_CLA
Copy_of_normalized_projection_volume-brain_subcortical_for_CLA
Copy_of_normalized_projection_volume-brain_subcortical_for_CLA, best_injection_list
Copy_of_normalized_projection_volume_brain_for_CLA, best_injection_list
Copy_of_normalized_projection_volume_brain_for_CLA
Copy_of_normalized_projection_volume-brain_subcortical_for_CLA, best_injection_list
Copy_of_normalized_projection_volume-brain_subcortical_for_CLA, best_injection_list
Copy_of_normalized_projection_volume-brain_subcortical_for_CLA
Copy_of_normalized_projection_volume-brain_subcortical_for_CLA
Copy_of_normalized_projection_volume-brain_subcortical_for_CLA

Copy_of_normalized_projection_volume-brain_subcortical_for_CLA
Copy_of_normalized_projection_volume_brain_for_CLA
Copy_of_normalized_projection_volume-brain_subcortical_for_CLA
Copy_of_normalized_projection_volume-brain_subcortical_for_CLA
Copy_of_normalized_projection_volume-brain_subcortical_for_CLA
Copy_of_normalized_projection_volume-brain_subcortical_for_CLA
Copy_of_normalized_projection_volume-brain_subcortical_for_CLA, best_injection_list
Copy_of_normalized_projection_volume-brain_subcortical_for_CLA, best_injection_list
Copy_of_normalized_projection_volume-brain_subcortical_for_CLA
Copy_of_normalized_projection_volume_brain_for_CLA
Copy_of_normalized_projection_volume-brain_subcortical_for_CLA
Copy_of_normalized_projection_volume_brain_for_CLA
Copy_of_normalized_projection_volume_brain_for_CLA
Copy_of_normalized_projection_volume-brain_subcortical_for_CLA
Copy_of_normalized_projection_volume-brain_subcortical_for_CLA
Copy_of_normalized_projection_volume-brain_subcortical_for_CLA
Copy_of_normalized_projection_volume-brain_subcortical_for_CLA
Copy_of_normalized_projection_volume-brain_subcortical_for_CLA
Copy_of_normalized_projection_volume-brain_subcortical_for_CLA
Copy_of_normalized_projection_volume-brain_subcortical_for_CLA
Copy_of_normalized_projection_volume-brain_subcortical_for_CLA
Copy_of_normalized_projection_volume_brain_for_CLA
Copy_of_normalized_projection_volume_brain_for_CLA
Copy_of_normalized_projection_volume-brain_subcortical_for_CLA
Copy_of_normalized_projection_volume_brain_for_CLA
Copy_of_normalized_projection_volume_brain_for_CLA, best_injection_list
Copy_of_normalized_projection_volume_brain_for_CLA
Copy_of_normalized_projection_volume_brain_for_CLA
Copy_of_normalized_projection_volume_brain_for_CLA
Copy_of_normalized_projection_volume-brain_subcortical_for_CLA
Copy_of_normalized_projection_volume-brain_subcortical_for_CLA, best_injection_list
Copy_of_normalized_projection_volume-brain_subcortical_for_CLA, best_injection_list
Copy_of_normalized_projection_volume-brain_subcortical_for_CLA
Copy_of_normalized_projection_volume-brain_subcortical_for_CLA
Copy_of_normalized_projection_volume_brain_for_CLA
Copy_of_normalized_projection_volume_brain_for_CLA
Copy_of_normalized_projection_volume_brain_for_CLA
Copy_of_normalized_projection_volume-brain_subcortical_for_CLA
Copy_of_normalized_projection_volume-brain_subcortical_for_CLA, best_injection_list
Copy_of_normalized_projection_volume-brain_subcortical_for_CLA
Copy_of_normalized_projection_volume-brain_subcortical_for_CLA
Copy_of_normalized_projection_volume-brain_subcortical_for_CLA
Copy_of_normalized_projection_volume-brain_subcortical_for_CLA
Copy_of_normalized_projection_volume-brain_subcortical_for_CLA
Copy_of_normalized_projection_volume_brain_for_CLA
Copy_of_normalized_projection_volume_brain_for_CLA
Copy_of_normalized_projection_volume_brain_for_CLA

Copy_of_normalized_projection_volume_brain_for_CLA
Copy_of_normalized_projection_volume-brain_subcortical_for_CLA, best_injection_list
Copy_of_normalized_projection_volume-brain_subcortical_for_CLA
Copy_of_normalized_projection_volume-brain_subcortical_for_CLA
Copy_of_normalized_projection_volume_brain_for_CLA, best_injection_list
Copy_of_normalized_projection_volume-brain_subcortical_for_CLA
Copy_of_normalized_projection_volume-brain_subcortical_for_CLA
Copy_of_normalized_projection_volume-brain_subcortical_for_CLA
Copy_of_normalized_projection_volume-brain_subcortical_for_CLA
Copy_of_normalized_projection_volume-brain_subcortical_for_CLA
Copy_of_normalized_projection_volume-brain_subcortical_for_CLA
Copy_of_normalized_projection_volume-brain_subcortical_for_CLA
Copy_of_normalized_projection_volume-brain_subcortical_for_CLA
Copy_of_normalized_projection_volume_brain_for_CLA
Copy_of_normalized_projection_volume-brain_subcortical_for_CLA
Copy_of_normalized_projection_volume_brain_for_CLA
Copy_of_normalized_projection_volume_brain_for_CLA
Copy_of_normalized_projection_volume-brain_subcortical_for_CLA
Copy_of_normalized_projection_volume-brain_subcortical_for_CLA
Copy_of_normalized_projection_volume-brain_subcortical_for_CLA
Copy_of_normalized_projection_volume-brain_subcortical_for_CLA
Copy_of_normalized_projection_volume_brain_for_CLA, best_injection_list
Copy_of_normalized_projection_volume_brain_for_CLA, best_injection_list
Copy_of_normalized_projection_volume_brain_for_CLA
Copy_of_normalized_projection_volume_brain_for_CLA, best_injection_list
Copy_of_normalized_projection_volume_brain_for_CLA
Copy_of_normalized_projection_volume-brain_subcortical_for_CLA
Copy_of_normalized_projection_volume-brain_subcortical_for_CLA
Copy_of_normalized_projection_volume-brain_subcortical_for_CLA
Copy_of_normalized_projection_volume-brain_subcortical_for_CLA
Copy_of_normalized_projection_volume-brain_subcortical_for_CLA
Copy_of_normalized_projection_volume-brain_subcortical_for_CLA
Copy_of_normalized_projection_volume-brain_subcortical_for_CLA
Copy_of_normalized_projection_volume_brain_for_CLA
Copy_of_normalized_projection_volume-brain_subcortical_for_CLA
Copy_of_normalized_projection_volume_brain_for_CLA, best_injection_list
Copy_of_normalized_projection_volume-brain_subcortical_for_CLA
Copy_of_normalized_projection_volume-brain_subcortical_for_CLA
Copy_of_normalized_projection_volume_brain_for_CLA
Copy_of_normalized_projection_volume_brain_for_CLA
Copy_of_normalized_projection_volume-brain_subcortical_for_CLA
Copy_of_normalized_projection_volume-brain_subcortical_for_CLA
Copy_of_normalized_projection_volume-brain_subcortical_for_CLA
Copy_of_normalized_projection_volume-brain_subcortical_for_CLA
Copy_of_normalized_projection_volume_brain_for_CLA
Copy_of_normalized_projection_volume-brain_subcortical_for_CLA
Copy_of_normalized_projection_volume-brain_subcortical_for_CLA

Copy_of_normalized_projection_volume_brain_for_CLA
Copy_of_normalized_projection_volume-brain_subcortical_for_CLA
Copy_of_normalized_projection_volume_brain_for_CLA
Copy_of_normalized_projection_volume-brain_subcortical_for_CLA
Copy_of_normalized_projection_volume-brain_subcortical_for_CLA
Copy_of_normalized_projection_volume-brain_subcortical_for_CLA
Copy_of_normalized_projection_volume_brain_for_CLA
Copy_of_normalized_projection_volume_brain_for_CLA
Copy_of_normalized_projection_volume_brain_for_CLA
Copy_of_normalized_projection_volume_brain_for_CLA
Copy_of_normalized_projection_volume_brain_for_CLA
Copy_of_normalized_projection_volume_brain_for_CLA
Copy_of_normalized_projection_volume_brain_for_CLA
Copy_of_normalized_projection_volume_brain_for_CLA
Copy_of_normalized_projection_volume_brain_for_CLA
Copy_of_normalized_projection_volume_brain_for_CLA
Copy_of_normalized_projection_volume_brain_for_CLA
Copy_of_normalized_projection_volume-brain_subcortical_for_CLA
Copy_of_normalized_projection_volume-brain_subcortical_for_CLA
Copy_of_normalized_projection_volume_brain_for_CLA, best_injection_list
Copy_of_normalized_projection_volume_brain_for_CLA
Copy_of_normalized_projection_volume-brain_subcortical_for_CLA
Copy_of_normalized_projection_volume-brain_subcortical_for_CLA
Copy_of_normalized_projection_volume_brain_for_CLA
Copy_of_normalized_projection_volume_brain_for_CLA
Copy_of_normalized_projection_volume_brain_for_CLA
Copy_of_normalized_projection_volume_brain_for_CLA
Copy_of_normalized_projection_volume_brain_for_CLA
Copy_of_normalized_projection_volume_brain_for_CLA
Copy_of_normalized_projection_volume_brain_for_CLA
Copy_of_normalized_projection_volume_brain_for_CLA
Copy_of_normalized_projection_volume_brain_for_CLA
Copy_of_normalized_projection_volume_brain_for_CLA
Copy_of_normalized_projection_volume_brain_for_CLA
Copy_of_normalized_projection_volume_brain_for_CLA
Copy_of_normalized_projection_volume-brain_subcortical_for_CLA
Copy_of_normalized_projection_volume-brain_subcortical_for_CLA
Copy_of_normalized_projection_volume-brain_subcortical_for_CLA
Copy_of_normalized_projection_volume_brain_for_CLA
Copy_of_normalized_projection_volume-brain_subcortical_for_CLA
Copy_of_normalized_projection_volume-brain_subcortical_for_CLA
Copy_of_normalized_projection_volume-brain_subcortical_for_CLA
Copy_of_normalized_projection_volume-brain_subcortical_for_CLA
Copy_of_normalized_projection_volume_brain_for_CLA
Copy_of_normalized_projection_volume_brain_for_CLA
Copy_of_normalized_projection_volume_brain_for_CLA, best_injection_list
Copy_of_normalized_projection_volume_brain_for_CLA

Copy_of_normalized_projection_volume_brain_for_CLA
Copy_of_normalized_projection_volume_brain_for_CLA, best_injection_list
Copy_of_normalized_projection_volume-brain_subcortical_for_CLA
Copy_of_normalized_projection_volume-brain_subcortical_for_CLA
Copy_of_normalized_projection_volume-brain_subcortical_for_CLA
Copy_of_normalized_projection_volume-brain_subcortical_for_CLA
Copy_of_normalized_projection_volume-brain_subcortical_for_CLA
Copy_of_normalized_projection_volume-brain_subcortical_for_CLA
Copy_of_normalized_projection_volume-brain_subcortical_for_CLA
Copy_of_normalized_projection_volume-brain_subcortical_for_CLA
Copy_of_normalized_projection_volume_brain_for_CLA
Copy_of_normalized_projection_volume-brain_subcortical_for_CLA
Copy_of_normalized_projection_volume-brain_subcortical_for_CLA
Copy_of_normalized_projection_volume-brain_subcortical_for_CLA
Copy_of_normalized_projection_volume-brain_subcortical_for_CLA
Copy_of_normalized_projection_volume-brain_subcortical_for_CLA
Copy_of_normalized_projection_volume_brain_for_CLA
Copy_of_normalized_projection_volume_brain_for_CLA
Copy_of_normalized_projection_volume_brain_for_CLA
Copy_of_normalized_projection_volume_brain_for_CLA
Copy_of_normalized_projection_volume-brain_subcortical_for_CLA
Copy_of_normalized_projection_volume-brain_subcortical_for_CLA
Copy_of_normalized_projection_volume-brain_subcortical_for_CLA
Copy_of_normalized_projection_volume-brain_subcortical_for_CLA
Copy_of_normalized_projection_volume-brain_subcortical_for_CLA
Copy_of_normalized_projection_volume-brain_subcortical_for_CLA, best_injection_list
Copy_of_normalized_projection_volume-brain_subcortical_for_CLA
Copy_of_normalized_projection_volume-brain_subcortical_for_CLA
Copy_of_normalized_projection_volume-brain_subcortical_for_CLA, best_injection_list
Copy_of_normalized_projection_volume-brain_subcortical_for_CLA
Copy_of_normalized_projection_volume-brain_subcortical_for_CLA
Copy_of_normalized_projection_volume-brain_subcortical_for_CLA
Copy_of_normalized_projection_volume-brain_subcortical_for_CLA
Copy_of_normalized_projection_volume-brain_subcortical_for_CLA
Copy_of_normalized_projection_volume_brain_for_CLA
Copy_of_normalized_projection_volume-brain_subcortical_for_CLA
Copy_of_normalized_projection_volume-brain_subcortical_for_CLA
Copy_of_normalized_projection_volume-brain_subcortical_for_CLA
Copy_of_normalized_projection_volume-brain_subcortical_for_CLA
Copy_of_normalized_projection_volume-brain_subcortical_for_CLA
Copy_of_normalized_projection_volume-brain_subcortical_for_CLA
Copy_of_normalized_projection_volume-brain_subcortical_for_CLA
Copy_of_normalized_projection_volume-brain_subcortical_for_CLA
Copy_of_normalized_projection_volume-brain_subcortical_for_CLA
Copy_of_normalized_projection_volume_brain_for_CLA
Copy_of_normalized_projection_volume_brain_for_CLA, best_injection_list
Copy_of_normalized_projection_volume_brain_for_CLA

Copy_of_normalized_projection_volume_brain_for_CLA
Copy_of_normalized_projection_volume-brain_subcortical_for_CLA
Copy_of_normalized_projection_volume_brain_for_CLA, best_injection_list
Copy_of_normalized_projection_volume_brain_for_CLA
Copy_of_normalized_projection_volume-brain_subcortical_for_CLA
Copy_of_normalized_projection_volume_brain_for_CLA
Copy_of_normalized_projection_volume-brain_subcortical_for_CLA
Copy_of_normalized_projection_volume-brain_subcortical_for_CLA
Copy_of_normalized_projection_volume_brain_for_CLA
Copy_of_normalized_projection_volume-brain_subcortical_for_CLA
Copy_of_normalized_projection_volume_brain_for_CLA
Copy_of_normalized_projection_volume-brain_subcortical_for_CLA
Copy_of_normalized_projection_volume-brain_subcortical_for_CLA
Copy_of_normalized_projection_volume-brain_subcortical_for_CLA
Copy_of_normalized_projection_volume-brain_subcortical_for_CLA
Copy_of_normalized_projection_volume-brain_subcortical_for_CLA
Copy_of_normalized_projection_volume_brain_for_CLA
Copy_of_normalized_projection_volume_brain_for_CLA
Copy_of_normalized_projection_volume-brain_subcortical_for_CLA
Copy_of_normalized_projection_volume_brain_for_CLA
Copy_of_normalized_projection_volume_brain_for_CLA
Copy_of_normalized_projection_volume-brain_subcortical_for_CLA
Copy_of_normalized_projection_volume-brain_subcortical_for_CLA
Copy_of_normalized_projection_volume-brain_subcortical_for_CLA
Copy_of_normalized_projection_volume-brain_subcortical_for_CLA
Copy_of_normalized_projection_volume-brain_subcortical_for_CLA
Copy_of_normalized_projection_volume-brain_subcortical_for_CLA
Copy_of_normalized_projection_volume-brain_subcortical_for_CLA
Copy_of_normalized_projection_volume-brain_subcortical_for_CLA
Copy_of_normalized_projection_volume-brain_subcortical_for_CLA, best_injection_list
Copy_of_normalized_projection_volume-brain_subcortical_for_CLA
Copy_of_normalized_projection_volume-brain_subcortical_for_CLA
Copy_of_normalized_projection_volume_brain_for_CLA
Copy_of_normalized_projection_volume_brain_for_CLA
Copy_of_normalized_projection_volume_brain_for_CLA
Copy_of_normalized_projection_volume-brain_subcortical_for_CLA
Copy_of_normalized_projection_volume-brain_subcortical_for_CLA
Copy_of_normalized_projection_volume-brain_subcortical_for_CLA
Copy_of_normalized_projection_volume-brain_subcortical_for_CLA
Copy_of_normalized_projection_volume-brain_subcortical_for_CLA
Copy_of_normalized_projection_volume-brain_subcortical_for_CLA
Copy_of_normalized_projection_volume_brain_for_CLA
Copy_of_normalized_projection_volume_brain_for_CLA
Copy_of_normalized_projection_volume-brain_subcortical_for_CLA
Copy_of_normalized_projection_volume_brain_for_CLA, best_injection_list
Copy_of_normalized_projection_volume_brain_for_CLA, best_injection_list
Copy_of_normalized_projection_volume_brain_for_CLA, best_injection_list

Copy_of_normalized_projection_volume_brain_for_CLA
Copy_of_normalized_projection_volume_brain_for_CLA, best_injection_list
Copy_of_normalized_projection_volume_brain_for_CLA
Copy_of_normalized_projection_volume_brain_for_CLA
Copy_of_normalized_projection_volume_brain_for_CLA, best_injection_list
Copy_of_normalized_projection_volume_brain_for_CLA
Copy_of_normalized_projection_volume_brain_for_CLA
Copy_of_normalized_projection_volume_brain_for_CLA
CLA_layers_NPV
Copy_of_normalized_projection_volume_brain_for_CLA, best_injection_list
Copy_of_normalized_projection_volume_brain_for_CLA
Copy_of_normalized_projection_volume_brain_for_CLA
Copy_of_normalized_projection_volume_brain_for_CLA
Copy_of_normalized_projection_volume_brain_for_CLA
Copy_of_normalized_projection_volume_brain_for_CLA, best_injection_list
Copy_of_normalized_projection_volume_brain_for_CLA
Copy_of_normalized_projection_volume_brain_for_CLA
Copy_of_normalized_projection_volume_brain_for_CLA
Copy_of_normalized_projection_volume_brain_for_CLA
Copy_of_normalized_projection_volume_brain_for_CLA
Copy_of_normalized_projection_volume_brain_for_CLA
Copy_of_normalized_projection_volume_brain_for_CLA
Copy_of_normalized_projection_volume_brain_for_CLA
Copy_of_normalized_projection_volume_brain_for_CLA
Copy_of_normalized_projection_volume_brain_for_CLA
Copy_of_normalized_projection_volume_brain_for_CLA, best_injection_list
Copy_of_normalized_projection_volume_brain_for_CLA
Copy_of_normalized_projection_volume_brain_for_CLA
Copy_of_normalized_projection_volume_brain_for_CLA
Copy_of_normalized_projection_volume_brain_for_CLA
Copy_of_normalized_projection_volume_brain_for_CLA
Copy_of_normalized_projection_volume_brain_for_CLA
Copy_of_normalized_projection_volume_brain_for_CLA
Copy_of_normalized_projection_volume_brain_for_CLA, best_injection_list
Copy_of_normalized_projection_volume_brain_for_CLA, best_injection_list
Copy_of_normalized_projection_volume_brain_for_CLA
Copy_of_normalized_projection_volume-brain_subcortical_for_CLA
Copy_of_normalized_projection_volume_brain_for_CLA
Copy_of_normalized_projection_volume_brain_for_CLA
Copy_of_normalized_projection_volume_brain_for_CLA
Copy_of_normalized_projection_volume_brain_for_CLA
Copy_of_normalized_projection_volume_brain_for_CLA
Copy_of_normalized_projection_volume_brain_for_CLA
Copy_of_normalized_projection_volume_brain_for_CLA
Copy_of_normalized_projection_volume_brain_for_CLA
Copy_of_normalized_projection_volume_brain_for_CLA
Copy_of_normalized_projection_volume-brain_subcortical_for_CLA

Copy_of_normalized_projection_volume-brain_subcortical_for_CLA
Copy_of_normalized_projection_volume-brain_subcortical_for_CLA
Copy_of_normalized_projection_volume-brain_subcortical_for_CLA, best_injection_list
Copy_of_normalized_projection_volume-brain_subcortical_for_CLA
Copy_of_normalized_projection_volume-brain_subcortical_for_CLA
Copy_of_normalized_projection_volume-brain_subcortical_for_CLA
Copy_of_normalized_projection_volume-brain_subcortical_for_CLA
Copy_of_normalized_projection_volume-brain_subcortical_for_CLA, best_injection_list
Copy_of_normalized_projection_volume-brain_subcortical_for_CLA
Copy_of_normalized_projection_volume-brain_subcortical_for_CLA
Copy_of_normalized_projection_volume-brain_subcortical_for_CLA
Copy_of_normalized_projection_volume_brain_for_CLA
Copy_of_normalized_projection_volume_brain_for_CLA
Copy_of_normalized_projection_volume-brain_subcortical_for_CLA
Copy_of_normalized_projection_volume_brain_for_CLA, best_injection_list
Copy_of_normalized_projection_volume_brain_for_CLA
Copy_of_normalized_projection_volume_brain_for_CLA
Copy_of_normalized_projection_volume-brain_subcortical_for_CLA
Copy_of_normalized_projection_volume_brain_for_CLA
Copy_of_normalized_projection_volume_brain_for_CLA
Copy_of_normalized_projection_volume_brain_for_CLA
Copy_of_normalized_projection_volume_brain_for_CLA
Copy_of_normalized_projection_volume_brain_for_CLA
Copy_of_normalized_projection_volume_brain_for_CLA
Copy_of_normalized_projection_volume_brain_for_CLA
Copy_of_normalized_projection_volume_brain_for_CLA
Copy_of_normalized_projection_volume_brain_for_CLA
Copy_of_normalized_projection_volume_brain_for_CLA
Copy_of_normalized_projection_volume_brain_for_CLA
Copy_of_normalized_projection_volume_brain_for_CLA
Copy_of_normalized_projection_volume_brain_for_CLA
Copy_of_normalized_projection_volume_brain_for_CLA
Copy_of_normalized_projection_volume_brain_for_CLA
Copy_of_normalized_projection_volume_brain_for_CLA
Copy_of_normalized_projection_volume_brain_for_CLA, best_injection_list
Copy_of_normalized_projection_volume-brain_subcortical_for_CLA
Copy_of_normalized_projection_volume_brain_for_CLA
Copy_of_normalized_projection_volume_brain_for_CLA, best_injection_list
Copy_of_normalized_projection_volume_brain_for_CLA
Copy_of_normalized_projection_volume_brain_for_CLA
Copy_of_normalized_projection_volume_brain_for_CLA
Copy_of_normalized_projection_volume_brain_for_CLA
Copy_of_normalized_projection_volume-brain_subcortical_for_CLA
Copy_of_normalized_projection_volume-brain_subcortical_for_CLA
Copy_of_normalized_projection_volume_brain_for_CLA
Copy_of_normalized_projection_volume_brain_for_CLA
Copy_of_normalized_projection_volume-brain_subcortical_for_CLA

Copy_of_normalized_projection_volume-brain_subcortical_for_CLA
Copy_of_normalized_projection_volume-brain_subcortical_for_CLA
Copy_of_normalized_projection_volume-brain_subcortical_for_CLA
Copy_of_normalized_projection_volume-brain_subcortical_for_CLA
Copy_of_normalized_projection_volume-brain_subcortical_for_CLA
Copy_of_normalized_projection_volume-brain_subcortical_for_CLA
Copy_of_normalized_projection_volume-brain_subcortical_for_CLA, best_injection_list
Copy_of_normalized_projection_volume-brain_subcortical_for_CLA, best_injection_list
Copy_of_normalized_projection_volume-brain_subcortical_for_CLA
Copy_of_normalized_projection_volume-brain_subcortical_for_CLA
Copy_of_normalized_projection_volume_brain_for_CLA
Copy_of_normalized_projection_volume-brain_subcortical_for_CLA
Copy_of_normalized_projection_volume-brain_subcortical_for_CLA
Copy_of_normalized_projection_volume-brain_subcortical_for_CLA
Copy_of_normalized_projection_volume-brain_subcortical_for_CLA
Copy_of_normalized_projection_volume-brain_subcortical_for_CLA
Copy_of_normalized_projection_volume-brain_subcortical_for_CLA
Copy_of_normalized_projection_volume-brain_subcortical_for_CLA
Copy_of_normalized_projection_volume-brain_subcortical_for_CLA
Copy_of_normalized_projection_volume-brain_subcortical_for_CLA
Copy_of_normalized_projection_volume_brain_for_CLA
Copy_of_normalized_projection_volume-brain_subcortical_for_CLA
Copy_of_normalized_projection_volume-brain_subcortical_for_CLA
Copy_of_normalized_projection_volume-brain_subcortical_for_CLA
Copy_of_normalized_projection_volume-brain_subcortical_for_CLA
Copy_of_normalized_projection_volume-brain_subcortical_for_CLA
Copy_of_normalized_projection_volume_brain_for_CLA
Copy_of_normalized_projection_volume_brain_for_CLA
CLA_layers_NPV
Copy_of_normalized_projection_volume_brain_for_CLA
Copy_of_normalized_projection_volume_brain_for_CLA
Copy_of_normalized_projection_volume_brain_for_CLA
Copy_of_normalized_projection_volume_brain_for_CLA
Copy_of_normalized_projection_volume_brain_for_CLA, best_injection_list
Copy_of_normalized_projection_volume-brain_subcortical_for_CLA
Copy_of_normalized_projection_volume-brain_subcortical_for_CLA, best_injection_list
Copy_of_normalized_projection_volume-brain_subcortical_for_CLA
Copy_of_normalized_projection_volume_brain_for_CLA
Copy_of_normalized_projection_volume-brain_subcortical_for_CLA
Copy_of_normalized_projection_volume-brain_subcortical_for_CLA
Copy_of_normalized_projection_volume-brain_subcortical_for_CLA
Copy_of_normalized_projection_volume-brain_subcortical_for_CLA
Copy_of_normalized_projection_volume-brain_subcortical_for_CLA
Copy_of_normalized_projection_volume-brain_subcortical_for_CLA
Copy_of_normalized_projection_volume-brain_subcortical_for_CLA
Copy_of_normalized_projection_volume-brain_subcortical_for_CLA, best_injection_list
Copy_of_normalized_projection_volume-brain_subcortical_for_CLA

Copy_of_normalized_projection_volume-brain_subcortical_for_CLA
Copy_of_normalized_projection_volume-brain_subcortical_for_CLA
Copy_of_normalized_projection_volume-brain_subcortical_for_CLA
Copy_of_normalized_projection_volume-brain_subcortical_for_CLA
Copy_of_normalized_projection_volume-brain_subcortical_for_CLA
Copy_of_normalized_projection_volume-brain_subcortical_for_CLA
Copy_of_normalized_projection_volume-brain_subcortical_for_CLA
Copy_of_normalized_projection_volume-brain_subcortical_for_CLA
Copy_of_normalized_projection_volume_brain_for_CLA
Copy_of_normalized_projection_volume-brain_subcortical_for_CLA
Copy_of_normalized_projection_volume-brain_subcortical_for_CLA
Copy_of_normalized_projection_volume-brain_subcortical_for_CLA
Copy_of_normalized_projection_volume-brain_subcortical_for_CLA
Copy_of_normalized_projection_volume-brain_subcortical_for_CLA
Copy_of_normalized_projection_volume-brain_subcortical_for_CLA
Copy_of_normalized_projection_volume-brain_subcortical_for_CLA
Copy_of_normalized_projection_volume-brain_subcortical_for_CLA
Copy_of_normalized_projection_volume-brain_subcortical_for_CLA, best_injection_list
Copy_of_normalized_projection_volume-brain_subcortical_for_CLA
Copy_of_normalized_projection_volume-brain_subcortical_for_CLA
Copy_of_normalized_projection_volume-brain_subcortical_for_CLA
Copy_of_normalized_projection_volume-brain_subcortical_for_CLA
Copy_of_normalized_projection_volume-brain_subcortical_for_CLA, best_injection_list
Copy_of_normalized_projection_volume-brain_subcortical_for_CLA
Copy_of_normalized_projection_volume-brain_subcortical_for_CLA
Copy_of_normalized_projection_volume-brain_subcortical_for_CLA
Copy_of_normalized_projection_volume-brain_subcortical_for_CLA
Copy_of_normalized_projection_volume-brain_subcortical_for_CLA
Copy_of_normalized_projection_volume-brain_subcortical_for_CLA
Copy_of_normalized_projection_volume-brain_subcortical_for_CLA
Copy_of_normalized_projection_volume-brain_subcortical_for_CLA
Copy_of_normalized_projection_volume-brain_subcortical_for_CLA
Copy_of_normalized_projection_volume_brain_for_CLA
Copy_of_normalized_projection_volume-brain_subcortical_for_CLA
Copy_of_normalized_projection_volume-brain_subcortical_for_CLA
Copy_of_normalized_projection_volume-brain_subcortical_for_CLA
Copy_of_normalized_projection_volume-brain_subcortical_for_CLA
Copy_of_normalized_projection_volume-brain_subcortical_for_CLA
Copy_of_normalized_projection_volume-brain_subcortical_for_CLA
Copy_of_normalized_projection_volume-brain_subcortical_for_CLA
Copy_of_normalized_projection_volume-brain_subcortical_for_CLA
Copy_of_normalized_projection_volume-brain_subcortical_for_CLA
Copy_of_normalized_projection_volume-brain_subcortical_for_CLA
Copy_of_normalized_projection_volume_brain_for_CLA
Copy_of_normalized_projection_volume_brain_for_CLA
Copy_of_normalized_projection_volume-brain_subcortical_for_CLA
Copy_of_normalized_projection_volume-brain_subcortical_for_CLA, best_injection_list

Copy_of_normalized_projection_volume-brain_subcortical_for_CLA
Copy_of_normalized_projection_volume_brain_for_CLA
Copy_of_normalized_projection_volume-brain_subcortical_for_CLA
Copy_of_normalized_projection_volume-brain_subcortical_for_CLA
Copy_of_normalized_projection_volume_brain_for_CLA
Copy_of_normalized_projection_volume-brain_subcortical_for_CLA
Copy_of_normalized_projection_volume-brain_subcortical_for_CLA
Copy_of_normalized_projection_volume-brain_subcortical_for_CLA
Copy_of_normalized_projection_volume-brain_subcortical_for_CLA
Copy_of_normalized_projection_volume-brain_subcortical_for_CLA
Copy_of_normalized_projection_volume-brain_subcortical_for_CLA
Copy_of_normalized_projection_volume-brain_subcortical_for_CLA
Copy_of_normalized_projection_volume-brain_subcortical_for_CLA
Copy_of_normalized_projection_volume-brain_subcortical_for_CLA
Copy_of_normalized_projection_volume-brain_subcortical_for_CLA
Copy_of_normalized_projection_volume-brain_subcortical_for_CLA
Copy_of_normalized_projection_volume-brain_subcortical_for_CLA
Copy_of_normalized_projection_volume-brain_subcortical_for_CLA
Copy_of_normalized_projection_volume-brain_subcortical_for_CLA
Copy_of_normalized_projection_volume_brain_for_CLA
Copy_of_normalized_projection_volume_brain_for_CLA
Copy_of_normalized_projection_volume_brain_for_CLA
Copy_of_normalized_projection_volume_brain_for_CLA
Copy_of_normalized_projection_volume_brain_for_CLA
Copy_of_normalized_projection_volume_brain_for_CLA
Copy_of_normalized_projection_volume-brain_subcortical_for_CLA
Copy_of_normalized_projection_volume-brain_subcortical_for_CLA
Copy_of_normalized_projection_volume-brain_subcortical_for_CLA
Copy_of_normalized_projection_volume-brain_subcortical_for_CLA
Copy_of_normalized_projection_volume-brain_subcortical_for_CLA
Copy_of_normalized_projection_volume-brain_subcortical_for_CLA
Copy_of_normalized_projection_volume-brain_subcortical_for_CLA
Copy_of_normalized_projection_volume-brain_subcortical_for_CLA
Copy_of_normalized_projection_volume-brain_subcortical_for_CLA
Copy_of_normalized_projection_volume-brain_subcortical_for_CLA
Copy_of_normalized_projection_volume-brain_subcortical_for_CLA
Copy_of_normalized_projection_volume-brain_subcortical_for_CLA
Copy_of_normalized_projection_volume-brain_subcortical_for_CLA
Copy_of_normalized_projection_volume-brain_subcortical_for_CLA
Copy_of_normalized_projection_volume-brain_subcortical_for_CLA
Copy_of_normalized_projection_volume-brain_subcortical_for_CLA
Copy_of_normalized_projection_volume-brain_subcortical_for_CLA
Copy_of_normalized_projection_volume-brain_subcortical_for_CLA
Copy_of_normalized_projection_volume-brain_subcortical_for_CLA
Copy_of_normalized_projection_volume-brain_subcortical_for_CLA
Copy_of_normalized_projection_volume-brain_subcortical_for_CLA
Copy_of_normalized_projection_volume-brain_subcortical_for_CLA

Copy_of_normalized_projection_volume-brain_subcortical_for_CLA
Copy_of_normalized_projection_volume-brain_subcortical_for_CLA
Copy_of_normalized_projection_volume-brain_subcortical_for_CLA
Copy_of_normalized_projection_volume-brain_subcortical_for_CLA
Copy_of_normalized_projection_volume-brain_subcortical_for_CLA
Copy_of_normalized_projection_volume-brain_subcortical_for_CLA
Copy_of_normalized_projection_volume-brain_subcortical_for_CLA
Copy_of_normalized_projection_volume-brain_subcortical_for_CLA
Copy_of_normalized_projection_volume-brain_subcortical_for_CLA
Copy_of_normalized_projection_volume-brain_subcortical_for_CLA
Copy_of_normalized_projection_volume-brain_subcortical_for_CLA
Copy_of_normalized_projection_volume-brain_subcortical_for_CLA
Copy_of_normalized_projection_volume-brain_subcortical_for_CLA
Copy_of_normalized_projection_volume-brain_subcortical_for_CLA
Copy_of_normalized_projection_volume-brain_subcortical_for_CLA
Copy_of_normalized_projection_volume-brain_subcortical_for_CLA
Copy_of_normalized_projection_volume-brain_subcortical_for_CLA
Copy_of_normalized_projection_volume-brain_subcortical_for_CLA
Copy_of_normalized_projection_volume-brain_subcortical_for_CLA
Copy_of_normalized_projection_volume-brain_subcortical_for_CLA
Copy_of_normalized_projection_volume-brain_subcortical_for_CLA
Copy_of_normalized_projection_volume-brain_subcortical_for_CLA
Copy_of_normalized_projection_volume-brain_subcortical_for_CLA
Copy_of_normalized_projection_volume-brain_subcortical_for_CLA
Copy_of_normalized_projection_volume-brain_subcortical_for_CLA
Copy_of_normalized_projection_volume_brain_for_CLA
Copy_of_normalized_projection_volume_brain_for_CLA
Copy_of_normalized_projection_volume_brain_for_CLA
Copy_of_normalized_projection_volume_brain_for_CLA
Copy_of_normalized_projection_volume_brain_for_CLA
Copy_of_normalized_projection_volume_brain_for_CLA
Copy_of_normalized_projection_volume-brain_subcortical_for_CLA
Copy_of_normalized_projection_volume-brain_subcortical_for_CLA
Copy_of_normalized_projection_volume-brain_subcortical_for_CLA, best_injection_list
Copy_of_normalized_projection_volume_brain_for_CLA
Copy_of_normalized_projection_volume_brain_for_CLA, best_injection_list
Copy_of_normalized_projection_volume_brain_for_CLA
Copy_of_normalized_projection_volume_brain_for_CLA
Copy_of_normalized_projection_volume_brain_for_CLA
Copy_of_normalized_projection_volume-brain_subcortical_for_CLA
Copy_of_normalized_projection_volume-brain_subcortical_for_CLA
Copy_of_normalized_projection_volume-brain_subcortical_for_CLA
Copy_of_normalized_projection_volume-brain_subcortical_for_CLA, best_injection_list
Copy_of_normalized_projection_volume-brain_subcortical_for_CLA
Copy_of_normalized_projection_volume-brain_subcortical_for_CLA
Copy_of_normalized_projection_volume-brain_subcortical_for_CLA
Copy_of_normalized_projection_volume-brain_subcortical_for_CLA

Copy_of_normalized_projection_volume_brain_for_CLA
Copy_of_normalized_projection_volume-brain_subcortical_for_CLA
Copy_of_normalized_projection_volume-brain_subcortical_for_CLA
Copy_of_normalized_projection_volume-brain_subcortical_for_CLA
Copy_of_normalized_projection_volume-brain_subcortical_for_CLA
Copy_of_normalized_projection_volume-brain_subcortical_for_CLA
Copy_of_normalized_projection_volume-brain_subcortical_for_CLA
Copy_of_normalized_projection_volume-brain_subcortical_for_CLA
Copy_of_normalized_projection_volume-brain_subcortical_for_CLA
Copy_of_normalized_projection_volume-brain_subcortical_for_CLA
Copy_of_normalized_projection_volume-brain_subcortical_for_CLA
Copy_of_normalized_projection_volume-brain_subcortical_for_CLA
Copy_of_normalized_projection_volume-brain_subcortical_for_CLA
Copy_of_normalized_projection_volume-brain_subcortical_for_CLA
Copy_of_normalized_projection_volume-brain_subcortical_for_CLA
Copy_of_normalized_projection_volume-brain_subcortical_for_CLA
Copy_of_normalized_projection_volume-brain_subcortical_for_CLA
Copy_of_normalized_projection_volume-brain_subcortical_for_CLA
Copy_of_normalized_projection_volume-brain_subcortical_for_CLA
Copy_of_normalized_projection_volume-brain_subcortical_for_CLA
Copy_of_normalized_projection_volume-brain_subcortical_for_CLA
Copy_of_normalized_projection_volume-brain_subcortical_for_CLA
Copy_of_normalized_projection_volume-brain_subcortical_for_CLA
Copy_of_normalized_projection_volume-brain_subcortical_for_CLA
Copy_of_normalized_projection_volume-brain_subcortical_for_CLA
Copy_of_normalized_projection_volume-brain_subcortical_for_CLA
Copy_of_normalized_projection_volume-brain_subcortical_for_CLA
Copy_of_normalized_projection_volume-brain_subcortical_for_CLA
Copy_of_normalized_projection_volume-brain_subcortical_for_CLA
Copy_of_normalized_projection_volume-brain_subcortical_for_CLA
Copy_of_normalized_projection_volume-brain_subcortical_for_CLA
Copy_of_normalized_projection_volume-brain_subcortical_for_CLA
Copy_of_normalized_projection_volume-brain_subcortical_for_CLA, best_injection_list
Copy_of_normalized_projection_volume-brain_subcortical_for_CLA
Copy_of_normalized_projection_volume-brain_subcortical_for_CLA
Copy_of_normalized_projection_volume-brain_subcortical_for_CLA
Copy_of_normalized_projection_volume-brain_subcortical_for_CLA
Copy_of_normalized_projection_volume-brain_subcortical_for_CLA
Copy_of_normalized_projection_volume-brain_subcortical_for_CLA
Copy_of_normalized_projection_volume-brain_subcortical_for_CLA
Copy_of_normalized_projection_volume-brain_subcortical_for_CLA
Copy_of_normalized_projection_volume-brain_subcortical_for_CLA
Copy_of_normalized_projection_volume-brain_subcortical_for_CLA
Copy_of_normalized_projection_volume-brain_subcortical_for_CLA
Copy_of_normalized_projection_volume-brain_subcortical_for_CLA
Copy_of_normalized_projection_volume-brain_subcortical_for_CLA
Copy_of_normalized_projection_volume-brain_subcortical_for_CLA

Copy_of_normalized_projection_volume-brain_subcortical_for_CLA
Copy_of_normalized_projection_volume-brain_subcortical_for_CLA
Copy_of_normalized_projection_volume-brain_subcortical_for_CLA
Copy_of_normalized_projection_volume-brain_subcortical_for_CLA
Copy_of_normalized_projection_volume-brain_subcortical_for_CLA
Copy_of_normalized_projection_volume-brain_subcortical_for_CLA, best_injection_list
Copy_of_normalized_projection_volume_brain_for_CLA
Copy_of_normalized_projection_volume_brain_for_CLA
Copy_of_normalized_projection_volume_brain_for_CLA
Copy_of_normalized_projection_volume_brain_for_CLA
Copy_of_normalized_projection_volume_brain_for_CLA
Copy_of_normalized_projection_volume_brain_for_CLA, best_injection_list
Copy_of_normalized_projection_volume-brain_subcortical_for_CLA
Copy_of_normalized_projection_volume-brain_subcortical_for_CLA
Copy_of_normalized_projection_volume-brain_subcortical_for_CLA
Copy_of_normalized_projection_volume-brain_subcortical_for_CLA
Copy_of_normalized_projection_volume_brain_for_CLA
Copy_of_normalized_projection_volume-brain_subcortical_for_CLA
Copy_of_normalized_projection_volume-brain_subcortical_for_CLA
Copy_of_normalized_projection_volume-brain_subcortical_for_CLA
Copy_of_normalized_projection_volume_brain_for_CLA
Copy_of_normalized_projection_volume_brain_for_CLA
Copy_of_normalized_projection_volume_brain_for_CLA
Copy_of_normalized_projection_volume-brain_subcortical_for_CLA
Copy_of_normalized_projection_volume-brain_subcortical_for_CLA
Copy_of_normalized_projection_volume-brain_subcortical_for_CLA
Copy_of_normalized_projection_volume-brain_subcortical_for_CLA
Copy_of_normalized_projection_volume-brain_subcortical_for_CLA
Copy_of_normalized_projection_volume-brain_subcortical_for_CLA
Copy_of_normalized_projection_volume_brain_for_CLA
Copy_of_normalized_projection_volume-brain_subcortical_for_CLA
Copy_of_normalized_projection_volume-brain_subcortical_for_CLA
Copy_of_normalized_projection_volume_brain_for_CLA
Copy_of_normalized_projection_volume_brain_for_CLA
Copy_of_normalized_projection_volume-brain_subcortical_for_CLA
Copy_of_normalized_projection_volume-brain_subcortical_for_CLA
Copy_of_normalized_projection_volume-brain_subcortical_for_CLA
Copy_of_normalized_projection_volume-brain_subcortical_for_CLA
Copy_of_normalized_projection_volume-brain_subcortical_for_CLA
Copy_of_normalized_projection_volume-brain_subcortical_for_CLA
Copy_of_normalized_projection_volume-brain_subcortical_for_CLA
Copy_of_normalized_projection_volume_brain_for_CLA
Copy_of_normalized_projection_volume_brain_for_CLA
Copy_of_normalized_projection_volume-brain_subcortical_for_CLA
Copy_of_normalized_projection_volume-brain_subcortical_for_CLA
Copy_of_normalized_projection_volume-brain_subcortical_for_CLA
Copy_of_normalized_projection_volume_brain_for_CLA

Copy_of_normalized_projection_volume_brain_for_CLA
Copy_of_normalized_projection_volume_brain_for_CLA
Copy_of_normalized_projection_volume_brain_for_CLA
Copy_of_normalized_projection_volume_brain_for_CLA
Copy_of_normalized_projection_volume_brain_for_CLA
Copy_of_normalized_projection_volume-brain_subcortical_for_CLA
Copy_of_normalized_projection_volume-brain_subcortical_for_CLA
Copy_of_normalized_projection_volume-brain_subcortical_for_CLA
Copy_of_normalized_projection_volume-brain_subcortical_for_CLA
Copy_of_normalized_projection_volume-brain_subcortical_for_CLA
Copy_of_normalized_projection_volume-brain_subcortical_for_CLA
Copy_of_normalized_projection_volume-brain_subcortical_for_CLA
Copy_of_normalized_projection_volume-brain_subcortical_for_CLA
Copy_of_normalized_projection_volume-brain_subcortical_for_CLA
Copy_of_normalized_projection_volume_brain_for_CLA
Copy_of_normalized_projection_volume-brain_subcortical_for_CLA
Copy_of_normalized_projection_volume-brain_subcortical_for_CLA
Copy_of_normalized_projection_volume-brain_subcortical_for_CLA
Copy_of_normalized_projection_volume-brain_subcortical_for_CLA
Copy_of_normalized_projection_volume-brain_subcortical_for_CLA
Copy_of_normalized_projection_volume-brain_subcortical_for_CLA
Copy_of_normalized_projection_volume-brain_subcortical_for_CLA
Copy_of_normalized_projection_volume-brain_subcortical_for_CLA
Copy_of_normalized_projection_volume-brain_subcortical_for_CLA
Copy_of_normalized_projection_volume-brain_subcortical_for_CLA
Copy_of_normalized_projection_volume-brain_subcortical_for_CLA
Copy_of_normalized_projection_volume-brain_subcortical_for_CLA
Copy_of_normalized_projection_volume-brain_subcortical_for_CLA, best_injection_list
Copy_of_normalized_projection_volume-brain_subcortical_for_CLA
Copy_of_normalized_projection_volume_brain_for_CLA, best_injection_list
Copy_of_normalized_projection_volume_brain_for_CLA
Copy_of_normalized_projection_volume_brain_for_CLA
Copy_of_normalized_projection_volume_brain_for_CLA
Copy_of_normalized_projection_volume_brain_for_CLA
Copy_of_normalized_projection_volume_brain_for_CLA
Copy_of_normalized_projection_volume_brain_for_CLA
Copy_of_normalized_projection_volume_brain_for_CLA
Copy_of_normalized_projection_volume_brain_for_CLA
Copy_of_normalized_projection_volume-brain_subcortical_for_CLA
Copy_of_normalized_projection_volume-brain_subcortical_for_CLA
Copy_of_normalized_projection_volume-brain_subcortical_for_CLA
Copy_of_normalized_projection_volume-brain_subcortical_for_CLA
Copy_of_normalized_projection_volume-brain_subcortical_for_CLA
Copy_of_normalized_projection_volume-brain_subcortical_for_CLA
Copy_of_normalized_projection_volume_brain_for_CLA
Copy_of_normalized_projection_volume_brain_for_CLA, best_injection_list
Copy_of_normalized_projection_volume_brain_for_CLA

Copy_of_normalized_projection_volume-brain_subcortical_for_CLA
Copy_of_normalized_projection_volume-brain_subcortical_for_CLA
Copy_of_normalized_projection_volume_brain_for_CLA
Copy_of_normalized_projection_volume_brain_for_CLA
Copy_of_normalized_projection_volume-brain_subcortical_for_CLA
Copy_of_normalized_projection_volume-brain_subcortical_for_CLA
Copy_of_normalized_projection_volume-brain_subcortical_for_CLA
Copy_of_normalized_projection_volume-brain_subcortical_for_CLA, CLA_layers_NPV
Copy_of_normalized_projection_volume_brain_for_CLA, best_injection_list
Copy_of_normalized_projection_volume-brain_subcortical_for_CLA
Copy_of_normalized_projection_volume-brain_subcortical_for_CLA, CLA_layers_NPV
Copy_of_normalized_projection_volume-brain_subcortical_for_CLA, best_injection_list
Copy_of_normalized_projection_volume-brain_subcortical_for_CLA
Copy_of_normalized_projection_volume-brain_subcortical_for_CLA
Copy_of_normalized_projection_volume_brain_for_CLA
Copy_of_normalized_projection_volume-brain_subcortical_for_CLA
Copy_of_normalized_projection_volume-brain_subcortical_for_CLA
Copy_of_normalized_projection_volume-brain_subcortical_for_CLA
Copy_of_normalized_projection_volume-brain_subcortical_for_CLA
Copy_of_normalized_projection_volume-brain_subcortical_for_CLA
Copy_of_normalized_projection_volume-brain_subcortical_for_CLA
Copy_of_normalized_projection_volume-brain_subcortical_for_CLA
Copy_of_normalized_projection_volume_brain_for_CLA
Copy_of_normalized_projection_volume_brain_for_CLA
Copy_of_normalized_projection_volume-brain_subcortical_for_CLA
Copy_of_normalized_projection_volume-brain_subcortical_for_CLA
Copy_of_normalized_projection_volume-brain_subcortical_for_CLA, best_injection_list
Copy_of_normalized_projection_volume_brain_for_CLA
Copy_of_normalized_projection_volume_brain_for_CLA
Copy_of_normalized_projection_volume_brain_for_CLA, CLA_layers_NPV
Copy_of_normalized_projection_volume-brain_subcortical_for_CLA, CLA_layers_NPV
CLA_layers_NPV
Copy_of_normalized_projection_volume_brain_for_CLA, best_injection_list
Copy_of_normalized_projection_volume-brain_subcortical_for_CLA
Copy_of_normalized_projection_volume-brain_subcortical_for_CLA
Copy_of_normalized_projection_volume-brain_subcortical_for_CLA
Copy_of_normalized_projection_volume-brain_subcortical_for_CLA
Copy_of_normalized_projection_volume-brain_subcortical_for_CLA, best_injection_list
Copy_of_normalized_projection_volume_brain_for_CLA
Copy_of_normalized_projection_volume-brain_subcortical_for_CLA
Copy_of_normalized_projection_volume_brain_for_CLA
Copy_of_normalized_projection_volume_brain_for_CLA, best_injection_list
Copy_of_normalized_projection_volume-brain_subcortical_for_CLA, best_injection_list
Copy_of_normalized_projection_volume_brain_for_CLA
Copy_of_normalized_projection_volume-brain_subcortical_for_CLA
Copy_of_normalized_projection_volume-brain_subcortical_for_CLA
Copy_of_normalized_projection_volume-brain_subcortical_for_CLA, best_injection_list

Copy_of_normalized_projection_volume-brain_subcortical_for_CLA
Copy_of_normalized_projection_volume-brain_subcortical_for_CLA
Copy_of_normalized_projection_volume-brain_subcortical_for_CLA
Copy_of_normalized_projection_volume-brain_subcortical_for_CLA
Copy_of_normalized_projection_volume_brain_for_CLA
Copy_of_normalized_projection_volume_brain_for_CLA, best_injection_list
Copy_of_normalized_projection_volume_brain_for_CLA
Copy_of_normalized_projection_volume_brain_for_CLA, best_injection_list
Copy_of_normalized_projection_volume-brain_subcortical_for_CLA
Copy_of_normalized_projection_volume_brain_for_CLA
Copy_of_normalized_projection_volume-brain_subcortical_for_CLA
Copy_of_normalized_projection_volume-brain_subcortical_for_CLA
Copy_of_normalized_projection_volume_brain_for_CLA
Copy_of_normalized_projection_volume_brain_for_CLA
Copy_of_normalized_projection_volume-brain_subcortical_for_CLA
Copy_of_normalized_projection_volume_brain_for_CLA
Copy_of_normalized_projection_volume-brain_subcortical_for_CLA
Copy_of_normalized_projection_volume-brain_subcortical_for_CLA
Copy_of_normalized_projection_volume_brain_for_CLA
Copy_of_normalized_projection_volume-brain_subcortical_for_CLA
Copy_of_normalized_projection_volume_brain_for_CLA
Copy_of_normalized_projection_volume_brain_for_CLA
Copy_of_normalized_projection_volume_brain_for_CLA, best_injection_list
Copy_of_normalized_projection_volume-brain_subcortical_for_CLA, best_injection_list
Copy_of_normalized_projection_volume-brain_subcortical_for_CLA
Copy_of_normalized_projection_volume-brain_subcortical_for_CLA
Copy_of_normalized_projection_volume-brain_subcortical_for_CLA
Copy_of_normalized_projection_volume-brain_subcortical_for_CLA
Copy_of_normalized_projection_volume-brain_subcortical_for_CLA
Copy_of_normalized_projection_volume-brain_subcortical_for_CLA
Copy_of_normalized_projection_volume-brain_subcortical_for_CLA
Copy_of_normalized_projection_volume-brain_subcortical_for_CLA
Copy_of_normalized_projection_volume-brain_subcortical_for_CLA
Copy_of_normalized_projection_volume-brain_subcortical_for_CLA
Copy_of_normalized_projection_volume-brain_subcortical_for_CLA
Copy_of_normalized_projection_volume-brain_subcortical_for_CLA, best_injection_list
Copy_of_normalized_projection_volume_brain_for_CLA
Copy_of_normalized_projection_volume-brain_subcortical_for_CLA, best_injection_list
Copy_of_normalized_projection_volume-brain_subcortical_for_CLA
Copy_of_normalized_projection_volume-brain_subcortical_for_CLA
Copy_of_normalized_projection_volume-brain_subcortical_for_CLA
Copy_of_normalized_projection_volume-brain_subcortical_for_CLA
Copy_of_normalized_projection_volume-brain_subcortical_for_CLA
Copy_of_normalized_projection_volume-brain_subcortical_for_CLA
Copy_of_normalized_projection_volume-brain_subcortical_for_CLA
Copy_of_normalized_projection_volume-brain_subcortical_for_CLA
Copy_of_normalized_projection_volume-brain_subcortical_for_CLA

Copy_of_normalized_projection_volume-brain_subcortical_for_CLA
Copy_of_normalized_projection_volume-brain_subcortical_for_CLA
Copy_of_normalized_projection_volume-brain_subcortical_for_CLA
Copy_of_normalized_projection_volume-brain_subcortical_for_CLA, best_injection_list
Copy_of_normalized_projection_volume_brain_for_CLA, best_injection_list
Copy_of_normalized_projection_volume_brain_for_CLA
Copy_of_normalized_projection_volume-brain_subcortical_for_CLA, best_injection_list
Copy_of_normalized_projection_volume-brain_subcortical_for_CLA
Copy_of_normalized_projection_volume-brain_subcortical_for_CLA
Copy_of_normalized_projection_volume-brain_subcortical_for_CLA
Copy_of_normalized_projection_volume-brain_subcortical_for_CLA
Copy_of_normalized_projection_volume-brain_subcortical_for_CLA
Copy_of_normalized_projection_volume-brain_subcortical_for_CLA
Copy_of_normalized_projection_volume-brain_subcortical_for_CLA
Copy_of_normalized_projection_volume-brain_subcortical_for_CLA, best_injection_list
Copy_of_normalized_projection_volume_brain_for_CLA
Copy_of_normalized_projection_volume-brain_subcortical_for_CLA
Copy_of_normalized_projection_volume-brain_subcortical_for_CLA
Copy_of_normalized_projection_volume-brain_subcortical_for_CLA
Copy_of_normalized_projection_volume-brain_subcortical_for_CLA
Copy_of_normalized_projection_volume-brain_subcortical_for_CLA
Copy_of_normalized_projection_volume-brain_subcortical_for_CLA
Copy_of_normalized_projection_volume-brain_subcortical_for_CLA, best_injection_list
Copy_of_normalized_projection_volume-brain_subcortical_for_CLA
Copy_of_normalized_projection_volume-brain_subcortical_for_CLA, best_injection_list
Copy_of_normalized_projection_volume_brain_for_CLA
Copy_of_normalized_projection_volume-brain_subcortical_for_CLA, best_injection_list
Copy_of_normalized_projection_volume-brain_subcortical_for_CLA
Copy_of_normalized_projection_volume-brain_subcortical_for_CLA
Copy_of_normalized_projection_volume-brain_subcortical_for_CLA
Copy_of_normalized_projection_volume_brain_for_CLA
Copy_of_normalized_projection_volume-brain_subcortical_for_CLA
Copy_of_normalized_projection_volume-brain_subcortical_for_CLA
Copy_of_normalized_projection_volume-brain_subcortical_for_CLA
Copy_of_normalized_projection_volume_brain_for_CLA
Copy_of_normalized_projection_volume_brain_for_CLA
Copy_of_normalized_projection_volume_brain_for_CLA
Copy_of_normalized_projection_volume_brain_for_CLA
Copy_of_normalized_projection_volume-brain_subcortical_for_CLA
Copy_of_normalized_projection_volume-brain_subcortical_for_CLA
Copy_of_normalized_projection_volume-brain_subcortical_for_CLA
Copy_of_normalized_projection_volume-brain_subcortical_for_CLA
Copy_of_normalized_projection_volume-brain_subcortical_for_CLA
Copy_of_normalized_projection_volume_brain_for_CLA
Copy_of_normalized_projection_volume-brain_subcortical_for_CLA
Copy_of_normalized_projection_volume_brain_for_CLA
Copy_of_normalized_projection_volume_brain_for_CLA

Copy_of_normalized_projection_volume_brain_for_CLA
Copy_of_normalized_projection_volume-brain_subcortical_for_CLA
Copy_of_normalized_projection_volume-brain_subcortical_for_CLA
Copy_of_normalized_projection_volume-brain_subcortical_for_CLA
Copy_of_normalized_projection_volume-brain_subcortical_for_CLA
Copy_of_normalized_projection_volume_brain_for_CLA
Copy_of_normalized_projection_volume_brain_for_CLA, best_injection_list
Copy_of_normalized_projection_volume_brain_for_CLA, best_injection_list
Copy_of_normalized_projection_volume_brain_for_CLA
Copy_of_normalized_projection_volume_brain_for_CLA
Copy_of_normalized_projection_volume_brain_for_CLA, best_injection_list
Copy_of_normalized_projection_volume_brain_for_CLA
Copy_of_normalized_projection_volume_brain_for_CLA
Copy_of_normalized_projection_volume-brain_subcortical_for_CLA, best_injection_list
Copy_of_normalized_projection_volume-brain_subcortical_for_CLA
Copy_of_normalized_projection_volume-brain_subcortical_for_CLA
Copy_of_normalized_projection_volume-brain_subcortical_for_CLA
Copy_of_normalized_projection_volume-brain_subcortical_for_CLA
Copy_of_normalized_projection_volume-brain_subcortical_for_CLA
Copy_of_normalized_projection_volume_brain_for_CLA
Copy_of_normalized_projection_volume_brain_for_CLA
Copy_of_normalized_projection_volume_brain_for_CLA, best_injection_list
Copy_of_normalized_projection_volume_brain_for_CLA
Copy_of_normalized_projection_volume_brain_for_CLA
Copy_of_normalized_projection_volume_brain_for_CLA
Copy_of_normalized_projection_volume_brain_for_CLA
Copy_of_normalized_projection_volume_brain_for_CLA
Copy_of_normalized_projection_volume_brain_for_CLA
Copy_of_normalized_projection_volume_brain_for_CLA
Copy_of_normalized_projection_volume_brain_for_CLA
Copy_of_normalized_projection_volume_brain_for_CLA
Copy_of_normalized_projection_volume_brain_for_CLA
Copy_of_normalized_projection_volume_brain_for_CLA
Copy_of_normalized_projection_volume_brain_for_CLA
Copy_of_normalized_projection_volume_brain_for_CLA
Copy_of_normalized_projection_volume_brain_for_CLA
Copy_of_normalized_projection_volume-brain_subcortical_for_CLA
Copy_of_normalized_projection_volume_brain_for_CLA
Copy_of_normalized_projection_volume_brain_for_CLA
Copy_of_normalized_projection_volume_brain_for_CLA
Copy_of_normalized_projection_volume_brain_for_CLA
Copy_of_normalized_projection_volume_brain_for_CLA
Copy_of_normalized_projection_volume-brain_subcortical_for_CLA
Copy_of_normalized_projection_volume-brain_subcortical_for_CLA
Copy_of_normalized_projection_volume-brain_subcortical_for_CLA
Copy_of_normalized_projection_volume-brain_subcortical_for_CLA
Copy_of_normalized_projection_volume-brain_subcortical_for_CLA

Copy_of_normalized_projection_volume-brain_subcortical_for_CLA
Copy_of_normalized_projection_volume-brain_subcortical_for_CLA
Copy_of_normalized_projection_volume-brain_subcortical_for_CLA
Copy_of_normalized_projection_volume_brain_for_CLA
Copy_of_normalized_projection_volume_brain_for_CLA
Copy_of_normalized_projection_volume_brain_for_CLA
Copy_of_normalized_projection_volume_brain_for_CLA
Copy_of_normalized_projection_volume_brain_for_CLA
Copy_of_normalized_projection_volume_brain_for_CLA
Copy_of_normalized_projection_volume_brain_for_CLA
Copy_of_normalized_projection_volume_brain_for_CLA
Copy_of_normalized_projection_volume_brain_for_CLA
Copy_of_normalized_projection_volume_brain_for_CLA
Copy_of_normalized_projection_volume_brain_for_CLA
Copy_of_normalized_projection_volume-brain_subcortical_for_CLA
Copy_of_normalized_projection_volume-brain_subcortical_for_CLA
Copy_of_normalized_projection_volume-brain_subcortical_for_CLA, best_injection_list
Copy_of_normalized_projection_volume_brain_for_CLA, best_injection_list
Copy_of_normalized_projection_volume_brain_for_CLA
Copy_of_normalized_projection_volume_brain_for_CLA
Copy_of_normalized_projection_volume-brain_subcortical_for_CLA
Copy_of_normalized_projection_volume-brain_subcortical_for_CLA
Copy_of_normalized_projection_volume_brain_for_CLA
Copy_of_normalized_projection_volume_brain_for_CLA
Copy_of_normalized_projection_volume-brain_subcortical_for_CLA
Copy_of_normalized_projection_volume_brain_for_CLA
Copy_of_normalized_projection_volume_brain_for_CLA
Copy_of_normalized_projection_volume_brain_for_CLA
Copy_of_normalized_projection_volume-brain_subcortical_for_CLA
Copy_of_normalized_projection_volume-brain_subcortical_for_CLA
Copy_of_normalized_projection_volume-brain_subcortical_for_CLA
Copy_of_normalized_projection_volume-brain_subcortical_for_CLA
Copy_of_normalized_projection_volume_brain_for_CLA
Copy_of_normalized_projection_volume-brain_subcortical_for_CLA
Copy_of_normalized_projection_volume-brain_subcortical_for_CLA
Copy_of_normalized_projection_volume_brain_for_CLA
Copy_of_normalized_projection_volume_brain_for_CLA
Copy_of_normalized_projection_volume-brain_subcortical_for_CLA
Copy_of_normalized_projection_volume_brain_for_CLA
Copy_of_normalized_projection_volume_brain_for_CLA
Copy_of_normalized_projection_volume_brain_for_CLA
Copy_of_normalized_projection_volume_brain_for_CLA
Copy_of_normalized_projection_volume_brain_for_CLA
Copy_of_normalized_projection_volume_brain_for_CLA
Copy_of_normalized_projection_volume_brain_for_CLA
Copy_of_normalized_projection_volume_brain_for_CLA
Copy_of_normalized_projection_volume_brain_for_CLA

Copy_of_normalized_projection_volume_brain_for_CLA
Copy_of_normalized_projection_volume_brain_for_CLA
Copy_of_normalized_projection_volume_brain_for_CLA
Copy_of_normalized_projection_volume_brain_for_CLA
Copy_of_normalized_projection_volume_brain_for_CLA
Copy_of_normalized_projection_volume_brain_for_CLA
Copy_of_normalized_projection_volume_brain_for_CLA
Copy_of_normalized_projection_volume_brain_for_CLA
Copy_of_normalized_projection_volume_brain_for_CLA
Copy_of_normalized_projection_volume_brain_for_CLA
Copy_of_normalized_projection_volume_brain_for_CLA
Copy_of_normalized_projection_volume_brain_for_CLA
Copy_of_normalized_projection_volume_brain_for_CLA
Copy_of_normalized_projection_volume_brain_for_CLA
Copy_of_normalized_projection_volume_brain_for_CLA, best_injection_list
Copy_of_normalized_projection_volume-brain_subcortical_for_CLA, best_injection_list
Copy_of_normalized_projection_volume-brain_subcortical_for_CLA
Copy_of_normalized_projection_volume_brain_for_CLA, best_injection_list
Copy_of_normalized_projection_volume_brain_for_CLA
Copy_of_normalized_projection_volume-brain_subcortical_for_CLA
Copy_of_normalized_projection_volume_brain_for_CLA
Copy_of_normalized_projection_volume_brain_for_CLA
Copy_of_normalized_projection_volume_brain_for_CLA
Copy_of_normalized_projection_volume_brain_for_CLA, best_injection_list
Copy_of_normalized_projection_volume_brain_for_CLA, best_injection_list
Copy_of_normalized_projection_volume_brain_for_CLA, best_injection_list
Copy_of_normalized_projection_volume-brain_subcortical_for_CLA
Copy_of_normalized_projection_volume-brain_subcortical_for_CLA, best_injection_list
Copy_of_normalized_projection_volume-brain_subcortical_for_CLA
Copy_of_normalized_projection_volume-brain_subcortical_for_CLA
Copy_of_normalized_projection_volume-brain_subcortical_for_CLA
Copy_of_normalized_projection_volume_brain_for_CLA
Copy_of_normalized_projection_volume_brain_for_CLA
Copy_of_normalized_projection_volume_brain_for_CLA, best_injection_list
Copy_of_normalized_projection_volume_brain_for_CLA
Copy_of_normalized_projection_volume_brain_for_CLA, best_injection_list
Copy_of_normalized_projection_volume_brain_for_CLA
Copy_of_normalized_projection_volume_brain_for_CLA, best_injection_list
Copy_of_normalized_projection_volume_brain_for_CLA, best_injection_list
Copy_of_normalized_projection_volume_brain_for_CLA
Copy_of_normalized_projection_volume_brain_for_CLA
Copy_of_normalized_projection_volume-brain_subcortical_for_CLA, best_injection_list
Copy_of_normalized_projection_volume-brain_subcortical_for_CLA, best_injection_list
Copy_of_normalized_projection_volume-brain_subcortical_for_CLA
Copy_of_normalized_projection_volume_brain_for_CLA
Copy_of_normalized_projection_volume-brain_subcortical_for_CLA, best_injection_list
Copy_of_normalized_projection_volume-brain_subcortical_for_CLA, best_injection_list

Copy_of_normalized_projection_volume_brain_for_CLA, best_injection_list
Copy_of_normalized_projection_volume_brain_for_CLA, best_injection_list
Copy_of_normalized_projection_volume_brain_for_CLA, best_injection_list
Copy_of_normalized_projection_volume_brain_for_CLA
Copy_of_normalized_projection_volume_brain_for_CLA
Copy_of_normalized_projection_volume_brain_for_CLA
Copy_of_normalized_projection_volume_brain_for_CLA
Copy_of_normalized_projection_volume_brain_for_CLA
Copy_of_normalized_projection_volume_brain_for_CLA
Copy_of_normalized_projection_volume_brain_for_CLA
Copy_of_normalized_projection_volume_brain_for_CLA
Copy_of_normalized_projection_volume_brain_for_CLA, best_injection_list
Copy_of_normalized_projection_volume_brain_for_CLA
Copy_of_normalized_projection_volume_brain_for_CLA, best_injection_list
Copy_of_normalized_projection_volume_brain_for_CLA, best_injection_list
Copy_of_normalized_projection_volume_brain_for_CLA
Copy_of_normalized_projection_volume_brain_for_CLA
Copy_of_normalized_projection_volume_brain_for_CLA
Copy_of_normalized_projection_volume_brain_for_CLA, best_injection_list
Copy_of_normalized_projection_volume_brain_for_CLA, best_injection_list
Copy_of_normalized_projection_volume_brain_for_CLA
Copy_of_normalized_projection_volume_brain_for_CLA, best_injection_list
Copy_of_normalized_projection_volume_brain_for_CLA, best_injection_list
Copy_of_normalized_projection_volume_brain_for_CLA
Copy_of_normalized_projection_volume_brain_for_CLA
Copy_of_normalized_projection_volume_brain_for_CLA
Copy_of_normalized_projection_volume_brain_for_CLA
Copy_of_normalized_projection_volume_brain_for_CLA
Copy_of_normalized_projection_volume_brain_for_CLA
Copy_of_normalized_projection_volume_brain_for_CLA
Copy_of_normalized_projection_volume_brain_for_CLA
Copy_of_normalized_projection_volume_brain_for_CLA
Copy_of_normalized_projection_volume_brain_for_CLA
Copy_of_normalized_projection_volume_brain_for_CLA, best_injection_list
Copy_of_normalized_projection_volume_brain_for_CLA
Copy_of_normalized_projection_volume_brain_for_CLA
Copy_of_normalized_projection_volume_brain_for_CLA
Copy_of_normalized_projection_volume_brain_for_CLA
Copy_of_normalized_projection_volume_brain_for_CLA
Copy_of_normalized_projection_volume_brain_for_CLA
Copy_of_normalized_projection_volume_brain_for_CLA
Copy_of_normalized_projection_volume_brain_for_CLA
Copy_of_normalized_projection_volume_brain_for_CLA
Copy_of_normalized_projection_volume_brain_for_CLA
Copy_of_normalized_projection_volume_brain_for_CLA
Copy_of_normalized_projection_volume-brain_subcortical_for_CLA
Copy_of_normalized_projection_volume_brain_for_CLA

Copy_of_normalized_projection_volume_brain_for_CLA
Copy_of_normalized_projection_volume_brain_for_CLA, best_injection_list
Copy_of_normalized_projection_volume_brain_for_CLA
Copy_of_normalized_projection_volume_brain_for_CLA, best_injection_list
Copy_of_normalized_projection_volume_brain_for_CLA
Copy_of_normalized_projection_volume_brain_for_CLA
Copy_of_normalized_projection_volume_brain_for_CLA
Copy_of_normalized_projection_volume_brain_for_CLA
Copy_of_normalized_projection_volume_brain_for_CLA
Copy_of_normalized_projection_volume_brain_for_CLA
Copy_of_normalized_projection_volume_brain_for_CLA, best_injection_list
Copy_of_normalized_projection_volume_brain_for_CLA, best_injection_list
Copy_of_normalized_projection_volume_brain_for_CLA
Copy_of_normalized_projection_volume_brain_for_CLA
Copy_of_normalized_projection_volume_brain_for_CLA
Copy_of_normalized_projection_volume-brain_subcortical_for_CLA
Copy_of_normalized_projection_volume_brain_for_CLA
Copy_of_normalized_projection_volume_brain_for_CLA
Copy_of_normalized_projection_volume_brain_for_CLA
Copy_of_normalized_projection_volume-brain_subcortical_for_CLA, best_injection_list
Copy_of_normalized_projection_volume_brain_for_CLA
Copy_of_normalized_projection_volume-brain_subcortical_for_CLA, best_injection_list
Copy_of_normalized_projection_volume-brain_subcortical_for_CLA
Copy_of_normalized_projection_volume_brain_for_CLA, best_injection_list
Copy_of_normalized_projection_volume_brain_for_CLA
Copy_of_normalized_projection_volume_brain_for_CLA
Copy_of_normalized_projection_volume_brain_for_CLA
Copy_of_normalized_projection_volume-brain_subcortical_for_CLA
Copy_of_normalized_projection_volume_brain_for_CLA
Copy_of_normalized_projection_volume_brain_for_CLA, best_injection_list
Copy_of_normalized_projection_volume_brain_for_CLA
Copy_of_normalized_projection_volume_brain_for_CLA, best_injection_list
Copy_of_normalized_projection_volume_brain_for_CLA
Copy_of_normalized_projection_volume_brain_for_CLA
Copy_of_normalized_projection_volume_brain_for_CLA
Copy_of_normalized_projection_volume_brain_for_CLA, best_injection_list
Copy_of_normalized_projection_volume_brain_for_CLA
Copy_of_normalized_projection_volume_brain_for_CLA, best_injection_list
Copy_of_normalized_projection_volume_brain_for_CLA
Copy_of_normalized_projection_volume_brain_for_CLA
Copy_of_normalized_projection_volume_brain_for_CLA
Copy_of_normalized_projection_volume_brain_for_CLA
Copy_of_normalized_projection_volume_brain_for_CLA, best_injection_list
Copy_of_normalized_projection_volume_brain_for_CLA
Copy_of_normalized_projection_volume_brain_for_CLA
Copy_of_normalized_projection_volume_brain_for_CLA
Copy_of_normalized_projection_volume_brain_for_CLA

Copy_of_normalized_projection_volume_brain_for_CLA
Copy_of_normalized_projection_volume_brain_for_CLA, best_injection_list
Copy_of_normalized_projection_volume_brain_for_CLA, best_injection_list
Copy_of_normalized_projection_volume_brain_for_CLA
Copy_of_normalized_projection_volume_brain_for_CLA
Copy_of_normalized_projection_volume_brain_for_CLA
Copy_of_normalized_projection_volume_brain_for_CLA
Copy_of_normalized_projection_volume_brain_for_CLA, best_injection_list
Copy_of_normalized_projection_volume_brain_for_CLA
Copy_of_normalized_projection_volume_brain_for_CLA, best_injection_list
Copy_of_normalized_projection_volume-brain_subcortical_for_CLA, best_injection_list
Copy_of_normalized_projection_volume_brain_for_CLA
Copy_of_normalized_projection_volume_brain_for_CLA, best_injection_list
Copy_of_normalized_projection_volume_brain_for_CLA
Copy_of_normalized_projection_volume_brain_for_CLA
Copy_of_normalized_projection_volume_brain_for_CLA
Copy_of_normalized_projection_volume_brain_for_CLA
Copy_of_normalized_projection_volume_brain_for_CLA
Copy_of_normalized_projection_volume_brain_for_CLA
Copy_of_normalized_projection_volume_brain_for_CLA
Copy_of_normalized_projection_volume_brain_for_CLA
Copy_of_normalized_projection_volume_brain_for_CLA
Copy_of_normalized_projection_volume_brain_for_CLA
Copy_of_normalized_projection_volume_brain_for_CLA
Copy_of_normalized_projection_volume_brain_for_CLA
Copy_of_normalized_projection_volume_brain_for_CLA
Copy_of_normalized_projection_volume_brain_for_CLA
Copy_of_normalized_projection_volume_brain_for_CLA
Copy_of_normalized_projection_volume_brain_for_CLA
Copy_of_normalized_projection_volume_brain_for_CLA
Copy_of_normalized_projection_volume_brain_for_CLA
Copy_of_normalized_projection_volume_brain_for_CLA
Copy_of_normalized_projection_volume_brain_for_CLA
Copy_of_normalized_projection_volume-brain_subcortical_for_CLA
Copy_of_normalized_projection_volume_brain_for_CLA, best_injection_list
Copy_of_normalized_projection_volume_brain_for_CLA, best_injection_list
Copy_of_normalized_projection_volume-brain_subcortical_for_CLA, best_injection_list
Copy_of_normalized_projection_volume-brain_subcortical_for_CLA, best_injection_list
Copy_of_normalized_projection_volume-brain_subcortical_for_CLA
Copy_of_normalized_projection_volume-brain_subcortical_for_CLA
Copy_of_normalized_projection_volume-brain_subcortical_for_CLA
Copy_of_normalized_projection_volume_brain_for_CLA
Copy_of_normalized_projection_volume_brain_for_CLA
Copy_of_normalized_projection_volume_brain_for_CLA
Copy_of_normalized_projection_volume_brain_for_CLA
Copy_of_normalized_projection_volume_brain_for_CLA
Copy_of_normalized_projection_volume_brain_for_CLA

Copy_of_normalized_projection_volume_brain_for_CLA
Copy_of_normalized_projection_volume_brain_for_CLA
Copy_of_normalized_projection_volume_brain_for_CLA, best_injection_list
Copy_of_normalized_projection_volume_brain_for_CLA
Copy_of_normalized_projection_volume_brain_for_CLA
Copy_of_normalized_projection_volume-brain_subcortical_for_CLA
Copy_of_normalized_projection_volume_brain_for_CLA
Copy_of_normalized_projection_volume_brain_for_CLA
Copy_of_normalized_projection_volume_brain_for_CLA
Copy_of_normalized_projection_volume_brain_for_CLA
Copy_of_normalized_projection_volume_brain_for_CLA
Copy_of_normalized_projection_volume_brain_for_CLA
Copy_of_normalized_projection_volume-brain_subcortical_for_CLA
Copy_of_normalized_projection_volume_brain_for_CLA
Copy_of_normalized_projection_volume_brain_for_CLA
Copy_of_normalized_projection_volume_brain_for_CLA
Copy_of_normalized_projection_volume_brain_for_CLA
Copy_of_normalized_projection_volume_brain_for_CLA, best_injection_list
Copy_of_normalized_projection_volume_brain_for_CLA, best_injection_list
Copy_of_normalized_projection_volume_brain_for_CLA
Copy_of_normalized_projection_volume_brain_for_CLA, best_injection_list
Copy_of_normalized_projection_volume_brain_for_CLA
Copy_of_normalized_projection_volume_brain_for_CLA
Copy_of_normalized_projection_volume_brain_for_CLA
Copy_of_normalized_projection_volume_brain_for_CLA
Copy_of_normalized_projection_volume_brain_for_CLA
Copy_of_normalized_projection_volume_brain_for_CLA, best_injection_list
Copy_of_normalized_projection_volume_brain_for_CLA
Copy_of_normalized_projection_volume_brain_for_CLA, best_injection_list
Copy_of_normalized_projection_volume_brain_for_CLA
Copy_of_normalized_projection_volume_brain_for_CLA
Copy_of_normalized_projection_volume_brain_for_CLA
Copy_of_normalized_projection_volume_brain_for_CLA
Copy_of_normalized_projection_volume_brain_for_CLA
Copy_of_normalized_projection_volume-brain_subcortical_for_CLA
Copy_of_normalized_projection_volume-brain_subcortical_for_CLA, best_injection_list
Copy_of_normalized_projection_volume_brain_for_CLA
Copy_of_normalized_projection_volume_brain_for_CLA
Copy_of_normalized_projection_volume-brain_subcortical_for_CLA
Copy_of_normalized_projection_volume_brain_for_CLA
Copy_of_normalized_projection_volume_brain_for_CLA
Copy_of_normalized_projection_volume_brain_for_CLA
Copy_of_normalized_projection_volume_brain_for_CLA, best_injection_list
Copy_of_normalized_projection_volume_brain_for_CLA, best_injection_list
Copy_of_normalized_projection_volume_brain_for_CLA
Copy_of_normalized_projection_volume_brain_for_CLA
Copy_of_normalized_projection_volume_brain_for_CLA

Copy_of_normalized_projection_volume_brain_for_CLA
Copy_of_normalized_projection_volume_brain_for_CLA
Copy_of_normalized_projection_volume_brain_for_CLA
Copy_of_normalized_projection_volume_brain_for_CLA
Copy_of_normalized_projection_volume_brain_for_CLA
Copy_of_normalized_projection_volume_brain_for_CLA, best_injection_list
Copy_of_normalized_projection_volume_brain_for_CLA, best_injection_list
Copy_of_normalized_projection_volume_brain_for_CLA, best_injection_list
Copy_of_normalized_projection_volume_brain_for_CLA
Copy_of_normalized_projection_volume_brain_for_CLA, best_injection_list
Copy_of_normalized_projection_volume_brain_for_CLA, best_injection_list
Copy_of_normalized_projection_volume_brain_for_CLA
Copy_of_normalized_projection_volume-brain_subcortical_for_CLA
Copy_of_normalized_projection_volume_brain_for_CLA
Copy_of_normalized_projection_volume_brain_for_CLA, best_injection_list
Copy_of_normalized_projection_volume_brain_for_CLA
Copy_of_normalized_projection_volume_brain_for_CLA
Copy_of_normalized_projection_volume_brain_for_CLA, best_injection_list
Copy_of_normalized_projection_volume_brain_for_CLA, best_injection_list
Copy_of_normalized_projection_volume_brain_for_CLA
Copy_of_normalized_projection_volume_brain_for_CLA
Copy_of_normalized_projection_volume_brain_for_CLA, best_injection_list
Copy_of_normalized_projection_volume_brain_for_CLA
Copy_of_normalized_projection_volume_brain_for_CLA
Copy_of_normalized_projection_volume_brain_for_CLA, best_injection_list
Copy_of_normalized_projection_volume_brain_for_CLA
Copy_of_normalized_projection_volume_brain_for_CLA
Copy_of_normalized_projection_volume_brain_for_CLA
Copy_of_normalized_projection_volume_brain_for_CLA
Copy_of_normalized_projection_volume_brain_for_CLA
Copy_of_normalized_projection_volume_brain_for_CLA
Copy_of_normalized_projection_volume_brain_for_CLA
Copy_of_normalized_projection_volume_brain_for_CLA, best_injection_list
Copy_of_normalized_projection_volume_brain_for_CLA
Copy_of_normalized_projection_volume_brain_for_CLA
Copy_of_normalized_projection_volume_brain_for_CLA, best_injection_list
Copy_of_normalized_projection_volume-brain_subcortical_for_CLA
Copy_of_normalized_projection_volume_brain_for_CLA
Copy_of_normalized_projection_volume_brain_for_CLA, best_injection_list
Copy_of_normalized_projection_volume_brain_for_CLA, best_injection_list
Copy_of_normalized_projection_volume_brain_for_CLA
Copy_of_normalized_projection_volume_brain_for_CLA, best_injection_list
Copy_of_normalized_projection_volume_brain_for_CLA
Copy_of_normalized_projection_volume_brain_for_CLA
Copy_of_normalized_projection_volume_brain_for_CLA
Copy_of_normalized_projection_volume_brain_for_CLA, best_injection_list
Copy_of_normalized_projection_volume_brain_for_CLA

Copy_of_normalized_projection_volume_brain_for_CLA
Copy_of_normalized_projection_volume_brain_for_CLA, best_injection_list
Copy_of_normalized_projection_volume_brain_for_CLA
Copy_of_normalized_projection_volume_brain_for_CLA, best_injection_list
Copy_of_normalized_projection_volume_brain_for_CLA, best_injection_list
Copy_of_normalized_projection_volume_brain_for_CLA, best_injection_list
Copy_of_normalized_projection_volume_brain_for_CLA
Copy_of_normalized_projection_volume_brain_for_CLA
Copy_of_normalized_projection_volume_brain_for_CLA, best_injection_list
Copy_of_normalized_projection_volume-brain_subcortical_for_CLA
Copy_of_normalized_projection_volume_brain_for_CLA, best_injection_list
Copy_of_normalized_projection_volume_brain_for_CLA
Copy_of_normalized_projection_volume_brain_for_CLA
Copy_of_normalized_projection_volume_brain_for_CLA
Copy_of_normalized_projection_volume_brain_for_CLA
Copy_of_normalized_projection_volume_brain_for_CLA
Copy_of_normalized_projection_volume-brain_subcortical_for_CLA
Copy_of_normalized_projection_volume_brain_for_CLA
Copy_of_normalized_projection_volume_brain_for_CLA
Copy_of_normalized_projection_volume_brain_for_CLA
Copy_of_normalized_projection_volume_brain_for_CLA
Copy_of_normalized_projection_volume_brain_for_CLA, best_injection_list
Copy_of_normalized_projection_volume-brain_subcortical_for_CLA
Copy_of_normalized_projection_volume_brain_for_CLA
Copy_of_normalized_projection_volume_brain_for_CLA
Copy_of_normalized_projection_volume_brain_for_CLA
Copy_of_normalized_projection_volume_brain_for_CLA
Copy_of_normalized_projection_volume_brain_for_CLA
Copy_of_normalized_projection_volume_brain_for_CLA, best_injection_list
Copy_of_normalized_projection_volume_brain_for_CLA
Copy_of_normalized_projection_volume_brain_for_CLA
Copy_of_normalized_projection_volume_brain_for_CLA, best_injection_list
Copy_of_normalized_projection_volume_brain_for_CLA
Copy_of_normalized_projection_volume_brain_for_CLA
cla_projection_volume_20191023, restricted_injections
cla_projection_volume_20191023
cla_projection_volume_20191023
cla_projection_volume_20191023
cla_projection_volume_20191023
cla_projection_volume_20191023
cla_projection_volume_20191023
cla_projection_volume_20191023, restricted_injections
cla_projection_volume_20191023, restricted_injections
cla_projection_volume_20191023, restricted_injections
cla_projection_volume_20191023, restricted_injections
cla_projection_volume_20191023
cla_projection_volume_20191023

cla_projection_volume_20191023

| included in Tabl | key resource name | key resource source | key resource stock number |
|---|---|---|---|
| | Mouse: C57Bl/6J | The Jackson Laboratory | JAX: 000664 |
| | Mouse: C57Bl/6J | The Jackson Laboratory | JAX: 000664 |
| | Mouse: C57Bl/6J | The Jackson Laboratory | JAX: 000664 |
| | Mouse: C57Bl/6J | The Jackson Laboratory | JAX: 000664 |
| | Mouse: C57Bl/6J | The Jackson Laboratory | JAX: 000664 |
| | Mouse: C57Bl/6J | The Jackson Laboratory | JAX: 000664 |
| | Mouse: C57Bl/6J | The Jackson Laboratory | JAX: 000664 |
| | Mouse: C57Bl/6J | The Jackson Laboratory | JAX: 000664 |
| | Mouse: C57Bl/6J | The Jackson Laboratory | JAX: 000664 |
| | Mouse: C57Bl/6J | The Jackson Laboratory | JAX: 000664 |
| | Mouse: C57Bl/6J | The Jackson Laboratory | JAX: 000664 |
| | Mouse: C57Bl/6J | The Jackson Laboratory | JAX: 000664 |
| | Mouse: C57Bl/6J | The Jackson Laboratory | JAX: 000664 |
| | Mouse: C57Bl/6J | The Jackson Laboratory | JAX: 000664 |
| | Mouse: C57Bl/6J | The Jackson Laboratory | JAX: 000664 |
| | Mouse: C57Bl/6J | The Jackson Laboratory | JAX: 000664 |
| | Mouse: C57Bl/6J | The Jackson Laboratory | JAX: 000664 |
| | Mouse: C57Bl/6J | The Jackson Laboratory | JAX: 000664 |
| Y | Mouse: C57Bl/6J | The Jackson Laboratory | JAX: 000664 |
| | Mouse: C57Bl/6J | The Jackson Laboratory | JAX: 000664 |
| | Mouse: C57Bl/6J | The Jackson Laboratory | JAX: 000664 |
| | Mouse: C57Bl/6J | The Jackson Laboratory | JAX: 000664 |
| | Mouse: C57Bl/6J | The Jackson Laboratory | JAX: 000664 |
| | Mouse: C57Bl/6J | The Jackson Laboratory | JAX: 000664 |
| | Mouse: C57Bl/6J | The Jackson Laboratory | JAX: 000664 |
| | Mouse: C57Bl/6J | The Jackson Laboratory | JAX: 000664 |
| | Mouse: C57Bl/6J | The Jackson Laboratory | JAX: 000664 |
| Y | Mouse: C57BL/6J | The Jackson Laboratory | JAX: 000664 |
| Y | Mouse: C57BL/6J | The Jackson Laboratory | JAX: 000664 |
| | Mouse: C57Bl/6J | The Jackson Laboratory | JAX: 000664 |
| | Mouse: C57Bl/6J | The Jackson Laboratory | JAX: 000664 |
| | Mouse: C57Bl/6J | The Jackson Laboratory | JAX: 000664 |
| | Mouse: C57Bl/6J | The Jackson Laboratory | JAX: 000664 |
| | Mouse: C57Bl/6J | The Jackson Laboratory | JAX: 000664 |
| | Mouse: C57Bl/6J | The Jackson Laboratory | JAX: 000664 |
| | Mouse: C57Bl/6J | The Jackson Laboratory | JAX: 000664 |
| | Mouse: C57Bl/6J | The Jackson Laboratory | JAX: 000664 |
| | Mouse: C57Bl/6J | The Jackson Laboratory | JAX: 000664 |
| | Mouse: Oxt-IRES-Cre | Bradford Lowell | N/A |
| | Mouse: Oxt-IRES-Cre | Bradford Lowell | N/A |
| | Mouse: C57Bl/6J | The Jackson Laboratory | JAX: 000664 |
| | Mouse: C57Bl/6J | The Jackson Laboratory | JAX: 000664 |
| | Mouse: C57Bl/6J | The Jackson Laboratory | JAX: 000664 |
| | Mouse: B6(Cg)-Etv1tm1.1 | The Jackson Laboratory | JAX: 013048 |
| | Mouse: C57Bl/6J | The Jackson Laboratory | JAX: 000664 |
| | Mouse: B6.129-Leprtm2( | The Jackson Laboratory | JAX: 008320 |

| | | | |
|---|---|---|---|
| Y | Mouse: C57Bl/6J | The Jackson Laboratory | JAX: 000664 |
| | Mouse: C57Bl/6J | The Jackson Laboratory | JAX: 000664 |
| | Mouse: B6.129-Leprtm2( | The Jackson Laboratory | JAX: 008320 |
| | Mouse: C57Bl/6J | The Jackson Laboratory | JAX: 000664 |
| | Mouse: STOCK Tg(Drd1-cr | MMRRC | MMRRC: 017264 |
| | Mouse: C57Bl/6J | The Jackson Laboratory | JAX: 000664 |
| | Mouse: C57Bl/6J | The Jackson Laboratory | JAX: 000664 |
| | Mouse: C57Bl/6J | The Jackson Laboratory | JAX: 000664 |
| | Mouse: C57Bl/6J | The Jackson Laboratory | JAX: 000664 |
| | Mouse: C57Bl/6J | The Jackson Laboratory | JAX: 000664 |
| | Mouse: STOCK Tg(Pmch-c | The Jackson Laboratory | JAX: 014099 |
| | Mouse: C57Bl/6J | The Jackson Laboratory | JAX: 000664 |
| | Mouse: C57Bl/6J | The Jackson Laboratory | JAX: 000664 |
| | Mouse: STOCK Tg(Pmch-c | The Jackson Laboratory | JAX: 014099 |
| | Mouse: C57Bl/6J | The Jackson Laboratory | JAX: 000664 |
| | Mouse: STOCK Tg(Gal-cre | MMRRC | MMRRC: 031060 |
| | Mouse: C57Bl/6J | The Jackson Laboratory | JAX: 000664 |
| | Mouse: STOCK Tg(Gal-cre | MMRRC | MMRRC: 031060 |
| | Mouse: C57Bl/6J | The Jackson Laboratory | JAX: 000664 |
| | Mouse: C57BL/6-Tg(Grik4 | The Jackson Laboratory | JAX: 006474 |
| | Mouse: B6.FVB-Tg(Pomc- | The Jackson Laboratory | JAX: 010714 |
| | Mouse: C57Bl/6J | The Jackson Laboratory | JAX: 000664 |
| | Mouse: C57Bl/6J | The Jackson Laboratory | JAX: 000664 |
| | Mouse: C57Bl/6J | The Jackson Laboratory | JAX: 000664 |
| | Mouse: C57Bl/6J | The Jackson Laboratory | JAX: 000664 |
| | Mouse: B6;C3-Tg(Scnn1a- | The Jackson Laboratory | JAX: 009613 |
| | Mouse: C57BL/6-Tg(Grik4 | The Jackson Laboratory | JAX: 006474 |
| | Mouse: B6;C3-Tg(Scnn1a- | The Jackson Laboratory | JAX: 009613 |
| | Mouse: C57Bl/6J | The Jackson Laboratory | JAX: 000664 |
| | Mouse: C57Bl/6J | The Jackson Laboratory | JAX: 000664 |
| | Mouse: C57Bl/6J | The Jackson Laboratory | JAX: 000664 |
| | Mouse: C57Bl/6J | The Jackson Laboratory | JAX: 000664 |
| | Mouse: C57BL/6-Tg(Grik4 | The Jackson Laboratory | JAX: 006474 |
| | Mouse: C57Bl/6J | The Jackson Laboratory | JAX: 000664 |
| | Mouse: C57Bl/6J | The Jackson Laboratory | JAX: 000664 |
| | Mouse: C57BL/6-Tg(Grik4 | The Jackson Laboratory | JAX: 006474 |
| | Mouse: STOCK Tg(Slc6a4- | MMRRC | MMRRC: 030071 |
| | Mouse: C57Bl/6J | The Jackson Laboratory | JAX: 000664 |
| | Mouse: STOCK Tg(Slc6a4- | MMRRC | MMRRC: 030071 |
| | Mouse: C57Bl/6J | The Jackson Laboratory | JAX: 000664 |
| | Mouse: FVB-Tg(Nr5a1-cre | The Jackson Laboratory | JAX: 006364 |
| | Mouse: C57Bl/6J | The Jackson Laboratory | JAX: 000664 |
| | Mouse: C57Bl/6J | The Jackson Laboratory | JAX: 000664 |
| | Mouse: C57Bl/6J | The Jackson Laboratory | JAX: 000664 |
| | Mouse: C57Bl/6J | The Jackson Laboratory | JAX: 000664 |
| | Mouse: C57Bl/6J | The Jackson Laboratory | JAX: 000664 |
| | Mouse: B6(Cg)-Etv1tm1.1 | The Jackson Laboratory | JAX: 013048 |

| | | | |
|---|---|---|---|
| | Mouse: C57Bl/6J | The Jackson Laboratory | JAX: 000664 |
| | Mouse: C57Bl/6J | The Jackson Laboratory | JAX: 000664 |
| | Mouse: C57Bl/6J | The Jackson Laboratory | JAX: 000664 |
| | Mouse: C57Bl/6J | The Jackson Laboratory | JAX: 000664 |
| | Mouse: C57Bl/6J | The Jackson Laboratory | JAX: 000664 |
| | Mouse: C57Bl/6J | The Jackson Laboratory | JAX: 000664 |
| | Mouse: B6(Cg)-Etv1tm1.1 | The Jackson Laboratory | JAX: 013048 |
| | Mouse: FVB-Tg(Nr5a1-cre | The Jackson Laboratory | JAX: 006364 |
| | Mouse: C57Bl/6J | The Jackson Laboratory | JAX: 000664 |
| | Mouse: C57Bl/6J | The Jackson Laboratory | JAX: 000664 |
| | Mouse: C57Bl/6J | The Jackson Laboratory | JAX: 000664 |
| | Mouse: FVB-Tg(Nr5a1-cre | The Jackson Laboratory | JAX: 006364 |
| | Mouse: C57Bl/6J | The Jackson Laboratory | JAX: 000664 |
| | Mouse: C57Bl/6J | The Jackson Laboratory | JAX: 000664 |
| | Mouse: STOCK Tg(Gal-cre | MMRRC | MMRRC: 031060 |
| | Mouse: C57Bl/6J | The Jackson Laboratory | JAX: 000664 |
| | Mouse: C57Bl/6J | The Jackson Laboratory | JAX: 000664 |
| | Mouse: C57Bl/6J | The Jackson Laboratory | JAX: 000664 |
| | Mouse: C57Bl/6J | The Jackson Laboratory | JAX: 000664 |
| | Mouse: C57Bl/6J | The Jackson Laboratory | JAX: 000664 |
| | Mouse: C57Bl/6J | The Jackson Laboratory | JAX: 000664 |
| | Mouse: C57Bl/6J | The Jackson Laboratory | JAX: 000664 |
| | Mouse: C57Bl/6J | The Jackson Laboratory | JAX: 000664 |
| | Mouse: C57Bl/6J | The Jackson Laboratory | JAX: 000664 |
| Y | Mouse: C57Bl/6J | The Jackson Laboratory | JAX: 000664 |
| | Mouse: B6.Cg-Erbb4tm1.: | The Jackson Laboratory | JAX: 012360 |
| | Mouse: C57Bl/6J | The Jackson Laboratory | JAX: 000664 |
| | Mouse: C57Bl/6J | The Jackson Laboratory | JAX: 000664 |
| | Mouse: C57Bl/6J | The Jackson Laboratory | JAX: 000664 |
| | Mouse: C57Bl/6J | The Jackson Laboratory | JAX: 000664 |
| | Mouse: C57Bl/6J | The Jackson Laboratory | JAX: 000664 |
| | Mouse: B6.FVB(Cg)-Tg(Nt: | MMRRC | MMRRC: 030648 |
| | Mouse: C57Bl/6J | The Jackson Laboratory | JAX: 000664 |
| | Mouse: C57Bl/6J | The Jackson Laboratory | JAX: 000664 |
| | Mouse: C57Bl/6J | The Jackson Laboratory | JAX: 000664 |
| | Mouse: B6.FVB(Cg)-Tg(Nt: | MMRRC | MMRRC: 030648 |
| | Mouse: C57Bl/6J | The Jackson Laboratory | JAX: 000664 |
| | Mouse: C57Bl/6J | The Jackson Laboratory | JAX: 000664 |
| | Mouse: C57Bl/6J | The Jackson Laboratory | JAX: 000664 |
| | Mouse: C57Bl/6J | The Jackson Laboratory | JAX: 000664 |
| | Mouse: STOCK Tg(Rbp4-cı | MMRRC | MMRRC: 031125 |
| | Mouse: C57Bl/6J | The Jackson Laboratory | JAX: 000664 |
| | Mouse: C57Bl/6J | The Jackson Laboratory | JAX: 000664 |
| | Mouse: STOCK Tg(Rbp4-cı | MMRRC | MMRRC: 031125 |
| | Mouse: STOCK Tg(Rbp4-cı | MMRRC | MMRRC: 031125 |
| | Mouse: C57Bl/6J | The Jackson Laboratory | JAX: 000664 |
| | Mouse: C57Bl/6J | The Jackson Laboratory | JAX: 000664 |

| | | | |
|---|---|---|---|
| | Mouse: C57Bl/6J | The Jackson Laboratory | JAX: 000664 |
| | Mouse: C57Bl/6J | The Jackson Laboratory | JAX: 000664 |
| | Mouse: C57Bl/6J | The Jackson Laboratory | JAX: 000664 |
| | Mouse: C57Bl/6J | The Jackson Laboratory | JAX: 000664 |
| | Mouse: C57Bl/6J | The Jackson Laboratory | JAX: 000664 |
| | Mouse: C57Bl/6J | The Jackson Laboratory | JAX: 000664 |
| | Mouse: STOCK Tg(Syt6-cre | MMRRC | MMRRC: 032012 |
| | Mouse: C57Bl/6J | The Jackson Laboratory | JAX: 000664 |
| | Mouse: C57Bl/6J | The Jackson Laboratory | JAX: 000664 |
| | Mouse: STOCK Tg(Syt6-cre | MMRRC | MMRRC: 032012 |
| | Mouse: C57Bl/6J | The Jackson Laboratory | JAX: 000664 |
| | Mouse: STOCK Tg(Syt6-cre | MMRRC | MMRRC: 032012 |
| | Mouse: C57Bl/6J | The Jackson Laboratory | JAX: 000664 |
| | Mouse: C57Bl/6J | The Jackson Laboratory | JAX: 000664 |
| | Mouse: STOCK Tg(Syt6-cre | MMRRC | MMRRC: 032012 |
| | Mouse: C57Bl/6J | The Jackson Laboratory | JAX: 000664 |
| | Mouse: C57Bl/6J | The Jackson Laboratory | JAX: 000664 |
| | Mouse: C57Bl/6J | The Jackson Laboratory | JAX: 000664 |
| | Mouse: STOCK Tg(Syt6-cre | MMRRC | MMRRC: 032012 |
| | Mouse: C57Bl/6J | The Jackson Laboratory | JAX: 000664 |
| | Mouse: C57Bl/6J | The Jackson Laboratory | JAX: 000664 |
| | Mouse: C57Bl/6J | The Jackson Laboratory | JAX: 000664 |
| Y | Mouse: STOCK Tg(Rbp4-cr | MMRRC | MMRRC: 031125 |
| | Mouse: C57Bl/6J | The Jackson Laboratory | JAX: 000664 |
| | Mouse: C57Bl/6J | The Jackson Laboratory | JAX: 000664 |
| Y | Mouse: C57Bl/6J | The Jackson Laboratory | JAX: 000664 |
| Y | Mouse: STOCK Tg(Rbp4-cr | MMRRC | MMRRC: 031125 |
| | Mouse: C57Bl/6J | The Jackson Laboratory | JAX: 000664 |
| | Mouse: C57Bl/6J | The Jackson Laboratory | JAX: 000664 |
| | Mouse: STOCK Tg(Syt6-cre | MMRRC | MMRRC: 032012 |
| | Mouse: C57Bl/6J | The Jackson Laboratory | JAX: 000664 |
| | Mouse: C57Bl/6J | The Jackson Laboratory | JAX: 000664 |
| | Mouse: C57Bl/6J | The Jackson Laboratory | JAX: 000664 |
| | Mouse: C57Bl/6J | The Jackson Laboratory | JAX: 000664 |
| | Mouse: STOCK Tg(Syt6-cre | MMRRC | MMRRC: 032012 |
| | Mouse: C57Bl/6J | The Jackson Laboratory | JAX: 000664 |
| | Mouse: C57Bl/6J | The Jackson Laboratory | JAX: 000664 |
| | Mouse: C57Bl/6J | The Jackson Laboratory | JAX: 000664 |
| | Mouse: C57Bl/6J | The Jackson Laboratory | JAX: 000664 |
| | Mouse: C57Bl/6J | The Jackson Laboratory | JAX: 000664 |
| | Mouse: C57Bl/6J | The Jackson Laboratory | JAX: 000664 |
| | Mouse: C57Bl/6J | The Jackson Laboratory | JAX: 000664 |
| | Mouse: C57Bl/6J | The Jackson Laboratory | JAX: 000664 |
| | Mouse: B6;129S6-Chattm | The Jackson Laboratory | JAX: 006410 |
| | Mouse: B6;129S6-Chattm | The Jackson Laboratory | JAX: 006410 |
| | Mouse: C57Bl/6J | The Jackson Laboratory | JAX: 000664 |
| | Mouse: C57Bl/6J | The Jackson Laboratory | JAX: 000664 |

| | | | |
|---|---|---|---|
| | Mouse: STOCK Tg(Kcnc2-( | The Jackson Laboratory | JAX: 008582 |
| | Mouse: STOCK Tg(Kcnc2-( | The Jackson Laboratory | JAX: 008582 |
| | Mouse: STOCK Tg(Slc6a5- | MMRRC | MMRRC: 030730 |
| | Mouse: C57Bl/6J | The Jackson Laboratory | JAX: 000664 |
| | Mouse: C57Bl/6J | The Jackson Laboratory | JAX: 000664 |
| | Mouse: C57Bl/6J | The Jackson Laboratory | JAX: 000664 |
| | Mouse: C57Bl/6J | The Jackson Laboratory | JAX: 000664 |
| | Mouse: B6;129P2-Pvalbtr | The Jackson Laboratory | JAX: 008069 |
| | Mouse: C57Bl/6J | The Jackson Laboratory | JAX: 000664 |
| | Mouse: B6;129P2-Pvalbtr | The Jackson Laboratory | JAX: 008069 |
| | Mouse: C57Bl/6J | The Jackson Laboratory | JAX: 000664 |
| Y | Mouse: C57Bl/6J | The Jackson Laboratory | JAX: 000664 |
| | Mouse: B6.FVB(Cg)-Tg(Dr | MMRRC | MMRRC: 032108 |
| | Mouse: C57Bl/6J | The Jackson Laboratory | JAX: 000664 |
| | Mouse: B6.FVB(Cg)-Tg(Dr | MMRRC | MMRRC: 032108 |
| | Mouse: C57Bl/6J | The Jackson Laboratory | JAX: 000664 |
| | Mouse: C57Bl/6J | The Jackson Laboratory | JAX: 000664 |
| | Mouse: C57Bl/6J | The Jackson Laboratory | JAX: 000664 |
| | Mouse: C57Bl/6J | The Jackson Laboratory | JAX: 000664 |
| | Mouse: C57Bl/6J | The Jackson Laboratory | JAX: 000664 |
| | Mouse: C57Bl/6J | The Jackson Laboratory | JAX: 000664 |
| | Mouse: STOCK Tg(Sim1-cr | MMRRC | MMRRC: 031742 |
| | Mouse: C57Bl/6J | The Jackson Laboratory | JAX: 000664 |
| | Mouse: C57Bl/6J | The Jackson Laboratory | JAX: 000664 |
| | Mouse: C57Bl/6J | The Jackson Laboratory | JAX: 000664 |
| | Mouse: C57Bl/6J | The Jackson Laboratory | JAX: 000664 |
| | Mouse: C57Bl/6J | The Jackson Laboratory | JAX: 000664 |
| | Mouse: STOCK Tg(Ucn3-c | MMRRC | MMRRC: 032078 |
| | Mouse: C57Bl/6J | The Jackson Laboratory | JAX: 000664 |
| | Mouse: STOCK Tg(Syt17-c | MMRRC | MMRRC: 034355 |
| | Mouse: C57Bl/6J | The Jackson Laboratory | JAX: 000664 |
| | Mouse: C57Bl/6J | The Jackson Laboratory | JAX: 000664 |
| | Mouse: STOCK Tg(Syt17-c | MMRRC | MMRRC: 034355 |
| | Mouse: C57Bl/6J | The Jackson Laboratory | JAX: 000664 |
| | Mouse: C57Bl/6J | The Jackson Laboratory | JAX: 000664 |
| | Mouse: C57Bl/6J | The Jackson Laboratory | JAX: 000664 |
| | Mouse: C57Bl/6J | The Jackson Laboratory | JAX: 000664 |
| | Mouse: C57Bl/6J | The Jackson Laboratory | JAX: 000664 |
| | Mouse: C57Bl/6J | The Jackson Laboratory | JAX: 000664 |
| | Mouse: C57Bl/6J | The Jackson Laboratory | JAX: 000664 |
| | Mouse: STOCK Tg(Syt17-c | MMRRC | MMRRC: 034355 |
| | Mouse: C57Bl/6J | The Jackson Laboratory | JAX: 000664 |
| | Mouse: B6.Cg-Erbb4tm1.: | The Jackson Laboratory | JAX: 012360 |
| Y | Mouse: C57Bl/6J | The Jackson Laboratory | JAX: 000664 |
| | Mouse: C57Bl/6J | The Jackson Laboratory | JAX: 000664 |
| | Mouse: B6.Cg-Erbb4tm1.: | The Jackson Laboratory | JAX: 012360 |
| | Mouse: C57Bl/6J | The Jackson Laboratory | JAX: 000664 |

| | | | |
|---|---|---|---|
| | Mouse: C57Bl/6J | The Jackson Laboratory | JAX: 000664 |
| | Mouse: B6.Cg-Erbb4tm1.: | The Jackson Laboratory | JAX: 012360 |
| | Mouse: C57Bl/6J | The Jackson Laboratory | JAX: 000664 |
| | Mouse: C57Bl/6J | The Jackson Laboratory | JAX: 000664 |
| | Mouse: STOCK Tg(Slc6a5- | MMRRC | MMRRC: 030730 |
| | Mouse: B6.Cg-Tg(Slc6a4-c | MMRRC | MMRRC: 031028 |
| | Mouse: C57Bl/6J | The Jackson Laboratory | JAX: 000664 |
| | Mouse: C57Bl/6J | The Jackson Laboratory | JAX: 000664 |
| | Mouse: C57Bl/6J | The Jackson Laboratory | JAX: 000664 |
| Y | Mouse: C57Bl/6J | The Jackson Laboratory | JAX: 000664 |
| | Mouse: C57Bl/6J | The Jackson Laboratory | JAX: 000664 |
| | Mouse: B6.Cg-Tg(Slc6a4-c | MMRRC | MMRRC: 031028 |
| | Mouse: C57Bl/6J | The Jackson Laboratory | JAX: 000664 |
| | Mouse: C57Bl/6J | The Jackson Laboratory | JAX: 000664 |
| | Mouse: C57Bl/6J | The Jackson Laboratory | JAX: 000664 |
| | Mouse: C57BL/6J | The Jackson Laboratory | JAX: 000664 |
| | Mouse: B6.Cg-Tg(Slc6a4-c | MMRRC | MMRRC: 031028 |
| | Mouse: STOCK Tg(Pmch-c | The Jackson Laboratory | JAX: 014099 |
| | Mouse: C57Bl/6J | The Jackson Laboratory | JAX: 000664 |
| | Mouse: C57Bl/6J | The Jackson Laboratory | JAX: 000664 |
| | Mouse: C57Bl/6J | The Jackson Laboratory | JAX: 000664 |
| Y | Mouse: C57Bl/6J | The Jackson Laboratory | JAX: 000664 |
| | Mouse: Pnmt-Cre | Steven Ebert | N/A |
| | Mouse: C57Bl/6J | The Jackson Laboratory | JAX: 000664 |
| | Mouse: C57Bl/6J | The Jackson Laboratory | JAX: 000664 |
| | Mouse: C57Bl/6J | The Jackson Laboratory | JAX: 000664 |
| | Mouse: C57Bl/6J | The Jackson Laboratory | JAX: 000664 |
| | Mouse: C57Bl/6J | The Jackson Laboratory | JAX: 000664 |
| | Mouse: C57Bl/6J | The Jackson Laboratory | JAX: 000664 |
| | Mouse: C57Bl/6J | The Jackson Laboratory | JAX: 000664 |
| | Mouse: C57Bl/6J | The Jackson Laboratory | JAX: 000664 |
| | Mouse: C57Bl/6J | The Jackson Laboratory | JAX: 000664 |
| | Mouse: STOCK Agrptm1(c | The Jackson Laboratory | JAX: 012899 |
| | Mouse: C57Bl/6J | The Jackson Laboratory | JAX: 000664 |
| | Mouse: C57Bl/6J | The Jackson Laboratory | JAX: 000664 |
| | Mouse: C57Bl/6J | The Jackson Laboratory | JAX: 000664 |
| | Mouse: C57Bl/6J | The Jackson Laboratory | JAX: 000664 |
| | Mouse: STOCK Agrptm1(c | The Jackson Laboratory | JAX: 012899 |
| | Mouse: C57Bl/6J | The Jackson Laboratory | JAX: 000664 |
| | Mouse: C57Bl/6J | The Jackson Laboratory | JAX: 000664 |
| | Mouse: C57Bl/6J | The Jackson Laboratory | JAX: 000664 |
| | Mouse: C57Bl/6J | The Jackson Laboratory | JAX: 000664 |
| Y | Mouse: C57Bl/6J | The Jackson Laboratory | JAX: 000664 |
| | Mouse: STOCK Tg(Cdhr1-( | MMRRC | MMRRC: 030952 |
| | Mouse: STOCK Tg(Cdhr1-( | MMRRC | MMRRC: 030952 |
| | Mouse: Oxt-IRES-Cre | Bradford Lowell | N/A |
| Y | Mouse: C57Bl/6J | The Jackson Laboratory | JAX: 000664 |

| | | | |
|---|---|---|---|
| Y | Mouse: C57Bl/6J | The Jackson Laboratory | JAX: 000664 |
| | Mouse: C57Bl/6J | The Jackson Laboratory | JAX: 000664 |
| | Mouse: C57Bl/6J | The Jackson Laboratory | JAX: 000664 |
| | Mouse: C57Bl/6J | The Jackson Laboratory | JAX: 000664 |
| | Mouse: Oxt-IRES-Cre | Bradford Lowell | N/A |
| | Mouse: C57Bl/6J | The Jackson Laboratory | JAX: 000664 |
| | Mouse: C57Bl/6J | The Jackson Laboratory | JAX: 000664 |
| | Mouse: C57Bl/6J | The Jackson Laboratory | JAX: 000664 |
| | Mouse: Oxt-IRES-Cre | Bradford Lowell | N/A |
| | Mouse: C57Bl/6J | The Jackson Laboratory | JAX: 000664 |
| | Mouse: C57Bl/6J | The Jackson Laboratory | JAX: 000664 |
| | Mouse: C57Bl/6J | The Jackson Laboratory | JAX: 000664 |
| | Mouse: C57Bl/6J | The Jackson Laboratory | JAX: 000664 |
| | Mouse: C57Bl/6J | The Jackson Laboratory | JAX: 000664 |
| | Mouse: C57Bl/6J | The Jackson Laboratory | JAX: 000664 |
| | Mouse: C57Bl/6J | The Jackson Laboratory | JAX: 000664 |
| | Mouse: C57Bl/6J | The Jackson Laboratory | JAX: 000664 |
| | Mouse: C57Bl/6J | The Jackson Laboratory | JAX: 000664 |
| | Mouse: C57Bl/6J | The Jackson Laboratory | JAX: 000664 |
| | Mouse: B6.Cg-Tg(Pcp2-cre | MMRRC | MMRRC: 030868 |
| | Mouse: C57Bl/6J | The Jackson Laboratory | JAX: 000664 |
| | Mouse: C57Bl/6J | The Jackson Laboratory | JAX: 000664 |
| | Mouse: C57Bl/6J | The Jackson Laboratory | JAX: 000664 |
| | Mouse: B6.Cg-Tg(Pcp2-cre | MMRRC | MMRRC: 030868 |
| | Mouse: C57Bl/6J | The Jackson Laboratory | JAX: 000664 |
| | Mouse: C57Bl/6J | The Jackson Laboratory | JAX: 000664 |
| | Mouse: C57Bl/6J | The Jackson Laboratory | JAX: 000664 |
| | Mouse: Oxt-IRES-Cre | Bradford Lowell | N/A |
| | Mouse: Oxt-IRES-Cre | Bradford Lowell | N/A |
| | Mouse: C57Bl/6J | The Jackson Laboratory | JAX: 000664 |
| | Mouse: C57Bl/6J | The Jackson Laboratory | JAX: 000664 |
| | Mouse: C57Bl/6J | The Jackson Laboratory | JAX: 000664 |
| | Mouse: Pnmt-Cre | Steven Ebert | N/A |
| | Mouse: STOCK Tg(Slc6a5- | MMRRC | MMRRC: 030730 |
| | Mouse: B6.FVB(Cg)-Tg(Dr | MMRRC | MMRRC: 032108 |
| | Mouse: STOCK Tg(Sim1-cr | MMRRC | MMRRC: 031742 |
| | Mouse: B6;C3-Tg(Scnn1a- | The Jackson Laboratory | JAX: 009112 |
| | Mouse: B6;C3-Tg(Scnn1a- | The Jackson Laboratory | JAX: 009112 |
| | Mouse: STOCK Gad2tm2( | The Jackson Laboratory | JAX: 010802 |
| | Mouse: Pnmt-Cre | Steven Ebert | N/A |
| | Mouse: STOCK Gad2tm2( | The Jackson Laboratory | JAX: 010802 |
| | Mouse: B6;C3-Tg(Scnn1a- | The Jackson Laboratory | JAX: 009613 |
| | Mouse: B6;C3-Tg(Scnn1a- | The Jackson Laboratory | JAX: 009613 |
| | Mouse: B6;C3-Tg(Scnn1a- | The Jackson Laboratory | JAX: 009613 |
| | Mouse: B6;129-Thtm1(cr | The Jackson Laboratory | JAX: 008532 |
| | Mouse: STOCK Gad2tm2( | The Jackson Laboratory | JAX: 010802 |
| | Mouse: STOCK Tg(Gal-cre | MMRRC | MMRRC: 031060 |

| | | | |
|---|---|---|---|
| | Mouse: B6.Cg-Tg(A93003 | The Jackson Laboratory | JAX: 017346 |
| | Mouse: B6.Cg-Tg(A93003 | The Jackson Laboratory | JAX: 017346 |
| | Mouse: B6(Cg)-Etv1tm1.1 | The Jackson Laboratory | JAX: 013048 |
| | Mouse: B6(Cg)-Etv1tm1.1 | The Jackson Laboratory | JAX: 013048 |
| | Mouse: B6(Cg)-Etv1tm1.1 | The Jackson Laboratory | JAX: 013048 |
| | Mouse: B6.FVB(Cg)-Tg(Nt | MMRRC | MMRRC: 030648 |
| | Mouse: B6.FVB(Cg)-Tg(Nt | MMRRC | MMRRC: 030648 |
| | Mouse: B6.FVB(Cg)-Tg(Dr | MMRRC | MMRRC: 032108 |
| | Mouse: B6.FVB(Cg)-Tg(Nt | MMRRC | MMRRC: 030648 |
| Y | Mouse: STOCK Tg(Rbp4-c | MMRRC | MMRRC: 031125 |
| | Mouse: B6.Cg-Erbb4tm1. | The Jackson Laboratory | JAX: 012360 |
| | Mouse: B6.Cg-Erbb4tm1. | The Jackson Laboratory | JAX: 012360 |
| | Mouse: STOCK Tg(Syt6-cr | MMRRC | MMRRC: 032012 |
| | Mouse: STOCK Tg(Syt6-cr | MMRRC | MMRRC: 032012 |
| | Mouse: STOCK Tg(Syt6-cr | MMRRC | MMRRC: 032012 |
| | Mouse: STOCK Tg(Syt6-cr | MMRRC | MMRRC: 032012 |
| | Mouse: B6;129S6-Chattm | The Jackson Laboratory | JAX: 006410 |
| | Mouse: C57Bl/6J | The Jackson Laboratory | JAX: 000664 |
| | Mouse: C57Bl/6J | The Jackson Laboratory | JAX: 000664 |
| | Mouse: C57Bl/6J | The Jackson Laboratory | JAX: 000664 |
| | Mouse: C57Bl/6J | The Jackson Laboratory | JAX: 000664 |
| | Mouse: C57Bl/6J | The Jackson Laboratory | JAX: 000664 |
| | Mouse: C57Bl/6J | The Jackson Laboratory | JAX: 000664 |
| | Mouse: C57Bl/6J | The Jackson Laboratory | JAX: 000664 |
| | Mouse: C57Bl/6J | The Jackson Laboratory | JAX: 000664 |
| Y | Mouse: C57Bl/6J | The Jackson Laboratory | JAX: 000664 |
| | Mouse: C57Bl/6J | The Jackson Laboratory | JAX: 000664 |
| | Mouse: C57Bl/6J | The Jackson Laboratory | JAX: 000664 |
| Y | Mouse: C57Bl/6J | The Jackson Laboratory | JAX: 000664 |
| | Mouse: C57Bl/6J | The Jackson Laboratory | JAX: 000664 |
| | Mouse: STOCK Tg(Syt6-cr | MMRRC | MMRRC: 032012 |
| | Mouse: B6;129S6-Chattm | The Jackson Laboratory | JAX: 006410 |
| | Mouse: STOCK Tg(Pdzk1ip | MMRRC | MMRRC: 030851 |
| | Mouse: STOCK Tg(Pdzk1ip | MMRRC | MMRRC: 030851 |
| | Mouse: FVB-Tg(Nr5a1-cre | The Jackson Laboratory | JAX: 006364 |
| | Mouse: B6.FVB(Cg)-Tg(Nt | MMRRC | MMRRC: 030648 |
| | Mouse: STOCK Tg(Syt6-cr | MMRRC | MMRRC: 032012 |
| | Mouse: STOCK Corttm1(c | The Jackson Laboratory | JAX: 010910 |
| | Mouse: B6.FVB(Cg)-Tg(Nt | MMRRC | MMRRC: 030648 |
| | Mouse: STOCK Tg(Slc6a5- | MMRRC | MMRRC: 030730 |
| | Mouse: STOCK Gad2tm2( | The Jackson Laboratory | JAX: 010802 |
| | Mouse: STOCK Tg(Hdc-cr | MMRRC | MMRRC: 032079 |
| | Mouse: STOCK Tg(Hdc-cr | MMRRC | MMRRC: 032079 |
| | Mouse: STOCK Tg(Gng7-c | MMRRC | MMRRC: 031181 |
| | Mouse: STOCK Tg(Gng7-c | MMRRC | MMRRC: 031181 |
| | Mouse: STOCK Tg(Gng7-c | MMRRC | MMRRC: 031181 |
| | Mouse: STOCK Tg(Gng7-c | MMRRC | MMRRC: 031181 |

| | | |
|---|---|---|
| Mouse: B6.FVB(Cg)-Tg(Dr | MMRRC | MMRRC: 032108 |
| Mouse: B6.FVB(Cg)-Tg(Dr | MMRRC | MMRRC: 032108 |
| Mouse: B6;129P2-Pvalbtr | The Jackson Laboratory | JAX: 008069 |
| Mouse: B6;129P2-Pvalbtr | The Jackson Laboratory | JAX: 008069 |
| Mouse: B6.FVB(Cg)-Tg(Nt | MMRRC | MMRRC: 030648 |
| Mouse: STOCK Tg(Syt6-cr | MMRRC | MMRRC: 032012 |
| Mouse: B6.FVB-Tg(Pomc- | The Jackson Laboratory | JAX: 010714 |
| Mouse: B6.Cg-Tg(A93003 | The Jackson Laboratory | JAX: 017346 |
| Mouse: C57Bl/6J | The Jackson Laboratory | JAX: 000664 |
| Mouse: C57Bl/6J | The Jackson Laboratory | JAX: 000664 |
| Mouse: C57Bl/6J | The Jackson Laboratory | JAX: 000664 |
| Mouse: C57Bl/6J | The Jackson Laboratory | JAX: 000664 |
| Mouse: C57Bl/6J | The Jackson Laboratory | JAX: 000664 |
| Mouse: C57Bl/6J | The Jackson Laboratory | JAX: 000664 |
| Mouse: C57Bl/6J | The Jackson Laboratory | JAX: 000664 |
| Mouse: C57Bl/6J | The Jackson Laboratory | JAX: 000664 |
| Mouse: C57Bl/6J | The Jackson Laboratory | JAX: 000664 |
| Mouse: C57Bl/6J | The Jackson Laboratory | JAX: 000664 |
| Mouse: C57Bl/6J | The Jackson Laboratory | JAX: 000664 |
| Mouse: C57Bl/6J | The Jackson Laboratory | JAX: 000664 |
| Mouse: C57Bl/6J | The Jackson Laboratory | JAX: 000664 |
| Mouse: C57Bl/6J | The Jackson Laboratory | JAX: 000664 |
| Mouse: C57Bl/6J | The Jackson Laboratory | JAX: 000664 |
| Mouse: C57Bl/6J | The Jackson Laboratory | JAX: 000664 |
| Mouse: C57Bl/6J | The Jackson Laboratory | JAX: 000664 |
| Mouse: C57Bl/6J | The Jackson Laboratory | JAX: 000664 |
| Mouse: C57Bl/6J | The Jackson Laboratory | JAX: 000664 |
| Mouse: C57Bl/6J | The Jackson Laboratory | JAX: 000664 |
| Mouse: C57Bl/6J | The Jackson Laboratory | JAX: 000664 |
| Mouse: C57Bl/6J | The Jackson Laboratory | JAX: 000664 |
| Mouse: C57Bl/6J | The Jackson Laboratory | JAX: 000664 |
| Mouse: C57Bl/6J | The Jackson Laboratory | JAX: 000664 |
| Mouse: B6(Cg)-Etv1tm1.1 | The Jackson Laboratory | JAX: 013048 |
| Mouse: STOCK Tg(Sim1-cr | MMRRC | MMRRC: 031742 |
| Mouse: STOCK Tg(Syt6-cr | MMRRC | MMRRC: 032012 |
| Mouse: STOCK Tg(Sim1-cr | MMRRC | MMRRC: 031742 |
| Mouse: STOCK Tg(Drd1-c | MMRRC | MMRRC: 017264 |
| Mouse: STOCK Tg(Syt6-cr | MMRRC | MMRRC: 032012 |
| Mouse: STOCK Tg(Syt6-cr | MMRRC | MMRRC: 032012 |
| Mouse: STOCK Tg(Pdzk1ip | MMRRC | MMRRC: 030851 |
| Mouse: B6.Cg-Tg(A93003 | The Jackson Laboratory | JAX: 017346 |
| Mouse: B6.FVB(Cg)-Tg(Nt | MMRRC | MMRRC: 030648 |
| Mouse: B6.FVB(Cg)-Tg(Nt | MMRRC | MMRRC: 030648 |
| Mouse: B6.FVB(Cg)-Tg(Nt | MMRRC | MMRRC: 030648 |
| Mouse: B6.FVB(Cg)-Tg(Nt | MMRRC | MMRRC: 030648 |
| Mouse: B6.FVB(Cg)-Tg(Dr | MMRRC | MMRRC: 032108 |
| Mouse: STOCK Tg(Syt6-cr | MMRRC | MMRRC: 032012 |
| Mouse: STOCK Tg(Syt6-cr | MMRRC | MMRRC: 032012 |

| | | | |
|---|---|---|---|
| | Mouse: STOCK Ccktm1.1( | The Jackson Laboratory | JAX: 012706 |
| | Mouse: B6.FVB(Cg)-Tg(Nt | MMRRC | MMRRC: 030648 |
| | Mouse: STOCK Tg(Sim1-cr | MMRRC | MMRRC: 031742 |
| | Mouse: STOCK Ccktm1.1( | The Jackson Laboratory | JAX: 012706 |
| | Mouse: STOCK Ccktm1.1( | The Jackson Laboratory | JAX: 012706 |
| | Mouse: STOCK Tg(Syt6-cr | MMRRC | MMRRC: 032012 |
| | Mouse: STOCK Tg(Syt6-cr | MMRRC | MMRRC: 032012 |
| | Mouse: STOCK Tg(Dbh-cr | MMRRC | MMRRC: 032081 |
| | Mouse: B6.FVB(Cg)-Tg(Nt | MMRRC | MMRRC: 030648 |
| | Mouse: B6.FVB(Cg)-Tg(Nt | MMRRC | MMRRC: 030648 |
| | Mouse: B6.FVB(Cg)-Tg(Nt | MMRRC | MMRRC: 030648 |
| | Mouse: B6.FVB(Cg)-Tg(Nt | MMRRC | MMRRC: 030648 |
| | Mouse: B6.FVB(Cg)-Tg(Nt | MMRRC | MMRRC: 030648 |
| | Mouse: STOCK Tg(Cartpt- | The Jackson Laboratory | JAX: 009615 |
| | Mouse: C57Bl/6J | The Jackson Laboratory | JAX: 000664 |
| | Mouse: C57Bl/6J | The Jackson Laboratory | JAX: 000664 |
| | Mouse: C57Bl/6J | The Jackson Laboratory | JAX: 000664 |
| | Mouse: B6.FVB(Cg)-Tg(Nt | MMRRC | MMRRC: 030648 |
| | Mouse: B6.FVB(Cg)-Tg(Nt | MMRRC | MMRRC: 030648 |
| | Mouse: B6.FVB(Cg)-Tg(Nt | MMRRC | MMRRC: 030648 |
| | Mouse: B6.FVB-Tg(Pomc- | The Jackson Laboratory | JAX: 010714 |
| Y | Mouse: STOCK Tg(Rbp4-c | MMRRC | MMRRC: 031125 |
| Y | Mouse: STOCK Tg(Rbp4-c | MMRRC | MMRRC: 031125 |
| | Mouse: FVB-Tg(Nr5a1-cre | The Jackson Laboratory | JAX: 006364 |
| | Mouse: B6;C3-Tg(Scnn1a- | The Jackson Laboratory | JAX: 009613 |
| | Mouse: STOCK Tg(Drd1-c | MMRRC | MMRRC: 017264 |
| | Mouse: STOCK Tg(Drd1-c | MMRRC | MMRRC: 017264 |
| | Mouse: STOCK Tg(Dbh-cr | MMRRC | MMRRC: 032081 |
| | Mouse: B6.129P2-Gabra6 | MMRRC | MMRRC: 015968 |
| | Mouse: STOCK Tg(Syt17-c | MMRRC | MMRRC: 034355 |
| | Mouse: STOCK Tg(Pdzk1ip | MMRRC | MMRRC: 030851 |
| | Mouse: STOCK Tg(Pomc1- | The Jackson Laboratory | JAX: 005965 |
| | Mouse: STOCK Tg(Pomc1- | The Jackson Laboratory | JAX: 005965 |
| | Mouse: B6;129P2-Pvalbtr | The Jackson Laboratory | JAX: 008069 |
| | Mouse: STOCK Tg(Slc6a5- | MMRRC | MMRRC: 030730 |
| | Mouse: STOCK Tg(Slc6a5- | MMRRC | MMRRC: 030730 |
| | Mouse: STOCK Tg(Slc6a5- | MMRRC | MMRRC: 030730 |
| | Mouse: STOCK Tg(Slc6a5- | MMRRC | MMRRC: 030730 |
| | Mouse: STOCK Tg(Slc6a5- | MMRRC | MMRRC: 030730 |
| | Mouse: STOCK Tg(Slc6a5- | MMRRC | MMRRC: 030730 |
| | Mouse: Sst-Cre | Allen Institute for Brain S | N/A |
| | Mouse: B6.129-Leprtm2( | The Jackson Laboratory | JAX: 008320 |
| | Mouse: B6.129-Leprtm2( | The Jackson Laboratory | JAX: 008320 |
| | Mouse: STOCK Tg(Gnrh1- | The Jackson Laboratory | JAX: 021207 |
| | Mouse: STOCK Tg(Gnrh1- | The Jackson Laboratory | JAX: 021207 |
| | Mouse: STOCK Tg(Drd1-c | MMRRC | MMRRC: 017264 |
| | Mouse: STOCK Tg(Drd1-c | MMRRC | MMRRC: 017264 |

| | | | |
|---|---|---|---|
| | Mouse: STOCK Slc6a3tm1 | The Jackson Laboratory | JAX: 020080 |
| | Mouse: STOCK Slc6a3tm1 | The Jackson Laboratory | JAX: 020080 |
| | Mouse: B6.FVB(Cg)-Tg(Dr | MMRRC | MMRRC: 032108 |
| | Mouse: STOCK Slc6a3tm1 | The Jackson Laboratory | JAX: 020080 |
| | Mouse: STOCK Tg(Gabrr3 | MMRRC | MMRRC: 030709 |
| | Mouse: STOCK Tg(Syt17-c | MMRRC | MMRRC: 034355 |
| Y | Mouse: STOCK Tg(Rbp4-cr | MMRRC | MMRRC: 031125 |
| | Mouse: B6;129S6-Chattm | The Jackson Laboratory | JAX: 006410 |
| | Mouse: B6;129S6-Chattm | The Jackson Laboratory | JAX: 006410 |
| | Mouse: B6.FVB(Cg)-Tg(Nt | MMRRC | MMRRC: 030648 |
| | Mouse: B6;C3-Tg(Scnn1a- | The Jackson Laboratory | JAX: 009112 |
| | Mouse: B6;C3-Tg(Scnn1a- | The Jackson Laboratory | JAX: 009112 |
| | Mouse: B6;C3-Tg(Scnn1a- | The Jackson Laboratory | JAX: 009112 |
| | Mouse: STOCK Gad2tm2( | The Jackson Laboratory | JAX: 010802 |
| | Mouse: STOCK Gad2tm2( | The Jackson Laboratory | JAX: 010802 |
| | Mouse: STOCK Tg(Sim1-cr | MMRRC | MMRRC: 031742 |
| | Mouse: STOCK Tg(Sim1-cr | MMRRC | MMRRC: 031742 |
| | Mouse: STOCK Tg(Sim1-cr | MMRRC | MMRRC: 031742 |
| | Mouse: STOCK Corttm1(c | The Jackson Laboratory | JAX: 010910 |
| | Mouse: B6.Cg-Erbb4tm1. | The Jackson Laboratory | JAX: 012360 |
| | Mouse: B6;129S6-Chattm | The Jackson Laboratory | JAX: 006410 |
| | Mouse: STOCK Tg(Ucn3-c | MMRRC | MMRRC: 032078 |
| | Mouse: B6.Cg-Erbb4tm1. | The Jackson Laboratory | JAX: 012360 |
| | Mouse: STOCK Corttm1(c | The Jackson Laboratory | JAX: 010910 |
| | Mouse: B6;129S6-Chattm | The Jackson Laboratory | JAX: 006410 |
| | Mouse: STOCK Tg(Pdzk1ip | MMRRC | MMRRC: 030851 |
| Y | Mouse: STOCK Tg(Rbp4-cr | MMRRC | MMRRC: 031125 |
| | Mouse: STOCK Tg(Rbp4-cr | MMRRC | MMRRC: 031125 |
| | Mouse: STOCK Tg(Rbp4-cr | MMRRC | MMRRC: 031125 |
| Y | Mouse: STOCK Tg(Rbp4-cr | MMRRC | MMRRC: 031125 |
| Y | Mouse: STOCK Tg(Rbp4-cr | MMRRC | MMRRC: 031125 |
| | Mouse: STOCK Slc6a3tm1 | The Jackson Laboratory | JAX: 020080 |
| | Mouse: STOCK Slc6a3tm1 | The Jackson Laboratory | JAX: 020080 |
| | Mouse: STOCK Tg(Grm2-c | MMRRC | MMRRC: 034611 |
| | Mouse: STOCK Tg(Grm2-c | MMRRC | MMRRC: 034611 |
| | Mouse: B6;C3-Tg(Scnn1a- | The Jackson Laboratory | JAX: 009613 |
| | Mouse: B6;C3-Tg(Scnn1a- | The Jackson Laboratory | JAX: 009613 |
| | Mouse: B6;C3-Tg(Scnn1a- | The Jackson Laboratory | JAX: 009613 |
| | Mouse: B6.Cg-Tg(A93003 | The Jackson Laboratory | JAX: 017346 |
| | Mouse: B6.Cg-Tg(A93003 | The Jackson Laboratory | JAX: 017346 |
| | Mouse: B6.Cg-Tg(A93003 | The Jackson Laboratory | JAX: 017346 |
| | Mouse: B6.Cg-Tg(A93003 | The Jackson Laboratory | JAX: 017346 |
| | Mouse: B6;C3-Tg(Scnn1a- | The Jackson Laboratory | JAX: 009613 |
| | Mouse: B6;C3-Tg(Scnn1a- | The Jackson Laboratory | JAX: 009613 |
| | Mouse: B6;C3-Tg(Scnn1a- | The Jackson Laboratory | JAX: 009613 |
| | Mouse: B6;C3-Tg(Scnn1a- | The Jackson Laboratory | JAX: 009613 |
| | Mouse: B6;C3-Tg(Scnn1a- | The Jackson Laboratory | JAX: 009613 |

| | | | |
|---|---|---|---|
| | Mouse: B6.Cg-Tg(A93003{ | The Jackson Laboratory | JAX: 017346 |
| | Mouse: B6.Cg-Tg(A93003{ | The Jackson Laboratory | JAX: 017346 |
| | Mouse: B6.Cg-Tg(A93003{ | The Jackson Laboratory | JAX: 017346 |
| | Mouse: Oxt-IRES-Cre | Bradford Lowell | N/A |
| | Mouse: B6.FVB(Cg)-Tg(Nt | MMRRC | MMRRC: 030648 |
| | Mouse: B6;129P2-Pvalbtr | The Jackson Laboratory | JAX: 008069 |
| | Mouse: FVB-Tg(Nr5a1-cre | The Jackson Laboratory | JAX: 006364 |
| | Mouse: STOCK Tg(Ucn3-c | MMRRC | MMRRC: 032078 |
| | Mouse: B6.Cg-Erbb4tm1.: | The Jackson Laboratory | JAX: 012360 |
| | Mouse: STOCK Tg(Cnnm2 | MMRRC | MMRRC: 030951 |
| | Mouse: STOCK Agrptm1(c | The Jackson Laboratory | JAX: 012899 |
| | Mouse: STOCK Tg(Pdzk1ip | MMRRC | MMRRC: 030851 |
| | Mouse: B6;129S6-Chattm | The Jackson Laboratory | JAX: 006410 |
| | Mouse: B6;129P2-Pvalbtr | The Jackson Laboratory | JAX: 008069 |
| | Mouse: B6;129P2-Pvalbtr | The Jackson Laboratory | JAX: 008069 |
| | Mouse: B6;129P2-Pvalbtr | The Jackson Laboratory | JAX: 008069 |
| | Mouse: STOCK Tg(Slc6a4- | MMRRC | MMRRC: 030071 |
| | Mouse: B6(Cg)-Otoftm1.1 | MMRRC | MMRRC: 032781 |
| | Mouse: Crh-IRES-Cre_BL | Bradford Lowell | N/A |
| | Mouse: Crh-IRES-Cre_BL | Bradford Lowell | N/A |
| | Mouse: STOCK Corttm1(c | The Jackson Laboratory | JAX: 010910 |
| | Mouse: STOCK Corttm1(c | The Jackson Laboratory | JAX: 010910 |
| | Mouse: STOCK Tg(Cnnm2 | MMRRC | MMRRC: 030951 |
| | Mouse: C57BL/6-Tg(Grik4 | The Jackson Laboratory | JAX: 006474 |
| | Mouse: STOCK Gad2tm2( | The Jackson Laboratory | JAX: 010802 |
| | Mouse: STOCK Gad2tm2( | The Jackson Laboratory | JAX: 010802 |
| | Mouse: STOCK Tg(Syt17-c | MMRRC | MMRRC: 034355 |
| | Mouse: STOCK Gad2tm2( | The Jackson Laboratory | JAX: 010802 |
| | Mouse: STOCK Gad2tm2( | The Jackson Laboratory | JAX: 010802 |
| | Mouse: STOCK Gad2tm2( | The Jackson Laboratory | JAX: 010802 |
| | Mouse: B6(Cg)-Etv1tm1.1 | The Jackson Laboratory | JAX: 013048 |
| | Mouse: B6(Cg)-Etv1tm1.1 | The Jackson Laboratory | JAX: 013048 |
| | Mouse: C57BL/6-Tg(Grik4 | The Jackson Laboratory | JAX: 006474 |
| | Mouse: STOCK Slc17a6tm | The Jackson Laboratory | JAX: 016963 |
| | Mouse: B6.129-Leprtm2( | The Jackson Laboratory | JAX: 008320 |
| | Mouse: Crh-IRES-Cre_BL | Bradford Lowell | N/A |
| | Mouse: FVB-Tg(Nr5a1-cre | The Jackson Laboratory | JAX: 006364 |
| | Mouse: B6;C3-Tg(Wfs1-cr | The Jackson Laboratory | JAX: 009103 |
| | Mouse: STOCK Tg(Rbp4-cı | MMRRC | MMRRC: 031125 |
| Y | Mouse: STOCK Tg(Rbp4-cı | MMRRC | MMRRC: 031125 |
| | Mouse: B6.FVB(Cg)-Tg(Dr | MMRRC | MMRRC: 032108 |
| | Mouse: B6.FVB(Cg)-Tg(Dr | MMRRC | MMRRC: 032108 |
| | Mouse: B6(Cg)-Cux2tm1.: | MMRRC | MMRRC: 031778 |
| | Mouse: STOCK Tg(Gal-cre | MMRRC | MMRRC: 031060 |
| | Mouse: B6(Cg)-Cux2tm1.: | MMRRC | MMRRC: 031778 |
| | Mouse: STOCK Tg(Gal-cre | MMRRC | MMRRC: 031060 |
| | Mouse: STOCK Tg(Gal-cre | MMRRC | MMRRC: 031060 |

| | | | |
|---|---|---|---|
| | Mouse: STOCK Tg(Gnrh1- | The Jackson Laboratory | JAX: 021207 |
| | Mouse: STOCK Tg(Gal-cre | MMRRC | MMRRC: 031060 |
| | Mouse: STOCK Tg(Syt6-cr | MMRRC | MMRRC: 032012 |
| | Mouse: STOCK Tg(Syt6-cr | MMRRC | MMRRC: 032012 |
| | Mouse: STOCK Tg(Syt6-cr | MMRRC | MMRRC: 032012 |
| Y | Mouse: STOCK Tg(Rbp4-c | MMRRC | MMRRC: 031125 |
| | Mouse: STOCK Tg(Rbp4-c | MMRRC | MMRRC: 031125 |
| | Mouse: STOCK Tg(Rbp4-c | MMRRC | MMRRC: 031125 |
| | Mouse: B6.Cg-Tg(A93003 | The Jackson Laboratory | JAX: 017346 |
| | Mouse: B6.Cg-Tg(A93003 | The Jackson Laboratory | JAX: 017346 |
| | Mouse: STOCK Tg(Slc6a4- | MMRRC | MMRRC: 030071 |
| | Mouse: STOCK Tg(Syt6-cr | MMRRC | MMRRC: 032012 |
| | Mouse: STOCK Tg(Syt6-cr | MMRRC | MMRRC: 032012 |
| | Mouse: B6.Cg-Tg(Pcp2-cr | MMRRC | MMRRC: 030868 |
| | Mouse: B6.Cg-Tg(Pcp2-cr | MMRRC | MMRRC: 030868 |
| | Mouse: B6.Cg-Tg(Slc6a4-c | MMRRC | MMRRC: 031028 |
| | Mouse: C57BL/6-Tg(Grik4 | The Jackson Laboratory | JAX: 006474 |
| | Mouse: C57BL/6-Tg(Grik4 | The Jackson Laboratory | JAX: 006474 |
| | Mouse: C57BL/6-Tg(Grik4 | The Jackson Laboratory | JAX: 006474 |
| | Mouse: STOCK Gad2tm2( | The Jackson Laboratory | JAX: 010802 |
| | Mouse: STOCK Gad2tm2( | The Jackson Laboratory | JAX: 010802 |
| | Mouse: STOCK Tg(Syt6-cr | MMRRC | MMRRC: 032012 |
| | Mouse: STOCK Tg(Cartpt- | The Jackson Laboratory | JAX: 009615 |
| | Mouse: STOCK Tg(Cartpt- | The Jackson Laboratory | JAX: 009615 |
| | Mouse: B6;129P2-Pvalbtr | The Jackson Laboratory | JAX: 008069 |
| | Mouse: B6(Cg)-Calb2tm1 | The Jackson Laboratory | JAX: 010774 |
| | Mouse: B6(Cg)-Calb2tm1 | The Jackson Laboratory | JAX: 010774 |
| | Mouse: B6(Cg)-Calb2tm1 | The Jackson Laboratory | JAX: 010774 |
| | Mouse: STOCK Tg(Slc6a5- | MMRRC | MMRRC: 030730 |
| | Mouse: B6(Cg)-Calb2tm1 | The Jackson Laboratory | JAX: 010774 |
| | Mouse: STOCK Tg(Drd1-c | MMRRC | MMRRC: 017264 |
| | Mouse: B6;129S6-Chattm | The Jackson Laboratory | JAX: 006410 |
| | Mouse: STOCK Tg(Hdc-cr | MMRRC | MMRRC: 032079 |
| | Mouse: STOCK Slc17a6tm | The Jackson Laboratory | JAX: 016963 |
| | Mouse: STOCK Tg(Gabrr3 | MMRRC | MMRRC: 030709 |
| | Mouse: B6.FVB(Cg)-Tg(Nt | MMRRC | MMRRC: 030648 |
| | Mouse: STOCK Tg(Gabrr3 | MMRRC | MMRRC: 030709 |
| | Mouse: Crh-IRES-Cre_BL | Bradford Lowell | N/A |
| | Mouse: STOCK Tg(Drd1-c | MMRRC | MMRRC: 017264 |
| | Mouse: STOCK Tg(Gabrr3 | MMRRC | MMRRC: 030709 |
| | Mouse: STOCK Tg(Gabrr3 | MMRRC | MMRRC: 030709 |
| | Mouse: STOCK Tg(Sim1-cr | MMRRC | MMRRC: 031742 |
| | Mouse: STOCK Tg(Sim1-cr | MMRRC | MMRRC: 031742 |
| | Mouse: STOCK Tg(Sim1-cr | MMRRC | MMRRC: 031742 |
| | Mouse: STOCK Tg(Pdzk1ip | MMRRC | MMRRC: 030851 |
| | Mouse: STOCK Tg(Syt17-c | MMRRC | MMRRC: 034355 |
| | Mouse: B6(Cg)-Etv1tm1.1 | The Jackson Laboratory | JAX: 013048 |

| | | | |
|---|---|---|---|
| | Mouse: STOCK Slc32a1tm | The Jackson Laboratory | JAX: 016962 |
| | Mouse: STOCK Tg(Pmch-c | The Jackson Laboratory | JAX: 014099 |
| | Mouse: STOCK Slc6a3tm1 | The Jackson Laboratory | JAX: 020080 |
| | Mouse: STOCK Ccktm1.1( | The Jackson Laboratory | JAX: 012706 |
| | Mouse: STOCK Slc32a1tm | The Jackson Laboratory | JAX: 016962 |
| | Mouse: STOCK Slc32a1tm | The Jackson Laboratory | JAX: 016962 |
| | Mouse: STOCK Slc32a1tm | The Jackson Laboratory | JAX: 016962 |
| | Mouse: STOCK Slc32a1tm | The Jackson Laboratory | JAX: 016962 |
| | Mouse: B6(Cg)-Crhtm1(cr | The Jackson Laboratory | JAX: 012704 |
| | Mouse: STOCK Tg(Prkcd-g | MMRRC | MMRRC: 011559 |
| | Mouse: B6.Cg-Tg(A93003 | The Jackson Laboratory | JAX: 017346 |
| | Mouse: B6.Cg-Tg(A93003 | The Jackson Laboratory | JAX: 017346 |
| | Mouse: B6;C3-Tg(Scnn1a- | The Jackson Laboratory | JAX: 009112 |
| | Mouse: B6.129-Leprtm2( | The Jackson Laboratory | JAX: 008320 |
| | Mouse: STOCK Tg(Cartpt- | The Jackson Laboratory | JAX: 009615 |
| | Mouse: C57Bl/6J | The Jackson Laboratory | JAX: 000664 |
| Y | Mouse: C57BL/6J | The Jackson Laboratory | JAX: 000664 |
| | Mouse: C57Bl/6J | The Jackson Laboratory | JAX: 000664 |
| | Mouse: C57Bl/6J | The Jackson Laboratory | JAX: 000664 |
| | Mouse: C57Bl/6J | The Jackson Laboratory | JAX: 000664 |
| | Mouse: C57Bl/6J | The Jackson Laboratory | JAX: 000664 |
| | Mouse: C57Bl/6J | The Jackson Laboratory | JAX: 000664 |
| | Mouse: C57Bl/6J | The Jackson Laboratory | JAX: 000664 |
| | Mouse: C57Bl/6J | The Jackson Laboratory | JAX: 000664 |
| | Mouse: C57Bl/6J | The Jackson Laboratory | JAX: 000664 |
| | Mouse: C57Bl/6J | The Jackson Laboratory | JAX: 000664 |
| | Mouse: C57Bl/6J | The Jackson Laboratory | JAX: 000664 |
| | Mouse: C57Bl/6J | The Jackson Laboratory | JAX: 000664 |
| | Mouse: C57Bl/6J | The Jackson Laboratory | JAX: 000664 |
| | Mouse: C57Bl/6J | The Jackson Laboratory | JAX: 000664 |
| | Mouse: C57Bl/6J | The Jackson Laboratory | JAX: 000664 |
| | Mouse: C57Bl/6J | The Jackson Laboratory | JAX: 000664 |
| | Mouse: C57Bl/6J | The Jackson Laboratory | JAX: 000664 |
| | Mouse: C57Bl/6J | The Jackson Laboratory | JAX: 000664 |
| | Mouse: C57Bl/6J | The Jackson Laboratory | JAX: 000664 |
| | Mouse: C57Bl/6J | The Jackson Laboratory | JAX: 000664 |
| | Mouse: C57Bl/6J | The Jackson Laboratory | JAX: 000664 |
| | Mouse: C57Bl/6J | The Jackson Laboratory | JAX: 000664 |
| | Mouse: C57Bl/6J | The Jackson Laboratory | JAX: 000664 |
| | Mouse: B6(Cg)-Etv1tm1.1 | The Jackson Laboratory | JAX: 013048 |
| | Mouse: B6(Cg)-Calb2tm1( | The Jackson Laboratory | JAX: 010774 |
| | Mouse: STOCK Tg(Cdhr1- | MMRRC | MMRRC: 030952 |
| | Mouse: B6.FVB(Cg)-Tg(Dr | MMRRC | MMRRC: 032108 |
| | Mouse: B6.FVB(Cg)-Tg(Dr | MMRRC | MMRRC: 032108 |
| | Mouse: STOCK Tg(Hdc-cre | MMRRC | MMRRC: 032079 |
| | Mouse: STOCK Tg(Pomc1- | The Jackson Laboratory | JAX: 005965 |
| | Mouse: STOCK Slc6a3tm1 | The Jackson Laboratory | JAX: 020080 |

| | | |
|---|---|---|
| Mouse: STOCK Tg(Grm2-c | MMRRC | MMRRC: 034611 |
| Mouse: STOCK Tg(Grm2-c | MMRRC | MMRRC: 034611 |
| Mouse: STOCK Tg(Gal-cre | MMRRC | MMRRC: 031060 |
| Mouse: STOCK Tg(Gal-cre | MMRRC | MMRRC: 031060 |
| Mouse: STOCK Tg(Gal-cre | MMRRC | MMRRC: 031060 |
| Mouse: STOCK Tg(Gpr26- | MMRRC | MMRRC: 033032 |
| Mouse: B6(Cg)-Cux2tm1. | MMRRC | MMRRC: 031778 |
| Mouse: STOCK Tg(Gal-cre | MMRRC | MMRRC: 031060 |
| Mouse: STOCK Tg(Gal-cre | MMRRC | MMRRC: 031060 |
| Mouse: B6.Cg-Tg(A93003 | The Jackson Laboratory | JAX: 017346 |
| Mouse: B6;129P2-Pvalbtr | The Jackson Laboratory | JAX: 008069 |
| Mouse: STOCK Tg(Syt6-cr | MMRRC | MMRRC: 032012 |
| Mouse: FVB-Tg(Nr5a1-cre | The Jackson Laboratory | JAX: 006364 |
| Mouse: FVB-Tg(Nr5a1-cre | The Jackson Laboratory | JAX: 006364 |
| Mouse: Esr1-2A-Cre | David Anderson | N/A |
| Mouse: Esr1-2A-Cre | David Anderson | N/A |
| Mouse: Esr1-2A-Cre | David Anderson | N/A |
| Mouse: STOCK Tg(Cartpt- | The Jackson Laboratory | JAX: 009615 |
| Mouse: STOCK Tg(Cartpt- | The Jackson Laboratory | JAX: 009615 |
| Mouse: STOCK Tg(Kcnc2- | The Jackson Laboratory | JAX: 008582 |
| Mouse: STOCK Tg(Pdzk1ip | MMRRC | MMRRC: 030851 |
| Mouse: FVB-Tg(Nr5a1-cre | The Jackson Laboratory | JAX: 006364 |
| Mouse: STOCK Tg(Syt6-cr | MMRRC | MMRRC: 032012 |
| Mouse: FVB-Tg(Nr5a1-cre | The Jackson Laboratory | JAX: 006364 |
| Mouse: B6;129S6-Chattm | The Jackson Laboratory | JAX: 006410 |
| Mouse: B6;129S6-Chattm | The Jackson Laboratory | JAX: 006410 |
| Mouse: B6;129S6-Chattm | The Jackson Laboratory | JAX: 006410 |
| Mouse: C57BL/6-Tg(Grik4 | The Jackson Laboratory | JAX: 006474 |
| Mouse: C57BL/6-Tg(Grik4 | The Jackson Laboratory | JAX: 006474 |
| Mouse: FVB-Tg(Nr5a1-cre | The Jackson Laboratory | JAX: 006364 |
| Mouse: FVB-Tg(Nr5a1-cre | The Jackson Laboratory | JAX: 006364 |
| Mouse: FVB-Tg(Nr5a1-cre | The Jackson Laboratory | JAX: 006364 |
| Mouse: STOCK Tg(Syt6-cr | MMRRC | MMRRC: 032012 |
| Mouse: STOCK Tg(Syt6-cr | MMRRC | MMRRC: 032012 |
| Mouse: STOCK Tg(Syt6-cr | MMRRC | MMRRC: 032012 |
| Mouse: STOCK Tg(Syt6-cr | MMRRC | MMRRC: 032012 |
| Mouse: STOCK Tg(Sim1-cr | MMRRC | MMRRC: 031742 |
| Mouse: FVB-Tg(Nr5a1-cre | The Jackson Laboratory | JAX: 006364 |
| Mouse: FVB-Tg(Nr5a1-cre | The Jackson Laboratory | JAX: 006364 |
| Mouse: Pnmt-Cre | Steven Ebert | N/A |
| Mouse: B6;C3-Tg(Scnn1a- | The Jackson Laboratory | JAX: 009112 |
| Mouse: B6;C3-Tg(Scnn1a- | The Jackson Laboratory | JAX: 009112 |
| Mouse: B6.Cg-Tg(A93003 | The Jackson Laboratory | JAX: 017346 |
| Mouse: STOCK Tg(Gal-cre | MMRRC | MMRRC: 031060 |
| Mouse: STOCK Tg(Gal-cre | MMRRC | MMRRC: 031060 |
| Mouse: STOCK Tg(Gal-cre | MMRRC | MMRRC: 031060 |
| Mouse: B6(Cg)-Cux2tm1. | MMRRC | MMRRC: 031778 |

| | | |
|---|---|---|
| Mouse: C57BL/6-Tg(Grik4 | The Jackson Laboratory | JAX: 006474 |
| Mouse: STOCK Tg(Cartpt-c | The Jackson Laboratory | JAX: 009615 |
| Mouse: STOCK Tg(Syt6-cre | MMRRC | MMRRC: 032012 |
| Mouse: Oxt-IRES-Cre | Bradford Lowell | N/A |
| Mouse: STOCK Tg(Satb2-c | MMRRC | MMRRC: 032908 |
| Mouse: STOCK Tg(Prkcd-g | MMRRC | MMRRC: 011559 |
| Mouse: B6;C3-Tg(Scnn1a- | The Jackson Laboratory | JAX: 009613 |
| Mouse: STOCK Tg(Syt6-cre | MMRRC | MMRRC: 032012 |
| Mouse: STOCK Tg(Syt6-cre | MMRRC | MMRRC: 032012 |
| Mouse: C57BL/6-Tg(Grik4 | The Jackson Laboratory | JAX: 006474 |
| Mouse: STOCK Tg(Grp-cre | MMRRC | MMRRC: 031183 |
| Mouse: STOCK Tg(Grp-cre | MMRRC | MMRRC: 031183 |
| Mouse: STOCK Tg(Ucn3-c | MMRRC | MMRRC: 032078 |
| Mouse: B6.129-Leprtm2( | The Jackson Laboratory | JAX: 008320 |
| Mouse: STOCK Viptm1(cre | The Jackson Laboratory | JAX: 010908 |
| Mouse: STOCK Viptm1(cre | The Jackson Laboratory | JAX: 010908 |
| Mouse: C57Bl/6J | The Jackson Laboratory | JAX: 000664 |
| Mouse: STOCK Tg(Vipr2-c | MMRRC | MMRRC: 034281 |
| Mouse: C57Bl/6J | The Jackson Laboratory | JAX: 000664 |
| Mouse: STOCK Tg(Vipr2-c | MMRRC | MMRRC: 034281 |
| Mouse: B6.Cg-Erbb4tm1. | The Jackson Laboratory | JAX: 012360 |
| Mouse: C57Bl/6J | The Jackson Laboratory | JAX: 000664 |
| Mouse: B6;129S-Tac1tm1 | The Jackson Laboratory | JAX: 021877 |
| Mouse: B6;129S-Tac1tm1 | The Jackson Laboratory | JAX: 021877 |
| Mouse: C57Bl/6J | The Jackson Laboratory | JAX: 000664 |
| Mouse: C57Bl/6J | The Jackson Laboratory | JAX: 000664 |
| Mouse: C57Bl/6J | The Jackson Laboratory | JAX: 000664 |
| Mouse: C57Bl/6J | The Jackson Laboratory | JAX: 000664 |
| Mouse: C57Bl/6J | The Jackson Laboratory | JAX: 000664 |
| Mouse: C57Bl/6J | The Jackson Laboratory | JAX: 000664 |
| Mouse: C57Bl/6J | The Jackson Laboratory | JAX: 000664 |
| Mouse: C57Bl/6J | The Jackson Laboratory | JAX: 000664 |
| Mouse: B6.129P2-Gabra6 | MMRRC | MMRRC: 015968 |
| Mouse: B6.129P2-Gabra6 | MMRRC | MMRRC: 015968 |
| Mouse: B6.FVB(Cg)-Tg(Nt | MMRRC | MMRRC: 030648 |
| Mouse: B6.FVB(Cg)-Tg(Nt | MMRRC | MMRRC: 030648 |
| Mouse: B6;129S-Tac1tm1 | The Jackson Laboratory | JAX: 021877 |
| Mouse: B6;129S-Tac1tm1 | The Jackson Laboratory | JAX: 021877 |
| Mouse: B6;129S-Tac1tm1 | The Jackson Laboratory | JAX: 021877 |
| Mouse: STOCK Tg(Gpr26-c | MMRRC | MMRRC: 033032 |
| Mouse: STOCK Tg(Gpr26-c | MMRRC | MMRRC: 033032 |
| Mouse: STOCK Tg(Gpr26-c | MMRRC | MMRRC: 033032 |
| Mouse: STOCK Tg(Gpr26-c | MMRRC | MMRRC: 033032 |
| Mouse: STOCK Tg(Sim1-cr | MMRRC | MMRRC: 031742 |
| Mouse: STOCK Tg(Sim1-cr | MMRRC | MMRRC: 031742 |
| Mouse: B6;C3-Tg(Scnn1a- | The Jackson Laboratory | JAX: 009613 |
| Mouse: B6;C3-Tg(Scnn1a- | The Jackson Laboratory | JAX: 009613 |

| | | |
|---|---|---|
| Mouse: B6(Cg)-Etv1tm1.1 | The Jackson Laboratory | JAX: 013048 |
| Mouse: B6(Cg)-Cux2tm3. | MMRRC | MMRRC: 032779 |
| Mouse: B6;C3-Tg(Scnn1a- | The Jackson Laboratory | JAX: 009613 |
| Mouse: B6;C3-Tg(Scnn1a- | The Jackson Laboratory | JAX: 009613 |
| Mouse: B6(Cg)-Etv1tm1.1 | The Jackson Laboratory | JAX: 013048 |
| Mouse: B6(Cg)-Etv1tm1.1 | The Jackson Laboratory | JAX: 013048 |
| Mouse: B6.129-Leprtm2( | The Jackson Laboratory | JAX: 008320 |
| Mouse: STOCK Slc17a6tm | The Jackson Laboratory | JAX: 016963 |
| Mouse: C57BL/6-Tg(Grik4 | The Jackson Laboratory | JAX: 006474 |
| Mouse: STOCK Tg(Pdzk1ip | MMRRC | MMRRC: 030851 |
| Mouse: STOCK Tg(Pdzk1ip | MMRRC | MMRRC: 030851 |
| Mouse: STOCK Tg(Pdzk1ip | MMRRC | MMRRC: 030851 |
| Mouse: STOCK Viptm1(cr | The Jackson Laboratory | JAX: 010908 |
| Mouse: STOCK Tg(Cartpt- | The Jackson Laboratory | JAX: 009615 |
| Mouse: STOCK Slc6a3tm1 | The Jackson Laboratory | JAX: 020080 |
| Mouse: B6;129S-Nos1tm | The Jackson Laboratory | JAX: 014541 |
| Mouse: B6;C3-Tg(Scnn1a- | The Jackson Laboratory | JAX: 009613 |
| Mouse: STOCK Tg(Rbp4-c | MMRRC | MMRRC: 031125 |
| Mouse: STOCK Tg(Syt6-cr | MMRRC | MMRRC: 032012 |
| Mouse: STOCK Tg(Grm2-c | MMRRC | MMRRC: 034611 |
| Mouse: B6.Cg-Erbb4tm1. | The Jackson Laboratory | JAX: 012360 |
| Mouse: B6.Cg-Erbb4tm1. | The Jackson Laboratory | JAX: 012360 |
| Mouse: STOCK Tg(Grm2-c | MMRRC | MMRRC: 034611 |
| Mouse: B6(Cg)-Crhtm1(cr | The Jackson Laboratory | JAX: 012704 |
| Mouse: B6(Cg)-Otoftm1.1 | MMRRC | MMRRC: 032781 |
| Mouse: STOCK Tg(Vipr2-c | MMRRC | MMRRC: 034281 |
| Mouse: STOCK Tg(Vipr2-c | MMRRC | MMRRC: 034281 |
| Mouse: STOCK Tg(Slc6a5- | MMRRC | MMRRC: 030730 |
| Mouse: STOCK Viptm1(cr | The Jackson Laboratory | JAX: 010908 |
| Mouse: STOCK Tg(Rbp4-c | MMRRC | MMRRC: 031125 |
| Mouse: STOCK Tg(Rbp4-c | MMRRC | MMRRC: 031125 |
| Mouse: FVB-Tg(Nr5a1-cre | The Jackson Laboratory | JAX: 006364 |
| Mouse: FVB-Tg(Nr5a1-cre | The Jackson Laboratory | JAX: 006364 |
| Mouse: B6(Cg)-Cux2tm1. | MMRRC | MMRRC: 031778 |
| Mouse: STOCK Tg(Syt6-cr | MMRRC | MMRRC: 032012 |
| Mouse: STOCK Tg(Gal-cre | MMRRC | MMRRC: 031060 |
| Mouse: STOCK Tg(Gal-cre | MMRRC | MMRRC: 031060 |
| Mouse: STOCK Tg(Gal-cre | MMRRC | MMRRC: 031060 |
| Mouse: STOCK Tg(Syt6-cr | MMRRC | MMRRC: 032012 |
| Mouse: B6.Cg-Tg(A93003 | The Jackson Laboratory | JAX: 017346 |
| Mouse: B6(Cg)-Cux2tm1. | MMRRC | MMRRC: 031778 |
| Mouse: STOCK Tg(Gal-cre | MMRRC | MMRRC: 031060 |
| Mouse: B6;129P2-Pvalbtm | The Jackson Laboratory | JAX: 008069 |
| Mouse: STOCK Tg(Grp-cre | MMRRC | MMRRC: 031183 |
| Mouse: STOCK Tg(Grp-cre | MMRRC | MMRRC: 031183 |
| Mouse: B6(Cg)-Cux2tm1. | MMRRC | MMRRC: 031778 |
| Mouse: B6;C3-Tg(Wfs1-c | The Jackson Laboratory | JAX: 009103 |

| | | | |
|---|---|---|---|
| | Mouse: C57BL/6-Tg(Grik4 | The Jackson Laboratory | JAX: 006474 |
| | Mouse: STOCK Tg(Cdhr1- | MMRRC | MMRRC: 030952 |
| | Mouse: B6.Cg-Trib2tm1.1 | The Jackson Laboratory | JAX: 022865 |
| Y | Mouse: STOCK Tg(Rbp4-c | MMRRC | MMRRC: 031125 |
| | Mouse: STOCK Tg(Rbp4-c | MMRRC | MMRRC: 031125 |
| | Mouse: STOCK Tg(Grm2-c | MMRRC | MMRRC: 034611 |
| | Mouse: STOCK Tg(Rbp4-c | MMRRC | MMRRC: 031125 |
| | Mouse: B6;129P2-Pvalbtr | The Jackson Laboratory | JAX: 008069 |
| | Mouse: STOCK Tg(Vipr2-c | MMRRC | MMRRC: 034281 |
| | Mouse: Crh-IRES-Cre_BL | Bradford Lowell | N/A |
| | Mouse: STOCK Tg(Sim1-cr | MMRRC | MMRRC: 031742 |
| | Mouse: STOCK Tg(Kcnc2-( | The Jackson Laboratory | JAX: 008582 |
| | Mouse: STOCK Ssttm2.1( | The Jackson Laboratory | JAX: 013044 |
| | Mouse: B6(Cg)-Calb2tm1 | The Jackson Laboratory | JAX: 010774 |
| | Mouse: B6(Cg)-Calb2tm1 | The Jackson Laboratory | JAX: 010774 |
| | Mouse: B6(Cg)-Calb2tm1 | The Jackson Laboratory | JAX: 010774 |
| | Mouse: STOCK Tg(Slc18a2 | MMRRC | MMRRC: 034814 |
| | Mouse: B6(Cg)-Etv1tm1.1 | The Jackson Laboratory | JAX: 013048 |
| | Mouse: STOCK Tg(Kcnc2-( | The Jackson Laboratory | JAX: 008582 |
| | Mouse: STOCK Gad2tm2( | The Jackson Laboratory | JAX: 010802 |
| | Mouse: STOCK Viptm1(cr | The Jackson Laboratory | JAX: 010908 |
| | Mouse: B6;129S6-Chattm | The Jackson Laboratory | JAX: 006410 |
| | Mouse: C57BL/6-Tg(Grik4 | The Jackson Laboratory | JAX: 006474 |
| | Mouse: STOCK Tg(Grp-cre | MMRRC | MMRRC: 031183 |
| | Mouse: STOCK Tg(Grp-cre | MMRRC | MMRRC: 031183 |
| | Mouse: STOCK Tg(Grp-cre | MMRRC | MMRRC: 031183 |
| | Mouse: Crh-IRES-Cre_BL | Bradford Lowell | N/A |
| | Mouse: Crh-IRES-Cre_BL | Bradford Lowell | N/A |
| | Mouse: Crh-IRES-Cre_BL | Bradford Lowell | N/A |
| | Mouse: STOCK Tg(Rbp4-c | MMRRC | MMRRC: 031125 |
| | Mouse: Crh-IRES-Cre_BL | Bradford Lowell | N/A |
| | Mouse: STOCK Tg(Rbp4-c | MMRRC | MMRRC: 031125 |
| | Mouse: STOCK Tg(Rbp4-c | MMRRC | MMRRC: 031125 |
| | Mouse: C57BL/6-Tg(Grik4 | The Jackson Laboratory | JAX: 006474 |
| | Mouse: STOCK Tg(Grm2-c | MMRRC | MMRRC: 034611 |
| | Mouse: STOCK Tg(Grm2-c | MMRRC | MMRRC: 034611 |
| | Mouse: B6(Cg)-Calb2tm1 | The Jackson Laboratory | JAX: 010774 |
| | Mouse: B6(Cg)-Calb2tm1 | The Jackson Laboratory | JAX: 010774 |
| | Mouse: B6(Cg)-Calb2tm1 | The Jackson Laboratory | JAX: 010774 |
| | Mouse: B6(Cg)-Calb2tm1 | The Jackson Laboratory | JAX: 010774 |
| | Mouse: STOCK Tg(Gal-cre | MMRRC | MMRRC: 031060 |
| Y | Mouse: STOCK Tg(Rbp4-c | MMRRC | MMRRC: 031125 |
| | Mouse: B6;C3-Tg(Scnn1a- | The Jackson Laboratory | JAX: 009613 |
| | Mouse: STOCK Tg(Pdzk1ip | MMRRC | MMRRC: 030851 |
| | Mouse: STOCK Tg(Pdzk1ip | MMRRC | MMRRC: 030851 |
| | Mouse: STOCK Tg(Rbp4-c | MMRRC | MMRRC: 031125 |
| | Mouse: B6.Cg-Avptm1.1( | The Jackson Laboratory | JAX: 023530 |

| | | |
|---|---|---|
| Mouse: B6(Cg)-Cux2tm1.: | MMRRC | MMRRC: 031778 |
| Mouse: B6(Cg)-Cux2tm1.: | MMRRC | MMRRC: 031778 |
| Mouse: B6(Cg)-Cux2tm1.: | MMRRC | MMRRC: 031778 |
| Mouse: B6(Cg)-Cux2tm1.: | MMRRC | MMRRC: 031778 |
| Mouse: STOCK Tg(Grp-cre | MMRRC | MMRRC: 031183 |
| Mouse: STOCK Ccktm1.1( | The Jackson Laboratory | JAX: 012706 |
| Mouse: STOCK Tg(Htr2a-c | MMRRC | MMRRC: 031150 |
| Mouse: STOCK Tg(Htr2a-c | MMRRC | MMRRC: 031150 |
| Mouse: B6;129S6-Chattm | The Jackson Laboratory | JAX: 006410 |
| Mouse: B6;C3-Tg(Scnn1a- | The Jackson Laboratory | JAX: 009613 |
| Mouse: B6;C3-Tg(Scnn1a- | The Jackson Laboratory | JAX: 009613 |
| Mouse: B6;C3-Tg(Scnn1a- | The Jackson Laboratory | JAX: 009613 |
| Mouse: B6;C3-Tg(Scnn1a- | The Jackson Laboratory | JAX: 009613 |
| Mouse: B6(Cg)-Calb2tm1( | The Jackson Laboratory | JAX: 010774 |
| Mouse: B6(Cg)-Calb2tm1( | The Jackson Laboratory | JAX: 010774 |
| Mouse: B6(Cg)-Calb2tm1( | The Jackson Laboratory | JAX: 010774 |
| Mouse: B6(Cg)-Cux2tm1.: | MMRRC | MMRRC: 031778 |
| Mouse: B6(Cg)-Cux2tm1.: | MMRRC | MMRRC: 031778 |
| Mouse: B6(Cg)-Cux2tm1.: | MMRRC | MMRRC: 031778 |
| Mouse: STOCK Tg(Grp-cre | MMRRC | MMRRC: 031183 |
| Mouse: B6;129S4-Ntrk1tr | MMRRC | MMRRC: 015500 |
| Mouse: STOCK Tg(Grp-cre | MMRRC | MMRRC: 031183 |
| Mouse: STOCK Tg(Syt17-c | MMRRC | MMRRC: 034355 |
| Mouse: Adcyap1-2A-Cre | Allen Institute for Brain S | N/A |
| Mouse: B6(Cg)-Cux2tm3.: | MMRRC | MMRRC: 032779 |
| Mouse: B6.Cg-Erbb4tm1.: | The Jackson Laboratory | JAX: 012360 |
| Mouse: Adcyap1-2A-Cre | Allen Institute for Brain S | N/A |
| Mouse: B6(Cg)-Crhtm1(cr | The Jackson Laboratory | JAX: 012704 |
| Mouse: B6(Cg)-Crhtm1(cr | The Jackson Laboratory | JAX: 012704 |
| Mouse: STOCK Tg(Gal-cre) | MMRRC | MMRRC: 031060 |
| Mouse: STOCK Tg(Kiss1-cı | The Jackson Laboratory | JAX: 023426 |
| Mouse: STOCK Tg(Kiss1-cı | The Jackson Laboratory | JAX: 023426 |
| Mouse: STOCK Tg(Rbp4-cı | MMRRC | MMRRC: 031125 |
| Mouse: B6;129S-Tac2tm1 | The Jackson Laboratory | JAX: 021878 |
| Mouse: B6;129S-Tac2tm1 | The Jackson Laboratory | JAX: 021878 |
| Mouse: STOCK Tg(Sim1-cr | MMRRC | MMRRC: 031742 |
| Mouse: STOCK Tg(Sim1-cr | MMRRC | MMRRC: 031742 |
| Mouse: STOCK Tg(Rbp4-cı | MMRRC | MMRRC: 031125 |
| Mouse: STOCK Tg(Rbp4-cı | MMRRC | MMRRC: 031125 |
| Mouse: STOCK Tg(Rbp4-cı | MMRRC | MMRRC: 031125 |
| Mouse: STOCK Tg(Syt17-c | MMRRC | MMRRC: 034355 |
| Mouse: B6;C3-Tg(Scnn1a- | The Jackson Laboratory | JAX: 009613 |
| Mouse: B6;C3-Tg(Scnn1a- | The Jackson Laboratory | JAX: 009613 |
| Mouse: B6;C3-Tg(Scnn1a- | The Jackson Laboratory | JAX: 009613 |
| Mouse: STOCK Tg(Slc6a5- | MMRRC | MMRRC: 030730 |
| Mouse: STOCK Tg(Gpr26-( | MMRRC | MMRRC: 033032 |
| Mouse: B6;129P2-Pvalbtr | The Jackson Laboratory | JAX: 008069 |

| | | | |
|---|---|---|---|
| | Mouse: B6;129P2-Pvalbtr | The Jackson Laboratory | JAX: 008069 |
| | Mouse: STOCK Tg(Htr2a-c | MMRRC | MMRRC: 031150 |
| | Mouse: STOCK Tg(Htr2a-c | MMRRC | MMRRC: 031150 |
| | Mouse: STOCK Viptm1(cr | The Jackson Laboratory | JAX: 010908 |
| | Mouse: B6;129S4-Ntrk1tr | MMRRC | MMRRC: 015500 |
| | Mouse: STOCK Tg(Htr2a-c | MMRRC | MMRRC: 031150 |
| Y | Mouse: STOCK Tg(Rbp4-c | MMRRC | MMRRC: 031125 |
| | Mouse: B6;129S4-Ntrk1tr | MMRRC | MMRRC: 015500 |
| | Mouse: B6;C3-Tg(Scnn1a- | The Jackson Laboratory | JAX: 009112 |
| | Mouse: STOCK Tg(Rbp4-c | MMRRC | MMRRC: 031125 |
| | Mouse: STOCK Tg(Pomc1- | The Jackson Laboratory | JAX: 005965 |
| | Mouse: STOCK Tg(Slc18a2 | MMRRC | MMRRC: 034814 |
| | Mouse: STOCK Tg(Slc18a2 | MMRRC | MMRRC: 034814 |
| | Mouse: B6;C3-Tg(Scnn1a- | The Jackson Laboratory | JAX: 009112 |
| | Mouse: STOCK Slc6a3tm1 | The Jackson Laboratory | JAX: 020080 |
| | Mouse: B6(Cg)-Cux2tm1. | MMRRC | MMRRC: 031778 |
| | Mouse: B6(Cg)-Cux2tm1. | MMRRC | MMRRC: 031778 |
| | Mouse: STOCK Tg(Syt17-c | MMRRC | MMRRC: 034355 |
| | Mouse: STOCK Slc17a6tm | The Jackson Laboratory | JAX: 016963 |
| | Mouse: B6(Cg)-Cux2tm1. | MMRRC | MMRRC: 031778 |
| | Mouse: STOCK Tg(Gabrr3 | MMRRC | MMRRC: 030709 |
| | Mouse: STOCK Slc32a1tm | The Jackson Laboratory | JAX: 016962 |
| | Mouse: B6(Cg)-Cux2tm1. | MMRRC | MMRRC: 031778 |
| | Mouse: STOCK Slc32a1tm | The Jackson Laboratory | JAX: 016962 |
| | Mouse: STOCK Slc32a1tm | The Jackson Laboratory | JAX: 016962 |
| | Mouse: STOCK Slc32a1tm | The Jackson Laboratory | JAX: 016962 |
| | Mouse: STOCK Tg(Syt6-cr | MMRRC | MMRRC: 032012 |
| | Mouse: B6.FVB(Cg)-Tg(Nt | MMRRC | MMRRC: 030648 |
| | Mouse: STOCK Tg(Pdzk1ip | MMRRC | MMRRC: 030851 |
| | Mouse: Esr1-2A-Cre | David Anderson | N/A |
| | Mouse: Esr1-2A-Cre | David Anderson | N/A |
| | Mouse: Esr1-2A-Cre | David Anderson | N/A |
| | Mouse: Esr1-2A-Cre | David Anderson | N/A |
| | Mouse: STOCK Tg(Syt6-cr | MMRRC | MMRRC: 032012 |
| | Mouse: B6;129S6-Chattm | The Jackson Laboratory | JAX: 006410 |
| | Mouse: B6;129S6-Chattm | The Jackson Laboratory | JAX: 006410 |
| | Mouse: B6;129S6-Chattm | The Jackson Laboratory | JAX: 006410 |
| | Mouse: B6;129S6-Chattm | The Jackson Laboratory | JAX: 006410 |
| | Mouse: B6.Cg-Tg(A93003 | The Jackson Laboratory | JAX: 017346 |
| | Mouse: B6.Cg-Tg(A93003 | The Jackson Laboratory | JAX: 017346 |
| | Mouse: B6.Cg-Tg(A93003 | The Jackson Laboratory | JAX: 017346 |
| | Mouse: B6.Cg-Tg(A93003 | The Jackson Laboratory | JAX: 017346 |
| | Mouse: B6.Cg-Tg(A93003 | The Jackson Laboratory | JAX: 017346 |
| | Mouse: Crh-IRES-Cre_BL | Bradford Lowell | N/A |
| Y | Mouse: STOCK Tg(Grm2-c | MMRRC | MMRRC: 034611 |
| | Mouse: STOCK Tg(Grm2-c | MMRRC | MMRRC: 034611 |
| | Mouse: STOCK Tg(Grm2-c | MMRRC | MMRRC: 034611 |

| | | |
|---|---|---|
| Mouse: B6(Cg)-Cux2tm3.: | MMRRC | MMRRC: 032779 |
| Mouse: B6(Cg)-Cux2tm3.: | MMRRC | MMRRC: 032779 |
| Mouse: B6(Cg)-Calb2tm1( | The Jackson Laboratory | JAX: 010774 |
| Mouse: C57BL/6-Tg(Grik4 | The Jackson Laboratory | JAX: 006474 |
| Mouse: Oxt-IRES-Cre | Bradford Lowell | N/A |
| Mouse: B6;129S4-Ntrk1tr | MMRRC | MMRRC: 015500 |
| Mouse: B6;129S4-Ntrk1tr | MMRRC | MMRRC: 015500 |
| Mouse: B6;129S-Tac2tm1 | The Jackson Laboratory | JAX: 021878 |
| Mouse: STOCK Tg(Cartpt-c | The Jackson Laboratory | JAX: 009615 |
| Mouse: B6;129S-Tac2tm1 | The Jackson Laboratory | JAX: 021878 |
| Mouse: STOCK Tg(Grp-cre | MMRRC | MMRRC: 031183 |
| Mouse: STOCK Tg(Grp-cre | MMRRC | MMRRC: 031183 |
| Mouse: STOCK Tg(Grp-cre | MMRRC | MMRRC: 031183 |
| Mouse: STOCK Tg(Grp-cre | MMRRC | MMRRC: 031183 |
| Mouse: STOCK Tg(Grp-cre | MMRRC | MMRRC: 031183 |
| Mouse: STOCK Tg(Grp-cre | MMRRC | MMRRC: 031183 |
| Mouse: STOCK Tg(Slc18a2 | MMRRC | MMRRC: 034814 |
| Mouse: STOCK Tg(Slc18a2 | MMRRC | MMRRC: 034814 |
| Mouse: STOCK Tg(Gpr26-c | MMRRC | MMRRC: 033032 |
| Mouse: STOCK Tg(Gpr26-c | MMRRC | MMRRC: 033032 |
| Mouse: B6.Cg-Tg(Ins2-cre | The Jackson Laboratory | JAX: 003573 |
| Mouse: STOCK Tg(Syt6-cre | MMRRC | MMRRC: 032012 |
| Mouse: STOCK Tg(Slc18a2 | MMRRC | MMRRC: 034814 |
| Mouse: STOCK Tg(Gpr26-c | MMRRC | MMRRC: 033032 |
| Mouse: FVB-Tg(Nr5a1-cre | The Jackson Laboratory | JAX: 006364 |
| Mouse: B6;129S4-Ntrk1tr | MMRRC | MMRRC: 015500 |
| Mouse: B6;129S4-Ntrk1tr | MMRRC | MMRRC: 015500 |
| Mouse: B6;129S4-Ntrk1tr | MMRRC | MMRRC: 015500 |
| Mouse: B6;129S4-Ntrk1tr | MMRRC | MMRRC: 015500 |
| Mouse: B6;129S4-Ntrk1tr | MMRRC | MMRRC: 015500 |
| Mouse: STOCK Tg(Prkcd-g | MMRRC | MMRRC: 011559 |
| Mouse: STOCK Tg(Prkcd-g | MMRRC | MMRRC: 011559 |
| Mouse: STOCK Tg(Prkcd-g | MMRRC | MMRRC: 011559 |
| Mouse: B6;129P2-Pvalbtr | The Jackson Laboratory | JAX: 008069 |
| Mouse: FVB-Tg(Nr5a1-cre | The Jackson Laboratory | JAX: 006364 |
| Mouse: B6.Cg-Trib2tm1.1 | The Jackson Laboratory | JAX: 022865 |
| Mouse: STOCK Tg(Gal-cre | MMRRC | MMRRC: 031060 |
| Mouse: STOCK Tg(Gal-cre | MMRRC | MMRRC: 031060 |
| Mouse: STOCK Tg(Cartpt-c | The Jackson Laboratory | JAX: 009615 |
| Mouse: FVB-Tg(Nr5a1-cre | The Jackson Laboratory | JAX: 006364 |
| Mouse: B6.Cg-Trib2tm1.1 | The Jackson Laboratory | JAX: 022865 |
| Mouse: FVB-Tg(Nr5a1-cre | The Jackson Laboratory | JAX: 006364 |
| Mouse: STOCK Tg(Gal-cre | MMRRC | MMRRC: 031060 |
| Mouse: STOCK Tg(Gal-cre | MMRRC | MMRRC: 031060 |
| Mouse: STOCK Tg(Rbp4-cı | MMRRC | MMRRC: 031125 |
| Mouse: STOCK Tg(Rbp4-cı | MMRRC | MMRRC: 031125 |
| Mouse: STOCK Tg(Rbp4-cı | MMRRC | MMRRC: 031125 |

| | | |
|---|---|---|
| Mouse: B6(Cg)-Cux2tm1. | MMRRC | MMRRC: 031778 |
| Mouse: B6(Cg)-Cux2tm1. | MMRRC | MMRRC: 031778 |
| Mouse: STOCK Gad2tm2( | The Jackson Laboratory | JAX: 010802 |
| Mouse: STOCK Slc32a1tm | The Jackson Laboratory | JAX: 016962 |
| Mouse: STOCK Gad2tm2( | The Jackson Laboratory | JAX: 010802 |
| Mouse: B6.FVB(Cg)-Tg(Th | MMRRC | MMRRC: 031029 |
| Mouse: B6.FVB(Cg)-Tg(Th | MMRRC | MMRRC: 031029 |
| Mouse: STOCK Gad2tm2( | The Jackson Laboratory | JAX: 010802 |
| Mouse: STOCK Tg(Grp-cre | MMRRC | MMRRC: 031183 |
| Mouse: STOCK Tg(Grp-cre | MMRRC | MMRRC: 031183 |
| Mouse: C57Bl/6J | The Jackson Laboratory | JAX: 000664 |
| Mouse: STOCK Tg(Slc18a2 | MMRRC | MMRRC: 034814 |
| Mouse: B6(Cg)-Cux2tm1. | MMRRC | MMRRC: 031778 |
| Mouse: STOCK Gad2tm2( | The Jackson Laboratory | JAX: 010802 |
| Mouse: C57Bl/6J | The Jackson Laboratory | JAX: 000664 |
| Mouse: STOCK Tg(Htr2a-c | MMRRC | MMRRC: 031150 |
| Mouse: Crh-IRES-Cre_BL | Bradford Lowell | N/A |
| Mouse: STOCK Tg(Sim1-cr | MMRRC | MMRRC: 031742 |
| Mouse: Crh-IRES-Cre_BL | Bradford Lowell | N/A |
| Mouse: Crh-IRES-Cre_BL | Bradford Lowell | N/A |
| Mouse: Crh-IRES-Cre_BL | Bradford Lowell | N/A |
| Mouse: Crh-IRES-Cre_BL | Bradford Lowell | N/A |
| Mouse: STOCK Slc17a6tm | The Jackson Laboratory | JAX: 016963 |
| Mouse: B6.FVB(Cg)-Tg(Nt | MMRRC | MMRRC: 030648 |
| Mouse: B6.FVB(Cg)-Tg(Nt | MMRRC | MMRRC: 030648 |
| Mouse: B6.FVB(Cg)-Tg(Nt | MMRRC | MMRRC: 030648 |
| Mouse: STOCK Tg(Pdzk1ip | MMRRC | MMRRC: 030851 |
| Mouse: STOCK Tg(Pdzk1ip | MMRRC | MMRRC: 030851 |
| Mouse: B6;129P2-Pvalbtm | The Jackson Laboratory | JAX: 008069 |
| Mouse: STOCK Tg(Syt6-cre | MMRRC | MMRRC: 032012 |
| Mouse: B6;C3-Tg(Scnn1a- | The Jackson Laboratory | JAX: 009613 |
| Mouse: B6;129S-Slc17a7t | The Jackson Laboratory | JAX: 023527 |
| Mouse: B6;129S-Slc17a7t | The Jackson Laboratory | JAX: 023527 |
| Mouse: B6;129S-Slc17a7t | The Jackson Laboratory | JAX: 023527 |
| Mouse: B6;129S-Slc17a7t | The Jackson Laboratory | JAX: 023527 |
| Mouse: B6;129S6-Chattm | The Jackson Laboratory | JAX: 006410 |
| Mouse: B6(Cg)-Crhtm1(cr | The Jackson Laboratory | JAX: 012704 |
| Mouse: B6(Cg)-Crhtm1(cr | The Jackson Laboratory | JAX: 012704 |
| Mouse: STOCK Tg(Pcdh9- | MMRRC | MMRRC: 036084 |
| Mouse: B6(Cg)-Calb2tm1( | The Jackson Laboratory | JAX: 010774 |
| Mouse: B6(Cg)-Calb2tm1( | The Jackson Laboratory | JAX: 010774 |
| Mouse: B6.Cg-Avptm1.1( | The Jackson Laboratory | JAX: 023530 |
| Mouse: B6.Cg-Avptm1.1( | The Jackson Laboratory | JAX: 023530 |
| Mouse: B6.Cg-Tg(A93003 | The Jackson Laboratory | JAX: 017346 |
| Mouse: B6.Cg-Avptm1.1( | The Jackson Laboratory | JAX: 023530 |
| Mouse: B6(Cg)-Calb2tm1( | The Jackson Laboratory | JAX: 010774 |
| Mouse: C57Bl/6J | The Jackson Laboratory | JAX: 000664 |

| | | |
|---|---|---|
| Mouse: STOCK Slc32a1tm | The Jackson Laboratory | JAX: 016962 |
| Mouse: B6.Cg-Erbb4tm1. | The Jackson Laboratory | JAX: 012360 |
| Mouse: B6;129S-Fezf1tm | The Jackson Laboratory | JAX: 013048 |
| Mouse: B6;129S-Fezf1tm | The Jackson Laboratory | JAX: 013048 |
| Mouse: STOCK Tg(Prkcd-g | MMRRC | MMRRC: 011559 |
| Mouse: STOCK Tg(Prkcd-g | MMRRC | MMRRC: 011559 |
| Mouse: STOCK Tg(Prkcd-g | MMRRC | MMRRC: 011559 |
| Mouse: STOCK Tg(Prkcd-g | MMRRC | MMRRC: 011559 |
| Mouse: STOCK Tg(Prkcd-g | MMRRC | MMRRC: 011559 |
| Mouse: STOCK Tg(Gpr26- | MMRRC | MMRRC: 033032 |
| Mouse: STOCK Tg(Gpr26- | MMRRC | MMRRC: 033032 |
| Mouse: STOCK Tg(Gpr26- | MMRRC | MMRRC: 033032 |
| Mouse: STOCK Tg(Efr3a-c | MMRRC | MMRRC: 036660 |
| Mouse: STOCK Tg(Efr3a-c | MMRRC | MMRRC: 036660 |
| Mouse: STOCK Tg(Efr3a-c | MMRRC | MMRRC: 036660 |
| Mouse: STOCK Tg(Slc6a5- | MMRRC | MMRRC: 030730 |
| Mouse: STOCK Tg(Efr3a-c | MMRRC | MMRRC: 036660 |
| Mouse: STOCK Tg(Efr3a-c | MMRRC | MMRRC: 036660 |
| Mouse: STOCK Tg(Efr3a-c | MMRRC | MMRRC: 036660 |
| Mouse: STOCK Tg(Chrna2 | MMRRC | MMRRC: 036502 |
| Mouse: STOCK Tg(Chrna2 | MMRRC | MMRRC: 036502 |
| Mouse: STOCK Tg(Efr3a-c | MMRRC | MMRRC: 036660 |
| Mouse: STOCK Tg(Efr3a-c | MMRRC | MMRRC: 036660 |
| Mouse: STOCK Tg(Oxtr-cr | MMRRC | MMRRC: 036545 |
| Mouse: STOCK Tg(Oxtr-cr | MMRRC | MMRRC: 036545 |
| Mouse: STOCK Tg(Ppp1r1 | MMRRC | MMRRC: 036205 |
| Mouse: STOCK Tg(Ppp1r1 | MMRRC | MMRRC: 036205 |
| Mouse: STOCK Tg(Ppp1r1 | MMRRC | MMRRC: 036205 |
| Mouse: STOCK Tg(Ppp1r1 | MMRRC | MMRRC: 036205 |
| Mouse: B6;129S-Nos1tm | The Jackson Laboratory | JAX: 014541 |
| Mouse: B6(Cg)-Etv1tm1.1 | The Jackson Laboratory | JAX: 013048 |
| Mouse: STOCK Tg(Ppp1r1 | MMRRC | MMRRC: 036205 |
| Mouse: STOCK Tg(Prkcd-g | MMRRC | MMRRC: 011559 |
| Mouse: STOCK Tg(Prkcd-g | MMRRC | MMRRC: 011559 |
| Mouse: B6;129S-Nos1tm | The Jackson Laboratory | JAX: 014541 |
| Mouse: B6;129S-Nos1tm | The Jackson Laboratory | JAX: 014541 |
| Mouse: B6;129S-Nos1tm | The Jackson Laboratory | JAX: 014541 |
| Mouse: B6.Cg-Tg(Slc6a4-c | MMRRC | MMRRC: 031028 |
| Mouse: STOCK Tg(Vipr2-c | MMRRC | MMRRC: 034281 |
| Mouse: B6.Cg-Avptm1.1( | The Jackson Laboratory | JAX: 023530 |
| Mouse: C57BL/6-Tg(Grik4 | The Jackson Laboratory | JAX: 006474 |
| Mouse: C57BL/6-Tg(Grik4 | The Jackson Laboratory | JAX: 006474 |
| Mouse: C57BL/6-Tg(Grik4 | The Jackson Laboratory | JAX: 006474 |
| Mouse: B6;C3-Tg(Scnn1a- | The Jackson Laboratory | JAX: 009613 |
| Mouse: B6;C3-Tg(Scnn1a- | The Jackson Laboratory | JAX: 009613 |
| Mouse: B6;C3-Tg(Scnn1a- | The Jackson Laboratory | JAX: 009613 |
| Mouse: B6;C3-Tg(Scnn1a- | The Jackson Laboratory | JAX: 009613 |

| | | | |
|---|---|---|---|
| | Mouse: STOCK Tg(Slc18a2 | MMRRC | MMRRC: 034814 |
| | Mouse: STOCK Tg(Vipr2-c | MMRRC | MMRRC: 034281 |
| | Mouse: STOCK Tg(Vipr2-c | MMRRC | MMRRC: 034281 |
| | Mouse: STOCK Tg(Ppp1r1 | MMRRC | MMRRC: 036205 |
| | Mouse: STOCK Tg(Ppp1r1 | MMRRC | MMRRC: 036205 |
| | Mouse: STOCK Tg(Ppp1r1 | MMRRC | MMRRC: 036205 |
| | Mouse: STOCK Tg(Ppp1r1 | MMRRC | MMRRC: 036205 |
| | Mouse: STOCK Tg(Ppp1r1 | MMRRC | MMRRC: 036205 |
| | Mouse: B6.Cg-Erbb4tm1.: | The Jackson Laboratory | JAX: 012360 |
| | Mouse: B6;129S6-Chattm | The Jackson Laboratory | JAX: 006410 |
| | Mouse: STOCK Tg(Ppp1r1 | MMRRC | MMRRC: 036205 |
| | Mouse: C57Bl/6J | The Jackson Laboratory | JAX: 000664 |
| | Mouse: STOCK Tg(Ppp1r1 | MMRRC | MMRRC: 036205 |
| | Mouse: STOCK Tg(Ppp1r1 | MMRRC | MMRRC: 036205 |
| | Mouse: C57Bl/6J | The Jackson Laboratory | JAX: 000664 |
| | Mouse: C57Bl/6J | The Jackson Laboratory | JAX: 000664 |
| | Mouse: C57Bl/6J | The Jackson Laboratory | JAX: 000664 |
| | Mouse: B6.FVB(Cg)-Tg(Th | MMRRC | MMRRC: 031029 |
| | Mouse: STOCK Tg(Rbp4-cı | MMRRC | MMRRC: 031125 |
| | Mouse: B6.FVB(Cg)-Tg(Th | MMRRC | MMRRC: 031029 |
| | Mouse: C57Bl/6J | The Jackson Laboratory | JAX: 000664 |
| | Mouse: STOCK Tg(Rbp4-cı | MMRRC | MMRRC: 031125 |
| | Mouse: B6(Cg)-Cux2tm1.: | MMRRC | MMRRC: 031778 |
| | Mouse: B6(Cg)-Cux2tm1.: | MMRRC | MMRRC: 031778 |
| | Mouse: C57Bl/6J | The Jackson Laboratory | JAX: 000664 |
| Y | Mouse: B6(Cg)-Cux2tm1.: | MMRRC | MMRRC: 031778 |
| | Mouse: Adcyap1-2A-Cre | Allen Institute for Brain S | N/A |
| | Mouse: B6(Cg)-Cux2tm1.: | MMRRC | MMRRC: 031778 |
| | Mouse: B6(Cg)-Cux2tm1.: | MMRRC | MMRRC: 031778 |
| | Mouse: C57Bl/6J | The Jackson Laboratory | JAX: 000664 |
| | Mouse: C57Bl/6J | The Jackson Laboratory | JAX: 000664 |
| Y | Mouse: B6(Cg)-Cux2tm1.: | MMRRC | MMRRC: 031778 |
| | Mouse: STOCK Tg(Pcdh9- | MMRRC | MMRRC: 036084 |
| | Mouse: C57Bl/6J | The Jackson Laboratory | JAX: 000664 |
| | Mouse: C57Bl/6J | The Jackson Laboratory | JAX: 000664 |
| | Mouse: C57Bl/6J | The Jackson Laboratory | JAX: 000664 |
| | Mouse: C57Bl/6J | The Jackson Laboratory | JAX: 000664 |
| | Mouse: C57Bl/6J | The Jackson Laboratory | JAX: 000664 |
| | Mouse: C57Bl/6J | The Jackson Laboratory | JAX: 000664 |
| | Mouse: C57Bl/6J | The Jackson Laboratory | JAX: 000664 |
| | Mouse: C57Bl/6J | The Jackson Laboratory | JAX: 000664 |
| Y | Mouse: STOCK Tg(Rbp4-cı | MMRRC | MMRRC: 031125 |
| | Mouse: C57Bl/6J | The Jackson Laboratory | JAX: 000664 |
| | Mouse: C57Bl/6J | The Jackson Laboratory | JAX: 000664 |
| | Mouse: C57Bl/6J | The Jackson Laboratory | JAX: 000664 |
| | Mouse: C57Bl/6J | The Jackson Laboratory | JAX: 000664 |
| | Mouse: C57Bl/6J | The Jackson Laboratory | JAX: 000664 |

| | | |
|---|---|---|
| Mouse: C57Bl/6J | The Jackson Laboratory | JAX: 000664 |
| Mouse: C57Bl/6J | The Jackson Laboratory | JAX: 000664 |
| Mouse: C57Bl/6J | The Jackson Laboratory | JAX: 000664 |
| Mouse: C57Bl/6J | The Jackson Laboratory | JAX: 000664 |
| Mouse: C57Bl/6J | The Jackson Laboratory | JAX: 000664 |
| Mouse: C57Bl/6J | The Jackson Laboratory | JAX: 000664 |
| Mouse: C57Bl/6J | The Jackson Laboratory | JAX: 000664 |
| Mouse: C57Bl/6J | The Jackson Laboratory | JAX: 000664 |
| Mouse: B6;129S4-Ntrk1tr | MMRRC | MMRRC: 015500 |
| Mouse: B6;129S4-Ntrk1tr | MMRRC | MMRRC: 015500 |
| Mouse: STOCK Ssttm2.1(c | The Jackson Laboratory | JAX: 013044 |
| Mouse: STOCK Ssttm2.1(c | The Jackson Laboratory | JAX: 013044 |
| Mouse: STOCK Ccktm1.1( | The Jackson Laboratory | JAX: 012706 |
| Mouse: FVB-Tg(Nr5a1-cre | The Jackson Laboratory | JAX: 006364 |
| Mouse: FVB-Tg(Nr5a1-cre | The Jackson Laboratory | JAX: 006364 |
| Mouse: B6;C3-Tg(Scnn1a- | The Jackson Laboratory | JAX: 009613 |
| Mouse: Crh-IRES-Cre_BL | Bradford Lowell | N/A |
| Mouse: B6.Cg-Erbb4tm1. | The Jackson Laboratory | JAX: 012360 |
| Mouse: B6.Cg-Erbb4tm1. | The Jackson Laboratory | JAX: 012360 |
| Mouse: Crh-IRES-Cre_BL | Bradford Lowell | N/A |
| Mouse: Crh-IRES-Cre_BL | Bradford Lowell | N/A |
| Mouse: Crh-IRES-Cre_BL | Bradford Lowell | N/A |
| Mouse: STOCK Tg(Efr3a-cı | MMRRC | MMRRC: 036660 |
| Mouse: STOCK Tg(Efr3a-cı | MMRRC | MMRRC: 036660 |
| Mouse: STOCK Tg(Htr2a-c | MMRRC | MMRRC: 031150 |
| Mouse: STOCK Tg(Htr2a-c | MMRRC | MMRRC: 031150 |
| Mouse: B6;129P2-Pvalbtr | The Jackson Laboratory | JAX: 008069 |
| Mouse: B6;129P2-Pvalbtr | The Jackson Laboratory | JAX: 008069 |
| Mouse: B6;129P2-Pvalbtr | The Jackson Laboratory | JAX: 008069 |
| Mouse: STOCK Tg(Prkcd-g | MMRRC | MMRRC: 011559 |
| Mouse: STOCK Tg(Chrna2 | MMRRC | MMRRC: 036502 |
| Mouse: STOCK Tg(Chrna2 | MMRRC | MMRRC: 036502 |
| Mouse: STOCK Tg(Gal-cre | MMRRC | MMRRC: 031060 |
| Mouse: STOCK Tg(Oxtr-cr | MMRRC | MMRRC: 036545 |
| Mouse: STOCK Tg(Chrna2 | MMRRC | MMRRC: 036502 |
| Mouse: STOCK Tg(Chrna2 | MMRRC | MMRRC: 036502 |
| Mouse: STOCK Tg(Chrnb4 | MMRRC | MMRRC: 036203 |
| Mouse: STOCK Tg(Chrna2 | MMRRC | MMRRC: 036502 |
| Mouse: Crh-IRES-Cre_BL | Bradford Lowell | N/A |
| Mouse: STOCK Tg(Efr3a-cı | MMRRC | MMRRC: 036660 |
| Mouse: Crh-IRES-Cre_BL | Bradford Lowell | N/A |
| Mouse: STOCK Tg(Efr3a-cı | MMRRC | MMRRC: 036660 |
| Mouse: STOCK Tg(Efr3a-cı | MMRRC | MMRRC: 036660 |
| Mouse: STOCK Tg(Efr3a-cı | MMRRC | MMRRC: 036660 |
| Mouse: STOCK Tg(Efr3a-cı | MMRRC | MMRRC: 036660 |
| Mouse: STOCK Tg(Oxtr-cr | MMRRC | MMRRC: 036545 |
| Mouse: STOCK Tg(Oxtr-cr | MMRRC | MMRRC: 036545 |

| | | | |
|---|---|---|---|
| | Mouse: STOCK Tg(Oxtr-cre | MMRRC | MMRRC: 036545 |
| | Mouse: B6(Cg)-Cux2tm1.: | MMRRC | MMRRC: 031778 |
| | Mouse: B6(Cg)-Cux2tm1.: | MMRRC | MMRRC: 031778 |
| | Mouse: STOCK Tg(Ppp1r1 | MMRRC | MMRRC: 036205 |
| | Mouse: STOCK Tg(Ppp1r1 | MMRRC | MMRRC: 036205 |
| | Mouse: STOCK Tg(Ppp1r1 | MMRRC | MMRRC: 036205 |
| | Mouse: STOCK Tg(Ppp1r1 | MMRRC | MMRRC: 036205 |
| | Mouse: STOCK Tg(Ppp1r1 | MMRRC | MMRRC: 036205 |
| | Mouse: STOCK Tg(Ppp1r1 | MMRRC | MMRRC: 036205 |
| | Mouse: STOCK Slc17a6tm | The Jackson Laboratory | JAX: 016963 |
| | Mouse: STOCK Tg(Htr2a-c | MMRRC | MMRRC: 031150 |
| Y | Mouse: B6(Cg)-Cux2tm1.: | MMRRC | MMRRC: 031778 |
| | Mouse: B6.Cg-Penktm1.1 | The Jackson Laboratory | JAX: 022862 |
| | Mouse: B6.Cg-Tg(Wfs1-cr | The Jackson Laboratory | JAX: 009614 |
| | Mouse: B6.Cg-Tg(Wfs1-cr | The Jackson Laboratory | JAX: 009614 |
| | Mouse: STOCK Tg(Htr2a-c | MMRRC | MMRRC: 031150 |
| | Mouse: STOCK Tg(Htr2a-c | MMRRC | MMRRC: 031150 |
| | Mouse: B6;C3-Tg(Scnn1a- | The Jackson Laboratory | JAX: 009613 |
| | Mouse: STOCK Tg(Ppp1r1 | MMRRC | MMRRC: 036205 |
| | Mouse: STOCK Tg(Ppp1r1 | MMRRC | MMRRC: 036205 |
| | Mouse: STOCK Tg(Ppp1r1 | MMRRC | MMRRC: 036205 |
| | Mouse: STOCK Tg(Ppp1r1 | MMRRC | MMRRC: 036205 |
| | Mouse: STOCK Tg(Ppp1r1 | MMRRC | MMRRC: 036205 |
| | Mouse: STOCK Tg(Ppp1r1 | MMRRC | MMRRC: 036205 |
| | Mouse: STOCK Tg(Ppp1r1 | MMRRC | MMRRC: 036205 |
| | Mouse: B6.Cg-Tg(A93003 | The Jackson Laboratory | JAX: 017346 |
| | Mouse: B6.Cg-Tg(A93003 | The Jackson Laboratory | JAX: 017346 |
| | Mouse: B6.Cg-Tg(A93003 | The Jackson Laboratory | JAX: 017346 |
| | Mouse: B6.Cg-Tg(A93003 | The Jackson Laboratory | JAX: 017346 |
| | Mouse: STOCK Tg(Grm2-c | MMRRC | MMRRC: 034611 |
| | Mouse: B6(Cg)-Cux2tm1.: | MMRRC | MMRRC: 031778 |
| | Mouse: B6(Cg)-Cux2tm1.: | MMRRC | MMRRC: 031778 |
| | Mouse: B6(Cg)-Cux2tm1.: | MMRRC | MMRRC: 031778 |
| | Mouse: STOCK Slc6a3tm1 | The Jackson Laboratory | JAX: 020080 |
| Y | Mouse: STOCK Tg(Rbp4-cr | MMRRC | MMRRC: 031125 |
| | Mouse: STOCK Tg(Rbp4-cr | MMRRC | MMRRC: 031125 |
| | Mouse: B6;129S-Slc17a7t | The Jackson Laboratory | JAX: 023527 |
| | Mouse: B6.Cg-Tg(Ins2-cre | The Jackson Laboratory | JAX: 003573 |
| | Mouse: B6.Cg-Tg(Ins2-cre | The Jackson Laboratory | JAX: 003573 |
| | Mouse: B6;129S4-Ntrk1tr | MMRRC | MMRRC: 015500 |
| | Mouse: STOCK Tg(Rbp4-cr | MMRRC | MMRRC: 031125 |
| Y | Mouse: STOCK Tg(Rbp4-cr | MMRRC | MMRRC: 031125 |
| | Mouse: STOCK Tg(Rbp4-cr | MMRRC | MMRRC: 031125 |
| | Mouse: STOCK Tg(Syt17-c | MMRRC | MMRRC: 034355 |
| | Mouse: STOCK Tg(Syt17-c | MMRRC | MMRRC: 034355 |
| | Mouse: STOCK Tg(Gpr26-c | MMRRC | MMRRC: 033032 |
| | Mouse: STOCK Tg(Gpr26-c | MMRRC | MMRRC: 033032 |

| | | |
|---|---|---|
| Mouse: STOCK Tg(Gpr26-[illegible] | MMRRC | MMRRC: 033032 |
| Mouse: FVB-Tg(Nr5a1-cre | The Jackson Laboratory | JAX: 006364 |
| Mouse: STOCK Slc17a6tm | The Jackson Laboratory | JAX: 016963 |
| Mouse: STOCK Slc17a6tm | The Jackson Laboratory | JAX: 016963 |
| Mouse: STOCK Slc17a6tm | The Jackson Laboratory | JAX: 016963 |
| Mouse: STOCK Slc17a6tm | The Jackson Laboratory | JAX: 016963 |
| Mouse: STOCK Tg(Syt17-c | MMRRC | MMRRC: 034355 |
| Mouse: STOCK Tg(Syt17-c | MMRRC | MMRRC: 034355 |
| Mouse: STOCK Tg(Syt17-c | MMRRC | MMRRC: 034355 |
| Mouse: B6;129S-Tac1tm1 | The Jackson Laboratory | JAX: 021877 |
| Mouse: STOCK Slc32a1tm | The Jackson Laboratory | JAX: 016962 |
| Mouse: STOCK Tg(Efr3a-c[illegible] | MMRRC | MMRRC: 036660 |
| Mouse: STOCK Tg(Efr3a-c[illegible] | MMRRC | MMRRC: 036660 |
| Mouse: STOCK Tg(Ppp1r1 | MMRRC | MMRRC: 036205 |
| Mouse: STOCK Gad2tm2([illegible] | The Jackson Laboratory | JAX: 010802 |
| Mouse: B6;129P2-Pvalbtr | The Jackson Laboratory | JAX: 008069 |
| Mouse: B6.Cg-Avptm1.1([illegible] | The Jackson Laboratory | JAX: 023530 |
| Mouse: B6.FVB(Cg)-Tg(Th | MMRRC | MMRRC: 031029 |
| Mouse: STOCK Tg(Ppp1r1 | MMRRC | MMRRC: 036205 |
| Mouse: B6(Cg)-Cux2tm3.[illegible] | MMRRC | MMRRC: 032779 |
| Mouse: B6.FVB(Cg)-Tg(Th | MMRRC | MMRRC: 031029 |
| Mouse: B6(Cg)-Etv1tm1.1 | The Jackson Laboratory | JAX: 013048 |
| Mouse: STOCK Tg(Efr3a-c[illegible] | MMRRC | MMRRC: 036660 |
| Mouse: B6(Cg)-Etv1tm1.1 | The Jackson Laboratory | JAX: 013048 |
| Mouse: B6.Cg-Tg(A93003[illegible] | The Jackson Laboratory | JAX: 017346 |
| Mouse: B6.Cg-Tg(A93003[illegible] | The Jackson Laboratory | JAX: 017346 |
| Mouse: B6.Cg-Tg(A93003[illegible] | The Jackson Laboratory | JAX: 017346 |
| Mouse: STOCK Tg(Efr3a-c[illegible] | MMRRC | MMRRC: 036660 |
| Mouse: STOCK Tg(Oxtr-cr[illegible] | MMRRC | MMRRC: 036545 |
| Mouse: STOCK Tg(Ppp1r1 | MMRRC | MMRRC: 036205 |
| Mouse: STOCK Tg(Ppp1r1 | MMRRC | MMRRC: 036205 |
| Mouse: STOCK Tg(Rbp4-c[illegible] | MMRRC | MMRRC: 031125 |
| Mouse: STOCK Tg(Dbh-cr[illegible] | MMRRC | MMRRC: 032081 |
| Mouse: STOCK Tg(Dbh-cr[illegible] | MMRRC | MMRRC: 032081 |
| Mouse: B6.FVB(Cg)-Tg(Nt[illegible] | MMRRC | MMRRC: 030648 |
| Mouse: B6.FVB(Cg)-Tg(Nt[illegible] | MMRRC | MMRRC: 030648 |
| Mouse: STOCK Tg(Slc18a2 | MMRRC | MMRRC: 034814 |
| Mouse: B6.Cg-Avptm1.1([illegible] | The Jackson Laboratory | JAX: 023530 |
| Mouse: STOCK Tg(Slc18a2 | MMRRC | MMRRC: 034814 |
| Mouse: STOCK Tg(Slc18a2 | MMRRC | MMRRC: 034814 |
| Mouse: STOCK Tg(Grm2-c | MMRRC | MMRRC: 034611 |
| Mouse: STOCK Tg(Grm2-c | MMRRC | MMRRC: 034611 |
| Mouse: STOCK Tg(Grm2-c | MMRRC | MMRRC: 034611 |
| Mouse: STOCK Tg(Grm2-c | MMRRC | MMRRC: 034611 |
| Mouse: B6;129S-Rorbtm1 | The Jackson Laboratory | JAX: 023526 |
| Mouse: B6;129S-Rorbtm1 | The Jackson Laboratory | JAX: 023526 |
| Mouse: B6;129S-Rorbtm1 | The Jackson Laboratory | JAX: 023526 |

| | | | |
|---|---|---|---|
| | Mouse: B6;129S-Rorbtm1 | The Jackson Laboratory | JAX: 023526 |
| | Mouse: Adcyap1-2A-Cre | Allen Institute for Brain S | N/A |
| | Mouse: Adcyap1-2A-Cre | Allen Institute for Brain S | N/A |
| | Mouse: Adcyap1-2A-Cre | Allen Institute for Brain S | N/A |
| | Mouse: STOCK Tg(Gpr26-c | MMRRC | MMRRC: 033032 |
| | Mouse: C57BL/6-Tg(Grik4 | The Jackson Laboratory | JAX: 006474 |
| | Mouse: C57BL/6-Tg(Grik4 | The Jackson Laboratory | JAX: 006474 |
| | Mouse: STOCK Slc17a6tm | The Jackson Laboratory | JAX: 016963 |
| | Mouse: STOCK Slc17a6tm | The Jackson Laboratory | JAX: 016963 |
| | Mouse: STOCK Tg(Slc18a2 | MMRRC | MMRRC: 034814 |
| | Mouse: STOCK Tg(Slc18a2 | MMRRC | MMRRC: 034814 |
| | Mouse: STOCK Tg(Ppp1r1 | MMRRC | MMRRC: 036205 |
| | Mouse: STOCK Ccktm1.1( | The Jackson Laboratory | JAX: 012706 |
| | Mouse: B6.Cg-Tg(A93003 | The Jackson Laboratory | JAX: 017346 |
| | Mouse: STOCK Tg(Slc18a2 | MMRRC | MMRRC: 034814 |
| Y | Mouse: B6(Cg)-Cux2tm1. | MMRRC | MMRRC: 031778 |
| | Mouse: B6(Cg)-Cux2tm1. | MMRRC | MMRRC: 031778 |
| | Mouse: B6.FVB(Cg)-Tg(Th | MMRRC | MMRRC: 031029 |
| | Mouse: STOCK Gad2tm2( | The Jackson Laboratory | JAX: 010802 |
| | Mouse: STOCK Gad2tm2( | The Jackson Laboratory | JAX: 010802 |
| | Mouse: STOCK Gad2tm2( | The Jackson Laboratory | JAX: 010802 |
| | Mouse: B6.Cg-Trib2tm1.1 | The Jackson Laboratory | JAX: 022865 |
| | Mouse: B6.Cg-Trib2tm1.1 | The Jackson Laboratory | JAX: 022865 |
| | Mouse: B6.Cg-Trib2tm1.1 | The Jackson Laboratory | JAX: 022865 |
| | Mouse: B6.Cg-Trib2tm1.1 | The Jackson Laboratory | JAX: 022865 |
| | Mouse: STOCK Tg(Syt6-cr | MMRRC | MMRRC: 032012 |
| | Mouse: STOCK Gad2tm2( | The Jackson Laboratory | JAX: 010802 |
| | Mouse: B6.Cg-Avptm1.1( | The Jackson Laboratory | JAX: 023530 |
| | Mouse: B6;129S-Fezf1tm | The Jackson Laboratory | JAX: 013048 |
| | Mouse: STOCK Nkx2-1tm | The Jackson Laboratory | JAX: 022861 |
| | Mouse: STOCK Nkx2-1tm | The Jackson Laboratory | JAX: 022861 |
| | Mouse: STOCK Nkx2-1tm | The Jackson Laboratory | JAX: 022861 |
| | Mouse: STOCK Nkx2-1tm | The Jackson Laboratory | JAX: 022861 |
| | Mouse: STOCK Tg(Htr2a-c | MMRRC | MMRRC: 031150 |
| | Mouse: STOCK Tg(Htr2a-c | MMRRC | MMRRC: 031150 |
| Y | Mouse: STOCK Tg(Rbp4-c | MMRRC | MMRRC: 031125 |
| | Mouse: B6;C3-Tg(Scnn1a- | The Jackson Laboratory | JAX: 009613 |
| | Mouse: B6;C3-Tg(Scnn1a- | The Jackson Laboratory | JAX: 009613 |
| | Mouse: STOCK Tg(Sim1-c | MMRRC | MMRRC: 031742 |
| | Mouse: STOCK Tg(Sim1-c | MMRRC | MMRRC: 031742 |
| | Mouse: STOCK Slc17a6tm | The Jackson Laboratory | JAX: 016963 |
| | Mouse: STOCK Gad2tm2( | The Jackson Laboratory | JAX: 010802 |
| | Mouse: STOCK Gad2tm2( | The Jackson Laboratory | JAX: 010802 |
| | Mouse: STOCK Tg(Grm2-c | MMRRC | MMRRC: 034611 |
| | Mouse: B6;129S-Rorbtm1 | The Jackson Laboratory | JAX: 023526 |
| | Mouse: B6(Cg)-Calb2tm1( | The Jackson Laboratory | JAX: 010774 |
| | Mouse: B6(Cg)-Calb2tm1( | The Jackson Laboratory | JAX: 010774 |

| | | |
|---|---|---|
| Mouse: B6(Cg)-Cux2tm1.1 | MMRRC | MMRRC: 031778 |
| Mouse: B6(Cg)-Cux2tm1.1 | MMRRC | MMRRC: 031778 |
| Mouse: B6(Cg)-Cux2tm1.1 | MMRRC | MMRRC: 031778 |
| Mouse: B6;129S4-Ntrk1tr | MMRRC | MMRRC: 015500 |
| Mouse: B6;129S4-Ntrk1tr | MMRRC | MMRRC: 015500 |
| Mouse: B6;129S4-Ntrk1tr | MMRRC | MMRRC: 015500 |
| Mouse: STOCK Tg(Efr3a-cı | MMRRC | MMRRC: 036660 |
| Mouse: STOCK Tg(Efr3a-cı | MMRRC | MMRRC: 036660 |
| Mouse: STOCK Tg(Efr3a-cı | MMRRC | MMRRC: 036660 |
| Mouse: STOCK Tg(Tlx3-cre | MMRRC | MMRRC: 041158 |
| Mouse: STOCK Tg(Tlx3-cre | MMRRC | MMRRC: 041158 |
| Mouse: STOCK Tg(Tlx3-cre | MMRRC | MMRRC: 041158 |
| Mouse: STOCK Ssttm2.1(c | The Jackson Laboratory | JAX: 013044 |
| Mouse: STOCK Ssttm2.1(c | The Jackson Laboratory | JAX: 013044 |
| Mouse: STOCK Ssttm2.1(c | The Jackson Laboratory | JAX: 013044 |
| Mouse: STOCK Ssttm2.1(c | The Jackson Laboratory | JAX: 013044 |
| Mouse: STOCK Ssttm2.1(c | The Jackson Laboratory | JAX: 013044 |
| Mouse: STOCK Ssttm2.1(c | The Jackson Laboratory | JAX: 013044 |
| Mouse: STOCK Tg(Syt17-c | MMRRC | MMRRC: 034355 |
| Mouse: STOCK Tg(Grp-cre | MMRRC | MMRRC: 031183 |
| Mouse: STOCK Tg(Grp-cre | MMRRC | MMRRC: 031183 |
| Mouse: STOCK Slc17a6tm | The Jackson Laboratory | JAX: 016963 |
| Mouse: STOCK Slc17a6tm | The Jackson Laboratory | JAX: 016963 |
| Mouse: STOCK Tg(Chrna2- | MMRRC | MMRRC: 036502 |
| Mouse: STOCK Tg(Chrna2- | MMRRC | MMRRC: 036502 |
| Mouse: STOCK Tg(Chrna2- | MMRRC | MMRRC: 036502 |
| Mouse: STOCK Tg(Chrna2- | MMRRC | MMRRC: 036502 |
| Mouse: STOCK Tg(Chrna2- | MMRRC | MMRRC: 036502 |
| Mouse: STOCK Tg(Chrna2- | MMRRC | MMRRC: 036502 |
| Mouse: STOCK Tg(Chrna2- | MMRRC | MMRRC: 036502 |
| Mouse: STOCK Tg(Chrna2- | MMRRC | MMRRC: 036502 |
| Mouse: STOCK Tg(Chrna2- | MMRRC | MMRRC: 036502 |
| Mouse: STOCK Tg(Chrna2- | MMRRC | MMRRC: 036502 |
| Mouse: STOCK Tg(Chrna2- | MMRRC | MMRRC: 036502 |
| Mouse: STOCK Tg(Chrna2- | MMRRC | MMRRC: 036502 |
| Mouse: B6;129S-Fezf1tm1 | The Jackson Laboratory | JAX: 013048 |
| Mouse: STOCK Gad2tm2( | The Jackson Laboratory | JAX: 010802 |
| Mouse: C57BL/6-Tg(Grik4 | The Jackson Laboratory | JAX: 006474 |
| Mouse: STOCK Tg(Efr3a-cı | MMRRC | MMRRC: 036660 |
| Mouse: STOCK Tg(Prkcd-g | MMRRC | MMRRC: 011559 |
| Mouse: STOCK Tg(Prkcd-g | MMRRC | MMRRC: 011559 |
| Mouse: STOCK Tg(Prkcd-g | MMRRC | MMRRC: 011559 |
| Mouse: STOCK Tg(Prkcd-g | MMRRC | MMRRC: 011559 |
| Mouse: STOCK Tg(Oxtr-cre | MMRRC | MMRRC: 036545 |
| Mouse: B6.Cg-Tg(A930038 | The Jackson Laboratory | JAX: 017346 |
| Mouse: B6.Cg-Tg(A930038 | The Jackson Laboratory | JAX: 017346 |
| Mouse: B6.Cg-Tg(A930038 | The Jackson Laboratory | JAX: 017346 |

| | | |
|---|---|---|
| Mouse: B6.Cg-Tg(A93003 | The Jackson Laboratory | JAX: 017346 |
| Mouse: STOCK Tg(Grp-cre | MMRRC | MMRRC: 031183 |
| Mouse: STOCK Tg(Grp-cre | MMRRC | MMRRC: 031183 |
| Mouse: STOCK Tg(Ppp1r1 | MMRRC | MMRRC: 036205 |
| Mouse: STOCK Tg(Ppp1r1 | MMRRC | MMRRC: 036205 |
| Mouse: B6.Cg-Tg(Wfs1-cr | The Jackson Laboratory | JAX: 009614 |
| Mouse: STOCK Tg(Grm2-c | MMRRC | MMRRC: 034611 |
| Mouse: STOCK Tg(Ppp1r1 | MMRRC | MMRRC: 036205 |
| Mouse: STOCK Tg(Ppp1r1 | MMRRC | MMRRC: 036205 |
| Mouse: STOCK Tg(Ppp1r1 | MMRRC | MMRRC: 036205 |
| Mouse: B6.Cg-Trib2tm1.1 | The Jackson Laboratory | JAX: 022865 |
| Mouse: B6(Cg)-Cux2tm3. | MMRRC | MMRRC: 032779 |
| Mouse: B6(Cg)-Crhtm1(cr | The Jackson Laboratory | JAX: 012704 |
| Mouse: STOCK Tg(Grm2-c | MMRRC | MMRRC: 034611 |
| Mouse: STOCK Tg(Grm2-c | MMRRC | MMRRC: 034611 |
| Mouse: STOCK Tg(Grm2-c | MMRRC | MMRRC: 034611 |
| Mouse: STOCK Tg(Syt6-cr | MMRRC | MMRRC: 032012 |
| Mouse: B6.Cg-Tg(A93003 | The Jackson Laboratory | JAX: 017346 |
| Mouse: B6.Cg-Tg(A93003 | The Jackson Laboratory | JAX: 017346 |
| Mouse: B6.Cg-Tg(A93003 | The Jackson Laboratory | JAX: 017346 |
| Mouse: STOCK Tg(Slc18a2 | MMRRC | MMRRC: 034814 |
| Mouse: STOCK Tg(Slc18a2 | MMRRC | MMRRC: 034814 |
| Mouse: B6;129S6-Chattm | The Jackson Laboratory | JAX: 006410 |
| Mouse: B6;129S4-Ntrk1tr | MMRRC | MMRRC: 015500 |
| Mouse: B6;129S4-Ntrk1tr | MMRRC | MMRRC: 015500 |
| Mouse: B6;129S4-Ntrk1tr | MMRRC | MMRRC: 015500 |
| Mouse: B6;129S4-Ntrk1tr | MMRRC | MMRRC: 015500 |
| Mouse: B6;129S4-Ntrk1tr | MMRRC | MMRRC: 015500 |
| Mouse: STOCK Tg(Syt17-c | MMRRC | MMRRC: 034355 |
| Mouse: STOCK Tg(Syt17-c | MMRRC | MMRRC: 034355 |
| Mouse: STOCK Tg(Syt17-c | MMRRC | MMRRC: 034355 |
| Mouse: STOCK Tg(Syt17-c | MMRRC | MMRRC: 034355 |
| Mouse: Crh-IRES-Cre_BL | Bradford Lowell | N/A |
| Mouse: STOCK Tg(Syt17-c | MMRRC | MMRRC: 034355 |
| Mouse: B6;129S-Nos1tm | The Jackson Laboratory | JAX: 014541 |
| Mouse: B6;129P2-Pvalbtr | The Jackson Laboratory | JAX: 008069 |
| Mouse: B6;129S-Tac2tm1 | The Jackson Laboratory | JAX: 021878 |
| Mouse: STOCK Tg(Htr2a-c | MMRRC | MMRRC: 031150 |
| Mouse: STOCK Tg(Plxnd1- | MMRRC | MMRRC: 036631 |
| Mouse: STOCK Tg(Plxnd1- | MMRRC | MMRRC: 036631 |
| Mouse: STOCK Tg(Plxnd1- | MMRRC | MMRRC: 036631 |
| Mouse: STOCK Tg(Plxnd1- | MMRRC | MMRRC: 036631 |
| Mouse: STOCK Tg(Plxnd1- | MMRRC | MMRRC: 036631 |
| Mouse: STOCK Viptm1(cr | The Jackson Laboratory | JAX: 010908 |
| Mouse: STOCK Tg(Tlx3-cre | MMRRC | MMRRC: 041158 |
| Mouse: STOCK Tg(Tlx3-cre | MMRRC | MMRRC: 041158 |
| Mouse: STOCK Tg(Tlx3-cre | MMRRC | MMRRC: 041158 |

| | | | |
|---|---|---|---|
| | Mouse: STOCK Tg(Htr2a-c | MMRRC | MMRRC: 031150 |
| | Mouse: STOCK Tg(Htr2a-c | MMRRC | MMRRC: 031150 |
| Y | Mouse: B6(Cg)-Cux2tm1. | MMRRC | MMRRC: 031778 |
| | Mouse: B6(Cg)-Cux2tm1. | MMRRC | MMRRC: 031778 |
| | Mouse: STOCK Slc17a6tm | The Jackson Laboratory | JAX: 016963 |
| | Mouse: FVB-Tg(Nr5a1-cre | The Jackson Laboratory | JAX: 006364 |
| | Mouse: STOCK Gad2tm2( | The Jackson Laboratory | JAX: 010802 |
| | Mouse: STOCK Gad2tm2( | The Jackson Laboratory | JAX: 010802 |
| | Mouse: B6;129S-Rorbtm1 | The Jackson Laboratory | JAX: 023526 |
| | Mouse: STOCK Tg(Drd3-cr | MMRRC | MMRRC: 034610 |
| | Mouse: STOCK Tg(Drd3-cr | MMRRC | MMRRC: 034610 |
| | Mouse: B6.Cg-Calb1tm1. | The Jackson Laboratory | JAX: 023531 |
| | Mouse: Crh-IRES-Cre_BL | Bradford Lowell | N/A |
| | Mouse: B6;129S-Slc17a7t | The Jackson Laboratory | JAX: 023527 |
| | Mouse: B6;129S-Slc17a7t | The Jackson Laboratory | JAX: 023527 |
| | Mouse: B6;129S-Slc17a7t | The Jackson Laboratory | JAX: 023527 |
| | Mouse: STOCK Tg(Sim1-cr | MMRRC | MMRRC: 031742 |
| | Mouse: STOCK Tg(Sim1-cr | MMRRC | MMRRC: 031742 |
| | Mouse: B6;129S-Nos1tm | The Jackson Laboratory | JAX: 014541 |
| | Mouse: B6;129S-Nos1tm | The Jackson Laboratory | JAX: 014541 |
| | Mouse: STOCK Tg(Chrna2 | MMRRC | MMRRC: 036502 |
| | Mouse: STOCK Slc17a6tm | The Jackson Laboratory | JAX: 016963 |
| | Mouse: STOCK Slc17a6tm | The Jackson Laboratory | JAX: 016963 |
| | Mouse: STOCK Slc17a6tm | The Jackson Laboratory | JAX: 016963 |
| | Mouse: STOCK Gad2tm2( | The Jackson Laboratory | JAX: 010802 |
| | Mouse: STOCK Gad2tm2( | The Jackson Laboratory | JAX: 010802 |
| | Mouse: STOCK Gad2tm2( | The Jackson Laboratory | JAX: 010802 |
| | Mouse: STOCK Slc17a6tm | The Jackson Laboratory | JAX: 016963 |
| | Mouse: STOCK Slc17a6tm | The Jackson Laboratory | JAX: 016963 |
| | Mouse: STOCK Slc17a6tm | The Jackson Laboratory | JAX: 016963 |
| | Mouse: STOCK Slc17a6tm | The Jackson Laboratory | JAX: 016963 |
| | Mouse: STOCK Slc17a6tm | The Jackson Laboratory | JAX: 016963 |
| | Mouse: STOCK Tg(Efr3a-c | MMRRC | MMRRC: 036660 |
| | Mouse: STOCK Tg(Oxtr-cr | MMRRC | MMRRC: 036545 |
| | Mouse: STOCK Tg(Oxtr-cr | MMRRC | MMRRC: 036545 |
| | Mouse: B6;129S-Tac1tm1 | The Jackson Laboratory | JAX: 021877 |
| | Mouse: B6;129S-Tac1tm1 | The Jackson Laboratory | JAX: 021877 |
| | Mouse: B6;129S-Tac1tm1 | The Jackson Laboratory | JAX: 021877 |
| | Mouse: B6;129S-Tac1tm1 | The Jackson Laboratory | JAX: 021877 |
| | Mouse: B6;129S-Tac1tm1 | The Jackson Laboratory | JAX: 021877 |
| | Mouse: STOCK Slc17a6tm | The Jackson Laboratory | JAX: 016963 |
| | Mouse: B6(Cg)-Cux2tm1. | MMRRC | MMRRC: 031778 |
| | Mouse: B6(Cg)-Cux2tm1. | MMRRC | MMRRC: 031778 |
| | Mouse: B6(Cg)-Cux2tm1. | MMRRC | MMRRC: 031778 |
| | Mouse: STOCK Tg(Oxtr-cr | MMRRC | MMRRC: 036545 |
| Y | Mouse: B6(Cg)-Cux2tm1. | MMRRC | MMRRC: 031778 |
| | Mouse: B6(Cg)-Cux2tm1. | MMRRC | MMRRC: 031778 |

| | | | |
|---|---|---|---|
| | Mouse: B6(Cg)-Cux2tm1.: | MMRRC | MMRRC: 031778 |
| | Mouse: STOCK Tg(Rbp4-cı | MMRRC | MMRRC: 031125 |
| | Mouse: STOCK Tg(Tlx3-cre | MMRRC | MMRRC: 041158 |
| | Mouse: STOCK Tg(Tlx3-cre | MMRRC | MMRRC: 041158 |
| | Mouse: STOCK Tg(Tlx3-cre | MMRRC | MMRRC: 041158 |
| | Mouse: STOCK Tg(Tlx3-cre | MMRRC | MMRRC: 041158 |
| | Mouse: STOCK Tg(Tlx3-cre | MMRRC | MMRRC: 041158 |
| | Mouse: STOCK Tg(Chrna2 | MMRRC | MMRRC: 036502 |
| Y | Mouse: B6(Cg)-Cux2tm1.: | MMRRC | MMRRC: 031778 |
| Y | Mouse: STOCK Tg(Rbp4-cı | MMRRC | MMRRC: 031125 |
| | Mouse: STOCK Tg(Tlx3-cre | MMRRC | MMRRC: 041158 |
| | Mouse: STOCK Tg(Tlx3-cre | MMRRC | MMRRC: 041158 |
| | Mouse: STOCK Tg(Tlx3-cre | MMRRC | MMRRC: 041158 |
| | Mouse: STOCK Tg(Tlx3-cre | MMRRC | MMRRC: 041158 |
| | Mouse: STOCK Tg(Tlx3-cre | MMRRC | MMRRC: 041158 |
| | Mouse: STOCK Tg(Tlx3-cre | MMRRC | MMRRC: 041158 |
| | Mouse: STOCK Tg(Tlx3-cre | MMRRC | MMRRC: 041158 |
| | Mouse: STOCK Tg(Tlx3-cre | MMRRC | MMRRC: 041158 |
| | Mouse: B6.FVB(Cg)-Tg(Nt | MMRRC | MMRRC: 030648 |
| | Mouse: B6.FVB(Cg)-Tg(Nt | MMRRC | MMRRC: 030648 |
| | Mouse: B6.FVB(Cg)-Tg(Nt | MMRRC | MMRRC: 030648 |
| | Mouse: STOCK Tg(Rbp4-cı | MMRRC | MMRRC: 031125 |
| | Mouse: STOCK Tg(Tlx3-cre | MMRRC | MMRRC: 041158 |
| | Mouse: STOCK Tg(Plxnd1- | MMRRC | MMRRC: 036631 |
| | Mouse: STOCK Tg(Plxnd1- | MMRRC | MMRRC: 036631 |
| | Mouse: STOCK Tg(Drd3-cr | MMRRC | MMRRC: 034610 |
| | Mouse: STOCK Tg(Sim1-cr | MMRRC | MMRRC: 031742 |
| | Mouse: STOCK Tg(Sim1-cr | MMRRC | MMRRC: 031742 |
| | Mouse: STOCK Tg(Sim1-cr | MMRRC | MMRRC: 031742 |
| | Mouse: STOCK Tg(Sim1-cr | MMRRC | MMRRC: 031742 |
| | Mouse: STOCK Tg(Sim1-cr | MMRRC | MMRRC: 031742 |
| | Mouse: STOCK Tg(Sim1-cr | MMRRC | MMRRC: 031742 |
| | Mouse: B6(Cg)-Cux2tm1.: | MMRRC | MMRRC: 031778 |
| | Mouse: STOCK Tg(Chrna2 | MMRRC | MMRRC: 036502 |
| | Mouse: STOCK Tg(Tlx3-cre | MMRRC | MMRRC: 041158 |
| | Mouse: STOCK Tg(Chrna2 | MMRRC | MMRRC: 036502 |
| | Mouse: STOCK Tg(Chrna2 | MMRRC | MMRRC: 036502 |
| | Mouse: STOCK Tg(Chrna2 | MMRRC | MMRRC: 036502 |
| | Mouse: STOCK Tg(Chrna2 | MMRRC | MMRRC: 036502 |
| | Mouse: STOCK Tg(Sim1-cr | MMRRC | MMRRC: 031742 |
| | Mouse: STOCK Tg(Sim1-cr | MMRRC | MMRRC: 031742 |
| | Mouse: STOCK Tg(Sim1-cr | MMRRC | MMRRC: 031742 |
| | Mouse: STOCK Tg(Sim1-cr | MMRRC | MMRRC: 031742 |
| | Mouse: STOCK Tg(Sim1-cr | MMRRC | MMRRC: 031742 |
| | Mouse: STOCK Tg(Chrnb4 | MMRRC | MMRRC: 036203 |
| | Mouse: STOCK Tg(Chrnb4 | MMRRC | MMRRC: 036203 |
| | Mouse: Lypd6-Cre_KL156 | Nathaniel Heintz and Cha | N/A |

| | | | |
|---|---|---|---|
| | Mouse: STOCK Tg(Htr2a-c | MMRRC | MMRRC: 031150 |
| | Mouse: STOCK Tg(Htr2a-c | MMRRC | MMRRC: 031150 |
| | Mouse: Hcrt-Cre | Takeshi Sakurai | N/A |
| | Mouse: Hcrt-Cre | Takeshi Sakurai | N/A |
| | Mouse: Lypd6-Cre_KL156 | Nathaniel Heintz and Cha | N/A |
| | Mouse: Lypd6-Cre_KL156 | Nathaniel Heintz and Cha | N/A |
| | Mouse: Lypd6-Cre_KL156 | Nathaniel Heintz and Cha | N/A |
| | Mouse: Lypd6-Cre_KL156 | Nathaniel Heintz and Cha | N/A |
| | Mouse: Lypd6-Cre_KL156 | Nathaniel Heintz and Cha | N/A |
| | Mouse: Lypd6-Cre_KL156 | Nathaniel Heintz and Cha | N/A |
| | Mouse: B6.Cg-Tg(Ins2-cre | The Jackson Laboratory | JAX: 003573 |
| | Mouse: B6(Cg)-Cux2tm1. | MMRRC | MMRRC: 031778 |
| Y | Mouse: STOCK Tg(Rbp4-cr | MMRRC | MMRRC: 031125 |
| | Mouse: B6.Cg-Tg(Ins2-cre | The Jackson Laboratory | JAX: 003573 |
| | Mouse: STOCK Tg(Rbp4-cr | MMRRC | MMRRC: 031125 |
| | Mouse: STOCK Tg(Tlx3-cre | MMRRC | MMRRC: 041158 |
| | Mouse: STOCK Tg(Efr3a-cr | MMRRC | MMRRC: 036660 |
| | Mouse: B6.Cg-Tg(A930038 | The Jackson Laboratory | JAX: 017346 |
| | Mouse: STOCK Tg(Chrna2- | MMRRC | MMRRC: 036502 |
| | Mouse: STOCK Tg(Chrna2- | MMRRC | MMRRC: 036502 |
| | Mouse: STOCK Tg(Chrna2- | MMRRC | MMRRC: 036502 |
| | Mouse: STOCK Tg(Chrna2- | MMRRC | MMRRC: 036502 |
| | Mouse: B6.Cg-Tg(A930038 | The Jackson Laboratory | JAX: 017346 |
| | Mouse: STOCK Tg(Chrna2- | MMRRC | MMRRC: 036502 |
| | Mouse: B6.Cg-Tg(A930038 | The Jackson Laboratory | JAX: 017346 |
| | Mouse: STOCK Tg(Efr3a-cr | MMRRC | MMRRC: 036660 |
| | Mouse: B6.Cg-Tg(A930038 | The Jackson Laboratory | JAX: 017346 |
| | Mouse: B6.Cg-Tg(A930038 | The Jackson Laboratory | JAX: 017346 |
| | Mouse: STOCK Tg(Efr3a-cr | MMRRC | MMRRC: 036660 |
| | Mouse: STOCK Tg(Gpr26-c | MMRRC | MMRRC: 033032 |
| | Mouse: STOCK Tg(Gpr26-c | MMRRC | MMRRC: 033032 |
| | Mouse: B6;129S-Rasgrf2t | The Jackson Laboratory | JAX: 022864 |
| | Mouse: B6;129S-Rasgrf2t | The Jackson Laboratory | JAX: 022864 |
| | Mouse: B6;129S-Rasgrf2t | The Jackson Laboratory | JAX: 022864 |
| | Mouse: STOCK Tg(Gpr26-c | MMRRC | MMRRC: 033032 |
| | Mouse: STOCK Tg(Rbp4-cr | MMRRC | MMRRC: 031125 |
| | Mouse: B6;129S-Rorbtm1 | The Jackson Laboratory | JAX: 023526 |
| | Mouse: B6;129S-Rasgrf2t | The Jackson Laboratory | JAX: 022864 |
| | Mouse: STOCK Tg(Drd3-cr | MMRRC | MMRRC: 034610 |
| | Mouse: STOCK Tg(Sim1-cr | MMRRC | MMRRC: 031742 |
| | Mouse: B6.Cg-Tg(A930038 | The Jackson Laboratory | JAX: 017346 |
| | Mouse: B6.Cg-Tg(A930038 | The Jackson Laboratory | JAX: 017346 |
| | Mouse: B6.Cg-Tg(A930038 | The Jackson Laboratory | JAX: 017346 |
| | Mouse: B6.Cg-Tg(A930038 | The Jackson Laboratory | JAX: 017346 |
| | Mouse: B6(Cg)-Cux2tm1. | MMRRC | MMRRC: 031778 |
| | Mouse: B6.FVB(Cg)-Tg(Nt | MMRRC | MMRRC: 030648 |
| | Mouse: B6.Cg-Tg(A930038 | The Jackson Laboratory | JAX: 017346 |

| | | |
|---|---|---|
| Mouse: B6(Cg)-Calb2tm1( | The Jackson Laboratory | JAX: 010774 |
| Mouse: STOCK Tg(Cartpt- | The Jackson Laboratory | JAX: 009615 |
| Mouse: STOCK Tg(Efr3a-c | MMRRC | MMRRC: 036660 |
| Mouse: STOCK Tg(Drd3-cr | MMRRC | MMRRC: 034610 |
| Mouse: STOCK Gad2tm2( | The Jackson Laboratory | JAX: 010802 |
| Mouse: STOCK Tg(Kiss1-c | The Jackson Laboratory | JAX: 023426 |
| Mouse: B6;129S-Nos1tm1 | The Jackson Laboratory | JAX: 014541 |
| Mouse: B6;129S4-Ntrk1tr | MMRRC | MMRRC: 015500 |
| Mouse: STOCK Tg(Oxtr-cr | MMRRC | MMRRC: 036545 |
| Mouse: B6;C3-Tg(Scnn1a- | The Jackson Laboratory | JAX: 009112 |
| Mouse: B6;C3-Tg(Scnn1a- | The Jackson Laboratory | JAX: 009613 |
| Mouse: STOCK Tg(Oxtr-cr | MMRRC | MMRRC: 036545 |
| Mouse: STOCK Slc32a1tm | The Jackson Laboratory | JAX: 016962 |
| Mouse: STOCK Ssttm2.1( | The Jackson Laboratory | JAX: 013044 |
| Mouse: STOCK Tg(Syt17-c | MMRRC | MMRRC: 034355 |
| Mouse: STOCK Tg(Syt17-c | MMRRC | MMRRC: 034355 |
| Mouse: B6;129S-Tac1tm1 | The Jackson Laboratory | JAX: 021877 |
| Mouse: STOCK Viptm1(cr | The Jackson Laboratory | JAX: 010908 |
| Mouse: STOCK Tg(Efr3a-c | MMRRC | MMRRC: 036660 |
| Mouse: STOCK Tg(Efr3a-c | MMRRC | MMRRC: 036660 |
| Mouse: STOCK Tg(Syt6-cr | MMRRC | MMRRC: 032012 |
| Mouse: STOCK Tg(Efr3a-c | MMRRC | MMRRC: 036660 |
| Mouse: STOCK Tg(Efr3a-c | MMRRC | MMRRC: 036660 |
| Mouse: Lypd6-Cre_KL156 | Nathaniel Heintz and Ch | N/A |
| Mouse: B6;129S-Nos1tm1 | The Jackson Laboratory | JAX: 014541 |
| Mouse: B6;129S-Nos1tm1 | The Jackson Laboratory | JAX: 014541 |
| Mouse: STOCK Tg(Chrna2 | MMRRC | MMRRC: 036502 |
| Mouse: STOCK Tg(Chrna2 | MMRRC | MMRRC: 036502 |
| Mouse: B6(Cg)-Cux2tm1.1 | MMRRC | MMRRC: 031778 |
| Mouse: STOCK Tg(Sim1-cr | MMRRC | MMRRC: 031742 |
| Mouse: STOCK Tg(Syt6-cr | MMRRC | MMRRC: 032012 |
| Mouse: STOCK Tg(Syt6-cr | MMRRC | MMRRC: 032012 |
| Mouse: STOCK Tg(Syt6-cr | MMRRC | MMRRC: 032012 |
| Mouse: STOCK Tg(Syt6-cr | MMRRC | MMRRC: 032012 |
| Mouse: STOCK Ccktm1.1( | The Jackson Laboratory | JAX: 012706 |
| Mouse: STOCK Tg(Chrna2 | MMRRC | MMRRC: 036502 |
| Mouse: Lypd6-Cre_KL156 | Nathaniel Heintz and Ch | N/A |
| Mouse: STOCK Tg(Plxnd1- | MMRRC | MMRRC: 036631 |
| Mouse: B6;129P2-Pvalbtr | The Jackson Laboratory | JAX: 008069 |
| Mouse: B6;129P2-Pvalbtr | The Jackson Laboratory | JAX: 008069 |
| Mouse: STOCK Tg(Sim1-cr | MMRRC | MMRRC: 031742 |
| Mouse: STOCK Viptm1(cr | The Jackson Laboratory | JAX: 010908 |
| Mouse: B6.Cg-Tg(A93003 | The Jackson Laboratory | JAX: 017346 |
| Mouse: B6(Cg)-Calb2tm1( | The Jackson Laboratory | JAX: 010774 |
| Mouse: STOCK Tg(Dbh-cr | MMRRC | MMRRC: 032081 |
| Mouse: STOCK Tg(Drd3-cr | MMRRC | MMRRC: 034610 |
| Mouse: STOCK Tg(Drd3-cr | MMRRC | MMRRC: 034610 |

| | | |
|---|---|---|
| Mouse: STOCK Tg(Drd3-cr | MMRRC | MMRRC: 034610 |
| Mouse: STOCK Tg(Drd3-cr | MMRRC | MMRRC: 034610 |
| Mouse: STOCK Tg(Drd3-cr | MMRRC | MMRRC: 034610 |
| Mouse: STOCK Tg(Efr3a-cı | MMRRC | MMRRC: 036660 |
| Mouse: STOCK Tg(Gabrr3 | MMRRC | MMRRC: 030709 |
| Mouse: STOCK Gad2tm2( | The Jackson Laboratory | JAX: 010802 |
| Mouse: STOCK Gad2tm2( | The Jackson Laboratory | JAX: 010802 |
| Mouse: STOCK Tg(Gal-cre | MMRRC | MMRRC: 031060 |
| Mouse: STOCK Tg(Grm2-c | MMRRC | MMRRC: 034611 |
| Mouse: STOCK Tg(Grm2-c | MMRRC | MMRRC: 034611 |
| Mouse: STOCK Tg(Htr2a-c | MMRRC | MMRRC: 031150 |
| Mouse: C57BL/6-Tg(Grik4 | The Jackson Laboratory | JAX: 006474 |
| Mouse: STOCK Tg(Htr2a-c | MMRRC | MMRRC: 031150 |
| Mouse: STOCK Tg(Htr2a-c | MMRRC | MMRRC: 031150 |
| Mouse: B6;129S-Nos1tm: | The Jackson Laboratory | JAX: 014541 |
| Mouse: STOCK Tg(Oxtr-cr | MMRRC | MMRRC: 036545 |
| Mouse: STOCK Tg(Rbp4-cı | MMRRC | MMRRC: 031125 |
| Mouse: STOCK Tg(Rbp4-cı | MMRRC | MMRRC: 031125 |
| Mouse: STOCK Tg(Rbp4-cı | MMRRC | MMRRC: 031125 |
| Mouse: STOCK Slc17a6tm | The Jackson Laboratory | JAX: 016963 |
| Mouse: STOCK Slc17a6tm | The Jackson Laboratory | JAX: 016963 |
| Mouse: STOCK Tg(Slc18a2 | MMRRC | MMRRC: 034814 |
| Mouse: STOCK Tg(Slc18a2 | MMRRC | MMRRC: 034814 |
| Mouse: STOCK Tg(Slc6a5- | MMRRC | MMRRC: 030730 |
| Mouse: STOCK Ssttm2.1(c | The Jackson Laboratory | JAX: 013044 |
| Mouse: STOCK Ssttm2.1(c | The Jackson Laboratory | JAX: 013044 |
| Mouse: STOCK Ssttm2.1(c | The Jackson Laboratory | JAX: 013044 |
| Mouse: B6;129S-Tac1tm1 | The Jackson Laboratory | JAX: 021877 |
| Mouse: B6;129S-Tac1tm1 | The Jackson Laboratory | JAX: 021877 |
| Mouse: B6;129S-Tac1tm1 | The Jackson Laboratory | JAX: 021877 |
| Mouse: B6;129S-Tac1tm1 | The Jackson Laboratory | JAX: 021877 |
| Mouse: B6;129S-Tac2tm1 | The Jackson Laboratory | JAX: 021878 |
| Mouse: B6;129S-Rorbtm1 | The Jackson Laboratory | JAX: 023526 |
| Mouse: B6.FVB(Cg)-Tg(Th | MMRRC | MMRRC: 031029 |
| Mouse: Adcyap1-2A-Cre | Allen Institute for Brain S | N/A |
| Mouse: B6(Cg)-Calb2tm1( | The Jackson Laboratory | JAX: 010774 |
| Mouse: B6(Cg)-Calb2tm1( | The Jackson Laboratory | JAX: 010774 |
| Mouse: B6(Cg)-Calb2tm1( | The Jackson Laboratory | JAX: 010774 |
| Mouse: B6(Cg)-Calb2tm1( | The Jackson Laboratory | JAX: 010774 |
| Mouse: B6(Cg)-Calb2tm1( | The Jackson Laboratory | JAX: 010774 |
| Mouse: STOCK Tg(Drd3-cr | MMRRC | MMRRC: 031741 |
| Mouse: STOCK Tg(Drd3-cr | MMRRC | MMRRC: 031741 |
| Mouse: STOCK Tg(Drd3-cr | MMRRC | MMRRC: 031741 |
| Mouse: STOCK Tg(Efr3a-cı | MMRRC | MMRRC: 036660 |
| Mouse: STOCK Tg(Efr3a-cı | MMRRC | MMRRC: 036660 |
| Mouse: STOCK Tg(Efr3a-cı | MMRRC | MMRRC: 036660 |
| Mouse: STOCK Tg(Efr3a-cı | MMRRC | MMRRC: 036660 |

| | | |
|---|---|---|
| Mouse: STOCK Tg(Gal-cre | MMRRC | MMRRC: 031060 |
| Mouse: STOCK Tg(Gpr26- | MMRRC | MMRRC: 033032 |
| Mouse: STOCK Tg(Gpr26- | MMRRC | MMRRC: 033032 |
| Mouse: STOCK Tg(Gpr26- | MMRRC | MMRRC: 033032 |
| Mouse: STOCK Tg(Gpr26- | MMRRC | MMRRC: 033032 |
| Mouse: STOCK Tg(Grm2-c | MMRRC | MMRRC: 034611 |
| Mouse: STOCK Tg(Grp-cre | MMRRC | MMRRC: 031183 |
| Mouse: B6;129S-Nos1tm | The Jackson Laboratory | JAX: 014541 |
| Mouse: B6;129S-Nos1tm | The Jackson Laboratory | JAX: 014541 |
| Mouse: B6;129S-Nos1tm | The Jackson Laboratory | JAX: 014541 |
| Mouse: STOCK Tg(Oxtr-cr | MMRRC | MMRRC: 036545 |
| Mouse: STOCK Tg(Oxtr-cr | MMRRC | MMRRC: 036545 |
| Mouse: STOCK Tg(Oxtr-cr | MMRRC | MMRRC: 036545 |
| Mouse: STOCK Tg(Plxnd1- | MMRRC | MMRRC: 036631 |
| Mouse: STOCK Tg(Plxnd1- | MMRRC | MMRRC: 036631 |
| Mouse: STOCK Tg(Plxnd1- | MMRRC | MMRRC: 036631 |
| Mouse: B6;129P2-Pvalbtr | The Jackson Laboratory | JAX: 008069 |
| Mouse: STOCK Tg(Rbp4-c | MMRRC | MMRRC: 031125 |
| Mouse: STOCK Tg(Rbp4-c | MMRRC | MMRRC: 031125 |
| Mouse: B6;129S-Rorbtm1 | The Jackson Laboratory | JAX: 023526 |
| Mouse: B6;129S-Rorbtm1 | The Jackson Laboratory | JAX: 023526 |
| Mouse: B6;129S-Rorbtm1 | The Jackson Laboratory | JAX: 023526 |
| Mouse: B6;129S-Rorbtm1 | The Jackson Laboratory | JAX: 023526 |
| Mouse: B6;129S-Rorbtm1 | The Jackson Laboratory | JAX: 023526 |
| Mouse: B6;129S-Rorbtm1 | The Jackson Laboratory | JAX: 023526 |
| Mouse: B6;129S-Slc17a7t | The Jackson Laboratory | JAX: 023527 |
| Mouse: STOCK Tg(Slc18a2 | MMRRC | MMRRC: 034814 |
| Mouse: STOCK Tg(Slc18a2 | MMRRC | MMRRC: 034814 |
| Mouse: STOCK Tg(Slc18a2 | MMRRC | MMRRC: 034814 |
| Mouse: STOCK Tg(Slc18a2 | MMRRC | MMRRC: 034814 |
| Mouse: STOCK Tg(Slc18a2 | MMRRC | MMRRC: 034814 |
| Mouse: STOCK Tg(Slc18a2 | MMRRC | MMRRC: 034814 |
| Mouse: STOCK Tg(Slc18a2 | MMRRC | MMRRC: 034814 |
| Mouse: STOCK Tg(Slc18a2 | MMRRC | MMRRC: 034814 |
| Mouse: STOCK Tg(Slc18a2 | MMRRC | MMRRC: 034814 |
| Mouse: B6(Cg)-Calb2tm1 | The Jackson Laboratory | JAX: 010774 |
| Mouse: B6;129S6-Chattm | The Jackson Laboratory | JAX: 006410 |
| Mouse: B6(Cg)-Crhtm1(cr | The Jackson Laboratory | JAX: 012704 |
| Mouse: STOCK Tg(Gabrr3 | MMRRC | MMRRC: 030709 |
| Mouse: C57BL/6-Tg(Grik4 | The Jackson Laboratory | JAX: 006474 |
| Mouse: STOCK Tg(Htr2a-c | MMRRC | MMRRC: 031150 |
| Mouse: STOCK Tg(Htr2a-c | MMRRC | MMRRC: 031150 |
| Mouse: STOCK Tg(Kiss1-c | The Jackson Laboratory | JAX: 023426 |
| Mouse: STOCK Nkx2-1tm | The Jackson Laboratory | JAX: 022861 |
| Mouse: STOCK Tg(Plxnd1- | MMRRC | MMRRC: 036631 |
| Mouse: B6;C3-Tg(Scnn1a- | The Jackson Laboratory | JAX: 009112 |
| Mouse: STOCK Slc17a6tm | The Jackson Laboratory | JAX: 016963 |

| | | |
|---|---|---|
| Mouse: STOCK Gad2tm2( | The Jackson Laboratory | JAX: 010802 |
| Mouse: STOCK Gad2tm2( | The Jackson Laboratory | JAX: 010802 |
| Mouse: STOCK Gad2tm2( | The Jackson Laboratory | JAX: 010802 |
| Mouse: STOCK Gad2tm2( | The Jackson Laboratory | JAX: 010802 |
| Mouse: STOCK Slc17a6tm | The Jackson Laboratory | JAX: 016963 |
| Mouse: STOCK Slc17a6tm | The Jackson Laboratory | JAX: 016963 |
| Mouse: STOCK Gad2tm2( | The Jackson Laboratory | JAX: 010802 |
| Mouse: STOCK Slc17a6tm | The Jackson Laboratory | JAX: 016963 |
| Mouse: STOCK Gad2tm2( | The Jackson Laboratory | JAX: 010802 |
| Mouse: STOCK Gad2tm2( | The Jackson Laboratory | JAX: 010802 |
| Mouse: STOCK Gad2tm2( | The Jackson Laboratory | JAX: 010802 |
| Mouse: STOCK Gad2tm2( | The Jackson Laboratory | JAX: 010802 |
| Mouse: STOCK Gad2tm2( | The Jackson Laboratory | JAX: 010802 |
| Mouse: STOCK Slc17a6tm | The Jackson Laboratory | JAX: 016963 |
| Mouse: STOCK Gad2tm2( | The Jackson Laboratory | JAX: 010802 |
| Mouse: STOCK Slc32a1tm | The Jackson Laboratory | JAX: 016962 |
| Mouse: STOCK Gad2tm2( | The Jackson Laboratory | JAX: 010802 |
| Mouse: STOCK Gad2tm2( | The Jackson Laboratory | JAX: 010802 |
| Mouse: STOCK Gad2tm2( | The Jackson Laboratory | JAX: 010802 |
| Mouse: STOCK Gad2tm2( | The Jackson Laboratory | JAX: 010802 |
| Mouse: STOCK Gad2tm2( | The Jackson Laboratory | JAX: 010802 |
| Mouse: STOCK Gad2tm2( | The Jackson Laboratory | JAX: 010802 |
| Mouse: STOCK Gad2tm2( | The Jackson Laboratory | JAX: 010802 |
| Mouse: STOCK Slc17a6tm | The Jackson Laboratory | JAX: 016963 |
| Mouse: STOCK Gad2tm2( | The Jackson Laboratory | JAX: 010802 |
| Mouse: STOCK Tg(Tlx3-cr | MMRRC | MMRRC: 041158 |
| Mouse: STOCK Tg(Tlx3-cr | MMRRC | MMRRC: 041158 |
| Mouse: STOCK Tg(Tlx3-cr | MMRRC | MMRRC: 041158 |
| Mouse: STOCK Tg(Tlx3-cr | MMRRC | MMRRC: 041158 |
| Mouse: STOCK Tg(Tlx3-cr | MMRRC | MMRRC: 041158 |
| Mouse: B6.FVB(Cg)-Tg(Nt | MMRRC | MMRRC: 030648 |
| Mouse: Adcyap1-2A-Cre | Allen Institute for Brain | N/A |
| Mouse: STOCK Tg(Chrna2 | MMRRC | MMRRC: 036502 |
| Mouse: Lypd6-Cre_KL156 | Nathaniel Heintz and Ch | N/A |
| Mouse: B6;129S-Rasgrf2t | The Jackson Laboratory | JAX: 022864 |
| Mouse: B6;129S-Rasgrf2t | The Jackson Laboratory | JAX: 022864 |
| Mouse: B6;129S-Rasgrf2t | The Jackson Laboratory | JAX: 022864 |
| Mouse: STOCK Tg(Rbp4-c | MMRRC | MMRRC: 031125 |
| Mouse: STOCK Ssttm2.1( | The Jackson Laboratory | JAX: 013044 |
| Mouse: B6.FVB(Cg)-Tg(Th | MMRRC | MMRRC: 031029 |
| Mouse: B6;129S-Rorbtm1 | The Jackson Laboratory | JAX: 023526 |
| Mouse: B6;129S-Rorbtm1 | The Jackson Laboratory | JAX: 023526 |
| Mouse: STOCK Slc17a6tm | The Jackson Laboratory | JAX: 016963 |
| Mouse: STOCK Slc32a1tm | The Jackson Laboratory | JAX: 016962 |
| Mouse: STOCK Slc17a6tm | The Jackson Laboratory | JAX: 016963 |
| Mouse: STOCK Nkx2-1tm | The Jackson Laboratory | JAX: 022861 |
| Mouse: STOCK Slc17a6tm | The Jackson Laboratory | JAX: 016963 |

| | | |
|---|---|---|
| Mouse: B6.FVB(Cg)-Tg(Nt | MMRRC | MMRRC: 030648 |
| Mouse: STOCK Slc32a1tm | The Jackson Laboratory | JAX: 016962 |
| Mouse: STOCK Gad2tm2( | The Jackson Laboratory | JAX: 010802 |
| Mouse: STOCK Slc32a1tm | The Jackson Laboratory | JAX: 016962 |
| Mouse: STOCK Tg(Htr2a-c | MMRRC | MMRRC: 031150 |
| Mouse: B6;129S-Nos1tm | The Jackson Laboratory | JAX: 014541 |
| Mouse: STOCK Tg(Chrna2 | MMRRC | MMRRC: 036502 |
| Mouse: B6;129S-Nos1tm | The Jackson Laboratory | JAX: 014541 |
| Mouse: STOCK Nkx2-1tm | The Jackson Laboratory | JAX: 022861 |
| Mouse: STOCK Tg(Rbp4-c | MMRRC | MMRRC: 031125 |
| Mouse: STOCK Ssttm2.1( | The Jackson Laboratory | JAX: 013044 |
| Mouse: B6.FVB(Cg)-Tg(Th | MMRRC | MMRRC: 031029 |
| Mouse: STOCK Tg(Chrna2 | MMRRC | MMRRC: 036502 |
| Mouse: STOCK Tg(Chrna2 | MMRRC | MMRRC: 036502 |
| Mouse: STOCK Tg(Drd3-cr | MMRRC | MMRRC: 034610 |
| Mouse: STOCK Tg(Drd3-cr | MMRRC | MMRRC: 034610 |
| Mouse: STOCK Tg(Gpr26- | MMRRC | MMRRC: 033032 |
| Mouse: B6;129S-Nos1tm | The Jackson Laboratory | JAX: 014541 |
| Mouse: STOCK Ssttm2.1( | The Jackson Laboratory | JAX: 013044 |
| Mouse: STOCK Tg(Syt6-cr | MMRRC | MMRRC: 032012 |
| Mouse: B6.FVB(Cg)-Tg(Th | MMRRC | MMRRC: 031029 |
| Mouse: STOCK Tg(Chrna2 | MMRRC | MMRRC: 036502 |
| Mouse: STOCK Tg(Gpr26- | MMRRC | MMRRC: 033032 |
| Mouse: B6;129S-Nos1tm | The Jackson Laboratory | JAX: 014541 |
| Mouse: B6;129S-Nos1tm | The Jackson Laboratory | JAX: 014541 |
| Mouse: STOCK Tg(Rbp4-c | MMRRC | MMRRC: 031125 |
| Mouse: B6;129S-Nos1tm | The Jackson Laboratory | JAX: 014541 |
| Mouse: STOCK Tg(Gpr26- | MMRRC | MMRRC: 033032 |
| Mouse: STOCK Tg(Drd3-cr | MMRRC | MMRRC: 034610 |
| Mouse: B6;129S-Nos1tm | The Jackson Laboratory | JAX: 014541 |
| Mouse: Lypd6-Cre_KL156 | Nathaniel Heintz and Ch | N/A |
| Mouse: STOCK Gad2tm2( | The Jackson Laboratory | JAX: 010802 |
| Mouse: STOCK Slc17a6tm | The Jackson Laboratory | JAX: 016963 |
| Mouse: STOCK Gad2tm2( | The Jackson Laboratory | JAX: 010802 |
| Mouse: STOCK Gad2tm2( | The Jackson Laboratory | JAX: 010802 |
| Mouse: STOCK Gad2tm2( | The Jackson Laboratory | JAX: 010802 |
| Mouse: STOCK Slc32a1tm | The Jackson Laboratory | JAX: 016962 |
| Mouse: STOCK Slc32a1tm | The Jackson Laboratory | JAX: 016962 |
| Mouse: STOCK Slc32a1tm | The Jackson Laboratory | JAX: 016962 |
| Mouse: STOCK Slc32a1tm | The Jackson Laboratory | JAX: 016962 |
| Mouse: STOCK Slc32a1tm | The Jackson Laboratory | JAX: 016962 |
| Mouse: STOCK Gad2tm2( | The Jackson Laboratory | JAX: 010802 |
| Mouse: STOCK Slc32a1tm | The Jackson Laboratory | JAX: 016962 |
| Mouse: STOCK Gad2tm2( | The Jackson Laboratory | JAX: 010802 |
| Mouse: STOCK Slc32a1tm | The Jackson Laboratory | JAX: 016962 |
| Mouse: STOCK Gad2tm2( | The Jackson Laboratory | JAX: 010802 |
| Mouse: STOCK Gad2tm2( | The Jackson Laboratory | JAX: 010802 |

| | | |
|---|---|---|
| Mouse: STOCK Slc17a6tm | The Jackson Laboratory | JAX: 016963 |
| Mouse: STOCK Slc17a6tm | The Jackson Laboratory | JAX: 016963 |
| Mouse: STOCK Slc17a6tm | The Jackson Laboratory | JAX: 016963 |
| Mouse: STOCK Ssttm2.1(c | The Jackson Laboratory | JAX: 013044 |
| Mouse: STOCK Ssttm2.1(c | The Jackson Laboratory | JAX: 013044 |
| Mouse: STOCK Slc17a6tm | The Jackson Laboratory | JAX: 016963 |
| Mouse: STOCK Tg(Htr3a-c | MMRRC | MMRRC: 036680 |
| Mouse: STOCK Tg(Htr3a-c | MMRRC | MMRRC: 036680 |
| Mouse: STOCK Tg(Htr3a-c | MMRRC | MMRRC: 036680 |
| Mouse: STOCK Tg(Htr3a-c | MMRRC | MMRRC: 036680 |
| Mouse: STOCK Tg(Htr3a-c | MMRRC | MMRRC: 036680 |
| Mouse: STOCK Tg(Colgalt2 | MMRRC | MMRRC: 036504 |
| Mouse: STOCK Tg(Htr3a-c | MMRRC | MMRRC: 036680 |
| Mouse: STOCK Tg(Htr3a-c | MMRRC | MMRRC: 036680 |
| Mouse: STOCK Tg(Htr3a-c | MMRRC | MMRRC: 036680 |
| Mouse: B6;129S-Penktm2 | The Jackson Laboratory | JAX: 025112 |
| Mouse: B6;129S-Penktm2 | The Jackson Laboratory | JAX: 025112 |
| Mouse: B6.FVB(Cg)-Tg(Th | MMRRC | MMRRC: 031029 |
| Mouse: B6.Cg-Erbb4tm1.1 | The Jackson Laboratory | JAX: 012360 |
| Mouse: STOCK Ccktm1.1( | The Jackson Laboratory | JAX: 012706 |
| Mouse: Adcyap1-2A-Cre | Allen Institute for Brain S | N/A |
| Mouse: STOCK Tg(Slc18a2 | MMRRC | MMRRC: 034814 |
| Mouse: STOCK Tg(Slc18a2 | MMRRC | MMRRC: 034814 |
| Mouse: Adcyap1-2A-Cre | Allen Institute for Brain S | N/A |
| Mouse: STOCK Tg(Ppp1r1 | MMRRC | MMRRC: 036205 |
| Mouse: STOCK Tg(Ppp1r1 | MMRRC | MMRRC: 036205 |
| Mouse: B6;129S6-Chattm | The Jackson Laboratory | JAX: 006410 |
| Mouse: STOCK Nkx2-1tm1 | The Jackson Laboratory | JAX: 022861 |
| Mouse: STOCK Nkx2-1tm1 | The Jackson Laboratory | JAX: 022861 |
| Mouse: B6(Cg)-Etv1tm1.1 | The Jackson Laboratory | JAX: 013048 |
| Mouse: B6;129S6-Chattm | The Jackson Laboratory | JAX: 006410 |
| Mouse: B6;129S6-Chattm | The Jackson Laboratory | JAX: 006410 |
| Mouse: B6;129S-Nos1tm1 | The Jackson Laboratory | JAX: 014541 |
| Mouse: B6;129S-Nos1tm1 | The Jackson Laboratory | JAX: 014541 |
| Mouse: STOCK Tg(Rbp4-cr | MMRRC | MMRRC: 031125 |
| Mouse: B6;129S4-Ntrk1tr | MMRRC | MMRRC: 015500 |
| Mouse: STOCK Tg(Htr2a-c | MMRRC | MMRRC: 031150 |
| Mouse: STOCK Tg(Ppp1r1 | MMRRC | MMRRC: 036205 |
| Mouse: STOCK Tg(Ppp1r1 | MMRRC | MMRRC: 036205 |
| Mouse: STOCK Tg(Pcdh9- | MMRRC | MMRRC: 036084 |
| Mouse: STOCK Tg(Pcdh9- | MMRRC | MMRRC: 036084 |
| Mouse: B6.Cg-Trib2tm1.1 | The Jackson Laboratory | JAX: 022865 |
| Mouse: B6.Cg-Trib2tm1.1 | The Jackson Laboratory | JAX: 022865 |
| Mouse: STOCK Gad2tm2( | The Jackson Laboratory | JAX: 010802 |
| Mouse: STOCK Ssttm2.1(c | The Jackson Laboratory | JAX: 013044 |
| Mouse: B6;129S-Fezf1tm1 | The Jackson Laboratory | JAX: 013048 |
| Mouse: STOCK Tg(Drd3-cr | MMRRC | MMRRC: 034610 |

| | | |
|---|---|---|
| Mouse: STOCK Tg(Drd3-cr | MMRRC | MMRRC: 034610 |
| Mouse: STOCK Tg(Drd3-cr | MMRRC | MMRRC: 034610 |
| Mouse: STOCK Tg(Efr3a-cı | MMRRC | MMRRC: 036660 |
| Mouse: STOCK Tg(Efr3a-cı | MMRRC | MMRRC: 036660 |
| Mouse: STOCK Slc32a1tm | The Jackson Laboratory | JAX: 016962 |
| Mouse: STOCK Slc17a6tm | The Jackson Laboratory | JAX: 016963 |
| Mouse: STOCK Slc17a6tm | The Jackson Laboratory | JAX: 016963 |
| Mouse: STOCK Slc17a6tm | The Jackson Laboratory | JAX: 016963 |
| Mouse: B6;129S4-Ntrk1tr | MMRRC | MMRRC: 015500 |
| Mouse: B6;129S4-Ntrk1tr | MMRRC | MMRRC: 015500 |
| Mouse: B6;129S4-Ntrk1tr | MMRRC | MMRRC: 015500 |
| Mouse: B6;129S4-Ntrk1tr | MMRRC | MMRRC: 015500 |
| Mouse: B6(Cg)-Calb2tm1( | The Jackson Laboratory | JAX: 010774 |
| Mouse: STOCK Slc17a6tm | The Jackson Laboratory | JAX: 016963 |
| Mouse: B6(Cg)-Cux2tm3.: | MMRRC | MMRRC: 032779 |
| Mouse: STOCK Ssttm2.1(c | The Jackson Laboratory | JAX: 013044 |
| Mouse: B6(Cg)-Cux2tm3.: | MMRRC | MMRRC: 032779 |
| Mouse: B6(Cg)-Cux2tm3.: | MMRRC | MMRRC: 032779 |
| Mouse: B6(Cg)-Cux2tm3.: | MMRRC | MMRRC: 032779 |
| Mouse: B6(Cg)-Crhtm1(cr | The Jackson Laboratory | JAX: 012704 |
| Mouse: B6;129S-Rasgrf2t | The Jackson Laboratory | JAX: 022864 |
| Mouse: B6;129S-Rasgrf2t | The Jackson Laboratory | JAX: 022864 |
| Mouse: STOCK Tg(Htr3a-c | MMRRC | MMRRC: 036680 |
| Mouse: STOCK Tg(Htr3a-c | MMRRC | MMRRC: 036680 |
| Mouse: B6.FVB(Cg)-Tg(Th | MMRRC | MMRRC: 031029 |
| Mouse: STOCK Slc17a6tm | The Jackson Laboratory | JAX: 016963 |
| Mouse: B6;129S-Rasgrf2t | The Jackson Laboratory | JAX: 022864 |
| Mouse: B6;129S-Rasgrf2t | The Jackson Laboratory | JAX: 022864 |
| Mouse: B6;129S-Rasgrf2t | The Jackson Laboratory | JAX: 022864 |
| Mouse: STOCK Tg(Colgalt2 | MMRRC | MMRRC: 036504 |
| Mouse: STOCK Tg(Colgalt2 | MMRRC | MMRRC: 036504 |
| Mouse: STOCK Tg(Selenov | MMRRC | MMRRC: 036190 |
| Mouse: STOCK Tg(Selenov | MMRRC | MMRRC: 036190 |
| Mouse: STOCK Tg(Selenov | MMRRC | MMRRC: 036190 |
| Mouse: B6.Cg-Ccn2tm1.1 | The Jackson Laboratory | JAX: 028535 |
| Mouse: STOCK Tg(Selenov | MMRRC | MMRRC: 036190 |
| Mouse: STOCK Tg(Selenov | MMRRC | MMRRC: 036190 |
| Mouse: B6.Cg-Calb1tm1.1 | The Jackson Laboratory | JAX: 023531 |
| Mouse: B6;129S-Penktm2 | The Jackson Laboratory | JAX: 025112 |
| Mouse: STOCK Tg(Dlg3-cr | MMRRC | MMRRC: 032809 |
| Mouse: B6;129S-Rasgrf2t | The Jackson Laboratory | JAX: 022864 |
| Mouse: B6;129S-Rasgrf2t | The Jackson Laboratory | JAX: 022864 |
| Mouse: B6;129S-Rasgrf2t | The Jackson Laboratory | JAX: 022864 |
| Mouse: B6;129S-Rasgrf2t | The Jackson Laboratory | JAX: 022864 |
| Mouse: B6.Cg-Ccn2tm1.1 | The Jackson Laboratory | JAX: 028535 |
| Mouse: STOCK Tg(Selenov | MMRRC | MMRRC: 036190 |
| Mouse: STOCK Tg(Selenov | MMRRC | MMRRC: 036190 |

| | | | |
|---|---|---|---|
| | Mouse: STOCK Tg(Dlg3-cr | MMRRC | MMRRC: 032809 |
| | Mouse: STOCK Tg(Dlg3-cr | MMRRC | MMRRC: 032809 |
| | Mouse: B6.Cg-Calb1tm1.1 | The Jackson Laboratory | JAX: 023531 |
| | Mouse: B6.Cg-Calb1tm1.1 | The Jackson Laboratory | JAX: 023531 |
| | Mouse: STOCK Tg(Dlg3-cr | MMRRC | MMRRC: 032809 |
| | Mouse: STOCK Tg(Dlg3-cr | MMRRC | MMRRC: 032809 |
| | Mouse: B6;129S-Rasgrf2t | The Jackson Laboratory | JAX: 022864 |
| Y | Mouse: B6.Cg-Ntng2tm1. | The Jackson Laboratory | JAX: 029588 |
| | Mouse: B6.Cg-Ntng2tm1. | The Jackson Laboratory | JAX: 029588 |
| | Mouse: B6.Cg-Gnb4tm1.1 | The Jackson Laboratory | JAX: 029587 |
| Y | Mouse: B6.Cg-Gnb4tm1.1 | The Jackson Laboratory | JAX: 029587 |
| | Mouse: STOCK Tg(Gpr26-c | MMRRC | MMRRC: 033032 |
| | Mouse: STOCK Tg(Colgalt2 | MMRRC | MMRRC: 036504 |
| | Mouse: B6;129S-Rorbtm1 | The Jackson Laboratory | JAX: 023526 |
| | Mouse: STOCK Tg(Sim1-cr | MMRRC | MMRRC: 031742 |
| | Mouse: STOCK Tg(Sim1-cr | MMRRC | MMRRC: 031742 |
| | Mouse: STOCK Tg(Sim1-cr | MMRRC | MMRRC: 031742 |
| | Mouse: STOCK Tg(Htr2a-c | MMRRC | MMRRC: 031150 |
| | Mouse: STOCK Tg(Htr2a-c | MMRRC | MMRRC: 031150 |
| | Mouse: B6;129S-Rorbtm1 | The Jackson Laboratory | JAX: 023526 |
| | Mouse: B6;129S-Tac2tm1 | The Jackson Laboratory | JAX: 021878 |
| | Mouse: B6;129S-Tac1tm1 | The Jackson Laboratory | JAX: 021877 |
| | Mouse: STOCK Tg(Selenov | MMRRC | MMRRC: 036190 |
| | Mouse: STOCK Tg(Selenov | MMRRC | MMRRC: 036190 |
| | Mouse: B6;C3-Tg(Scnn1a- | The Jackson Laboratory | JAX: 009112 |
| | Mouse: B6;129P2-Pvalbtr | The Jackson Laboratory | JAX: 008069 |
| | Mouse: B6.FVB(Cg)-Tg(Th | MMRRC | MMRRC: 031029 |
| | Mouse: B6.Cg-Npr3tm1.1 | The Jackson Laboratory | JAX: 031333 |
| | Mouse: B6.Cg-Npr3tm1.1 | The Jackson Laboratory | JAX: 031333 |
| Y | Mouse: B6.Cg-Ntng2tm1. | The Jackson Laboratory | JAX: 029588 |
| Y | Mouse: B6.Cg-Ntng2tm1. | The Jackson Laboratory | JAX: 029588 |
| | Mouse: B6.Cg-Gnb4tm1.1 | The Jackson Laboratory | JAX: 029587 |
| | Mouse: B6.Cg-Gnb4tm1.1 | The Jackson Laboratory | JAX: 029587 |
| | Mouse: STOCK Tg(Gal-cre | MMRRC | MMRRC: 031060 |
| | Mouse: STOCK Tg(Sim1-cr | MMRRC | MMRRC: 031742 |
| | Mouse: STOCK Tg(Sim1-cr | MMRRC | MMRRC: 031742 |
| | Mouse: STOCK Tg(Slc6a5- | MMRRC | MMRRC: 030730 |
| | Mouse: STOCK Tg(Chrnb4 | MMRRC | MMRRC: 036203 |
| | Mouse: STOCK Tg(Tlx3-cre | MMRRC | MMRRC: 041158 |
| | Mouse: STOCK Tg(Chrnb4 | MMRRC | MMRRC: 036203 |
| | Mouse: STOCK Tg(Drd3-cr | MMRRC | MMRRC: 034610 |
| | Mouse: STOCK Tg(Drd3-cr | MMRRC | MMRRC: 034610 |
| | Mouse: B6(Cg)-Cux2tm1.1 | MMRRC | MMRRC: 031778 |
| | Mouse: STOCK Tg(Drd3-cr | MMRRC | MMRRC: 034610 |
| | Mouse: STOCK Tg(Drd3-cr | MMRRC | MMRRC: 034610 |
| | Mouse: STOCK Tg(Gpr26-c | MMRRC | MMRRC: 033032 |
| | Mouse: STOCK Tg(Grp-cre | MMRRC | MMRRC: 031183 |

| | | |
|---|---|---|
| Mouse: STOCK Tg(Hdc-cre | MMRRC | MMRRC: 032079 |
| Mouse: STOCK Tg(Hdc-cre | MMRRC | MMRRC: 032079 |
| Mouse: STOCK Tg(Hdc-cre | MMRRC | MMRRC: 032079 |
| Mouse: STOCK Tg(Hdc-cre | MMRRC | MMRRC: 032079 |
| Mouse: STOCK Tg(Sim1-cr | MMRRC | MMRRC: 031742 |
| Mouse: STOCK Tg(Drd3-cr | MMRRC | MMRRC: 034610 |
| Mouse: STOCK Tg(Drd3-cr | MMRRC | MMRRC: 034610 |
| Mouse: STOCK Tg(Seleno | MMRRC | MMRRC: 036190 |
| Mouse: STOCK Tg(Prkcd-g | MMRRC | MMRRC: 011559 |
| Mouse: B6.Cg-Ccn2tm1.1 | The Jackson Laboratory | JAX: 028535 |
| Mouse: B6;129S-Rasgrf2t | The Jackson Laboratory | JAX: 022864 |
| Mouse: B6;129S-Rasgrf2t | The Jackson Laboratory | JAX: 022864 |
| Mouse: B6.Cg-Ccn2tm1.1 | The Jackson Laboratory | JAX: 028535 |
| Mouse: B6.Cg-Calb1tm1.1 | The Jackson Laboratory | JAX: 023531 |
| Mouse: B6.Cg-Calb1tm1.1 | The Jackson Laboratory | JAX: 023531 |
| Mouse: STOCK Tg(Seleno | MMRRC | MMRRC: 036190 |
| Mouse: STOCK Slc17a6tm | The Jackson Laboratory | JAX: 016963 |
| Mouse: STOCK Slc17a6tm | The Jackson Laboratory | JAX: 016963 |
| Mouse: STOCK Tg(Drd3-cr | MMRRC | MMRRC: 034610 |
| Mouse: B6.Cg-Ndnftm1.1 | The Jackson Laboratory | JAX: 028536 |
| Mouse: STOCK Tg(Drd3-cr | MMRRC | MMRRC: 034610 |
| Mouse: STOCK Tg(Drd3-cr | MMRRC | MMRRC: 034610 |
| Mouse: STOCK Tg(Drd3-cr | MMRRC | MMRRC: 034610 |
| Mouse: B6.Cg-Calb1tm1.1 | The Jackson Laboratory | JAX: 023531 |
| Mouse: B6.Cg-Calb1tm1.1 | The Jackson Laboratory | JAX: 023531 |
| Mouse: Lypd6-Cre_KL156 | Nathaniel Heintz and Cha | N/A |
| Mouse: STOCK Tg(Tlx3-cre | MMRRC | MMRRC: 041158 |
| Mouse: B6;129S-Pdyntm1 | The Jackson Laboratory | JAX: 030197 |
| Mouse: B6;129S-Pdyntm1 | The Jackson Laboratory | JAX: 030197 |
| Mouse: B6;129S-Pdyntm1 | The Jackson Laboratory | JAX: 030197 |
| Mouse: B6.129(SJL)-Kcng4 | The Jackson Laboratory | JAX: 029414 |
| Mouse: B6.129(SJL)-Kcng4 | The Jackson Laboratory | JAX: 029414 |
| Mouse: B6.129(SJL)-Kcng4 | The Jackson Laboratory | JAX: 029414 |
| Mouse: B6.Cg-Npr3tm1.1 | The Jackson Laboratory | JAX: 031333 |
| Mouse: B6.Cg-Npr3tm1.1 | The Jackson Laboratory | JAX: 031333 |
| Mouse: B6;129S-Slc17a8t | The Jackson Laboratory | JAX: 028534 |
| Mouse: STOCK Tg(Tlx3-cre | MMRRC | MMRRC: 041158 |
| Mouse: B6;129S-Slc17a8t | The Jackson Laboratory | JAX: 028534 |
| Mouse: B6;129S-Slc17a8t | The Jackson Laboratory | JAX: 028534 |
| Mouse: B6.Cg-Oxtrtm1.1( | The Jackson Laboratory | JAX: 031303 |
| Mouse: B6.Cg-Oxtrtm1.1( | The Jackson Laboratory | JAX: 031303 |
| Mouse: B6.Cg-Oxtrtm1.1( | The Jackson Laboratory | JAX: 031303 |
| Mouse: B6;129S-Cartpttm | The Jackson Laboratory | JAX: 028533 |
| Mouse: B6.Cg-Trib2tm1.1 | The Jackson Laboratory | JAX: 022865 |
| Mouse: Adcyap1-2A-Cre | Allen Institute for Brain S | N/A |
| Mouse: STOCK Tg(Htr2a-c | MMRRC | MMRRC: 031150 |
| Mouse: Adcyap1-2A-Cre | Allen Institute for Brain S | N/A |

| | | |
|---|---|---|
| Mouse: STOCK Tg(Htr2a-c | MMRRC | MMRRC: 031150 |
| Mouse: STOCK Tg(Htr2a-c | MMRRC | MMRRC: 031150 |
| Mouse: Adcyap1-2A-Cre | Allen Institute for Brain S | N/A |
| Mouse: Adcyap1-2A-Cre | Allen Institute for Brain S | N/A |
| Mouse: B6.Cg-Ccn2tm1.1 | The Jackson Laboratory | JAX: 028535 |
| Mouse: B6.Cg-Gnb4tm1.1 | The Jackson Laboratory | JAX: 030159 |
| Mouse: Tacr1-T2A-Cre | Allen Institute for Brain S | N/A |
| Mouse: Tacr1-T2A-Cre | Allen Institute for Brain S | N/A |
| Mouse: Tacr1-T2A-Cre | Allen Institute for Brain S | N/A |
| Mouse: B6;129S-Cartpttm | The Jackson Laboratory | JAX: 028533 |
| Mouse: B6;129S-Cartpttm | The Jackson Laboratory | JAX: 028533 |
| Mouse: B6;129S-Cartpttm | The Jackson Laboratory | JAX: 028533 |
| Mouse: B6;129S-Slc17a8t | The Jackson Laboratory | JAX: 028534 |
| Mouse: Tacr1-T2A-Cre | Allen Institute for Brain S | N/A |
| Mouse: B6.Cg-Ntng2tm1. | The Jackson Laboratory | JAX: 029588 |
| Mouse: B6.Cg-Ntng2tm1. | The Jackson Laboratory | JAX: 029588 |
| Mouse: B6;129S-Nos1tm1 | The Jackson Laboratory | JAX: 014541 |
| Mouse: B6;129S-Pdyntm1 | The Jackson Laboratory | JAX: 030197 |
| Mouse: Adcyap1-2A-Cre | Allen Institute for Brain S | N/A |
| Mouse: C57Bl/6J | The Jackson Laboratory | JAX: 000664 |
| Mouse: Adcyap1-2A-Cre | Allen Institute for Brain S | N/A |
| Mouse: Adcyap1-2A-Cre | Allen Institute for Brain S | N/A |
| Mouse: B6.Cg-Foxp2tm1. | The Jackson Laboratory | JAX: 030541 |
| Mouse: B6;129S-Htr1atm | The Jackson Laboratory | JAX: 030160 |
| Mouse: B6;129S-Slc17a8t | The Jackson Laboratory | JAX: 028534 |
| Mouse: B6;129S-Slc17a8t | The Jackson Laboratory | JAX: 028534 |
| Mouse: B6;129S-Esr2tm1 | The Jackson Laboratory | JAX: 030158 |
| Mouse: B6;129S-Htr1atm | The Jackson Laboratory | JAX: 030160 |
| Mouse: B6.Cg-Foxp2tm1. | The Jackson Laboratory | JAX: 030541 |
| Mouse: Tacr1-T2A-Cre | Allen Institute for Brain S | N/A |
| Mouse: B6;129S-Slc17a8t | The Jackson Laboratory | JAX: 028534 |
| Mouse: B6.Cg-Foxp2tm1. | The Jackson Laboratory | JAX: 030541 |
| Mouse: B6;129S-Nos1tm1 | The Jackson Laboratory | JAX: 014541 |
| Mouse: B6;129S-Nos1tm1 | The Jackson Laboratory | JAX: 014541 |
| Mouse: B6.Cg-Npr3tm1.1 | The Jackson Laboratory | JAX: 031333 |
| Mouse: B6.Cg-Npr3tm1.1 | The Jackson Laboratory | JAX: 031333 |
| Mouse: B6;129S-Penktm2 | The Jackson Laboratory | JAX: 025112 |
| Mouse: B6.Cg-Gnb4tm1.1 | The Jackson Laboratory | JAX: 030159 |
| Mouse: B6;129S-Nos1tm1 | The Jackson Laboratory | JAX: 014541 |
| Mouse: B6;129S-Nos1tm1 | The Jackson Laboratory | JAX: 014541 |
| Mouse: B6;129S-Nos1tm1 | The Jackson Laboratory | JAX: 014541 |
| Mouse: B6;129S-Nos1tm1 | The Jackson Laboratory | JAX: 014541 |
| Mouse: B6;129S-Esr2tm1 | The Jackson Laboratory | JAX: 030158 |
| Mouse: STOCK Tg(Colgalt2 | MMRRC | MMRRC: 036504 |
| Mouse: C57BL/6-Tg(Grik4 | The Jackson Laboratory | JAX: 006474 |
| Mouse: STOCK Tg(Syt6-cre | MMRRC | MMRRC: 032012 |
| Mouse: STOCK Tg(Syt6-cre | MMRRC | MMRRC: 032012 |

| | | |
|---|---|---|
| Mouse: STOCK Tg(Tlx3-cre | MMRRC | MMRRC: 041158 |
| Mouse: C57Bl/6J | The Jackson Laboratory | JAX: 000664 |
| Mouse: C57Bl/6J | The Jackson Laboratory | JAX: 000664 |
| Mouse: B6.Cg-Ntng2tm1. | The Jackson Laboratory | JAX: 029588 |
| Mouse: C57BL/6-Tg(Grik4 | The Jackson Laboratory | JAX: 006474 |
| Mouse: B6;129S-Penktm2 | The Jackson Laboratory | JAX: 025112 |
| Mouse: B6.Cg-Npr3tm1.1 | The Jackson Laboratory | JAX: 031333 |
| Mouse: B6;129S-Penktm2 | The Jackson Laboratory | JAX: 025112 |
| Mouse: B6.Cg-Npr3tm1.1 | The Jackson Laboratory | JAX: 031333 |
| Mouse: B6.Cg-Npr3tm1.1 | The Jackson Laboratory | JAX: 031333 |
| Mouse: B6.Cg-Npr3tm1.1 | The Jackson Laboratory | JAX: 031333 |
| Mouse: B6.Cg-Npr3tm1.1 | The Jackson Laboratory | JAX: 031333 |
| Mouse: B6;129S-Penktm2 | The Jackson Laboratory | JAX: 025112 |
| Mouse: Adcyap1-2A-Cre | Allen Institute for Brain S | N/A |
| Mouse: Adcyap1-2A-Cre | Allen Institute for Brain S | N/A |
| Mouse: STOCK Tg(Pomc1- | The Jackson Laboratory | JAX: 005965 |
| Mouse: STOCK Tg(Gabrr3 | MMRRC | MMRRC: 030709 |
| Mouse: STOCK Tg(Ppp1r1 | MMRRC | MMRRC: 036205 |
| Mouse: B6.Cg-Avptm1.1( | The Jackson Laboratory | JAX: 023530 |
| Mouse: STOCK Tg(Tlx3-cre | MMRRC | MMRRC: 041158 |
| Mouse: STOCK Tg(Tlx3-cre | MMRRC | MMRRC: 041158 |
| Mouse: B6.Cg-Npr3tm1.1 | The Jackson Laboratory | JAX: 031333 |
| Mouse: B6.Cg-Npr3tm1.1 | The Jackson Laboratory | JAX: 031333 |
| Mouse: B6.Cg-Oxtrtm1.1( | The Jackson Laboratory | JAX: 031303 |
| Mouse: STOCK Tg(Sim1-cr | MMRRC | MMRRC: 031742 |
| Mouse: B6.Cg-Oxtrtm1.1( | The Jackson Laboratory | JAX: 031303 |
| Mouse: STOCK Tg(Sim1-cr | MMRRC | MMRRC: 031742 |
| Mouse: STOCK Tg(Rbp4-cr | MMRRC | MMRRC: 031125 |
| Mouse: B6.Cg-Ccn2tm1.1 | The Jackson Laboratory | JAX: 028535 |
| Mouse: B6.Cg-Ccn2tm1.1 | The Jackson Laboratory | JAX: 028535 |
| Mouse: STOCK Tg(Sim1-cr | MMRRC | MMRRC: 031742 |
| Mouse: STOCK Tg(Sim1-cr | MMRRC | MMRRC: 031742 |
| Mouse: STOCK Tg(Rbp4-cr | MMRRC | MMRRC: 031125 |
| Mouse: Tg(Slc17a8-icre)1 | The Jackson Laboratory | JAX: 018147 |
| Mouse: STOCK Tg(Tlx3-cre | MMRRC | MMRRC: 041158 |
| Mouse: B6;129S-Rorbtm1 | The Jackson Laboratory | JAX: 023526 |
| Mouse: C57Bl/6J | The Jackson Laboratory | JAX: 000664 |
| Mouse: B6.Cg-Ccn2tm1.1 | The Jackson Laboratory | JAX: 028535 |
| Mouse: B6.Cg-Ccn2tm1.1 | The Jackson Laboratory | JAX: 028535 |
| Mouse: B6.Cg-Pvalbtm1.1 | The Jackson Laboratory | JAX: 021189 |
| Mouse: B6.Cg-Pvalbtm1.1 | The Jackson Laboratory | JAX: 021189 |
| Mouse: B6;129S-Penktm2 | The Jackson Laboratory | JAX: 025112 |
| Mouse: C57Bl/6J | The Jackson Laboratory | JAX: 000664 |
| Mouse: C57Bl/6J | The Jackson Laboratory | JAX: 000664 |
| Mouse: C57Bl/6J | The Jackson Laboratory | JAX: 000664 |
| Mouse: B6;C3-Tg(Scnn1a- | The Jackson Laboratory | JAX: 009112 |
| Mouse: STOCK Tg(Vipr2-c | The Jackson Laboratory | JAX: 031332 |

| | | |
|---|---|---|
| Mouse: STOCK Tg(Vipr2-c | The Jackson Laboratory | JAX: 031332 |
| Mouse: B6;C3-Tg(Scnn1a- | The Jackson Laboratory | JAX: 009112 |
| Mouse: C57Bl/6J | The Jackson Laboratory | JAX: 000664 |
| Mouse: Rorb-IRES2-Cre-n | Allen Institute for Brain S | N/A |
| Mouse: Rorb-IRES2-Cre-n | Allen Institute for Brain S | N/A |
| Mouse: Tg(Slc17a8-icre)1 | The Jackson Laboratory | JAX: 018147 |
| Mouse: B6;129S-Slc17a8t | The Jackson Laboratory | JAX: 028534 |
| Mouse: B6;129S-Rorbtm1 | The Jackson Laboratory | JAX: 023526 |
| Mouse: B6;129S-Penktm2 | The Jackson Laboratory | JAX: 025112 |
| Mouse: B6;129S-Penktm2 | The Jackson Laboratory | JAX: 025112 |
| Mouse: B6;129S-Penktm2 | The Jackson Laboratory | JAX: 025112 |
| Mouse: B6;129S-Slc17a8t | The Jackson Laboratory | JAX: 028534 |
| Mouse: B6;129S-Slc17a8t | The Jackson Laboratory | JAX: 028534 |
| Mouse: B6;129S-Rorbtm1 | The Jackson Laboratory | JAX: 023526 |
| Mouse: STOCK Ssttm2.1(c | The Jackson Laboratory | JAX: 013044 |
| Mouse: B6.Cg-Oxtrtm1.1( | The Jackson Laboratory | JAX: 031303 |
| Mouse: STOCK Nkx2-1tm1 | The Jackson Laboratory | JAX: 014552 |
| Mouse: STOCK Tg(Tlx3-cre | MMRRC | MMRRC: 041158 |
| Mouse: STOCK Tg(Sim1-cr | MMRRC | MMRRC: 031742 |
| Mouse: STOCK Tg(Sim1-cr | MMRRC | MMRRC: 031742 |
| Mouse: STOCK Ssttm2.1(c | The Jackson Laboratory | JAX: 013044 |
| Mouse: B6.Cg-Oxtrtm1.1( | The Jackson Laboratory | JAX: 031303 |
| Mouse: STOCK Tg(Tlx3-cre | MMRRC | MMRRC: 041158 |
| Mouse: STOCK Tg(Tlx3-cre | MMRRC | MMRRC: 041158 |
| Mouse: STOCK Nkx2-1tm1 | The Jackson Laboratory | JAX: 014552 |
| Mouse: B6(Cg)-Cux2tm1.1 | MMRRC | MMRRC: 031778 |
| Mouse: B6.FVB(Cg)-Tg(Nt | MMRRC | MMRRC: 030648 |
| Mouse: STOCK Tg(Sim1-cr | MMRRC | MMRRC: 031742 |
| Mouse: STOCK Ssttm2.1(c | The Jackson Laboratory | JAX: 013044 |
| Mouse: STOCK Ssttm2.1(c | The Jackson Laboratory | JAX: 013044 |
| Mouse: STOCK Ssttm2.1(c | The Jackson Laboratory | JAX: 013044 |
| Mouse: STOCK Nkx2-1tm1 | The Jackson Laboratory | JAX: 014552 |
| Mouse: STOCK Tg(Tlx3-cre | MMRRC | MMRRC: 041158 |
| Mouse: B6.Cg-Foxp2tm1.1 | The Jackson Laboratory | JAX: 030541 |
| Mouse: B6.Cg-Foxp2tm1.1 | The Jackson Laboratory | JAX: 030541 |
| Mouse: STOCK Tg(Syt6-cre | MMRRC | MMRRC: 032012 |
| Mouse: STOCK Tg(Syt6-cre | MMRRC | MMRRC: 032012 |
| Mouse: STOCK Ssttm2.1(c | The Jackson Laboratory | JAX: 013044 |
| Mouse: B6;129S-Penktm2 | The Jackson Laboratory | JAX: 025112 |
| Mouse: B6;129S-Penktm2 | The Jackson Laboratory | JAX: 025112 |
| Mouse: B6;129S-Penktm2 | The Jackson Laboratory | JAX: 025112 |
| Mouse: B6;129S-Rorbtm1 | The Jackson Laboratory | JAX: 023526 |
| Mouse: B6;129S-Rorbtm1 | The Jackson Laboratory | JAX: 023526 |
| Mouse: STOCK Tg(Colgalt2 | MMRRC | MMRRC: 036504 |
| Mouse: Rorb-IRES2-Cre-n | Allen Institute for Brain S | N/A |
| Mouse: STOCK Tg(Seleno | MMRRC | MMRRC: 036190 |
| Mouse: Rorb-IRES2-Cre-n | Allen Institute for Brain S | N/A |

| | | |
|---|---|---|
| Mouse: STOCK Tg(Colgalt2 | MMRRC | MMRRC: 036504 |
| Mouse: B6.Cg-Pvalbtm1.1 | The Jackson Laboratory | JAX: 021189 |
| Mouse: B6.Cg-Pvalbtm1.1 | The Jackson Laboratory | JAX: 021189 |
| Mouse: B6;129S-Penktm2 | The Jackson Laboratory | JAX: 025112 |
| Mouse: Rorb-IRES2-Cre-n | Allen Institute for Brain S | N/A |
| Mouse: Tg(Slc17a8-icre)1 | The Jackson Laboratory | JAX: 018147 |
| Mouse: B6;129S-Penktm2 | The Jackson Laboratory | JAX: 025112 |
| Mouse: B6;129S-Penktm2 | The Jackson Laboratory | JAX: 025112 |
| Mouse: STOCK Tg(Selenov | MMRRC | MMRRC: 036190 |
| Mouse: B6;129S-Rorbtm1 | The Jackson Laboratory | JAX: 023526 |
| Mouse: Rorb-IRES2-Cre-n | Allen Institute for Brain S | N/A |
| Mouse: B6;129S-Rorbtm1 | The Jackson Laboratory | JAX: 023526 |
| Mouse: B6.Cg-Tg(A930038 | The Jackson Laboratory | JAX: 017346 |
| Mouse: B6.FVB(Cg)-Tg(Nt | MMRRC | MMRRC: 030648 |
| Mouse: B6.129S2-Emx1tr | The Jackson Laboratory | JAX: 005628 |
| Mouse: B6;129S6-Chattm | The Jackson Laboratory | JAX: 006410 |
| Mouse: STOCK Tg(Prkcd-g | MMRRC | MMRRC: 011559 |
| Mouse: B6.129S2-Emx1tr | The Jackson Laboratory | JAX: 005628 |
| Mouse: B6;C3-Tg(Scnn1a- | The Jackson Laboratory | JAX: 009613 |
| Mouse: STOCK Tg(Prkcd-g | MMRRC | MMRRC: 011559 |
| Mouse: B6.129S2-Emx1tr | The Jackson Laboratory | JAX: 005628 |
| Mouse: B6.129S2-Emx1tr | The Jackson Laboratory | JAX: 005628 |
| Mouse: B6.129S2-Emx1tr | The Jackson Laboratory | JAX: 005628 |
| Mouse: B6.129S2-Emx1tr | The Jackson Laboratory | JAX: 005628 |
| Mouse: B6.129S2-Emx1tr | The Jackson Laboratory | JAX: 005628 |
| Mouse: B6.129S2-Emx1tr | The Jackson Laboratory | JAX: 005628 |
| Mouse: B6;C3-Tg(Scnn1a- | The Jackson Laboratory | JAX: 009112 |
| Mouse: B6.Cg-Tg(Slc6a4-c | MMRRC | MMRRC: 031028 |
| Mouse: STOCK Tg(Dbh-cr | MMRRC | MMRRC: 032081 |
| Mouse: STOCK Slc17a6tm | The Jackson Laboratory | JAX: 016963 |
| Mouse: STOCK Slc17a6tm | The Jackson Laboratory | JAX: 016963 |
| Mouse: STOCK Tg(Rbp4-cr | MMRRC | MMRRC: 031125 |
| Mouse: B6(Cg)-Cux2tm1.1 | MMRRC | MMRRC: 031778 |
| Mouse: STOCK Tg(Rbp4-cr | MMRRC | MMRRC: 031125 |
| Mouse: B6(Cg)-Cux2tm1.1 | MMRRC | MMRRC: 031778 |
| Mouse: B6(Cg)-Cux2tm1.1 | MMRRC | MMRRC: 031778 |
| Mouse: STOCK Tg(Rbp4-cr | MMRRC | MMRRC: 031125 |
| Mouse: STOCK Tg(Rbp4-cr | MMRRC | MMRRC: 031125 |
| Mouse: STOCK Tg(Rbp4-cr | MMRRC | MMRRC: 031125 |
| Mouse: B6.129S2-Emx1tr | The Jackson Laboratory | JAX: 005628 |
| Mouse: B6;C3-Tg(Scnn1a- | The Jackson Laboratory | JAX: 009613 |
| Mouse: B6;129S6-Chattm | The Jackson Laboratory | JAX: 006410 |
| Mouse: B6;129S6-Chattm | The Jackson Laboratory | JAX: 006410 |
| Mouse: STOCK Tg(Prkcd-g | MMRRC | MMRRC: 011559 |
| Mouse: B6.129S2-Emx1tr | The Jackson Laboratory | JAX: 005628 |
| Mouse: STOCK Slc17a6tm | The Jackson Laboratory | JAX: 016963 |
| Mouse: B6;129S6-Chattm | The Jackson Laboratory | JAX: 006410 |

| | | |
|---|---|---|
| Mouse: B6.129S2-Emx1tr | The Jackson Laboratory | JAX: 005628 |
| Mouse: B6.129S2-Emx1tr | The Jackson Laboratory | JAX: 005628 |
| Mouse: B6.129S2-Emx1tr | The Jackson Laboratory | JAX: 005628 |
| Mouse: STOCK Tg(Tlx3-cre | MMRRC | MMRRC: 041158 |
| Mouse: B6.Cg-Tg(A93003 | The Jackson Laboratory | JAX: 017346 |
| Mouse: B6.Cg-Tg(A93003 | The Jackson Laboratory | JAX: 017346 |
| Mouse: B6.Cg-Tg(A93003 | The Jackson Laboratory | JAX: 017346 |
| Mouse: STOCK Tg(Tlx3-cre | MMRRC | MMRRC: 041158 |
| Mouse: B6.Cg-Tg(A93003 | The Jackson Laboratory | JAX: 017346 |
| Mouse: B6(Cg)-Cux2tm1. | MMRRC | MMRRC: 031778 |
| Mouse: B6.129S2-Emx1tr | The Jackson Laboratory | JAX: 005628 |
| Mouse: STOCK Tg(Tlx3-cre | MMRRC | MMRRC: 041158 |
| Mouse: STOCK Tg(Tlx3-cre | MMRRC | MMRRC: 041158 |
| Mouse: B6(Cg)-Cux2tm1. | MMRRC | MMRRC: 031778 |
| Mouse: B6(Cg)-Cux2tm1. | MMRRC | MMRRC: 031778 |
| Mouse: B6.FVB(Cg)-Tg(Nt | MMRRC | MMRRC: 030648 |
| Mouse: STOCK Tg(Rbp4-cı | MMRRC | MMRRC: 031125 |
| Mouse: B6(Cg)-Cux2tm1. | MMRRC | MMRRC: 031778 |
| Mouse: STOCK Tg(Rbp4-cı | MMRRC | MMRRC: 031125 |
| Mouse: STOCK Tg(Rbp4-cı | MMRRC | MMRRC: 031125 |
| Mouse: B6.Cg-Tg(A93003 | The Jackson Laboratory | JAX: 017346 |
| Mouse: STOCK Tg(Tlx3-cre | MMRRC | MMRRC: 041158 |
| Mouse: STOCK Tg(Tlx3-cre | MMRRC | MMRRC: 041158 |
| Mouse: B6.FVB(Cg)-Tg(Nt | MMRRC | MMRRC: 030648 |
| Mouse: STOCK Tg(Tlx3-cre | MMRRC | MMRRC: 041158 |
| Mouse: STOCK Tg(Tlx3-cre | MMRRC | MMRRC: 041158 |
| Mouse: STOCK Tg(Rbp4-cı | MMRRC | MMRRC: 031125 |
| Mouse: STOCK Tg(Tlx3-cre | MMRRC | MMRRC: 041158 |
| Mouse: B6.129S2-Emx1tr | The Jackson Laboratory | JAX: 005628 |
| Mouse: B6.Cg-Tg(A93003 | The Jackson Laboratory | JAX: 017346 |
| Mouse: B6.129S2-Emx1tr | The Jackson Laboratory | JAX: 005628 |
| Mouse: B6.129S2-Emx1tr | The Jackson Laboratory | JAX: 005628 |
| Mouse: B6.FVB(Cg)-Tg(Nt | MMRRC | MMRRC: 030648 |
| Mouse: B6(Cg)-Cux2tm1. | MMRRC | MMRRC: 031778 |
| Mouse: B6.FVB(Cg)-Tg(Nt | MMRRC | MMRRC: 030648 |
| Mouse: B6.FVB(Cg)-Tg(Nt | MMRRC | MMRRC: 030648 |
| Mouse: STOCK Tg(Rbp4-cı | MMRRC | MMRRC: 031125 |
| Mouse: STOCK Tg(Tlx3-cre | MMRRC | MMRRC: 041158 |
| Mouse: B6.FVB(Cg)-Tg(Nt | MMRRC | MMRRC: 030648 |
| Mouse: B6.FVB(Cg)-Tg(Nt | MMRRC | MMRRC: 030648 |
| Mouse: B6.FVB(Cg)-Tg(Nt | MMRRC | MMRRC: 030648 |
| Mouse: B6.FVB(Cg)-Tg(Nt | MMRRC | MMRRC: 030648 |
| Mouse: B6.FVB(Cg)-Tg(Nt | MMRRC | MMRRC: 030648 |
| Mouse: B6.Cg-Tg(A93003 | The Jackson Laboratory | JAX: 017346 |
| Mouse: B6.FVB(Cg)-Tg(Nt | MMRRC | MMRRC: 030648 |
| Mouse: STOCK Slc17a6tm | The Jackson Laboratory | JAX: 016963 |
| Mouse: B6.Cg-Tg(A93003 | The Jackson Laboratory | JAX: 017346 |

| | | |
|---|---|---|
| Mouse: STOCK Tg(Rbp4-cı | MMRRC | MMRRC: 031125 |
| Mouse: STOCK Tg(Tlx3-cre | MMRRC | MMRRC: 041158 |
| Mouse: B6.129S2-Emx1tr | The Jackson Laboratory | JAX: 005628 |
| Mouse: STOCK Tg(Tlx3-cre | MMRRC | MMRRC: 041158 |
| Mouse: B6(Cg)-Cux2tm1.: | MMRRC | MMRRC: 031778 |
| Mouse: B6.FVB(Cg)-Tg(Nt: | MMRRC | MMRRC: 030648 |
| Mouse: B6.129S2-Emx1tr | The Jackson Laboratory | JAX: 005628 |
| Mouse: B6.129S2-Emx1tr | The Jackson Laboratory | JAX: 005628 |
| Mouse: B6.129S2-Emx1tr | The Jackson Laboratory | JAX: 005628 |
| Mouse: B6.129S2-Emx1tr | The Jackson Laboratory | JAX: 005628 |
| Mouse: B6.129S2-Emx1tr | The Jackson Laboratory | JAX: 005628 |
| Mouse: B6(Cg)-Cux2tm1.: | MMRRC | MMRRC: 031778 |
| Mouse: B6(Cg)-Cux2tm1.: | MMRRC | MMRRC: 031778 |
| Mouse: B6(Cg)-Cux2tm1.: | MMRRC | MMRRC: 031778 |
| Mouse: STOCK Tg(Rbp4-cı | MMRRC | MMRRC: 031125 |
| Mouse: STOCK Tg(Sim1-cr | MMRRC | MMRRC: 031742 |
| Mouse: B6(Cg)-Cux2tm1.: | MMRRC | MMRRC: 031778 |
| Mouse: B6(Cg)-Cux2tm1.: | MMRRC | MMRRC: 031778 |
| Mouse: B6(Cg)-Cux2tm1.: | MMRRC | MMRRC: 031778 |
| Mouse: STOCK Tg(Prkcd-g | MMRRC | MMRRC: 011559 |
| Mouse: B6.129S2-Emx1tr | The Jackson Laboratory | JAX: 005628 |
| Mouse: STOCK Tg(Prkcd-g | MMRRC | MMRRC: 011559 |
| Mouse: STOCK Slc17a6tm | The Jackson Laboratory | JAX: 016963 |
| Mouse: B6.129S2-Emx1tr | The Jackson Laboratory | JAX: 005628 |
| Mouse: B6.Cg-Tg(A93003 | The Jackson Laboratory | JAX: 017346 |
| Mouse: B6.Cg-Tg(A93003 | The Jackson Laboratory | JAX: 017346 |
| Mouse: B6(Cg)-Cux2tm1.: | MMRRC | MMRRC: 031778 |
| Mouse: B6.Cg-Tg(A93003 | The Jackson Laboratory | JAX: 017346 |
| Mouse: B6.129S2-Emx1tr | The Jackson Laboratory | JAX: 005628 |
| Mouse: B6.129S2-Emx1tr | The Jackson Laboratory | JAX: 005628 |
| Mouse: B6.Cg-Tg(A93003 | The Jackson Laboratory | JAX: 017346 |
| Mouse: B6.129S2-Emx1tr | The Jackson Laboratory | JAX: 005628 |
| Mouse: B6.Cg-Tg(A93003 | The Jackson Laboratory | JAX: 017346 |
| Mouse: B6.FVB(Cg)-Tg(Nt: | MMRRC | MMRRC: 030648 |
| Mouse: B6.FVB(Cg)-Tg(Nt: | MMRRC | MMRRC: 030648 |
| Mouse: B6.129S2-Emx1tr | The Jackson Laboratory | JAX: 005628 |
| Mouse: STOCK Tg(Tlx3-cre | MMRRC | MMRRC: 041158 |
| Mouse: B6.129S2-Emx1tr | The Jackson Laboratory | JAX: 005628 |
| Mouse: B6.129S2-Emx1tr | The Jackson Laboratory | JAX: 005628 |
| Mouse: STOCK Tg(Rbp4-cı | MMRRC | MMRRC: 031125 |
| Mouse: STOCK Tg(Rbp4-cı | MMRRC | MMRRC: 031125 |
| Mouse: B6.FVB(Cg)-Tg(Nt: | MMRRC | MMRRC: 030648 |
| Mouse: STOCK Tg(Tlx3-cre | MMRRC | MMRRC: 041158 |
| Mouse: STOCK Tg(Tlx3-cre | MMRRC | MMRRC: 041158 |
| Mouse: STOCK Tg(Sim1-cr | MMRRC | MMRRC: 031742 |
| Mouse: STOCK Tg(Sim1-cr | MMRRC | MMRRC: 031742 |
| Mouse: B6;C3-Tg(Scnn1a- | The Jackson Laboratory | JAX: 009613 |

| | | |
|---|---|---|
| Mouse: B6.FVB(Cg)-Tg(Nt | MMRRC | MMRRC: 030648 |
| Mouse: STOCK Tg(Tlx3-cre | MMRRC | MMRRC: 041158 |
| Mouse: STOCK Tg(Rbp4-cr | MMRRC | MMRRC: 031125 |
| Mouse: B6;C3-Tg(Scnn1a- | The Jackson Laboratory | JAX: 009613 |
| Mouse: B6.Cg-Tg(A93003 | The Jackson Laboratory | JAX: 017346 |
| Mouse: B6.Cg-Tg(A93003 | The Jackson Laboratory | JAX: 017346 |
| Mouse: B6.Cg-Tg(A93003 | The Jackson Laboratory | JAX: 017346 |
| Mouse: STOCK Tg(Tlx3-cre | MMRRC | MMRRC: 041158 |
| Mouse: B6.Cg-Tg(A93003 | The Jackson Laboratory | JAX: 017346 |
| Mouse: STOCK Tg(Rbp4-cr | MMRRC | MMRRC: 031125 |
| Mouse: B6;129S4-Ntrk1tr | MMRRC | MMRRC: 015500 |
| Mouse: STOCK Tg(Rbp4-cr | MMRRC | MMRRC: 031125 |
| Mouse: STOCK Tg(Tlx3-cre | MMRRC | MMRRC: 041158 |
| Mouse: STOCK Tg(Rbp4-cr | MMRRC | MMRRC: 031125 |
| Mouse: STOCK Tg(Rbp4-cr | MMRRC | MMRRC: 031125 |
| Mouse: B6.FVB(Cg)-Tg(Nt | MMRRC | MMRRC: 030648 |
| Mouse: B6.FVB(Cg)-Tg(Nt | MMRRC | MMRRC: 030648 |
| Mouse: STOCK Tg(Tlx3-cre | MMRRC | MMRRC: 041158 |
| Mouse: B6.Cg-Gnb4tm1.1 | The Jackson Laboratory | JAX: 030159 |
| Mouse: B6.129S2-Emx1tr | The Jackson Laboratory | JAX: 005628 |
| Mouse: B6;C3-Tg(Scnn1a- | The Jackson Laboratory | JAX: 009613 |
| Mouse: B6;C3-Tg(Scnn1a- | The Jackson Laboratory | JAX: 009613 |
| Mouse: B6;C3-Tg(Scnn1a- | The Jackson Laboratory | JAX: 009613 |
| Mouse: B6;C3-Tg(Scnn1a- | The Jackson Laboratory | JAX: 009613 |
| Mouse: B6;C3-Tg(Scnn1a- | The Jackson Laboratory | JAX: 009613 |
| Mouse: B6;C3-Tg(Scnn1a- | The Jackson Laboratory | JAX: 009613 |
| Mouse: B6;C3-Tg(Scnn1a- | The Jackson Laboratory | JAX: 009613 |
| Mouse: STOCK Tg(Tlx3-cre | MMRRC | MMRRC: 041158 |
| Mouse: B6;C3-Tg(Scnn1a- | The Jackson Laboratory | JAX: 009613 |
| Mouse: B6;C3-Tg(Scnn1a- | The Jackson Laboratory | JAX: 009613 |
| Mouse: B6;C3-Tg(Scnn1a- | The Jackson Laboratory | JAX: 009613 |
| Mouse: B6;C3-Tg(Scnn1a- | The Jackson Laboratory | JAX: 009613 |
| Mouse: B6.Cg-Tg(A93003 | The Jackson Laboratory | JAX: 017346 |
| Mouse: STOCK Tg(Prkcd-g | MMRRC | MMRRC: 011559 |
| Mouse: B6.129S2-Emx1tr | The Jackson Laboratory | JAX: 005628 |
| Mouse: B6.129S2-Emx1tr | The Jackson Laboratory | JAX: 005628 |
| Mouse: B6;129S6-Chattm | The Jackson Laboratory | JAX: 006410 |
| Mouse: B6;129S6-Chattm | The Jackson Laboratory | JAX: 006410 |
| Mouse: STOCK Tg(Prkcd-g | MMRRC | MMRRC: 011559 |
| Mouse: STOCK Slc17a6tm | The Jackson Laboratory | JAX: 016963 |
| Mouse: B6;C3-Tg(Scnn1a- | The Jackson Laboratory | JAX: 009112 |
| Mouse: B6.129S2-Emx1tr | The Jackson Laboratory | JAX: 005628 |
| Mouse: B6.129S2-Emx1tr | The Jackson Laboratory | JAX: 005628 |
| Mouse: B6.129S2-Emx1tr | The Jackson Laboratory | JAX: 005628 |
| Mouse: B6.129S2-Emx1tr | The Jackson Laboratory | JAX: 005628 |
| Mouse: B6.129S2-Emx1tr | The Jackson Laboratory | JAX: 005628 |
| Mouse: B6.129S2-Emx1tr | The Jackson Laboratory | JAX: 005628 |

| | | |
|---|---|---|
| Mouse: STOCK Tg(Rbp4-cı | MMRRC | MMRRC: 031125 |
| Mouse: STOCK Tg(Rbp4-cı | MMRRC | MMRRC: 031125 |
| Mouse: STOCK Tg(Rbp4-cı | MMRRC | MMRRC: 031125 |
| Mouse: B6(Cg)-Cux2tm1.: | MMRRC | MMRRC: 031778 |
| Mouse: B6(Cg)-Cux2tm1.: | MMRRC | MMRRC: 031778 |
| Mouse: STOCK Slc17a6tm | The Jackson Laboratory | JAX: 016963 |
| Mouse: B6(Cg)-Cux2tm1.: | MMRRC | MMRRC: 031778 |
| Mouse: B6.FVB(Cg)-Tg(Nt | MMRRC | MMRRC: 030648 |
| Mouse: B6.FVB(Cg)-Tg(Nt | MMRRC | MMRRC: 030648 |
| Mouse: B6.FVB(Cg)-Tg(Nt | MMRRC | MMRRC: 030648 |
| Mouse: B6(Cg)-Cux2tm1.: | MMRRC | MMRRC: 031778 |
| Mouse: STOCK Tg(Rbp4-cı | MMRRC | MMRRC: 031125 |
| Mouse: STOCK Tg(Dbh-cr | MMRRC | MMRRC: 032081 |
| Mouse: B6(Cg)-Cux2tm1.: | MMRRC | MMRRC: 031778 |
| Mouse: B6.Cg-Tg(A93003 | The Jackson Laboratory | JAX: 017346 |
| Mouse: B6.Cg-Tg(A93003 | The Jackson Laboratory | JAX: 017346 |
| Mouse: B6.129S2-Emx1tr | The Jackson Laboratory | JAX: 005628 |
| Mouse: B6.129S2-Emx1tr | The Jackson Laboratory | JAX: 005628 |
| Mouse: B6.129S2-Emx1tr | The Jackson Laboratory | JAX: 005628 |
| Mouse: B6.129S2-Emx1tr | The Jackson Laboratory | JAX: 005628 |
| Mouse: B6.129S2-Emx1tr | The Jackson Laboratory | JAX: 005628 |
| Mouse: B6.129S2-Emx1tr | The Jackson Laboratory | JAX: 005628 |
| Mouse: B6(Cg)-Cux2tm1.: | MMRRC | MMRRC: 031778 |
| Mouse: B6(Cg)-Cux2tm1.: | MMRRC | MMRRC: 031778 |
| Mouse: B6.129S2-Emx1tr | The Jackson Laboratory | JAX: 005628 |
| Mouse: B6.129S2-Emx1tr | The Jackson Laboratory | JAX: 005628 |
| Mouse: B6.129S2-Emx1tr | The Jackson Laboratory | JAX: 005628 |
| Mouse: STOCK Tg(Tlx3-cre | MMRRC | MMRRC: 041158 |
| Mouse: STOCK Tg(Tlx3-cre | MMRRC | MMRRC: 041158 |
| Mouse: STOCK Tg(Tlx3-cre | MMRRC | MMRRC: 041158 |
| Mouse: STOCK Tg(Tlx3-cre | MMRRC | MMRRC: 041158 |
| Mouse: STOCK Tg(Rbp4-cı | MMRRC | MMRRC: 031125 |
| Mouse: STOCK Tg(Rbp4-cı | MMRRC | MMRRC: 031125 |
| Mouse: B6;C3-Tg(Scnn1a- | The Jackson Laboratory | JAX: 009613 |
| Mouse: STOCK Tg(Prkcd-g | MMRRC | MMRRC: 011559 |
| Mouse: B6;129S6-Chattm | The Jackson Laboratory | JAX: 006410 |
| Mouse: B6(Cg)-Cux2tm1.: | MMRRC | MMRRC: 031778 |
| Mouse: B6(Cg)-Cux2tm1.: | MMRRC | MMRRC: 031778 |
| Mouse: STOCK Tg(Prkcd-g | MMRRC | MMRRC: 011559 |
| Mouse: B6.Cg-Tg(A93003 | The Jackson Laboratory | JAX: 017346 |
| Mouse: STOCK Tg(Tlx3-cre | MMRRC | MMRRC: 041158 |
| Mouse: STOCK Tg(Tlx3-cre | MMRRC | MMRRC: 041158 |
| Mouse: STOCK Tg(Rbp4-cı | MMRRC | MMRRC: 031125 |
| Mouse: STOCK Tg(Tlx3-cre | MMRRC | MMRRC: 041158 |
| Mouse: B6.FVB(Cg)-Tg(Nt | MMRRC | MMRRC: 030648 |
| Mouse: B6.129S2-Emx1tr | The Jackson Laboratory | JAX: 005628 |
| Mouse: B6.FVB(Cg)-Tg(Nt | MMRRC | MMRRC: 030648 |

| | | |
|---|---|---|
| Mouse: B6.FVB(Cg)-Tg(Nt | MMRRC | MMRRC: 030648 |
| Mouse: B6.FVB(Cg)-Tg(Nt | MMRRC | MMRRC: 030648 |
| Mouse: B6.FVB(Cg)-Tg(Nt | MMRRC | MMRRC: 030648 |
| Mouse: B6.FVB(Cg)-Tg(Nt | MMRRC | MMRRC: 030648 |
| Mouse: STOCK Tg(Rbp4-c | MMRRC | MMRRC: 031125 |
| Mouse: B6(Cg)-Cux2tm1. | MMRRC | MMRRC: 031778 |
| Mouse: B6(Cg)-Cux2tm1. | MMRRC | MMRRC: 031778 |
| Mouse: B6(Cg)-Cux2tm1. | MMRRC | MMRRC: 031778 |
| Mouse: B6.FVB(Cg)-Tg(Nt | MMRRC | MMRRC: 030648 |
| Mouse: STOCK Tg(Tlx3-cre | MMRRC | MMRRC: 041158 |
| Mouse: STOCK Tg(Tlx3-cre | MMRRC | MMRRC: 041158 |
| Mouse: B6.Cg-Tg(A93003 | The Jackson Laboratory | JAX: 017346 |
| Mouse: STOCK Tg(Dbh-cr | MMRRC | MMRRC: 032081 |
| Mouse: B6.Cg-Tg(A93003 | The Jackson Laboratory | JAX: 017346 |
| Mouse: B6.129S2-Emx1tr | The Jackson Laboratory | JAX: 005628 |
| Mouse: STOCK Tg(Tlx3-cre | MMRRC | MMRRC: 041158 |
| Mouse: B6.FVB(Cg)-Tg(Nt | MMRRC | MMRRC: 030648 |
| Mouse: STOCK Tg(Rbp4-c | MMRRC | MMRRC: 031125 |
| Mouse: STOCK Tg(Rbp4-c | MMRRC | MMRRC: 031125 |
| Mouse: B6.Cg-Tg(A93003 | The Jackson Laboratory | JAX: 017346 |
| Mouse: B6.129S2-Emx1tr | The Jackson Laboratory | JAX: 005628 |
| Mouse: B6(Cg)-Cux2tm1. | MMRRC | MMRRC: 031778 |
| Mouse: B6.FVB(Cg)-Tg(Nt | MMRRC | MMRRC: 030648 |
| Mouse: B6.FVB(Cg)-Tg(Nt | MMRRC | MMRRC: 030648 |
| Mouse: B6(Cg)-Cux2tm1. | MMRRC | MMRRC: 031778 |
| Mouse: B6.FVB(Cg)-Tg(Nt | MMRRC | MMRRC: 030648 |
| Mouse: B6.FVB(Cg)-Tg(Nt | MMRRC | MMRRC: 030648 |
| Mouse: B6.FVB(Cg)-Tg(Nt | MMRRC | MMRRC: 030648 |
| Mouse: B6.FVB(Cg)-Tg(Nt | MMRRC | MMRRC: 030648 |
| Mouse: STOCK Tg(Rbp4-c | MMRRC | MMRRC: 031125 |
| Mouse: B6.FVB(Cg)-Tg(Nt | MMRRC | MMRRC: 030648 |
| Mouse: B6.FVB(Cg)-Tg(Nt | MMRRC | MMRRC: 030648 |
| Mouse: B6.129S2-Emx1tr | The Jackson Laboratory | JAX: 005628 |
| Mouse: B6.129S2-Emx1tr | The Jackson Laboratory | JAX: 005628 |
| Mouse: STOCK Tg(Tlx3-cre | MMRRC | MMRRC: 041158 |
| Mouse: STOCK Tg(Tlx3-cre | MMRRC | MMRRC: 041158 |
| Mouse: STOCK Slc17a6tm | The Jackson Laboratory | JAX: 016963 |
| Mouse: B6.129S2-Emx1tr | The Jackson Laboratory | JAX: 005628 |
| Mouse: B6.129S2-Emx1tr | The Jackson Laboratory | JAX: 005628 |
| Mouse: B6(Cg)-Cux2tm1. | MMRRC | MMRRC: 031778 |
| Mouse: STOCK Tg(Rbp4-c | MMRRC | MMRRC: 031125 |
| Mouse: STOCK Tg(Rbp4-c | MMRRC | MMRRC: 031125 |
| Mouse: B6.129S2-Emx1tr | The Jackson Laboratory | JAX: 005628 |
| Mouse: B6.Cg-Tg(A93003 | The Jackson Laboratory | JAX: 017346 |
| Mouse: B6.129S2-Emx1tr | The Jackson Laboratory | JAX: 005628 |
| Mouse: B6.129S2-Emx1tr | The Jackson Laboratory | JAX: 005628 |
| Mouse: B6.Cg-Tg(A93003 | The Jackson Laboratory | JAX: 017346 |

| | | |
|---|---|---|
| Mouse: B6.129S2-Emx1tr | The Jackson Laboratory | JAX: 005628 |
| Mouse: B6(Cg)-Cux2tm1.: | MMRRC | MMRRC: 031778 |
| Mouse: STOCK Tg(Tlx3-cre | MMRRC | MMRRC: 041158 |
| Mouse: STOCK Tg(Tlx3-cre | MMRRC | MMRRC: 041158 |
| Mouse: STOCK Tg(Tlx3-cre | MMRRC | MMRRC: 041158 |
| Mouse: STOCK Tg(Tlx3-cre | MMRRC | MMRRC: 041158 |
| Mouse: STOCK Tg(Tlx3-cre | MMRRC | MMRRC: 041158 |
| Mouse: B6(Cg)-Cux2tm1.: | MMRRC | MMRRC: 031778 |
| Mouse: B6.129S2-Emx1tr | The Jackson Laboratory | JAX: 005628 |
| Mouse: B6;129S4-Ntrk1tr | MMRRC | MMRRC: 015500 |
| Mouse: STOCK Tg(Tlx3-cre | MMRRC | MMRRC: 041158 |
| Mouse: STOCK Tg(Rbp4-cı | MMRRC | MMRRC: 031125 |
| Mouse: B6.129S2-Emx1tr | The Jackson Laboratory | JAX: 005628 |
| Mouse: B6.FVB(Cg)-Tg(Nt | MMRRC | MMRRC: 030648 |
| Mouse: B6.FVB(Cg)-Tg(Nt | MMRRC | MMRRC: 030648 |
| Mouse: B6.129S2-Emx1tr | The Jackson Laboratory | JAX: 005628 |
| Mouse: B6.129S2-Emx1tr | The Jackson Laboratory | JAX: 005628 |
| Mouse: B6.FVB(Cg)-Tg(Nt | MMRRC | MMRRC: 030648 |
| Mouse: B6.Cg-Tg(A93003 | The Jackson Laboratory | JAX: 017346 |
| Mouse: STOCK Tg(Rbp4-cı | MMRRC | MMRRC: 031125 |
| Mouse: B6.Cg-Tg(A93003 | The Jackson Laboratory | JAX: 017346 |
| Mouse: STOCK Tg(Rbp4-cı | MMRRC | MMRRC: 031125 |
| Mouse: B6.Cg-Tg(Slc6a4-c | MMRRC | MMRRC: 031028 |
| Mouse: B6.FVB(Cg)-Tg(Nt | MMRRC | MMRRC: 030648 |
| Mouse: B6.Cg-Tg(A93003 | The Jackson Laboratory | JAX: 017346 |
| Mouse: STOCK Tg(Tlx3-cre | MMRRC | MMRRC: 041158 |
| Mouse: STOCK Tg(Tlx3-cre | MMRRC | MMRRC: 041158 |
| Mouse: B6(Cg)-Cux2tm1.: | MMRRC | MMRRC: 031778 |
| Mouse: B6.Cg-Gnb4tm1.1 | The Jackson Laboratory | JAX: 030159 |
| Mouse: STOCK Tg(Rbp4-cı | MMRRC | MMRRC: 031125 |
| Mouse: B6.129S2-Emx1tr | The Jackson Laboratory | JAX: 005628 |
| Mouse: B6.129S2-Emx1tr | The Jackson Laboratory | JAX: 005628 |
| Mouse: B6.129S2-Emx1tr | The Jackson Laboratory | JAX: 005628 |
| Mouse: STOCK Tg(Tlx3-cre | MMRRC | MMRRC: 041158 |
| Mouse: B6.Cg-Gnb4tm1.1 | The Jackson Laboratory | JAX: 029587 |
| Mouse: B6.Cg-Gnb4tm1.1 | The Jackson Laboratory | JAX: 029587 |
| Mouse: STOCK Tg(Colgalt | MMRRC | MMRRC: 036504 |
| Mouse: STOCK Tg(Rbp4-cı | MMRRC | MMRRC: 031125 |
| Mouse: B6.Cg-Gnb4tm1.1 | The Jackson Laboratory | JAX: 029587 |
| Mouse: B6.Cg-Gnb4tm1.1 | The Jackson Laboratory | JAX: 029587 |
| Mouse: B6.Cg-Gnb4tm1.1 | The Jackson Laboratory | JAX: 029587 |
| Mouse: B6.Cg-Gnb4tm1.1 | The Jackson Laboratory | JAX: 029587 |
| Mouse: B6.Cg-Gnb4tm1.1 | The Jackson Laboratory | JAX: 029587 |
| Mouse: B6.Cg-Gnb4tm1.1 | The Jackson Laboratory | JAX: 029587 |
| Mouse: B6.Cg-Ntng2tm1. | The Jackson Laboratory | JAX: 029588 |
| Mouse: B6.Cg-Ntng2tm1. | The Jackson Laboratory | JAX: 029588 |
| Mouse: B6.Cg-Gnb4tm1.1 | The Jackson Laboratory | JAX: 029587 |

Mouse: B6.Cg-Gnb4tm1.1The Jackson Laboratory   JAX: 029587
