## Supplementary Table 2 for "Regional and cell type-specific afferent and efferent projections of the mouse claustrum"

| image series id | CLA | dorsal to CLA | ventral to CLA | lateral to CLA | medial to CLA (CP) |
|---|---|---|---|---|---|
| 780505913 | 216 | 9 | 62 | 58 | 10 |
| 730766377 | 10 | 13 | 0 | 0 | 0 |
| 780508099 | 197 | 33 | 30 | 14 | 15 |
| 780510390 | 38 | 6 | 4 | 0 | 8 |
| 863469869 | 121 | 79 | 9 | 14 | 6 |

| total | % retrograde neurons in CLA |
|---|---|
| 355 | 60.8% |
| 23 | 43.5% |
| 289 | 68.2% |
| 56 | 67.9% |
| 229 | 52.8% |
