## Supplementary Table 3 for "Regional and cell type-specific afferent and efferent projections of the mouse claustrum"

**Supplementary Table 3. Summary of all mouse lines and injections into isocortical areas**

| | Mouse Lin | Generation Method | Type | Originating Lab (Donating Investigator) | Public Repository | Stock # | Layer | Projection Class |
|---|---|---|---|---|---|---|---|---|
| 1 | C57BL/6J | n/a | n/a | n/a | JAX | 000664 | L2-L6 | IT ET CT |
| 2 | Emx1-IRES | Knock-in | IRES | Kevin Jone | JAX | 005628 | L2-L6 | IT ET CT |
| 3 | Cux2-IRES | Knock-in | IRES | Ulrich Muel | MMRRC | 031778 | L2/3 | IT |
| 4 | Sepw1-Cre | Transgenic | BAC | Nathaniel H | MMRRC | 036190 | L2/3 | IT |
| 16 | Calb1-T2A | Knock-in | T2A | Allen Institu | JAX | 023531 | L2/3 | IT |
| 17 | Rasgrf2-T2 | Knock-in | T2A | Allen Institu | JAX | 022864 | L2/3 | IT |
| 18 | Grm2-Cre_ | Transgenic | BAC | Nathaniel H | MMRRC | 034611 | L2/3 | IT |
| 19 | Grp-Cre_K | Transgenic | BAC | Nathaniel H | MMRRC | 031183 | L2/3 | IT |
| 20 | Plxnd1-Cre | Transgenic | BAC | Nathaniel H | MMRRC | 036631 | L2/3 | IT |
| 30 | Htr2a-Cre_ | Transgenic | BAC | Nathaniel H | MMRRC | 031150 | L2/3, L5 | IT ET |
| 21 | Penk-IRES | Knock-in | IRES | Allen Institu | JAX | 025112 | L2/3, L6 | IT |
| 5 | Nr5a1-Cre | Transgenic | BAC | Brad Lowel | JAX | 006364 | L4 | IT |
| 6 | Scnn1a-Tg | Transgenic | BAC | Allen Institu | JAX | 009613 | L4/5 | IT |
| 7 | Rorb-IRES | Knock-in | IRES | Allen Institu | JAX | 023526 | L4/5 | IT |
| 22 | Slc18a2-Cr | Transgenic | BAC | Nathaniel H | MMRRC | 034814 | L4 | local |
| 23 | Rorb-IRES | Knock-in | IRES | Allen Institu | n/a | n/a | L4 | local |
| 8 | Rbp4-Cre_ | Transgenic | BAC | Nathaniel H | MMRRC | 031125 | L5 | IT ET |
| 25 | Drd3-Cre_ | Transgenic | BAC | Nathaniel H | MMRRC | 034610 | L5 | IT ET |
| 26 | Etv1-CreEF | Knock-in | Direct | Z. Josh Hu | JAX | 013048 | L5 | IT ET |
| 28 | Gng7-Cre_ | Transgenic | BAC | Nathaniel H | MMRRC | 031181 | L5 | IT ET |
| 29 | Gpr26-Cre | Transgenic | BAC | Nathaniel H | MMRRC | 033032 | L5 | IT ET |
| 32 | Trib2-2A-C | Knock-in | F2A | Allen Institu | JAX | 022865 | L5 | IT ET |
| 9 | Tlx3-Cre_F | Transgenic | BAC | Nathaniel H | MMRRC | 036547 | L5 | IT |
| 27 | Glt25d2-Cr | Transgenic | BAC | Nathaniel H | MMRRC | 036504 | L5 | IT |
| 10 | A93-Tg1-C | Transgenic | BAC | Allen Institu | JAX | 017346 | L5 | ET |
| 11 | Chrna2-Cre | Transgenic | BAC | Nathaniel H | MMRRC | 036502 | L5 | ET |
| 12 | Efr3a-Cre_ | Transgenic | BAC | Nathaniel H | MMRRC | 036660 | L5 | ET |
| 13 | Sim1-Cre_ | Transgenic | BAC | Nathaniel H | MMRRC | 031742 | L5 | ET |
| 24 | Chrnb4-Cre | Transgenic | BAC | Nathaniel H | MMRRC | 036503 | L5 | ET |
| 31 | Npr3-IRES | Knock-in | IRES | Allen Institu | JAX | 031333 | L5 | ET |
| 33 | Slc17a8-IR | Knock-in | IRES | Allen Institu | JAX | 028534 | L5 | local |
| 14 | Ntsr1-Cre_ | Transgenic | BAC | Nathaniel H | MMRRC | 030648 | L6 | CT |
| 15 | Syt6-Cre_K | Transgenic | BAC | Nathaniel H | MMRRC | 032012 | L6 | CT |
| 34 | Gnb4-IRES | Knock-in | IRES | Allen Institu | JAX | 030159 | L6 | local |
| 35 | Ntng2-IRES | Knock-in | IRES | Allen Institu | JAX | 029588 | L6 | IT |
| 36 | Oxtr-Cre_C | Transgenic | BAC | Nathaniel H | MMRRC | 036545 | L6b | IT / local |
| 37 | Ctgf-T2A-d | Knock-in | T2A | Allen Institu | JAX | 028535 | L6b | local |
| 38 | Calb2-IRES | Knock-in | IRES | Z. Josh Hu | JAX | 010774 | inh | local |
| 39 | Cort-T2A-C | Knock-in | T2A | Z. Josh Hu | JAX | 010910 | inh | local |
| 40 | Crh-IRES-C | Knock-in | IRES | Brad Lowel | n/a | n/a | inh | local |
| 41 | Erbb4-2A-C | Knock-in | F2A | Allen Institu | JAX | 012360 | inh | local |

| | | | | | | | | |
|---|---|---|---|---|---|---|---|---|
| **42** | Gad2-IRES | Knock-in | IRES | Z. Josh Hu | JAX | 010802 | **inh** | **local** |
| **43** | Htr3a-Cre_ | Transgenic | BAC | Nathaniel H | MMRRC | 036680 | **inh** | **local** |
| **44** | Nos1-CreE | Knock-in | Direct | Z. Josh Hu | JAX | 014541 | **inh** | **local** |
| **45** | Oxtr-T2A-C | Knock-in | T2A | Allen Institu | JAX | 031303 | **inh** | **local** |
| **46** | Pvalb-IRES | Knock-in | IRES | Silvia Arber | JAX | 008069 | **inh** | **local** |
| **47** | Pvalb-T2A- | Knock-in | T2A | Allen Institu | JAX | 021189 | **inh** | **local** |
| **48** | Sst-IRES-C | Knock-in | IRES | Z. Josh Hu | JAX | 013044 | **inh** | **IT** |
| **49** | Tac1-IRES | Knock-in | IRES | Allen Institu | JAX | 021877 | **inh** | **local** |
| **50** | Vip-IRES-C | Knock-in | IRES | Z. Josh Hu | JAX | 010908 | **inh** | **local** |
| | | | | | | | | **TOTAL** |

Note:

- (red) no axonal termnials in the CLA
- (grey) Axonal terminals in the CLA
- (blue) Axonal terminals in the CLA from injection site contamination

| prefrontal | | | | | | | | |
|---|---|---|---|---|---|---|---|---|
| FRP | ACAd | ACAv | PL | ILA | ORBl | ORBm | ORBvl | AId |
| 1 | 3 | 2 | 1 | 1 | 4 | 1 | 2 | 2 |
| | 5 | | 1 | | | 1 | | |
| 1 | 2 | 3 | 3 | 1 | 1 | 2 | 1 | 1 |
| | | 1 | | | 1 | | | |
| | 1 | 1 | | | 1 | | | |
| | | | | | 1 | | | |
| | 1 | 3 | 1 | | 1 | 1 | 1 | 1 |
| 1 | | 1 | 1 | | | 1 | 1 | 1 |
| | | | | | 1 | | | |
| | 2 | 2 | | | | | | |
| | 3 | 3 | 1 | 1 | 1 | 1 | 3 | 1 |
| | 1 | | | | | | | |
| | 2 | 1 | | | 1 | | | |
| | 1 | | | | | | | |
| | 2 | 1 | | | 1 | | | |
| | 1 | 1 | | | | | | |
| | 2 | | 1 | | | | 1 | 1 |
| | | 1 | 1 | | 1 | | | 1 |
| | 3 | 2 | 1 | 1 | 1 | | | |
| | 2 | 2 | | 2 | 1 | | 1 | 1 |
| | 2 | 2 | | | 1 | | | |
| | 1 | 1 | | | 1 | | | |
| | 2 | 1 | 1 | 1 | | | | 1 |
| 1 | 3 | 3 | | 1 | 1 | 2 | | 1 |
| | | | | 1 | 2 | | | |
| | | 1 | | | | | | |
| | 1 | | | | | | | |
| | 1 | | | | | | | |
| | 1 | | | | | 1 | | |
| | 2 | | | | | | | |

| | | | | | | | | |
|---|---|---|---|---|---|---|---|---|
| | 1 | | | | | | | |
| | 1 | | | | | | | |
| | | | | | 1 | | | |
| | | 1 | | | | | | |
| | 1 | | | | | | 1 | |
| | 1 | | | | | | | |
| 4 | 48 | 33 | 12 | 9 | 22 | 10 | 11 | 11 |

0 0

| lateral | | | | | | | | |
|---|---|---|---|---|---|---|---|---|
| AIv | AIp | GU | VISC | TEa | PERI | ECT | SSs | SSp-bfd |
| | | | 4 | | | 1 | 3 | 6 |
| | | | | | | | | 1 |
| | 1 | | | | | | 3 | 3 |
| | | | | | | | | 1 |
| | | | | | | | | 1 |
| | | | | | | | | 1 |
| | | | | | | | | 1 |
| | | | | | | | 1 | 2 |
| | | | | | | | 1 | 4 |
| | | | | | | | 1 | 1 |
| | | | | | | | 1 | 2 |
| 1 | 1 | 1 | | | | | 4 | 4 |
| | | | | | | | | 2 |
| | | | | | | | | 2 |
| | | | | | | | | 1 |
| | | | | | | | | 1 |
| | | | | | | | 1 | 3 |
| | 1 | 1 | | | | | 2 | 4 |
| | 1 | | | 1 | | | 2 | 1 |
| | | | | | | | 1 | 2 |
| | | | 1 | | | | 1 | 2 |
| | | | | 1 | | | 3 | 8 |
| | | | | | | | 2 | 2 |
| | 1 | | | | | | 2 | |
| 1 | 1 | | | | | | | |
| 1 | | | 1 | | | | 1 | |
| | | | | | | | | 1 |
| | | | | | | | | 1 |
| | | | | | | | | 1 |
| | | | | | | | 1 | |

| | | | | | | | | |
|---|---|---|---|---|---|---|---|---|
| | | | | | | | | 1 |
| | | | | | | | | 1 |
| | | | | | | | | 2 |
| | | | | | | | 1 | |
| 3 | 6 | 2 | 6 | 2 | 0 | 1 | 31 | 62 |

| | | |
|---|---|---|
| L2/3 | 0.001363 | 2.53E-07 |
| L4 IT | 0 | 0 |
| L4/5 IT | 0.000534 | 0 |
| L5 IT | 0.022841 | 0.005229 |
| L5 ET | 0 | 0 |
| L6 CT | 0 | 0 |
| L6 IT | | |

| somatomotor | | | | | | | | |
| --- | --- | --- | --- | --- | --- | --- | --- | --- |
| SSp-tr | SSp-ll | SSp-ul | SSp-un | SSp-n | SSp-m | MOp | MOs | VISal |
| 2 | 3 | 2 | | 3 | 5 | 8 | 9 | 1 |
| | | | | | | | | 1 |
| | 3 | 2 | | | 2 | 1 | 5 | 1 |
| | | | | 1 | | | 4 | |
| | | | | | | 1 | 1 | |
| 1 | | | | | | | 1 | |
| | | | | | | | 2 | |
| | | | | | | | 3 | |
| | | | | | | | 1 | |
| | | | | | | 1 | 2 | |
| | | | | 1 | | | 7 | |
| | 1 | | | 1 | 2 | 1 | 2 | |
| | | 2 | | 1 | 4 | 1 | | 2 |
| | 2 | | | 1 | 1 | | 4 | |
| 1 | 1 | 1 | | | 1 | | | |
| | | | | | | | 4 | |
| | 3 | 1 | | 2 | 1 | 1 | 7 | 3 |
| | | 1 | | | 1 | | | |
| | | | | | | | 2 | |
| | | | | 1 | | | | |
| 1 | 1 | | | 1 | | | 1 | |
| | 1 | | | | | | 1 | |
| 1 | 3 | 1 | | | 1 | 1 | 9 | 3 |
| | | | | | | | 2 | |
| 1 | 2 | 1 | | 2 | 1 | 2 | 5 | 2 |
| 1 | | 1 | | 1 | | 2 | 4 | |
| 1 | 1 | 1 | | 1 | 1 | 1 | 4 | |
| 1 | 1 | 1 | | 1 | 2 | 3 | 10 | |
| | | | | | | | 1 | |
| 1 | | | | 1 | | 1 | 1 | |
| | | | | | | | 1 | |
| 1 | 2 | | 1 | | 3 | 2 | 4 | 2 |
| | 1 | | | 2 | 2 | 3 | 6 | |
| | | | | 1 | | | | |
| | | | | | | | 1 | |
| | | | | | | | 1 | |
| | | | | | | | 1 | |
| | | | | 1 | | 1 | 1 | |
| | | | | 1 | | | 1 | |

| | | | | | | | | |
|---|---|---|---|---|---|---|---|---|
| | | | | | | | 1 | |
| | | | | 1 | | | 1 | |
| | | | | | | | 2 | |
| | | | 1 | | | | 1 | |
| | | | | | | | 2 | |
| | | | | | | | 1 | |
| | | | | 1 | | | 1 | |
| | | | | 1 | | | 2 | |
| 12 | 25 | 14 | 2 | 26 | 27 | 30 | 119 | 15 |

| visual | | | | | | me | | |
|---|---|---|---|---|---|---|---|---|
| VISl | VISp | VISpl | VISli | VISpor | VISrl | VISa | VISam | VISpm |
| 4 | 31 | 1 | 2 | 2 | | | 3 | 1 |
| 6 | 26 | | | 1 | 2 | | 2 | 4 |
| 4 | 11 | | 1 | 4 | 3 | | 4 | 1 |
| | 2 | | | | | 1 | | |
| | 1 | | | | | | | |
| | 8 | | | | | | | |
| 1 | 7 | | 1 | | 2 | | | |
| | 7 | | | 2 | 2 | | 2 | 4 |
| 2 | 6 | | | | | | | 2 |
| | 2 | | | | | | | |
| 2 | 8 | 2 | | 5 | 2 | 2 | 3 | 2 |
| 2 | 1 | | | 1 | 1 | | 1 | |
| | 1 | | | | | | | 1 |
| | 1 | | | | | | | |
| 1 | 2 | | | | | | 1 | |
| | 1 | | | | | | 1 | |
| 5 | 5 | 2 | 2 | 2 | 3 | 3 | 6 | 3 |
| 4 | 17 | | 1 | 2 | 2 | 2 | 1 | 1 |
| | 2 | | 2 | | | | | 1 |
| 3 | 3 | | | | | 1 | 1 | |
| | 2 | | | | 1 | | 1 | |
| | | | | | | 1 | | |
| | 1 | | | | | 1 | | |
| | 4 | | | | | | | |
| 4 | 16 | 1 | 1 | 6 | | 2 | 3 | 3 |
| 1 | 4 | 1 | 1 | 2 | 1 | | | 1 |
| 1 | | | | | | | | |
| | 1 | | | | | | | |
| | 4 | 1 | | | | | | |
| | 1 | | | | | | | |
| | 1 | | | | | | | |
| | 1 | | | | | | | |
| | 1 | | | | | | | |

| | | | | | | | | |
|---|---|---|---|---|---|---|---|---|
| | 1 | | | | | | | |
| | 1 | | | | | | | |
| | 3 | | | | | | | |
| | 2 | | | | | | | |
| | 1 | | | | | | | |
| | 2 | | | | | | | |
| | 1 | | | | | | 1 | |
| | 1 | | | | | | | |
| 40 | 190 | 8 | 11 | 27 | 19 | 13 | 30 | 24 |

| dial | | | auditory | | | | |
|---|---|---|---|---|---|---|---|
| RSPagl | RSPd | RSPv | AUDd | AUDp | AUDpo | AUDv | Expts |
| 1 | 2 | 3 | 2 | 4 | 1 | 1 | 122 |
| 2 | 2 | 3 | | 2 | 3 | | 62 |
| 3 | 1 | 3 | 1 | 2 | 3 | | 77 |
| | | 1 | 1 | | | | 13 |
| | | | | 1 | | | 4 |
| | | | | 1 | | | 7 |
| | | | | | | | 3 |
| | | | | | | | 12 |
| | | | | | | | 3 |
| | | | | | | | 9 |
| | | | | | | | 16 |
| | | | 1 | | | | 23 |
| | 1 | 2 | | 1 | | | 40 |
| | | | | 1 | | | 21 |
| | | | | | | | 7 |
| | | | | | | | 6 |
| 4 | 1 | 4 | | 1 | 2 | | 81 |
| | | 3 | | | | | 14 |
| | | 1 | 1 | | | | 12 |
| | | | | 1 | | | 4 |
| | 1 | 1 | | 1 | | | 16 |
| | | 2 | | 1 | | | 10 |
| 1 | 2 | | 1 | 1 | | | 64 |
| | | | | | | | 6 |
| 1 | 1 | 1 | | | 1 | | 66 |
| | | 3 | | | | | 31 |
| | 1 | 1 | | | | | 28 |
| | | | | 1 | | | 28 |
| | | 1 | | | | | 3 |
| | | 1 | 1 | | | | 11 |
| | | | | | | | 5 |
| 5 | 4 | | | 2 | 1 | | 81 |
| 1 | | 3 | | 1 | | | 46 |
| | | | | | | | 4 |
| | | | | | | | 3 |
| | | | | | | | 7 |
| | | | 1 | | | | 9 |
| | | | | 1 | | | 5 |
| | | | | 1 | | | 5 |
| | | | | 1 | | | 7 |
| | | | | | | | 6 |

| | | | | | | | |
|---|---|---|---|---|---|---|---|
| | | | | 1 | | | 5 |
| | | | | 1 | | | 5 |
| | | 1 | | 1 | | | 9 |
| | | | | | | | 2 |
| | | | 1 | | | | 5 |
| | | | | | | | 4 |
| | | 1 | | 1 | | | 9 |
| | | | | | | | 4 |
| | | | | 1 | | | 5 |
| 18 | 16 | 35 | 10 | 29 | 11 | 1 | 1025 |
