## Supplementary material for "Regional and cell type-specific afferent and efferent projections of the mouse claustrum": Supplimentary Table 4

| Mouse Lin | Generation Method | Type | Originating Lab (Donating Investigat | Public Repository | Stock # | | | |
|---|---|---|---|---|---|---|---|---|
| | | | | | | MOB | AOB | AON |
| A930038C( | Transgenic | BAC | Allen Institu | JAX | 017346 | 1 | | |
| Adcyap1-F: | Knock-in | F2A | Allen Institu | JAX | 030155 | | | |
| Agrp-IRES- | Knock-in | IRES | Bradford Lo | JAX | 012899 | | | |
| Avp-IRES2 | Knock-in | IRES | Allen Institu | JAX | 023530 | | | |
| C57BL/6J | n/a | n/a | n/a | JAX | 000664 | 1 | 2 | 3 |
| Calb1-T2A- | Knock-in | T2A | Allen Institu | JAX | 023531 | | | |
| Calb2-IRES | Knock-in | IRES | Z. Josh Hu | JAX | 010774 | | | 1 |
| Cart-IRES2 | Knock-in | IRES | Allen Institu | JAX | 028533 | | | |
| Cart-Tg1-C | Transgenic | BAC | Allen Institu | JAX | 009615 | | | |
| Cck-IRES-( | Knock-in | IRES | Z. Josh Hu | JAX | 012706 | | | |
| Cdhr1-Cre_ | Transgenic | BAC | Nathaniel H | MMRRC | 030952 | 3 | 1 | |
| Chat-IRES- | Knock-in | IRES | Bradford Lo | JAX | 006410 | | | |
| Chrna2-Cre | Transgenic | BAC | Nathaniel H | MMRRC | 036502 | | | 1 |
| Chrnb4-Cre | Transgenic | BAC | Nathaniel H | MMRRC | 036503 | | | |
| Cnnm2-Cre | Transgenic | BAC | Nathaniel H | MMRRC | 030951 | | | |
| Crh-IRES-( | Knock-in | IRES | Brad Lowel | n/a | n/a | | | |
| Crh-IRES-( | Knock-in | IRES | Z. Josh Hu | JAX | 012704 | | | |
| Ctgf-T2A-d | Knock-in | T2A | Allen Institu | JAX | 028535 | | | |
| Cux2-CreE | Knock-in | Direct | Ulrich Muel | MMRRC | 032779 | | | |
| Cux2-IRES | Knock-in | IRES | Ulrich Muel | MMRRC | 031778 | | | |
| Dbh-Cre_K | Transgenic | BAC | Nathaniel H | MMRRC | 032081 | | | |
| Dlg3-Cre_K | Transgenic | BAC | Nathaniel H | MMRRC | 032809 | | | |
| Drd1a-Cre | Knock-in | Direct | Richard Pa | n/a | n/a | | | |
| Drd2-Cre_E | Transgenic | BAC | Nathaniel H | MMRRC | 032108 | | | |
| Drd3-Cre_K | Transgenic | BAC | Nathaniel H | MMRRC | 034610 | | | |
| Drd3-Cre_K | Transgenic | BAC | Nathaniel H | MMRRC | 031741 | | | |
| Efr3a-Cre_ | Transgenic | BAC | Nathaniel H | MMRRC | 036660 | | 1 | |
| Emx1-IRES | Knock-in | IRES | Kevin Jone | JAX | 005628 | | | |
| Erbb4-T2A- | Knock-in | T2A | Allen Institu | JAX | 012360 | | | |
| Esr1-2A-Cr | Knock-in | F2A | David Ande | n/a | n/a | | | |
| Esr2-IRES2 | Knock-in | IRES | Allen Institu | JAX | 030158 | | | |
| Etv1-CreEF | Knock-in | Direct | Z. Josh Hu | JAX | 013048 | | | |
| Fezf1-T2A- | Knock-in | T2A | Allen Institu | JAX | 025110 | | | |
| Foxp2-IRE: | Knock-in | IRES | Richard Pa | JAX | 030541 | | | |
| Gabra6-IRE | Knock-in | IRES | William Wis | MMRRC | 015968 | | | |
| Gabrr3-Cre | Transgenic | BAC | Nathaniel H | MMRRC | 030709 | | | |
| Gad2-IRES | Knock-in | IRES | Z. Josh Hu | JAX | 010802 | | 1 | |
| Gal-Cre_KI | Transgenic | BAC | Nathaniel H | MMRRC | 031060 | | | |
| Glt25d2-Cr | Transgenic | BAC | Nathaniel H | MMRRC | 036504 | | | |
| Gnb4-IRES | Knock-in | IRES | Allen Institu | JAX | 029587 | | | |
| Gnrh1-Cre | Transgenic | BAC | Catherine D | JAX | 021207 | | | |
| Gpr26-Cre_ | Transgenic | BAC | Nathaniel H | MMRRC | 033032 | 1 | | |
| Grik4-Cre | Transgenic | BAC | Susumu To | JAX | 006474 | | | |

| | | | | | | | | |
|---|---|---|---|---|---|---|---|---|
| Grm2-Cre_ | Transgenic | BAC | Nathaniel H | MMRRC | 034611 | | | |
| Grp-Cre_K | Transgenic | BAC | Nathaniel H | MMRRC | 031183 | | | |
| Hcrt-Cre | Knock-in | Direct | Takeshi Sa | n/a | n/a | | | |
| Hdc-Cre_IN | Transgenic | BAC | Nathaniel H | MMRRC | 032079 | | | |
| Htr1a-IRES | Knock-in | IRES | Allen Institu | JAX | 030160 | | | |
| Htr2a-Cre_ | Transgenic | BAC | Nathaniel H | MMRRC | 031150 | | | 1 |
| Htr3a-Cre_ | Transgenic | BAC | Nathaniel H | MMRRC | 036680 | | | |
| Ins2-Cre_2 | Transgenic | Convention | Mark Magn | JAX | 003573 | | | |
| Kcnc2-Cre | Transgenic | Convention | Susumu To | JAX | 008582 | | | |
| Kcng4-Cre | Knock-in | Direct | Joshua Sar | JAX | 029414 | | | |
| Kiss1-Cre | Transgenic | BAC | Carol Elias | JAX | 023426 | | | |
| Lepr-IRES- | Knock-in | IRES | Jeffrey Frie | JAX | 008320 | | | |
| Lypd6-Cre_ | Transgenic | BAC | Nathaniel H | n/a | n/a | | | |
| Ndnf-IRES2 | Knock-in | IRES | Allen Institu | JAX | 028536 | | | |
| Nkx2-1-Cre | Knock-in | Direct | Z. Josh Hua | JAX | 014552 | | | |
| Nos1-CreE | Knock-in | Direct | Z. Josh Hua | JAX | 014541 | | | 1 |
| Npr3-IRES2 | Knock-in | IRES | Allen Institu | JAX | 031333 | | | |
| Nr5a1-Cre | Transgenic | BAC | Brad Lowel | JAX | 006364 | | | |
| Ntng2-IRES | Knock-in | IRES | Allen Institu | JAX | 029588 | | | |
| Ntrk1-IRES | Knock-in | IRES | Louis F. Re | MMRRC | 015500 | | | |
| Ntsr1-Cre_ | Transgenic | BAC | Nathaniel H | MMRRC | 030648 | | | |
| Nxph4-T2A | Knock-in | T2A | Allen Institu | JAX | 022861 | | | |
| Otof-Cre | Knock-in | Direct | Ulrich Muel | MMRRC | 032781 | | | |
| Oxt-IRES-C | Knock-in | IRES | Bradford Lo | n/a | n/a | | | |
| Oxtr-Cre_C | Transgenic | BAC | Nathaniel H | MMRRC | 036545 | | | |
| Oxtr-T2A-C | Knock-in | T2A | Allen Institu | JAX | 031303 | | | |
| Pcdh9-Cre_ | Transgenic | BAC | Nathaniel H | MMRRC | 036084 | | | |
| Pcp2-Cre_( | Transgenic | BAC | Nathaniel H | MMRRC | 030868 | | | |
| Pdyn-T2A-( | Knock-in | T2A | Allen Institu | JAX | 030197 | | | |
| Pdzk1ip1-C | Transgenic | BAC | Nathaniel H | MMRRC | 030851 | | | |
| Penk-F2A-( | Knock-in | F2A | Allen Institu | JAX | 022862 | | | |
| Penk-IRES | Knock-in | IRES | Allen Institu | JAX | 025112 | | | |
| Plxnd1-Cre | Transgenic | BAC | Nathaniel H | MMRRC | 036631 | | | |
| Pmch-Cre | Transgenic | BAC | Bradford Lo | JAX | 014099 | | | |
| Pnmt-Cre | Knock-in | Direct | Steven Ebe | n/a | n/a | | | |
| Pomc-Cre_ | Transgenic | Convention | Bradford Lo | JAX | 005965 | | | |
| Pomc-Cre_ | Transgenic | BAC | Susumu To | JAX | 010714 | | | |
| Ppp1r17-C | Transgenic | BAC | Nathaniel H | MMRRC | 036205 | | | |
| Prkcd-GluC | Knock-in | IRES | David Ande | MMRRC | 011559 | | | |
| Pvalb-IRES | Knock-in | IRES | Silvia Arber | JAX | 008069 | | | |
| Rasgrf2-T2 | Knock-in | T2A | Allen Institu | JAX | 022864 | | | |
| Rbp4-Cre_ | Transgenic | BAC | Nathaniel H | MMRRC | 031125 | | | |
| Rorb-IRES2 | Knock-in | IRES | Allen Institu | JAX | 023526 | | | |
| Satb2-Cre_ | Transgenic | BAC | Nathaniel H | MMRRC | 032908 | | | |
| Scnn1a-Tg | Transgenic | BAC | Allen Institu | JAX | 009112 | | | |
| Scnn1a-Tg | Transgenic | BAC | Allen Institu | JAX | 009613 | | | |
| Sim1-Cre_( | Transgenic | BAC | Nathaniel H | MMRRC | 031742 | | | |

| | | | | | | | | |
|---|---|---|---|---|---|---|---|---|
| Slc17a6-IR | Knock-in | IRES | Bradford Lo | JAX | 028863 | 1 | 1 | |
| Slc17a7-IR | Knock-in | IRES | Allen Institu | JAX | 023527 | | | 1 |
| Slc17a8-IR | Knock-in | IRES | Allen Institu | JAX | 028534 | | | |
| Slc18a2-Cr | Transgenic | BAC | Nathaniel H | MMRRC | 034814 | | | |
| Slc32a1-IR | Knock-in | IRES | Bradford Lo | JAX | 028862 | | | |
| Slc6a3-Cre | Knock-in | Direct | Xiaoxi Zhua | JAX | 020080 | 3 | | |
| Slc6a4-Cre | Transgenic | BAC | Nathaniel H | MMRRC | 031028 | | | |
| Slc6a4-Cre | Transgenic | BAC | Nathaniel H | MMRRC | 030071 | | | |
| Slc6a5-Cre | Transgenic | BAC | Nathaniel H | MMRRC | 030730 | | | |
| Sst-Cre | Knock-in | Direct | Allen Institu | n/a | n/a | | | |
| Sst-IRES-C | Knock-in | IRES | Z. Josh Hu | JAX | 013044 | | | |
| Syt17-Cre_ | Transgenic | BAC | Nathaniel H | MMRRC | 034355 | | 1 | 2 |
| Syt6-Cre_K | Transgenic | BAC | Nathaniel H | MMRRC | 032012 | 1 | | |
| Tac1-IRES | Knock-in | IRES | Allen Institu | JAX | 021877 | | | 1 |
| Tac2-IRES | Knock-in | IRES | Allen Institu | JAX | 021878 | | | |
| Tacr1-T2A- | Knock-in | T2A | Allen Institu | n/a | n/a | | | |
| Th-Cre_FI1 | Transgenic | BAC | Nathaniel H | MMRRC | 031029 | | | |
| Th-IRES-C | Knock-in | IRES | Jeremy Nat | JAX | 008532 | | | |
| Tlx3-Cre_P | Transgenic | BAC | Nathaniel H | MMRRC | 036547 | | | |
| Trib2-F2A-( | Knock-in | F2A | Allen Institu | JAX | 022865 | | | |
| Ucn3-Cre_ | Transgenic | BAC | Nathaniel H | MMRRC | 032078 | | | |
| Vip-IRES-C | Knock-in | IRES | Z. Josh Hu | JAX | 010908 | | | |
| Vipr2-Cre_ | Transgenic | BAC | Nathaniel H | MMRRC | 034281 | | | |
| Vipr2-IRES | Knock-in | IRES | Allen Institu | JAX | 031332 | | | |
| Wfs1-Tg2-( | Transgenic | BAC | Allen Institu | JAX | 009614 | | | |
| Wfs1-Tg3-( | Transgenic | BAC | Allen Institu | JAX | 009103 | | | |
| | | | | | **TOTAL** | **11** | **7** | **11** |

Notes: Red means no protections to the CLA
Gray means protections to the CLA
Blue means projection to the CLA from contaminated structures

| OLF | | | | | | | | |
|---|---|---|---|---|---|---|---|---|
| TT | DP | PIR | NLOT | COAa | COAp | PAA | TR | CA1 |
| | | 2 | 1 | | | | | |
| 1 | 1 | 7 | | | 1 | 1 | 1 | 6 |
| | | 1 | | | | | | |
| | | 1 | | | | | | |
| | | | | | | | | 1 |
| 1 | | | | | | | | |
| | | 3 | | | | | | |
| | | | | | 1 | | | |
| | | 1 | | | | | | 1 |
| | | | | | | | | 2 |
| | | | | | | | | 2 |
| | | 1 | | | | | | |
| | | 2 | | | | | | 1 |
| | | | | | 1 | | | |

| | | | | | | | | |
|---|---|---|---|---|---|---|---|---|
| | 1 | | | | | | | 1 |
| | | | | | | | | 1 |
| | | | | | | | | 1 |
| | | | | | | | | 4 |
| | | 1 | 1 | | 1 | | | |
| | | | 1 | | | | | 1 |

| | | | | | | | | |
|---|---|---|---|---|---|---|---|---|
| | | | 1 | | 1 | | | |
| | | | | | 1 | | | 2 |
| | | 1 | | | 1 | | | 1 |
| | 1 | 1 | | | | | | 2 |
| | | 1 | | | | | | 1 |
| | | | | | | | | 3 |
| 2 | 3 | 22 | 4 | 0 | 7 | 1 | 1 | 30 |

| | | | | | | HPF | | |
|---|---|---|---|---|---|---|---|---|
| CA2 | CA3 | DG | FC | IG | ENTl | ENTm | PAR | POST |
| | | 1 | | | 2 | | | |
| | 7 | 8 | | | | | 3 | |
| | | | | | | 1 | | |
| | | | | | 1 | | | |
| | | | | | 1 | | | |
| | | | | | | 4 | | |
| | | | | | 3 | | | |
| | 1 | | | | | | | |
| | | | | | 1 | 1 | | |
| | 1 | 1 | | | | | | |
| | | | | | | 1 | | |
| | 1 | 1 | | | | | | |
| | | 1 | | | | | | |
| | | | | | 1 | | | |
| | | | | | 2 | | | |
| | 4 | 4 | | | | 4 | | |

| | | | | | | | | |
|---|---|---|---|---|---|---|---|---|
| | | 4 | | | 1 | | | |
| | 4 | | | | | 1 | | |
| | | | | | 1 | | | |
| | | 1 | | | 2 | | | 1 |
| | 1 | | | | | | | |
| | | 1 | | | | | | |
| | 1 | 2 | | | | 1 | 1 | |
| | | | | | 1 | 1 | | |
| | | | | | 2 | | | |
| | 1 | 1 | | | | | | 1 |
| | | | | | | 2 | 1 | |
| | | | | | 1 | | | |
| | | 2 | | | | | | |
| | | 1 | | | | | | |
| 4 | 4 | 2 | | | | | | |
| 1 | 1 | | | | | | | |
| | | 3 | | | 3 | 1 | | |
| | | | | | | 1 | | |
| | | | | | | 2 | | |
| | | | | | 1 | | 1 | 2 |

| | | | | | | | | |
|---|---|---|---|---|---|---|---|---|
| | | | | | | | | 1 |
| 1 | 1 | 2 | | | | | | |
| | | | | | | 1 | | |
| 0 | 1 | 4 | | | 1 | 1 | | |
| | | | | | | 2 | | |
| | | | | | | | 1 | |
| **21** | **28** | **39** | **0** | **0** | **24** | **24** | **7** | **5** |

| | | | | | CTXsp | | | |
|---|---|---|---|---|---|---|---|---|
| PRE | SUB | ProS | HATA | APr | CLA | EPd | EPv | LA |
| | 1 | | | | | | | |
| 1 | 2 | 1 | | 1 | | 1 | | |
| | | | | | | | | 1 |
| | | 1 | | | | | | 1 |
| | 1 | | | | | | | |
| | | 1 | | | | | | |
| 1 | | | | | | 1 | | |
| | | | | | 1 | | | |
| | 1 | | | | | | | |

| | | | | | | | | |
|---|---|---|---|---|---|---|---|---|
| 1 | | | | | | | | |
| 1 | | | | | | 1 | | |
| | | | | | | | | 1 |
| | | | | | | | | 1 |
| 1 | 1 | | | | | | | |

| | | | | | | | | |
|---|---|---|---|---|---|---|---|---|
| | 1 | | | | | | | |
| | 1 | | | | | | | 1 |
| | 1 | | | | | | | |
| | 1 | | | | | | | |
| | 1 | | | | | | | |
| 1 | | | | | | | | |
| 6 | 11 | 3 | 0 | 1 | 1 | 3 | 0 | 5 |

| BLA | BMA | PA | CP | ACB | FS | OT | LSc | LSr |
|---|---|---|---|---|---|---|---|---|
| 3 | 4 | | 10 | 3 | | | 1 | 2 |
| | | | | | | | | 1 |
| | | | 5 | | | | | |
| 1 | | | | | | | | |
| | | | 4 | 2 | | | | |
| | | | 9 | 2 | | 1 | | |
| | | | | 1 | | | | |
| | | | 1 | | | | | 1 |
| | 1 | | | | | | | |
| | 1 | | | | | | | |
| | | | 1 | | | | | |
| | | | 2 | | | | | |
| | | | | | | | | 2 |
| | | | | 1 | | | | |

| | | | | | | | | |
|---|---|---|---|---|---|---|---|---|
| | | | | | | | 1 | |
| | | 1 | | | | | | |
| | | | 6 | 4 | | 1 | | |
| | | | 1 | | | | | |
| | | | 1 | | | | | |
| | | | 1 | | | | | |
| | | | 2 | | | | | |
| | | | 1 | | | | | 1 |
| | | | | | | | | 1 |
| | | 1 | | | | | | |
| | 1 | | | | | | | |

| | | | | | | | | |
|---|---|---|---|---|---|---|---|---|
| | | | 1 | | | | | 1 |
| | | | 1 | | | | 1 | |
| | | | | | | | 1 | 1 |
| | 1 | | | | | | | |
| | | | 1 | 1 | | | | |
| **4** | **8** | **2** | **47** | **14** | **0** | **2** | **4** | **10** |

| STR | | | | | | | | |
|---|---|---|---|---|---|---|---|---|
| LSv | SF | SH | AAA | BA | CEA | IA | MEA | GPe |
| | | | | | | | 1 | |
| 1 | 1 | | 1 | | 3 | | 3 | 1 |
| 1 | | | | | | | | |
| | | | | | | | 1 | |
| | | | | | | | 2 | |
| | | | | | 2 | | | |
| | | | | | 2 | | | |
| | | | | | | | | 1 |
| | | | | | 1 | | | |
| | | | | | | | 1 | |
| | | | | | 3 | | | |
| | | | | | | | 2 | |
| | | | | | 1 | | | |
| 1 | | | | | | | 1 | |

| | | | | | | | | |
|---|---|---|---|---|---|---|---|---|
| | | | | | | | 1 | 1 |
| | | | | | | | 1 | |
| | | | | | | | 1 | |
| | | | | | | | | 3 |
| | | | | | | | 1 | |
| | | | | | 1 | | | |
| | | | | | 1 | | | |
| | | | | | | | | 1 |
| | | | | | | | 4 | |

| | | | | | | | | |
|---|---|---|---|---|---|---|---|---|
| | | | | | | | 1 | |
| | | | | | 1 | | | |
| | | | | | | | 1 | |
| | | | | | 2 | | | |
| | | | | | 1 | | | |
| | | | | | 1 | | | |
| 3 | 1 | 0 | 1 | 0 | 19 | 0 | 21 | 7 |

| PAL | | | | | | | | |
|---|---|---|---|---|---|---|---|---|
| GPi | SI | MA | MS | NDB | TRS | BST | BAC | VAL |
| | 6 | | 1 | 1 | | 3 | | 1 |
| | | | | | 1 | | | |
| | | | | | | 1 | | |
| | 8 | | 1 | 3 | | | | |
| | 1 | | | | | | | |
| | | | | | | 1 | | |
| | 2 | | | | | 1 | | |
| | 1 | | | | | | | |
| | | | | | | 2 | | |
| | 2 | | 1 | | | | | |
| | | | 1 | | | | | |
| | | | | | | 1 | | |
| | | | | | | | | 1 |

| | | | | | | |
|---|---|---|---|---|---|---|
| 1 | | | | | | 2 |
| | | | | | | 1 |
| | 2 | | 1 | 2 | | |
| | | | 1 | | | |
| | 1 | | | | | |
| | | | | | | 2 |
| | | | | | | 1 |
| 1 | | | | | | 1 |
| | | | | | | 3 |

| | | | | | | | | |
|---|---|---|---|---|---|---|---|---|
| | 1 | | | | | | | 1 |
| 1 | | | | | | 1 | | |
| | | | | | 1 | | | |
| | | | | | | 1 | | |
| | | | 1 | | | | | |
| 3 | 24 | 0 | 7 | 6 | 2 | 21 | 0 | 3 |

| VM | VPL | VPLpc | VPM | VPMpc | PoT | SPFm | SPFp | SPA |
|---|---|---|---|---|---|---|---|---|
| 5 | | | 3 | | | | | |
| 1 | | | | | | | | |
| 1 | | | | | | | | |
| 1 | | | | | | | | |
| | 2 | | | | | | | |

| | | | | | | |
|---|---|---|---|---|---|---|
| | | | 1 | | | |
| | | | | | | 1 |
| | | | | | | 1 |
| 1 | | | 2 | | | |
| 1 | | | 1 | | | |
| | | | 1 | | | |
| | | | 1 | | | |

| | | | | | | | | |
|---|---|---|---|---|---|---|---|---|
| | | | 1 | | | | | |
| 10 | 2 | 0 | 10 | 0 | 0 | 2 | 0 | 0 |

| PP | MG | LGd | LP | PO | POL | SGN | AV | AM |
|---|---|---|---|---|---|---|---|---|
| | 1 | 1 | 2 | 3 | | 1 | 3 | 2 |
| | 1 | | 3 | | | | | |
| | | | | | | | 1 | |
| | | | | | | 1 | | |
| | | 1 | | | | | | |
| | 1 | 3 | | 1 | | | 1 | 2 |
| | | | | | | | 1 | |
| | | | 2 | 3 | | | | 1 |

| | | | | | | | | |
|---|---|---|---|---|---|---|---|---|
| | | | 1 | | | | | |
| | | | | | | | 1 | |
| | | 2 | | | | | | |
| | 1 | | 1 | | | | | |
| | | 3 | 4 | | | | 3 | 2 |
| | | | 1 | | | | | |
| | | 6 | | | | | | |

| | | | | | | | | |
|---|---|---|---|---|---|---|---|---|
| | | 3 | 1 | | | | | 1 |
| | | 1 | | | | | | |
| | | 3 | | 2 | | 1 | | |
| | | | | | | 1 | | |
| | | 2 | | | | | | |
| **0** | **4** | **25** | **15** | **9** | **0** | **4** | **10** | **8** |

| TH | | | | | | | | |
|---|---|---|---|---|---|---|---|---|
| AD | IAM | IAD | LD | IMD | MD | SMT | PR | PVT |
| | | | 3 | | 8 | 2 | | 3 |
| | | | 1 | | 2 | | | 2 |
| | | | | | 1 | | | |
| | | | | | 1 | | | |
| | | | | | | | | 1 |
| | | | | | 1 | | | |
| | | 1 | 2 | | 3 | | | 1 |
| | | | | | 2 | | | |

| | | | | | | | | |
|---|---|---|---|---|---|---|---|---|
| | | | | 1 | 4 | | | 1 |
| | | | | | 2 | | | |
| | | | | | 1 | | | 1 |
| | | | | | | | | 1 |
| | | | | | | | | 1 |
| | | | | | 2 | | | |
| | | | | | 2 | | | 1 |
| | | | 1 | | 4 | | | |
| | | | | | | | | 1 |

| | | | | | | | | |
|---|---|---|---|---|---|---|---|---|
| | | | 2 | | 5 | | | |
| 2 | | | | | | | | |
| | | | | 1 | 2 | | | |
| | | | | | 1 | | | |
| | | | | 1 | | | | |
| | | | | | 1 | 1 | | |
| 2 | 0 | 1 | 9 | 3 | 42 | 3 | 0 | 13 |

| PT | RE | Xi | RH | CM | PCN | CL | PF | PIL |
|---|---|---|---|---|---|---|---|---|
| | 1 | | | | | | | 1 |
| 3 | 4 | | | 2 | | | 2 | 1 |
| | 1 | | | | | | | |
| | 1 | | | 1 | | | | 2 |
| 1 | | | | | | | | |
| | 1 | | | | | | | |
| 2 | | | | | | | | |
| 3 | 1 | | | | | | | |
| | | | | | | | 1 | |
| | 1 | | | | | | | |

| | | | | | | | |
|---|---|---|---|---|---|---|---|
| | | | | 1 | 1 | | |
| | 1 | | | | | | |
| | | | | 1 | | | |
| | | | | 1 | | | |
| | 1 | | | | | | |
| | 1 | | | | | | |
| | | | | | | | 1 |

| | | | | | | | | |
|---|---|---|---|---|---|---|---|---|
| 2 | | | | | | | 2 | |
| 1 | | | | | | | | |
| | 1 | | | | | | | |
| | | | | | | | 1 | |
| **12** | **14** | **0** | **0** | **6** | **1** | **0** | **7** | **4** |

| RT | IGL | IntG | LGv | SubG | MH | LH | SO | ASO |
|---|---|---|---|---|---|---|---|---|
| 2 | | | 1 | | | 1 | | |
| | | | 1 | | | | | |
| | | | | | | 1 | | |
| 1 | | | | | | | | |
| | | | | | 1 | | | |
| 2 | | | | | | | | |
| 1 | | | | | | | | |

| | | | | | | |
|---|---|---|---|---|---|---|
| | | | | | 1 | 1 |
| | | | | | | 1 |
| 1 | | | | | | |
| 1 | | | | | | |
| | | | 1 | | | |
| | | | 1 | | | |
| | | | 1 | | | |
| 3 | | | | | | |

| | | | | | | | | |
|---|---|---|---|---|---|---|---|---|
| 2 | | | 1 | | | | | |
| 1 | | | 1 | | | | | |
| | | | | | 1 | 1 | | |
| | | | | | 1 | 1 | | |
| 14 | 0 | 0 | 7 | 0 | 4 | 6 | 0 | 0 |

| PVH | PVa | PVi | ARH | ADP | AVP | AVPV | DMH | MEPO |
|---|---|---|---|---|---|---|---|---|
| | | | 2 | | | | | |
| 3 | | | | | | | | |
| | | | | | | 1 | | |
| | | | | | | | 2 | |
| 1 | | | | 1 | | | | |
| | | | | | | | 1 | |
| 1 | | | | | | | 1 | |
| 1 | | | 2 | | | | 1 | |

| | | | | | | | |
|---|---|---|---|---|---|---|---|
| | | | 2 | | | | 1 |
| | | | 2 | | | | |
| | | | 2 | | | | 3 |
| | | | | | | | 1 |
| 1 | | | | | | | |
| | | | | | | | 1 |
| | | | | 1 | | | |
| 3 | | | | | | | |
| 1 | | | | | | | |
| | | | 3 | | | | |
| | | | 2 | | | | |

| | | | | | | | | |
|---|---|---|---|---|---|---|---|---|
| | | | | | | | 1 | |
| 1 | | | | | | | | |
| | | | 1 | | | | | |
| | | | 1 | | | | | |
| 1 | | | | | | | | |
| 13 | 0 | 0 | 17 | 2 | 0 | 1 | 12 | 0 |

| MPO | OV | PD | PS | PVp | PVpo | SBPV | SCH | SFO |
|---|---|---|---|---|---|---|---|---|
| | | | | 1 | | | | |
| | | | | | | | 1 | |
| 5 | | | | | | | | |
| 1 | | | | 1 | | | | |
| 1 | | | | 1 | | | | |

| | | | | | |
|---|---|---|---|---|---|
| 1 | | | | | |
| 1 | | | | | |
| 1 | | | | | |
| | | | | | 1 |

| | | | | | | | | |
|---|---|---|---|---|---|---|---|---|
| 1 | | | | | | | | |
| | | | | 1 | | | | |
| | | | | | | | 1 | |
| **11** | **0** | **0** | **0** | **4** | **0** | **1** | **2** | **0** |

| HY | | | | | | | | |
|---|---|---|---|---|---|---|---|---|
| VMPO | VLPO | AHN | LM | MM | SUM | TMd | TMv | MPN |
| | | | | 2 | | | | 1 |
| | | 6 | | 4 | 1 | | | 3 |
| | | | | | | | | 1 |
| | | | | 1 | | | | |
| | | 2 | | | | | | 2 |
| | | | | | | | | 1 |
| | | | | | | | | 1 |

| | | | | | | | | |
|---|---|---|---|---|---|---|---|---|
| | | | | 2 | | | | |
| | | | 2 | | 1 | | 1 | |
| | | 1 | | | | | | |
| | | | | | | | | 1 |
| | | 1 | 1 | | | | | |
| | | | | 1 | 2 | | | |
| | | | | 1 | | | | |
| | | 1 | | | | | | |
| | | 2 | | | | | | 1 |
| | | 1 | | | | | | |
| | | 3 | | 1 | | | | |

| | | | | | | | | |
|---|---|---|---|---|---|---|---|---|
| | | 1 | | | | | | |
| | | 1 | | | | | | |
| | | | | | | | | 1 |
| **0** | **0** | **19** | **3** | **12** | **4** | **0** | **1** | **12** |

| PMd | PMv | PVHd | VMH | PH | LHA | LPO | PST | PSTN |
|---|---|---|---|---|---|---|---|---|
| | | | | | | | | 1 |
| | | | 2 | 1 | | | | |
| | 1 | | | | 1 | | | |
| | 1 | | 4 | | 6 | 1 | | |
| | | | | 1 | 1 | | | |
| | | | | 1 | | | | |
| | | | | 1 | | | | |
| | | | | | | 1 | | |
| | | 1 | | | | | | |
| | | | 2 | | | | | |
| | | | 2 | | 1 | | | |
| | | | 2 | | 1 | | | |
| | | | | 1 | | 1 | | |
| | | | | 1 | | | | |

| | | | | | |
|---|---|---|---|---|---|
| | | | | 1 | 2 |
| | 2 | | | | |
| | | | | 1 | |
| | 1 | | 1 | | |
| | 2 | | | | 1 |
| | | | | | 1 |
| | | | 2 | | |
| | | 1 | | | |
| | | | 2 | | |
| | | 2 | | | 1 |
| | | | | 1 | |
| | 1 | | 1 | | |
| | | | | 1 | |
| | | | | | 1 |
| | | | | | 4 |
| | | | 1 | 3 | 3 |
| | | | 2 | | 1 |
| | | | | 3 | 5 |

| | | | | | | | | |
|---|---|---|---|---|---|---|---|---|
| | | | 2 | 2 | | 1 | | |
| | | | 1 | | 2 | | | |
| | 1 | | | | | | | |
| | 2 | 1 | | | | | | |
| | | | | 1 | 1 | | | |
| | | | | | | | | 1 |
| 0 | 11 | 5 | 24 | 19 | 32 | 4 | 0 | 2 |

| PeF | RCH | STN | TU | ZI | ME | SCs | IC | NB |
|---|---|---|---|---|---|---|---|---|
| | | | | | | | 1 | |
| | | | 1 | | | | | |
| | | 1 | | 4 | | 2 | 4 | |
| | | | | | | | 1 | |
| | | | | 1 | | | | |
| | | | 3 | | | | | |
| | | 1 | 2 | | | | | |
| | | | | 2 | | | | |
| | | | | 3 | | 1 | | |

| | | | | | | | |
|---|---|---|---|---|---|---|---|
| | | 1 | | | | | |
| | | | | 1 | | | |
| | | | | | | | 1 |
| | | | | | | | 1 |
| | | | 1 | | | | |
| | | | | | | 1 | 2 |
| | | | | | | 1 | |
| | | | | 1 | | | 1 |
| | | | | | | 3 | |
| | | | | 1 | | | |
| | | | 1 | | | | |

| | | | | | | | | |
|---|---|---|---|---|---|---|---|---|
| | | | | | | | 2 | |
| | | | | 2 | | | | |
| | | | 1 | 1 | | | 1 | |
| 0 | 0 | 3 | 9 | 16 | 0 | 8 | 14 | 0 |

| SAG | PBG | MEV | SCO | SNr | VTA | PN | RR | MRN |
|---|---|---|---|---|---|---|---|---|
| 1 | | | | 3 | 2 | | | 16 |
| | | | | | 1 | | | |
| | | | | | 1 | | | |
| | | | | | 1 | | | |
| | | | | | 2 | | | 1 |
| | | | | 1 | | | | 2 |

| | | | | | | | | |
|---|---|---|---|---|---|---|---|---|
| | | | | | 1 | | | |
| | | | | | | | | 1 |
| | | | | | 1 | | | 1 |
| | | | | 1 | | | | 1 |
| | | | | 2 | 1 | | | |

| | | | | | | | | |
|---|---|---|---|---|---|---|---|---|
| 1 | | | | | | | | |
| | | | | | 1 | | | |
| | | | | | 2 | | | |
| | | | | | 2 | | | |
| | | | | | 1 | | | 2 |
| | | | | | 1 | | | |
| **2** | **0** | **0** | **0** | **7** | **17** | **0** | **0** | **24** |

| | | | | | | | | |
|---|---|---|---|---|---|---|---|---|
| | | | | | | | MB | |
| SCm | PAG | APN | MPT | NOT | NPC | OP | PPT | CUN |
| | 1 | | | | | | | |
| 3 | 9 | 4 | | 1 | | 1 | | 1 |
| | 1 | | | | | | | |
| | 2 | | | | | | | |
| | 1 | | | | | | | |
| | | 1 | | | | | | |
| | 1 | | | | | | | |
| | 2 | 2 | | | 1 | | | |

| | | | | | | | |
|---|---|---|---|---|---|---|---|
| | 1 | | | | | | |
| | | 1 | | | | | |
| | | 1 | | | | | |
| | 1 | | | | | | |
| | 1 | | | | | | |
| 1 | | | | | | | |
| | 1 | | | | | | |
| 2 | | 2 | | | 1 | | 1 |
| 5 | 1 | | | | | | |
| 1 | 2 | | | | | | |
| 1 | | | | | | | |
| | 2 | | | | | | |

| | | | | | | | | |
|---|---|---|---|---|---|---|---|---|
| 1 | | 1 | | | 1 | | | |
| | 1 | | | | | | | |
| | | 1 | | | | | | |
| | 1 | | | | | | | |
| 1 | | | | | | | | |
| | 2 | | | | | | | |
| | 1 | | | | | | | |
| | | | | | | | | 1 |
| | 2 | | | | | | | |
| | 1 | | | | | | | |
| | 1 | | | | | | | |
| **15** | **35** | **13** | **0** | **1** | **3** | **1** | **1** | **2** |

| RN | III | MA3 | EW | IV | Pa4 | VTN | AT | LT |
|---|---|---|---|---|---|---|---|---|
| 1 | | | | | | | | |
| | 1 | | | | | | | |

| | |
|---|---|
| | 1 |
| 1 | |
| 1 | |

| | | | | | | | | |
|---|---|---|---|---|---|---|---|---|
| 3 | 2 | 0 | 0 | 0 | 0 | 0 | 0 | 0 |

| DT | MT | SNc | PPN | IF | IPN | RL | CLI | DR |
|---|---|---|---|---|---|---|---|---|
| | | | | | 1 | | | |
| | | | | | | | 2 | |
| | | | | | | | | 1 |
| | | | 1 | | | | | |
| | | | | | 1 | | | |
| | | | | | 1 | | | |
| | | | | | | | | 1 |
| | | | | | 1 | | 1 | |

| | | | | | |
|---|---|---|---|---|---|
| | | | 1 | | |
| | | | | | 1 |
| | | | | | 1 |
| | | | 1 | | |

| | | | | | | | | |
|---|---|---|---|---|---|---|---|---|
| | | | | | | | 1 | |
| | | | | | | | | 1 |
| | | | | 1 | | | | |
| | | | | | 1 | | | |
| | | | | | | | | 4 |
| | | | | | | | | 2 |
| 0 | 0 | 0 | 3 | 1 | 7 | 0 | 4 | 9 |

| NLL | PSV | PB | SOC | B | DTN | PDTg | PCG | PG |
|---|---|---|---|---|---|---|---|---|
| | | 2 | | | | | | |
| 2 | 2 | 5 | | | | | 2 | |
| | | 1 | | | | | | |
| | | 2 | | | | | | |
| | | | | | | | 2 | |
| | | 1 | | | | | | |
| | | 1 | | | | | | |
| | | 1 | | | | | | |

| | | | | | | | |
|---|---|---|---|---|---|---|---|
| | | 1 | | | | | |
| | | 1 | | | | | |
| | | 1 | | | | | |
| | | 1 | | | | | |
| | 1 | | | | | | |
| | | 1 | | | | | |
| | | | | | | | 2 |
| | | 2 | 1 | | | | |
| | | 1 | | | | | |
| 1 | | | | | | | |
| | | 1 | | | | | |
| 1 | | | | | | | |

| | | | | | | | | |
|---|---|---|---|---|---|---|---|---|
| | | 2 | | | | | | |
| 1 | | | | | | | | |
| 1 | | | | | | | | |
| | 1 | | | | | | | |
| | | 1 | | | | | | |
| | | 1 | | | | | | |
| | | 2 | | | | | | |
| | | 1 | | | | | | |
| 6 | 4 | 29 | 1 | 0 | 0 | 0 | 6 | 0 |

| P | | | | | | | | |
|---|---|---|---|---|---|---|---|---|
| PRNc | SG | SUT | TRN | V | P5 | Acs5 | PC5 | I5 |
| 6 | | | 2 | | | | | |
| | | | | 1 | | | | |
| 1 | | | | | | | | |
| 1 | | | | | | | | |
| | | | 2 | | | | | |
| | | 1 | | | | | | |

| | | | | |
|---|---|---|---|---|
| 1 | | | | |
| | | | | 1 |
| 1 | | | | 1 |
| | | | | 1 |

| | | | | | | | | |
|---|---|---|---|---|---|---|---|---|
| 1 | | | | | | | | |
| **11** | **0** | **1** | **4** | **4** | **0** | **0** | **0** | **0** |

| CS | LC | LDT | NI | PRNr | RPO | SLC | SLD | AP |
|---|---|---|---|---|---|---|---|---|
| 1 | | | | 3 | | | | |
| | | 1 | | | | | | |
| 1 | | | | | | | 1 | |
| | | 2 | | | | | | |
| 1 | | | | | | | | |

| | | |
|---|---|---|
| | | 1 |

| | | | | | | | | |
|---|---|---|---|---|---|---|---|---|
| | | | | 1 | | | | |
| 1 | | | 1 | | | | | |
| 1 | | | | | 1 | | | |
| 1 | | | | | | | | |
| | | | | 1 | | | | |
| | | | | | | | 1 | |
| 6 | 0 | 4 | 1 | 5 | 1 | 0 | 2 | 0 |

| DCO | VCO | CU | GR | ECU | NTB | NTS | SPVC | SPVI |
|---|---|---|---|---|---|---|---|---|
| | | | 1 | | | | 1 | |
| | 3 | | | | | 1 | 3 | 2 |
| | | | | | | | 1 | |
| 2 | | | | | | | | |
| | | | 1 | | | | | |
| 1 | | | | | | 1 | | |
| | | | | | | 1 | | |

| | | | | | | |
|---|---|---|---|---|---|---|
| | | | | | 1 | 1 |
| | 1 | | | | | |
| 1 | | | | | | 1 |
| | | | | | | 1 |
| 1 | | | | | | |
| | 1 | | | | | |
| | | | | | | 1 |

| | | | | | | | | |
|---|---|---|---|---|---|---|---|---|
| 1 | | | | | | | | |
| | | | | | | 1 | | |
| | | | | | | | 1 | |
| 1 | | | | | | 1 | 1 | 2 |
| | | | 1 | | | | | |
| | | | | | | | 1 | |
| | | | | | | 1 | | |
| | | | | | | | 1 | |
| **7** | **5** | **0** | **3** | **0** | **1** | **10** | **9** | **4** |

| SPVO | Pa5 | VI | VII | ACVII | AMB | DMX | GRN | ICB |
|---|---|---|---|---|---|---|---|---|
| | | | | | | | 1 | |
| 1 | | | 2 | | | | 6 | |
| | | 1 | 3 | | | | | |
| | | | | | | | 1 | |
| | | | | | | | 1 | |
| | | | 1 | | | | | |

| | | | | | | |
|---|---|---|---|---|---|---|
| | | | | | 1 | |
| | | | | | | 1 |
| | | | 1 | | | |
| | | | 1 | | | |

| | | | | | | | | |
|---|---|---|---|---|---|---|---|---|
| | | | | | | | 1 | |
| | | | | | | 1 | | |
| | | | | | | 1 | | |
| | | | | | | | 3 | |
| | | | | | | 1 | | |
| **1** | **0** | **1** | **8** | **0** | **0** | **4** | **14** | **0** |

| MY | | | | | | | | |
|---|---|---|---|---|---|---|---|---|
| IO | IRN | ISN | LIN | LRN | MARN | MDRN | MDRN d | MDRN v |
| | 6 | | | | | 4 | | |
| | 1 | | | | | | | |
| 1 | | | | 1 | | | | |
| | | | | 1 | | 1 | | |
| | | | | | | 1 | | |
| | | | | 1 | | | |
| 1 | | | | | | | | |

| | | | | | |
|---|---|---|---|---|---|
| | | | | | 1 |
| | | | | 1 | |

| | | | | | | | | |
|---|---|---|---|---|---|---|---|---|
| | 1 | | | | | 1 | | |
| | | | | | | 1 | | |
| | 1 | | | | 2 | | | |
| | 1 | | | | | | | |
| | | | | 1 | | | | |
| 2 | 10 | 0 | 0 | 4 | 4 | 8 | 0 | 0 |

| PARN | PAS | PGRN d | PGRN l | NR | PRP | PPY | LAV | MV |
|---|---|---|---|---|---|---|---|---|
| | | | | | | | | 1 |
| 2 | | | 4 | | 1 | | | 4 |
| | | | 1 | | | | | |
| | | | | | | | | 1 |
| 1 | | | 1 | | | | | 1 |
| 2 | | | | | | | | |
| | | | 2 | | | | | |

| | | | | | | | | |
|---|---|---|---|---|---|---|---|---|
| | | | | | | | | 1 |
| 1 | | | | | | | | 1 |
| | | | 1 | | | | | |
| | | | 1 | | | | | |

| | | | | | | | | |
|---|---|---|---|---|---|---|---|---|
| 1 | | | | | | | | 1 |
| | | | | | | | | 1 |
| | | | | | | | | 1 |
| 7 | 0 | 0 | 10 | 0 | 1 | 0 | 0 | 12 |

| SPIV | SUV | x | XII | y | RM | RPA | RO | LING |
|---|---|---|---|---|---|---|---|---|
| 3 | | | 1 | | | | | |
| | | | 3 | | | | | |
| | | | 1 | | | | | |

| | | | |
|---|---|---|---|
| | | | 1 |

| | | | | | | | | |
|---|---|---|---|---|---|---|---|---|
| | | | | | | | 1 | |
| | | | | | | | 2 | |
| | | | | | | | 2 | |
| | | | 1 | | | | | |
| 3 | 0 | 0 | 7 | 0 | 0 | 0 | 5 | 0 |

| CB | | | | | | | | |
|---|---|---|---|---|---|---|---|---|
| CENT | CUL | DEC | FOTU | PYR | UVU | NOD | SIM | AN |

| | | | | | | | | |
|---|---|---|---|---|---|---|---|---|
| 0 | 0 | 0 | 0 | 0 | 0 | 0 | 0 | 0 |

| PRM | COPY | PFL | FL | FN | IP | DN | VeCB | Expts |
|---|---|---|---|---|---|---|---|---|
| | | | | | | | | 18 |
| | | | | | | | | 18 |
| | | | | | | | | 3 |
| | | | | | | | | 7 |
| | | | | 1 | 2 | | | 301 |
| | | | | | | | | 4 |
| | | | | | | | | 25 |
| | | | | | | | | 4 |
| | | | | | | | | 10 |
| | | | | | | | | 9 |
| | | | | | | | | 4 |
| | | | | | | | | 33 |
| | | | | | | | | 7 |
| | | | | | | | | 1 |
| | | | | | | | | 2 |
| | | | | | | | | 15 |
| | | | | | | | | 9 |
| | | | | | | | | 1 |
| | | | | | | | | 8 |
| | | | | | | | | 5 |
| | | | | | | | | 8 |
| | | | | | | | | 5 |
| | | | | | | | | 8 |
| | | | | | | | | 13 |
| | | | | | | | | 12 |
| | | | | | | | | 2 |
| | | | | | | | | 8 |
| | | | | | | | | 1 |
| | | | | | | | | 12 |
| | | | | | | | | 6 |
| | | | | | | | | 2 |
| | | | | | | | | 7 |
| | | | | | | | | 5 |
| | | | | | | | | 5 |
| | | | | | | | | 0 |
| | | | | | 1 | | | 7 |
| | | | | | | 1 | | 52 |
| | | | | | | | | 30 |
| | | | | | | | | 1 |
| | | | | | | | | 2 |
| | | | | | | | | 3 |
| | | | | | 1 | 1 | | 11 |
| | | | | | | | | 27 |

| | | | | | | | | |
|---|---|---|---|---|---|---|---|---|
| | | | | | | | | 19 |
| | | | | | | | | 12 |
| | | | | | | | | 3 |
| | | | | | | | | 7 |
| | | | | | | | | 2 |
| | | | | | | 1 | | 20 |
| | | | | | | | | 5 |
| | | | | | | | | 5 |
| | | | | | | | | 4 |
| | | | | | | | | 3 |
| | | | | | | | | 4 |
| | | | | | | | | 8 |
| | | | | | | | | 12 |
| | | | | | | | | 1 |
| | | | | | | | | 3 |
| | | | | | | | | 20 |
| | | | | | | | | 2 |
| | | | | | | | | 3 |
| | | | | | | | | 2 |
| | | | | | | | | 29 |
| | | | | | 1 | | | 2 |
| | | | | | 1 | | | 8 |
| | | | | | | | | 2 |
| | | | | | | | | 7 |
| | | | | | | | | 9 |
| | | | | | | | | 5 |
| | | | | | | | | 4 |
| | | | | | | | | 0 |
| | | | | | | | | 4 |
| | | | | | | | | 15 |
| | | | | | | | | 1 |
| | | | | | | | | 2 |
| | | | | | | | | 10 |
| | | | | | | | | 4 |
| | | | | | | | | 4 |
| | | | | | | | | 5 |
| | | | | | | | | 3 |
| | | | | | | | | 44 |
| | | | | | | | | 26 |
| | | | | | | | | 18 |
| | | | | | | | | 12 |
| | | | | | | | | 18 |
| | | | | | | | | 4 |
| | | | | | | | | 1 |
| | | | | | | | | 15 |
| | | | | | | | | 7 |
| | | | | | | | | 23 |

| | | | | | | | | |
|---|---|---|---|---|---|---|---|---|
| | | | | | 1 | 1 | | 52 |
| | | | | | | | | 9 |
| | | | | | | | | 6 |
| | | | | | | | | 19 |
| | | | | | 1 | | | 21 |
| | | | | | | | | 13 |
| | | | | | | | | 5 |
| | | | | | | | | 2 |
| | | | | | | | | 15 |
| | | | | | | | | 4 |
| | | | | | | | | 14 |
| | | | | | | | | 9 |
| | | | | | | | | 6 |
| | | | | | | | | 13 |
| | | | | | | | | 7 |
| | | | | | | | | 5 |
| | | | | | | | | 12 |
| | | | | | | | | 1 |
| | | | | | | | | 1 |
| | | | | | | | | 1 |
| | | | | | | | | 4 |
| | | | | | | | | 4 |
| | | | | | | | | 8 |
| | | | | | | | | 2 |
| | | | | | | | | 2 |
| | | | | | | | | 2 |
| **0** | **0** | **0** | **0** | **1** | **8** | **4** | **0** | **1365** |
