## Supplementary Table 5 for "Regional and cell type-specific afferent and efferent projections of the mouse claustrum"

| ID | Cell Name | unsupervised cluster |
|---|---|---|
| 1 | 17781_4698_X17857_Y11456.swc | 7 |
| 2 | 236174_3729_x13645_y9551.semi_r.swc | 7 |
| 3 | 236174_3829_x16301_y26647.semi_r.swc | 7 |
| 4 | 17782_3651_x35286_y18512.semi_r.swc | 1 |
| 5 | 17781_3320_X16423_Y14748.swc | 1 |
| 6 | 17781_3668_X17825_Y13313.swc | 1 |
| 7 | 236174_3215_x11999_y11133.semi_r.swc | 1 |
| 8 | 236174_3970_x13439_y8678.semi_r.swc | 1 |
| 9 | 236174_3229_X12413_Y11831.swc | 4 |
| 10 | 236174_3276-x12423-y11836_xy25z25.swc | 4 |
| 11 | 236174_3729_x12692_y9419.semi_r.swc | 4 |
| 12 | 236174_3529_x12805_y10541.semi_r.swc | 5 |
| 13 | 236174_4229_x13663_y8589.semi_r.swc | 5 |
| 14 | 236174_3425-x14960-y25777_xy25z25.swc | 5 |
| 15 | 236174_3929_x13599_y9165.semi_r.swc | 5 |
| 16 | 236174_4429_x13147_y8003.semi_r.swc | 5 |
| 17 | 236174_3729_x15151_y26698.semi_r.swc | 5 |
| 18 | 236174_4243-x16082-y27546_xy25z25.swc | 5 |
| 19 | 236174_4891-x13696-y7671_xy25z25.swc | 5 |
| 20 | 236174_4288-x15993-y27586_xy25z25.swc | 4 |
| 21 | 236174_3590-x15778-y26936_xy25z25.swc | 3 |
| 22 | 236174_4029_x13079_y8858.semi_r.swc | 3 |
| 23 | 236174_3915-x15856-y26875_xy25z25.swc | 3 |
| 24 | 236174_4445-x13724-y8174_xy25z25.swc | 3 |
| 25 | 236174_3442-x15385-y25403_xy25z25.swc | 3 |
| 26 | 236174_4709-x16524-y27926_xy25z25.swc | 3 |
| 27 | 236174_3732-x13018-y9645_xy25z25.swc | 2 |
| 28 | 236174_4406-x13402-y8151_xy25z25.swc | 2 |
| 29 | 236174_5220-x13868-y7456_xy25z25.swc | 2 |
| 30 | 236174_5496-x13847-y7315_xy25z25.swc | 2 |
| 31 | 236174_3829_x13590_y9284.semi_r.swc | 6 |
| 32 | 17781_4095_x17570_y12460.semi_r.swc | 6 |
| 33 | 236174_4729_x16869_y27809.semi_r.swc | 6 |
| 34 | 236174_3729_x15443_y26410.semi_r.swc | 6 |
| 35 | 236174_4113-x16041-y27245_xy25z25.swc | 6 |
| 36 | 236174_3757-x15816-y26363_xy25z25.swc | 6 |
| 37 | 236174_4283-x13211-y8428_xy25z25.swc | 6 |
| 38 | 236174_4647_x16405_y27845.semi_r.swc | 6 |
| 39 | 236174_4266_x13848_y8550.semi_r.swc | 6 |
| 40 | 236174_5138_x16501_y28259.semi_r.swc | 6 |
| 41 | 236174_5338_x13590_y7348.semi_r.swc | 6 |
| 42 | 236174_4315-x16369-y27510_xy25z25.swc | 6 |
| 43 | 17109_2401_x9695_y9693.semi_r.swc | 9 |
| 44 | 236174_3429_x12632_y10625.semi_r.swc | 9 |

| | | |
|---|---|---|
| 45 | 17109_2601_x10213_y8783.semi_r.swc | 9 |
| 46 | 17109_3101_x10824_y7188.semi_r.swc | 9 |
| 47 | 236174_3447_x12562_y10626.semi_r.swc | 9 |
| 48 | 17782_3672_x34784_y19432.semi_r.swc | 8 |
| 49 | 17782_3284_x11909_y16428.semi_r.swc | 8 |
| 50 | 17782_3487_x11014_y17041.semi_r.swc | 8 |
| 51 | 236174_3536_x15159_y25525.semi_r.swc | 8 |
| 52 | 236174_3328_X11950_Y11335.swc | 8 |

| mannually annotated cluster | Morphological fearure | |
|---|---|---|
| 7 | Ipsilateral features | |
| 7 | 1 | transverse difussing length ratio from ACA & RSP |
| 7 | 2 | axon length ratio from the left side of the soma |
| 7 | 3 | axon length ratio from the right side of the soma |
| 1 | 4 | axon length ratio from the anterior side of the soma |
| 1 | 5 | anterior projection ratio |
| 1 | 6 | posterior projecttion ratio |
| 1 | 7 | posterior projecting path efficiency |
| 4 | 8 | VISl-VISpl-ENT axon length ratio |
| 4 | 9 | VISl-VISpl-ENT cluster width |
| 4 | 10 | cluster width extended from RSP |
| 5 | 11 | crown forming axon length |
| 5 | | |
| 5 | | |
| 5 | Bilateral features | |
| 5 | 12 | Projections to RSP |
| 5 | 13 | Axon length ratio in RSP |
| 5 | | |
| 5 | | |
| 5 | | |
| 3 | | |
| 3 | | |
| 3 | | |
| 3 | | |
| 3 | | |
| 3 | | |
| 2 | | |
| 2 | | |
| 2 | | |
| 2 | | |
| 6 | | |
| 6 | | |
| 6 | | |
| 6 | | |
| 6 | | |
| 6 | | |
| 6 | | |
| 6 | | |
| 6 | | |
| 6 | | |
| 6 | | |
| 6 | | |
| 9 | | |
| 9 | | |

| |
|---|
| 9 |
| 9 |
| 9 |
| 8 |
| 8 |
| 8 |
| 8 |
| 8 |
