## Supplementary Table 6 for "Regional and cell type-specific afferent and efferent projections of the mouse claustrum"

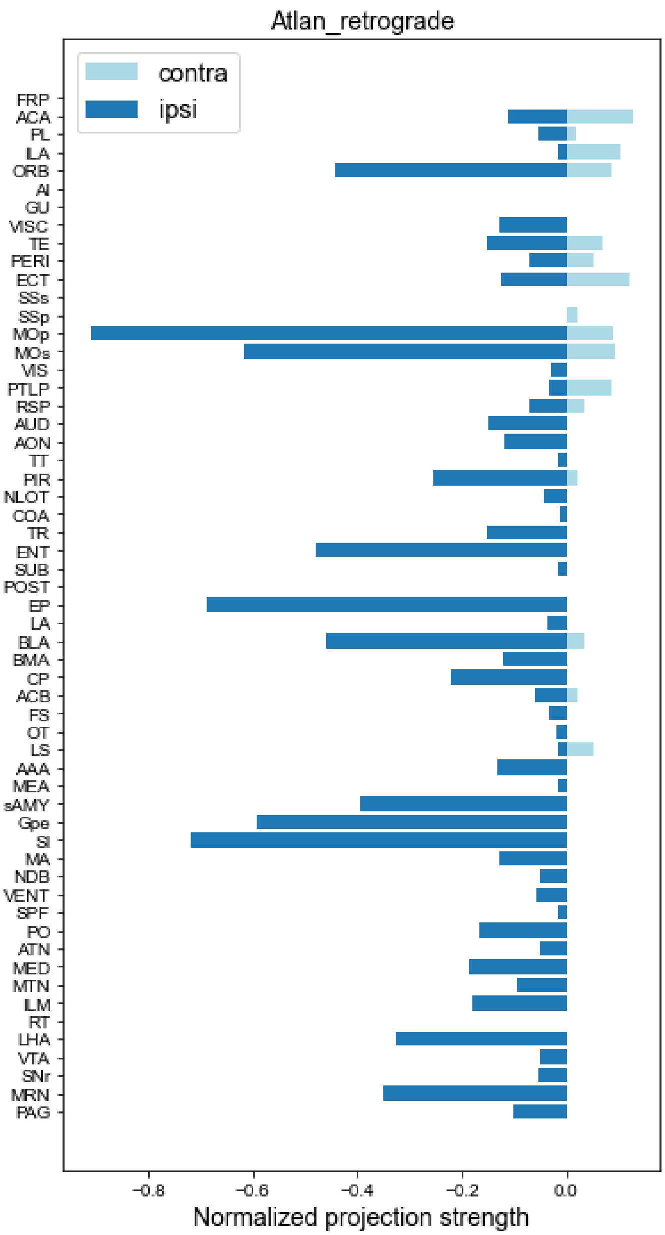


Atlan_retrograde

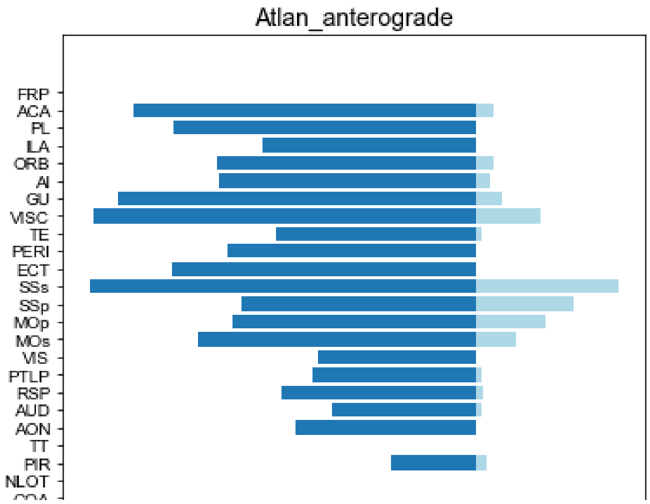


Atlan_anterograde

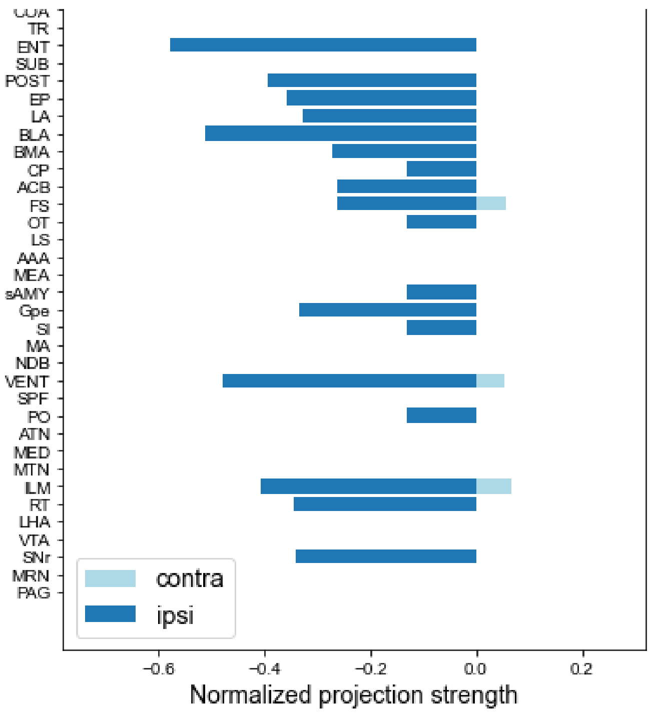


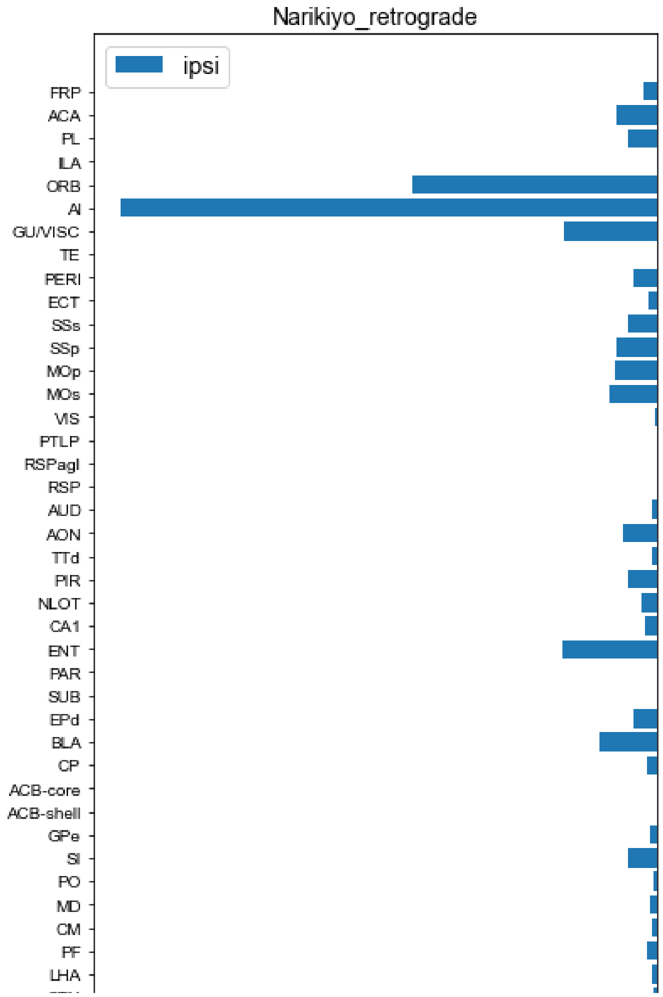


Narikiyo_retrograde
Average % of total Rabies
(n=4).

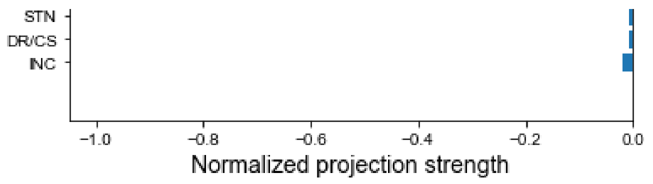


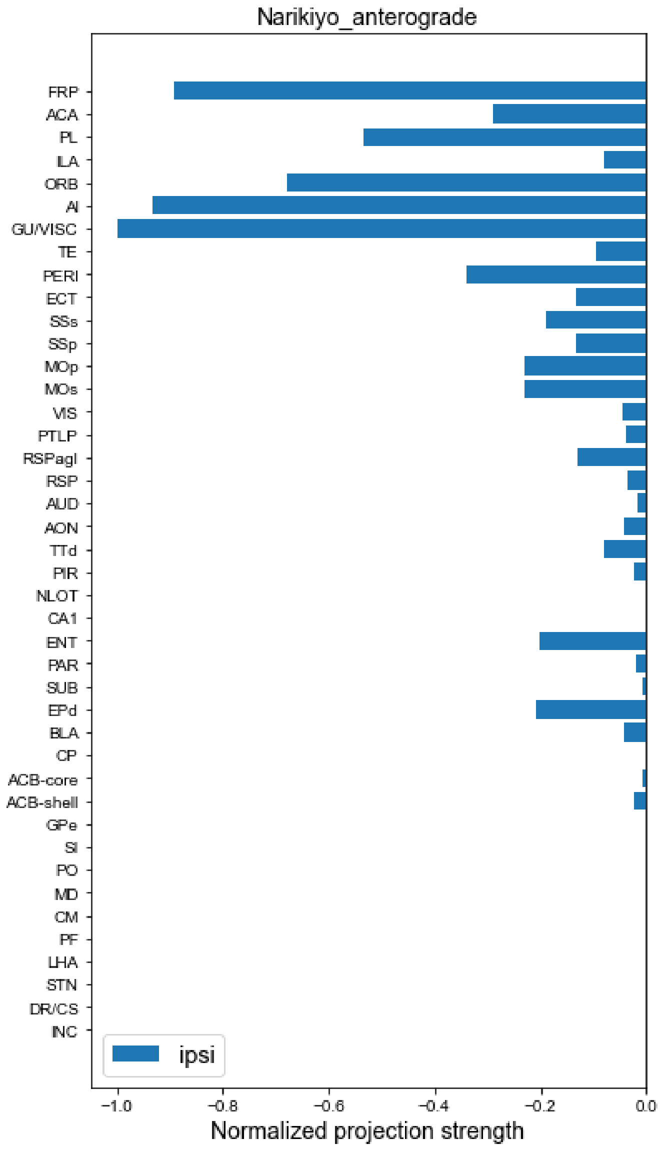


Narikiyo_anterograde
Average % of total sypGFP
(n=4).

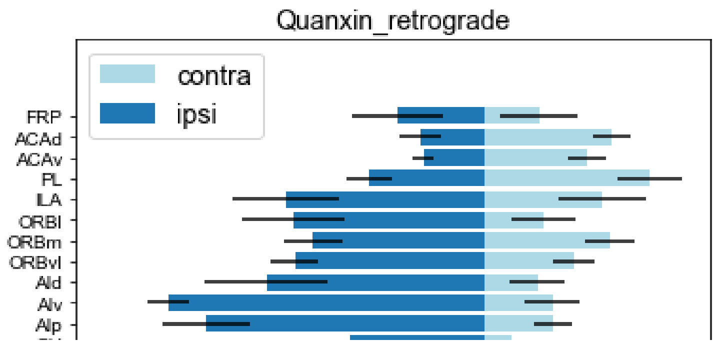

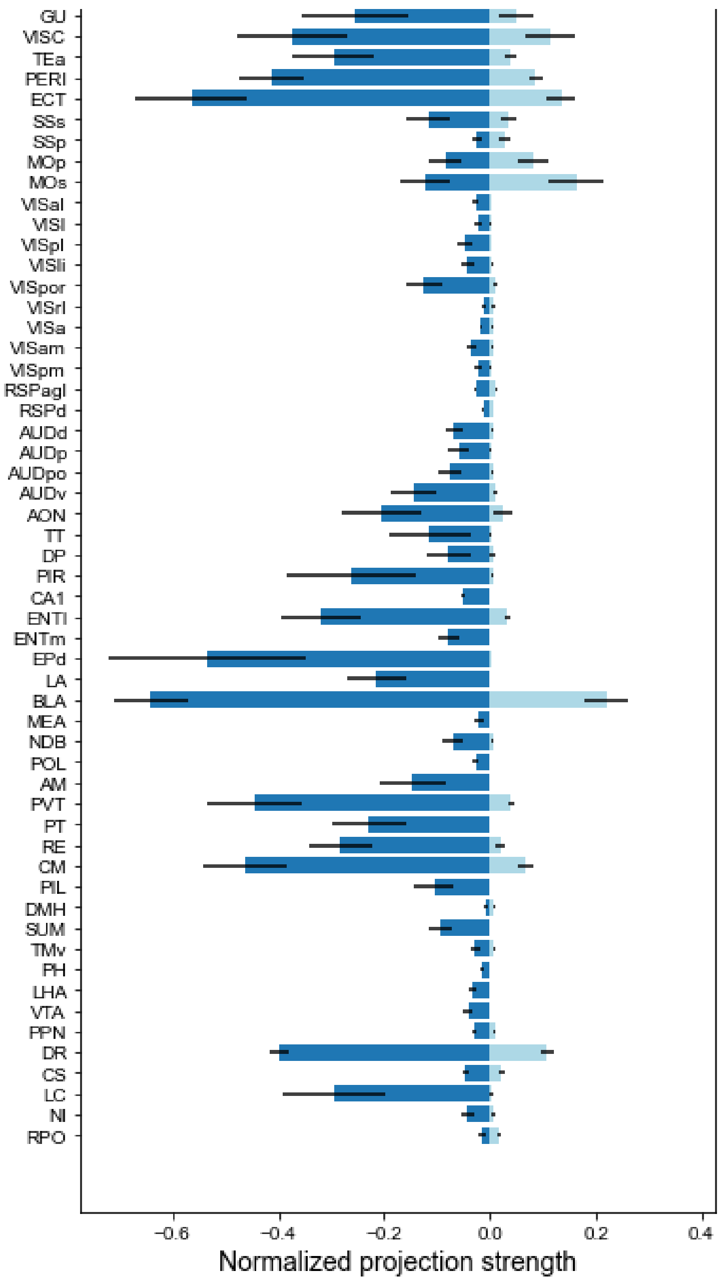


Quanxin_retrograde

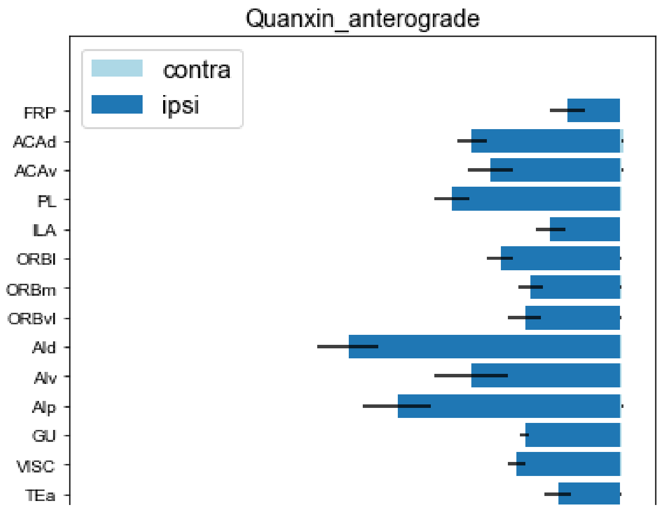

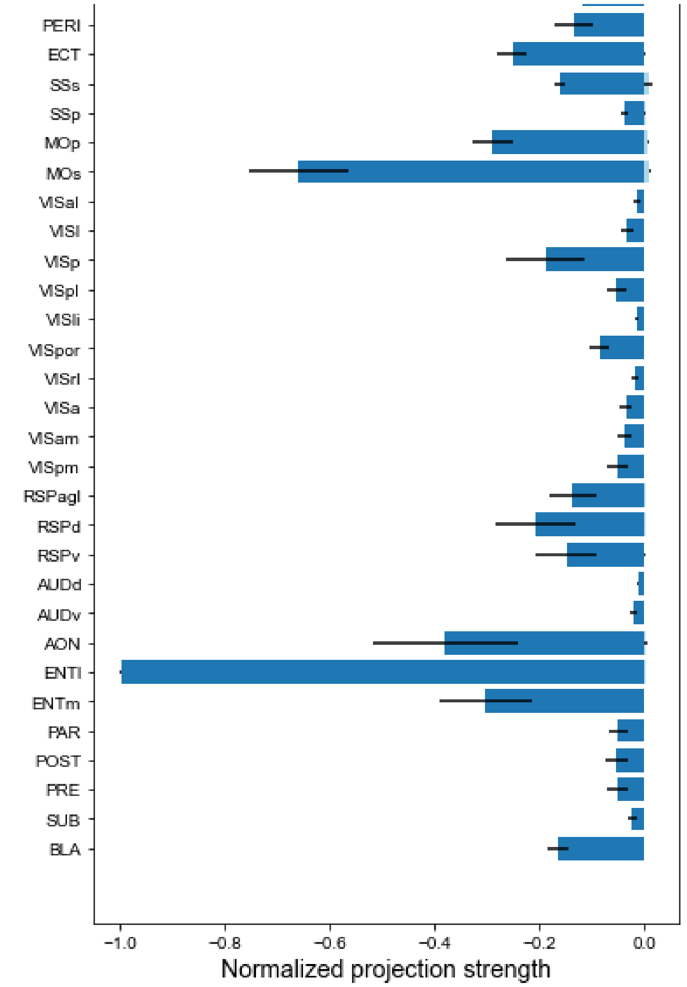


Quanxin_anterograde

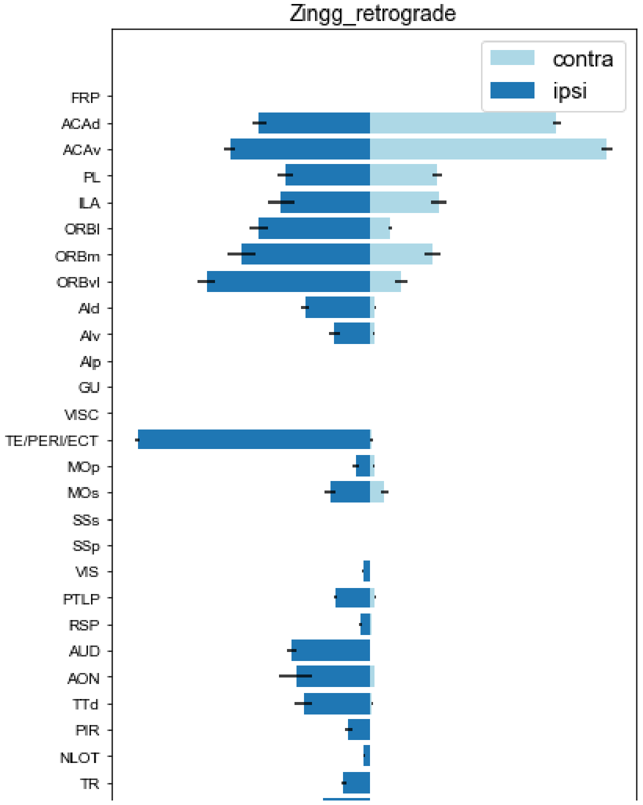


Zingg_retrograde

Number of cells projecting to CLA from ipsilateral and contralateral brain regions (n = 4 animals)

Rules of conversion from region names in the original paper to CCFv3 names shown in "Zingg_region_matching".

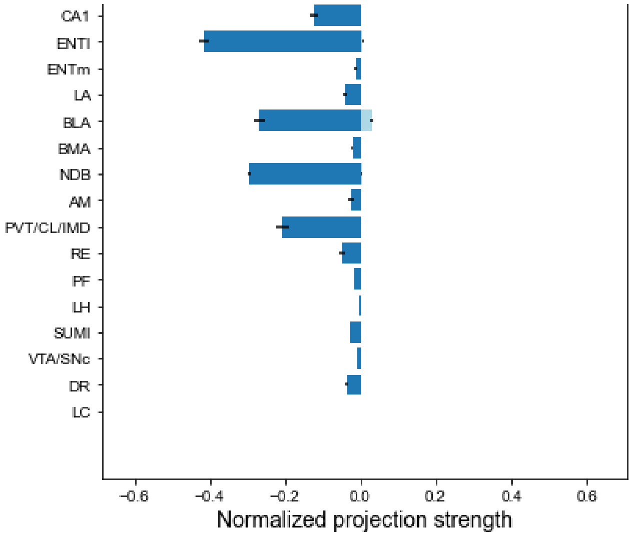


Absent regions in the original paper were filled with 0.

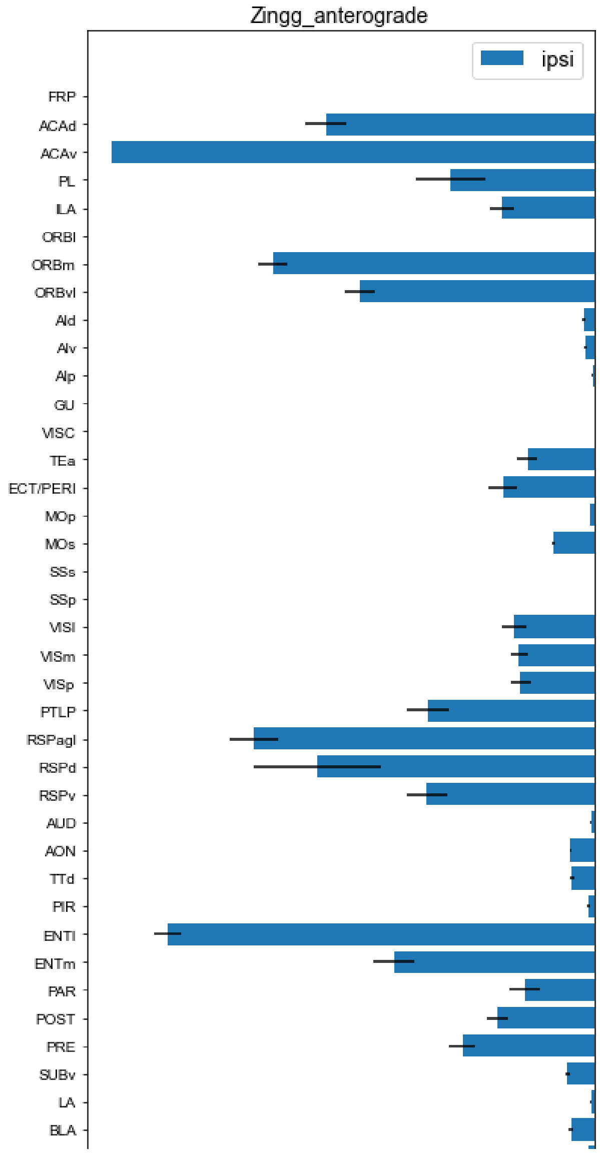


Zingg_anterograde

Number of cells projecting to CLA from ipsilateral and contralateral brain regions (n = 4 animals)

Rules of conversion from region names in the original paper to CCFv3 names shown in "Zingg_region_matching".

Absent regions in the original paper were filled with 0.

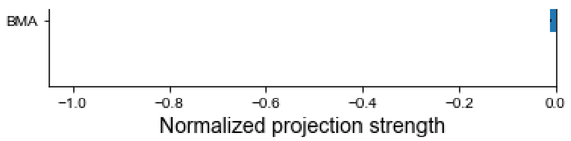
BMA
−1.0
−0.8
−0.6
−0.4
−0.2
0.0
Normalized projection strength

| | | FRP | ACA | PL | ILA |
|---|---|---|---|---|---|
| Raw projection strength | ipsilateral | 0 | 0.011984 | 0.005658 | 0.001812 |
| | controlater | 0 | 0.013544 | 0.001812 | 0.011092 |
| | Sum | 0 | 0.025528 | 0.007469 | 0.012904 |
| Normalized projection strength | ipsilateral | 0 | -0.11258 | -0.05315 | -0.01702 |
| | controlater | 0 | 0.127238 | 0.017019 | 0.104207 |

| | | FRP | ACA | PL | ILA |
|---|---|---|---|---|---|
| Raw projection strength | ipsilateral | 0 | 2.46129 | 2.174242 | 1.541667 |
| | controlater | 0 | 0.128629 | 0 | 0 |
| | Sum | 0 | 2.589919 | 2.174242 | 1.541667 |
| Normalized projection strength | ipsilateral | 0 | -0.64624 | -0.57088 | -0.40478 |
| | controlater | 0 | 0.033773 | 0 | 0 |

| | | FRP | ACA | PL | ILA |
|---|---|---|---|---|---|
| Raw projection strength | ipsilateral | 0.871462 | 2.678022 | 1.960159 | 0 |
| Normalized projection strength | ipsilateral | -0.02491 | -0.07656 | -0.05604 | 0 |

| | | FRP | ACA | PL | ILA |
|---|---|---|---|---|---|
| Raw projection strength | ipsilateral | 13.3273 | 4.341513 | 7.987564 | 1.210745 |
| Normalized projection strength | ipsilateral | -0.8904 | -0.29006 | -0.53365 | -0.08089 |

Raw projection strength
Normalized projection volume (NPV) values for each brain region.
NPV: the volume of detected projection

| | Experiment | FRP | ACAd | ACAv |
|---|---|---|---|---|
| Ipsilateral | 780505913 | 0.098477 | 0.149005 | 0.174787 |
| | 780508099 | 0.051462 | 0.04014 | 0.070335 |
| | 780510390 | 0.00447 | 0.008065 | 0.016724 |
| | 863469869 | 0.080408 | 0.055823 | 0.059898 |
| | 730766377 | 0.128956 | 0.067917 | 0.022122 |
| | 780505913 | 0.034931 | 0.250561 | 0.291951 |
| | 780508099 | 0.016683 | 0.103813 | 0.107637 |

NPV: the volume of detected projection signals in all voxels in a structure (in mm3), divided by the total volume of detected signal in the manually annotated injection site.

| | | | | |
|---|---|---|---|---|
| Contralatera | 780510390 | 0.00197 | 0.030266 | 0.036497 |
| | 863469869 | 0.034702 | 0.137205 | 0.103088 |
| | 730766377 | 0.102207 | 0.096081 | 0.024772 |
| Sum | 780505913 | 0.133408 | 0.399566 | 0.466738 |
| | 780508099 | 0.068145 | 0.143953 | 0.177972 |
| | 780510390 | 0.00644 | 0.038331 | 0.053221 |
| | 863469869 | 0.11511 | 0.193028 | 0.162986 |
| | 730766377 | 0.231163 | 0.163998 | 0.046894 |

Normalized projection strength

| | Experimen | FRP | ACAd | ACAv |
|---|---|---|---|---|
| Ipsilateral | 780505913 | 0.0995966 | 0.150699 | 0.1767741 |
| | 780508099 | 0.0959568 | 0.0748456 | 0.1311476 |
| | 780510390 | 0.022858 | 0.0412416 | 0.0855207 |
| | 863469869 | 0.147578 | 0.1024555 | 0.1099346 |
| | 730766377 | 0.4622476 | 0.243451 | 0.0792971 |
| Contralatera | 780505913 | 0.0353281 | 0.2534096 | 0.2952701 |
| | 780508099 | 0.0311074 | 0.1935712 | 0.2007015 |
| | 780510390 | 0.0100739 | 0.1547698 | 0.1866329 |
| | 863469869 | 0.0636908 | 0.2518211 | 0.189204 |
| | 730766377 | 0.3663648 | 0.344406 | 0.0887962 |
| Sum | 780505913 | 0.134925 | 0.404109 | 0.472044 |
| | 780508099 | 0.127064 | 0.268417 | 0.331849 |
| | 780510390 | 0.032932 | 0.196011 | 0.272154 |
| | 863469869 | 0.211269 | 0.354277 | 0.299139 |
| | 730766377 | 0.828612 | 0.587857 | 0.168093 |

| | | FRP | ACAd | ACAv |
|---|---|---|---|---|
| ipsilateral | Mean | 0.165647 | 0.122539 | 0.116535 |
| controlater | | 0.101313 | 0.239596 | 0.192121 |
| ipsilateral | Std | 0.153562 | 0.070285 | 0.035311 |
| controlater | | 0.133623 | 0.064248 | 0.065465 |

Raw projection strength
Normalized projection volume (NPV) values for each brain region.
NPV: the volume of detected projection signals in all voxels in a structure (in mm3), divided by the total volume of detected signal in the manually annotated injection site.

| | Experimen | FRP | ACAd | ACAv |
|---|---|---|---|---|
| Ipsilateral | 485903475 | 0.441235 | 2.016716 | 1.825853 |
| | 485846989 | 0.233785 | 1.429705 | 1.444136 |
| | 513775257 | 0.148496 | 0.64598 | 0.647831 |
| | 514505957 | 0.326472 | 0.404364 | 0.187351 |
| | 513773998 | 0.197135 | 0.451028 | 0.440182 |
| Contralatera | 485903475 | 0.001319 | 0.044511 | 0.053501 |
| | 485846989 | 0.001737 | 0.030021 | 0.024671 |
| | 513775257 | 0.000252 | 0.012128 | 0.011599 |
| | 514505957 | 0.000295 | 0.000478 | 8.19E-05 |
| | 513773998 | 0.001702 | 0.008083 | 0.004258 |
| | 485903475 | 0.442554 | 2.061227 | 1.879354 |

annotated injection site.

| | | | | |
|---|---|---|---|---|
| Sum | 485846989 | 0.235522 | 1.459726 | 1.468807 |
| | 513775257 | 0.148748 | 0.658108 | 0.65943 |
| | 514505957 | 0.326767 | 0.404842 | 0.187433 |
| | 513773998 | 0.198837 | 0.459111 | 0.44444 |

Normalized projection strength

| | Experimen | FRP | ACAd | ACAv |
|---|---|---|---|---|
| Ipsilateral | 485903475 | 0.072595 | 0.331804 | 0.300402 |
| | 485846989 | 0.056277 | 0.344158 | 0.347632 |
| | 513775257 | 0.060145 | 0.26164 | 0.26239 |
| | 514505957 | 0.217659 | 0.26959 | 0.124907 |
| | 513773998 | 0.090554 | 0.20718 | 0.202198 |
| Contralatera | 485903475 | 0.000217 | 0.007323 | 0.008802 |
| | 485846989 | 0.000418 | 0.007227 | 0.005939 |
| | 513775257 | 0.000102 | 0.004912 | 0.004698 |
| | 514505957 | 0.000197 | 0.000319 | 5.46E-05 |
| | 513773998 | 0.000782 | 0.003713 | 0.001956 |

| | | FRP | ACAd | ACAv |
|---|---|---|---|---|
| ipsilateral | Mean | 0.099446 | 0.282875 | 0.247506 |
| controlater | | 0.000343 | 0.004699 | 0.00429 |
| ipsilateral | Std | 0.060303 | 0.050014 | 0.077625 |
| controlater | | 0.000242 | 0.002588 | 0.003054 |

Raw projection strength

| | Experimen | FRP | ACAd | ACAv |
|---|---|---|---|---|
| Ipsilateral | 1 | 0 | 421 | 517 |
| | 2 | 0 | 243 | 296 |
| | 3 | 0 | 190 | 271 |
| | 4 | 0 | 377 | 465 |
| Contralatera | 1 | 0 | 727 | 926 |
| | 2 | 0 | 372 | 424 |
| | 3 | 0 | 390 | 516 |
| | 4 | 0 | 589 | 786 |
| Sum | 1 | 0 | 1148 | 1443 |
| | 2 | 0 | 615 | 720 |
| | 3 | 0 | 580 | 787 |
| | 4 | 0 | 966 | 1251 |

| | Experimen | FRP | ACAd | ACAv |
|---|---|---|---|---|
| Ipsilateral | 1 | 0 | 0.291753 | 0.358281 |
| | 2 | 0 | 0.3375 | 0.411111 |
| | 3 | 0 | 0.241423 | 0.344346 |
| | 4 | 0 | 0.301359 | 0.371703 |
| | 1 | 0 | 0.503812 | 0.641719 |

| | | | | | |
|---|---|---|---|---|---|
| Normalized projection strength | Contralatera | 2 | 0 | 0.516667 | 0.588889 |
| | | 3 | 0 | 0.495553 | 0.655654 |
| | | 4 | 0 | 0.470823 | 0.628297 |
| | | | FRP | ACAd | ACAv |
| | ipsilateral | Mean | 0 | 0.293009 | 0.37136 |
| | controlater | | 0 | 0.496714 | 0.62864 |
| | ipsilateral | Std | 0 | 0.034321 | 0.024905 |
| | controlater | | 0 | 0.016734 | 0.024905 |

| | | Experimen | FRP | ACAd | ACAv |
|---|---|---|---|---|---|
| Raw projection strength | Ipsilateral | **1** | 0 | 0.0818 | 0.1411 |
| | | **2** | 0 | 0.0528 | 0.1147 |
| | | **3** | 0 | 0.0699 | 0.1118 |
| | | Experimen | FRP | ACAd | ACAv |
| Normalized projection strength | Ipsilateral | **1** | 0 | 0.579731 | 1 |
| | | **2** | 0 | 0.460331 | 1 |
| | | **3** | 0 | 0.625224 | 1 |
| | ipsilateral | Mean | 0 | 0.068167 | 0.122533 |
| | ipsilateral | Std | 0 | 0.069534 | 0 |

| ORB | Al | GU | VISC | TE | PERl | ECT | SSs | SSp |
|---|---|---|---|---|---|---|---|---|
| 0.046987 | 0 | 0 | 0.013544 | 0.01629 | 0.007469 | 0.01335 | 0 | 0 |
| 0.009058 | 0 | 0 | 0 | 0.007246 | 0.005435 | 0.012681 | 0 | 0.002257 |
| 0.056045 | 0 | 0 | 0.013544 | 0.023536 | 0.012904 | 0.026031 | 0 | 0.002257 |
| -0.44141 | 0 | 0 | -0.12724 | -0.15303 | -0.07017 | -0.12541 | 0 | 0 |
| 0.085094 | 0 | 0 | 0 | 0.068075 | 0.051056 | 0.119132 | 0 | 0.021206 |

| ORB | Al | GU | VISC | TE | PERl | ECT | SSs | SSp |
|---|---|---|---|---|---|---|---|---|
| 1.868519 | 1.849707 | 2.571429 | 2.75 | 1.432353 | 1.785088 | 2.194444 | 2.782297 | 1.689071 |
| 0.127273 | 0.100529 | 0.19 | 0.467611 | 0.041667 | 0 | 0 | 1.026316 | 0.695946 |
| 1.995791 | 1.950236 | 2.761429 | 3.217611 | 1.47402 | 1.785088 | 2.194444 | 3.808612 | 2.385017 |
| -0.4906 | -0.48566 | -0.67516 | -0.72205 | -0.37608 | -0.4687 | -0.57618 | -0.73053 | -0.44349 |
| 0.033417 | 0.026395 | 0.049887 | 0.122777 | 0.01094 | 0 | 0 | 0.269472 | 0.18273 |

| ORB | AI | GU/VISC | TE | PERl | ECT | SSs | SSp | MOp |
|---|---|---|---|---|---|---|---|---|
| 15.99939 | 34.98091 | 6.047172 | 0.050419 | 1.559792 | 0.61108 | 1.859978 | 2.642092 | 2.80839 |
| -0.45737 | -1 | -0.17287 | -0.00144 | -0.04459 | -0.01747 | -0.05317 | -0.07553 | -0.08028 |

| ORB | AI | GU/VISC | TE | PERl | ECT | SSs | SSp | MOp |
|---|---|---|---|---|---|---|---|---|
| 10.16891 | 13.97384 | 14.9677 | 1.440798 | 5.083525 | 1.967404 | 2.823272 | 2.000388 | 3.46607 |
| -0.67939 | -0.9336 | -1 | -0.09626 | -0.33963 | -0.13144 | -0.18862 | -0.13365 | -0.23157 |

| PL | ILA | ORBl | ORBm | ORBvl | AId | AIv | AIp | GU |
|---|---|---|---|---|---|---|---|---|
| 0.257745 | 0.520189 | 0.399235 | 0.462824 | 0.454057 | 0.306168 | 0.665558 | 0.584171 | 0.158911 |
| 0.105146 | 0.301728 | 0.159389 | 0.225441 | 0.201531 | 0.129311 | 0.354158 | 0.295115 | 0.074509 |
| 0.024732 | 0.075317 | 0.031985 | 0.052732 | 0.049026 | 0.038472 | 0.094437 | 0.105465 | 0.024079 |
| 0.091151 | 0.191973 | 0.150923 | 0.137923 | 0.157877 | 0.332448 | 0.312098 | 0.379007 | 0.328426 |
| 0.096816 | 0.016659 | 0.188388 | 0.059686 | 0.118096 | 0.198524 | 0.168362 | 0.071751 | 0.070136 |
| 0.293032 | 0.462456 | 0.096213 | 0.352765 | 0.235631 | 0.057662 | 0.130417 | 0.095809 | 0.031773 |
| 0.152482 | 0.155092 | 0.023387 | 0.160588 | 0.080687 | 0.013471 | 0.031467 | 0.064733 | 0.009056 |

| | | | | | | | | |
|---|---|---|---|---|---|---|---|---|
| 0.038065 | 0.027404 | 0.003917 | 0.026898 | 0.018002 | 0.001727 | 0.003279 | 0.017016 | 0.004305 |
| 0.138068 | 0.095698 | 0.037526 | 0.08532 | 0.051482 | 0.070042 | 0.070607 | 0.140202 | 0.087361 |
| 0.143888 | 0.010783 | 0.090588 | 0.061179 | 0.071936 | 0.074868 | 0.08201 | 0.02127 | 0.004404 |
| 0.550777 | 0.982645 | 0.495448 | 0.815589 | 0.689688 | 0.36383 | 0.795975 | 0.67998 | 0.190684 |
| 0.257628 | 0.45682 | 0.182776 | 0.386029 | 0.282218 | 0.142782 | 0.385625 | 0.359848 | 0.083565 |
| 0.062797 | 0.102721 | 0.035902 | 0.07963 | 0.067028 | 0.040199 | 0.097716 | 0.122481 | 0.028384 |
| 0.229219 | 0.287671 | 0.188449 | 0.223243 | 0.209359 | 0.40249 | 0.382705 | 0.519209 | 0.415787 |
| 0.240704 | 0.027442 | 0.278976 | 0.120865 | 0.190032 | 0.273392 | 0.250372 | 0.093021 | 0.07454 |

| PL | ILA | ORBl | ORBm | ORBvl | AId | AIv | AIp | GU |
|---|---|---|---|---|---|---|---|---|
| 0.2606753 | 0.5261029 | 0.4037738 | 0.4680858 | 0.4592191 | 0.3096488 | 0.6731246 | 0.5908123 | 0.1607176 |
| 0.1960567 | 0.5626063 | 0.297199 | 0.4203605 | 0.3757775 | 0.2411151 | 0.660368 | 0.5502756 | 0.1389305 |
| 0.1264708 | 0.3851448 | 0.1635601 | 0.269653 | 0.2507018 | 0.1967324 | 0.4829178 | 0.5393112 | 0.1231316 |
| 0.1672953 | 0.3523404 | 0.2769987 | 0.2531389 | 0.2897618 | 0.6101631 | 0.5728135 | 0.6956159 | 0.6027813 |
| 0.3470406 | 0.0597148 | 0.6752839 | 0.2139467 | 0.4233196 | 0.7116168 | 0.6034999 | 0.2571942 | 0.2514051 |
| 0.2963634 | 0.4677136 | 0.0973068 | 0.3567755 | 0.2383098 | 0.0583175 | 0.1318997 | 0.0968982 | 0.0321342 |
| 0.2843201 | 0.2891867 | 0.0436077 | 0.2994346 | 0.1504501 | 0.0251182 | 0.0586738 | 0.1207021 | 0.0168859 |
| 0.1946511 | 0.1401345 | 0.0200302 | 0.137547 | 0.0920559 | 0.0088313 | 0.0167677 | 0.0870139 | 0.0220143 |
| 0.2534051 | 0.1756407 | 0.0688739 | 0.1565933 | 0.0944882 | 0.1285526 | 0.1295896 | 0.2573217 | 0.1603392 |
| 0.515772 | 0.0386521 | 0.3247161 | 0.2192984 | 0.2578573 | 0.2683672 | 0.2939679 | 0.0762431 | 0.0157863 |
| 0.557039 | 0.993816 | 0.501081 | 0.824861 | 0.697529 | 0.367966 | 0.805024 | 0.687711 | 0.192852 |
| 0.480377 | 0.851793 | 0.340807 | 0.719795 | 0.526228 | 0.266233 | 0.719042 | 0.670978 | 0.155816 |
| 0.321122 | 0.525279 | 0.18359 | 0.4072 | 0.342758 | 0.205564 | 0.499686 | 0.626325 | 0.145146 |
| 0.4207 | 0.527981 | 0.345873 | 0.409732 | 0.38425 | 0.738716 | 0.702403 | 0.952938 | 0.763121 |
| 0.862813 | 0.098367 | 1 | 0.433245 | 0.681177 | 0.979984 | 0.897468 | 0.333437 | 0.267191 |

| PL | ILA | ORBl | ORBm | ORBvl | AId | AIv | AIp | GU |
|---|---|---|---|---|---|---|---|---|
| 0.219508 | 0.377182 | 0.363363 | 0.325037 | 0.359756 | 0.413855 | 0.598545 | 0.526642 | 0.255393 |
| 0.308902 | 0.222266 | 0.110907 | 0.23393 | 0.166632 | 0.097837 | 0.12618 | 0.127636 | 0.049432 |
| 0.077319 | 0.177778 | 0.1736 | 0.100127 | 0.078717 | 0.207383 | 0.06847 | 0.145596 | 0.179284 |
| 0.109261 | 0.146531 | 0.109958 | 0.083506 | 0.069983 | 0.094653 | 0.094578 | 0.066487 | 0.055754 |

| PL | ILA | ORBl | ORBm | ORBvl | AId | AIv | AIp | GU |
|---|---|---|---|---|---|---|---|---|
| 1.782951 | 0.693587 | 1.217304 | 1.027388 | 1.265235 | 2.429602 | 1.467801 | 3.146529 | 0.99564 |
| 1.253824 | 0.780888 | 0.763711 | 0.807838 | 0.895378 | 1.91445 | 1.054615 | 1.298178 | 0.721297 |
| 0.660266 | 0.39773 | 0.537675 | 0.43456 | 0.449598 | 1.120018 | 0.716812 | 0.948138 | 0.44191 |
| 0.649223 | 0.073038 | 0.346122 | 0.140579 | 0.11008 | 0.890375 | 0.187153 | 0.466685 | 0.268252 |
| 0.667544 | 0.324759 | 0.673915 | 0.47876 | 0.492018 | 1.481342 | 1.107319 | 1.304365 | 0.466825 |
| 0.006345 | 0.001333 | 0.000796 | 0.002264 | 0.00437 | 0.009102 | 0.001915 | 0.014056 | 0.009252 |
| 0.014449 | 0.003705 | 0.001054 | 0.008082 | 0.005517 | 0.00532 | 0.000169 | 0.001604 | 0.021867 |
| 0.004874 | 0.001199 | 0.002254 | 0.002323 | 0.001937 | 0.010104 | 0.006382 | 0.010637 | 0.007074 |
| 2.03E-05 | 0 | 5.06E-06 | 0.00031 | 8.61E-05 | 0.005643 | 0.007078 | 0.005863 | 0.005634 |
| 0.007916 | 0.002181 | 0.005964 | 0.005507 | 0.002639 | 0.012837 | 0.013445 | 0.013022 | 0.00854 |
| 1.789296 | 0.69492 | 1.2181 | 1.029652 | 1.269605 | 2.438704 | 1.469716 | 3.160585 | 1.004892 |

| | | | | | | | | |
|---|---|---|---|---|---|---|---|---|
| 1.268273 | 0.784593 | 0.764765 | 0.81592 | 0.900895 | 1.91977 | 1.054784 | 1.299782 | 0.743164 |
| 0.66514 | 0.398929 | 0.539929 | 0.436883 | 0.451535 | 1.130122 | 0.723194 | 0.958775 | 0.448984 |
| 0.649243 | 0.073038 | 0.346127 | 0.140889 | 0.110166 | 0.896018 | 0.194231 | 0.472548 | 0.273886 |
| 0.67546 | 0.32694 | 0.679879 | 0.484267 | 0.494657 | 1.494179 | 1.120764 | 1.317387 | 0.475365 |

| PL | ILA | ORBl | ORBm | ORBvl | AId | AIv | AIp | GU |
|---|---|---|---|---|---|---|---|---|
| 0.293344 | 0.114114 | 0.200279 | 0.169033 | 0.208165 | 0.399735 | 0.241493 | 0.517689 | 0.16381 |
| 0.30182 | 0.187975 | 0.18384 | 0.194463 | 0.215535 | 0.460846 | 0.253867 | 0.312497 | 0.17363 |
| 0.267426 | 0.161092 | 0.217774 | 0.176009 | 0.1821 | 0.453639 | 0.290329 | 0.384023 | 0.178986 |
| 0.432838 | 0.048694 | 0.23076 | 0.093724 | 0.07339 | 0.593614 | 0.124775 | 0.311139 | 0.178844 |
| 0.306637 | 0.149178 | 0.309563 | 0.219919 | 0.226009 | 0.680456 | 0.508648 | 0.599161 | 0.214436 |
| 0.001044 | 0.000219 | 0.000131 | 0.000372 | 0.000719 | 0.001498 | 0.000315 | 0.002313 | 0.001522 |
| 0.003478 | 0.000892 | 0.000254 | 0.001945 | 0.001328 | 0.001281 | 4.07E-05 | 0.000386 | 0.005264 |
| 0.001974 | 0.000486 | 0.000913 | 0.000941 | 0.000785 | 0.004092 | 0.002585 | 0.004308 | 0.002865 |
| 1.35E-05 | 0 | 3.37E-06 | 0.000207 | 5.74E-05 | 0.003762 | 0.004719 | 0.003909 | 0.003756 |
| 0.003636 | 0.001002 | 0.00274 | 0.00253 | 0.001212 | 0.005897 | 0.006176 | 0.005982 | 0.003923 |

| PL | ILA | ORBl | ORBm | ORBvl | AId | AIv | AIp | GU |
|---|---|---|---|---|---|---|---|---|
| 0.320413 | 0.132211 | 0.228443 | 0.17063 | 0.18104 | 0.517658 | 0.283822 | 0.424902 | 0.181941 |
| 0.002029 | 0.00052 | 0.000808 | 0.001199 | 0.00082 | 0.003306 | 0.002767 | 0.00338 | 0.003466 |
| 0.057819 | 0.048029 | 0.043548 | 0.042293 | 0.055745 | 0.103445 | 0.125359 | 0.115086 | 0.017159 |
| 0.001394 | 0.000383 | 0.001015 | 0.000901 | 0.000448 | 0.001727 | 0.002405 | 0.001898 | 0.001238 |

| PL | ILA | ORBl | ORBm | ORBvl | AId | AIv | AIp | GU |
|---|---|---|---|---|---|---|---|---|
| 299 | 448 | 450 | 631 | 597 | 242 | 183 | 0 | 0 |
| 204 | 165 | 258 | 243 | 360 | 101 | 39 | 0 | 0 |
| 149 | 113 | 203 | 203 | 323 | 153 | 64 | 0 | 0 |
| 273 | 324 | 318 | 412 | 511 | 229 | 138 | 0 | 0 |
| 255 | 353 | 93 | 309 | 71 | 22 | 15 | 0 | 0 |
| 154 | 125 | 43 | 106 | 94 | 9 | 8 | 0 | 0 |
| 141 | 114 | 35 | 98 | 68 | 16 | 11 | 0 | 0 |
| 193 | 215 | 68 | 224 | 83 | 15 | 14 | 0 | 0 |
| 554 | 801 | 543 | 940 | 668 | 264 | 198 | 0 | 0 |
| 358 | 290 | 301 | 349 | 454 | 110 | 47 | 0 | 0 |
| 290 | 227 | 238 | 301 | 391 | 169 | 75 | 0 | 0 |
| 466 | 539 | 386 | 636 | 594 | 244 | 152 | 0 | 0 |

| PL | ILA | ORBl | ORBm | ORBvl | AId | AIv | AIp | GU |
|---|---|---|---|---|---|---|---|---|
| 0.207207 | 0.310464 | 0.31185 | 0.437283 | 0.413721 | 0.167706 | 0.126819 | 0 | 0 |
| 0.283333 | 0.229167 | 0.358333 | 0.3375 | 0.5 | 0.140278 | 0.054167 | 0 | 0 |
| 0.189327 | 0.143583 | 0.257942 | 0.257942 | 0.410419 | 0.194409 | 0.081321 | 0 | 0 |
| 0.218225 | 0.258993 | 0.254197 | 0.329337 | 0.408473 | 0.183054 | 0.110312 | 0 | 0 |
| 0.176715 | 0.244629 | 0.064449 | 0.214137 | 0.049203 | 0.015246 | 0.010395 | 0 | 0 |

| | | | | | | | | |
|---|---|---|---|---|---|---|---|---|
| 0.213889 | 0.173611 | 0.059722 | 0.147222 | 0.130556 | 0.0125 | 0.011111 | 0 | 0 |
| 0.179161 | 0.144854 | 0.044473 | 0.124524 | 0.086404 | 0.02033 | 0.013977 | 0 | 0 |
| 0.154277 | 0.171863 | 0.054357 | 0.179057 | 0.066347 | 0.01199 | 0.011191 | 0 | 0 |

| PL | ILA | ORBl | ORBm | ORBvl | Ald | Alv | Alp | GU |
|---|---|---|---|---|---|---|---|---|
| 0.224523 | 0.235552 | 0.29558 | 0.340515 | 0.433153 | 0.171362 | 0.093155 | 0 | 0 |
| 0.181011 | 0.183739 | 0.05575 | 0.166235 | 0.083127 | 0.015017 | 0.011669 | 0 | 0 |
| 0.035486 | 0.06054 | 0.042813 | 0.063868 | 0.038639 | 0.020294 | 0.027784 | 0 | 0 |
| 0.021316 | 0.036957 | 0.007426 | 0.033765 | 0.030383 | 0.003308 | 0.001368 | 0 | 0 |

| PL | ILA | ORBl | ORBm | ORBvl | Ald | Alv | Alp | GU |
|---|---|---|---|---|---|---|---|---|
| 0.0283 | 0.0186 | 0 | 0.0844 | 0.0667 | 0.0033 | 0.0037 | 0.0002 | 0 |
| 0.0263 | 0.0241 | 0 | 0.0785 | 0.0494 | 0.0017 | 0.0012 | 0.0002 | 0 |
| 0.0518 | 0.0252 | 0 | 0.0794 | 0.0619 | 0.0026 | 0.0018 | 0.0007 | 0 |

| PL | ILA | ORBl | ORBm | ORBvl | Ald | Alv | Alp | GU |
|---|---|---|---|---|---|---|---|---|
| 0.200567 | 0.131821 | 0 | 0.598157 | 0.472714 | 0.023388 | 0.026223 | 0.001417 | 0 |
| 0.229294 | 0.210113 | 0 | 0.684394 | 0.430689 | 0.014821 | 0.010462 | 0.001744 | 0 |
| 0.463327 | 0.225403 | 0 | 0.710197 | 0.553667 | 0.023256 | 0.0161 | 0.006261 | 0 |
| 0.035467 | 0.022633 | 0 | 0.080767 | 0.059333 | 0.002533 | 0.002233 | 0.000367 | 0 |
| 0.117681 | 0.040989 | 0 | 0.047907 | 0.051037 | 0.004008 | 0.00652 | 0.00221 | 0 |

| MOp | MOs | VIS | PTLP | RSP | AUD | AON | TT | PIR |
|---|---|---|---|---|---|---|---|---|
| 0.096957 | 0.065534 | 0.003163 | 0.003623 | 0.007469 | 0.015816 | 0.012667 | 0.001812 | 0.027088 |
| 0.009489 | 0.009935 | 0 | 0.009058 | 0.003623 | 0 | 0 | 0 | 0.002257 |
| 0.106446 | 0.075469 | 0.003163 | 0.012681 | 0.011092 | 0.015816 | 0.012667 | 0.001812 | 0.029345 |
| -0.91085 | -0.61566 | -0.02972 | -0.03404 | -0.07017 | -0.14858 | -0.119 | -0.01702 | -0.25448 |
| 0.089147 | 0.093335 | 0 | 0.085094 | 0.034038 | 0 | 0 | 0 | 0.021206 |

| MOp | MOs | VIS | PTLP | RSP | AUD | AON | TT | PIR |
|---|---|---|---|---|---|---|---|---|
| 1.75 | 2.001974 | 1.136364 | 1.176471 | 1.404672 | 1.031676 | 1.295455 | 0 | 0.611111 |
| 0.506757 | 0.289474 | 0 | 0.041667 | 0.046875 | 0.038194 | 0 | 0 | 0.075 |
| 2.256757 | 2.291447 | 1.136364 | 1.218137 | 1.451547 | 1.069871 | 1.295455 | 0 | 0.686111 |
| -0.45948 | -0.52564 | -0.29837 | -0.3089 | -0.36881 | -0.27088 | -0.34014 | 0 | -0.16046 |
| 0.133055 | 0.076005 | 0 | 0.01094 | 0.012308 | 0.010028 | 0 | 0 | 0.019692 |

| MOs | VIS | PTLP | RSPagl | RSP | AUD | AON | TTd | PIR |
|---|---|---|---|---|---|---|---|---|
| 3.082619 | 0.160202 | 0 | 0 | 0 | 0.42303 | 2.193913 | 0.409577 | 1.92903 |
| -0.08812 | -0.00458 | 0 | 0 | 0 | -0.01209 | -0.06272 | -0.01171 | -0.05515 |

| MOs | VIS | PTLP | RSPagl | RSP | AUD | AON | TTd | PIR |
|---|---|---|---|---|---|---|---|---|
| 3.434618 | 0.656077 | 0.603902 | 1.93023 | 0.546173 | 0.258541 | 0.623524 | 1.169745 | 0.348061 |
| -0.22947 | -0.04383 | -0.04035 | -0.12896 | -0.03649 | -0.01727 | -0.04166 | -0.07815 | -0.02325 |

| VISC | TEa | PERI | ECT | SSs | SSp | MOp | MOs | VISal |
|---|---|---|---|---|---|---|---|---|
| 0.201469 | 0.286146 | 0.442927 | 0.54948 | 0.051579 | 0.014104 | 0.051934 | 0.097402 | 0.029353 |
| 0.185359 | 0.26275 | 0.30343 | 0.42169 | 0.052243 | 0.00983 | 0.024125 | 0.030763 | 0.022854 |
| 0.059208 | 0.064626 | 0.079657 | 0.127239 | 0.018629 | 0.002891 | 0.005855 | 0.007159 | 0.005048 |
| 0.398914 | 0.167026 | 0.224148 | 0.329599 | 0.14109 | 0.031304 | 0.086128 | 0.084394 | 0.015666 |
| 0.07709 | 0.016947 | 0.065092 | 0.062946 | 0.022005 | 0.00516 | 0.03876 | 0.075169 | 0.002544 |
| 0.079619 | 0.035212 | 0.077364 | 0.122729 | 0.025067 | 0.024499 | 0.045666 | 0.113067 | 0.003183 |
| 0.051358 | 0.036335 | 0.047777 | 0.097865 | 0.013834 | 0.010659 | 0.026618 | 0.055065 | 0.002136 |

| | | | | | | | | |
|---|---|---|---|---|---|---|---|---|
| 0.019732 | 0.009873 | 0.016752 | 0.031448 | 0.004863 | 0.003411 | 0.005456 | 0.013127 | 0.000518 |
| 0.145937 | 0.018923 | 0.065324 | 0.084371 | 0.046392 | 0.036709 | 0.088884 | 0.112563 | 0.002787 |
| 0.006757 | 0.000989 | 0.016694 | 0.013464 | 0.003061 | 0.002728 | 0.032435 | 0.089905 | 0.000787 |
| 0.281088 | 0.321358 | 0.520291 | 0.672209 | 0.076646 | 0.038603 | 0.0976 | 0.210469 | 0.032536 |
| 0.236717 | 0.299085 | 0.351207 | 0.519555 | 0.066077 | 0.020489 | 0.050743 | 0.085828 | 0.02499 |
| 0.07894 | 0.074499 | 0.096409 | 0.158687 | 0.023492 | 0.006302 | 0.011311 | 0.020286 | 0.005566 |
| 0.544851 | 0.185949 | 0.289472 | 0.41397 | 0.187482 | 0.068013 | 0.175012 | 0.196957 | 0.018453 |
| 0.083847 | 0.017936 | 0.081786 | 0.07641 | 0.025066 | 0.007888 | 0.071195 | 0.165074 | 0.003331 |

| VISC | TEa | PERI | ECT | SSs | SSp | MOp | MOs | VISal |
|---|---|---|---|---|---|---|---|---|
| 0.2037595 | 0.2893991 | 0.4479625 | 0.5557269 | 0.0521654 | 0.0142643 | 0.0525244 | 0.0985093 | 0.0296867 |
| 0.345623 | 0.4899274 | 0.5657799 | 0.7862891 | 0.097413 | 0.0183292 | 0.0449838 | 0.0573611 | 0.0426139 |
| 0.302769 | 0.3304748 | 0.4073381 | 0.6506558 | 0.0952622 | 0.0147836 | 0.0299404 | 0.0366086 | 0.0258137 |
| 0.7321525 | 0.3065535 | 0.4113932 | 0.6049342 | 0.2589515 | 0.0574542 | 0.1580762 | 0.1548937 | 0.0287528 |
| 0.276332 | 0.0607472 | 0.2333247 | 0.2256323 | 0.0788778 | 0.0184962 | 0.1389367 | 0.2694461 | 0.0091191 |
| 0.0805242 | 0.0356123 | 0.0782435 | 0.1241243 | 0.025352 | 0.0247775 | 0.0461852 | 0.1143524 | 0.0032192 |
| 0.0957629 | 0.0677508 | 0.0890857 | 0.1824805 | 0.0257951 | 0.0198749 | 0.0496323 | 0.102675 | 0.0039828 |
| 0.1009026 | 0.0504871 | 0.0856639 | 0.1608141 | 0.0248677 | 0.0174427 | 0.0279001 | 0.0671269 | 0.0026489 |
| 0.2678475 | 0.0347306 | 0.1198933 | 0.1548515 | 0.0851462 | 0.0673744 | 0.1631345 | 0.2065941 | 0.0051152 |
| 0.0242207 | 0.0035451 | 0.0598403 | 0.0482622 | 0.0109723 | 0.0097786 | 0.1162645 | 0.3222679 | 0.002821 |
| 0.284284 | 0.325011 | 0.526206 | 0.679851 | 0.077517 | 0.039042 | 0.09871 | 0.212862 | 0.032906 |
| 0.441386 | 0.557678 | 0.654866 | 0.96877 | 0.123208 | 0.038204 | 0.094616 | 0.160036 | 0.046597 |
| 0.403672 | 0.380962 | 0.493002 | 0.81147 | 0.12013 | 0.032226 | 0.057841 | 0.103736 | 0.028463 |
| 1 | 0.341284 | 0.531287 | 0.759786 | 0.344098 | 0.124829 | 0.321211 | 0.361488 | 0.033868 |
| 0.300553 | 0.064292 | 0.293165 | 0.273895 | 0.08985 | 0.028275 | 0.255201 | 0.591714 | 0.01194 |

| VISC | TEa | PERI | ECT | SSs | SSp | MOp | MOs | VISal |
|---|---|---|---|---|---|---|---|---|
| 0.372127 | 0.29542 | 0.41316 | 0.564648 | 0.116534 | 0.024666 | 0.084892 | 0.123364 | 0.027197 |
| 0.113852 | 0.038425 | 0.086545 | 0.134107 | 0.034427 | 0.02785 | 0.080623 | 0.162603 | 0.003557 |
| 0.185826 | 0.137327 | 0.10666 | 0.186095 | 0.073024 | 0.016487 | 0.052795 | 0.083468 | 0.010724 |
| 0.081675 | 0.021183 | 0.019501 | 0.046803 | 0.025965 | 0.020348 | 0.050999 | 0.092156 | 0.000904 |

| VISC | TEa | PERI | ECT | SSs | SSp | MOp | MOs | VISal |
|---|---|---|---|---|---|---|---|---|
| 1.263211 | 1.184278 | 0.745667 | 1.557112 | 1.122088 | 0.31822 | 2.170432 | 4.366504 | 0.163108 |
| 0.69322 | 0.514981 | 0.382745 | 0.929766 | 0.701875 | 0.205833 | 1.504558 | 2.492973 | 0.079301 |
| 0.404625 | 0.284918 | 0.226965 | 0.530791 | 0.352177 | 0.076944 | 0.619531 | 1.164495 | 0.056038 |
| 0.369772 | 0.083771 | 0.395867 | 0.529129 | 0.210506 | 0.040866 | 0.269762 | 1.430938 | 0.00134 |
| 0.434522 | 0.22676 | 0.227322 | 0.459481 | 0.356864 | 0.063236 | 0.652643 | 1.194746 | 0.007834 |
| 0.005089 | 0.000648 | 0.001718 | 0.003146 | 0.007714 | 0.00804 | 0.026269 | 0.055462 | 0 |
| 0.008243 | 0.002215 | 0.001533 | 0.004653 | 0.008585 | 0.000477 | 0.027499 | 0.055002 | 0 |
| 0.004397 | 0.002381 | 0.000174 | 0.001633 | 0.003595 | 0.001101 | 0.007676 | 0.017176 | 0 |
| 0.004548 | 0.002296 | 0.000778 | 0.000342 | 0.049164 | 0.004307 | 0.01536 | 0.009607 | 0 |
| 0.006097 | 0.002709 | 0.000211 | 0.00337 | 0.005086 | 0.001576 | 0.020556 | 0.038248 | 0 |
| 1.2683 | 1.184926 | 0.747385 | 1.560258 | 1.129802 | 0.32626 | 2.196701 | 4.421966 | 0.163108 |

| | | | | | | | | |
|---|---|---|---|---|---|---|---|---|
| 0.701463 | 0.517196 | 0.384278 | 0.934419 | 0.71046 | 0.20631 | 1.532057 | 2.547975 | 0.079301 |
| 0.409022 | 0.287299 | 0.227139 | 0.532424 | 0.355772 | 0.078045 | 0.627207 | 1.181671 | 0.056038 |
| 0.37432 | 0.086067 | 0.396645 | 0.529471 | 0.25967 | 0.045173 | 0.285122 | 1.440545 | 0.00134 |
| 0.440619 | 0.229469 | 0.227533 | 0.462851 | 0.36195 | 0.064812 | 0.673199 | 1.232994 | 0.007834 |

| VISC | TEa | PERI | ECT | SSs | SSp | MOp | MOs | VISal |
|---|---|---|---|---|---|---|---|---|
| 0.207832 | 0.194846 | 0.122682 | 0.256187 | 0.184614 | 0.052356 | 0.357095 | 0.718408 | 0.026836 |
| 0.166872 | 0.123966 | 0.092134 | 0.223813 | 0.168955 | 0.049548 | 0.362177 | 0.600108 | 0.019089 |
| 0.163885 | 0.1154 | 0.091927 | 0.214985 | 0.142642 | 0.031164 | 0.250928 | 0.471653 | 0.022697 |
| 0.246527 | 0.05585 | 0.263925 | 0.352771 | 0.140345 | 0.027245 | 0.179851 | 0.954008 | 0.000893 |
| 0.199598 | 0.104162 | 0.104421 | 0.211063 | 0.163926 | 0.029048 | 0.299792 | 0.548808 | 0.003599 |
| 0.000837 | 0.000107 | 0.000283 | 0.000518 | 0.001269 | 0.001323 | 0.004322 | 0.009125 | 0 |
| 0.001984 | 0.000533 | 0.000369 | 0.00112 | 0.002067 | 0.000115 | 0.00662 | 0.01324 | 0 |
| 0.001781 | 0.000964 | 7.05E-05 | 0.000661 | 0.001456 | 0.000446 | 0.003109 | 0.006957 | 0 |
| 0.003032 | 0.001531 | 0.000519 | 0.000228 | 0.032778 | 0.002871 | 0.010241 | 0.006405 | 0 |
| 0.002801 | 0.001244 | 9.69E-05 | 0.001548 | 0.002336 | 0.000724 | 0.009442 | 0.017569 | 0 |

| VISC | TEa | PERI | ECT | SSs | SSp | MOp | MOs | VISal |
|---|---|---|---|---|---|---|---|---|
| 0.196943 | 0.118845 | 0.135018 | 0.251764 | 0.160096 | 0.037872 | 0.289968 | 0.658597 | 0.014623 |
| 0.002087 | 0.000876 | 0.000268 | 0.000815 | 0.007981 | 0.001096 | 0.006747 | 0.010659 | 0 |
| 0.03027 | 0.044717 | 0.065424 | 0.052939 | 0.016668 | 0.010788 | 0.068508 | 0.168088 | 0.010434 |
| 0.000783 | 0.000506 | 0.000168 | 0.000466 | 0.012404 | 0.000972 | 0.002779 | 0.004209 | 0 |

| VISC | TE/PERI/ECT | MOp | MOs | SSs | SSp | VIS | PTLP | RSP |
|---|---|---|---|---|---|---|---|---|
| 0 | 899 | 52 | 125 | 0 | 0 | 30 | 137 | 42 |
| 0 | 430 | 42 | 103 | 0 | 0 | 12 | 63 | 8 |
| 0 | 490 | 15 | 80 | 0 | 0 | 15 | 78 | 22 |
| 0 | 777 | 38 | 109 | 0 | 0 | 19 | 105 | 35 |
| 0 | 15 | 26 | 31 | 0 | 0 | 0 | 24 | 8 |
| 0 | 0 | 4 | 53 | 0 | 0 | 0 | 5 | 5 |
| 0 | 0 | 8 | 25 | 0 | 0 | 0 | 10 | 3 |
| 0 | 9 | 18 | 39 | 0 | 0 | 0 | 23 | 5 |
| 0 | 914 | 78 | 156 | 0 | 0 | 30 | 161 | 50 |
| 0 | 430 | 46 | 156 | 0 | 0 | 12 | 68 | 13 |
| 0 | 490 | 23 | 105 | 0 | 0 | 15 | 88 | 25 |
| 0 | 786 | 56 | 148 | 0 | 0 | 19 | 128 | 40 |

| VISC | TE/PERI/ECT | MOp | MOs | SSs | SSp | VIS | PTLP | RSP |
|---|---|---|---|---|---|---|---|---|
| 0 | 0.623008 | 0.036036 | 0.086625 | 0 | 0 | 0.02079 | 0.094941 | 0.029106 |
| 0 | 0.597222 | 0.058333 | 0.143056 | 0 | 0 | 0.016667 | 0.0875 | 0.011111 |
| 0 | 0.622618 | 0.01906 | 0.101652 | 0 | 0 | 0.01906 | 0.099111 | 0.027954 |
| 0 | 0.621103 | 0.030376 | 0.08713 | 0 | 0 | 0.015188 | 0.083933 | 0.027978 |
| 0 | 0.010395 | 0.018018 | 0.021483 | 0 | 0 | 0 | 0.016632 | 0.005544 |

| | | | | | | | | |
|---|---|---|---|---|---|---|---|---|
| 0 | 0 | 0.005556 | 0.073611 | 0 | 0 | 0 | 0.006944 | 0.006944 |
| 0 | 0 | 0.010165 | 0.031766 | 0 | 0 | 0 | 0.012706 | 0.003812 |
| 0 | 0.007194 | 0.014388 | 0.031175 | 0 | 0 | 0 | 0.018385 | 0.003997 |
| VISC | TE/PERI/ECT | MOp | MOs | SSs | SSp | VIS | PTLP | RSP |
| 0 | 0.615988 | 0.035951 | 0.104616 | 0 | 0 | 0.017926 | 0.091371 | 0.024037 |
| 0 | 0.004397 | 0.012032 | 0.039509 | 0 | 0 | 0 | 0.013667 | 0.005074 |
| 0 | 0.010858 | 0.014295 | 0.022999 | 0 | 0 | 0.002155 | 0.005978 | 0.007477 |
| 0 | 0.004541 | 0.004659 | 0.020108 | 0 | 0 | 0 | 0.004392 | 0.001272 |

| VISC | TEa | ECT/PERI | MOp | MOs | SSs | SSp | VISl | VISm |
|---|---|---|---|---|---|---|---|---|
| 0 | 0.0127 | 0.0232 | 0.0013 | 0.0129 | 0 | 0 | 0.0155 | 0.0167 |
| 0 | 0.0197 | 0.0168 | 0.0007 | 0.0093 | 0 | 0 | 0.0222 | 0.0212 |
| 0 | 0.0171 | 0.0284 | 0.001 | 0.0089 | 0 | 0 | 0.0214 | 0.018 |
| VISC | TEa | ECT/PERI | MOp | MOs | SSs | SSp | VISl | VISm |
| 0 | 0.090007 | 0.164422 | 0.009213 | 0.091425 | 0 | 0 | 0.109851 | 0.118356 |
| 0 | 0.171752 | 0.146469 | 0.006103 | 0.081081 | 0 | 0 | 0.193548 | 0.18483 |
| 0 | 0.152952 | 0.254025 | 0.008945 | 0.079606 | 0 | 0 | 0.191413 | 0.161002 |
| 0 | 0.0165 | 0.0228 | 0.001 | 0.010367 | 0 | 0 | 0.0197 | 0.018633 |
| 0 | 0.034957 | 0.047045 | 0.001407 | 0.005258 | 0 | 0 | 0.038962 | 0.027498 |

| NLOT | COA | TR | ENT | SUB | POST | EP | LA | BLA |
|---|---|---|---|---|---|---|---|---|
| 0.004752 | 0.001129 | 0.016304 | 0.050876 | 0.001812 | 0 | 0.073363 | 0.003846 | 0.048884 |
| 0 | 0 | 0 | 0 | 0 | 0 | 0 | 0 | 0.003623 |
| 0.004752 | 0.001129 | 0.016304 | 0.050876 | 0.001812 | 0 | 0.073363 | 0.003846 | 0.052508 |
| -0.04464 | -0.0106 | -0.15317 | -0.47795 | -0.01702 | 0 | -0.6892 | -0.03613 | -0.45924 |
| 0 | 0 | 0 | 0 | 0 | 0 | 0 | 0 | 0.034038 |

| NLOT | COA | TR | ENT | SUB | POST | EP | LA | BLA |
|---|---|---|---|---|---|---|---|---|
| 0 | 0 | 0 | 2.198864 | 0 | 1.5 | 1.368293 | 1.25 | 1.944444 |
| 0 | 0 | 0 | 0 | 0 | 0 | 0 | | |
| 0 | 0 | 0 | 2.198864 | 0 | 1.5 | 1.368293 | 1.25 | 1.944444 |
| 0 | 0 | 0 | -0.57734 | 0 | -0.39384 | -0.35926 | -0.3282 | -0.51054 |
| 0 | 0 | 0 | 0 | 0 | 0 | 0 | 0 | 0 |

| NLOT | CA1 | ENT | PAR | SUB | EPd | BLA | CP | ACB-core |
|---|---|---|---|---|---|---|---|---|
| 1.072332 | 0.824973 | 6.220958 | 0 | 0 | 1.525038 | 3.754191 | 0.714566 | 0 |
| -0.03065 | -0.02358 | -0.17784 | 0 | 0 | -0.0436 | -0.10732 | -0.02043 | 0 |

| NLOT | CA1 | ENT | PAR | SUB | EPd | BLA | CP | ACB-core |
|---|---|---|---|---|---|---|---|---|
| 0 | 0 | 3.026369 | 0.297356 | 0.10129 | 3.130584 | 0.628602 | 0.025553 | 0.108993 |
| 0 | 0 | -0.20219 | -0.01987 | -0.00677 | -0.20916 | -0.042 | -0.00171 | -0.00728 |

| VISl | VISp | VISpl | VISli | VISpor | VISrl | VISa | VISam | VISpm |
|---|---|---|---|---|---|---|---|---|
| 0.020723 | 0.004806 | 0.04243 | 0.051769 | 0.141794 | 0.014641 | 0.022139 | 0.040607 | 0.024777 |
| 0.022167 | 0.003102 | 0.046593 | 0.041509 | 0.106489 | 0.009247 | 0.011032 | 0.031328 | 0.021105 |
| 0.005179 | 0.000967 | 0.011848 | 0.006449 | 0.028502 | 0.001062 | 0.002551 | 0.006843 | 0.007122 |
| 0.013697 | 0.001327 | 0.023933 | 0.026156 | 0.066365 | 0.006524 | 0.009691 | 0.013195 | 0.008585 |
| 0.000517 | 2.48E-05 | 0.001248 | 0.001463 | 0.003742 | 0.001064 | 0.005023 | 0.004778 | 0.000956 |
| 0.002423 | 0 | 0.004741 | 0.003102 | 0.015643 | 0.012184 | 0.004077 | 0.005099 | 0.001294 |
| 0.00075 | 0 | 0.001313 | 0.006849 | 0.009734 | 0.003873 | 0.004811 | 0.003129 | 0.001692 |

| | | | | | | | | |
|---|---|---|---|---|---|---|---|---|
| 0.00024 | 0 | 0.001132 | 0.00049 | 0.002503 | 0.00108 | 0.001476 | 0.001443 | 4.18E-05 |
| 0.000178 | 0 | 0.00043 | 0.000559 | 0.003662 | 0.002146 | 0.001231 | 0.00192 | 0.000388 |
| 0 | 0 | 0 | 3.86E-06 | 0 | 0.000477 | 0.001515 | 0.000595 | 0.000157 |
| 0.023146 | 0.004806 | 0.047171 | 0.054871 | 0.157437 | 0.026825 | 0.026216 | 0.045706 | 0.026071 |
| 0.022917 | 0.003102 | 0.047906 | 0.048358 | 0.116223 | 0.01312 | 0.015843 | 0.034457 | 0.022797 |
| 0.005419 | 0.000967 | 0.01298 | 0.006939 | 0.031005 | 0.002142 | 0 | 0.008286 | 0.007164 |
| 0.013875 | 0.001327 | 0.024363 | 0.026715 | 0.070027 | 0.00867 | 0 | 0.015115 | 0 |
| 0.000517 | 2.48E-05 | 0.001248 | 0.001467 | 0.003742 | 0.001541 | 0.006538 | 0 | 0.001113 |

| VISl | VISp | VISpl | VISli | VISpor | VISrl | VISa | VISam | VISpm |
|---|---|---|---|---|---|---|---|---|
| 0.0209586 | 0.0048606 | 0.0429124 | 0.0523576 | 0.143406 | 0.0148075 | 0.0223907 | 0.0410687 | 0.0250587 |
| 0.0413329 | 0.005784 | 0.086878 | 0.0773983 | 0.1985609 | 0.0172421 | 0.0205704 | 0.0584146 | 0.0393527 |
| 0.0264836 | 0.0049449 | 0.0605865 | 0.0329779 | 0.1457493 | 0.0054307 | 0.0130449 | 0.0349927 | 0.0364194 |
| 0.025139 | 0.0024355 | 0.0439258 | 0.0480058 | 0.1218039 | 0.0119739 | 0.0177865 | 0.0242176 | 0.0157566 |
| 0.0018532 | 8.89E-05 | 0.0044735 | 0.0052442 | 0.0134133 | 0.0038139 | 0.0180051 | 0.0171269 | 0.0034268 |
| 0.0024505 | 0 | 0.0047949 | 0.0031373 | 0.0158208 | 0.0123225 | 0.0041234 | 0.005157 | 0.0013087 |
| 0.0013985 | 0 | 0.0024482 | 0.0127707 | 0.0181502 | 0.0072217 | 0.0089707 | 0.0058344 | 0.0031549 |
| 0.0012273 | 0 | 0.0057887 | 0.0025057 | 0.0127995 | 0.0055227 | 0.0075477 | 0.007379 | 0.0002138 |
| 0.0003267 | 0 | 0.0007892 | 0.001026 | 0.0067211 | 0.0039387 | 0.0022593 | 0.0035239 | 0.0007121 |
| 0 | 0 | 0 | 1.384E-05 | 0 | 0.0017098 | 0.0054306 | 0.0021328 | 0.0005628 |
| 0.023409 | 0.004861 | 0.047707 | 0.055495 | 0.159227 | 0.02713 | 0.026514 | 0.046226 | 0.026367 |
| 0.042731 | 0.005784 | 0.089326 | 0.090169 | 0.216711 | 0.024464 | 0.029541 | 0.064249 | 0.042508 |
| 0.027711 | 0.004945 | 0.066375 | 0.035484 | 0.158549 | 0.010953 | 0.020593 | 0.042372 | 0.036633 |
| 0.025466 | 0.002436 | 0.044715 | 0.049032 | 0.128525 | 0.015913 | 0.020046 | 0.027742 | 0.016469 |
| 0.001853 | 8.89E-05 | 0.004474 | 0.005258 | 0.013413 | 0.005524 | 0.023436 | 0.01926 | 0.00399 |

| VISl | VISp | VISpl | VISli | VISpor | VISrl | VISa | VISam | VISpm |
|---|---|---|---|---|---|---|---|---|
| 0.023153 | 0.003623 | 0.047755 | 0.043197 | 0.124587 | 0.010654 | 0.01836 | 0.035164 | 0.024003 |
| 0.001081 | 0 | 0.002764 | 0.003891 | 0.010698 | 0.006143 | 0.005666 | 0.004805 | 0.00119 |
| 0.012681 | 0.00209 | 0.026858 | 0.023755 | 0.06106 | 0.005224 | 0.003157 | 0.014287 | 0.013284 |
| 0.000864 | 0 | 0.002232 | 0.004573 | 0.00658 | 0.003585 | 0.002388 | 0.001822 | 0.001044 |

| VISl | VISp | VISpl | VISli | VISpor | VISrl | VISa | VISam | VISpm |
|---|---|---|---|---|---|---|---|---|
| 0.403597 | 2.176811 | 0.588285 | 0.150757 | 0.843701 | 0.231271 | 0.411257 | 0.364693 | 0.53244 |
| 0.166889 | 1.208218 | 0.328436 | 0.067988 | 0.416989 | 0.081113 | 0.197538 | 0.243655 | 0.335486 |
| 0.098867 | 0.569094 | 0.150642 | 0.045779 | 0.200801 | 0.059781 | 0.104331 | 0.118376 | 0.160717 |
| 0.011343 | 0.044599 | 0.019532 | 0.00668 | 0.094473 | 0.00045 | 0.013134 | 0.007035 | 0.003467 |
| 0.016821 | 0.06933 | 0.038259 | 0.010196 | 0.091525 | 0.010232 | 0.025084 | 0.025023 | 0.029953 |
| 0 | 0 | 0 | 0 | 0 | 0 | 0 | 0 | 0 |
| 0 | 0 | 0 | 0 | 0 | 0 | 0 | 0 | 0 |
| 0 | 0 | 0 | 0 | 0 | 0 | 0 | 0 | 0 |
| 0 | 0 | 0 | 0 | 0 | 0 | 0 | 0 | 0 |
| 0 | 0 | 0 | 0 | 0 | 0 | 0 | 0 | 0 |
| 0.403597 | 2.176811 | 0.588285 | 0.150757 | 0.843701 | 0.231271 | 0.411257 | 0.364693 | 0.53244 |

| | | | | | | | | |
|---|---|---|---|---|---|---|---|---|
| 0.166889 | 1.208218 | 0.328436 | 0.067988 | 0.416989 | 0.081113 | 0.197538 | 0.243655 | 0.335486 |
| 0.098867 | 0.569094 | 0.150642 | 0.045779 | 0.200801 | 0.059781 | 0.104331 | 0.118376 | 0.160717 |
| 0.011343 | 0.044599 | 0.019532 | 0.00668 | 0.094473 | 0.00045 | 0.013134 | 0.007035 | 0.003467 |
| 0.016821 | 0.06933 | 0.038259 | 0.010196 | 0.091525 | 0.010232 | 0.025084 | 0.025023 | 0.029953 |

| VISl | VISp | VISpl | VISli | VISpor | VISrl | VISa | VISam | VISpm |
|---|---|---|---|---|---|---|---|---|
| 0.066403 | 0.358144 | 0.096789 | 0.024804 | 0.138812 | 0.03805 | 0.067663 | 0.060002 | 0.087601 |
| 0.040173 | 0.290842 | 0.079061 | 0.016366 | 0.100377 | 0.019525 | 0.047551 | 0.058653 | 0.080758 |
| 0.040044 | 0.230499 | 0.061014 | 0.018542 | 0.08133 | 0.024213 | 0.042257 | 0.047946 | 0.065095 |
| 0.007562 | 0.029734 | 0.013022 | 0.004454 | 0.062985 | 0.0003 | 0.008756 | 0.00469 | 0.002311 |
| 0.007727 | 0.031847 | 0.017574 | 0.004684 | 0.042042 | 0.0047 | 0.011522 | 0.011494 | 0.013759 |
| 0 | 0 | 0 | 0 | 0 | 0 | 0 | 0 | 0 |
| 0 | 0 | 0 | 0 | 0 | 0 | 0 | 0 | 0 |
| 0 | 0 | 0 | 0 | 0 | 0 | 0 | 0 | 0 |
| 0 | 0 | 0 | 0 | 0 | 0 | 0 | 0 | 0 |
| 0 | 0 | 0 | 0 | 0 | 0 | 0 | 0 | 0 |

| VISl | VISp | VISpl | VISli | VISpor | VISrl | VISa | VISam | VISpm |
|---|---|---|---|---|---|---|---|---|
| 0.032382 | 0.188213 | 0.053492 | 0.01377 | 0.085109 | 0.017358 | 0.03555 | 0.036557 | 0.049905 |
| 0 | 0 | 0 | 0 | 0 | 0 | 0 | 0 | 0 |
| 0.022364 | 0.134732 | 0.033205 | 0.008008 | 0.033093 | 0.013646 | 0.02243 | 0.023712 | 0.035143 |
| 0 | 0 | 0 | 0 | 0 | 0 | 0 | 0 | 0 |

| AUD | AON | TTd | PIR | NLOT | TR | CA1 | ENTl | ENTm |
|---|---|---|---|---|---|---|---|---|
| 291 | 430 | 232 | 126 | 24 | 89 | 201 | 618 | 10 |
| 168 | 84 | 177 | 39 | 7 | 64 | 99 | 316 | 14 |
| 140 | 103 | 122 | 28 | 13 | 48 | 77 | 304 | 9 |
| 265 | 298 | 176 | 59 | 18 | 78 | 148 | 503 | 13 |
| 0 | 21 | 11 | 0 | 0 | 0 | 0 | 2 | 0 |
| 0 | 11 | 0 | 0 | 0 | 0 | 0 | 9 | 0 |
| 0 | 9 | 5 | 0 | 0 | 0 | 0 | 5 | 0 |
| 0 | 12 | 12 | 0 | 0 | 0 | 0 | 6 | 0 |
| 291 | 451 | 243 | 126 | 24 | 89 | 201 | 620 | 10 |
| 168 | 95 | 177 | 39 | 7 | 64 | 99 | 325 | 14 |
| 140 | 112 | 127 | 28 | 13 | 48 | 77 | 309 | 9 |
| 265 | 310 | 188 | 59 | 18 | 78 | 148 | 509 | 13 |

| AUD | AON | TTd | PIR | NLOT | TR | CA1 | ENTl | ENTm |
|---|---|---|---|---|---|---|---|---|
| 0.201663 | 0.29799 | 0.160776 | 0.087318 | 0.016632 | 0.061677 | 0.139293 | 0.428274 | 0.00693 |
| 0.233333 | 0.116667 | 0.245833 | 0.054167 | 0.009722 | 0.088889 | 0.1375 | 0.438889 | 0.019444 |
| 0.177891 | 0.130877 | 0.155019 | 0.035578 | 0.016518 | 0.060991 | 0.09784 | 0.386277 | 0.011436 |
| 0.211831 | 0.238209 | 0.140687 | 0.047162 | 0.014388 | 0.06235 | 0.118305 | 0.402078 | 0.010392 |
| 0 | 0.014553 | 0.007623 | 0 | 0 | 0 | 0 | 0.001386 | 0 |

| | | | | | | | | |
|---|---|---|---|---|---|---|---|---|
| 0 | 0.015278 | 0 | 0 | 0 | 0 | 0 | 0.0125 | 0 |
| 0 | 0.011436 | 0.006353 | 0 | 0 | 0 | 0 | 0.006353 | 0 |
| 0 | 0.009592 | 0.009592 | 0 | 0 | 0 | 0 | 0.004796 | 0 |

| AUD | AON | TTd | PIR | NLOT | TR | CA1 | ENTl | ENTm |
|---|---|---|---|---|---|---|---|---|
| 0.206179 | 0.195936 | 0.175579 | 0.056056 | 0.014315 | 0.068477 | 0.123235 | 0.41388 | 0.01205 |
| 0 | 0.012715 | 0.005892 | 0 | 0 | 0 | 0 | 0.006259 | 0 |
| 0.019937 | 0.075363 | 0.041216 | 0.019231 | 0.002798 | 0.011795 | 0.016812 | 0.02082 | 0.004583 |
| 0 | 0.002309 | 0.003592 | 0 | 0 | 0 | 0 | 0.004026 | 0 |

| VISp | PTLP | RSPagl | RSPd | RSPv | AUD | AON | TTd | PIR |
|---|---|---|---|---|---|---|---|---|
| 0.0147 | 0.0396 | 0.0878 | 0.0421 | 0.0366 | 0.0008 | 0.0066 | 0.0076 | 0.0011 |
| 0.0203 | 0.0352 | 0.0936 | 0.0944 | 0.0496 | 0.0002 | 0.0059 | 0.0039 | 0.0013 |
| 0.0194 | 0.0495 | 0.0754 | 0.0664 | 0.0387 | 0.0009 | 0.005 | 0.0054 | 0.0019 |
| VISp | PTLP | RSPagl | RSPd | RSPv | AUD | AON | TTd | PIR |
| 0.104181 | 0.280652 | 0.622254 | 0.29837 | 0.259391 | 0.00567 | 0.046775 | 0.053863 | 0.007796 |
| 0.176983 | 0.306888 | 0.816042 | 0.823017 | 0.432432 | 0.001744 | 0.051439 | 0.034002 | 0.011334 |
| 0.173524 | 0.442755 | 0.674419 | 0.593918 | 0.346154 | 0.00805 | 0.044723 | 0.048301 | 0.016995 |
| 0.018133 | 0.041433 | 0.0856 | 0.067633 | 0.041633 | 0.000633 | 0.005833 | 0.005633 | 0.001433 |
| 0.033534 | 0.071044 | 0.081875 | 0.214758 | 0.070644 | 0.0026 | 0.00281 | 0.008366 | 0.003789 |

| BMA | CP | ACB | FS | OT | LS | AAA | MEA | sAMY |
|---|---|---|---|---|---|---|---|---|
| 0.012904 | 0.023551 | 0.006563 | 0.003623 | 0.002034 | 0.001812 | 0.014033 | 0.001812 | 0.042098 |
| 0 | 0 | 0.002257 | 0 | 0 | 0.005435 | 0 | 0 | 0 |
| 0.012904 | 0.023551 | 0.008821 | 0.003623 | 0.002034 | 0.007246 | 0.014033 | 0.001812 | 0.042098 |
| -0.12123 | -0.22124 | -0.06166 | -0.03404 | -0.01911 | -0.01702 | -0.13183 | -0.01702 | -0.39549 |
| 0 | 0 | 0.021206 | 0 | 0 | 0.051056 | 0 | 0 | 0 |

| BMA | CP | ACB | FS | OT | LS | AAA | MEA | sAMY |
|---|---|---|---|---|---|---|---|---|
| 1.041667 | 0.5 | 1 | 1 | 0.5 | 0 | 0 | 0 | 0.5 |
| | 0 | 0 | 0.214286 | 0 | 0 | 0 | 0 | 0 |
| 1.041667 | 0.5 | 1 | 1.214286 | 0.5 | 0 | 0 | 0 | 0.5 |
| -0.2735 | -0.13128 | -0.26256 | -0.26256 | -0.13128 | 0 | 0 | 0 | -0.13128 |
| 0 | 0 | 0 | 0.056263 | 0 | 0 | 0 | 0 | 0 |

| ACB-shell | GPe | SI | PO | MD | CM | PF | LHA | STN |
|---|---|---|---|---|---|---|---|---|
| 0 | 0.47149 | 1.851476 | 0.247704 | 0.48302 | 0.349156 | 0.691726 | 0.384936 | 0.217543 |
| 0 | -0.01348 | -0.05293 | -0.00708 | -0.01381 | -0.00998 | -0.01977 | -0.011 | -0.00622 |

| ACB-shell | GPe | SI | PO | MD | CM | PF | LHA | STN |
|---|---|---|---|---|---|---|---|---|
| 0.351354 | 0 | 0 | 0 | 0 | 0 | 0 | 0 | 0 |
| -0.02347 | 0 | 0 | 0 | 0 | 0 | 0 | 0 | 0 |

| RSPagl | RSPd | RSPv | AUDd | AUDp | AUDpo | AUDv | AON | TT |
|---|---|---|---|---|---|---|---|---|
| 0.029102 | 0.009283 | 0.006303 | 0.072685 | 0.054018 | 0.068985 | 0.136281 | 0.433017 | 0.376667 |
| 0.017886 | 0.00661 | 0.00255 | 0.0567 | 0.057924 | 0.068832 | 0.137391 | 0.120581 | 0.053046 |
| 0.005822 | 0.002332 | 0.000789 | 0.016315 | 0.014233 | 0.018466 | 0.033151 | 0.038017 | 0.006749 |
| 0.01279 | 0.004727 | 0.001324 | 0.034761 | 0.031283 | 0.043839 | 0.073153 | 0.068837 | 0.022842 |
| 0.005934 | 0.005453 | 0.005185 | 0.003958 | 0.001813 | 0.001732 | 0.005227 | 0.011516 | 0.004339 |
| 0.006596 | 0.003861 | 0.001916 | 0.007026 | 0.002179 | 0.009498 | 0.006516 | 0.08209 | 0.001982 |
| 0.007979 | 0.003699 | 0.00144 | 0.002931 | 0.001201 | 0.006249 | 0.00967 | 0.006004 | 0.000651 |

| | | | | | | | | |
|---|---|---|---|---|---|---|---|---|
| 0.002763 | 0.001471 | 0.000282 | 0.000778 | 4.07E-05 | 0.000122 | 0.002708 | 0.001177 | 0.000404 |
| 0.005162 | 0.003359 | 0.001513 | 0.002224 | 8.21E-05 | 0.001756 | 0.004234 | 0.004117 | 0.000256 |
| 0.00176 | 0.001012 | 0.000269 | 0.000648 | 0 | 0 | 0.000332 | 0.003228 | 0 |
| 0.035698 | 0.013144 | 0.008219 | 0.079711 | 0.056197 | 0.078483 | 0.142797 | 0.515107 | 0.378649 |
| 0.025865 | 0.010309 | 0.00399 | 0.059631 | 0.059125 | 0.075081 | 0.147061 | 0.126585 | 0.053697 |
| 0 | 0.003803 | 0.001071 | 0.017093 | 0.014274 | 0.018588 | 0.035859 | 0.039194 | 0.007153 |
| 0 | 0.008086 | 0.002837 | 0.036985 | 0.031365 | 0.045595 | 0.077387 | 0.072954 | 0.023098 |
| 0 | 0.006465 | 0.005454 | 0.004606 | 0 | 0.001732 | 0.005559 | 0.014744 | 0.004339 |

| RSPagl | RSPd | RSPv | AUDd | AUDp | AUDpo | AUDv | AON | TT |
|---|---|---|---|---|---|---|---|---|
| 0.0294329 | 0.0093885 | 0.0063747 | 0.0735113 | 0.0546321 | 0.0697693 | 0.1378304 | 0.4379399 | 0.3809493 |
| 0.0333505 | 0.0123251 | 0.0047548 | 0.1057236 | 0.1080059 | 0.1283451 | 0.2561812 | 0.224837 | 0.0989103 |
| 0.0297717 | 0.011925 | 0.0040347 | 0.0834292 | 0.0727826 | 0.0944287 | 0.1695226 | 0.1944057 | 0.034512 |
| 0.0234743 | 0.0086758 | 0.00243 | 0.0637991 | 0.0574157 | 0.0804605 | 0.1342624 | 0.126341 | 0.0419234 |
| 0.0212706 | 0.0195465 | 0.0185858 | 0.0141876 | 0.0064988 | 0.0062084 | 0.0187364 | 0.0412795 | 0.0155533 |
| 0.006671 | 0.0039049 | 0.0019378 | 0.0071059 | 0.0022038 | 0.009606 | 0.0065901 | 0.0830233 | 0.0020045 |
| 0.0148778 | 0.0068972 | 0.002685 | 0.0054652 | 0.0022394 | 0.011652 | 0.0180308 | 0.0111951 | 0.0012139 |
| 0.014129 | 0.0075222 | 0.001442 | 0.0039784 | 0.0002081 | 0.0006239 | 0.0138478 | 0.0060188 | 0.0020659 |
| 0.0094741 | 0.006165 | 0.0027769 | 0.0040818 | 0.0001507 | 0.0032229 | 0.0077709 | 0.0075562 | 0.0004699 |
| 0.0063088 | 0.0036276 | 0.0009642 | 0.0023228 | 0 | 0 | 0.0011901 | 0.0115709 | 0 |
| 0.036104 | 0.013293 | 0.008312 | 0.080617 | 0.056836 | 0.079375 | 0.14442 | 0.520963 | 0.382954 |
| 0.048228 | 0.019222 | 0.00744 | 0.111189 | 0.110245 | 0.139997 | 0.274212 | 0.236032 | 0.100124 |
| 0.043901 | 0.019447 | 0.005477 | 0.087408 | 0.072991 | 0.095053 | 0.18337 | 0.200424 | 0.036578 |
| 0.032948 | 0.014841 | 0.005207 | 0.067881 | 0.057566 | 0.083683 | 0.142033 | 0.133897 | 0.042393 |
| 0.027579 | 0.023174 | 0.01955 | 0.01651 | 0.006499 | 0.006208 | 0.019926 | 0.05285 | 0.015553 |

| RSPagl | RSPd | RSPv | AUDd | AUDp | AUDpo | AUDv | AON | TT |
|---|---|---|---|---|---|---|---|---|
| 0.02746 | 0.012372 | 0.007236 | 0.06813 | 0.059867 | 0.075842 | 0.143307 | 0.204961 | 0.11437 |
| 0.010292 | 0.005623 | 0.001961 | 0.004591 | 0.00096 | 0.005021 | 0.009486 | 0.023873 | 0.001151 |
| 0.00443 | 0.003853 | 0.005815 | 0.030347 | 0.032759 | 0.040023 | 0.076233 | 0.132483 | 0.136157 |
| 0.003617 | 0.001578 | 0.0007 | 0.001604 | 0.001032 | 0.004749 | 0.005868 | 0.029651 | 0.00082 |

| RSPagl | RSPd | RSPv | AUDd | AUDp | AUDpo | AUDv | AON | PIR |
|---|---|---|---|---|---|---|---|---|
| 1.387355 | 2.242169 | 1.60853 | 0.092367 | 0.093822 | 0.081363 | 0.253331 | 0.943011 | 0 |
| 0.863265 | 1.245842 | 0.873614 | 0.056618 | 0.05842 | 0.047889 | 0.074451 | 1.958228 | 0 |
| 0.411062 | 0.677783 | 0.560077 | 0.029469 | 0.022884 | 0.021789 | 0.047203 | 0.979395 | 0 |
| 0.040174 | 0.032847 | 0.026567 | 0.015612 | 0.001084 | 8.84E-05 | 0.012582 | 0.135618 | 0 |
| 0.113666 | 0.146523 | 0.052181 | 0.012506 | 0.008449 | 0.004191 | 0.036867 | 1.703207 | 0 |
| 0.007269 | 0.008084 | 0.008795 | 0 | 0 | 0 | 0.000398 | 0.012448 | 0.003189 |
| 0.004575 | 0.012791 | 0.003716 | 0 | 0 | 0 | 0.000204 | 0.000298 | 0.00098 |
| 0.004606 | 0.006357 | 0.003257 | 0 | 0 | 0 | 0.000492 | 0.006365 | 0.002575 |
| 0.007705 | 0.003536 | 0.000155 | 0 | 0 | 0 | 0.00092 | 0 | 0.039114 |
| 0.001419 | 0.000636 | 0.000141 | 0 | 0 | 0 | 0.000488 | 0.020637 | 0.01719 |
| 1.394624 | 2.250253 | 1.617325 | 0.092367 | 0.093822 | 0.081363 | 0.253729 | 0.955459 | 0.003189 |

| | | | | | | | | |
|---|---|---|---|---|---|---|---|---|
| 0.86784 | 1.258633 | 0.87733 | 0.056618 | 0.05842 | 0.047889 | 0.074655 | 1.958526 | 0.00098 |
| 0.415668 | 0.68414 | 0.563334 | 0.029469 | 0.022884 | 0.021789 | 0.047695 | 0.98576 | 0.002575 |
| 0.047879 | 0.036383 | 0.026722 | 0.015612 | 0.001084 | 8.84E-05 | 0.013502 | 0.135618 | 0.039114 |
| 0.115085 | 0.147159 | 0.052322 | 0.012506 | 0.008449 | 0.004191 | 0.037355 | 1.723844 | 0.01719 |

| RSPagl | RSPd | RSPv | AUDd | AUDp | AUDpo | AUDv | AON | PIR |
|---|---|---|---|---|---|---|---|---|
| 0.228257 | 0.368898 | 0.264647 | 0.015197 | 0.015436 | 0.013386 | 0.04168 | 0.155151 | 0 |
| 0.207805 | 0.299899 | 0.210296 | 0.013629 | 0.014063 | 0.011528 | 0.017922 | 0.471384 | 0 |
| 0.166492 | 0.274521 | 0.226847 | 0.011936 | 0.009269 | 0.008825 | 0.019119 | 0.396683 | 0 |
| 0.026784 | 0.021899 | 0.017712 | 0.010409 | 0.000723 | 5.89E-05 | 0.008388 | 0.090417 | 0 |
| 0.052213 | 0.067305 | 0.023969 | 0.005745 | 0.003881 | 0.001925 | 0.016935 | 0.78237 | 0 |
| 0.001196 | 0.00133 | 0.001447 | 0 | 0 | 0 | 6.55E-05 | 0.002048 | 0.000525 |
| 0.001101 | 0.003079 | 0.000895 | 0 | 0 | 0 | 4.91E-05 | 7.17E-05 | 0.000236 |
| 0.001866 | 0.002575 | 0.001319 | 0 | 0 | 0 | 0.000199 | 0.002578 | 0.001043 |
| 0.005137 | 0.002357 | 0.000103 | 0 | 0 | 0 | 0.000613 | 0 | 0.026077 |
| 0.000652 | 0.000292 | 6.48E-05 | 0 | 0 | 0 | 0.000224 | 0.00948 | 0.007896 |

| RSPagl | RSPd | RSPv | AUDd | AUDp | AUDpo | AUDv | AON | PIR |
|---|---|---|---|---|---|---|---|---|
| 0.13631 | 0.206504 | 0.148694 | 0.011383 | 0.008674 | 0.007145 | 0.020809 | 0.379201 | 0 |
| 0.00199 | 0.001927 | 0.000766 | 0 | 0 | 0 | 0.00023 | 0.002835 | 0.007155 |
| 0.081909 | 0.136511 | 0.105887 | 0.003245 | 0.00568 | 0.005262 | 0.0111 | 0.246959 | 0 |
| 0.00162 | 0.000996 | 0.000586 | 0 | 0 | 0 | 0.000204 | 0.003479 | 0.009877 |

| | | | | | PVT/CL/IM | | | |
|---|---|---|---|---|---|---|---|---|
| LA | BLA | BMA | NDB | AM | D | RE | PF | LH |
| 60 | 412 | 28 | 410 | 22 | 279 | 101 | 24 | 5 |
| 39 | 204 | 18 | 223 | 33 | 187 | 35 | 9 | 1 |
| 28 | 218 | 13 | 232 | 15 | 152 | 23 | 13 | 2 |
| 41 | 285 | 30 | 366 | 19 | 235 | 66 | 23 | 4 |
| 0 | 51 | 0 | 6 | 0 | 0 | 0 | 0 | 0 |
| 0 | 26 | 0 | 0 | 0 | 0 | 0 | 0 | 0 |
| 0 | 16 | 0 | 3 | 0 | 0 | 0 | 0 | 0 |
| 0 | 35 | 0 | 4 | 0 | 0 | 0 | 0 | 0 |
| 60 | 463 | 28 | 416 | 22 | 279 | 101 | 24 | 5 |
| 39 | 230 | 18 | 223 | 33 | 187 | 35 | 9 | 1 |
| 28 | 234 | 13 | 235 | 15 | 152 | 23 | 13 | 2 |
| 41 | 320 | 30 | 370 | 19 | 235 | 66 | 23 | 4 |

| | | | | | PVT/CL/IM | | | |
|---|---|---|---|---|---|---|---|---|
| LA | BLA | BMA | NDB | AM | D | RE | PF | LH |
| 0.04158 | 0.285516 | 0.019404 | 0.28413 | 0.015246 | 0.193347 | 0.069993 | 0.016632 | 0.003465 |
| 0.054167 | 0.283333 | 0.025 | 0.309722 | 0.045833 | 0.259722 | 0.048611 | 0.0125 | 0.001389 |
| 0.035578 | 0.277001 | 0.016518 | 0.29479 | 0.01906 | 0.193139 | 0.029225 | 0.016518 | 0.002541 |
| 0.032774 | 0.227818 | 0.023981 | 0.292566 | 0.015188 | 0.18785 | 0.052758 | 0.018385 | 0.003197 |
| 0 | 0.035343 | 0 | 0.004158 | 0 | 0 | 0 | 0 | 0 |

| | | | | | | | | |
|---|---|---|---|---|---|---|---|---|
| 0 | 0.036111 | 0 | 0 | 0 | 0 | 0 | 0 | 0 |
| 0 | 0.02033 | 0 | 0.003812 | 0 | 0 | 0 | 0 | 0 |
| 0 | 0.027978 | 0 | 0.003197 | 0 | 0 | 0 | 0 | 0 |
| | | | | | PVT/CL/IM | | | |
| LA | BLA | BMA | NDB | AM | D | RE | PF | LH |
| 0.041025 | 0.268417 | 0.021226 | 0.295302 | 0.023832 | 0.208514 | 0.050147 | 0.016009 | 0.002648 |
| 0 | 0.029941 | 0 | 0.002792 | 0 | 0 | 0 | 0 | 0 |
| 0.008227 | 0.023648 | 0.003439 | 0.009226 | 0.012799 | 0.029647 | 0.014498 | 0.002157 | 0.000801 |
| 0 | 0.006393 | 0 | 0.001648 | 0 | 0 | 0 | 0 | 0 |

| ENTl | ENTm | PAR | POST | PRE | SUBv | LA | BLA | BMA |
|---|---|---|---|---|---|---|---|---|
| 0.1184 | 0.0449 | 0.0127 | 0.0212 | 0.0326 | 0.0081 | 0.0014 | 0.0081 | 0.0014 |
| 0.0986 | 0.0549 | 0.0245 | 0.0256 | 0.0381 | 0.0071 | 0.0008 | 0.0041 | 0.0017 |
| 0.1063 | 0.0495 | 0.0145 | 0.025 | 0.0284 | 0.0049 | 0.0003 | 0.0054 | 0.0007 |
| ENTl | ENTm | PAR | POST | PRE | SUBv | LA | BLA | BMA |
| 0.839121 | 0.318214 | 0.090007 | 0.150248 | 0.231042 | 0.057406 | 0.009922 | 0.057406 | 0.009922 |
| 0.859634 | 0.47864 | 0.213601 | 0.223191 | 0.332171 | 0.061901 | 0.006975 | 0.035745 | 0.014821 |
| 0.950805 | 0.442755 | 0.129696 | 0.223614 | 0.254025 | 0.043828 | 0.002683 | 0.048301 | 0.006261 |
| 0.107767 | 0.049767 | 0.017233 | 0.023933 | 0.033033 | 0.0067 | 0.000833 | 0.005867 | 0.001267 |
| 0.048541 | 0.068746 | 0.051522 | 0.034486 | 0.043285 | 0.007682 | 0.002972 | 0.00888 | 0.003507 |

| Gpe | SI | MA | NDB | VENT | SPF | PO | ATN | MED |
|---|---|---|---|---|---|---|---|---|
| 0.063054 | 0.076433 | 0.01381 | 0.005198 | 0.006103 | 0.001812 | 0.017627 | 0.005435 | 0.01969 |
| 0 | 0 | 0 | 0 | 0 | 0 | 0 | 0 | 0 |
| 0.063054 | 0.076433 | 0.01381 | 0.005198 | 0.006103 | 0.001812 | 0.017627 | 0.005435 | 0.01969 |
| -0.59235 | -0.71804 | -0.12973 | -0.04883 | -0.05734 | -0.01702 | -0.1656 | -0.05106 | -0.18498 |
| 0 | 0 | 0 | 0 | 0 | 0 | 0 | 0 | 0 |

| Gpe | SI | MA | NDB | VENT | SPF | PO | ATN | MED |
|---|---|---|---|---|---|---|---|---|
| 1.272727 | 0.5 | 0 | 0 | 1.831169 | 0 | 0.5 | 0 | 0 |
| 0 | 0 | 0 | 0 | 0.207143 | 0 | 0 | 0 | 0 |
| 1.272727 | 0.5 | 0 | 0 | 2.038312 | 0 | 0.5 | 0 | 0 |
| -0.33417 | -0.13128 | 0 | 0 | -0.4808 | 0 | -0.13128 | 0 | 0 |
| 0 | 0 | 0 | 0 | 0.054388 | 0 | 0 | 0 | 0 |

| DR/CS | INC |
|---|---|
| 0.260363 | 0.663288 |
| -0.00744 | -0.01896 |

| DR/CS | INC |
|---|---|
| 0 | 0 |
| 0 | 0 |

| DP | PIR | CA1 | ENTl | ENTm | EPd | LA | BLA | MEA |
|---|---|---|---|---|---|---|---|---|
| 0.217497 | 0.648806 | 0.048951 | 0.440043 | 0.094016 | 0.98735 | 0.238302 | 0.642882 | 0.036148 |
| 0.044724 | 0.172794 | 0.028412 | 0.240256 | 0.055789 | 0.426412 | 0.174455 | 0.394583 | 0.019781 |
| 0.005695 | 0.027451 | 0.00936 | 0.064863 | 0.020275 | 0.095962 | 0.052864 | 0.146757 | 0.002403 |
| 0.025889 | 0.088754 | 0.02349 | 0.154229 | 0.039765 | 0.174444 | 0.110668 | 0.363204 | 0.008072 |
| 0.004154 | 0.008237 | 0.016341 | 0.022786 | 0.004026 | 0.016821 | 0.009437 | 0.111521 | 0.000899 |
| 0.021679 | 0.006061 | 0 | 0.035844 | 0 | 0.001409 | 0 | 0.26035 | 0 |
| 0.000495 | 0.001765 | 0 | 0.020471 | 0 | 0.00093 | 0 | 0.141721 | 0 |

| | | | | | | | | |
|---|---|---|---|---|---|---|---|---|
| 0 | 0.00155 | 0 | 0.006263 | 0 | 0.000171 | 0 | 0.048798 | 0 |
| 0.001566 | 0.002472 | 0 | 0.024517 | 0 | 0.003198 | 0 | 0.136545 | 0 |
| 0 | 0.000263 | 0 | 0.003659 | 0 | 0.000209 | 0 | 0.020375 | 0 |
| 0.239176 | 0.654867 | 0.048951 | 0.475887 | 0.094016 | 0.988759 | 0.238302 | 0.903232 | 0.036148 |
| 0.045219 | 0.174559 | 0.028412 | 0.260727 | 0.055789 | 0.427342 | 0.174455 | 0.536304 | 0.019781 |
| 0.005695 | 0.029001 | 0 | 0.071126 | 0.020275 | 0.096133 | 0.052864 | 0.195555 | 0.002403 |
| 0.027455 | 0.091226 | 0.02349 | 0.178746 | 0.039765 | 0.177642 | 0.110668 | 0.499749 | 0.008072 |
| 0.004154 | 0.0085 | 0.016341 | 0.026445 | 0.004026 | 0.01703 | 0.009437 | 0.131896 | 0.000899 |

| DP | PIR | CA1 | ENTl | ENTm | EPd | LA | BLA | MEA |
|---|---|---|---|---|---|---|---|---|
| 0.2199697 | 0.6561821 | 0.0495075 | 0.4450458 | 0.0950848 | 0.998575 | 0.2410112 | 0.6501908 | 0.036559 |
| 0.083393 | 0.3221941 | 0.0529774 | 0.4479847 | 0.104025 | 0.7950938 | 0.3252913 | 0.735745 | 0.0368839 |
| 0.0291222 | 0.1403748 | 0.0478638 | 0.3316867 | 0.1036793 | 0.4907162 | 0.270328 | 0.7504641 | 0.0122881 |
| 0.0475157 | 0.1628959 | 0.0431127 | 0.2830664 | 0.0729833 | 0.3201683 | 0.2031161 | 0.6666116 | 0.0148151 |
| 0.0148902 | 0.0295258 | 0.0585749 | 0.0816773 | 0.0144313 | 0.0602955 | 0.0338273 | 0.3997512 | 0.0032225 |
| 0.0219255 | 0.0061299 | 0 | 0.0362515 | 0 | 0.001425 | 0 | 0.2633099 | 0 |
| 0.000923 | 0.003291 | 0 | 0.0381705 | 0 | 0.0017341 | 0 | 0.264255 | 0 |
| 0 | 0.0079262 | 0 | 0.0320268 | 0 | 0.0008744 | 0 | 0.2495359 | 0 |
| 0.0028742 | 0.004537 | 0 | 0.0449976 | 0 | 0.0058695 | 0 | 0.2506098 | 0 |
| 0 | 0.0009427 | 0 | 0.0131158 | 0 | 0.0007492 | 0 | 0.073035 | 0 |
| 0.241895 | 0.662312 | 0.049508 | 0.481297 | 0.095085 | 1 | 0.241011 | 0.913501 | 0.036559 |
| 0.084316 | 0.325485 | 0.052977 | 0.486155 | 0.104025 | 0.796828 | 0.325291 | 1 | 0.036884 |
| 0.029122 | 0.148301 | 0.047864 | 0.363714 | 0.103679 | 0.491591 | 0.270328 | 1 | 0.012288 |
| 0.05039 | 0.167433 | 0.043113 | 0.328064 | 0.072983 | 0.326038 | 0.203116 | 0.917221 | 0.014815 |
| 0.01489 | 0.030469 | 0.058575 | 0.094793 | 0.014431 | 0.061045 | 0.033827 | 0.472786 | 0.003222 |

| DP | PIR | CA1 | ENTl | ENTm | EPd | LA | BLA | MEA |
|---|---|---|---|---|---|---|---|---|
| 0.078978 | 0.262235 | 0.050407 | 0.317892 | 0.078041 | 0.53297 | 0.214715 | 0.640553 | 0.020754 |
| 0.005145 | 0.004565 | 0 | 0.032912 | 0 | 0.00213 | 0 | 0.220149 | 0 |
| 0.074136 | 0.218019 | 0.005173 | 0.134382 | 0.033748 | 0.333463 | 0.09886 | 0.126412 | 0.013596 |
| 0.008456 | 0.002387 | 0 | 0.010747 | 0 | 0.001904 | 0 | 0.073814 | 0 |

| ENTl | ENTm | PAR | POST | PRE | SUB | ProS | BLA |
|---|---|---|---|---|---|---|---|
| 6.072411 | 2.832519 | 0.501341 | 0.610597 | 0.553388 | 0.169398 | 0.070595 | 1.213927 |
| 4.147422 | 1.774693 | 0.316874 | 0.30884 | 0.30398 | 0.134549 | 0.03169 | 0.434479 |
| 2.46543 | 0.946348 | 0.169845 | 0.194739 | 0.179649 | 0.105126 | 0.037529 | 0.409771 |
| 1.492886 | 0.098354 | 0.012542 | 0.003062 | 0.002965 | 0.000674 | 0 | 0.213559 |
| 2.175421 | 0.373428 | 0.020839 | 0.019293 | 0.034514 | 0.01883 | 0.020096 | 0.435222 |
| 0.005615 | 0 | 0 | 0 | 0 | 0 | 0 | 0 |
| 0.006787 | 0 | 0 | 0 | 0 | 0 | 0 | 0 |
| 0.003534 | 0 | 0 | 0 | 0 | 0 | 0 | 0 |
| 0.007037 | 0 | 0 | 0 | 0 | 0 | 0 | 0 |
| 0.001564 | 0 | 0 | 0 | 0 | 0 | 0 | 0 |
| 6.078026 | 2.832519 | 0.501341 | 0.610597 | 0.553388 | 0.169398 | 0.070595 | 1.213927 |

| | | | | | | | |
|---|---|---|---|---|---|---|---|
| 4.154209 | 1.774693 | 0.316874 | 0.30884 | 0.30398 | 0.134549 | 0.03169 | 0.434479 |
| 2.468964 | 0.946348 | 0.169845 | 0.194739 | 0.179649 | 0.105126 | 0.037529 | 0.409771 |
| 1.499923 | 0.098354 | 0.012542 | 0.003062 | 0.002965 | 0.000674 | 0 | 0.213559 |
| 2.176985 | 0.373428 | 0.020839 | 0.019293 | 0.034514 | 0.01883 | 0.020096 | 0.435222 |

| ENTl | ENTm | PAR | POST | PRE | SUB | ProS | BLA |
|---|---|---|---|---|---|---|---|
| 0.999076 | 0.466026 | 0.082484 | 0.10046 | 0.091047 | 0.027871 | 0.011615 | 0.199724 |
| 0.998366 | 0.427204 | 0.076278 | 0.074344 | 0.073174 | 0.032389 | 0.007628 | 0.104588 |
| 0.998569 | 0.383298 | 0.068792 | 0.078875 | 0.072763 | 0.042579 | 0.0152 | 0.165969 |
| 0.995308 | 0.065573 | 0.008362 | 0.002041 | 0.001977 | 0.000449 | 0 | 0.14238 |
| 0.999282 | 0.171534 | 0.009572 | 0.008862 | 0.015854 | 0.00865 | 0.009231 | 0.19992 |
| 0.000924 | 0 | 0 | 0 | 0 | 0 | 0 | 0 |
| 0.001634 | 0 | 0 | 0 | 0 | 0 | 0 | 0 |
| 0.001431 | 0 | 0 | 0 | 0 | 0 | 0 | 0 |
| 0.004692 | 0 | 0 | 0 | 0 | 0 | 0 | 0 |
| 0.000718 | 0 | 0 | 0 | 0 | 0 | 0 | 0 |

| ENTl | ENTm | PAR | POST | PRE | SUB | ProS | BLA |
|---|---|---|---|---|---|---|---|
| 0.99812 | 0.302727 | 0.049098 | 0.052916 | 0.050963 | 0.022387 | 0.008735 | 0.162516 |
| 0.00188 | 0 | 0 | 0 | 0 | 0 | 0 | 0 |
| 0.001444 | 0.156273 | 0.033054 | 0.039805 | 0.035235 | 0.015542 | 0.005057 | 0.036212 |
| 0.001444 | 0 | 0 | 0 | 0 | 0 | 0 | 0 |

| SUMl | VTA/SNc | DR | LC |
|---|---|---|---|
| 44 | 13 | 56 | 1 |
| 20 | 5 | 34 | 0 |
| 24 | 4 | 24 | 0 |
| 36 | 8 | 45 | 1 |
| 0 | 0 | 0 | 0 |
| 0 | 0 | 0 | 0 |
| 0 | 0 | 0 | 0 |
| 0 | 0 | 0 | 0 |
| 44 | 13 | 56 | 1 |
| 20 | 5 | 34 | 0 |
| 24 | 4 | 24 | 0 |
| 36 | 8 | 45 | 1 |

| SUMl | VTA/SNc | DR | LC |
|---|---|---|---|
| 0.030492 | 0.009009 | 0.038808 | 0.000693 |
| 0.027778 | 0.006944 | 0.047222 | 0 |
| 0.030496 | 0.005083 | 0.030496 | 0 |
| 0.028777 | 0.006395 | 0.035971 | 0.000799 |
| 0 | 0 | 0 | 0 |

| | | | |
|---|---|---|---|
| 0 | 0 | 0 | 0 |
| 0 | 0 | 0 | 0 |
| 0 | 0 | 0 | 0 |
| SUMl | VTA/SNc | DR | LC |
| 0.029386 | 0.006858 | 0.038124 | 0.000373 |
| 0 | 0 | 0 | 0 |
| 0.001163 | 0.001414 | 0.006043 | 0.000375 |
| 0 | 0 | 0 | 0 |

| MTN | ILM | RT | LHA | VTA | SNr | MRN | PAG |
|---|---|---|---|---|---|---|---|
| 0.009964 | 0.019216 | 0 | 0.034557 | 0.005435 | 0.005658 | 0.037275 | 0.01087 |
| 0 | 0 | 0 | 0 | 0 | 0 | 0 | 0 |
| 0.009964 | 0.019216 | 0 | 0.034557 | 0.005435 | 0.005658 | 0.037275 | 0.01087 |
| -0.0936 | -0.18052 | 0 | -0.32464 | -0.05106 | -0.05315 | -0.35017 | -0.10211 |
| 0 | 0 | 0 | 0 | 0 | 0 | 0 | 0 |

| MTN | ILM | RT | LHA | VTA | SNr | MRN | PAG |
|---|---|---|---|---|---|---|---|
| 0 | 1.55 | 1.3125 | 0 | 0 | 1.3 | 0 | 0 |
| 0 | 0.25 | 0 | 0 | 0 | 0 | 0 | 0 |
| 0 | 1.8 | 1.3125 | 0 | 0 | 1.3 | 0 | 0 |
| 0 | -0.40697 | -0.34461 | 0 | 0 | -0.34133 | 0 | 0 |
| 0 | 0.065641 | 0 | 0 | 0 | 0 | 0 | 0 |

| NDB | POL | AM | PVT | PT | RE | CM | PIL | DMH |
|---|---|---|---|---|---|---|---|---|
| 0.108636 | 0.016413 | 0.103262 | 0.459674 | 0.360574 | 0.421101 | 0.443341 | 0.050738 | 0.01669 |
| 0.059522 | 0.015495 | 0.052069 | 0.337254 | 0.191644 | 0.19263 | 0.219715 | 0.077953 | 0.004623 |
| 0.010975 | 0.004172 | 0.017238 | 0.095493 | 0.036951 | 0.046578 | 0.050547 | 0.025686 | 0.003229 |
| 0.025693 | 0.025299 | 0.038665 | 0.270229 | 0.108021 | 0.14925 | 0.276103 | 0.106887 | 0.002415 |
| 0.008063 | 0.005626 | 0.103597 | 0.040277 | 0.009306 | 0.031964 | 0.192813 | 0.002164 | 1.31E-05 |
| 0.009661 | 0 | 0 | 0.039061 | 0 | 0.048273 | 0.06684 | 0 | 0.007902 |
| 0.003426 | 0 | 0 | 0.027431 | 0 | 0.008873 | 0.036586 | 0 | 0.005768 |

| | | | | | | | | |
|---|---|---|---|---|---|---|---|---|
| 0.000228 | 0 | 0 | 0.008615 | 0 | 0.002668 | 0.004727 | 0 | 0.001388 |
| 0.002329 | 0 | 0 | 0.018674 | 0 | 0.008804 | 0.034494 | 0 | 0.002929 |
| 0.001077 | 0 | 0 | 0.008245 | 0 | 0.000998 | 0.030027 | 0 | 0.000428 |
| 0.118297 | 0.016413 | 0.103262 | 0.498735 | 0.360574 | 0.469374 | 0.510181 | 0.050738 | 0.024592 |
| 0.062948 | 0.015495 | 0.052069 | 0.364685 | 0.191644 | 0.201503 | 0.256301 | 0.077953 | 0.010391 |
| 0.011203 | 0.004172 | 0.017238 | 0.104108 | 0.036951 | 0.049246 | 0.055274 | 0.025686 | 0.004617 |
| 0.028022 | 0.025299 | 0.038665 | 0.288903 | 0.108021 | 0.158054 | 0.310597 | 0.106887 | 0.005344 |
| 0.00914 | 0.005626 | 0.103597 | 0.048522 | 0.009306 | 0.032962 | 0.22284 | 0.002164 | 0.000441 |
| NDB | POL | AM | PVT | PT | RE | CM | PIL | DMH |
| 0.1098711 | 0.0165996 | 0.104436 | 0.4648999 | 0.3646733 | 0.4258884 | 0.4483813 | 0.0513148 | 0.0168797 |
| 0.1109856 | 0.0288922 | 0.0970886 | 0.6288486 | 0.3573421 | 0.3591806 | 0.4096837 | 0.1453523 | 0.0086201 |
| 0.0561223 | 0.0213342 | 0.0881491 | 0.4883179 | 0.1889545 | 0.2381836 | 0.2584797 | 0.1313492 | 0.016512 |
| 0.047156 | 0.0464329 | 0.0709644 | 0.4959686 | 0.1982579 | 0.2739281 | 0.5067496 | 0.1961766 | 0.0044324 |
| 0.0289021 | 0.0201666 | 0.3713474 | 0.1443744 | 0.0333577 | 0.1145762 | 0.6911455 | 0.0077569 | 4.696E-05 |
| 0.0097708 | 0 | 0 | 0.0395051 | 0 | 0.0488218 | 0.0675999 | 0 | 0.0079918 |
| 0.0063882 | 0 | 0 | 0.0511482 | 0 | 0.0165447 | 0.0682188 | 0 | 0.0107551 |
| 0.0011659 | 0 | 0 | 0.0440541 | 0 | 0.0136432 | 0.0241722 | 0 | 0.0070977 |
| 0.0042746 | 0 | 0 | 0.0342736 | 0 | 0.0161585 | 0.0633091 | 0 | 0.0053758 |
| 0.0038605 | 0 | 0 | 0.0295545 | 0 | 0.0035774 | 0.1076329 | 0 | 0.0015342 |
| 0.119642 | 0.0166 | 0.104436 | 0.504405 | 0.364673 | 0.47471 | 0.515981 | 0.051315 | 0.024872 |
| 0.117374 | 0.028892 | 0.097089 | 0.679997 | 0.357342 | 0.375725 | 0.477902 | 0.145352 | 0.019375 |
| 0.057288 | 0.021334 | 0.088149 | 0.532372 | 0.188955 | 0.251827 | 0.282652 | 0.131349 | 0.02361 |
| 0.051431 | 0.046433 | 0.070964 | 0.530242 | 0.198258 | 0.290087 | 0.570059 | 0.196177 | 0.009808 |
| 0.032763 | 0.020167 | 0.371347 | 0.173929 | 0.033358 | 0.118154 | 0.798778 | 0.007757 | 0.001581 |
| NDB | POL | AM | PVT | PT | RE | CM | PIL | DMH |
| 0.070607 | 0.026685 | 0.146397 | 0.444482 | 0.228517 | 0.282351 | 0.462888 | 0.10639 | 0.009298 |
| 0.005092 | 0 | 0 | 0.039707 | 0 | 0.019749 | 0.066187 | 0 | 0.006551 |
| 0.033678 | 0.010655 | 0.113029 | 0.160652 | 0.123047 | 0.106467 | 0.140605 | 0.067771 | 0.006622 |
| 0.00287 | 0 | 0 | 0.007516 | 0 | 0.015279 | 0.026448 | 0 | 0.003053 |

| SUM | TMv | VMH | PH | LHA | VTA | PPN | DR | PCG |
|---|---|---|---|---|---|---|---|---|
| 0.127237 | 0.03544 | 0.013064 | 0.01979 | 0.052452 | 0.042112 | 0.024893 | 0.45631 | 0.009815 |
| 0.079486 | 0.02593 | 0.012274 | 0.008834 | 0.022098 | 0.030743 | 0.017105 | 0.212158 | 0.004344 |
| 0.012807 | 0.007118 | 0.000692 | 0.004423 | 0.005648 | 0.003197 | 0.005988 | 0.075152 | 0.00079 |
| 0.037416 | 0.011457 | 0.003326 | 0.006913 | 0.017409 | 0.031016 | 0.012874 | 0.206108 | 0.001513 |
| 0.016381 | 0.000945 | 0.000659 | 0.000948 | 0.003152 | 0.008353 | 0.009655 | 0.103802 | 0 |
| 0 | 0.01169 | 0 | 0 | 0 | 0 | 0.007726 | 0.147952 | 0 |
| 0 | 0.0031 | 0 | 0 | 0 | 0 | 0.002911 | 0.058072 | 0 |

| | | | | | | | | |
|---|---|---|---|---|---|---|---|---|
| 0 | 0.001261 | 0 | 0 | 0 | 0 | 0.001677 | 0.015706 | 0 |
| 0 | 0.00071 | 0 | 0 | 0 | 0 | 0.003096 | 0.050947 | 0 |
| 0 | 0.001703 | 0 | 0 | 0 | 0 | 0.003524 | 0.02961 | 0 |
| 0.127237 | 0.04713 | 0.013064 | 0.01979 | 0.052452 | 0.042112 | 0.032619 | 0.604262 | 0.009815 |
| 0.079486 | 0.02903 | 0.012274 | 0.008834 | 0.022098 | 0.030743 | 0.020016 | 0.27023 | 0.004344 |
| 0.012807 | 0.008379 | 0.000692 | 0.004423 | 0.005648 | 0.003197 | 0.007665 | 0.090858 | 0.00079 |
| 0.037416 | 0.012167 | 0.003326 | 0.006913 | 0.017409 | 0.031016 | 0.01597 | 0.257055 | 0.001513 |
| 0.016381 | 0.002648 | 0.000659 | 0.000948 | 0.003152 | 0.008353 | 0.013179 | 0.133412 | 0 |
| SUM | TMv | VMH | PH | LHA | VTA | PPN | DR | PCG |
| 0.1286835 | 0.0358429 | 0.0132125 | 0.020015 | 0.0530483 | 0.0425908 | 0.025176 | 0.4614977 | 0.0099266 |
| 0.1482107 | 0.0483494 | 0.0228863 | 0.016472 | 0.0412042 | 0.0573238 | 0.0318942 | 0.3955928 | 0.0080999 |
| 0.0654905 | 0.036399 | 0.0035386 | 0.0226177 | 0.0288819 | 0.0163483 | 0.0306205 | 0.3843011 | 0.0040398 |
| 0.068672 | 0.0210278 | 0.0061044 | 0.0126879 | 0.0319519 | 0.0569257 | 0.0236285 | 0.3782832 | 0.0027769 |
| 0.0587183 | 0.0033874 | 0.0023622 | 0.0033981 | 0.0112985 | 0.0299416 | 0.0346087 | 0.3720822 | 0 |
| 0 | 0.0118229 | 0 | 0 | 0 | 0 | 0.0078138 | 0.149634 | 0 |
| 0 | 0.0057803 | 0 | 0 | 0 | 0 | 0.0054279 | 0.1082819 | 0 |
| 0 | 0.0064483 | 0 | 0 | 0 | 0 | 0.0085756 | 0.080315 | 0 |
| 0 | 0.0013031 | 0 | 0 | 0 | 0 | 0.0056823 | 0.0935063 | 0 |
| 0 | 0.0061045 | 0 | 0 | 0 | 0 | 0.0126319 | 0.1061382 | 0 |
| 0.128684 | 0.047666 | 0.013213 | 0.020015 | 0.053048 | 0.042591 | 0.03299 | 0.611132 | 0.009927 |
| 0.148211 | 0.05413 | 0.022886 | 0.016472 | 0.041204 | 0.057324 | 0.037322 | 0.503875 | 0.0081 |
| 0.065491 | 0.042847 | 0.003539 | 0.022618 | 0.028882 | 0.016348 | 0.039196 | 0.464616 | 0.00404 |
| 0.068672 | 0.022331 | 0.006104 | 0.012688 | 0.031952 | 0.056926 | 0.029311 | 0.47179 | 0.002777 |
| 0.058718 | 0.009492 | 0.002362 | 0.003398 | 0.011298 | 0.029942 | 0.047241 | 0.47822 | 0 |
| SUM | TMv | VMH | PH | LHA | VTA | PPN | DR | PCG |
| 0.093955 | 0.029001 | 0.009621 | 0.015038 | 0.033277 | 0.040626 | 0.029186 | 0.398351 | 0.004969 |
| 0 | 0.006292 | 0 | 0 | 0 | 0 | 0.008026 | 0.107575 | 0 |
| 0.036989 | 0.015463 | 0.007628 | 0.006712 | 0.013841 | 0.015824 | 0.004142 | 0.032512 | 0.003598 |
| 0 | 0.003341 | 0 | 0 | 0 | 0 | 0.0026 | 0.023292 | 0 |

| CS | LC | NI | RPO | IP |
|---|---|---|---|---|
| 0.067173 | 0.299076 | 0.077568 | 0.034461 | 0 |
| 0.024664 | 0.306459 | 0.03085 | 0.010318 | 0 |
| 0.006324 | 0.03591 | 0.007873 | 0.002651 | 0 |
| 0.022606 | 0.19652 | 0.007791 | 0.000902 | 0 |
| 0.011489 | 0.015805 | 0.007006 | 0 | 0 |
| 0.028589 | 0.001804 | 0.021105 | 0.015482 | 3.48E-05 |
| 0.009615 | 0.009572 | 0.00458 | 0.004691 | 0 |

| | | | | |
|---|---|---|---|---|
| 0.00178 | 0 | 0 | 0.004535 | 2.99E-07 |
| 0.009387 | 9.11E-19 | 0.00264 | 0.008014 | 8.03E-06 |
| 0.009617 | 0 | 0 | 0.006174 | 0 |
| 0.095762 | 0.30088 | 0.098673 | 0.049943 | 3.48E-05 |
| 0.034279 | 0.316031 | 0.03543 | 0.015009 | 0 |
| 0.008104 | 0.03591 | 0.007873 | 0.007186 | 2.99E-07 |
| 0.031993 | 0.19652 | 0.010431 | 0.008916 | 8.03E-06 |
| 0.021106 | 0.015805 | 0.007006 | 0.006174 | 0 |
| CS | LC | NI | RPO | IP |
| 0.0679367 | 0.3024761 | 0.0784499 | 0.0348528 | 0 |
| 0.0459888 | 0.5714278 | 0.0575233 | 0.0192391 | 0 |
| 0.0323387 | 0.1836312 | 0.0402598 | 0.0135563 | 0 |
| 0.0414902 | 0.3606858 | 0.0142993 | 0.0016555 | 0 |
| 0.0411828 | 0.0566536 | 0.0251133 | 0 | 0 |
| 0.028914 | 0.0018245 | 0.0213449 | 0.015658 | 3.52E-05 |
| 0.0179283 | 0.0178481 | 0.0085399 | 0.0087469 | 0 |
| 0.0091023 | 0 | 0 | 0.0231904 | 1.529E-06 |
| 0.0172286 | 1.672E-18 | 0.0048454 | 0.0147086 | 1.474E-05 |
| 0.0344725 | 0 | 0 | 0.0221309 | 0 |
| 0.096851 | 0.304301 | 0.099795 | 0.050511 | 3.52E-05 |
| 0.063917 | 0.589276 | 0.066063 | 0.027986 | 0 |
| 0.041441 | 0.183631 | 0.04026 | 0.036747 | 1.53E-06 |
| 0.058719 | 0.360686 | 0.019145 | 0.016364 | 1.47E-05 |
| 0.075655 | 0.056654 | 0.025113 | 0.022131 | 0 |
| CS | LC | NI | RPO | IP |
| 0.045787 | 0.294975 | 0.043129 | 0.013861 | 0 |
| 0.021529 | 0.003935 | 0.006946 | 0.016887 | 1.03E-05 |
| 0.011926 | 0.173202 | 0.022885 | 0.012734 | 0 |
| 0.009033 | 0.006993 | 0.007884 | 0.005287 | 1.36E-05 |
