## Supplementary Table 7 for "Regional and cell type-specific afferent and efferent projections of the mouse claustrum"

| Major brain divisions | full name | Acronym |
|---|---|---|
| Isocortex | Frontal poleal cortex | FRP |
| Isocortex | Primary motor area | MOp |
| Isocortex | Secondary motor area | MOs |
| Isocortex | Primary somatosensory area | SSp |
| Isocortex | Primary somatosensory area, nose | SSp-n |
| Isocortex | Primary somatosensory area, barrel field | SSp-bfd |
| Isocortex | Primary somatosensory area, lower limb | SSp-ll |
| Isocortex | Primary somatosensory area, mouth | SSp-m |
| Isocortex | Primary somatosensory area, upper limb | SSp-ul |
| Isocortex | Primary somatosensory area, trunk | SSp-tr |
| Isocortex | Primary somatosensory area, unassigned | SSp-un |
| Isocortex | Supplemental somatosensory area | SSs |
| Isocortex | Gustatory areas | GU |
| Isocortex | Visceral area | VISC |
| Isocortex | Dorsal auditory area | AUDd |
| Isocortex | Primary auditory area | AUDp |
| Isocortex | Posterior auditory area | AUDpo |
| Isocortex | Ventral auditory area | AUDv |
| Isocortex | Anterolateral visual area | VISal |
| Isocortex | Anteromedial visual area | VISam |
| Isocortex | Lateral visual area | VISl |
| Isocortex | Primary visual area | VISp |
| Isocortex | Posterolateral visual area | VISpl |
| Isocortex | posteromedial visual area | VISpm |
| Isocortex | Laterointermediate area | VISli |
| Isocortex | Postrhinal area | VISpor |
| Isocortex | Anterior cingulate area, dorsal part | ACAd |
| Isocortex | Anterior cingulate area, ventral part | ACAv |
| Isocortex | Prelimbic area | PL |
| Isocortex | Infralimbic area | ILA |
| Isocortex | Orbital area | ORB |
| Isocortex | Orbital area, lateral part | ORBl |
| Isocortex | Orbital area, medial part | ORBm |
| Isocortex | Orbital area, ventrolateral part | ORBvl |
| Isocortex | Agranular insular area | AI |
| Isocortex | Agranular insular area, dorsal part | AId |
| Isocortex | Agranular insular area, posterior part | AIp |
| Isocortex | Agranular insular area, ventral part | AIv |
| Isocortex | Retrosplenial area | RSP |
| Isocortex | Retrosplenial area, lateral agranular part | RSPagl |
| Isocortex | Retrosplenial area, dorsal part | RSPd |
| Isocortex | Retrosplenial area, ventral part | RSPv |
| Isocortex | Posterior parietal association areas | PTLp |
| Isocortex | Anterior area | VISa |
| Isocortex | Rostrolateral visual area | VISrl |
| Isocortex | Temporal association areas | TEa |

| | | |
|---|---|---|
| Isocortex | Perirhinal area | PERI |
| Isocortex | Ectorhinal area | ECT |
| Ofactory areas | Main olfactory bulb | MOB |
| Ofactory areas | Anterior olfactory nucleus | AON |
| Ofactory areas | Taenia tecta | TT |
| Ofactory areas | Taenia tecta, dorsal part | TTd |
| Ofactory areas | Taenia tecta, ventral part | TTv |
| Ofactory areas | Dorsal peduncular area | DP |
| Ofactory areas | Piriform area | PIR |
| Ofactory areas | Nucleus of the lateral olfactory tract | NLOT |
| Ofactory areas | Cortical amygdalar area, anterior part | COAa |
| Ofactory areas | Cortical amygdalar area, posterior part | COAp |
| Ofactory areas | Piriform-amygdalar area | PAA |
| Ofactory areas | Postpiriform transition area | TR |
| Hippocampal formation | Field CA1 | CA1 |
| Hippocampal formation | Entorhinal area | ENT |
| Hippocampal formation | Entorhinal area, lateral part | ENTl |
| Hippocampal formation | Entorhinal area, medial part, dorsal zone | ENTm |
| Hippocampal formation | Parasubiculum | PAR |
| Hippocampal formation | Postsubiculum | POST |
| Hippocampal formation | Presubiculum | PRE |
| Hippocampal formation | Subiculum | SUB |
| Hippocampal formation | Prosubiculum | ProS |
| Hippocampal formation | Hippocampo-amygdalar transition area | HATA |
| Hippocampal formation | Area prostriata | APr |
| Hippocampal formation | Claustrum | CLA |
| Hippocampal formation | Endopiriform nucleus | EP |
| Hippocampal formation | Endopiriform nucleus, dorsal part | EPd |
| Hippocampal formation | Endopiriform nucleus, ventral part | EPv |
| Hippocampal formation | Lateral amygdalar nucleus | LA |
| Hippocampal formation | Basolateral amygdalar nucleus | BLA |
| Hippocampal formation | Basomedial amygdalar nucleus | BMA |
| Hippocampal formation | Posterior amygdalar nucleus | PA |
| Striatum | Caudoputamen | CP |
| Striatum | Nucleus accumbens | ACB |
| Striatum | Anterior amygdalar area | AAA |
| Striatum | Central amygdalar nucleus | CEA |
| Striatum | Medial amygdalar nucleus | MEA |
| Pallidum | Globus pallidus, external segment | GPe |
| Pallidum | Globus pallidus, internal segment | GPi |
| Pallidum | Substantia innominata | SI |
| Pallidum | Magnocellular nucleus | MA |
| Pallidum | Medial septal nucleus | MS |
| Pallidum | Diagonal band nucleus | NDB |
| Pallidum | Triangular nucleus of septum | TRS |
| Pallidum | Bed nuclei of the stria terminalis | BST |
| Thalamus | Ventral medial nucleus of the thalamus | VM |

| | | |
|---|---|---|
| Thalamus | Posterior triangular thalamic nucleus | PoT |
| Thalamus | Subparafascicular nucleus, magnocellular part | SPFm |
| Thalamus | Subparafascicular nucleus, parvicellular part | SPFp |
| Thalamus | Subparafascicular area | SPA |
| Thalamus | Peripeduncular nucleus | PP |
| Thalamus | Lateral posterior nucleus of the thalamus | LP |
| Thalamus | Posterior complex of the thalamus | PO |
| Thalamus | Posterior limiting nucleus of the thalamus | POL |
| Thalamus | Suprageniculate nucleus | SGN |
| Thalamus | Anteroventral nucleus of thalamus | AV |
| Thalamus | Anteromedial nucleus | AM |
| Thalamus | Anterodorsal nucleus | AD |
| Thalamus | Interanteromedial nucleus of the thalamus | IAM |
| Thalamus | Interanterodorsal nucleus of the thalamus | IAD |
| Thalamus | Lateral dorsal nucleus of thalamus | LD |
| Thalamus | Intermediodorsal nucleus of the thalamus | IMD |
| Thalamus | Mediodorsal nucleus of thalamus | MD |
| Thalamus | Submedial nucleus of the thalamus | SMT |
| Thalamus | Perireunensis nucleus | PR |
| Thalamus | Paraventricular nucleus of the thalamus | PVT |
| Thalamus | Parataenial nucleus | PT |
| Thalamus | Nucleus of reuniens | RE |
| Thalamus | Xiphoid thalamic nucleus | Xi |
| Thalamus | Rhomboid nucleus | RH |
| Thalamus | Central medial nucleus of the thalamus | CM |
| Thalamus | Paracentral nucleus | PCN |
| Thalamus | Central lateral nucleus of the thalamus | CL |
| Thalamus | Parafascicular nucleus | PF |
| Thalamus | Posterior intralaminar thalamic nucleus | PIL |
| Thalamus | Lateral habenula | LH |
| Hypothalamus | Paraventricular hypothalamic nucleus | PVH |
| Hypothalamus | Arcuate hypothalamic nucleus | ARH |
| Hypothalamus | Dorsomedial nucleus of the hypothalamus | DMH |
| Hypothalamus | Medial preoptic area | MPO |
| Hypothalamus | Anterior hypothalamic nucleus | AHN |
| Hypothalamus | Supramammillary nucleus | SUM |
| Hypothalamus | Tuberomammillary nucleus, dorsal part | TMd |
| Hypothalamus | Tuberomammillary nucleus, ventral part | TMv |
| Hypothalamus | Dorsal premammillary nucleus | PMd |
| Hypothalamus | Ventral premammillary nucleus | PMv |
| Hypothalamus | Paraventricular hypothalamic nucleus, descendin | PVHd |
| Hypothalamus | Ventromedial hypothalamic nucleus | VMH |
| Hypothalamus | Posterior hypothalamic nucleus | PH |
| Hypothalamus | Lateral hypothalamic area | LHA |
| Hypothalamus | Lateral preoptic area | LPO |
| Hypothalamus | Parasubthalamic nucleus | PSTN |
| Hypothalamus | Retrochiasmatic area | RCH |

| | | |
|---|---|---|
| Hypothalamus | Subthalamic nucleus | STN |
| Hypothalamus | Tuberal nucleus | TU |
| Midbrain | Nucleus of the brachium of the inferior colliculus | NB |
| Midbrain | Substantia nigra, reticular part | SNr |
| Midbrain | Ventral tegmental area | VTA |
| Midbrain | Midbrain reticular nucleus, retrorubral area | RR |
| Midbrain | Midbrain reticular nucleus | MRN |
| Midbrain | Superior colliculus, motor related | SCm |
| Midbrain | Periaqueductal gray | PAG |
| Midbrain | Substantia nigra, compact part | SNc |
| Midbrain | Pedunculopontine nucleus | PPN |
| Midbrain | Rostral linear nucleus raphe | RL |
| Midbrain | Central linear nucleus raphe | CLI |
| Midbrain | Dorsal nucleus raphe | DR |
| Pons | Parabrachial nucleus | PB |
| Pons | Pontine central gray | PCG |
| Pons | Pontine reticular nucleus, caudal part | PRNc |
| Pons | Superior central nucleus raphe | CS |
| Pons | Locus ceruleus | LC |
| Pons | Nucleus incertus | NI |
| Pons | Laterodorsal tegmental nucleus | LDT |
| Pons | Pontine reticular nucleus | PRNr |
| Pons | Nucleus raphe pontis | RPO |
| Pons | Subceruleus nucleus | SLC |
| Pons | Sublaterodorsal nucleus | SLD |
| Medulla | Nucleus of the solitary tract | NTS |
| Cerebellum | Fastigial nucleus | FN |
| | Interposed nucleus | IP |
| | Dentate nucleus | DN |
